# Targeting Regulatory Factors Associated with the *Drosophila Myc cis*-Elements by Reporter Expression, Gel Shift Assay, and Mass Spectrometric Protein Identification

**DOI:** 10.1101/2025.04.04.647281

**Authors:** Jasmine Kharazmi, Thomas Brody, Cameron Moshfegh

**Affiliations:** Department of Reproductive Endocrinology and Plastic and Hand Surgery, Campus Schlieren, University of Zurich, 8952 Schlieren Switzerland; Neural Cell-Fate Determinants Section, National Institute of Neurological Disorders and Stroke, US National Institutes of Health, Bethesda, Maryland, USA.; Laboratory of Applied Mechanobiology, Department of Health Sciences and Technology, ETH Zurich, Switzerland

**Keywords:** Reporter Assay, Electrophoretic-Mobility Shift Assay, Solid Surface Magnetic Enrichment, Liquid Chromatography Mass Spectrometry, *Drosophila Myc*, Transcription

## Abstract

Transcription factor MYC is highly responsive to mitogenic stimuli to modulate the expression of several targets involved in cell growth, protein biogenesis, proliferation, and differentiation. However, MYC’s transcriptional regulation remains incompletely understood. Our previous work demonstrated that key regulatory elements controlling *Drosophila Myc* reside within its 5’ UTR and intergenic regions. In this study, we developed a highly sensitive and selective Solid Surface Magnetic Enrichment Protocol (SSMEP) to purify DNA-protein complexes formed on *Myc cis*-elements. Using analysis of truncated reporters, Electrophoretic-Mobility Shift Assay (EMSA), and mass spectrometric protein identification, we identified a cis-regulatory module (CRM) within the proximal 5’ UTR that appears sufficient to activate *lacZ* reporter expression in the ovary and embryos. Furthermore, combining CRMs from either the upstream region or the intergenic Downstream Promoter Element (DPE) region with the proximal 5’-UTR regulatory unit recapitulates expression patterns previously observed in larval tissues, ovaries, and embryos. The DPE itself exhibits promoter activity when fused in-frame to an adjacent binding site cluster, and drives expression mimicking that of wild-type *Myc*. Further *in vivo* validation is necessary to confirm the functional relevance of the associated factors and identified signaling cascades as *Myc* regulators. Additionally, this highly selective and sensitive protocol has potential applications in other model organisms and various fields of biomedical and synthetic biology research.

## Introduction

Developmental patterning of gene expression arises from complex interactions between transcription factors associated with enhancers and promoters (BERMAN *et al*. 2004; SMITH *et al*. 2023), orchestrating cell growth, proliferation, apoptosis, and differentiation throughout an organism’s development (EDGAR *et al*. 2001; PEARSON AND CREWS 2014; JIMENO-MARTIN *et al*. 2022). During early development, the stage of cell fate specification and organ shaping, gene activity must be tightly controlled in space and time to confer the organism’s normal body size and proper organ function (SMALL AND ARNOSTI 2020; RAY-JONES AND SPIVAKOV 2021).

As an evolutionarily conserved and powerful early super oncogene, MYC activity necessitates tight transcriptional and post-transcriptional control throughout development, adult life (CENCIONI *et al*. 2023), pathological conditions, and evolutionary history (GALLANT *et al*. 1996; LEE *et al*. 2012; LI *et al*. 2013; HSIEH *et al*. 2019; THEVENON *et al*. 2020; REITER *et al*. 2023). Binding of Myc/Max heterodimers to E-box sequences activates target genes that increase S-phase cyclin expression, enhance DNA replication (DANESHVAR *et al*. 2011), and promote cell growth and cell cycle progression (DE LA COVA AND JOHNSTON 2006; PROCHOWNIK 2022). Conversely, when Myc binds to other Myc-associated bHLHzip factors like Miz1, it activates cell cycle arrest genes (FIORESI *et al*. 2022), counteracting its own proliferative effects. Furthermore, Mad/Max or Mnt/Max heterodimers compete with Myc/Max for the same E-box binding sites, thereby repressing Myc/Max-driven transcriptional activation (GRANDORI *et al*. 2000; OSTER *et al*. 2000; LUSCHER 2001; ZHOU AND HURLIN 2001; LOO *et al*. 2005). Beyond these core regulatory functions, *Drosophila Myc*, similar to its human ortholog MYC, plays significant roles in regenerative growth, proliferation, and stem cell biology. For instance, Myc collaborates with Wingless signaling in tissue regeneration following surgical injury or apoptosis within imaginal discs (SMITH-BOLTON *et al*. 2009). Additionally, somatic stem cells (SSCs) within terminally differentiated organs maintain and regenerate tissues post-damage (QUINN *et al*. 2013). Consistent with this, MYC family proteins are crucial for replenishing organs after cancer treatment or injury, participating in processes such as midgut maintenance (REN *et al*. 2013), neuroblast maintenance, neural stem cell proliferation (RUST *et al*. 2018; SAMUELS *et al*. 2020), and epidermis regeneration (ZANET *et al*. 2005).

Since deregulated *MYC* expression is implicated in a wide range of diseases, from neurodegenerative cases (MARINKOVIC AND MARINKOVIC 2021; ANATSKAYA AND VINOGRADOV 2022) to cancer (DHANASEKARAN *et al*. 2022; CENCIONI *et al*. 2023) and diabetes (LUO *et al*. 2021; ROSSELOT *et al*. 2021), tightly controlled *MYC* regulation plays a pivotal role in organismal physiology and homeostasis throughout development and adult life. Evolutionary conservation and functional similarities between *Drosophila Myc* and human *cMYC* (GALLANT *et al*. 1996) have made the fruit fly, with its short life cycle and well-established genetics, a valuable model organism for studying *Myc* biology. *Myc* patterning is highly responsive to signaling molecules and mitogenic stimulation during both development and disease states. Literature indicates that signaling pathways such as Dpp, Wingless, Notch (JOHNSTON *et al*. 1999; HERRANZ *et al*. 2008), Hippo (NETO-SILVA *et al*. 2010), and Ecdysone (DELANOUE *et al*. 2010) can positively and/or negatively regulate *Myc* expression, thereby maintaining the balance between cell growth/proliferation and differentiation. Despite intensive studies of *Myc* regulation, many questions about the sophisticated and multi-level signaling cascades influencing *Myc* transcription remain unanswered.

In our previous work, we identified various *cis*-regulatory modules (CRMs) as Conserved Sequence Blocks (CSBs) throughout the *Myc* gene body, along with two main transcription start sites: a TATA-box containing promoter at the proximal 5’-end and a downstream promoter element (DPE) located in the 2^nd^ intron (KHARAZMI *et al*. 2012). In this study, we analyzed further truncations of *Myc-lacZ* transgenes containing the promoter and other conserved regulatory elements. Our findings confirm that *Myc*’s dynamic expression during oogenesis and early embryogenesis stems from transcription initiation at multiple sites. This includes a third newly identified start site at the 3’-end of intron 1 within the proximal 5’-UTR, containing the strong developmental enhancer P37/P38 (ChrX:3,374,908..3,374,960). We also determined that high expression levels in larval discs and brain require at least one additional late or differentiation enhancer beyond the regulatory elements sufficient for embryo and ovary expression. Analysis of DNA-protein interaction complexes identified key regulatory factors associated with *Myc cis*-elements, primarily cell cycle regulators and components of established signaling pathways including noncanonical/canonical Wingless, Notch, DPP, Hedgehog, MAPK, JAK/STAT, insulin-like receptor (INR), and Hippo. The relationships between these associated factors and their connection to developmental and metabolic signals relevant to *Myc* transcriptional and post-transcriptional regulation will be discussed.

## Material and Methods

### I. Generation of *lacZ* reporter fly strains

Constructs pJ2.1, pJ7, pJ8, pJ8.4, pJ8.5, J2.1, J7, J8, J8.4, J8.5 (Table S1.1) and intermediate vectors pCaSpeR4, pCaSpeR-NLSlacZ, SKII-dmyc5, and SKII-In2 were engineered and generated previously (KHARAZMI *et al*. 2012). Inserts for the new transgenes J7.1-J7.3 were derived from genomic sequences in SKII-dmyc5 and subsequently subcloned into pCaSpeR-NLSlacZ.

The J7 promoter region (ChrX:3,373,200-3,375,005) includes 100 base pairs upstream of the main transcription start site, noncoding exon1, intron1 and noncoding sequences of exon 2 (Table S1.1). Transgene J7.1, derived from the J7 construct, lacks sequences upstream of the main transcription start site containing the TATA1-GC1-Inr1 elements. To generate transgene J7.1, the intermediate plasmid SKII-dmyc5 and the vector pCaSpeR-NLSlacZ were digested separately with SacII and Acc65I. Following isolation on a 1.5% agarose gel, the 5’-SacII/Acc65I-3’ linearized and dephosphorylated vector was ligated overnight with the 1806 bp genomic 5’-SacII/Acc65I-3’ insert (ChrX:3,373,200-3,375,005) to create transgene J7.1 (Table S1.1).

To obtain the promoter sequences for transgene J7.2, a fragment of 2164bp from the *Myc* 5’ noncoding sequences was PCR amplified with the primers J7.2-F and J7.2-R (Table S1.2) and the forward primer introduced a SacII restriction site at the 5’-end of the PCR fragment, and the plasmid SKII-dmyc5 was used as template. PCR conditions consisted of an initial denaturation (98°C, 30s), followed by 30 cycles of denaturation (98°C, 10s), annealing (58°C, 30s), and extension (72°C, 2 min), with a final extension (72°C, 5 min) before holding at 4°C. The PCR product was then digested with 5’ SacII-Acc65 3’ to isolate a 1295 bp genomic insert fragment containing 14 bp from the 3’ end of exon1, In1 and the noncoding region of exon 2 (ChrX:3,373,710-3,375,005). This insert was subcloned into the 5’ SacII-Acc65I 3’ linearized and dephosphorylated pCaSpeR-NLSlacZ vector upstream of the lacZ reporter to generate transgene J7.2. The promoter region of J7.2 contains 100 bp upstream of the main transcription start site and noncoding exon 1.

To generate the J7.3 promoter, a 641 bp fragment was amplified from the SKII-dmyc5 plasmid using primers J7.3-F and J7.3-R (Table S1.2). The resulting genomic fragment (ChrX:3,373,088..3,373,728) was blunt-ligated (using CIAP, Roche Diagnostics, Indianapolis, IN for vector preparation) into the Acc65I linearized, Klenow-treated, and dephosphorylated pCaSpeR-NLSlacZ vector upstream of the lacZ reporter to create transgene J7.3. To obtain the genomic sequences for the J7.3 promoter, primer set J7.4-F and J7.4-R (Table S1.2) was used to amplify a 1562 bp fragment from *Myc* genomic sequences within the pJ2.1 enhancer region, which contains an E-box with the sequence 5’-CAGCTG-3’ (Table S1.3, primers IR5-P1 and Az5-P2-IR3). The J7.4-F primer introduced a SacII restriction site at the 5’ end of the PCR fragment, while primer J7.4-R introduced an Acc65I site at the 3’ end. Following digestion, the resulting 1527 bp genomic insert fragment (ChrX:3,369,648..3,371,174) with 5’ SacII-Acc65I 3’ ends was ligated into the 5’ SacII-Acc65I 3’ linearized and dephosphorylated pCaSpeR-NLSlacZ vector to create the intermediate plasmid pJ7.4-temp.

To create the composite promoter element in transgene pJ7.4, the pJ7 promoter (ChrX:3,373,047..3,375,005) (Table S1.1) was amplified using primer pair J7-F and J7-R (Table S1.2), introducing an Acc65I site at the 5’-end of the fragment. The reverse primer annealed 116 bp downstream of the *Myc* fragment within the multiple cloning site (MCS) and amplified the Acc65I site present at the 3’ end of the genomic fragment in the MCS. After digestion with Acc65I, the resulting 5’ Acc65I-Acc65I 3’ insert was ligated into the Acc65I linearized and dephosphorylated pJ7.4-temp vector at the 3’-end of the J2.1 enhancer region, creating transgene pJ7.4 with its composite promoter element (Table S1.1).

For transgene J7.5, a 932 bp sequence from the J8 enhancer region, containing a conserved 12 bp cluster (5’-CGC**GTGGGAA**AA-3’, which includes the *Hairless* (H) consensus binding site **GTGGGAA**), was first amplified using primers In2E-F-Not and In2E-R-Asp (Tables S1.2 and S1.3, primers IR5-P29 and IR5-P29-Az3). These primers introduced a NotI site at the 5’-end and an Acc65I site at the 3’-end of the fragment. This 5’ NotI-Acc65I 3’ genomic insert fragment (ChrX:3,379,324..3,380,255) was swapped with the full-length In2 in the J8 transgene to create the intermediate product pJ7.5-temp. Next, plasmid pJ7.4 was digested with Acc65I to isolate the pJ7 promoter sequence insert. This insert was then introduced into the Acc65I linearized and dephosphorylated pJ7.5-temp vector, downstream of the J8 enhancer sequences and upstream of the lacZ reporter, yielding the final transgene J7.5 (Table S1.1).

For detailed protocols concerning transfections, plasmid DNA preparations, the phage ɸC31 integrase transgenesis system, and the random P-element transgenesis system, please refer to the Materials and Methods section in *Kharazmi et al., 2012*.

The transgenes J2.1, J7, J7.1-J7.5, J8, J8.4, and J8.5, containing *Myc* genomic sequences, were sequenced at both distal and proximal sites. Distal sequencing used primers pcaF and Acc651_F (Table S1.2), while proximal sequencing employed primers pCAR and pZR (Table S1.2), with all reactions oriented toward the *Myc* gene. All genomic fragments, intermediate plasmids, engineered transforming vectors, and final transgenes were sequenced by Microsynth AG (Balgach, Switzerland). Apart from standard primers provided by Microsynth, all other oligonucleotides and sequencing primers were designed using the DNASTAR Lasergene module PrimerSelect and subsequently synthesized by Microsynth AG.

### II. *D. melanogaster* stocks and X-Gal staining assays

Transgenic lines J2.1, J8, J8.4, and J8.5 were generated using random P-element transgenesis and the phage ɸC31 integrase transgenesis system (KHARAZMI *et al*. 2012), while J7.1-J7.5 strains were generated by random P-element insertion into embryos of genotype *y[1] w[1118]* (performed by Rainbow Transgenic Flies Inc, CA). A *dpp-lacZ* fly stock (KHARAZMI *et al*. 2012) served as a positive control, and the *w[1118]* fly stock served as the negative control (Table S1.4). All fly stocks were maintained on standard medium at 25°C, and crosses were performed at the same temperature. Larval and adult female tissues for lacZ staining were dissected from homozygous F2 generations. X-Gal staining for transgenic lines J2.1, J8, J8.4, and J8.5 followed our standardized protocol (SONG *et al*. 2004; KHARAZMI AND MOSHFEGH 2013), while staining for *Myc-lacZ* lines J7-J7.5 followed a protocol adapted from the Walter Gehrig laboratory.

### III. Protein extraction from *Drosophila* embryos

Wild-type Canton-S flies (Bloomington *Drosophila* Stock Center; donor: Kyoto Stock Center, Kyoto Institute of Technology, Bloomington Stock Number #64349, originally Stock Number #1) (Table S1.4) were maintained at 25°C and 75-80% humidity in population cages and fed with yeast paste. Embryos were collected on apple-agar petri dishes over a 0-12 hour period after egg laying and stored at 4°C. Embryos were accumulated from up to three successive days (72 hours total) and stored at 4°C before processing. For harvesting, embryos were washed from the petri dishes using embryo collection solution (0.7% w/v NaCl, 0.04% v/v Triton X-100), followed by five extensive water washes to eliminate all traces of yeast. Embryos were then dechorionated by immersion in 50% v/v bleach solution for 2.5 minutes at room temperature, followed by three thorough rinses with running water for 5 minutes each. After drying with paper towels, the embryos were transferred to a pre-chilled beaker, and cold (4°C) Buffer I was added at a ratio of 3 ml per gram of embryos. Buffer I composition was: 15 mM HEPES (K+, pH 7.6), 10 mM KCl, 5 mM MgCl₂, 0.1 mM EDTA, 0.5 mM EGTA, 350 mM sucrose, 1 mM dithiothreitol, 1 mM sodium metabisulfite, 0.2 mM phenylmethylsulfonyl fluoride, and 1 mM benzamidine-HCl. The embryos were then dispersed in the buffer using a glass rod.

We disrupted the embryos using a 40-ml Wheaton Dounce homogenizer in a 4°C room until the suspension became very milky. The homogenate was immediately filtered through a single layer of Miracloth (Calbiochem cat. #475855) into a Sorvall GSA centrifuge bottle, and nuclei were pelleted in a Sorvall GSA rotor at 8000 rpm for 15 minutes at 4°C. The supernatant, designated as the Soluble Cytoplasmic Fraction (SCF), was carefully removed and stored at −80°C. After wiping lipids from the bottle walls with Kimwipes, we resuspended the tan-colored loose nuclei pellet in chilled Buffer I (3 ml/g embryos), taking care to avoid disturbing the yellow yolk pellet. The nuclei were gently dispersed using a 40-ml Wheaton Dounce homogenizer with a loose B pestle, applying only a single stroke; this suspension was then transferred to clean centrifuge bottles and re-pelleted in a Sorvall GSA rotor at 8000 rpm for 15 minutes at 4°C. Following supernatant removal and lipid clean-up, we gently swirled the nuclei in Buffer I (1 ml/g embryos). After homogenizing the nuclei with two to three strokes of the pestle, we repelleted them at 8000 rpm for 15 minutes at 4°C. Following supernatant removal, we suspended the nuclei in HEMG/0.1M KCl buffer (0.5 ml/g nuclei) composed of 25 mM HEPES (K+, pH 7.6), 0.1 M KCl, 12.5 mM MgCl₂, 0.1 mM EDTA, 20% (v/v) glycerol, 1.5 mM dithiothreitol, 1 mM sodium metabisulfite, 0.1 mM phenylmethylsulfonyl fluoride, and 1 mM benzamidine-HCl). We incubated this nuclear suspension on ice for 20 minutes (maximum 60 minutes) with occasional swirling and shaking, then subjected the mixture to ultracentrifugation in a Beckman SW28 rotor at 24,000 rpm (100,000 g) for 1 hour at 4°C. Following centrifugation, we carefully removed the topmost thin gray-white lipid layer with a spatula and discarded it. The yellow liquid layer underneath, comprising the Soluble Nuclear Fraction (SNF), was carefully aspirated, immediately divided into 0.1 to 1.0 ml aliquots, quick-frozen in liquid nitrogen, and stored at −80°C; the third gray layer and the pellet at the bottom, composed of DNA and nuclear debris, remained untouched. For protein quantification of SCF and SNF fractions, we loaded the extracts onto a 10% SDS-PAGE gel, performed electrophoresis for 1 hour at 80V, stained the gel with Coomassie Blue (LUO *et al*. 2006; HARRIS *et al*. 2007), and imaged it using the Odyssey Classic Infrared Imaging System at 700 nm. Additionally, protein concentration was determined using the Pierce™ BCA Protein Assay Kit (cat# 23225, Thermo Scientific™).

### IV. Designing of EMSA oligonucleotide pairs using Myc-CRMs as targets for protein binding

EMSA primers were designed from the *Myc* conserved *cis*-regulatory sequences within the proximal upstream and large intron regions. A dTCF/Pan site, a Wnt-dependent *cis*-regulatory module (W-CRM), served as positive control (Table S1.3). A sequence from the noncoding region of the *Mdm2* gene of mouse strain C57BL/6J (gi|372099100|ref|NC_000076.6|:117686685-117712961) was processed using the sequence shuffling tool (http://www.bioinformatics.org/sms2/shuffle_dna.html) to design a scrambled oligonucleotide as a negative control. For the visualization of double-stranded (ds) oligonucleotides, one strand was modified with the non-hazardous near-infrared fluorescence IRDye Dyomics 781 at one end and the complementary strand was modified with the same IRDye at the opposite end, so that after annealing the two dye molecules, one at the 5’-end and the other at the 3’-end, were in close proximity. To minimize steric hindrance for subsequent protein binding after magnetic bead attachment, each oligonucleotide pair was ordered twice; in one pair the reactive Azide moiety (N-hydroxysuccinimide (NHS) ester) was attached to one strand at the free 5’-end, and in the second pair the Azide group was attached to the 3’-end of the complementary strand. Since it was not clear whether proteins bind to one or both strands, this strategy helped to preserve bound proteins in the DNA-protein complexes during multiple wash steps. The oligonucleotides were synthesized at Microsynth AG. Considering key determining factors that significantly influence hybridization efficiency, namely low salt concentration and the rate of temperature decrease (Table S1.5), the final annealing reaction mix contained oligonucleotides at 1 pmol/µl (1:1 molar ratio) and 50 mM NaCl. The thermal cycler was programmed as follows: one cycle of initial denaturation at 95°C for 5 min; *40 cycles of hybridization starting at 95°C (**1°C/cycle) for 1 min each; annealing at 55°C for 30 min; *20 cycles of hybridization starting at 55°C (1°C/cycle) for 1 min each; followed by a hold at 4°C (*The number of cycles in hybridization steps was determined by the Tm of the strand pairs; the protocol described is for an oligonucleotide pair with a Tm of 55°C in 50 mM NaCl, pH 8.0); (**decrease of cycler’s heating 1°C per min). Double). To verify annealing, double-stranded oligonucleotides were subjected to electrophoresis in the Mini-PROTEAN system (Bio-Rad Laboratories Inc.) under non-denaturing conditions on a 5% Precast TBE PAGE gel (Bio-Rad Laboratories Inc., cat. #4565014) for 1 hour at 70V and visualized while still wet using the Odyssey Classic Infrared Imager at 800 nm wavelength.

### V. Electrophoretic-Mobility Shift Assay (EMSA) using double-stranded azide and Dyomics 781 modified oligonucleotides

We added 1.35 picomole of double-stranded targets (dsAzide-DNA-Dyo781) (Table S1.5) to binding mixtures containing: **2 μL** of 10x binding buffer (100 mM Tris, 500 mM KCl, 10 mM DTT; pH 7.5); **2 μL** of 25 mM DTT, 2.5% Tween®-20; 1 μL of poly(dI.dc) (1 μg/μL in 10 mM Tris, 1 mM EDTA, pH 7.5); **1 μL** of 100 mM MgCl₂; **1 μL** of 1% NP (Nonylphenoxypolyethoxyethanol); **200 μg** protein (SCF in cytoplasmic samples or SNF in nuclear samples); and ultrapure water to a final volume of **20 μL**. Reactions were incubated in the dark for 30 minutes. After adding **2 μL** of 10x Orange Loading Dye (Odyssey® Infrared EMSA Kit), we subjected the mixtures and PageRuler protein marker (Thermo Fisher, cat. #26616) to electrophoresis using the Mini-PROTEAN system under non-denaturing conditions on a 5% Precast TBE PAGE gel for 1 hour at 80V. We monitored the gels while still wet using the Odyssey Classic Infrared Imager at 800 nm wavelength. The acquired EMSA images were processed in Adobe Photoshop CS6 by adjusting image brightness and contrast.

### VI. Creation of chimeric carboxylic magnetic Bead-PEG-DNA construct

Based on existing technology, we developed a new protocol for enriching proteins from whole cell lysates or embryonic extracts for subsequent identification by mass spectrometry (MS) analysis. We built a chimeric construct consisting of a magnetic bead covalently attached to a Dibenzocyclooctyne-Poly Ethyl Glycol-amine (DBCO-PEG23-amine) linker, which was then attached to a ds DNA oligonucleotide via a copper-free Click Chemistry reaction. The hydrophilic PEG linker provides sufficient steric freedom on the bead surface, allowing adequate spatial configuration for protein binding. The oligonucleotide was modified at one end with an IRDye for infrared detection and at the other end with an azide group for covalent bonding with the PEG-linker via the Click reaction.

#### A). Activation and covalent coupling of ligand to the magnetic microspheres

Dynabeads® MyOne™ Carboxylic Acid (Invitrogen by Thermo Fisher cat. #65001) were resuspended by rolling the vial for >30 minutes, and 1 mL (10 mg) was transferred to a new tube. The tube was placed on a magnet rack for 2 minutes and the supernatant was aspirated. Magnetic beads were washed twice in 1 mL of 100 mM MES (2-[N-morpholino]ethane sulfonic acid, MW 213.25) buffer (pH 6.00) and resuspended in 100 μL of the same buffer (final concentration 10 μg/μL). In a separate 2 mL tube, we combined 140 μL magnetic beads (10 μg/μL), 84 μL of 1250 mM EDC (N-Ethyl-N’-(3-dimethylaminopropyl) carbodiimide hydrochloride, MW 191.7) (Thermo Fisher, cat.# 77149) (75 nmol EDC/1 μg bead), 42 μL of 100 mM linker Dibenzocyclooctyne-Poly Ethyl Glycol-amine (DBCO-PEG23-amine, free-end amine salt) (BroadPharm San Diego, CA, cat. #BP-25103; formula C67H113N3O25) (3 nmol PEG/1 μg bead), and 1x PBS (pH 7.5)/0.1% Triton X-100 to a final volume of 300 μL. The reaction mix was placed in a HulaMixer™ Sample Mixer (Thermo Fisher, cat.# 15920D) and incubated overnight at room temperature in the dark with gentle rotation (7 rpm) for 12 to 16 hours. After activation of carboxylic sites on the magnetic beads by EDC and formation of amide bonds between activated carboxylic groups and the amine group of DBCO-PEG23-amine, the Dynabeads® magnetic beads were washed 3 times for 10 minutes each with storage buffer (100 mM PBS, 0.15 M NaCl, pH 7.5, 0.1% Tween®-20) to remove excess EDC and PEG molecules.

#### B). Inactivation of the unbound carboxylic sites on Dynabeads® Magnetic Beads

The unbound carboxylic sites on Dynabeads® magnetic beads were quenched by incubation with 1 mL of 50 mM Tris buffer (pH 7.4) containing 0.01% Tween®-20 for >30 minutes. The ligand-bound magnetic beads (DBCO-Beads) were then gently washed 3 times with 1 mL of storage buffer, resuspended in the same buffer at a concentration of 10 μg/μL, and stored at 4°C for less than one week until further use.

#### C). Copper-free click chemistry (alkyne-azide) reaction

D). For the click chemistry reaction (covalent tethering of azide-modified DNA molecules to coated beads), 140 μL DBCO-Beads (10 μg/μL) were washed twice in 1 mL of 100 mM MES buffer (pH 6.00) and resuspended in 140 μL of storage buffer (final concentration 10 μg/μL).

We added 7 μL of double-stranded oligonucleotide (InfraRed Dyomics 781/Azide end-modified, 0.45 pmol/μL) to the beads, bringing the final volume to 300 μL with storage buffer. The mixture was incubated for 4.5 hours in a HulaMixer™ at room temperature in the dark with 7 rpm rotation. After the click chemistry reaction, the magnetic Bead-PEG-DNA construct was washed 4 times for 10 minutes each with storage buffer to remove all traces of unbound DNA molecules. The construct was resuspended in 500 μL of storage buffer and stored at 4°C for less than two days until use in protein binding experiments.

### VII. Enrichment of DNA/protein complex using magnetic Bead-PEG-DNA pulldown construct

The magnetic bead-PEG-DNA construct in storage buffer was carefully resuspended. After placement of the tube on a magnetic rack, the supernatant was aspirated. The beads were then rinsed once with storage buffer and resuspended in 140 μL of the same buffer to a final concentration of 10 μg/μL. We set up protein binding reactions for the soluble cytoplasmic fraction (SCF) and soluble nuclear fraction (SNF) samples using components from the Odyssey® Infrared EMSA Kit (Part #829-07910) with the following components per reaction: **a)** Magnetic bead-PEG-DNA construct (10 μg/ μL): **1400 μg**; **b)** 10x binding buffer: **10 μL**; **c)** 25 mM DTT, 2.5% Tween®-20: **10 μL**; **c)** poly (dI.dc), 1 μg/ μL: **5 μL**; **d)** 1% NP: **5 μ**L; **e)** 100 mM MgCl2: **5 μL**; **f)** Protein (SCF in cytoplasmic samples and SNF in nuclear samples): **200 μg**; **g)** ultrapure water was added to a final volume of **100 μL**. Sample tubes were incubated in the dark for 30 minutes by occasional manual flipping to keep the mixture in suspension. After incubation and aspiration of supernatants, the Beads-PEG DNA/protein complexes were gently washed 4 times for 5 minutes with storage buffer. Once more the residual liquid traces were removed, and the samples were stored at −80℃.

### VIII. Sample processing and protein identification by mass spectrometry (LC-MS/MS)

#### a) Sample preparation

Proteins from each sample (Table S1.6) were denatured and reduced by suspending the beads (both unmodified beads and Beads-PEG-DNA/protein complexes) in 8 M Urea, 100 mM Ammonium bicarbonate containing 5 mM DTT and incubating for 1 h at room temperature. The reduced/denatured proteins were alkylated at cysteine residues by adding iodoacetamide to a final concentration of 15 mM and incubating for 1 h. The reduced/alkylated samples were diluted 8-fold with 100 mM ammonium bicarbonate, and supernatants were collected after centrifugation at 20,000 x g for 5 min. Subsequently, samples were treated with 0.2 μg trypsin for 16 h at 37°C. The resulting tryptic peptides were isolated and desalted by micropurification (reverse phase) prior to MS analysis. We processed Myc-CRM samples, SCF and SNF extracts (starting materials), a Scrambled oligo (Scm) as negative control, a W-CRM configuration as positive control (ARCHBOLD *et al*. 2014), along with raw beads to identify potential contaminants. **b) MS analysis**: Peptides were dissolved in 0.1% formic acid and loaded onto a trap column (2 cm x 100 µm inner diameter). The peptides were then eluted from the trap column and separated on a 15-cm analytical column (75 µm inner diameter). Both columns contained ReproSil-Pur C18-AQ 3 µm resin (Dr. Maisch GmbH, Ammerbuch-Entringen, Germany). Peptides were eluted using a flow rate of 250 nL/min and a 30 min gradient from 5% to 35% phase B (0.1% formic acid and 90% acetonitrile). Samples were analyzed using a high-resolution LC-MS/MS setup consisting of an EASY-nLC 1200 system (Thermo Scientific) connected to a Thermo Eclipse Tribrid Mass Spectrometer (Thermo Scientific. **c) MS data processing**: The acquired MS raw data was processed using Proteome Discoverer 2.5 (Thermo Scientific) software. The data was searched against the *Drosophila melanogaster* (Fruit fly) proteome (https://www.uniprot.org/proteomes/UP000000803). To filter non-*Drosophila* contaminants, the data was searched against a contaminant database from the Max Planck Institute (MPI) (Germany) (https://lotus1.gwdg.de/mpg/mmbc/maxquant_input.nsf/7994124a4298328fc125748d0048fee2/$FILE/contaminants.fasta). Search parameters included: trypsin as protease; minimum peptide length of 6 residues; Precursor Mass Tolerance of 10 ppm; Fragment Mass Tolerance of 0.02 Da; and dynamic modifications including oxidation on Met and acetylation of the N-terminus. The search results were used for precursor ion quantification based on summed abundances of unique peptides without normalization.

### IX. Software processing of proteomics raw data

After filtering MS data and removing contaminants using Thermo Fisher Scientific™ Proteome Discoverer™ 2.5 software, protein candidates identified with high confidence (Protein FDR Confidence; False Discovery Rate (FDR) < 1%) were processed using the software tool ComplexBrowser (MICHALAK *et al*. 2019), accessible as a web service at https://computproteomics.bmb.sdu.dk/app_direct/ComplexBrowser/. ComplexBrowser is a tool for identifying protein complexes in experimental datasets using curated information from CORUM (Comprehensive Resource of Mammalian Protein Complexes) and complex portal databases, incorporating quality control metrics and interactive visualizations, with data normalization performed as recommended (MICHALAK *et al*. 2019). For *Drosophila* protein complexes, we manually curated protein complexes from our data and primarily used ComplexBrowser for quality control of our protein candidates. For each sample pair (including Negative Control, Myc-CRMs, and Positive Control), ComplexBrowser was used to generate boxplots of log2 abundance intensities with median normalization, histograms of the coefficient of variation (CV) of abundance intensities to assess replicate variation, and scatter plots comparing log abundance intensities with Pearson correlation values. Statistical analysis via volcano plots represented changes in protein abundance between conditions. Volcano plots were generated using the open-source online tool VolcaNoseR (GOEDHART AND LUIJSTERBURG 2020), accessible at https://huygens.science.uva.nl/VolcaNoseR2/. These plots show log2 abundance ratios for all protein candidates in each sample group compared to the Scm negative control on the x-axis and −log10(q-value) on the y-axis. Significance thresholds were set at −log10(q-value) > 2 (equivalent to q-value < 0.01) and an absolute abundance ratio > 1.5.

We utilized various web-based tools and community resources to manually identify protein complexes from our protein hits, including UniProt (https://www.uniprot.org/), The Interactive Fly (https://www.sdbonline.org/sites/fly/aimain/1aahome.htm), String (https://string-db.org/), and particularly the comprehensive resource FlyBase (https://flybase.org/reports/FBgn0013263.htm), along with relevant scientific literature.

## Results

### I. *Myc* P1/P2 distal enhancer patterns larval brain and discs while the intron 1 3’-end P37/P38 enhancer drives *Myc* expression in adult female tissues

Our previous work showed that the J2.1 transgene, containing 7.2 kb of upstream noncoding region, controls *Myc* expression in both larval and adult female tissues (KHARAZMI *et al*. 2012). In contrast, the smallest J7 transgene (1.928 kb in size; ChrX: 3,373,088..3,375,005), consisting of 100 bp upstream of exon 1, noncoding exon 1, intron 1, and the noncoding 5’-region of exon 2, is sufficient for expression in embryos and ovary but shows no activity in larval brain and imaginal tissues (KHARAZMI *et al*. 2012). These findings suggested that sequences within the J2.1 transgene likely contain a late enhancer for tissue-specific patterning in larval imaginal discs and brain, and further indicated that the 1.928 kb sequences within the proximal 5’-UTR might contain regulatory elements important for oogenesis and embryogenesis (Figure1 A-B, Table S1.1).

In this work, we examined both J2.1 and J7 transgenes in greater detail to isolate enhancer elements responsible for *Myc* expression in ovary, larval discs, and brain. We tested the functionality of specific noncoding Myc-CRMs within the J7 transgene (Figure 1B)—specifically the TATA-box containing P31/P32 (44 bp) and the P37/P38 sequence (53 bp upstream of exon 1)—in embryos and ovaries. We first deleted the P1 promoter harboring the Inr/TATA-box-containing Myc-CRM P31/P32 to create the J7.1 transgene (Figure 1C). As shown for transgene J7.2, we further truncated this region by removing 1295 bp (Figure 1D). Notably, *Myc-lacZ* expression in embryos and ovaries remained unchanged in these three transgenes (J7, J7.1, J7.2). However, with the J7.3 transgene, where we deleted intron 1 and noncoding exon 2 sequences (including the P37/P38 enhancer cluster) while retaining the upstream sequences containing the Myc-CRM P31/P32 within the P1 promoter and the entire exon 1, no detectable expression was observed in adult female tissues (Figure 1E). These results indicate that endogenous *Myc* expression in adult female tissues operates independently of P1 promoter activity. Furthermore, they suggest that the 100 bp sequence upstream of exon 1 is not required for *Myc-lacZ* activity in embryos and ovary. We conclude that the P37/P38 enhancer, a 53 base pair sequence (ChrX:3,374,908..3,374,960) (Table S1.3), is both necessary and sufficient for the endogenous patterning of *Myc* expression in highly proliferating embryonic and ovarian cells during early developmental stages. Previous studies have demonstrated that enhancers can undergo histone modifications, bind to RNA polymerase II, and initiate transcription in lineage-specific cell types during early development and differentiation (CATARINO *et al*. 2017; BARRAL AND DEJARDIN 2023; LINDHORST AND HALFON 2023). The transcriptional capacity of enhancers is particularly evident during early development, cellular differentiation, disease, and evolution (ANDERSSON *et al*. 2014; CATARINO *et al*. 2017; DAO *et al*. 2017; CATARINO AND STARK 2018; LINDHORST AND HALFON 2023). Based on previous findings and our current results, we refer to the Myc-CRM P37/P38 enhancer cluster (Figure 1, A-D and F; Table S1.3) as the “*Myc* oocyte element.”

**Figure 1.**
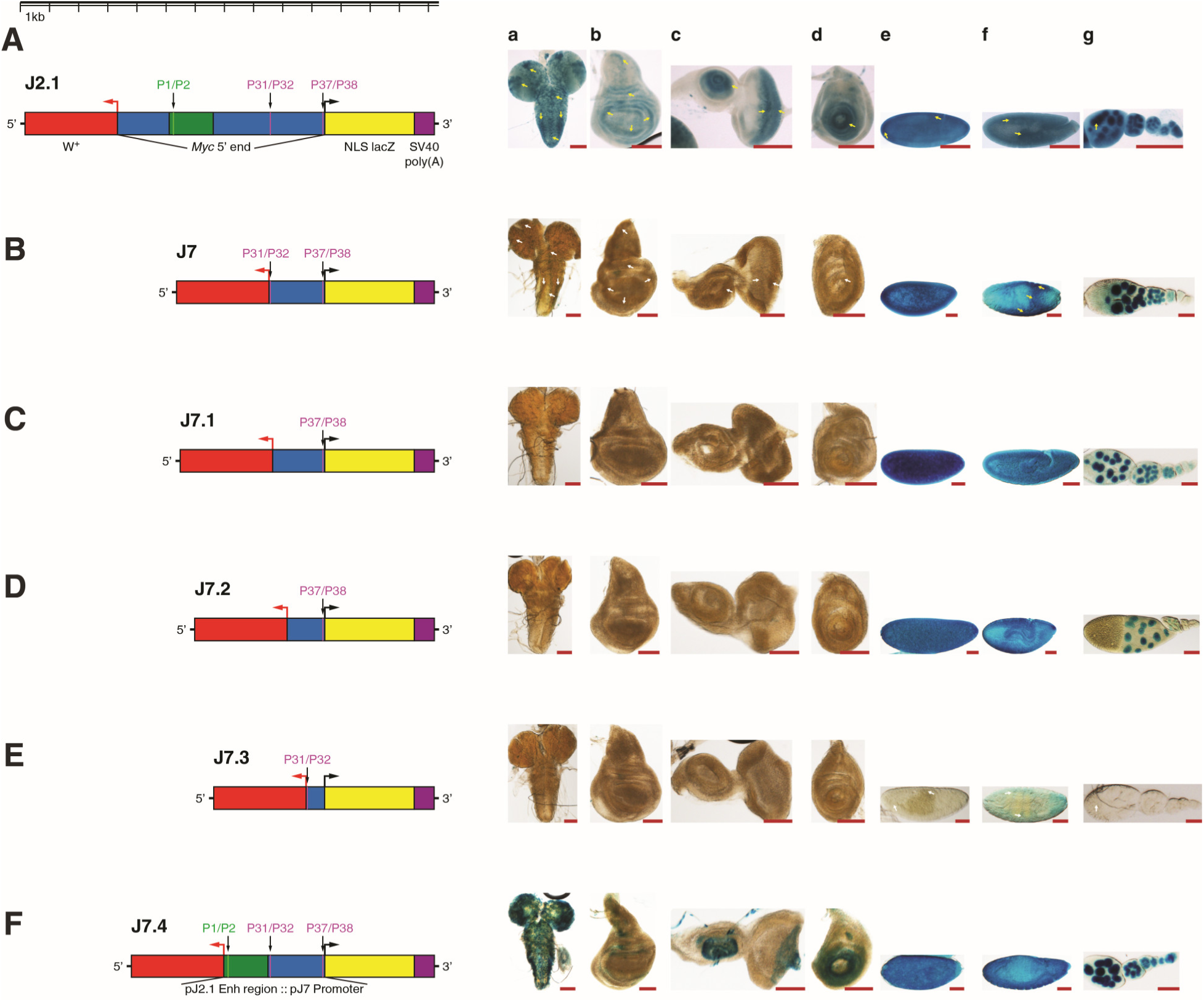
Partitioning *Myc* 5’ regulatory sequences detects a late developmental enhancer and an oocyte element. **(A)** J2.1 contains a 7.159 kb upstream regulatory region (*myc* 5’ end, indicated by blue and green), lacking open reading frame (ORF) sequences from the *Myc* 5’-end. These sequences include the J2.1 enhancer region (green) containing the upstream enhancer P1/P2 cluster. Downstream of this enhancer region are the TATA-box-containing Myc-CRM P31/P32 and the proximal oocyte element P37/P38 (blue). **(B)** J7 is the smallest truncation derived from J2.1, 1.923 kb in size. Sequences upstream of the P31/P32 Myc-CRM and the P1/P2 enhancer are deleted. **(C-E)** J7.1-J7.3 are truncations derived from J7. J7.1 and J7.2 contain the P37/P38 enhancer. However, J7.3 contains the P31/P32 promoter, but the P37/P38 enhancer is deleted. **(F)** The regulatory region in J7.4 is a composite element created by fusing sequences containing the P1/P2 enhancer (green) in-frame with the J7 regulatory sequences. Panels **(a-d)** show *lacZ* reporter expression patterns in larval discs and brain; panels **(e-g)** show expression in embryos and ovary. The **J2.1** transgene expresses *lacZ* in all tested tissues, and the activity reflects the *Myc* endogenous mRNA distribution (yellow arrowheads) (Kharazmi *et al*. 2012). In transgenes **J7**, **J7.1**, and **J7.2**, *lacZ* expression is abolished in the larval discs and brain (a-d) but retained in embryos and ovary (e-g). Truncation **J7.3** fails to express lacZ in any of the tested tissues. In the composite **J7.4** transgene containing Myc-CRMs P1/P2, P31/P32, and P37/P38, but lacking the intervening sequences between the first two elements (compared to J2.1), reporter activity is restored in the larval brain and leg disc, embryos, and ovary; however, expression in the eye and wing imaginal discs is weaker than that observed with the J2.1 transgene. **Abbreviations**: **a**, brain; **b**, wing disc; **c**, eye-antennal disc; **d**, leg disc; **e** and **f**, embryos; **g**, ovary. **Note**: The sequences and coordinates of the Myc-CRMs P1/P2, P31/P32, and P37/P38 are indicated in Supplemental Table S1.3 and S1.5. β-Gal staining was done overnight at 25°C on discs and brain taken from 3^rd^ instar larvae and adult female embryos and ovary (**g**). Scale bar in (**a–g**) indicates 100 µm.

As a next step, we tested the P1/P2 enhancer within the conserved regulatory region of the J2.1 transgene for its tissue specificity in larval brain and imaginal discs. Our approach was to fuse the J2.1 enhancer region harboring the P1/P2 element to the J7 promoter in-frame and test this J7.4 transgene for functionality in larval tissues (Figure 1F). Interestingly, staining of larval and adult female tissues revealed endogenous *Myc* patterning in all examined tissues. However, the expression in eye and wing discs was not as strong as observed with the J2.1 transgene. This result suggests that the upstream P1/P2 enhancer cluster is necessary for *Myc* patterning in larval tissues and that the proper distance between the enhancer and promoter is required for strong transcriptional activation, as seen in the J2.1 transgene. Furthermore, our findings indicate that the P1/P2 enhancer functions as a late developmental enhancer, active during the onset of differentiation in larval limb, eye, wing, and brain patterning, as well as during pupal development.

### II. Initiator-Downstream Promoter Element (Inr-DPE) requires the action of neighboring enhancer cluster for *Myc* patterning

Transcriptional interactions between enhancers and core promoters are crucial for gene expression in space and time during development (Kutach and Kadonaga 2000; Butler and Kadonaga 2001; Levine 2010; Shir-Shapira *et al*. 2019; Dreos *et al*. 2021; Yokoshi *et al*. 2022). Multiple DNA regulatory motifs have been identified, among which the TATA-box is the most extensively studied element (Goldberg *et al*. 1979; Kadonaga 2012). Additionally, a conserved TATA-less regulatory element, namely the Downstream Promoter Element (DPE), was first identified and extensively studied in *Drosophila* (Burke and Kadonaga 1997; Kadonaga 2002; Ohler and Wassarman 2010; Kadonaga 2012), and later in humans (Dreos *et al*. 2021). Previously, by scanning *Myc* 8 kb intron 2 sequences with the phylogenetic footprinting tools *cis*-Decoder-*EvoPrinter* (Brody *et al*. 2007; Yavatkar *et al*. 2008), we identified multiple conserved sequence blocks (Myc-CRMs) in this region. A search using the Lasergene Bioinformatic tool GeneQuest module identified a TATA-less initiator-promoter element comparable to the DPE consensus sequence found in many *Drosophila* genes; this element proved to be active unidirectionally (Kharazmi *et al*. 2012). The transgene J8, containing the 8 kb full-length intron 2 sequences within which the P29/P30 enhancer cluster and the TATA-less DPE promoter reside, drove *Myc-lacZ* expression in third instar larval imaginal discs, the brain, and adult female tissues (Figure 2A). The two truncations derived from the J8 transgene—J8.4, containing only the enhancer P29/P30, and J8.5, retaining only the DPE—failed to express *lacZ* in the tested tissues (Figure 2, B-C). These results suggested to us that neither the enhancer nor the DPE was capable of initiating a transcriptional burst independently. We hypothesize that the core promoter element DPE functions as a regulatory unit to modify the activity of the enhancer’s transcriptional dynamics throughout development. Indeed, a core promoter element can serve as a modulator of a developmental enhancer in differential gene activation during highly proliferative stages of early development (YOKOSHI *et al*. 2022).

**Figure 2.**
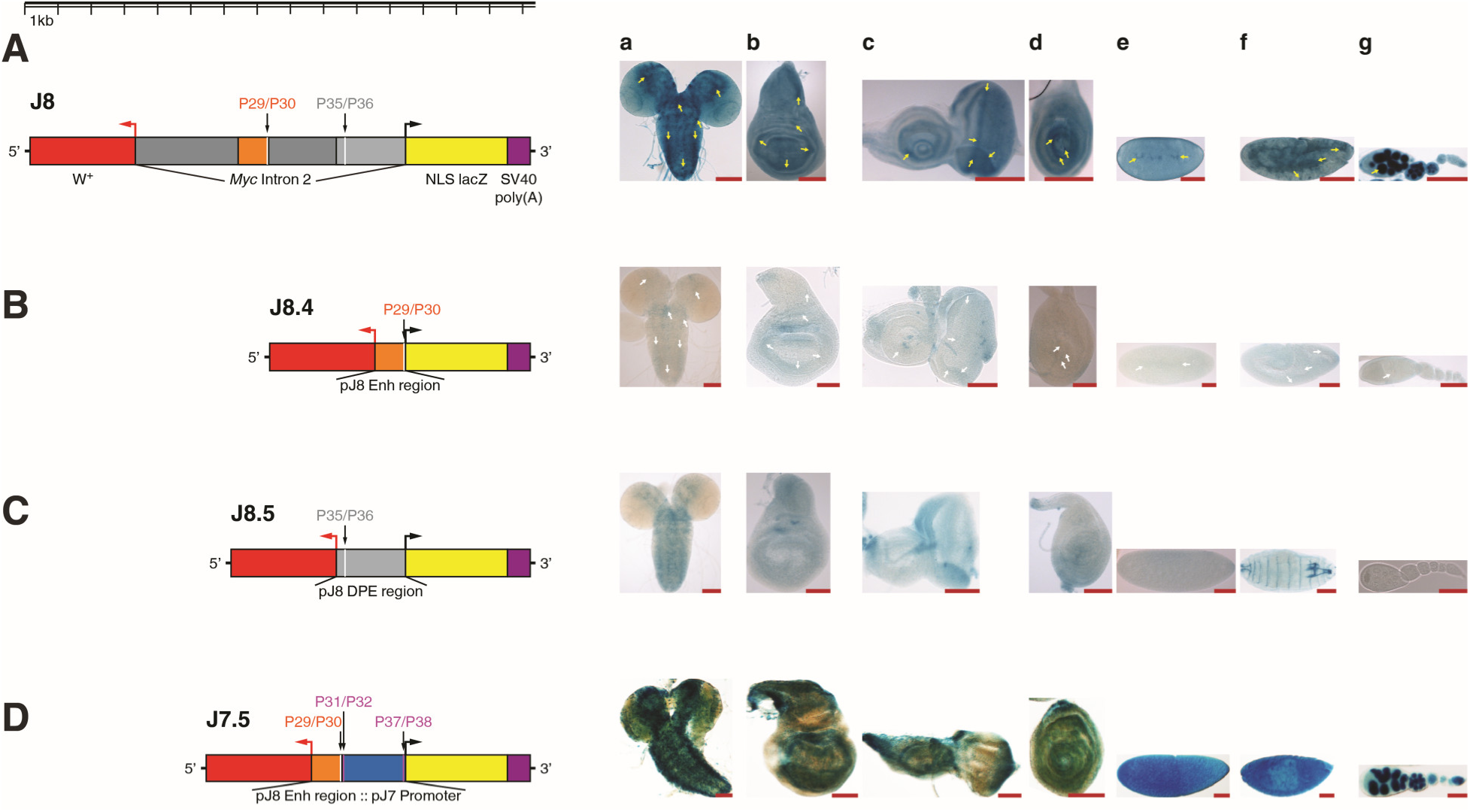
The Downstream Promoter Element (DPE) within the large intron requires the adjacent enhancer cluster P29/P30 for transcription activation. Reporters J8, J8.4, and J8.5 are adapted from the previous work (KHARAZMI *et al*. 2012). **(A)** J8 contains the 8 kb full-length intron 2 sequences (indicated by dark gray, orange, white, and light gray). The 932 bp enhancer region (indicated by orange) contains multiple conserved sequence blocks. The enhancer cluster P29/P30 (45 bp in length) resides within the enhancer region; P29/P30 contains a conserved cluster of 12 bp, 5’-CGCGTGGGAAAA-3’ (the Hairless binding site GTGGGAA is in the cluster). The 2.5 kb sequences at the 3’-end of intron 2 contain the DPE-containing Myc-CRM P35/P36 (47 bp in length). **(B)** J8.4 contains only the enhancer region (indicated by orange) including the enhancer cluster P29/P30 (indicated by white). **(C)** J8.5 contains only the downstream sequences of intron 2 (indicated by light gray) including the DPE-containing Myc-CRM P35/P36 (indicated by white). **(D)** The composite regulatory unit in the J7.5 transgene is under control of the J8 enhancer region containing the P29/P30 enhancer cluster fused upstream of the J7 promoter (indicated by white and blue). Panel (**a-g**) shows expression patterns of the *lacZ* reporter in the tested larval discs and brain (**a-d**), and in embryos and ovary (**e-g**). In the J8 transgene, expression in larval imaginal discs and brain (**a-d**) and in adult female embryos (**e-f**) and ovary (**g**) reflects the *Myc* endogenous mRNA patterning (yellow arrowheads) (KHARAZMI *et al*. 2012). The J8.4 truncation containing only the enhancer region and the J8.5 transgene under control of the DPE-containing downstream sequences do not mimic the endogenous *Myc* patterning (white arrowheads). The composite promoter in J7.5, consisting of the J8 enhancer region including the Myc-CRM P29/P30 subcloned upstream of the J7 promoter, rescued *lacZ* expression in larval discs and brain and adult female tissues compared to the expression observed with J8.4 and J8.5. **Abbreviations**: **a**, brain; **b**, wing disc; **c**, eye-antennal disc; **d**, leg disc; **e** and **f**, embryos; **g**, ovary. **Note**: The sequences and coordinates of the Myc-CRMs P29/P30, P31/P32, P35/P36, and P37/P38 are indicated in Supplemental Table S1.3 and S1.5. β-Galactosidase staining was performed overnight at 25°C on tissues dissected from 3^rd^ instar larvae (discs and brain) and adult females (embryos and ovary). Scale bar in (**a–g**) indicates 100 µm.

We next tested whether the transcriptional activity of the enhancer P29/P30 depends exclusively on a DPE element, or alternatively whether a TATA-box promoter might also serve as a regulatory module for *Myc-lacZ* expression in all tissues, similar to the full-length reporter construct J8 (Figure 2A). To address this, the enhancer P29/P30 was inserted upstream of the TATA-box promoter in the J7 context to generate the reporter J7.5 (Figure 1B, Figure 2D). Remarkably, J7.5 recapitulated almost all aspects of the endogenous *Myc* activity in larval brain and discs, as well as in early embryos and ovary (Figure 2D). For β-galactosidase staining of tissues from positive and negative control flies, see Figure S1a-g in Supplemental Figures.

### III. Selection of *Myc cis*-elements as targets for the identification of *Myc* regulators and W-CRM as a positive control

Previously we identified *in silico* conserved *cis*-Regulatory Modules (CRMs) and promoter sequences within the noncoding regions of the *Myc* locus that drove positive *lacZ* reporter expression in embryos, ovary, and larval brain and imaginal tissues (Kharazmi *et al*. 2012; Kharazmi and Moshfegh 2013). Based on this *lacZ* activity, we selected five oligonucleotides corresponding to functional Myc-CRMs as targets to test for the assembly of associated gene regulatory proteins. For the negative control, we used a scrambled oligonucleotide derived from the noncoding region of the *Mdm2* gene of Mus musculus strain C57BL/6J (details in Materials and Methods, Section IV). As a positive control, we used a binding site for the Wingless/Wg pathway effector dTCF/Pangolin (W-CRM) (Figure 3; Table S1.3, primers IR5-Pos1 and Az5-Pos2-IR3). dTCF/Pan is a final effector of the Wingless pathway driving target gene expression. This dTCF/Pan site consists of an HMG domain with a basic tail (BT), similar to the murine LEF-1 site, for binding the HMG domain (Archbold *et al*. 2014). *Drosophila* dTCF/Pan shares 92% homology with the HMG domain of LEF1. The HMG site is attached to a spacer followed by the C-clamp sequence (Archbold *et al*. 2014). Normally, the dTCF/Pangolin transcription factor is cytoplasmic; upon Wingless signaling activation, it translocates into the nucleus to drive *wingless/wg* target gene expression (Schweizer *et al*. 2003; Archbold *et al*. 2014). DNA-protein complexes formed on the *Myc* targets, as well as complexes formed with the positive and negative controls, were subjected to MS analysis for protein identification. The analysis procedure and results are presented in the following sections.

**Figure 3.**
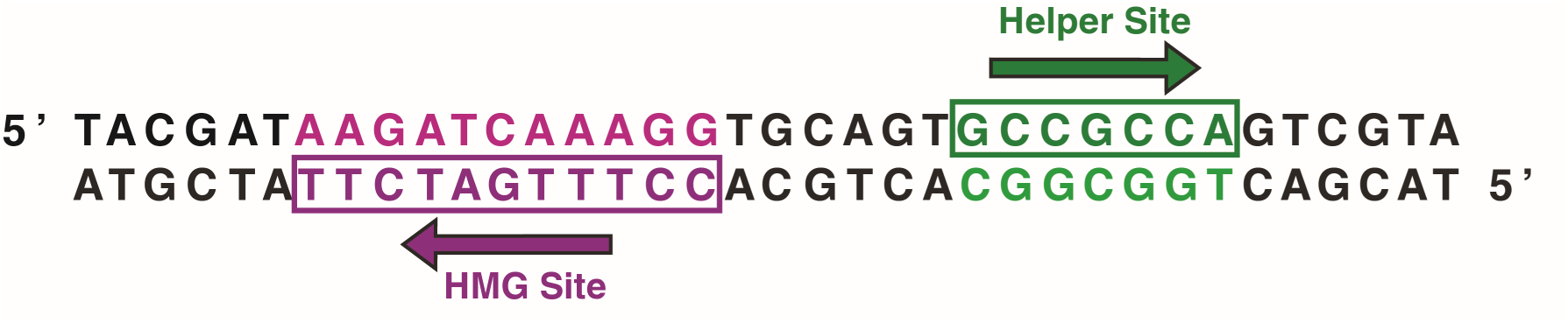
A variant of W-CRM containing HMG and Helper sites was used as positive control for protein identification studies. HMG-Helper site pair orientation has been termed Akimbo (AK) with six bp spacing (AK6) (Archbold *et al*. 2014). A right-pointing arrow indicates the consensus sequence is read 5’ to 3’ on the “top” strand; a left-pointing arrow indicates the consensus is read 5’ to 3’ on the “bottom” strand. The sequences shown for HMG (magenta) and Helper site (green) are identical to those used for DNA binding experiments and synthetic Wnt reporter constructs, and this sequence showed the highest binding affinity to TCF/Pan *in vitro*. Figure adapted from (Archbold *et al*. 2014).

### IV. Quality control of oligonucleotides and protein extracts prior to method development

*Myc* target sequences, positive control, and negative control oligonucleotides were annealed to form double-stranded molecules (Figure 4a-c, Table S1.5). Annealed oligonucleotides were pegylated (SHCHEPINOV *et al*. 1997) for further control experiments prior to protein identification (Figure 4d). To test whether the click chemistry reaction works and the azide moiety at the end of an oligonucleotide can be covalently bound to the DBCO group of DBCO-PEG23, we ran both pegylated and unpegylated oligonucleotides on a 5% PAGE gel and demonstrated that pegylated DNA migrates more slowly than unpegylated oligonucleotide (Figure 4e). Furthermore, through Electrophoretic-Mobility-Shift-Assay (EMSA) using the positive control, we showed that proteins associate with pegylated DNA as efficiently as non-pegylated DNA (Figure 4f). To determine optimal conditions for using modified oligonucleotides and nuclear extracts, subsequent control experiments involved keeping the oligo concentration constant while incrementally increasing protein concentration and observing the intensity of DNA-protein supershift bands. For the positive control, increasing the concentration of soluble cytoplasmic fraction (SCF) or soluble nuclear fraction (SNF) enhanced the intensity of supershifts containing DNA-protein complexes for both extracts, and the 2.25 picomole DNA was fully complexed by proteins in each case (Figure 4g, lane 2-9). Similar results were obtained for the *Myc* target P31/P32 with both protein fractions (Figure 4h, lane 2-7). However, although the amount of DNA used per lane (1.35 picomole) was less than for the positive control, the proportion of unbound oligonucleotides migrating near the bottom of the gel was higher and more visible compared to the positive control. Based on the intensity of supershifts observed with 100 µg and 150 µg of SCF (Figure 4h, lane 3-4) and SNF fractions (Figure 4h, lane 6-7), 200 µg of each fraction was used for each protein identification sample (see Material and Methods, section VII).

**Figure 4.**
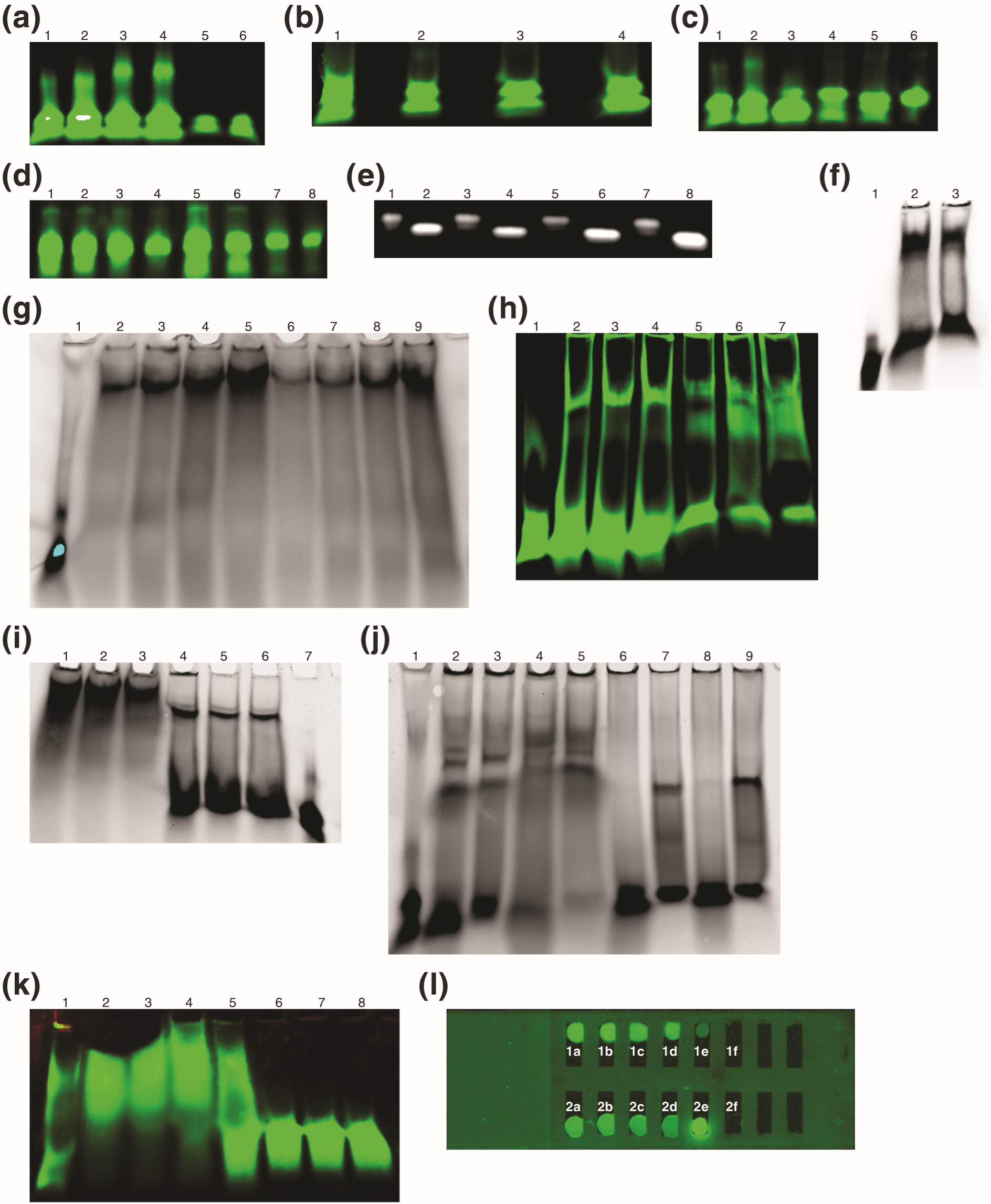
Annealing of the oligonucleotides to double-stranded molecules and EMSA assays after incubation with *Drosophila* embryonic protein extracts. (a) Annealing of IR-labeled positive control oligonucleotides. **1-4**: azide-labeled; **5-6**: no azide labeling. Amounts loaded: **1, 3, 5** = 0.9 pmol; **2, 4, 6** = 1.35 pmol. **(b) Annealing of IR- and azide-labeled Myc CRM oligonucleotides (1.35 pmol per lane)**. **1**: P31/P32; **2**: P32/P31; **3**: P37/P38; **4**: P38/P37. **(c) Annealing of various IR- and azide-labeled oligonucleotides (1.35 pmol per lane)**. **1**: Pos1/Pos2; **2**: Pos2/Pos1; **3**: Scm1/Scm2; **4**: Scm2/Scm1; **5**: P1/P2; **6**: P2/P1. **(d) Annealing and pegylation check for Myc CRMs**. **1, 3, 5, 7** = 0.45 pmol; **2, 4, 6, 8** = 0.225 pmol. Oligos: **1-2**: P31/P32; **3-4**: P32/P31; **5-6**: P35/P36; **7-8**: P36/P35. **(e) Comparison of migration**: unpegylated vs. pegylated positive control DNA. **1**: Pos2/Pos1-PEG23 (0.2 pmol); **2**: Pos2/Pos1 (0.4 pmol); **3**: Pos2/Pos1-PEG23 (0.3 pmol); **4**: Pos2/Pos1 (0.4 pmol); **5**: Pos1/Pos2-PEG23 (0.2 pmol); **6**: Pos1/Pos2 (0.4 pmol); **7**: Pos1/Pos2-PEG23 (0.3 pmol); **8**: Pos1/Pos2 (0.6 pmol). **(f) EMSA comparing SNF binding to non-pegylated vs. pegylated positive control (0.4 pmol per lane)**. **1**: Pos2/Pos1 DNA only; **2**: Pos2/Pos1 + 100 µg SNF; **3**: Pos2/Pos1-PEG23 + 100 µg SNF. **(g) EMSA titration: Binding of increasing SCF/SNF concentrations to positive control (Pos1/Pos2, 2.25 pmol per lane)**. **1**: DNA only; Lanes 2-5: + 100, 200, 300, 400 µg SCF; Lanes 6-9: + 100, 200, 300, 400 µg SNF. (h) EMSA titration: Binding of increasing SCF/SNF concentrations to *Myc* target P31/P32 (1.35 pmol per lane). **1**: P31/P32-IR3 only; **2-4**: + 70, 100, 150 µg SCF; **5-7**: + 70, 100, 150 µg SNF. **(i) EMSA titration**: Binding of decreasing SCF/SNF concentrations to *Myc* target P37/P38 (0.45 pmol per lane). **1-3**: + 500, 400, 300 µg SCF; **4-6**: + 500, 400, 300 µg SNF; **7**: P37/P38 DNA only. (j) EMSA specificity test: Binding of SCF/SNF to *Myc* target P1/P2 vs. Scm negative control (1.35 pmol per lane). **1**: Scm1/Scm2 only; **2, 4**: Scm1/Scm2 + 70, 150 µg SCF; **3, 5**: P1/P2 + 70, 150 µg SCF; **6, 8**: Scm1/Scm2 + 70, 150 µg SNF; **7, 9**: P1/P2 + 70, 150 µg SNF. **(k) Competition EMSA**: IR-labeled P1/P2 (hot probe, 1.35 pmol) vs. unlabeled P3/P4 (cold competitor) with 130 µg SNF. **1**: Probe only; **2**: Probe + SNF (no competitor); Lanes 3-8: Probe + SNF + increasing molar excess of competitor (**3**: 1x; **4**: 5x; **5**: 15x; **6**: 30x; **7**: 45x; **8**: 60x). **(l) Visualization of Bead-PEG23-DNA constructs (IR-labeled) on glass slide before protein binding**. Panels **1a, 1b**: Bead-PEG23-P37/P38 construct; Panels **1c, 1d**: Bead-PEG23-P38/P37 construct (3 µL loaded, from 0.3 pmol in 300 µL total). Panel **1e**: Raw beads (30 µg); Panel **1f**: Binding buffer only (30 µL). Panels **2a, 2b**: Supernatant from Bead-PEG23-P37/P38 prep; Panels **2c, 2d**: Supernatant from Bead-PEG23-P38/P37 prep (30 µL loaded, from 0.3 pmol input in 300 µL total). Panel **2e**: Free P37/P38 oligo (0.3 pmol); Panel **2f**: Bead buffer only (30 µL). Run conditions for gel in (e): 5% non-denaturing TBE PAGE (Bio-Rad), 2 hours at 40V. Run conditions for other gels (annealing checks and EMSAs): 5% non-denaturing TBE PAGE (Bio-Rad), 55 minutes at 100V.

In the next experiment, similar to the tests with the positive control and target P31/P32, we kept the oligo P37/P38 concentration constant but decreased the amount of SCF and SNF in increments, expecting to observe a decrease in the intensity of shifted bands for both extracts. For supershifts obtained with the SCF fraction, intensity decreased, and essentially all oligo molecules were associated with proteins, with no free DNA detectable near the bottom of the gel (Figure 4i, lane 1-3). For the SNF fraction, the supershifts became weaker with decreasing protein amounts, and a substantial portion of the DNA remained unbound and migrated near the bottom of the gel (Figure 4i, lane 4-6). This indicated that 0.45 picomole of oligonucleotide is adequate for isolating DNA-protein complexes for protein identification. To confirm the specificity of proteins forming supershifts with the *Myc* target, we performed SCF and SNF protein binding assays with the *Myc* target P1/P2, alongside the scrambled oligo (Scm1/Scm2, short Scm) as a negative control. The SCF fraction showed supershifts for both the negative control Scm (Figure 4j, lane 2 and 4) and the *Myc* target IR-P1/P2 (Figure 4j, lane 3 and 5). However, no binding was observed for the negative control Scm with the SNF fraction (Figure 4j, lane 6 and 8). For the *Myc* target IR-P1/P2 (Figure 4j, lane 7 and 9), the supershifts for both concentrations of 70 and 150 µg of SNF were evident and sharp, with the band intensity increasing with higher protein concentration. We conclude that the binding of factors to Scm is nonspecific. To further confirm the specificity of factor association in SNF with *Myc* sequences, we performed a competition EMSA with the *Myc* target IR5-P1/P2-IR3. Like other oligonucleotides used in this study, IR5-P1/P2-IR3 was end-labeled with the infrared dye Dyomics 781 (referred to as hot probe). An unlabeled primer pair P3/P4 (referred to as cold probe or competitor) shares the same sequence as IR5-P1/P2-IR3 (Table S1.3). With a 15-fold molar excess of competitor, the intensity of the hot probe IR5-P1/P2-IR3 began decreasing (Figure 4k, lane 5), and with a 30-fold molar excess of cold probe, the hot probe IR5-P1/P2-IR3 was fully competed out (Figure 4k, lane 6). This result further confirms the specific interaction between the DNA sequence and factors in the SNF fraction.

After performing these control experiments and obtaining positive results, we used the Soluble Nuclear Fraction (SNF), which is enriched with transcription factors, as the starting material for protein identification. The scrambled oligonucleotide (Scm) served as the negative control, and W-CRM dTCF/Pan served as the positive control. Since pegylated DNA effectively recruited factors to the DNA sequence of interest, the created chimeric Magnetic Bead-PEG-DNA pulldown construct (see Methods, Section VI for construction details) and controls were placed on a glass slide with chambers separated by silicon, and the IR-labeled DNA (in this experiment, oligo P37/P38) was visualized using a Licor InfraRed imager (Figure 4(l), panel 1a-1f and 2a-2f). For each target and control oligo, a corresponding pulldown construct was built and subsequently used to isolate factors complexed by the DNA (see Figure S2 A-D in Supplemental Figures).

### V. Magnetic Solid Surface Enrichment (MSSE) of *Myc* regulators and associated cofactors for identification by mass spectrometry

Affinity purification mass-spectrometry, footprinting, gel shift methodology, and similar approaches have often proven insufficient for reducing background noise and enhancing selectivity, sensitivity, and specificity of protein identification experiments in whole cell lysates by MS analysis. This obstacle is more evident for identifying factors present at low stoichiometric levels, such as microproteins, hormone peptides, membrane-bound receptors, and gene regulatory proteins. Methods involving immobilization of nucleic acid-protein or protein-protein complexes on a solid surface followed by washing off unbound candidates and extraction of enriched proteins decrease background noise caused by chimeric spectra and hence false discovery rate (FDR); consequently, the accuracy of identified proteins increases significantly with high confidence (LEITNER *et al*. 2010). In our study, we developed and used a chimeric Carboxylic Magnetic Bead-PEG-DNA construct (Figure 5; see also Materials and Methods, Section VI) to pull down proteins specifically enriched on the target DNA sequences. The hydrophilic PEG-spacer increases water solubility, reduces steric effects, provides adequate spacing between the DNA molecule and the magnetic bead, and potentially enhances protein binding to the target sequence (SHCHEPINOV *et al*. 1997). We tested this methodology for identifying *Myc* regulators, and it yielded optimal results with high selectivity and sensitivity in protein identification. Furthermore, this basic approach can be readily adapted for various experimental goals, including detecting other molecules such as antigens or antibodies. Finally, the construct could potentially be used to study specific protein-miRNA interactions by replacing the DNA oligonucleotide with a relevant miRNA sequence or structure, such as a hairpin.

**Figure 5.**
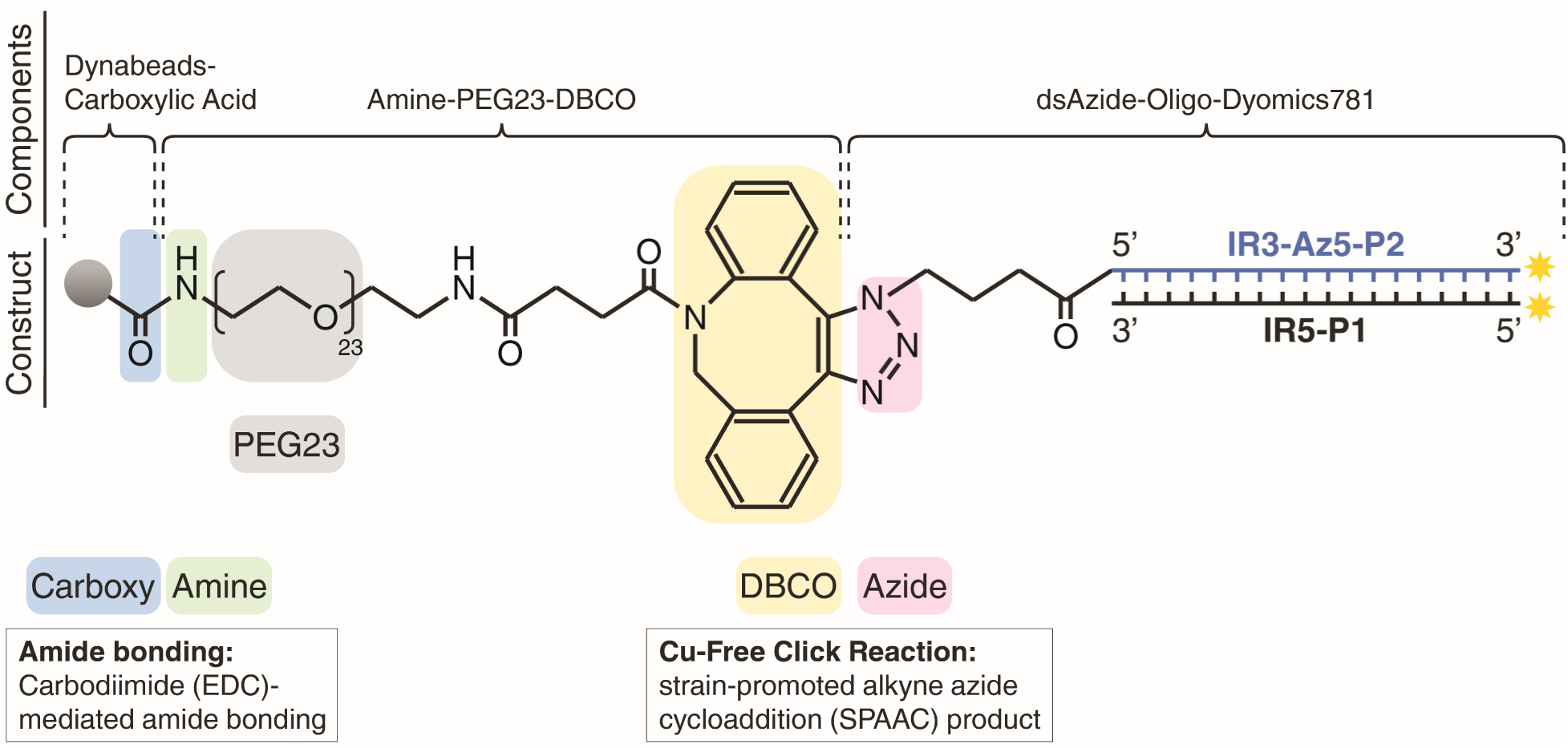
Magnetic Solid Surface Construct (MSSC) designed and used for protein enrichment. Amine-PEG23-DBCO is covalently bound to carbodiimide-activated microspheric Dynabeads® MyOne™ Carboxylic acid beads via covalent amide bonding. The chimeric structure underwent a copper-free click chemistry reaction (Jewett and Bertozzi 2010; Eeftens *et al*. 2015) for covalent bonding between its reactive alkyne, the DBCO moiety (Dibenzocyclooctyne), and the reactive Azide (N-hydroxysuccinimide (NHS) ester) end-modified oligonucleotide to yield the Bead-PEG23-Oligonucleotide construct. The resulting solid surface was used to biomagnetically separate and affinity-purify *Myc* regulators from *Drosophila* embryonic nuclear protein extracts (see Materials and Methods, Section VI for more detail).

As an example, this methodology could be applied to understand the role of oncogenes such as *MYC* in microRNA biology. Research has demonstrated that MYC plays a significant role in regulating oncogenic miRNAs, particularly the miR-17-92 complex (He *et al*. 2005; Treiber *et al*. 2017). The purpose of this study, as will be described in later sections, was to use the Bead-PEG23-Oligonucleotide construct to identify *Myc* transcriptional regulators.

### VI. Soluble Nuclear Fraction (SNF) and raw beads as components of *in vitro* DNA-protein interaction

Generally, proteins interact with DNA either by sequence-specific binding, as is the case for transcription factors, or they may bind non-specifically as is the case for some replication proteins (NILKANTA AND BAGCHI 2018). Several factors favor DNA-protein interactions, including hydrogen bonds, low salt concentration, appropriate pH, and the presence of divalent ions such as Zinc (Zn^2+^) and Magnesium (Mg^2+^) (Phizicky and Fields 1995; Dey *et al*. 2012). For *in vitro* studies of DNA-protein interactions, it is essential to maintain conditions mimicking the *in vivo* environment as closely as possible by performing experiments with non-denaturing agents. Before incubating different Myc-CRMs with SNF, we assessed increasing concentrations of the nuclear fraction for quality and quantity on a non-denaturing 5% Precast PAGE gel using low salt 0.5% TBE running buffer. After Coomassie Blue staining, a protein smear with increasing intensity was observed in each lane (see Figure S2E, 1-9 in Supplemental Figures). Previous studies have shown that in-gel digestion for protein quantification and/or identification by MS analysis often suffers from background noise, resulting in low sensitivity and selectivity in the obtained MS data and interpretation of the dataset (Goodman *et al*. 2018). A primary reason for poor result quality may be contamination of DNA-protein complexes with unbound background proteins migrating alongside the DNA-protein complex on the gel. We addressed this problem by switching to a magnetic bead solid surface support for protein identification. This approach enabled us to obtain highly selective datasets with low background noise and low false discovery rate (FDR) for all samples. Proteins identified by Mass spectrometric analysis and the processing of protein categories will be presented in the subsequent sections.

### VII. MS data obtained for raw beads, Scm, embryonic extracts, and the Myc-CRMs

The raw beads, crude nuclear extract, and DNA-protein complexes were subjected to MS analysis for protein identification. The processing and analysis of the obtained dataset were performed using Thermo Fisher Scientific™ Proteome Discoverer™ 2.5 software and the software tool ComplexBrowser (Material and Methods, Section IX). We have performed the exact same data analysis of the blank sample (raw beads), both control samples the Scrambled (Scm) oligonucleotide (Negative Control), the positive control (Pos), and the Myc-CRM samples.

The blank bead analysis identified 26 proteins annotated to *D. melanogaster* (see Table 2 and Image 2 in File S2). Twelve of these proteins showed low scores with low confidence of identification (ID). Additionally, the identification of 10 of these proteins was based on a single unique peptide, which would normally be excluded prior to publication. Three proteins (Actin, Albumin and ATPase) were identified by peptides that are not unique to *Drosophila melanogaster* and could represent contamination from a different source. One protein (Collagen alpha I(III) chain, Accession P04258) was unique to the Blank Beads and not found in SCF and SNF extracts or samples including Scm. This protein contains 1466 residues with a molecular weight of 138.4 kDa. The collagen contamination likely originates from the surface coating of the magnetic beads that leaches off during sample preparation (e.g., boiling). We tested the beads for protein contamination prior to their use for sample preparation. After treating the beads with 8 M urea, followed by boiling and running on a 1% SDS-PAGE gel, a few artifactual bands were visible upon intensive exposure of the gel to UV light (see Figure S2F in Supplemental Figures). Our overall conclusion is that raw beads can be effectively used for biomagnetic separation and affinity purification of proteins and nucleic acids, provided the collagen contaminant is filtered from the obtained MS data.

The original unbiased data file with the ungrouped replicates contained 3872 proteins (see Table 3 and Image 3 in File S3). Contaminants originating from the beads were removed from the original dataset (see Table 2 and Image 2 in File S2). Since the ComplexBrowser software requires ungrouped data input, we used the data file with ungrouped replicates for the ComplexBrowser analysis. In this data file, after applying the filter for *Drosophila* species and removing non-*Drosophila* contaminants (see Table 4 and Image 4 in File S4), we obtained 2224 proteins (see Table 5 and Image 5 in File S5). The unbiased data file with the grouped replicates resulted in 2264 proteins (see Table 6 and Image 6 in File S6).

### VIII. Heat map analysis of proteins associated with *Myc* targets and control sequences

The unbiased proteome result of 2264 proteins from the grouped replicates (see Table 6 and Image 6 in File S6) for the nuclear samples underwent multiple general data quality filters. One example filter was ‘mass over charge ratio for sample is > 10x higher than for the scrambled control’. Further filters included: a false detection rate threshold (’protein FDR confidence combined has at least level high’); a confidence threshold against the Scrambled negative control (’found in sample groups has at most confidence Peak Found in sample groups Scm’); and restriction to the *Drosophila* species proteome (’species map is true in species *Drosophila melanogaster* OX 7227’). As a result, this yielded 1001 protein candidates for further analysis (see Table 7 and Image 7 in File S7). These results were useful for excluding contaminants and served as a reference for correlating protein binding with the functional activity of Myc-CRMs previously characterized by phylogenetic footprinting (Kharazmi *et al*. 2012) and *Myc-lacZ* expression profiles (Figure 1, A-F and Figure 2, A-D). We processed the results for target-associated proteins identified using pegylated DNA (Figure 4(f)) by generating heat maps (see Figure S3-S4 in Supplemental Figures) using Proteome Discoverer 2.5 (Thermo Scientific) software and looked for clusters of proteins potentially related to the observed *Myc*-*lacZ* expression patterns in tested tissues (Figure 1 and Figure 2). Although the *in vitro* association of factors with Myc-CRMs will require future *in vivo* validation, the patterns observed *in vitro*, when considered alongside the expression analysis results, suggest specificity for the tested targets as potential *Myc* regulators.

### IX. Functional classification of proteins associated with Myc-CRMs and the positive control W-CRM

Following quality control and visualization of functionally similar sample pairs (Figure 6-9), the proteins associated with each *Myc* target pair and the positive control pair were manually searched against databases (see Methods IX) to identify the protein complexes to which specific factors belong (Table S8). Based on biological activity and molecular function, the identified proteins were categorized and subdivided into groups (Figure 10; see also supporting information, Table A-H in File S9). We differentiated between unique factors associated with each target and those associated with more than one tested target. We confirmed the identification of factors previously known from the literature to control *Myc* expression, corroborating their roles as relevant *Myc* regulators. Such factors include, but are not limited to, components of the Hippo, Wnt/Wg, Dpp, Hedgehog, and Notch signaling pathways. Factors newly discovered or not yet fully characterized in this context represent potential candidates for future studies, such as *in vivo* biochemical assays and structural biological studies. In the heat map grouping (see Figure S3-S4 in Supplemental Figures), the *Myc* distal enhancer P1/P2 and the positive control, dTCF/Pan site, fall into the same group. In a previous study and the present work, we demonstrated that the absence of upstream regulatory sequences containing the P1/P2 enhancer cluster leads to abrogation of expression in larval imaginal tissues and brain (Kharazmi *et al*. 2012) (Figure 1B).

**Figure 6.**
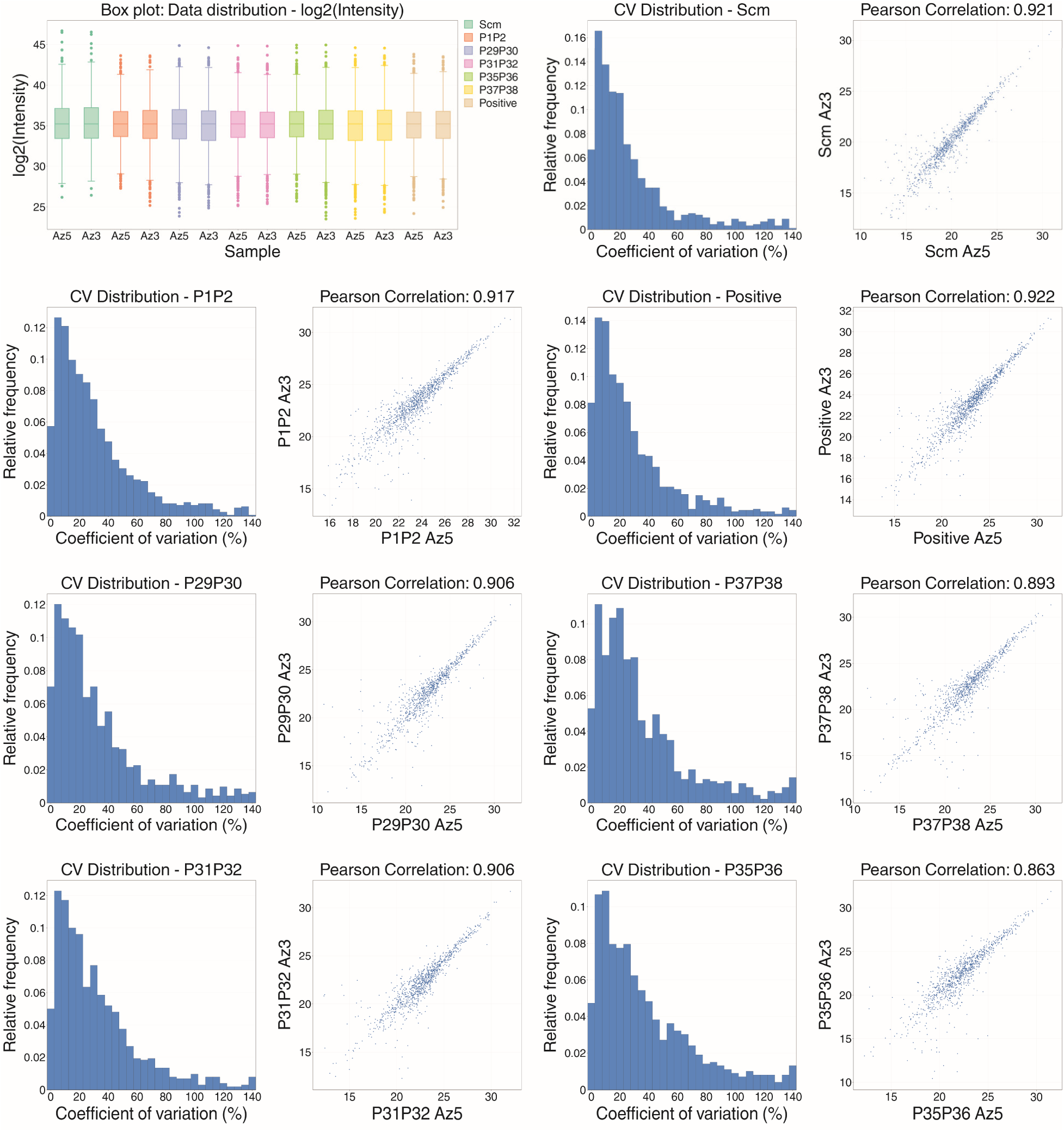
Quality control analysis visualization using ComplexBrowser for label-free proteomics data. Box plots show log2-transformed intensities for identified proteins across duplicate samples for the negative control (Scm), Myc-CRMs, and positive control (Positive). Histograms show the Coefficient of Variation (CV) distribution for protein intensities between replicates for each sample condition (Scm, Positive, P1P2, P29P30, P31P32, P35P36, and P37P38). Scatter plots compare log2 intensities between the two replicates for each of the seven sample conditions, with Pearson correlation coefficients shown. **Note**: The Az5 and Az3 designations associated with sample names represent the end-modification of the oligonucleotide at its 5’-end and 3’-end, respectively (see Methods, Section IV for details). The Azide (N-hydroxysuccinimide (NHS) ester)) moieties were removed from the DNA-protein complexes during protein purification prior to MS analysis. See Methods, Sections IV, VI, and VIII for further details on oligonucleotide design and protein sample preparation.

**Figure 7.**
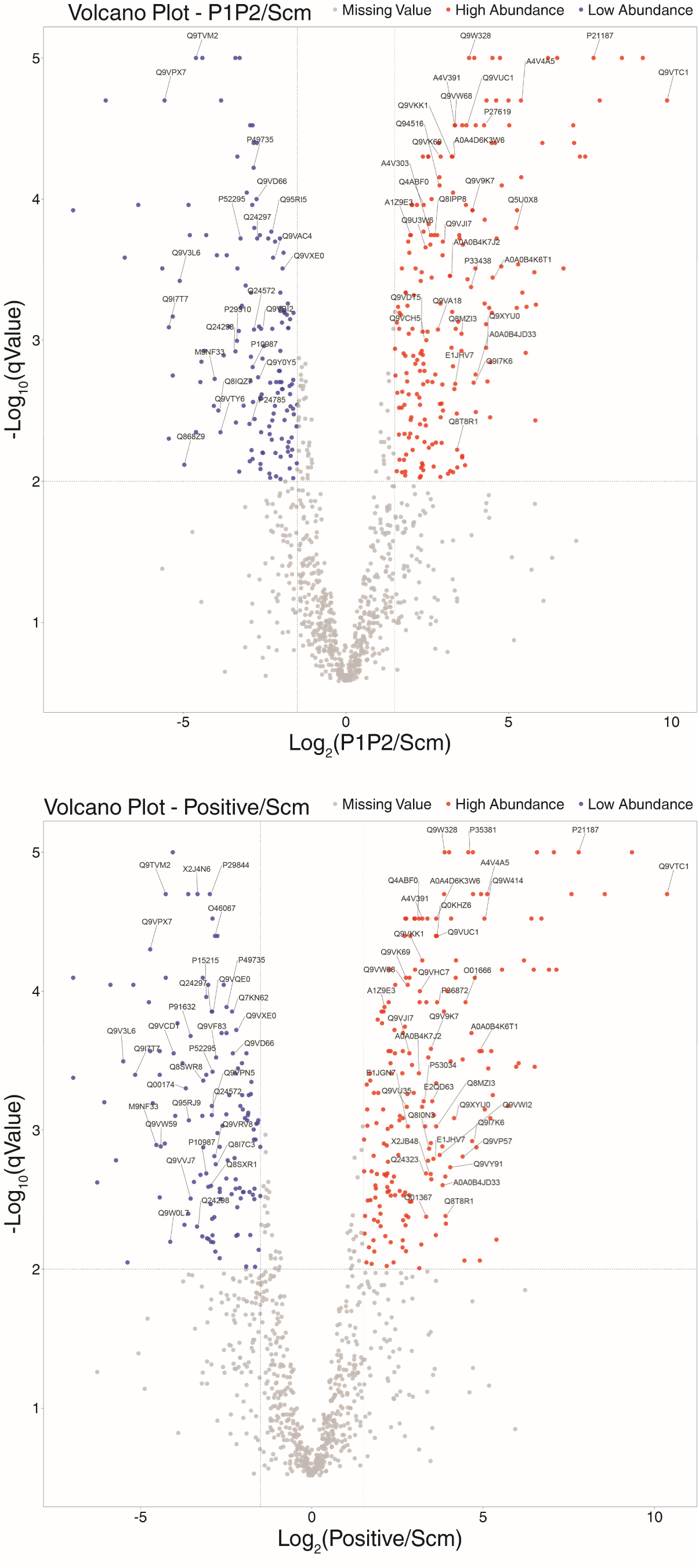
Volcano plots comparing Myc-CRM P1/P2 and positive control samples to the negative control. Volcano plots display log2(abundance ratio) comparing the indicated sample group (P1/P2 or Positive) versus the negative control (Scm) on the x-axis, plotted against −log10(q-value) on the y-axis. Significance thresholds are indicated by lines corresponding to q-value < 0.01 (−log10(q-value) > 2) and absolute abundance ratio > 1.5.

**Figure 8.**
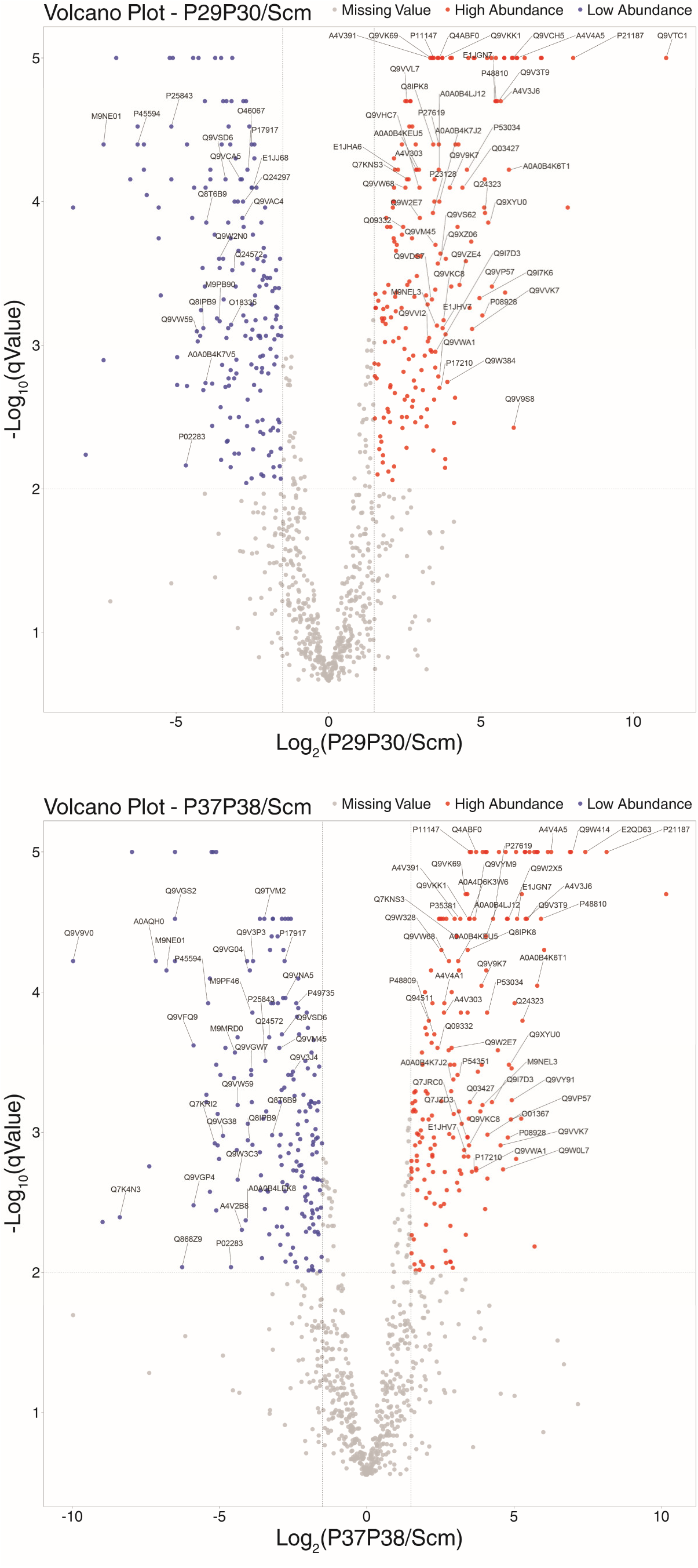
Volcano plots comparing Myc-CRM P29/P30 and P37/P38 samples to the negative control. Volcano plots display log2(abundance ratio) comparing the indicated sample group (P29/P30 or P37/P38) versus the negative control (Scm) on the x-axis, plotted against −log10(q-value) on the y-axis. Significance thresholds are indicated by lines corresponding to q-value < 0.01 (−log10(q-value) > 2) and absolute abundance ratio > 1.5.

**Figure 9.**
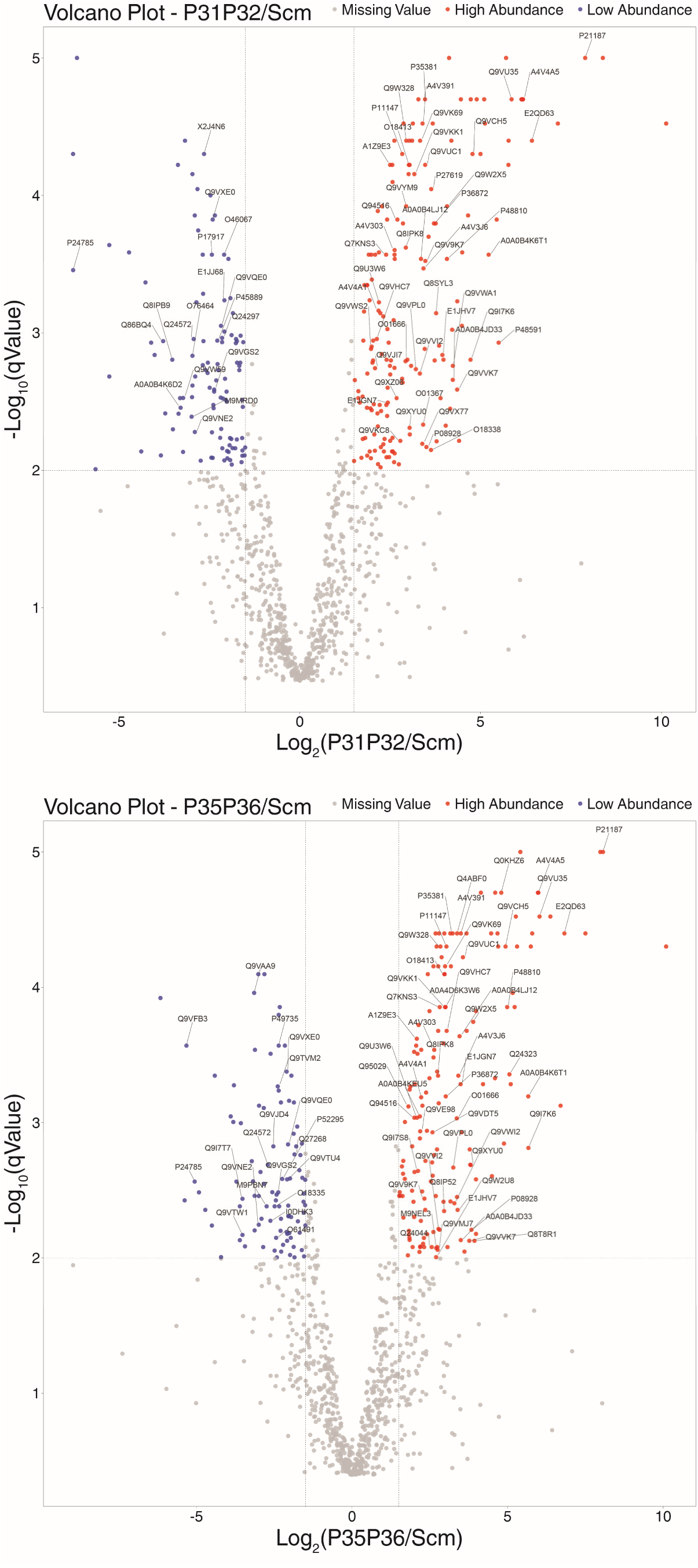
Volcano plots comparing Myc-CRM P31/P32 and P35/P36 samples to the negative control. Volcano plots display log2(abundance ratio) comparing the indicated sample group (P31/P32 or P35/P36) versus the negative control (Scm) on the x-axis, plotted against −log10(q-value) on the y-axis. Significance thresholds are indicated by lines corresponding to q-value < 0.01 (−log10(q-value) > 2) and absolute abundance ratio > 1.5.

**Figure 10.**
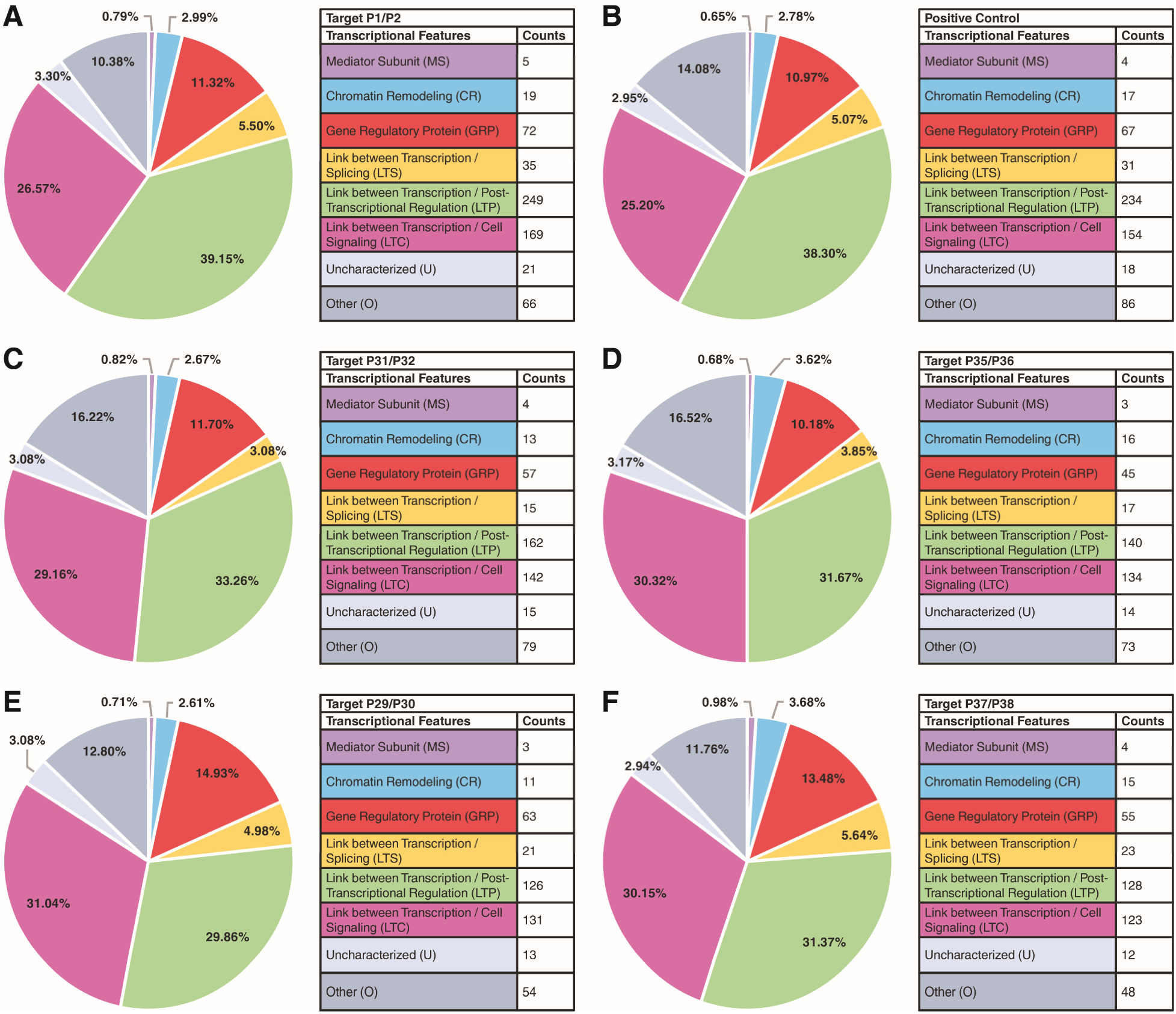
Functional classification of factors associated with Myc-CRMs and the positive control. Sample pairs identified as functionally similar by Heat Map analysis (see Figure S3-S4 in Supplemental Figures) of the filtered high-confidence proteins (see Table 7 and Image 7 in File S7) are depicted adjacent to each other. Analysis of the 1001 regulatory factors revealed the following approximate percentage ranges across the different samples: 0.65-0.98% belong to the mediator complex; 2.61-3.68% to chromatin remodeling; 10.18-14.93% are gene-specific proteins; 3.08-5.64% belong to the splicing machinery; 29.86-39.15% are posttranscriptional regulators; 25.20-30.32% are cell signaling components; 2.94-3.30% comprise uncharacterized factors; and 10.38-16.52% fall into other regulatory factor categories not included above. **A**: Upstream 5’-end P1/P2 enhancer cluster; **B**: Positive Control (W-CRM); **C**: TATA-Box-containing Myc-CRM P31/P32; **D**: DPE-containing Myc-CRM P35/P36; **E**: Intron 2 enhancer cluster P29/P30 (near DPE); **F**: 5’-UTR containing the P37/P38 enhancer cluster (oocyte element). See Table A-H in File S9 for detailed pie chart data analysis.

It has been shown that Wnt morphogens play a crucial role during early embryogenesis but also at later stages of development, including differentiation, embryonic segmentation, organogenesis, and tissue homeostasis (Herranz *et al*. 2008; Archbold *et al*. 2014; Stroebele and Erives 2016; Franz *et al*. 2017). We hypothesize that the Myc-CRM P1/P2 is important for larval development and organ shape during morphogenesis, and therefore we searched for factors supporting this role. The two enhancer elements P1/P2 and dTCF/Pan show similar activity profiles, being active during early cell proliferation and later during pattern formation. Another important similar group comprises the two initiation sites, namely the TATA-Box-containing target P31/P32 and the DPE-containing *Myc* target P35/P36. Experiments with *Myc-lacZ* reporter assays demonstrated that P31/P32 and the DPE function as two separable and important transcription units within the *Myc* gene (Kharazmi *et al*. 2012; Kharazmi and Moshfegh 2013), a result consistent with the heat map grouping in this study (see Figure S3-S4 in Supplemental Figures). The two developmental super enhancers, the 5’-UTR P37/P38 and the downstream P29/P30, form another similar group. However, reporter studies showed that the activity of P29/P30 is dependent on either the DPE-containing P35/P36 *cis*-element (Figure 2A) or the TATA-containing promoter P31/P32 (Figure 2D), whereas P37/P38 can function autonomously and express *Myc* during periods of high cell proliferation in early embryos and ovary (Figure 1, C-D). In the heat map, the starting material (SNF embryonic extract, representing the total input protein pool) clusters near the P1/P2 and positive control group, which are associated with a relatively high number of factors compared to other target groups (see Figure S3-S4 in Supplemental Figures). The negative control (Scm) clusters near the P31/P32 and P35/P36 group, which associated with fewer proteins compared to the enhancer groups (P1/P2, P29/P30, P37/P38) (see Figure S3-S4 in Supplemental Figures).

### X. Regulatory factors associated with the positive control W-CRM

A major group of canonical Wnt/β-catenin regulators belong to the T-Cell Factor (TCF) family of proteins (Cadigan 2012). The DNA recognition motif for TCF/Pangolin containing an HMG-Helper site was reported to respond to Wingless signaling in a tissue-dependent manner (Archbold *et al*. 2014), making it an ideal candidate for our positive control.

The potential Wnt regulators associated with the W-CRM in our analysis (Figure 11; see also supporting information, Table 3 in File S3 and Table S8) included: casein kinase IIα (*CKIIα*), known for roles in the cell cycle (Ducat *et al*. 2008) and regulation of Hedgehog and Wnt signaling (Glover *et al*. 1983; Dahmus *et al*. 1984; Willert *et al*. 1997; Siomi *et al*. 2002; Jia *et al*. 2010) during proliferation, differentiation, and tissue patterning (Karandikar *et al*. 2004; Bose *et al*. 2006; Wong *et al*. 2011);Sorting nexin 6 (*Snx6*), a component of the retromer/tubulation complex (Gene Ontology Curators 2002) and a positive regulator of Wnt secretion (Zhang *et al*. 2011); Galla-1 (*galla-1*), a component of the crb-galla-Xpd (CGX) complex required for proper mitotic chromosome segregation, cell proliferation in developing wing discs, and cell polarity (Yeom *et al*. 2015); the adaptor molecule Cullin 1 (*Cul1*), part of the Cullin-RING and SCF ubiquitin ligase complexes (Smelkinson *et al*. 2007) involved in ubiquitinating cell cycle-specific proteins (Sarikas *et al*. 2011) and negatively regulating the Wnt pathway (Swarup and Verheyen 2011; Roberts *et al*. 2012); Twins (*tws*), a component of the TWS-Protein phosphatase Pp2A complex that regulates Wnt signaling by stabilizing Armadillo/TCF during transcription (Bajpai *et al*. 2004), and influences cell fate specification and patterning in the wing disc (Uemura *et al*. 1993); Another transcription unit (*Atu*), involved in forming the nucleoplasmic Wg/Wnt transactivation complex, pol II pre-initiation, transcription elongation (GO Reference Genome Project 2011), and hemocyte proliferation (Sinenko *et al*. 2010); and Rap1 GTPase (*Rap1*), a small GTPase from the Ras superfamily with important roles in planar cell polarity-related morphogenetic events including imaginal disc patterning (O’Keefe *et al*. 2009), border follicle cell migration (Chang *et al*. 2018), and response to cAMP signaling (GO Reference Genome Project 2011).

**Figure 11.**
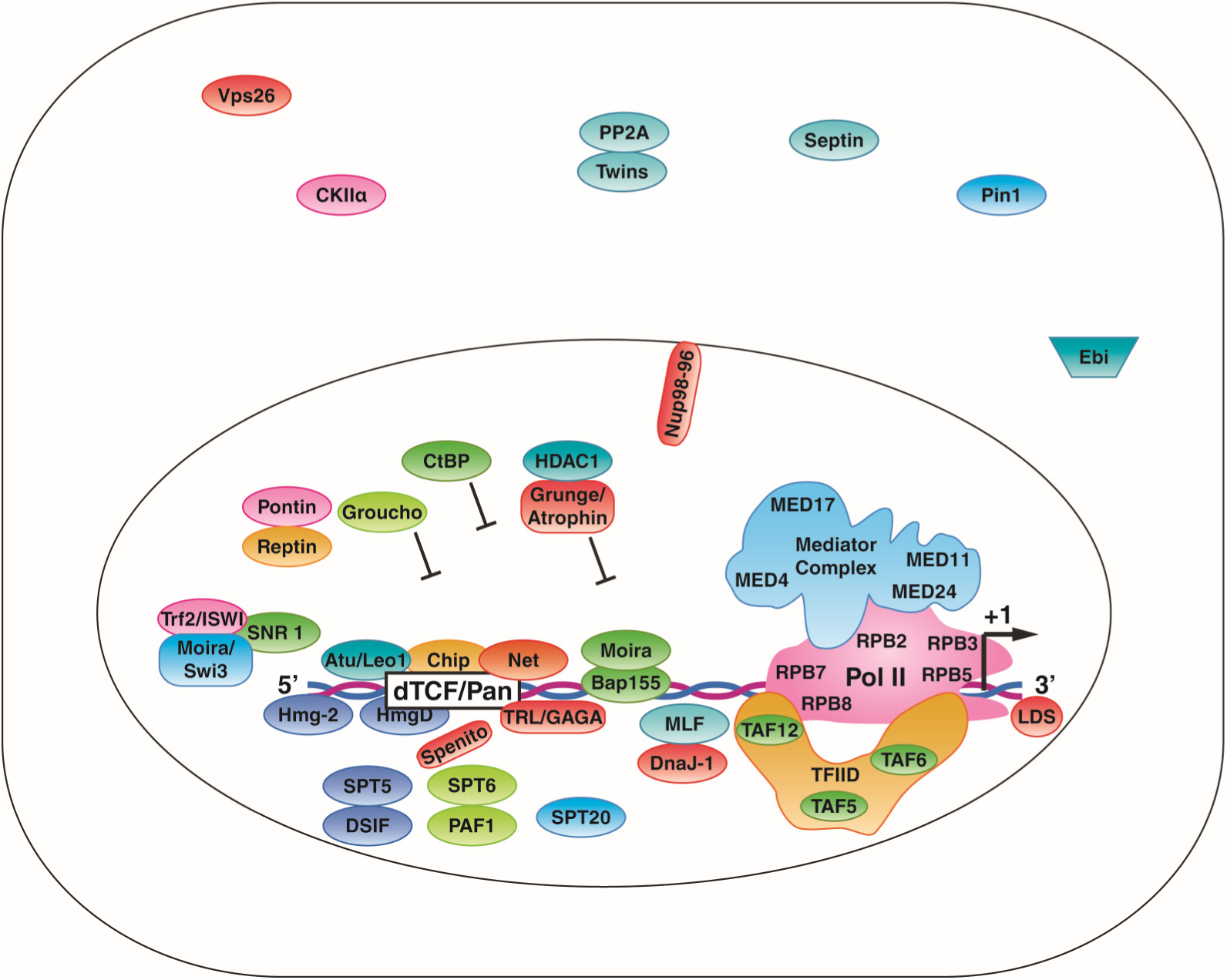
Schematic illustration of some identified factors associated with the positive control (W-CRM). These factors include known Wnt/Wg regulators such as High Mobility Group proteins HmgD and Hmg-2; transcription factors like Myeloid leukemia factor (Mlf), Chip (chi), Another transcription unit (Atu), Net (Net), Spenito (nito), and Trithorax-like (Trl); general repressors CtBP, Groucho, and Atrophin-Grunge (Gug, Atro), considered components of the Wnt enhanceosome; as well as components of the Mediator complex, RNA pol II subunits, and the TFIID complex (see Table 3 and Image 3 in File S3, Table S8). MED24 was exclusively associated with the Heat Map pair comprising the positive control and the *Myc* upstream enhancer cluster P1/P2. MED24 participates in larval salivary gland histolysis during metamorphosis, pupal development (WANG *et al*. 2010), and the response to ecdysone signaling (IHRY AND BASHIRULLAH 2014).

Some of the identified transcriptional regulatory factors have previously been reported as components of the Wnt enhanceosome: the cofactor Chip (*chi*) (Fiedler *et al*. 2015; van Tienen *et al*. 2017), the E-box binding repressor Net (*Net, CG11450*) (Brentrup *et al*. 2000; Peyrefitte *et al*. 2001), the global corepressor Groucho (*gro*) (Cavallo *et al*. 1998; Arce *et al*. 2009), and the corepressor Atrophin-Grunge (*Gug, Atro*) (Haecker *et al*. 2007). Armadillo (a segment polarity protein) was identified in the initial list but did not meet enrichment criteria in this analysis.

#### Transcription factor Mad, myeloid leukemia factor Mlf, Misshapen Msn, and Sumo were among the factors associated with the positive control

Mothers against dpp (Mad) is a conserved transcription factor and a component of the BMP/Dpp signaling pathway involved in inducing cell cycle arrest (Firth and Baker 2005) during morphogenesis and organ development (Lecuit *et al*. 1996; Wiersdorff *et al*. 1996; Marquez *et al*. 2001; Kirilly *et al*. 2005; Dworkin and Gibson 2006). Mad is required by Wingless signaling during wing disc morphogenesis and embryonic segmentation (EIVERS *et al*. 2009). Mad functions as a transcription repressor and interacts with components of Wingless signaling, such as Schnurri (*Shn*), during fate specification and patterning (Yao *et al*. 2006).

Myeloid leukemia factor (*Mlf*) interacts with DNA-bound transcription factors and protein-folding chaperones (Miller *et al*. 2017) and modulates transcription via interaction with the Hedgehog regulator Su(fu) during morphogenesis, cell proliferation (Fouix *et al*. 2003), and blood cell development (Bras *et al*. 2012; Miller *et al*. 2017). Whether Mlf directly regulates Wnt/Wg signaling during development and disease remains to be determined.

The *Drosophila* Sumo protein and the protein kinase Misshapen (*Msn*) might be potential regulators of the Wnt/Wg signaling pathway. Small ubiquitin-like modifier (*Sumo*, *smt3*), the sole member of the Sumo family proteins in *Drosophila*, is involved in epigenetic control of transcription via ubiquitin-like conjugation of proteins (Cappadocia and Lima 2018; Yau *et al*. 2020). Sumo responds to TGF-β signaling (MILES *et al*. 2008) during embryonic patterning (Nie et al., 2009). Furthermore, Sumo functions as a Hedgehog signaling modifier (Zhang *et al*. 2017), and during wing disc morphogenesis interacts with the protein kinase Misshapen and several other signaling pathways, including Wingless, to establish boundary formation between different cell populations in the developing wing disc (Rosales-Vega *et al*. 2023).

#### Chromatin modifying factors may also act as potential regulators of Wnt/Wg

Beyond the transcriptional regulators described above, our study provides strong evidence that regulation of *Wnt/wg* expression involves chromatin modification. Wnt/Wg signaling facilitates diffuse chromatin architectural changes at target gene loci, and chromatin and histone modifying enzymes have been documented to interact with and regulate the Wnt/Wg signaling pathway during development and disease (Liu *et al*. 2008; Parker *et al*. 2008; Curtis and Griffin 2012; Konig and Shcherbata 2015; Takahashi and Takada 2023). In our analysis, we identified a subset of chromatin and histone modifiers associated with the positive control W-CRM. These associated factors and their corresponding protein complexes likely function as potential *Wnt/wg* regulators.

#### High mobility group proteins HmgD and Hmg-2 were found associated with the tested W-CRM and are known to induce chromatin changes

High mobility group (HMG) protein D (HmgD), a Wnt/Wg signaling effector (Archbold *et al*. 2014), induces changes to chromatin-DNA architecture during gene expression (Wagner *et al*. 1992; Churchill *et al*. 1995; Dragan *et al*. 2003). Hmg-2 (*Hmg-2*), another High mobility group protein, is involved in chromatin-DNA structural changes and gene expression (GO Reference Genome Project 2011). Both HmgD and Hmg-2 likely participate in regulating *Drosophila wingless*/*wg* gene expression.

The X-linked wings apart-like (*wapl*) protein is involved in chromatin organization and preservation of chromosome structure by facilitating sister chromatid adhesion in specific heterochromatic regions, which is essential for normal mitosis (Verni *et al*. 2000). Wapl functions as a Wnt/Wg modifier during wing size determination in the Mediterranean fruit fly (medfly) (Cho *et al*. 2013).

#### Components of the NuRF, SWI/SNF (Brahma), and Cdc73/PAF1 chromatin remodeling complexes were identified and are potentially involved in regulating Wnt/wg target genes

Nucleosome remodeling factor NURF-38 (Hamiche *et al*. 1999; Mizuguchi *et al*. 2001) interacts with the NuRF complex and is required for homeotic gene expression and larval hematopoiesis (Sinenko *et al*. 2010). Armadillo, a cofactor of Wingless signaling, has been shown to interact with and recruit the NuRF complex to promoters of Wingless signaling target genes to positively regulate their transcription in cell cultures, suggesting a potential role for NURF-38 in regulating *wingless/wg* transcription (Song *et al*. 2009).

Snf5-related 1 (Snr1), a component of the SWI/SNF chromatin-remodeling complex (Brahma), works alongside the Wnt/Wg signaling pathway to induce higher-order chromatin remodeling, establishing gene-specific expression patterns and controlling growth and morphogenesis (Parker *et al*. 2008; Ramanujam *et al*. 2021).

Moira, BAP155 (*mor*, *CG18740*), a Swi3 component of the Brahma complex, regulates trithorax homeotic gene transcription (Crosby *et al*. 1999). Moira participates in mitotic cell cycle regulation (Brumby *et al*. 2002), partly via negative regulation of Armadillo-dTCF target genes in *Drosophila* (Collins and Treisman 2000) and TCF/β-catenin targets in mammals (Nollet *et al*. 1999), as well as wing disc margin morphogenesis (Terriente-Felix and de Celis 2009; Curtis *et al*. 2011).

Antimeros (*atms*) and Another transcription unit (*Atu*), as components of the Cdc73/PAF1 complex, interact with the basal transcription machinery and bound transcription factors to regulate histone modifications and pol II pause release (Adelman *et al*. 2006). Atu participates in the assembly of the Wnt/Wg transactivation complex and the initiation of pol II transcription (GO Reference Genome Project 2011), suggesting a potential role for Antimeros as a regulator of *Wnt/wg* targets.

C-terminal binding protein (*CtBP*), isoform G (A0A0B4KHB2) and isoform F (A0A0B4KGG9) interact with repressive transcription factors and block developmental enhancers during transcription (Zhang and Arnosti 2011). CtBP regulates *Wnt/wg* target genes during embryonic segmentation and wing disc morphogenesis (Fang *et al*. 2006; Bhambhani *et al*. 2011), suggesting a potential role for this protein in the regulation of *wingless/wg* transcription.

#### The SNF2L nucleosome remodeling factor Imitation SWI (Iswi), a known downregulator of wingless target genes, might also regulate wingless transcription

*Iswi*, a component of the NuRF, ACF, CHRAC, and RSF protein complexes (Corona *et al*. 2000; Eberharter *et al*. 2001; Badenhorst *et al*. 2005), is involved in pol II transcription via chromatin remodeling (Ito *et al*. 1999; Burgio *et al*. 2008; Emelyanov *et al*. 2010), chromatin assembly (Ito *et al*. 1997; Ito *et al*. 1999), and nucleosome displacement (Ito *et al*. 1997; Varga-Weisz *et al*. 1997) during homeotic gene expression and development (Badenhorst *et al*. 2002; Doyen *et al*. 2015). Iswi represses *wg* target genes by localizing to TCF binding sites within target gene sequences (Liu *et al*. 2008). This suggests that Iswi together with TCF might regulate Wingless transcription.

#### Components of the extracellular matrix (ECM), such as secreted heparan sulfates, may act as cofactors in the Wnt/Wg signaling pathway

ECM protein Perlecan, encoded by terribly reduced optic lobes (*trol*), is a secreted heparan sulfate that plays crucial roles in cell fate specification, tissue patterning, and maintenance of apico-basal polarity in epithelial cells (Schneider *et al*. 2006). It is involved in multiple signaling cascades, including Hedgehog (Park *et al*. 2003b), Wnt/Wg, and TGF-β (Lindner *et al*. 2007). Trol might therefore play a role in regulating *wingless/wg* expression.

#### Molecules involved in morphogen transport might also be potential regulators of Wnt/wg expression

Adaptor protein complex 2, α subunit (*AP-2α*) is involved in WNT5A-dependent internalization of FZD2, FZD4, and FZD5 in the noncanonical Wnt signaling pathway (Narayanan and Ramaswami 2001).

Vacuolar protein sorting 26 (*Vps26*) is a component of the retromer complex and is required for tetrameric cycling and sustaining Wingless and Dpp morphogen gradients during cell fate specification and tissue patterning (Port *et al*. 2008).

Phosphomannomutase type 2 (Pmm2) enzyme is involved in the synthesis of GDP-mannose and dolichol-phosphate-mannose. It functions as an effector molecule of the Wingless pathway for the correct positioning of wings in adult flies by maintaining N-linked glycosylation of the synaptic Mmp2 protein at the neuromuscular junction (NMJ) through Dlp/Wg signaling during development (Parkinson *et al*. 2016). AP-2α, Vps26, and Pmm2 likely play important roles in the regulation of the *wingless/wg* gene.

### XI. Factors associated with the investigated Myc-CRMs based on their molecular activities and biological processes

The *MYC* super-oncogene belongs to a family of regulatory genes controlling the expression of approximately 15% of transcribed genes (Orian *et al*. 2005; Caforio *et al*. 2018; Dhanasekaran *et al*. 2022). Among the factors identified associated with Myc-CRMs, we focused on identifying potential *Myc* regulators. Based on the results obtained from the heat map (see Figure S3-S4 in Supplemental Figures) and the Pie Chart analysis (Figure 10; see also supporting information, Table A-H in File S9), we subdivided the putative *Myc* regulators into different categories according to their roles at various levels of *Myc* expression. With our novel protein purification strategy, we captured most ubiquitous and tissue-specific factors, including components of the RNA polymerase transcription regulator complex such as general transcription factors (GTFs), mediators/cofactors, polymerases, gene-specific factors, pol II-specific transcription factors, chromatin/histone modifiers, and posttranscriptional regulators like RNA processing and gene silencing factors that play essential roles in controlling *Myc* mRNA levels and protein biogenesis during development and disease (Figure 12; see also supporting information, Table A-G in File S10). Among these subdivisions, many kinases were identified; these are potential targets of the transcription factor Myc, suggesting they may also function as *Myc* regulators (Boi *et al*. 2023). For each category, we will discuss the relevance of factor association with specific Myc-CRMs, their identities as potential *Myc* regulators based on known *Myc* biology, and their connections to developmental signals converging on the *Myc* promoter and cell cycle regulation. The ultimate goal is to characterize the early and/or late developmental enhancer elements and their trans-acting factors controlling *Myc* expression during cell growth, proliferation, and differentiation. Additionally, we will screen for potential new transcription initiation sites besides the previously identified TATA-containing P1 promoter and downstream DPE-containing P2 promoter (Kharazmi *et al*. 2012). Our analysis will also consider factors controlling *Myc* posttranscriptionally to maintain the balance between *Myc* mRNA levels, translation, ribosome biogenesis, and protein turnover.

**Figure 12.**
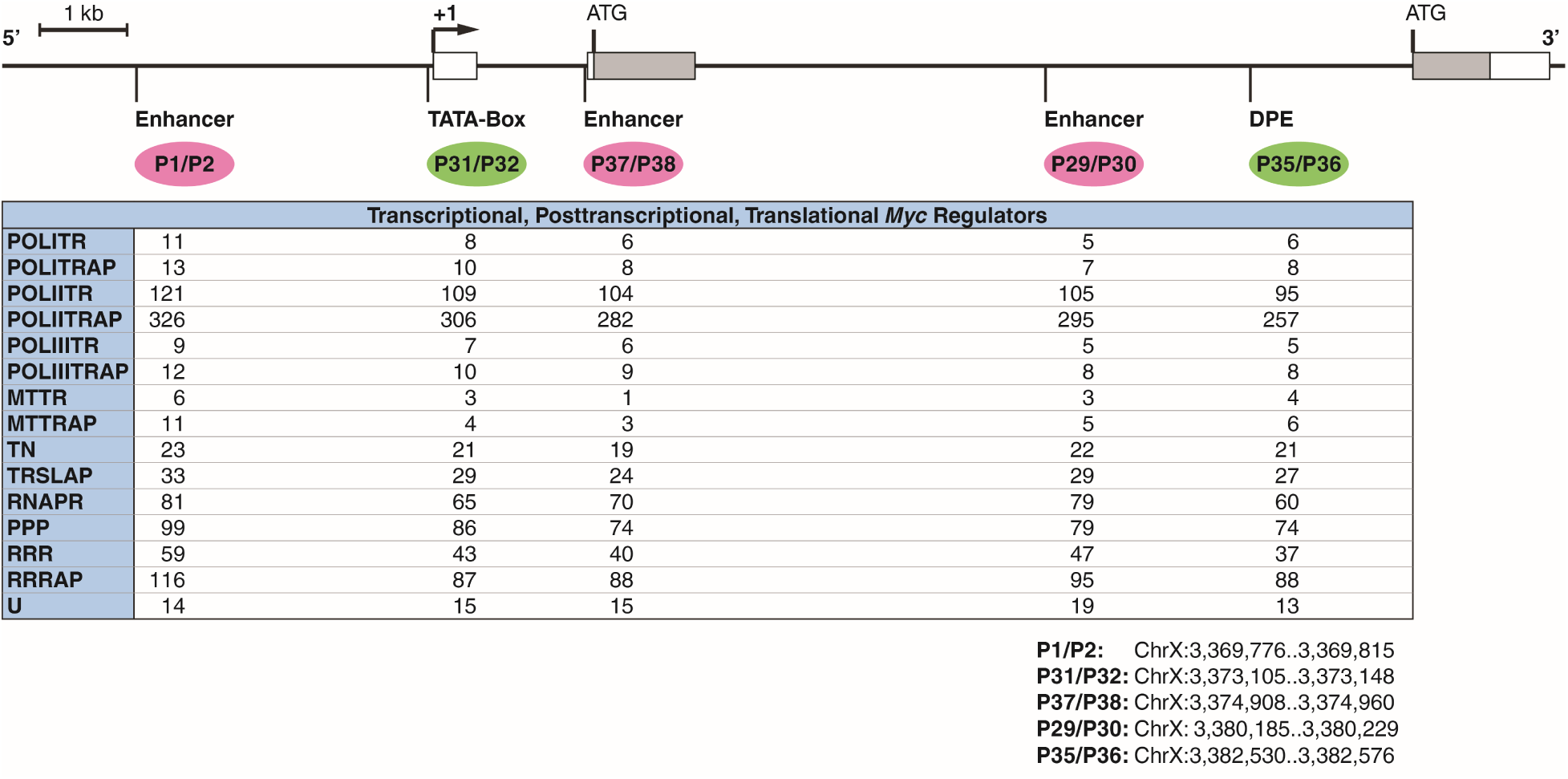
Subdivisions of factors identified associated with each Myc-CRM, categorized by molecular function and biological process. Investigated enhancers (upstream P1/P2, 5’-proximal P37/P38, intron 2 P29/P30) are indicated in magenta; promoter elements (TATA-box-containing P31/P32, DPE-containing P35/P36) are indicated in green. **Abbreviations**: **POLITR**, RNA Pol I Transcription; **POLITRAP**, RNA Pol I Transcription-Associated Processes; **POLIITR**, RNA Pol II Transcription; **POLIITRAP**, RNA Pol II Transcription-Associated Processes; **POLIIITR**, RNA Pol III Transcription; **POLIIITRAP**, RNA Pol III Transcription-Associated Processes; **MTTR**, Mitochondrial Transcription; **MTTRAP**, Mitochondrial Transcription-Associated Processes; **TN**, Translation; **TRSLAP**, Translation-Associated Processes; **RNAPR**, RNA Processing; **PPP**, Posttranscriptional/Posttranslational Processes; **RRR**, Replication-Repair-Recombination; **RRRAP**, Replication-Repair-Recombination-Associated Processes; **U**, Uncharacterized. **Note**: A complete list of potential *Myc* regulators, including factors involved in transcription, post-transcriptional/post-translational processes, hormone reception, and protein modification, is available in supplemental data (Table 3, Image 3 in File S3). See Table A-G in File S10 for analysis of transcriptional categories.

#### *Myc* cell cycle cofactors associated with the Myc-CRMs might act as potential regulators of *Myc* expression

Most of the factors described here represent *Myc* cell cycle cofactors (Figure 13; see also supporting information, Table A-F in File S11), which are tightly linked to cell growth and patterning (Edgar *et al*. 2001). It is well accepted that factors regulated by *Myc* feedback on Myc activity to tightly control *Myc* expression and influence wild-type body size during development. Our results further show that many conserved developmental signaling pathways converge on conserved noncoding sequences at the *Drosophila Myc* locus to regulate *Myc* expression in a manner dependent on developmental stage and cellular context. Many factors discussed in this Results Section demonstrate that signaling pathways including Wnt/Wg (Barker *et al*. 2000), Notch (Orian *et al*. 2007), TGF-β/Dpp (Barker *et al*. 2000), EGFR (Orian *et al*. 2007), and Hedgehog (Kenney *et al*. 2003; Hallikas *et al*. 2006) act as primary regulators of *Myc* (Figure 13), findings that have been confirmed by other researchers as well.

**Figure 13.**
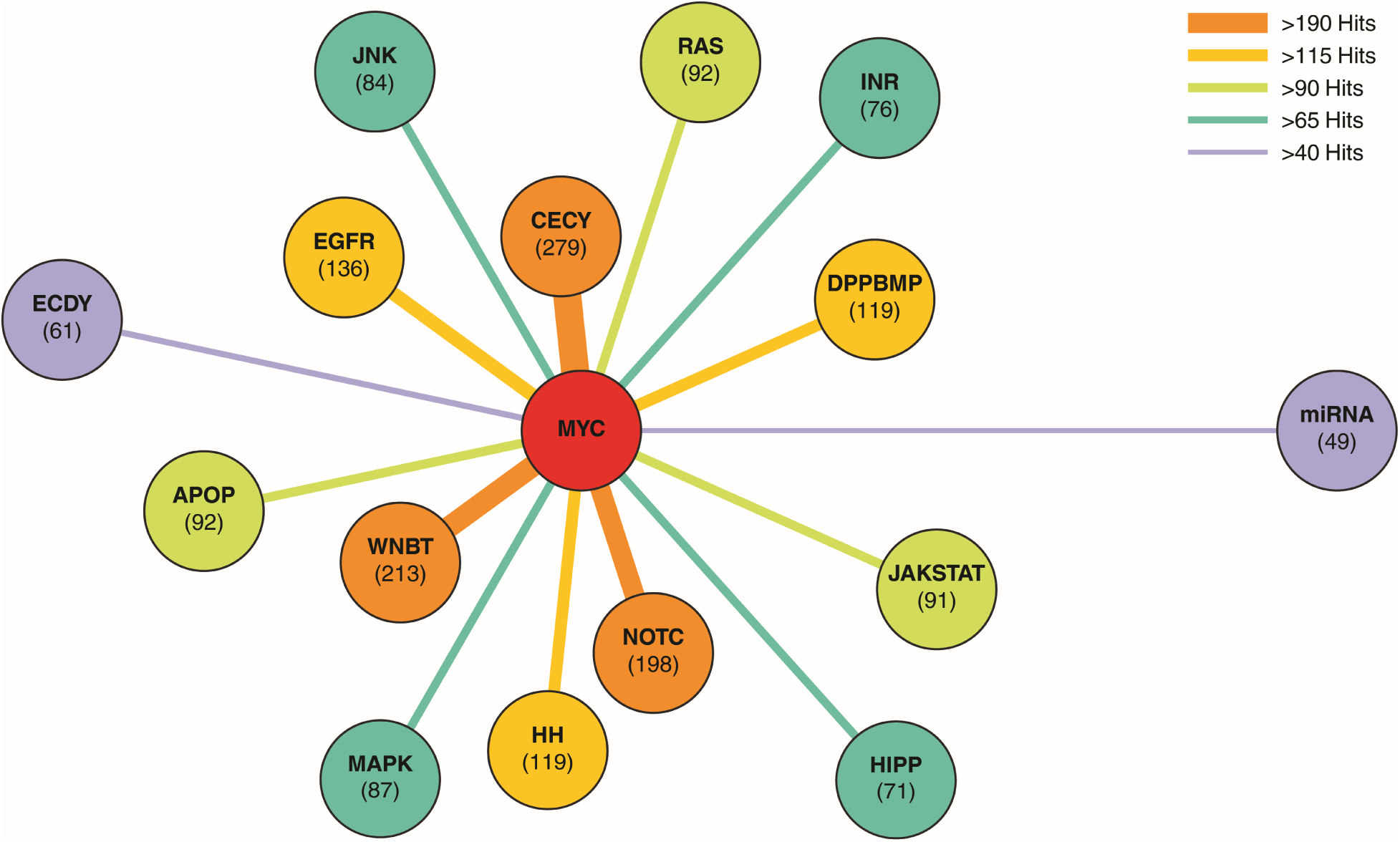
Graphical illustration of cell cycle factors and components of signaling pathways as potential regulators of *Myc* and *wingless/wg* expression. The factors identified as potential regulators of *Myc* predominantly belong to cell cycle regulators and signaling pathways, including Wingless (Wg), Notch (N), Hedgehog (HH), EGFR, BMP/DPP, MAPK, JAK/STAT, and RAS, followed by several other signaling pathways. All tested Myc-CRMs and the positive control (W-CRM) dynamically associate with the identified cell cycle factors and components of the listed pathways (see Table A-F in File S11). The numbers likely refer to protein counts per cell cycle category or pathway. **Abbreviations**: **CECY**, Cell Cycle; **WNBT**, canonical Wnt signaling (Wnt/β-Catenin/TCF); **NOTC**, Notch signaling pathway; **EGFR**, Epidermal Growth Factor Signaling Pathway; **DPPBMP**, DPP/BMP signaling pathway; **HH**, Hedgehog signaling pathway; **RAS**, RAS GTPase signaling pathway; **APOP**, Apoptosis pathway; **JAKSTAT**, JAK/STAT signaling pathway; **JNK**, Janus kinase signaling pathway; **INR**, Insulin-like receptor signaling pathway; **HIPP**, Hippo signaling pathway; **MAPK**, Mitogen-activated protein kinase signaling pathway; **ECDY**, Ecdysteroid signaling pathway; **miRNA**, miRNA signaling pathway.

#### Cyclin-dependent kinase 1 (Cdk1, also known as Cdc2) might control Myc levels during cell cycle progression, potentially contributing to cell size checkpoint control

Cdk1, regulated by the Cdc25/String phosphatase (Edgar *et al*. 2001), functions in association with Cyclin A, Cyclin B, and Cyclin B3. Phosphorylation of a wide range of cell cycle factors by Cdk1 regulates entry into and progression through mitosis at the G1/S (Lehner and O’Farrell 1990) and G2/M transitions (Lehner and O’Farrell 1990; Stern *et al*. 1993) in both growing and proliferating cells, as well as in differentiated larval cells lacking a G2/M transition (Edgar *et al*. 2001). The transcription factor Myc regulates String activity to control Cdk1 and balance cell division with differentiation in embryos and discs (Edgar *et al*. 2001). Previous research has shown that in metazoans, a Cdk1-linked cell size checkpoint may exist, influenced by the Myc/Max network through regulation of ribosomal protein biogenesis and proteasomal degradation of cell cycle proteins (Bonke *et al*. 2013).

#### The Wee1 tyrosine kinase, which regulates entry into mitosis, might also be involved in regulating Myc levels

Wee1, a non-specific protein kinase that belongs to cell cycle checkpoint factors (Kiessling *et al*. 2009), inhibits Cdc2 (Cdk1) to prevent premature entry into the mitotic phase of the cell cycle (Campbell *et al*. 1995). Transitions into the G1/S (Laranjeiro *et al*. 2013) and G2/M phases (Matsuo *et al*. 2003) are key determinants during normal cell growth and division, and aberrations in these checkpoints can lead to malignant transformation. The activities of both Wee1 kinase (Masri *et al*. 2013; Kelleher *et al*. 2014) and the growth regulator MYC (Bouchard *et al*. 1999; Perez-Roger *et al*. 1999) are under tight control by other proteins during cell cycle transitions to synchronize cell cycle phases and prevent transmission of damaged DNA (Feillet *et al*. 2014; Kelleher *et al*. 2014). Like MYC, the developmental protein Wee1 is essential for embryonic syncytial blastoderm development (Lee and Orr-Weaver 2003). We identified Wee1 associated with the upstream enhancer P1/P2 and the initiation sites—the TATA-box promoter (P31/P32) and the DPE (P35/P36). This result suggests that Wee1 kinase likely plays a crucial role in fine-tuning Myc levels during cell cycle phases in both proliferating and differentiating cells.

#### Cyclin-dependent kinase 5 (Cdk5) might be involved in controlling Myc expression during neuronal differentiation

Cdk5, a non-specific serine/threonine protein kinase, contributes to degenerative diseases such as retinal degeneration through phosphorylation of MEKK1, activation of the JNK cascade, and induction of apoptosis in response to endoplasmic stress signals (Kang *et al*. 2012). Cdk5 interacts with Rac GTPase to control axonogenesis in *Drosophila* embryos, and together with its interacting partner P35, is essential for the differentiation of post-mitotic neurons (Connell-Crowley *et al*. 2000). In mammals, Cdk5, which is frequently dysregulated in many cancer types, interacts physically with MYC. It promotes tumorigenesis through mechanisms including phosphorylation of the MYC protein and inhibition of MYC inhibitors like Bin1 (Ito *et al*. 2014; Zhang *et al*. 2019).

#### Tousled-like kinase (TLK) participates in cell cycle-dependent morphogenesis, a process potentially involving Myc control

TLK is a conserved non-specific serine/threonine protein kinase for which the conserved anti-silencing protein 1 (ASF1) serves as a substrate. TLK functions throughout the cell cycle, with activity levels peaking during S phase. TLK and ACF, as components of the chromatin assembly complex, phosphorylate each other during S phase in both mammals and *Drosophila* to coordinate replication and chromatin organization (Carrera *et al*. 2003), progression through cell cycle phases, and nuclear division (Carrera *et al*. 2003). Tousled-like kinase associates with components of Wingless signaling (such as Armadillo, DE-cadherin, and atypical protein kinase C (aPKC)) as well as Notch signaling to induce protein degradation and apoptosis in adherens junctions, facilitating cell migration and modulating morphogenesis (Yeh *et al*. 2015; Zhang *et al*. 2016).

#### P-element somatic inhibitor (Psi) plays a role in regulating Myc transcription

Psi, an RNA-binding splicing factor, is phosphorylated by TLK (Taliaferro *et al*. 2013). Psi has dual roles: RNA processing of transposons in a tissue-specific manner (Siebel *et al*. 1994) and transcriptional regulation. Psi activates *Myc* transcription through association with the basal transcription machinery and mediator complex at the *Myc* promoter (Guo *et al*. 2016). Psi and TLK were both associated with all tested *Myc* targets and likely function together in controlling *Myc* expression, predominantly during the S phase of the cell cycle.

#### SKP1-related A (SkpA) may target Myc for degradation during cell fate commitment and patterning

SkpA, a component of Skp, Cullin, and F-box (SCF)-containing ubiquitin ligase complexes (Murphy 2003), plays essential roles in centrosome duplication, chromosome condensation, endoreplication (Murphy 2003), and microtubule assembly during mitosis (Ducat *et al*. 2008). SkpA conveys information from signaling pathways to regulate cell fate specification and morphogenesis in developing tissues. The SCF/Slimb complex induces degradation of Kibra protein, a negative regulator of the Hippo pathway, thereby promoting Hippo activity independently of Yorkie-dependent transcription, which in turn controls tissue patterning and growth (Tokamov *et al*. 2021). During neuronal remodeling and nervous system development, the SCF/E3 ligase complex targets components of insulin receptor, Pi3K, and Myc/TOR pathways for degradation to establish axon guidance and dendrite pruning (Wong *et al*. 2013). In conjunction with different signaling pathways, the SCF/E3 ligase complex targets Myc and Cyclin E for degradation to downregulate the cell cycle and initiate tissue patterning (Shcherbata *et al*. 2004; Nicholson *et al*. 2011).

#### Cell division cycle 6 protein (CDC6), a potential inhibitor of Myc transcriptional activity, might control Myc expression during replication and repair processes

CDC6 is an essential component of the pre-replication complex, present at virtually all origins of replication in a cell (InterPro Project Members 2004). Additionally, CDC6 enhances coordination between DNA replication and repair in early S phase and during mitosis (Crevel *et al*. 2005; GO Reference Genome Project 2011). CDC6 may also influence apoptosis and transcription (Crevel *et al*. 2005). Both *in vitro* and *in vivo* studies have shown that CDC6 competes with the MYC binding partner Max for interaction with MYC, thereby inhibiting MYC/Max binding to target gene E-boxes and reducing their transcriptional activation (Takayama *et al*. 2000).

#### Cell division cycle protein 16 (CDC16) might target Myc for degradation and thus contribute to controlling Myc levels during cell fate specification

CDC16 is part of the anaphase promoting complex (APC/C) or Cyclosome and associates with Shattered (*shtd*), the largest subunit of the APC/E3 ubiquitin ligase, for its function (Martins *et al*. 2017). The APC/C mediates the degradation of Cyclins, String, and other cell cycle factors during the G1 phase to induce cell fate specification in differentiating cells through modulation of Wingless signaling and control of post-mitotic processes (Martins *et al*. 2017). For example, G1 arrest is established in the morphogenetic furrow of the developing eye disc via degradation of Wingless pathway components to promote photoreceptor cell differentiation (Martins *et al*. 2017).

#### Rad50 and Mre11 may function as potential regulators of Myc, particularly during the S phase of the cell cycle

Rad50 and Mre11 are two conserved proteins belonging to the MRE11-RAD50-NBS1 (MRN) complex (Ciapponi *et al*. 2006; Gao *et al*. 2009; Bosso *et al*. 2019; On *et al*. 2021), with essential roles in DNA double-strand break repair (Gorski *et al*. 2004), telomere maintenance, and chromosome organization (Ciapponi *et al*. 2004). Disruption of Rad50 function leads to retarded third instar larval development and lethality during pupariation due to defects in double-strand break repair and subsequent cell death induction (Gorski *et al*. 2004). Rad50 contributes to G-quadruplex DNA binding for the unwinding and stabilization of G4 structures (GO Reference Genome Project 2011). As a predominant DNA damage complex component, Rad50 senses double-strand breaks and replication stress during DNA synthesis and, together with other factors, induces apoptosis to eliminate damaged cells (Gorski *et al*. 2004). In eukaryotes, conserved and tightly controlled mechanisms regulate cell cycle factors such as Myc, conferring integrity and precise shape to each organ during development (Liu *et al*. 2015). Whether Rad50 directly regulates *Myc* expression to ensure normal cell cycle progression remains to be investigated.

#### Replication factors, including DNA polymerase delta subunits and origin recognition/priming components, might be involved in regulating Myc expression

DNA polymerase delta subunit 1 (PolD1), the catalytic subunit of Pol delta, plays a crucial role in high-fidelity genome replication, including lagging strand synthesis, DNA recombination, and repair (Peck *et al*. 1992; Chiang *et al*. 1993; Aoyagi *et al*. 1994). PolD1 functions in highly proliferative embryonic cells to drive replication during the earliest cell cycles (Peck *et al*. 1992; Ji *et al*. 2019). PolD1 might be involved in controlling *Myc* expression during early development.

DNA polymerase delta subunit 2 (PolD2, also known as Pol31) functions as the accessory component of both the pol delta and pol zeta complexes and supports the function of PolD3 (Pol32). PolD2 participates in both embryonic development and postembryonic processes in somatic cells (Ji *et al*. 2019).

DNA polymerase delta subunit 3 (PolD3, also known as Pol32) is a component of multi-subunit DNA polymerases involved in DNA replication and double-strand break repair (Tritto *et al*. 2015), and serves as a processivity factor during genome replication (Ji *et al*. 2019). Pol32 might associate with Myc during DNA replication and repair. Origin recognition complex subunit 6 (Orc6), the smallest component of the origin recognition complex (ORC) (Akhmetova *et al*. 2015), plays a crucial role in initiating replication in eukaryotic cells. Through interaction with Septin, Orc6 is essential for both replication and cytokinesis in *Drosophila* (Huijbregts *et al*. 2009).

DNA primase subunit 1 (Prim1), a subunit of the DNA polymerase alpha complex, together with the Prim2 subunit, is responsible for synthesizing short RNA primers required by DNA polymerase alpha to prime DNA synthesis at replication origins (Villani *et al*. 1980; Kaguni *et al*. 1983; Cotterill *et al*. 1987; Kuroda *et al*. 1990; Bakkenist and Cotterill 1994).

#### Chromosome fidelity factors, such as Cutlet, may also regulate Myc during DNA synthesis

Chromosome cohesion factor Cutlet (also known as CTF18-RFC, a component of one of the PCNA loader complexes, CTF18-replication factor C) is important for the fidelity of chromosome transmission. Cutlet participates in DNA clamp loading and positively regulates DNA polymerase activity during the S phase of the cell cycle (Gene Ontology Curators 2002). Cutlet might play a crucial role in regulating *Myc* expression during the cell cycle.

### Components of different signaling pathways as potential regulators of the *Myc* expression

Many of these factors belong to known signaling pathways that play essential roles in *Myc* expression (Figure 13). We analyzed these factors to characterize the complex network of pathways converging on *Myc* promoter and enhancer elements.

#### The Hedgehog signaling inhibitor Suppressor of Fused (Su(fu)) might regulate Myc expression

*Su(fu)* is a tumor suppressor protein kinase (Han *et al*. 2015) and a conserved negative regulator of the Hedgehog signaling pathway that inhibits transcription factors Ci/Gli (Methot and Basler 2000) by preventing recruitment of the coactivator p300/CBP to the transcription regulator complex (Wang *et al*. 2000; Shi *et al*. 2014; Han *et al*. 2015; Oh *et al*. 2015; Han *et al*. 2019). MYC has been shown to regulate the Hh component Gli (Yoon *et al*. 2013; Aberger and Ruiz i Altaba 2014) in certain tumor types. Mechanistically, a regulatory circuit might exist between *Drosophila Myc* and the Hedgehog signaling regulator Su(fu).

The protein 14-3-3ε, an interactor with Ras/MAPK and Hippo signaling, might also regulate *Myc* expression. (*14-3-3ε*) is involved in cell proliferation and differentiation, tissue patterning, and organ size determination through interaction with multiple signaling pathways, predominantly the Ras/MAPK (Ashton-Beaucage *et al*. 2014) and Hippo signaling pathways (Ren *et al*. 2010; Pojer *et al*. 2021). The transcriptional cofactor Yorkie (Yki/YAP) of the Hippo signaling pathway is a well-established coactivator of *Myc* transcription during cell growth and cell cycle progression through its translocation to the nucleus (Neto-Silva *et al*. 2010). Phosphorylation of Yorkie promotes binding of 14-3-3 epsilon to Yorkie, thereby inhibiting Yorkie nuclear translocation (Ren *et al*. 2010), suggesting an indirect role for 14-3-3 epsilon in *Myc* downregulation during cell fate initiation and organ size determination.

#### The Drosophila Sumo protein (Sumo, smt3) is implicated in Myc protein degradation via Sumoylation

Small ubiquitin-like modifier (*Sumo*, *smt3*), the sole member of the Sumo family proteins in *Drosophila*, responds to signals from TGF-β (Miles *et al*. 2008), Ras/MAPK (Nie *et al*. 2009), Smoothened (Zhang *et al*. 2017), and Toll-NF-κB (Kanoh *et al*. 2015) signaling pathways, and plays a crucial role during the syncytial blastoderm mitotic cell cycle in embryonic patterning (Nie *et al*. 2009). Sumo contributes to wing morphogenesis through modulation of Hedgehog signaling during larval development (Zhang *et al*. 2017). Sumo reportedly regulates Myc protein levels during development through protein Sumoylation and subsequent degradation (Gonzalez-Prieto *et al*. 2015).

#### Adaptor protein AP-2, potentially along with Host cell factor (HCF) and Myeloid leukemia factor (MLF), may be involved in cell growth and differentiation through Myc level modulation

The AP-2 subunit of the Adaptor protein complex participates in asymmetric cell division during nervous system development via modulation of Notch signaling (Bardin *et al*. 2004). Furthermore, AP-2α is required for intracellular vesicle trafficking (Gonzalez-Gaitan and Jackle 1997; Narayanan and Ramaswami 2001; Windler and Bilder 2010), signaling receptor internalization, gene expression regulation (Moore *et al*. 2020), and cell polarity maintenance (Lin *et al*. 2015). Notch signaling plays a pivotal role in regulating cell proliferation and differentiation by modulating the transcription of Notch targets such as *Myc* (Mazur *et al*. 2010). Numb protein together with AP-2, key determinants of cardiac progenitor differentiation, downregulate *Myc* in a Pim1 background during mitosis to initiate cardiomyocyte fate (Cottage *et al*. 2010). Mammalian AP-2 is involved in the repression of *c-MYC* target genes (Gaubatz *et al*. 1995) and inhibition of MYC-induced apoptosis (Moser *et al*. 1997).

Coactivator Host cell factor (HCF/*Hcf*) (Mahajan *et al*. 2003) is a component of the Ada2a complex (ATAC); Ash2-Sin3A-HCF interactions induce chromatin remodeling and histone modifications on promoters of genes involved in cell growth and proliferation (Guelman *et al*. 2006a; Suganuma and Workman 2008; Furrer *et al*. 2010; Suganuma *et al*. 2012). The DPE promoter requires Ada2 and DR1/NC2B to establish transcription initiation from this site (Hsu *et al*. 2008). DR1-Drap1/NC2B is a bifunctional basal transcription factor that can repress TATA-dependent transcription while activating DPE-dependent transcription to establish differential transcriptional bursting (Willy *et al*. 2000). Ash2 and HCF were associated with all examined Myc-CRMs, indicating that chromatin remodeling complexes containing Ash2-Sin3A-HCF components might differentially control *Myc* transcription from multiple sites.

DNA binding Myelodysplasia/myeloid leukemia factor (MLF) was associated with all tested Myc-CRMs. MLF is required for cell growth, proliferation, and differentiation in the developing eye via interaction with the transcription factor DREF (Jasper *et al*. 2002), and MLF/DnaJ-1 participates in lineage commitment of primary hematopoietic progenitors during blood cell development. Transcription factor MYC plays a crucial role during hematopoiesis (Lu *et al*. 2023), and in acute myeloid leukemia (AML), both MYC and MLF are overexpressed (Bras *et al*. 2012). We identified both MLF and DnaJ-1 proteins in our *Myc* pulldown samples, suggesting a putative role for MLF in regulating *Myc* expression during cell proliferation, differentiation, and patterning.

#### Mothers against dpp (Mad), which can downregulate Myc during the G1/S transition to induce cell differentiation, was identified associated with Myc-CRMs

Mad is a pivotal pol II-specific Mad homology domain transcription factor with diverse functions during development. As a core component of the Dpp/BMP signaling pathway, Mad responds to Dpp/BMP ligands decapentaplegic (*dpp*), Screw (*scw*), and glass bottom boat (*gbb*) to regulate the expression of Dpp-responsive target genes (Das *et al*. 1998b; Inoue *et al*. 1998; Dai *et al*. 2000; Muller *et al*. 2003; Yao *et al*. 2006; Weiss *et al*. 2010; Vuilleumier *et al*. 2022). Mad induces proliferation arrest during the G1/S transition (Firth and Baker 2005) and modulates differentiation and morphogenesis during limb development (Marquez *et al*. 2001), wing patterning (Lecuit *et al*. 1996; Dworkin and Gibson 2006), eye development (Wiersdorff *et al*. 1996; Liu *et al*. 2023b), and follicle cell morphogenesis (Kirilly *et al*. 2005). The Myc/Max/Mad network of transcription factors functions as transcriptional activators or repressors in a tissue-specific manner during cell proliferation and differentiation (Grandori *et al*. 2000). Myc and Mad are two opposing transcription factors that control the balance between cell fate determination and tissue size and shape (Okada and Shi 2018).

#### Pontin (pont) and Reptin (rept) may control Myc expression during the establishment of basolateral cell polarity

Pontin and Reptin belong to the family of RuvB-like DNA helicases (GO Reference Genome Project 2011) and play key roles in multiple signaling pathways. They are components of chromatin remodeling complexes such as Tip60, SWRI (Kusch *et al*. 2004), and Ino80 (Klymenko *et al*. 2006) that enhance DNA accessibility to transcription factors, replication machinery, and DNA damage/repair factors by modifying chromatin structure and nucleosome positioning (Kusch *et al*. 2004). Pontin (Tip49/Pontin) and Reptin (Tip48/Reptin)—with Pontin showing emphasized genetic interaction—are essential for normal Myc function during growth and proliferation (Bellosta *et al*. 2005). The evolutionarily conserved Pontin operates at the nexus of complex signaling cascades including Jun kinase, TNFα/Eiger (Wang *et al*. 2018a; Wang *et al*. 2018b), Wingless/Wg (Bauer *et al*. 2000), and the Myc pathway (Bellosta *et al*. 2005) to control the activation or repression of developmental genes during patterning. We identified guanylate kinase Discs large 1 (*dlg1*), the scaffolding protein Scribble (*scrib*), and the tumor suppressor protein Lethal giant larvae (LGL/*lgl*, *l(2)gl*, *CG2671*) associated with the examined Myc-CRMs (see Table 5 in File S5 and Table 6 in File S6). These proteins are components of the Scribble protein complex and play crucial roles in epithelial cell basolateral integrity and function (Khoury and Bilder 2020). The Scribble protein complex modulates signal transduction processes and controls cell proliferation (Khoury and Bilder 2020). Scribble function is essential for embryonic epithelium patterning (Gibson and Perrimon 2003), control of cell polarity and proliferation (Zeitler *et al*. 2004), stem cell maintenance and division (Zeng *et al*. 2010), and cell competition. Cell polarity proteins such as aPKC, Scribble, and LGL (Wakefield and Stuelten 2007; Kim *et al*. 2014), together with the growth regulator Myc, considered a key indicator of cell fitness, interact to balance epithelial cell morphogenesis and polarity for normal development (Fahey-Lozano *et al*. 2019).

#### Ecdysteroid receptor-related protein SMRTER regulates Myc expression during cell fate specification and metamorphosis

SMRTER (SMRT-related and ecdysteroid receptor interacting factor) (*Smr*), a chromatin binding factor (TSAI *et al*. 2004; TSUDA *et al*. 2006; HECK *et al*. 2012) is part of a conserved transcription repressor complex that together with the Sin3A/HDAC complex negatively regulates ecdysone receptor signaling during fly development (KOZLOVA AND THUMMEL 2000). SMRTER also interacts with components of other key developmental pathways; during follicle cell morphogenesis and wing disc patterning, it binds the Notch antagonist Suppressor of Hairless (Su(H)) to inhibit Notch target transcription (Heck *et al*. 2012). Additionally, as part of the Ebi/SMRTER/Su(H) corepressor complex, it modulates both EGFR and Notch signaling pathways to guide photoreceptor cell activity in the developing *Drosophila* eye (Tsuda *et al*. 2006). Given that Ecdysone signaling is itself an important regulator of *Myc*—influencing cell growth, proliferation, and cell fate specification to control developmental timing and tissue patterning (Delanoue *et al*. 2010; Rajan and Perrimon 2010; Gallant 2013).

#### The demethylase enzyme Little imaginal discs/Lysine demethylase 5 (Lid/Kdm5) potentially regulates Myc-dependent cell growth through direct binding to Myc

Little imaginal discs/Lysine demethylase 5 (*Kdm5*, *lid*, *CG9088*), a Trithorax Group protein, possesses histone H3 demethylase activity (Secombe *et al*. 2007; Lloret-Llinares *et al*. 2008; Tarayrah *et al*. 2015). Lid/Kdm5 belongs to the AT-rich interaction domain (ARID) family of transcriptional regulators (Liu and Secombe 2015) and binds DNA non-specifically to remodel chromatin (GO Reference Genome Project 2011) and control general transcription (Secombe *et al*. 2007; Liu and Secombe 2015). Lid/Kdm5 functions as a transcriptional modulator of JAK/STAT signaling during male germline stem cell differentiation and maintenance (Tarayrah *et al*. 2015). Lid regulates Myc-dependent cell growth through direct binding to Myc (Secombe *et al*. 2007), and this binding interaction between Lid homologs JARID1A, JARID1B, and MYC has also been observed in mammalian systems, indicating evolutionary conservation of the function of these proteins (Secombe *et al*. 2007).

#### Lysine ketoglutarate reductase (LKRSDH) might regulate Myc transcription, potentially during apoptosis

LKRSDH is a bifunctional enzyme that functions in the Lysine degradation pathway (Cakouros *et al*. 2008; St Clair *et al*. 2017) and, acting as a transcriptional corepressor, regulates the expression of apoptotic genes through histone methylation in response to ecdysone signaling (Cakouros *et al*. 2008). Similar to Lid (Kdm5, lid), LKRSDH plays roles in Myc-induced growth control (Secombe *et al*. 2007), suggesting it might have a feedforward function in regulating *Myc* transcription.

The nucleolar-localized and RNA recognition motif (RRM)-containing Modulo (*mod*) protein (Perrin *et al*. 1999) is a downstream target of *Myc* required during cell growth and proliferation in imaginal tissues but not in endoreplicating larval tissues (Perrin *et al*. 2003). Modulo, tightly controlled by Myc, plays an important role in biomass production and is required for nucleolus function (Perrin *et al*. 1998). Mutations in Modulo result in the Minute phenotype, characterized by flies with small body size and female sterility (Perrin *et al*. 1999). Here we identified Modulo associated with all investigated Myc-CRMs, suggesting a possible role for Modulo in regulating *Myc*.

#### Wingless cofactor ‘Short wing’ is a potential regulator of Myc and the chromatin remodeler ‘will die slowly’ controls Myc expression during growth and morphogenesis

Short wing, Sw (*sw*), a dynactin binding (MORGAN *et al*. 2011) protein, and a subunit of the cytoplasmic dynein motor complex (NURMINSKY *et al*. 1998) contributes to wing and eye development (MYSTER AND PEIFER 2001), mitotic cell cycle (DZHINDZHEV *et al*. 2005; MORALES-MULIA AND SCHOLEY 2005), fate determination in 16-cell syncytium stage withing the egg chamber (MCGRAIL AND HAYS 1997), and neurogenesis (ZHENG *et al*. 2008). Since Sw was associated with all investigated *Myc* targets, and both Myc and Sw proteins are essential during early embryonic development, cell fate specification, and oocyte maturation, there might be yet unknown direct or indirect regulatory circuits whereby Sw controls *Myc* expression.

Protein will die slowly (*wds*), a WD40 domain-containing chromatin remodeler that, in association with the Ada2a complex (ATAC), enables histone H3 acetylation and possibly H4 to regulate transcription (Suganuma and Workman 2008). As part of the Trx/MLL and NSL complexes (Pascual-Garcia *et al*. 2014), Wds plays a crucial role in the sustained transcriptional activation of neurogenic genes through interaction with the Myc-interacting protein Mad and Trx (Yao *et al*. 2018a). WD40 domain proteins have been implicated in *MYC* downregulation through ubiquitination and degradation of the MYC protein during tissue patterning (Gu *et al*. 2007).

#### Components of Ras/ERK, MAPK, and insulin signaling pathways are known regulators of Myc

Mitogen-activated protein kinase (*ERK*, MAPK, *rl*) is a core component of the RAS/MAPK pathway and binds to a wide range of proteins to phosphorylate downstream cytoplasmic and nuclear signal transducers and transcription factors. ERK plays a crucial role in varied cell fate decisions, as well as cell, tissue, and body size control (Brunner *et al*. 1994; O’Neill *et al*. 1994; Schweitzer *et al*. 1995; Oellers and Hafen 1996; Karim and Rubin 1999; Baker *et al*. 2001; Qiao *et al*. 2006) by interacting with components of multiple signaling pathways, including the epidermal growth factor receptor (EGFR) signaling pathway (Diaz-Benjumea and Hafen 1994; Schweitzer *et al*. 1995; Oellers and Hafen 1996; Schnepp *et al*. 1998; Reich and Shilo 2002; Molnar and de Celis 2013; Yang *et al*. 2016; Ogura *et al*. 2018), the ERK1 and ERK2 cascade (Oellers and Hafen 1996; Clemens *et al*. 2000; Roy *et al*. 2002; Douziech *et al*. 2006; Ashton-Beaucage *et al*. 2014; Goyal *et al*. 2017), fibroblast growth factor receptor (FGFR) signaling (Clemens *et al*. 2000; Ohshiro *et al*. 2002; Sato and Kornberg 2002; Gryzik and Muller 2004; Petit *et al*. 2004; Csiszar *et al*. 2010; Ukken *et al*. 2014; Du *et al*. 2018), insulin receptor signaling (Clemens *et al*. 2000; Xu *et al*. 2013; Slack *et al*. 2015), Wingless signaling(Sinenko *et al*. 2009; Dragojlovic-Munther and Martinez-Agosto 2013), Sevenless signaling (Biggs *et al*. 1994; Brunner *et al*. 1994; Karim *et al*. 1996; Kumar *et al*. 1998; Kitadate *et al*. 2007), and Torso (Tor) signaling pathways (Brunner *et al*. 1994; Ghiglione *et al*. 1999; Lim *et al*. 1999; Goyal *et al*. 2017). The Ras/ERK cascade modulates the spatiotemporal regulation of *Myc* in many different ways during development and malignant transformation (Sears *et al*. 1999; Prober and Edgar 2002; Grewal 2015). Ras/ERK and Myc pathways are growth-promoting and pro-survival signaling pathways. *Myc* expression was shown to be elevated for compensatory proliferation in damaged *Drosophila* imaginal tissues expressing the apoptosis activator head involution defective (hid), not solely in response to the apoptotic pathway but rather in conjunction with increased activity of the Ras/ERK cascade (Crucianelli *et al*. 2022).

MAPK-regulated corepressor-interacting protein (MCRIP) (*CG34159*) localizes to both cytoplasmic stress granules and the nucleus and exists in two variants, MCRIP1 and MCRIP2 (Ichikawa *et al*. 2015). When ERK concentration is low, the MCRIP1 isoform remains complexed with CtBP in the cytoplasm, permitting E-cadherin transcription and maintaining epithelial cell integrity. Elevated ERK levels, however, lead to MCRIP1 phosphorylation by ERK, causing its dissociation from CtBP and subsequent nuclear translocation where it interacts with the DNA-bound corepressor ZEB1 at the E-cadherin promoter. This MCRIP1/ZEB1 heterodimer formation results in epigenetic silencing and transcriptional inhibition of E-cadherin (Ichikawa *et al*. 2015), promoting epithelial to mesenchymal transition (EMT) during both development and disease states (Ichikawa *et al*. 2015). Notably, in adult malignancies, both the Zinc finger E-box binding protein ZEB1 and MYC are overexpressed in numerous cancer types (Katoh *et al*. 2013; Li *et al*. 2021; Fratini *et al*. 2023). The presence of an E-box motif (5’-CAGCTG-3’) within the P1/P2 enhancer cluster (Table S1.3) suggests that MCRIP may participate in either direct or indirect regulation of *Myc* transcription, potentially contributing to *Myc* dysregulation during tumorigenesis.

#### Ajuba (jub) may control Myc activity during the mitotic cell cycle and differentiation, potentially via interaction with the Hippo signaling pathway

The nuclear protein Ajuba (*jub*) functions as a transcriptional corepressor (GO Reference Genome Project 2011) and is involved in follicle cell development (Jia *et al*. 2015) and mitotic cell cycle control (Sabino *et al*. 2011). Through its protein sequestering ability (Rauskolb *et al*. 2014), Ajuba inhibits Hippo pathway activation by sequestering Yorkie (*Yki*), thereby inhibiting Hippo pathway activation during epithelial morphogenesis and tissue size determination (Das Thakur *et al*. 2010; Rauskolb *et al*. 2014). Hippo signaling is an important regulator of *Myc* during cell proliferation, differentiation, and patterning (Neto-Silva *et al*. 2010). Myc function is essential during embryonic epithelial morphogenesis and cell differentiation (Nair *et al*. 2006), dosage compensation (Mathews *et al*. 2017), and oocyte maturation (Maines *et al*. 2004). Since the corepressor Ajuba associates with the pol II transcription regulator complex (GO Reference Genome Project 2011), Ajuba might be involved in regulating *Myc* transcription.

Misshapen (*Msn*), a non-specific Ser/Thr protein kinase, is a negative regulator of Notch signaling (Mishra *et al*. 2015) and a positive regulator of Hippo signaling (Li *et al*. 2014; Li *et al*. 2015; Meng *et al*. 2015). Msn plays a crucial role during cell migration, epithelial morphogenesis, dorsal closure (Su *et al*. 1998; Kadrmas *et al*. 2004; Koppen *et al*. 2006), imaginal disc fusion, and thorax closure (Kadrmas *et al*. 2004). As a regulator of Hippo signaling, Misshapen is involved in Yorkie nuclear localization and Hippo target gene activation (Li *et al*. 2015), which suggests the Misshapen-Warts-Yorkie circuit as a potential regulator of *Myc* transcription.

#### The transcription factor Putzig (pzg) may regulate Myc transcription during DNA synthesis, cell division, and patterning

Nuclear C2H2 Zinc finger transcription factor Putzig (*pzg*) is involved in proliferation and growth via expression of mitotic cell cycle components and transcriptional stimulation of replication-required genes in a Trf2/DREF-dependent manner (Kugler and Nagel 2007; Kugler *et al*. 2011). Independent of DREF, Putzig can positively regulate Notch signaling in certain proliferating cell types by binding to target gene promoters and activating transcription (Kugler *et al*. 2011). In association with the nucleosome remodeling factor complex (NuRF), Putzig positively regulates ecdysone receptor (EcR) signaling targets during growth and metamorphosis while negatively regulating JAK/STAT signaling pathway targets to modulate the immune response (Kugler *et al*. 2011). Trf2/DREF is an important transcriptional regulator involved in controlling the transcription of cell cycle components such as Myc and E2F, thus playing an important role in cell division and tissue size control (Hyun *et al*. 2005). Putzig and Trf2 were both associated with the Myc-CRMs, suggesting they function in the transcriptional regulation of *Myc*.

#### MYC cofactor Moesin plays a role in the regulation of Drosophila Myc during oogenesis and body plan specification

Moesin (*Moe*), the sole *Drosophila* Erzin, Radixin, Moesin protein is involved in a variety of processes including cortical cytoskeleton stability (CARRENO *et al*. 2008; JAYANANDANAN *et al*. 2014), epithelial integrity (YANG *et al*. 2012), cell shape including ovarian follicular morphogenesis (POLESELLO *et al*. 2002; SOLINET *et al*. 2013), and determination of left/right body symmetry (TANIGUCHI *et al*. 2007). Moesin belongs to c-MYC networks of cell cycle and apoptosis proteins and is a MYC cofactor (Ciribilli *et al*. 2015). We conclude that the Myc cofactor Moesin might also be involved in regulating *Myc* expression during development.

#### Scribbler (sbb) might be a potential Myc regulator, potentially acting via interactions with multiple signaling pathways

Scribbler (*sbb*) (also known as Brakeless (*bks*); master of Thickveins (*mtv*)) is a transcriptional coregulator (Haecker *et al*. 2007) that interacts with the mediator complex (Crona *et al*. 2015) and the nuclear corepressor Atrophin-Grunge (*Gug*, *Atro*) to repress transcription in a Tailless (*tll*)-mediated manner during early embryonic segmentation (Haecker *et al*. 2007). In photoreceptor cells, Scribbler represses the function of Runt (*Run*) (Kaminker *et al*. 2002) to determine axon guidance and projection to the medulla (Haecker *et al*. 2007). Scribbler cooperates with Notch and Wingless signaling during wing disc morphogenesis and D/V boundary formation (Bejarano *et al*. 2008), and functions downstream of Hedgehog signaling to modulate the expression of TGF-β and Hedgehog target genes, determining many aspects of wing disc patterning (Dworkin and Gibson 2006). Whether Scribbler regulates *Myc* during development remains to be investigated.

#### Amun, a component of the Notch signaling pathway, might regulate Myc expression during processes such as sensory organ development

Amun, isoform A (*Amun*), detectable in association with protein-containing complexes, is a nuclear transcriptional regulator containing a glycosylase domain and is a component of the conserved Notch signaling pathway. It plays a crucial role during cell fate commitment, sensory organ development, chaeta development, compound eye morphogenesis, and wing disc patterning (Shalaby *et al*. 2009). Components of Notch signaling and Myc regulate the balance between proliferation and cell fate specification to propagate morphogenesis and tissue patterning (Orian *et al*. 2007; Kopecky *et al*. 2011). Glycosylation is essential for Notch components to transduce the signal and express target genes (Haines and Irvine 2003).

#### The adhesion-related molecule Ran-binding protein M (RanBPM) is a potential regulator of Myc during larval development

*RanBPM* functions as a bridge between extracellular adhesion molecules and membrane receptors (Dansereau and Lasko 2008) and contributes to germline stem cell maintenance, follicular cell adhesion within the egg chamber, and the female stem cell niche (Dansereau and Lasko 2008). RanBPM is part of the E3 ubiquitin ligase complex known as Glucose-Induced Degradation Deficient (GID), which is involved in the metabolic transition between sugar degradation and gluconeogenesis (Zavortink *et al*. 2020). *Drosophila Myc* is essential during larval growth, influencing processes including endoreplication, body size, and behavior, while the function of RanBPM in the larval mushroom body is critical for proper behavioral development (Demontis and Perrimon 2009; Scantlebury *et al*. 2010), suggesting a putative functional and regulatory link between these two genes.

#### The nucleoporin Nup98-96 may regulate Myc during the mitotic cell cycle

*Nup98-96*, a member of the nucleoporin protein family and precursor to Nup96 and Nup98 (Panda *et al*. 2014), associates with the nuclear membrane and chromatin (Kristo *et al*. 2017). It directs mRNA export from the nucleus (Kristo *et al*. 2017), facilitates the translocation of SMAD, Mad, and other proteins into and out of the nucleus (Chen and Xu 2010; Kristo *et al*. 2017; Wu *et al*. 2019), and is involved in transcriptional regulation (Capelson *et al*. 2010; Pascual-Garcia *et al*. 2014), predominantly upregulating developmentally active genes and genes regulating the cell cycle, such as *Myc* (Kalverda *et al*. 2010). Nucleoporins (Nups) enhance the function of actively transcribing genes via enhancer-promoter loop anchoring and also provide a scaffold for poised genes, enabling their immediate induction (Pascual-Garcia *et al*. 2014). These functions are important in processes such as the maintenance of germline stem cells and preservation of the germline stem cell niche, often in collaboration with components of pathways like TGF-β and EGFR signaling (Parrott *et al*. 2011). In oncogenic diseases, Nup98 collaborates with PHD finger and bromodomain transcription factors to upregulate the downstream target Pim1, leading to *MYC* overexpression (Noura *et al*. 2024).

#### The Wnt component transcription factor Net and the Notch antagonist Hairless are potential regulators of Myc transcription during development

The E-box binding (GO Reference Genome Project 2011) Myc-type basic helix-loop-helix (bHLH) transcription repressor Net (*Net, CG11450*) is an essential developmental protein involved in cell proliferation, fate induction, and differentiation (Peyrefitte *et al*. 2001). Net is a component of the Wnt enhanceosome and is important for wing vein specification and morphogenesis by antagonizing EGFR activities in intervein territories (Brentrup *et al*. 2000; De Celis 2003). The Wingless signaling pathway is known as a major regulator of *Myc* during cell proliferation, differentiation, and organ size determination (Wu and Johnston 2010). Both Net and Myc proteins belong to the basic helix-loop-helix (bHLH) family of transcription factors. A link may exist between Net and the downregulation of *Myc* during cell fate specification and differentiation through the involvement of signaling cascades like Notch, Wingless, and EGFR.

Corepressor Hairless (*H*) (Brou *et al*. 1994; Maier *et al*. 2011), a conserved silencer of Notch signaling, forms transcriptional repressor complexes consisting of (H/Su(H)) and/or (H/Su(H)/Gro/CtBP) proteins (Lyman and Yedvobnick 1995; Nagel *et al*. 2005; Maier *et al*. 2011; Smylla *et al*. 2019; Wolf *et al*. 2019) to modulate Notch signaling during sensory organ development (Nagel *et al*. 2000; Castro *et al*. 2005). The H/Su(H) complex controls the rate of cell proliferation and differentiation via Notch signaling during wing vein and wing margin morphogenesis (Nagel *et al*. 2005) and plays an essential role in wing disc Dorsal/Ventral compartment boundary formation by modulating the expression of Wingless (*wg*) and Vestigial (*vg*) (Koelzer and Klein 2006). Previous studies have shown that Hairless is involved in cell proliferation, cell death, differentiation initiation, and the regulation of *Myc* transcription (Muller *et al*. 2005; Orian *et al*. 2007; Herranz *et al*. 2008).

#### Components of immune pathways, such as Turandot Z (TotZ), were identified among the proteins associated with Myc-CRMs

The secreted peptide Totz functions in stress situations and in conjunction with the Toll, JAK/STAT, Imd, and Mekk1/JNK pathways (Brun *et al*. 2006) in response to UV damage, heat induction, paraquat-induced oxidative stress, and bacterial infection (Ekengren and Hultmark 2001). In our proteomics approach for identifying *Myc* regulators and related pathways, we identified a wide range of components from conserved signaling pathways, including Toll, JAK/STAT, Imd, and Mekk1/JNK (Figure 13; see also supporting information, Table A-F in File S11); these pathways have previously been documented by others as important regulators of *Myc* expression (Cordero *et al*. 2012; Zoranovic *et al*. 2013; Alpar *et al*. 2018; Fahey-Lozano *et al*. 2019; Schaub *et al*. 2019; Wang *et al*. 2019; Crucianelli *et al*. 2022). Turandot Z was highly enriched in the DPE-containing *Myc*-CRM P35/P36 sample (see Table 3 in File S3), indicating a potential role for TotZ in *Myc* regulation.

### XII. Proteins identified with the enhancer cluster P1/P2 are potential regulators of *Myc* during cell fate induction and differentiation

Next, we examined factors that were predominantly uniquely associated with the upstream enhancer cluster P1/P2, given that the absence of these conserved sequences abrogated expression in larval brain and imaginal tissues (Figure 1B-E). These factors included DOC3, Wdr62, Chip, Mael, Piwi, Nclb/dPWP, Grip91, γSnap1, Pmm2, Pka-C1, RecQ4, and Top2. Potential functions of these factors are described below.

#### The T-box transcription factor Drosocross3 (Doc3) may define the P1/P2 enhancer’s tissue specificity

Drosocross3 (Pflugfelder *et al*. 2017), found exclusively associated with P1/P2 in this study (Figure 14; see also supporting information, Table S8), is a target gene of the TGF-β/Dpp signaling pathway that plays a role in organogenesis during third instar larval and early pupal development (Hatton-Ellis *et al*. 2007). Doc3 is involved in pupal thorax closure (Lu *et al*. 2022) and plays essential roles in proliferation arrest, completion of differentiation, cell fate specification, larval tissue patterning, notum specification, nephrocyte differentiation, limb development (Pflugfelder *et al*. 2017), and cardiogenesis by mediating interplay between Dpp and Wingless inputs (Reim and Frasch 2005). This suggests it may act possibly by controlling *Myc* expression, a known target of the Dpp and Wingless signaling pathways.

**Figure 14.**
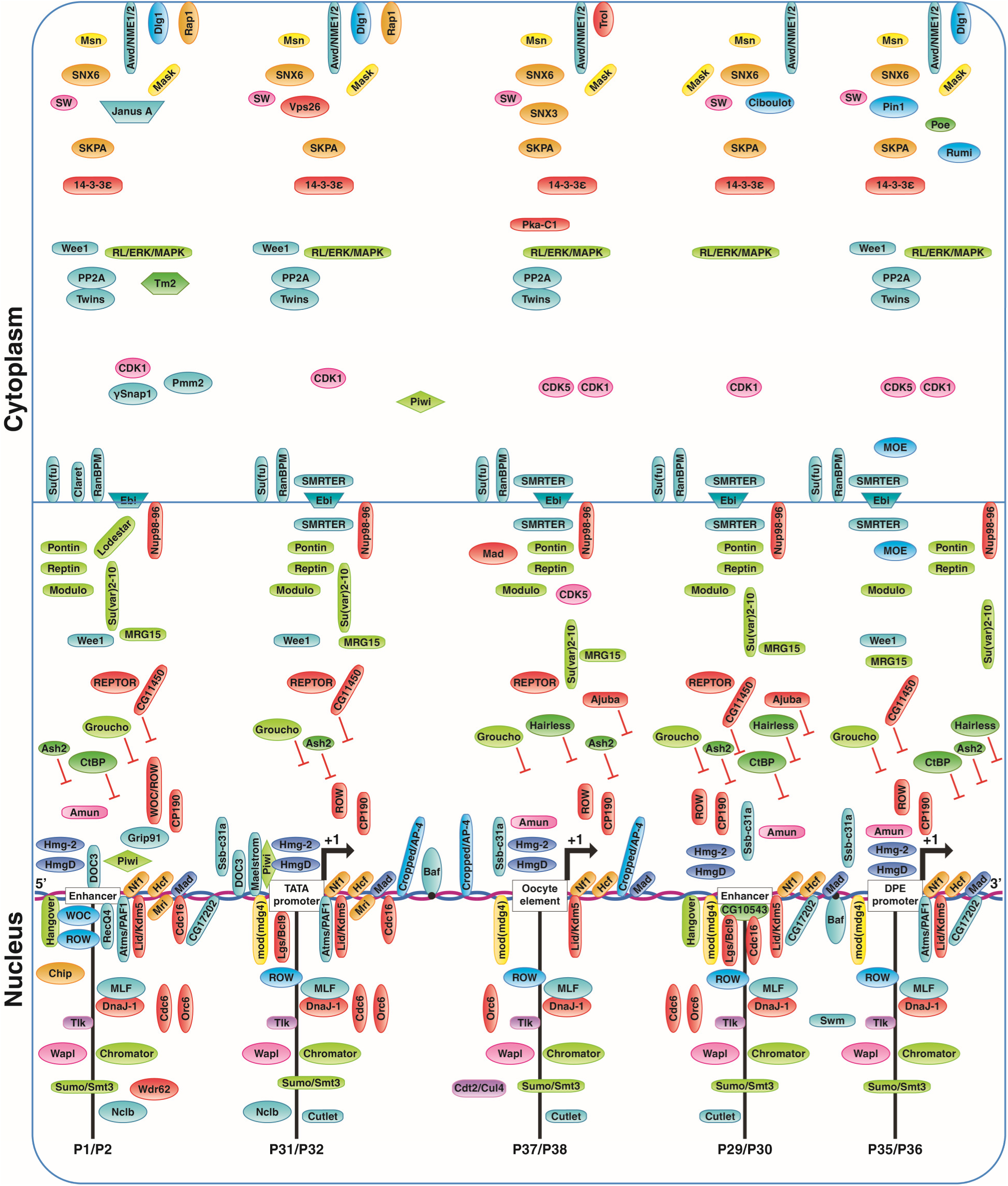
Schematic illustration of signaling pathway components, cell cycle factors, boundary elements, and other gene regulatory proteins potentially involved in *Myc* regulation. Drosocross 3 (Doc3)—an inhibitor of proliferation and pro-differentiation transcription factor—was exclusively associated with the upstream enhancer cluster P1/P2. Modulo (Myc partner for cell growth), High Mobility Group (HMG) proteins HmgD and Hmg-2, Myeloid leukemia factor (Mlf), Host cell factor (Hcf), and transcription factor Mad (which competes with Myc for binding to partners like Max) were associated with all tested *Myc* targets. Barrier to autointegration factor (BAF), with roles in cell cycle and oocyte karyosome formation, was exclusively associated with the start sites P37/P38 and P35/P36. Piwi (a piRNA-interacting factor) was associated with the upstream enhancer P1/P2 and the TATA promoter (P31/P32), while its interactor Maelstrom was only associated with the TATA-containing target. Hairless (a Notch antagonist) was associated with the DPE promoter (P35/P36), the nearby enhancer P29/P30, and the super enhancer cluster P37/P38 (the oocyte element). For more information about the association of these factors with the Myc-CRMs and their abundance ratios, see Table 3 in File S3, Table 7 in File S7, and Table S8.

#### WD repeat domain 62 (Wdr62) may regulate Myc transcription in neuronal progenitor cells

*Wdr62* protein (Figure 14; see also supporting information, Table S8) is a cytoplasmic protein involved in cytoskeleton regulation, the mitotic cell cycle (Ramdas Nair *et al*. 2016; Zeke *et al*. 2016), and centriole duplication (GO Reference Genome Project 2011). A study has shown that the JNK pathway phosphorylates its target Wdr62 in mitotically arrested proneural cell progenitors where Myc protein is downregulated. However, in parallel, phosphorylation of Aurora kinase A (AURK) leads to association between AURK and phosphorylated Wdr62, causing Wdr62 localization to microtubule poles during neuronal differentiation and brain development (Zeke *et al*. 2016). This finding suggests that Wdr62 association with the P1/P2 enhancer might regulate Myc levels during the differentiation of neuroprogenitor cells into brain neurons.

#### The transcription factor Chip (chi) may integrate Notch and Wingless signaling inputs to regulate Myc during cell fate specification and tissue patterning

The LIM-homeodomain-binding transcription factor and enhancer facilitator Chip (*chi*) (Figure 14; see also supporting information, Table S8) is essential for nervous system development, partnering with the Apterous protein (van Meyel *et al*. 2000) and various other Chip-binding partners. Protein complexes containing Chip integrate multiple signaling pathways, such as Notch and WNT/Wg, to induce sensory organ precursor (SOP) cell fate, chaeta morphogenesis, eye development (Biryukova and Heitzler 2005; Roignant *et al*. 2010), wing disc morphogenesis (Rincon-Limas *et al*. 2000), leg development, thoracic bristle formation (Zenvirt *et al*. 2008), and cardiogenesis (Roignant *et al*. 2010). Studies have shown that the *Drosophila* Chip (*chi*) homolog in mouse, LIM domain-binding protein 1 (Ldb1/*ldb1*), which plays essential roles during embryonic development, contains a binding site for MYC, and that Ldb1 physically interacts with MYC (Yamashita *et al*. 1998). Disruption of *ldb1* expression causes embryonic lethality and liver cancer in adult mice (Teufel *et al*. 2010). Microarray analysis revealed *MYC* expression in hepatocellular carcinoma cells, upregulation of the *MYC* downstream target Cyclin D1, and downregulation of p21 (Teufel *et al*. 2010).

#### Maelstrom (mael), potentially acting with Piwi, may regulate Myc expression

HMG box protein Maelstrom (*mael*) (Figure 14; see also supporting information, Table S8) is part of the meiotic complex and is involved in ncRNA-, piRNA-, and miRNA-mediated metabolic processes, gene silencing, heterochromatin formation (Findley *et al*. 2003; Lim and Kai 2007; Pek *et al*. 2009; Sienski *et al*. 2012), cell fate specification and differentiation (Clegg *et al*. 1997; Pek *et al*. 2012; Hayashi *et al*. 2014), and germline maintenance (Lim and Kai 2007; Pek *et al*. 2009). Furthermore, Mael induces malignant transformation in collaboration with Myc and Ras pathways (Kim *et al*. 2017).

P-element induced wimpy testis (*Piwi*) exhibits the ability to silence retrotransposons, partly through physical interaction with piRNAs. Piwi and the HMG protein Maelstrom colocalize with transcribing pol II at numerous genome instability hot spots embedded within transposons (Sienski *et al*. 2012). Repression of transposable elements by Piwi during transcription in ovarian somatic cells depends on the transcription factor Maelstrom, which in turn impacts chromatin organization and regulation (Sienski *et al*. 2012; Ilyin *et al*. 2017). The retrotransposon gypsy insulator is inserted into the first intron within the 5’-UTR of the *Myc* gene, 418 bp upstream of the translation initiation site (Gallant *et al*. 1996). Piwi was associated with the upstream enhancer P1/P2 and the TATA-containing promoter P31/P32, while Mael was only found in association with the P31/P32 promoter. Both Piwi and Mael proteins might play an important role in repressing the gypsy element during early embryonic development and oogenesis, potentially acting from the *Myc* TATA promoter (P31/P32) and the oocyte element (P37/P38) initiation sites.

#### The Drosophila chromatin-associated factor dPWP/Nclb may regulate Myc levels during animal growth in response to nutrients

The chromatin-associated factor (*nclb*, *dPWP*), also known as ‘no child left behind’, functions in the Myc and TORC1 signaling pathway to regulate animal growth in response to nutrients, primarily through pol I-mediated ribosomal gene expression and possibly through pol III transcription (Liu *et al*. 2017). In *Drosophila* larvae, dPWP regulates the localization of Cdk7 to the nucleolus; Cdk7 is a component of the pol I elongation factor TFIIH (Liu *et al*. 2017). Its function is crucial for germline stem cell maintenance in early germ cell cysts, female germ cell differentiation, and egg chamber development (Casper *et al*. 2011). Overexpression of dPWP has been reported in human squamous cell head and neck cancer, correlating with poor prognosis (Liu *et al*. 2017).

#### Gamma-tubulin ring protein 91 (Grip91) may be involved in Myc regulation during meiosis and mitosis

*Grip91* (Figure 14; see also supporting information, Table S8) is a component of the gamma-tubulin structural protein complex (Oegema *et al*. 1999) and plays a crucial role during male meiotic nuclear division and spindle organization (Barbosa *et al*. 2003). During mitosis, Grip91 facilitates spindle assembly, alignment, and segregation of duplicated chromosomes (Tariq *et al*. 2020). Both Grip91 and Myc are important proteins involved in the cell division cycle, cell proliferation, and maintenance of germline divisions.

#### γSnap1 may be indirectly involved in Myc regulation during cell proliferation and differentiation

γSoluble NSF attachment protein 1 (γ*Snap1*) (Figure 14; see also supporting information, Table S8) is an N-ethylmaleimide-sensitive fusion protein involved in synaptic vesicle internalization, protein transport, and activation of Notch signaling during wing D/V patterning (Bejarano *et al*. 2008). Activated Notch stimulates Wingless in the D/V boundary cells, and the combined action of Wingless and Notch signaling coordinates cell proliferation and differentiation in the wing primordium by controlling the activity of target genes such as *Myc* (Giraldez and Cohen 2003).

#### Phosphomannomutase (Pmm2) and Claret might regulate Myc expression during proliferation and tissue patterning

*Pmm2* (Figure 14; see also supporting information, Table S8) plays an essential role in GDP-mannose biosynthetic processes and protein glycosylation at neuromuscular junctions (NMJ). Synaptic Mmp2 glycosylation by Pmm2 occurs through modulation of Wnt/Wg signaling with the involvement of the heparan sulfate proteoglycan coreceptor Dally-like protein (Dlp), thus acting via multiple cascades to control patterning and differentiation during developmental processes and adult life (Parkinson *et al*. 2016). Dpp represses the protooncogene *Myc* and, in an autoregulatory loop, antagonizes the positive effects of Wingless/Wnt signaling on cell proliferation to limit stem cell number and Myc activity (Pennetier *et al*. 2012).

Claret (*ca*) (Figure 14; see also supporting information, Table S8) belongs to the Ras superfamily and the guanine nucleotide exchange factor family of proteins, similar to Ran-GTPase. Claret is essential for eye color biosynthetic processes and eye pigment precursor transport via vesicular trafficking pathways (Reaume *et al*. 1991; Ma *et al*. 2004; Harris *et al*. 2011), regulation of lipid storage (Wang *et al*. 2012), cell cycle progression, and centrosome duplication (Ren *et al*. 1994; Sazer and Dasso 2000). Studies have shown that v-Myc directly binds to the *Ran* promoter to drive expression of the *Ran* gene (Yuen *et al*. 2013). Whether Claret, a member of the Ran-GTPase family, might regulate *Drosophila Myc* expression, or alternatively, whether Myc might be involved in *claret* expression remains to be resolved.

#### cAMP-dependent protein kinase catalytic subunit 1 (Pka-C1) was associated with both the Myc upstream enhancer P1/P2 and the oocyte element P37/P38

Pka-C1 contributes to Anterior/Posterior (A/P) axis pattern specification in larval discs, wing morphogenesis (Pan and Rubin 1995), synaptic transmission (Yao and Wu 2001), and rhythmic behavior (Levine *et al*. 1994). Protein kinases such as PKA belong to *Myc* target genes, and Myc is a phosphorylation substrate for cAMP-dependent PKA species during transcriptional activity, suggesting a putative feedback regulatory loop between PKA kinases and Myc during cell proliferation and differentiation (Boi *et al*. 2023).

#### DNA helicase RecQ4, which plays an essential role during replication in highly proliferative cells, was associated exclusively with the Myc enhancer P1/P2

ATP-dependent DNA helicase (*RecQ4*) (Figure 14; see also supporting information, Table S8) is important for genome stability and DNA metabolism (including duplex unwinding and DNA double-strand break repair) (Capp *et al*. 2009), as well as larval development (Wu *et al*. 2008), the mitotic cell cycle, and oogenesis (Xu *et al*. 2009). Studies have shown that RecQ4 plays an important role in viability and fertility in follicle cells (related to chorion genes) and is actively involved in DNA replication during development (Wu *et al*. 2008). RecQ4 binding to the *Myc* upstream enhancer cluster could impact *Myc* regulation during oogenesis and larval development.

#### The Myc enhancer cluster P1/P2 appears to play an essential role in larval and pupal development

Most factors associated with the enhancer cluster P1/P2 contribute to cytoskeletal structure, tissue patterning, dorsal closure, segmentation, and larval and pupal development. We conclude that the *Myc* enhancer P1/P2 functions predominantly as a tissue-specific patterning enhancer and plays a crucial role at the onset of cell fate specification and differentiation during larval and pupal development. We reason that the factors associated with the P1/P2 enhancer and the basal transcription protein complex integrate developmental signals from pathways such as Wnt/Wg, Notch, Hedgehog, EGFR, and Dpp/BMP (Figure 14 and Figure 15) to drive transcription of target genes in specific cell types to pattern tissues (Gallant *et al*. 1996; Prober and Edgar 2002; Maines *et al*. 2004; Zanet *et al*. 2005; Wu and Johnston 2010; Tavares *et al*. 2017).

**Figure 15.**
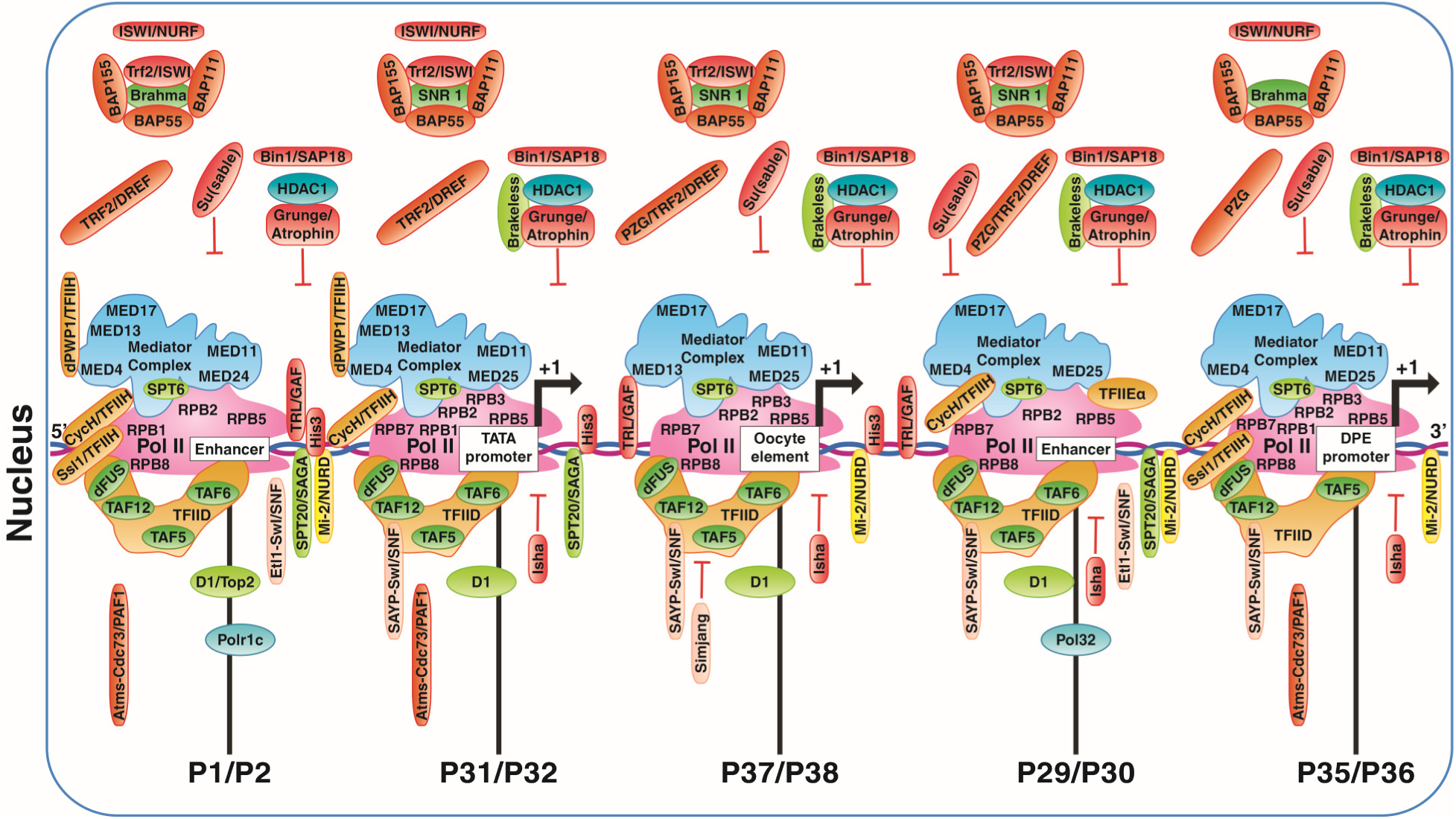
Core promoter and basal transcription factors, chromatin remodelers, and histone modifiers associated with the investigated Myc-CRMs. Subunits of the Mediator complex, transcription factor TFIID complex, and RNA pol II complex were generally detected at the three initiation sites P31/P32, P37/P38, and P35/P36. MED13/Skuld was also found associated with the P1/P2 enhancer cluster in addition to the start sites P31/P32 and P37/P38. Similarly, MED25 was associated with the enhancer cluster P29/P30 as well as the three initiation sites P31/P32, P37/P38, and P35/P36. Components of histone and chromatin modifiers, factors associated with boundary elements, and other transcription factors and cofactors are also illustrated. Most factors are discussed in the Results and Discussion sections. Detailed information regarding the association of these factors with Myc-CRMs and their abundance ratios is available in supplementary materials (Table 3 in File S3, Table 7 in File S7, and Table S8).

### XIII. Core Promoter Factors, Chromatin Modulators, and Gene-Specific Factors Associated with Myc-CRMs

#### RNA polymerase complex

RNA pol II subunit A, RPII215 (*RPB1*), is the largest subunit of the RNA pol II complex. The RPB1 subunit serves as a scaffold for the assembly of basal transcription factors and is involved in transcription initiation (Zehring *et al*. 1988; Brickey and Greenleaf 1995; InterPro Project Members 2004) and elongation (Brickey and Greenleaf 1995).

RNA pol II subunit B, Polr2B (*RpII140*, *RPB2*), is the second largest component of the pol II core complex and is essential for RNA polymerase transcriptional activity (Falkenburg *et al*. 1987; Aoyagi and Wassarman 2000) at nearly all start sites (Greenleaf 1983; Hamilton *et al*. 1993).

RNA pol II subunit C (*Polr2C*, *RPB3*) responds to heat shock and triggers transcription of the Hsp70 gene (Yao *et al*. 2007).

RNA pol II subunit E (*Polr2E*, *RPB5*) possesses polymerase activity for transcription from pol II, pol I, and pol III promoters, contributing to tRNA synthesis by pol III and transcription elongation by pol I (Aoyagi and Wassarman 2000; Gene Ontology Curators 2002; GO Reference Genome Project 2011).

RNA pol II subunit G (*Polr2G*, *RPB7*) binds to ssDNA and RNA and is involved in gene expression regulation, translation initiation (GO Reference Genome Project 2011), and pol II transcription (Aoyagi and Wassarman 2000).

RNA pol II, I, and III subunit H (*Polr2H*, *RPB8*), shared by pol I, II, and III, is capable of 5’-3’ polymerase activity (InterPro Project Members 2004) and participates in pol I, pol II, and pol III transcription (Aoyagi and Wassarman 2000; Gene Ontology Curators 2002; GO Reference Genome Project 2011). Polr2H is involved in tRNA transcription by pol III (Gene Ontology Curators 2002).

A wide array of literature confirms the binding of mammalian MYC and *Drosophila* Myc to proximal promoter regions and their interaction with the RNA pol II complex (Grewal *et al*. 2005; Raha *et al*. 2010b; de Pretis *et al*. 2017; Baluapuri *et al*. 2019).

#### TBP-TFIID subunits

We identified TBP-associated factors TAF5, TAF6, and TAF12 associated with the tested Myc-CRM samples. These TAFs were present in samples corresponding to the enhancer clusters P1/P2, P29/P30, P37/P38, and the TATA-containing promoter P31/P32. Notably, TAF6 was absent from the DPE-containing Myc-CRM P35/P36 sample (Figure 15). TAF5 and TAF6 associate with the TFIID complex (Hansen and Tjian 1995; Wright *et al*. 2006; GO Reference Genome Project 2011);TAF12 recruits TAF4, and TAF6 recruits TAF9 to the TBP/TFIID complex to provide stability to the pre-initiation complex (Yokomori *et al*. 1993). A core TFIID complex consists of TAF4, TAF5, TAF6, TAF9, and TAF12, forming a stable pre-initiation complex (Wright *et al*. 2006). We propose that robust transcriptional bursting from *Myc* start sites likely requires the assembly of these essential TFIID components. TAF5 and TAF6 are important mediators of promoter responses to protein complexes associated with enhancers (Scheer *et al*. 2012), regulating developmental transcription (Aoyagi and Wassarman 2000). Additionally, TAF6 and TAF12 are components of the SAGA complex, which possesses histone acetyltransferase (HAT) and ubiquitin protease capabilities (Guelman *et al*. 2006b). The HAT activity of SAGA and its interaction with TBP and bound transcription factors is essential for promoter opening and pol II promoter escape (Hansen and Tjian 1995; Wright *et al*. 2006). The SAGA protease activity degrades unwanted transcription factors and enhances tissue-specific transcription in certain cell types, inducing differentiation initiation. SAGA is enriched at the 5’-end of genes, indicating that paused SAGA can immediately facilitate reinitiation for further rounds of transcription (Weake *et al*. 2011). We conclude that transcription of *Myc* as an immediate early gene likely relies on the assembly of TFIID core components at its initiation sites and potentially the presence of a poised SAGA complex near the 5’-end. The pol II pre-initiation complex and SAGA, together with factors bound to upstream and downstream *Myc* enhancers, integrate information from developmental signals into transcriptional regulatory programs controlling *Myc* transcription.

#### Trf2/DREF and Trl/GAF as potential regulators of Myc

TBP-related factor 2 (Trf2, TLF), a general transcription factor associates with the DNA replication element binding factor DREF (BASHIRULLAH *et al*. 2007; SHIMA *et al*. 2007) (TRF2/DREF), recognizes and binds specifically to core promoter sequences (OHLER *et al*. 2002; KEDMI *et al*. 2014; WANG *et al*. 2014), and interacts with the basal transcription machinery and transcription factors-occupied enhancers to modulate gene expression. For example, via interaction with components of the ecdysteroid pathway, TRF2/DREF controls the regulation of developmental gene expression during spiracle morphogenesis (Shima *et al*. 2007) and pupal development (Hochheimer *et al*. 2002; Bashirullah *et al*. 2007). TRF2/DREF is an important transcriptional regulator of cell cycle components such as Myc and E2F, thus playing an important role in controlling cell division and tissue size (Hyun *et al*. 2005). *Drosophila Myc* is also a known regulator of DREF (Bellosta and Gallant 2010) in S2 cells, ovaries, and imaginal discs (Thao *et al*. 2008).

Trithorax-like (TRL), the GAGA transcription factor (GAF) (Trl/GAF), may also regulate *Myc*. Trl/GAF, a TrxG group C2H2 zinc finger transcription factor, binds specifically to core promoters (Agianian *et al*. 1999). Trl/GAF recruits the Polycomb group (PcG) complex, inducing chromatin remodeling (Farkas *et al*. 1994; Espinas *et al*. 1999; Bonchuk *et al*. 2011; Ogiyama *et al*. 2018). TrxG and Polycomb groups are both essential for chromatin-mediated transcription of homeotic and developmental genes (Matharu *et al*. 2010). In the ovaries, TRL/GAF is essential for follicular cell morphogenesis and germline cell differentiation; mutations cause female sterility (Ogirenko *et al*. 2008). Trl/GAF was enriched at the distal enhancer cluster P1/P2 and the initiation sites P31/P32 (TATA-containing) and P37/P38 (oocyte element). We propose that TRL/GAF might regulate *Myc* transcription, potentially involving the PcG complex during development.

#### Cabeza (caz/dFUS), a member of the TFIID complex, is a potential Myc regulator

Cabeza (*caz*, *dFUS*), also known as sarcoma-associated RNA binding fly homolog (SARFH) and P19 (Immanuel *et al*. 1995), a member of the TFIID group (hTAFII68/dCabeza) (Aoyagi and Wassarman 2000), is the sole *Drosophila* homolog of human TLS/FUS (Bertolotti *et al*. 1996; Mallik *et al*. 2018) and binds to ssDNA/RNA. Cabeza can integrate transcription initiation via association with the pre-initiation complex (PIC) (Bertolotti *et al*. 1996; Aoyagi and Wassarman 2000) and gene expression through mRNA processing (Herold *et al*. 2009) during development and adult life (Wang *et al*. 2011; Xia *et al*. 2012; Azuma *et al*. 2014; Shimamura *et al*. 2014; Azuma *et al*. 2018). Abnormalities in protein function result in the severe progressive neurodegenerative disease amyotrophic lateral sclerosis in humans (ALS) (Xia *et al*. 2012; Taylor *et al*. 2016; Mallik *et al*. 2018). In addition to being part of the TFIID complex, like FUS and TAF15, Cabeza belongs to the TET (TAF15-EWS-TLS) or FET (FUS-EWS-TLS) protein family (Chau *et al*. 2016). N-Myc downstream regulated gene-1 (*NDRG1*) is the binding partner of TAF15, colocalizes with TAF15 to the nucleus, and promotes cell proliferation (Li *et al*. 2023). Xrp1, an AT-hook binding transcription factor in *Drosophila* (Blanco *et al*. 2020), interacts genetically with *caz*, and the majority of Xrp1-interacting factors are involved in gene expression (Mallik *et al*. 2018). Cabeza might establish specific subcomplexes of TFIID to mediate differential activation of *Myc* expression by various classes of activators and TAFs during development and disease.

#### The High Mobility Group (HMG) transcription factors HmgD and Hmg-2 might regulate Myc

High mobility group transcription factors HmgD and Hmg-2 (Dragan *et al*. 2003), along with corepressors Groucho (Giagtzoglou *et al*. 2003; Helman *et al*. 2011) and CtBP, are downstream effectors of signaling pathways such as Wnt/Wg (Levanon *et al*. 1998) and TGF-β/Dpp (Dorfman *et al*. 2002). These effectors converge signaling inputs onto the transcription regulator complex at the promoter, mediating the context-dependent activation or repression of target genes like *Myc* and enabling controlled transcription during development (Barker *et al*. 2000; Duman-Scheel *et al*. 2004; Herranz *et al*. 2008).

#### The RNA polymerase II transcription coactivator Ssb-c31a may be involved in Myc transcription

The single-stranded DNA binding protein (*Ssb-c31a*) binds sequence-specifically to the NssBF element, a 26-nucleotide sequence within the long terminal repeat (LTR) of the *D. melanogaster* 1731 retrotransposon, and represses the promoter activity of this retrotransposon (Lacoste *et al*. 1995). As a general transcriptional coactivator, Ssb-c31a interacts with TAFs to connect upstream transcriptional activators to the general transcription machinery (InterPro Project Members 2004), which suggests potential involvement of Ssb-c31a in *Myc* regulation.

#### The insulator protein Isha was identified associated with samples P29/P30, P37/P38, P31/P32, and P35/P36

Suppressor of hairy wing (Su(Hw)) mRNA adaptor protein Isha (*Isha*, *CG4266*) associates physically with the *Su(Hw)* mRNA in vitro and in vivo. Isha binds to and recruits *Su(Hw)* mRNA to the insulator protein complex on chromatin and negatively regulates transcription by interacting with both the full RNA pol II complex and the elongating pol II complex (Bag *et al*. 2022). No direct evidence currently exists linking Isha to the regulation of *Myc* transcription.

#### Enhancer of yellow 3 (SAYP/e(y)3) might play a crucial role in forming the initiation complex on Myc promoters

The nuclear protein Enhancer of yellow 3 (*e(y)3*, *SAYP*), a component of the SWI/SNF (Brahma) chromatin remodeling complex (Chalkley *et al*. 2008; Barish *et al*. 2020) and the TFIID complex, activates yolk protein expression and functions as a transcriptional regulator (Shidlovskii *et al*. 2005). SAYP can activate euchromatic genes on polytene chromosomes during embryogenesis and oogenesis and repress activity from heterochromatic regions (Shidlovskii *et al*. 2005). TBP has been reported to interact with c-MYC to propagate transcription through assembly of the initiation complex at the start site (Wei *et al*. 2019). SAYP can combine Brahma with TFIID to form a coactivator supercomplex for efficient transcription initiation (Vorobyeva *et al*. 2009). SAYP might play a crucial role in transcription initiation from both the *Myc* TATA and DPE promoters.

#### Components of the polymerase-associated factor 1 complex (PAF1C) were associated with the P29/P30 enhancer cluster

Another transcription unit (*Atu*), an ortholog of human LEO1, is part of the Cdc73/PAF1 and PAF1C complexes (Chen *et al*. 2015; Goodman *et al*. 2019). PAF1C is recruited to phosphorylated RNA pol II through the PAF1 complex and is involved in releasing the proximal-promoter pause and in transcription elongation (GO Reference Genome Project 2011; Qiu *et al*. 2012). PAF1C might interact with DPE-bound Antimeros (a PAF1 component) to facilitate histone modifications and pol II pause release (Adelman *et al*. 2006). The 14-3-3zeta protein Leonardo (Leo/*14-3-3zeta, leonardo, leo*), existing as isoforms resulting from alternative mRNA splicing, functions as an effector of multiple signaling pathways, primarily the Ras/MAPK cascade and cell cycle control (Li *et al*. 1997; Yano *et al*. 2006; Ashton-Beaucage *et al*. 2014). Leo is a component of the RNA polymerase-associated factor 1 complex (PAF1C) and was found associated with both the enhancer cluster P29/P30 and the target P35/P36 (DPE-containing). Leo is involved in recruiting Myc to open promoters of target genes for *Myc*’s transcriptional activity during cell growth and proliferation (Gerlach *et al*. 2017). Taken together, Antimeros/PAF1, Atu/Cdc73/PAF1, and Leo/PAF1C might contribute to proximal pause release and regulation of transcription elongation from the *Myc* DPE promoter.

#### Insulator proteins, including Modifier of mdg4 (Mod(mdg4)) and CP190, might modulate Myc transcription

Modifier of mdg4 (*Mod(mdg4)*) functions in various processes including male meiosis, chromosome segregation (Soltani-Bejnood *et al*. 2007; Matsui *et al*. 2011), programmed cell death (Harvey *et al*. 1997), chromatin organization (Gerasimova *et al*. 1995), regulation of transcription (GO Reference Genome Project 2011), and enhancer blocking via interaction with Su(Hw) and CP190 (Gerasimova *et al*. 1995; Savitsky *et al*. 2016).

Centrosomal protein 190 kD (*CP190*), a BTB/POZ domain transcription factor (Bonchuk *et al*. 2011), functions as an insulator protein (Chodagam *et al*. 2005). CP190 is involved in establishing heterochromatic genomic boundaries with enhancer-blocking effects (Cuartero *et al*. 2014) through physical interaction with DNA-binding proteins such as CTCF, Su(Hw) (Mohan *et al*. 2007), and Mod(mdg4) (Pai *et al*. 2004; Capelson *et al*. 2010). The insulator activity of CP190/CTCF/Su(Hw) links homeotic gene regulation to body patterning during development (Mohan *et al*. 2007). Given the presence of the gypsy element within the 5’-UTR of the *Myc* (see Section XII), the function of CP190/CTCF/Su(Hw) insulator complex might be relevant to *Myc* transcriptional regulation during development.

#### The protein Relative Of Woc (ROW) might be involved in regulating Myc transcription

The putative zinc finger transcription factor Relative Of Woc (ROW/*row*) plays an essential role in the expression of the chromatin protein HP1c, itself essential for gene expression. For ROW binding to methylated H3 and chromatin, and for the regulation of gene expression, HP1c requires both Relative of Woc (ROW) and Without Children (WOC) proteins. Through an autoregulatory transcriptional feedback loop (Font-Burgada *et al*. 2008; Abel *et al*. 2009), HP1c can remodel chromatin and fine-tune pol II transcription. Myc has a global impact on chromatin structural changes within multiple genomic loci via the recruitment of chromatin modifying complexes during transcription (Knoepfler *et al*. 2006), which suggests a putative role for ROW and WOC in regulating *Myc* transcription.

#### Bicoid interacting protein 1 (Bin1/SAP18), associated with histone deacetylase complexes, recruits the Sin3A/HDAC1 repressor complex to silence target gene promoters

Bin1/SAP18 (*Bin1, SAP18*), the histone deacetylase complex subunit SAP18, facilitates the formation of the Sin3A-HDAC1 corepressor complex (Matyash *et al*. 2009) to silence the transcription of Bicoid target genes at the tip of the embryo during germband retraction (Zhu *et al*. 2001). SAP18 interacts with GAGA factor (GAF), and SAP18/GAF associates with Polycomb group (PcG) response elements (PREs) at the loci of silenced homeotic genes of the bithorax complex to activate transcription (Espinas *et al*. 2000). The SAP18-mSin3-Su(fu) repressor complex inhibits Hedgehog signaling through repression of Gli-mediated transcription and induction of cell fate commitment and morphogenesis (Cheng and Bishop 2002). MYC can recruit the Sin3A-HDAC1 protein complex to the promoters of certain target genes, such as the cell cycle inhibitor p21, to silence these genes (Liu *et al*. 2023a). Therefore, SAP18 function might be required for regulating *Myc* transcription during cell proliferation and differentiation in development.

#### Chromatin assembly factor 1 subunit p55 (Caf1-55), with roles in replication and transcription, may regulate Myc

Caf1-55 (also known as Nurf55/p55) was associated with all tested Myc-CRMs. *Drosophila* Caf-1 (*dCaf-1*), containing p55 as a main component and part of a multiprotein complex, is involved in DNA replication, chromatin assembly, and histone modification (Tyler *et al*. 1996). The Caf-1/p55 complex participates in nucleosome assembly onto newly synthesized DNA and, in collaboration with the histone deacetylase complex Sin3, induces H3K27me3 histone marks on chromatin during replication and transcription, contributing to the regulation of transcriptional patterning during development (Tyler *et al*. 1996; Spain *et al*. 2010). Caf-1 binds to transcriptional *cis*-regulatory regions and is involved in cell cycle regulation via replication-coupled histone modification and gene regulation (Yao *et al*. 2018b). Its association with H3, H4, and the Polycomb Repressive Complex 2 (PRC2) contributes to cell survival, segment identity, and tumor suppression (Anderson *et al*. 2011; Nowak *et al*. 2011). Caf1-55 was associated with all tested Myc-CRMs, suggesting a role for Caf1-55 in the epigenetic regulation of *Myc* transcription during development and disease.

#### Iswi and MRG15 (discussed below) are potential regulators of Myc expression in different biological settings

*Iswi*, an ATPase component of the NuRF complex (Badenhorst *et al*. 2005) and part of several other chromatin remodeling complexes (Corona *et al*. 2000; Eberharter *et al*. 2001), plays a crucial role during pol II transcription via induction of changes to chromatin structure, histone octamer sliding, and replication-dependent chromatin assembly (Ito *et al*. 1997; Varga-Weisz *et al*. 1997; Hamiche *et al*. 1999; Ito *et al*. 1999; Eberharter *et al*. 2001; Langst and Becker 2001; Burgio *et al*. 2008; Emelyanov *et al*. 2010). The NuRF complex interacts with the *c-MYC* promoter to modulate *c-MYC* transcription during hematopoietic stem cell fate commitment and differentiation (Xia *et al*. 2014), and *c-MYC* is often misexpressed in many cancer types (Zhan *et al*. 2019). In *Drosophila*, the NuRF complex components NURF-38 and NURF-55 interact with Myc/Max heterodimers to regulate growth through insulin receptor and mTOR signaling (Liu *et al*. 2020).

The MORF-related gene 15 (*MRG15*) belongs to the Tip60/NuA4-HAT chromatin remodeling complex and is involved in the acetylation of phosphorylated H2AV and its exchange with the unmodified form at DNA lesions (Kusch *et al*. 2004). MRG15 assists repair-dependent chromatin remodeling (Kusch *et al*. 2004), positive regulation of gene expression, and transcription (InterPro Project Members 2004; Huang *et al*. 2017). The Myc and c-MYC network proteins are substrates for Tip60/NuA4 within different biological settings, such as neuroblast differentiation, establishment of cell polarity, regulation of cell cycle genes and replication, and stem cell self-renewal (Patel *et al*. 2004; Ravens *et al*. 2015; Rust *et al*. 2018; Zhao *et al*. 2018), suggesting that MRG15 might regulate *Myc* expression as part of the Tip60/NuA4 complex.

#### Snf5-related 1 (Snr1/BAP45) might be involved in regulating Myc expression

Snr1 (also known as BAP45) functions as a tumor suppressor protein during the S phase of the cell cycle by suppressing *Myc* target genes, including Cyclin E (CycE) (Brumby *et al*. 2002). By binding to the SET domain of proteins and mediating methylation of lysine residues on histone tails, Snr1 maintains the expression of a subset of genes important for cell differentiation and tissue patterning, while simultaneously suppressing factors like Myc (Marenda *et al*. 2003; Marmorstein 2003; Terriente-Felix and de Celis 2009; Curtis *et al*. 2011; Koe *et al*. 2014).

#### Brahma associated protein 111 kD (BAP111/BAF57) is a potential regulator of Myc transcription from multiple initiation sites

The DNA binding protein Bap111 (also known as BAF57) is a component of the Brahma Associated Proteins (BAP) complex (Kal *et al*. 2000; Mohrmann *et al*. 2004; Chalkley *et al*. 2008; Barish *et al*. 2020; Mashtalir *et al*. 2020), the Polybromo-containing protein complex, and associates with High Mobility Group (HMG) box transcription factors (Papoulas *et al*. 2001). BAP111 is involved in chromatin remodeling and induction of transcriptional bursting at the promoters of developmental genes, potentially including *Myc*, through the integration of information from different signaling pathways (Collins and Treisman 2000; Marenda *et al*. 2003; Luo *et al*. 2011; Vinayagam *et al*. 2013; Song *et al*. 2017).

#### BAP155/Moira, potentially along with SPT20, may regulate Myc transcription from multiple sites

Brahma-associated protein 155 kDa (BAP155/Moira), a component of the SWI/SNF (Brahma) chromatin remodeling complex and the trithorax group (TrxG) (Crosby *et al*. 1999; Moller *et al*. 2005). BAP155 plays a crucial role in the transcriptional regulation of human and *Drosophila MYC*/*Myc* as key targets during embryonic development and disease (Shi *et al*. 2013; Guo *et al*. 2022; Chambers *et al*. 2023). In *Drosophila*, BAP155 is involved in wing disc patterning (Terriente-Felix and de Celis 2009; Curtis *et al*. 2011), negative regulation of neuroblast proliferation (Janssens *et al*. 2014), and negative regulation of the G1/S cell cycle transition via downregulation of S phase Cyclin E (Brumby *et al*. 2002).

SPT20 (*Spt20*) is a bifunctional component of the SAGA complex involved in histone acetylation, chromatin remodeling, and activation of pol II transcription (Gene Ontology Curators 2002; InterPro Project Members 2004; GO Reference Genome Project 2011). SPT20/SAGA contributes to tissue-specific protein degradation during patterning via ubiquitin protease activity (Weake *et al*. 2011) and differential transcription regulation through interaction with TBP and tissue-specific transcription factors (Weake *et al*. 2009). SPT20 was associated with the upstream enhancer cluster P1/P2, the downstream intronic enhancer P29/P30, and the TATA-containing P31/P32 Myc-CRMs, suggesting a putative role for SPT20 in regulating *Myc* transcription.

#### The histone methyltransferase complex component Set1/Ash2 is potentially involved in regulating Myc transcription

The DNA binding and transcriptional coregulator enzyme Ash2 (Perez-Lluch *et al*. 2011) (absent, small, or homeotic discs 2) (*ash2*) is a member of the histone methyltransferase (HMT) complex and the trithorax-group (TrxG) family. Ash2 plays an essential role in H3K4 trimethylation (H3K4me3) via interaction with the Sin3A/HCF complex to remodel chromatin and regulate gene expression, partly in response to ecdysone and EGFR signaling (LaJeunesse and Shearn 1995; Angulo *et al*. 2004; Beltran *et al*. 2007; Carbonell *et al*. 2013). Ash2 has been reported to contribute to the regulation of developmental genes such as *MYC* and has been found in protein complexes containing MYC (Luscher-Firzlaff *et al*. 2008; Wan *et al*. 2013). In protein interaction experiments, Ash2 coimmunoprecipitated with FUSE binding protein (FBP1/FUBP1); both are part of the SET1 (COMPASS) multiprotein complex, and FUBP1 interacts with the far upstream element (FUSE) of the *MYC* gene and modulates *MYC* regulation (Duncan *et al*. 1994; Zaman *et al*. 2014).

Thus, we identified an array of chromatin remodeling factors converging on *Myc cis*-elements, likely contributing to *Myc* regulation. Myc and its cofactors are important modifiers of chromatin architecture during cell growth (Marinho *et al*. 2011). Chromatin remodeling processes, genomic architectural changes, and *Myc* expression are tightly intertwined during development and disease.

### XIV. Factors regulating *Myc* from TATA promoter

Factors associated with the TATA promoter belong to a wide array of protein complex groups with vital functions governing chromatin dynamics, pol II function, trans-acting factor activity, and RNA processing. Proteins RPB1, RPB2, RPB3, RPB5, RPB7, and RPB8 constituted the RNA pol II complex. Subunits of the TFIID/MED complexes included TAF5, TAF6, TAF12, MED4, MED11, MED13/Skuld (TRAP), MED17, and MED25. Factors associated with TFIIH included Cyclin H (*CycH*) and dPWP. Other factors potentially important for transcription included Cabeza/dFUS, *SAYP*, Trf2, Bin1/SAP18, HDAC1, Snr1, mod(mdg4), CP190, Isha, SWI, MRG15, Set1/Ash2, Bap111, TRL, SPT20, and BAP155/Moira.

#### Basal transcription machinery regulating Myc from TATA promoter

The largest subunit of RNA pol II, RPB1, important for initiation (Zehring *et al*. 1988; Brickey and Greenleaf 1995; InterPro Project Members 2004) and elongation (Brickey and Greenleaf 1995), together with the second largest component Polr2B (RPB2), essential for transcriptional activity (Falkenburg *et al*. 1987; Aoyagi and Wassarman 2000) at nearly all start sites (Greenleaf 1983; Hamilton *et al*. 1993), confer stability to the pol II complex.

RPB3 is required for the expression of the heat-induced *Hsp70* gene (Yao *et al*. 2007). Bag1, a downstream target of *Myc*, together with the cofactor Hsp70, forms the Bag1/Hsp70 chaperone complex to counteract Myc’s apoptotic function and drive the cell cycle (Gennaro *et al*. 2019), suggesting evidence that the composition of RNA pol II components links *Myc* control to a multitude of Myc cofactors regulating cellular processes.

RPB7 interacts with open chromatin and, by binding to ssDNA/RNA, connects transcription termination to translation initiation (GO Reference Genome Project 2011) and RNA processing.

The composition of the identified subunits of the RNA pol II complex shows that a functional RNA pol II enzyme can be formed on the *Myc* TATA promoter in a context-dependent manner.

MED24, associated with the upstream enhancer cluster P1/P2, might be recruited to the other identified Mediator subunits (MED4, MED11, MED13/Skuld, MED17, and MED25) present at the TATA promoter for a proper response to *Myc* regulation during development. MED13/Skuld (TRAP) is one of the four subunits of the Mediator complex’s kinase module (containing Kto/Med12, Skd/Med13, Cdk8, CycC), where Cdk8-CycC and MED12-MED13 function as pairs during cell growth and tissue patterning (Loncle *et al*. 2007).

TAF5, TAF6, and TAF12 can potentially recruit further components to the TFIID complex. Depending on the composition of the TFIID, Mediator, and RNA pol II complexes, different pre-initiation complexes (PICs) likely form for the spatio-temporal expression of *Myc* from the TATA promoter.

We propose that a stable RNA pol II complex can potentially form on the *Myc* TATA promoter, contributing to pre-initiation complex stability (Figure 15).

#### TAF-TBP and TAF-Trf2 complexes may regulate Myc transcription from the TATA promoter

Transcription factor MYC functions as a differential regulator of transcription for many target genes to control growth, proliferation, and developmental programs (Kress *et al*. 2015; Zaytseva and Quinn 2017). Similarly, *Myc* patterning might be under the control of differential regulatory switches during cell division and differentiation (Kress *et al*. 2015). During neuronal stem cell renewal and differentiation, both TAF-TBP and TAF-Trf2 complexes function in *Drosophila*, and their target genes include different transcription factors and RNA binding proteins (Neves and Eisenman 2019). During spermatogenesis, TAF7L, paralogous to TAF7, interacts with Trf2 to regulate the expression of postmeiotic genes in a tissue-specific manner (Zhou *et al*. 2013; Herrera *et al*. 2014). Since TAFs are parts of the TFIID and SAGA complexes, and our proteomic analysis detected TAFs and Trf2 associated with the TATA promoter, we propose that selective regulatory programs involving both TAF-TBP and TAF-Trf2 might direct *Myc* transcription from the TATA promoter.

#### Promoter melting, initiation, and elongation on the Myc TATA promoter

Cyclin H (*CycH*), enriched on the TATA-containing P31/P32 Myc-CRM, comprises the Cdk-activating kinase (CAK) subunit of the TFIIH core subcomplex. The heterotrimeric TFIIH subcomplex Cdk7-CycH-Mat1 plays a fundamental role during transcription and cell cycle control (Peissert *et al*. 2020). TFIIH phosphorylation of the RNA pol II C-terminal domain (CTD) enhances Mediator dissociation from the pre-initiation complex (PIC) and pol II promoter escape (Larochelle *et al*. 2001; Wong *et al*. 2014). These activities, along with TFIIH’s roles in cell cycle control and DNA excision repair, suggest a potential role for TFIIH in regulating *Myc* transcription from the TATA promoter. Suppressor of stem-loop mutation Ssl1 (p44/Ssl1) is part of the TFIIH core subcomplex, and p44/Ssl1 associates with the XPD subunit of TFIIH to increase the helicase activity of XPD and facilitate promoter clearance (Wang *et al*. 1995). The P1/P2 enhancer, which was found associated with p44/Ssl1 in our dataset, might fold onto the TATA promoter and contribute to promoter clearance at the pre-initiation complex during *Myc* transcription.

Enhancer of yellow 3 (*SAYP/e(y)3*), a component of the SWI/SNF (Brahma) complex and the TFIID complex, is a multidomain-containing coactivator that can combine Brahma chromatin remodeling function with the TAF9-containing TFIID complex to form a coactivator supercomplex, enabling efficient transcription initiation during early development (Shidlovskii *et al*. 2005).

SPT6, a pol II elongation factor and histone H3 chaperone (Adelman *et al*. 2006), recruits the SPT5 component of the DSIF elongation factor to the transcription regulatory apparatus and facilitates Cdk7-dependent loading onto the RNA pol II complex (Baluapuri *et al*. 2019). MYC function is required during pol II pausing, pause release, and processivity of elongation for most, if not all, transcribed genes (Baluapuri *et al*. 2019). MYC binds directly to SPT5 and engages with promoters to load SPT5 onto RNA pol II in a Cdk7-dependent manner, facilitating processive transcription elongation (Baluapuri *et al*. 2019). Conversely, in MYC-overexpressing tumor cells, MYC sequesters SPT5 into inactive protein complexes to block the transcription of anti-proliferative genes such as P21 (Baluapuri *et al*. 2019). SPT6 was enriched in all tested *Myc*-CRM samples; however, SPT5, although identified in the initial dataset, was not significantly enriched (see Table 3 in File S3). We propose that SPT6 might interact with the elongation complex on the *Myc* promoter, potentially recruiting SPT5 for processive elongation of *Myc* transcripts.

Topoisomerase type II (Top2) is an important enzyme responsible for removing supercoils that emerge during meiotic and mitotic crossovers and transcription (Lee *et al*. 1989; Strick *et al*. 2000). Top2 accommodates the passage of DNA strands during transient double-strand break repair by unwinding and relaxing DNA to detangle chromosomes (Strick *et al*. 2000). Collaboration between MYC and Top2 has been well documented during development and disease (Dang 2012; Matias-Barrios and Dong 2023). Top2 and the chromatin condensation factor Barren interact *in vivo* with the Polycomb group (PcG) and bind to PcG response elements and bithorax binding sites to control epigenetic expression of homeotic genes (Lupo *et al*. 2001). Top2 and Barren were enriched exclusively in the upstream enhancer P1/P2 sample (see Table 3 in File S3, Table S8), which might indicate an involvement of Top2 in the epigenetic control of *Myc* expression from the TATA promoter via interaction with Barren and components of PcG.

The chromatin repressive complex component SAP18, which recruits HDAC1, was associated with *Myc cis*-elements and plays context-dependent roles during embryonic body patterning. SAP18 interacts with Kruppel and recruits an enhancer-specific Sin3A/HDAC1 repressor complex (Matyash *et al*. 2009) to repress the transcription of certain genes, such as Bicoid, during embryonic germband extension (Zhu *et al*. 2001). SAP18/GAF associates with Polycomb response elements (PREs) to activate transcription of silenced homeotic genes within the bithorax complex (Espinas *et al*. 2000). The SAP18-mSin3-Su(fu) repressor complex inhibits Gli-mediated transcription during morphogenesis (Cheng and Bishop 2002). MYC can recruit the Sin3A-HDAC1 complex to the promoters of certain targets to inhibit transcription (Liu *et al*. 2023a), suggesting a potential role for the SAP18-Sin3A-HDAC1 complex in regulating *Myc* transcription during development.

Set1/Ash2, part of the COMPASS complex, is involved in H3K4 mono- and dimethylation, thereby providing chromatin accessibility to transcription factors, adjusting transcription fidelity, and enhancing splicing events (Collins *et al*. 2019; Zhu *et al*. 2024). The mammalian far upstream element (FUSE) binding protein 1 (FBP1) interacts with Set1/Ash2 to activate transcription and DNA repair (Zaman *et al*. 2014). Psi, the sole *Drosophila* homolog of mammalian FBP1-3, associates with the pol II complex and *Myc* intergenic sites to activate transcription during cell growth and morphogenesis (Guo *et al*. 2016). FBP1 and Psi both bind ssDNA, suggesting a potential role for Set1/Ash2 in *Myc* regulation through activators such as Psi.

Half pint (*hfp*), an ssDNA/RNA binding protein and the *Drosophila* homolog of Fuse Interacting Repressor (FIR), associates with the transcription machinery. Hfp/FIR counteracts Psi function to inactivate *Myc* (Guo *et al*. 2016) and enhance alternative transcript processing. Ash2, Psi, and Hfp were found associated with all investigated Myc-CRMs.

#### Termination transcription initiated from the Myc TATA promoter

Cpsf160, AAUAAA recognition factor was associated with the three upstream enhancer clusters P1/P2 and plays a key role in RNA 3’-end formation (GLOVER-CUTTER *et al*. 2008) and pre-mRNA polyadenylation via interaction with the poly(A) polymerase complex (GLOVER-CUTTER *et al*. 2008). Cpsf160 and partners such as CPSF6, CPSF73, and CPSF100 colocalize with and bind to the *c-MYC* mRNA during 3’-end formation and mRNA degradation by the FBW7 proteasomal pathway (Glover-Cutter *et al*. 2008; Tran *et al*. 2014; Sim *et al*. 2024). Cpsf160 might be involved in transcription termination from the TATA promoter and in polyadenylating nascent *Myc* transcripts.

The translation factor Poly(A)-binding protein (PABP), found associated with all tested *Myc cis*-elements, couples transcription termination and pre-mRNA processing to translation initiation, predominantly during oogenesis and the maternal-to-zygotic transition (MZT) (Mount and Salz 2000; Clouse *et al*. 2008; Herold *et al*. 2009; Wang *et al*. 2017). PABP interacts with quadruplex G4 DNA-binding factors during transcription and alternative polyadenylation (Li *et al*. 2020), and the *MYC* promoter contains G4 structures known to play roles during replication, transcription, and polyadenylation (Siddiqui-Jain *et al*. 2002). PABP might be important for transcription termination and alternative polyadenylation of *Myc* transcripts.

### XV. Factors regulating *Myc* transcription from the DPE promoter

We aimed to identify factors binding to the downstream promoter element (DPE), contained within Myc-CRM P35/P36. Reporter studies previously showed that the enhancer P29/P30, located proximal to the DPE, is necessary for the DPE promoter to drive *Myc-lacZ* patterning (Figure 2A). Consistent with this, we identified a subset of factors associated with both of these *cis*-elements that might cooperatively initiate *Myc* transcription from the DPE promoter.

#### Pre-initiation complex (PIC) and initiation at the Myc DPE promoter

RNA pol II subunits previously identified associated with the TATA promoter were also present in the DPE-containing Myc-CRM P35/P36 sample, providing evidence for the assembly of a stable RNA pol II complex forming on the DPE promoter (Figure 15).

Associated TFIID subunits included TAF5 and TAF12, while associated Mediator subunits included MED4, MED11, MED17, and MED25. TAF6 and MED13/Skuld (TRAP), which were associated with the TATA-containing P31/P32 promoter, were notably absent from the DPE sample. Other components of the Mediator kinase module (often considered accessory), such as MED12/Kto, Cdk8, and CycC, were also absent from the DPE sample. The Mediator kinase module (containing MED12/Kto, MED13/Skuld, Cdk8, CycC) can function as a repressor; for example, MED12/Kto and MED13/Skuld repress the Hox gene *Ubx* during developmental switches, similar to PcG silencing of certain Hox genes (Gaytan de Ayala Alonso *et al*. 2007). This result suggests that repression of *Myc* transcription from the DPE promoter might not occur via the Mediator inhibitory kinase module.

#### The nearby Myc downstream enhancer cluster P29/P30 might modulate transcriptional bursting from the DPE promoter

We identified TAF6 and TFIIEα associated with the P29/P30 enhancer cluster proximal to the DPE. In humans and yeast, TAF6 reportedly recruits TAF9 to the PIC, and in flies, to the TFIID complex (Weake *et al*. 2009).

Transcription factor TFIIEα contacts TAF6 (Ohkuma *et al*. 1991), and TFIIEα/TFIIEβ interacts with RNA pol II and GTFs *in vitro* (Wang *et al*. 1997). TFIIE is essential for PIC formation, activation of the PIC via promoter melting, and the initiation of elongation (Okamoto *et al*. 1998). TFIIEα can either associate with TFIIH and enhance TFIIH function or function independently from TFIIH (Okamoto *et al*. 1998). TFIIE was shown to be among the class II transcription factors associating with pol II and other GTFs on the *c-MYC* promoter to trigger initiation *in vitro* (Szentirmay and Sawadogo 1991).

From the TFIIH complex, we identified Cyclin H, a component of the kinase subcomplex required for pol II-CTD phosphorylation, PIC destabilization, and pol II promoter escape (Larochelle *et al*. 2001; Wong *et al*. 2014). Another identified TFIIH subunit, p44/Ssl1, together with the XPD subunit, elevates TFIIH helicase activity and assists pol II promoter clearance (Wang *et al*. 1995).

We propose that TAF6 associated with the enhancer cluster P29/P30 might complement the GTFs and the pol II complex bound to the DPE to form a stable PIC. The factors TFIIEα, Cdk7-CycH/TFIIH, and Ssl1/TFIIH associated with the enhancer P29/P30 could potentially contribute to promoter melting, pol II CTD phosphorylation, dissociation of the Mediator/TFIID complex, activation of the PIC, pol II promoter escape, and transcription initiation.

Finally, we identified the general transcription factor TRF2 associated with the P29/P30 enhancer. TRF2 is a core promoter recognition factor and part of the TFIID complex (Aoyagi and Wassarman 2000; Andersen *et al*. 2017). This raises the possibility of a TAF-TRF2 association with the DPE promoter being involved in forming an initiation-competent RNA pol II complex.

#### Transcription elongation and termination from the DPE promoter

In addition to TFIIEα and the subunits of TFIIH, we identified several factors that might contribute to coordinating transcription steps from the DPE promoter. Cabeza/dFUS (hTAFII68/FUS), a member of the TFIID group (Aoyagi and Wassarman 2000) and an ssDNA/RNA binding protein, can facilitate transcription initiation via interaction with the pre-initiation complex (PIC) (Bertolotti *et al*. 1996; Aoyagi and Wassarman 2000). Cabeza was associated with all the Myc-CRMs.

Enhancer of yellow 3 (*SAYP/e(y)3*), a component of the SWI/SNF (Brahma) complex and a multidomain-containing coactivator, can couple chromatin remodeling with the TAF9-containing TFIID complex to form a coactivator supercomplex, enabling efficient transcription initiation during early development (Shidlovskii *et al*. 2005). SAYP was associated with both the TATA and DPE promoters.

PAF1 (Antimeros) interacts with the basal transcription machinery and sequence-specific transcription factors, recruits other chromatin factors like SPT6 and FACT, and is involved in H3K4 trimethylation and control of pol II pause release during elongation (Adelman *et al*. 2006; GO Reference Genome Project 2011). PAF1 was associated with the DPE promoter and might regulate *Myc* transcription from the DPE promoter.

Spt6, the central factor of the activated RNA pol II elongation complex, interacts with all four histones and can recruit SPT5, a member of DSIF; Spt6, potentially together with PAF1, contributes to the processivity of pol II elongation (Miller *et al*. 2023). Spt6 was associated with all tested Myc-CRMs, suggesting a crucial role for Spt6 in regulating *Myc* transcription from multiple sites.

Suppressor of sable (Su(sable)), CPSF6, and Poly(A)-binding protein PABP (*pAbp*) were associated with the DPE and the enhancer P29/P30, and CPSF160 was associated with the enhancer P29/P30. Wdr82 and CPSF100 appeared on the original identification list but were not significantly enriched in the final analysis (see Table 3 in File S3). The protein Su(sable) promotes transcription termination of aberrant RNAs (Voelker *et al*. 1991; Turnage *et al*. 2000; Kuan *et al*. 2004; Kuan *et al*. 2009; Brewer-Jensen *et al*. 2016). Su(sable) interacts with Wdr82 to promote transcription termination of heat shock-inducible genes containing repetitive sequences and degradation of RNAs via exosomes (Brewer-Jensen *et al*. 2016). Wdr82 is one of the unique components of the Compass complex (Wdr82-Set1/Compass) in *Drosophila* and enables methylation of H3K4 to activate transcription (Shilatifard 2012). Transcription regulation by MYC depends on MYC’s interaction with transcription regulator complexes and epigenetic factors (Gouw *et al*. 2019). MYC and Wdr82 interact physically, and this complex has been found in certain transcriptional complexes at the promoters of *MYC* targets (Morelli *et al*. 2023). Su(sable) and Wdr82 might be involved in regulating *Myc* transcription termination and the transition to RNA processing by CPSF and PABP complexes.

#### Several transcription factors and coregulators were associated with the P29/P30 enhancer cluster

CG10543, associated exclusively with P29/P30, is a pol II-specific C2H2 Zinc finger transcription factor that binds to a sequence containing a GGGA core in the DNA major groove (Wolfe *et al*. 2000). CG10543 can activate or repress transcription (Gene Ontology Curators 2002; InterPro Project Members 2004) and is essential for gravitaxis in *Drosophila* (Armstrong *et al*. 2006). Notably, the *Myc* target P29/P30 contains a Hairless consensus binding site within which the core sequence GGGA resides (Table S1.3). Hairless itself was associated with both the enhancer P29/P30 and the DPE-containing region P35/P36. CG10543 might be a *Myc* regulator, potentially through modulation of Notch signaling.

Putzig (*pzg*), a sequence-specific transcription factor (Yao *et al*. 2018b), is involved in chromatin activation and transcription of targets of signaling pathways including Notch, Ecdysone, and JAK/STAT in certain proliferating cells (Eggert *et al*. 2004; Kugler *et al*. 2011). For the activation of replication-related genes, the Putzig/Trf2/DREF complex binds to the promoters of these genes (Kugler and Nagel 2007). As mentioned previously, Trf2/DREF regulates key cell cycle components like *Myc* and E2F (Hyun *et al*. 2005). Putzig/Trf2/DREF might regulate *Myc*, possibly during the G1/S phase. Furthermore, a potential repressor complex—perhaps consisting of CG10543, Hairless, Groucho, and CtBP (which were also associated with the P29/P30 enhancer and/or the DPE region)— might counteract Notch activation to control *Myc* transcription during differentiation.

The histone H2A variant (His2Av in *Drosophila*) is present in about 25% of nucleosomes (InterPro Project Members 2004). At DNA lesion sites, phosphorylated His2Av is acetylated by the dTip60 chromatin remodeling complex and exchanged with the unmodified form during DNA repair (Kusch *et al*. 2004). His2Av is required for *Drosophila* adult stem cell differentiation and self-renewal (Morillo Prado *et al*. 2013), and histone variants have been implicated in occupying transcribing regions of the genome in both plants and animals (Conerly *et al*. 2010). In humans, the histone H2A variant H2AZ has been shown to occupy transcription start sites (TSSs). In this study, we identified His2Av associated with the *Myc* P1/P2 and P29/P30 enhancer clusters. In B-cell lymphomagenesis, H2AZ facilitates Myc-induced transcriptional transformation towards tumor cell development (Conerly *et al*. 2010). In *Drosophila*, His2Av was shown to positively regulate IMD signaling to modulate the immune response (Tang *et al*. 2021).

Turandot Z (*Totz*), a secreted peptide involved in the fly stress response (Ekengren and Hultmark 2001), was associated with the DPE promoter in our analysis. This finding, combined with the association of His2Av with the P29/P30 enhancer, suggests a potential collaboration between His2Av and TotZ. This interaction could occur via the transcription complex on the DPE promoter, potentially regulating *Myc* expression in response to various stress-induced stimuli.

The Activator Protein-4/Cropped is a sequence-specific pol II transcription factor (King-Jones *et al*. 1999) involved in growth regulation, cell proliferation, and organ size determination; its activity is modulated by the Sin3A chromatin modifier during larval somatic muscle development (Dobi *et al*. 2014). *AP-4* is a direct downstream target of MYC, and MYC binding to the E-box CACGTG in the first intron of the *AP-4* gene leads to the induction of mitotic cell cycle reentry and maintenance of cell proliferation in a progenitor-like manner (Jung *et al*. 2008). AP-4 overexpression has been reported in several cancer types, such as MYCN-expressing neuroblastoma cells and colorectal cancer with concomitant MYC expression (Jackstadt *et al*. 2013; Xue *et al*. 2016). AP-4 might play a role in regulating *Myc* transcription.

### XVI. Enhancer cluster P37/P38 might transcribe eRNAs in early embryos and ovary

Minimal sequences from the proximal 5’-UTR retaining the P37/P38 enhancer cluster recapitulated endogenous *Myc* expression in embryos and ovary across all tested truncations (Figure 1A-D and F). We therefore aimed to identify proteins associated with P37/P38 that are essential for early development (oogenesis and embryogenesis) to understand how this ‘oocyte element’ functions as a key developmental switch for *Myc* expression. Factors enriched on the P37/P38 target likely play predominant roles in early embryos, oocyte maturation, and cell fate induction. Transcription initiation from the *Myc* super enhancer oocyte element adds another layer to differential bursting from multiple sites.

#### Formation of pre-initiation/initiation complex on the enhancer cluster P37/P38

RPB2 is the second largest subunit with 5’-3’ polymerase activity and is present in all pol II complexes (Falkenburg *et al*. 1987; Aoyagi and Wassarman 2000). RPB3 localizes to the cytoplasm, nucleus, and polytene chromosomes (Yao *et al*. 2007; Pankotai *et al*. 2010) and is involved in transcription by RNA pol II (InterPro Project Members 2004; Yao *et al*. 2007). RPB5 and RPB8, required by RNA pol I, pol II, and pol III, contribute to the synthesis of mRNAs, rRNAs, and tRNAs (Aoyagi and Wassarman 2000; Gene Ontology Curators 2002; GO Reference Genome Project 2011). RPB7 not only enables pol II transcription (Aoyagi and Wassarman 2000) but also binds to mRNA and can initiate translation (GO Reference Genome Project 2011).

Similar to the TATA and DPE initiation sites, we identified TAF5, TAF6, and TAF12 associated with the P37/P38 enhancer cluster. Mediator subunits MED11, MED13/Skuld, MED17, and MED25 were associated with P37/P38. The Mediator complex serves as a hub for linking the pol II preinitiation complex at the start site to transcription factors to modulate gene regulation (Gu *et al*. 2002; Janody *et al*. 2003; Park *et al*. 2003a). MED13/Skuld and Sin3A, as global transcription regulators, are required for cell proliferation, differentiation, and tissue patterning during development (Dobi *et al*. 2014). Pygopus, a key nuclear component of the Wnt/Wg pathway, interacts with MED12-MED13 to associate with the pre-initiation complex, convey signaling stimuli to the RNA pol II complex, and trigger expression of Wnt/Wg target genes (Carrera *et al*. 2008). In other developmental settings, the function of MED13 and MED17 is important for inducing cell fate commitment, segment identity (Boube *et al*. 2000), and tissue patterning (Loncle *et al*. 2007; Bejarano *et al*. 2008; Carrera *et al*. 2008). MED25 plays a role in nervous system development (Koizumi *et al*. 2007) and immune response through NF-κB pathways (Valanne *et al*. 2010). The mRNA binding protein dFUS/Cabeza (*dFUS, caz*) (Lasko 2000), a member of the TFIID complex also found associated with the P37/P38 element, participates in transcription initiation (Aoyagi and Wassarman 2000), termination, and mRNA processing (Herold *et al*. 2009), highlighting its potential broad role in regulating gene expression from this enhancer.

#### Chromatin remodelers and other transcriptional regulators associated with P37/P38

Among these is the nuclear Zinc Finger protein Simjang (*simj*), a highly conserved repressor and regulatory component (p66) of the Mi-2/NURD chromatin remodeling complex (Kon *et al*. 2005). Simjang contributes to transcriptional repression by inducing chromatin remodeling and histone deacetylation (Kon *et al*. 2005) and is required for heart fate map induction (Kim *et al*. 2004) and brain synaptic development (Willemsen *et al*. 2013). Since both the NURD and Sin3 complexes influence *Myc* regulation (Harper *et al*. 1996; Alland *et al*. 1997; Satou *et al*. 2001; Kadamb *et al*. 2013; Hata *et al*. 2019), the Simjang/p66/NURD complex likely plays a role in controlling *Myc* expression during development.

Another factor associated with the oocyte element is the highly conserved DNA binding protein Barrier-to-autointegration factor (Baf), which participates in chromosome condensation and organization (GO Reference Genome Project 2011; Nikalayevich and Ohkura 2015). Specifically in the oocyte nucleus, Baf induces karyosome formation (Lancaster *et al*. 2007), a process aided by the coordinated functions of NURD and phosphorylated Baf in preventing chromosome attachment to the nuclear matrix (Lancaster *et al*. 2007; Nikalayevich and Ohkura 2015). Baf’s association with the P37/P38 element suggests it might modulate *Myc* expression during this specialized nuclear reorganization.

Additionally, we identified the uncharacterized protein CG6026 associated with the oocyte element P37/P38 (see Table 3 in File S3). While its structural features, motifs, and connection to *Myc* remain uninvestigated, CG6026 is annotated as a transcriptional coregulator involved in chromatin remodeling and pol II transcription during pseudohyphal growth (GO Reference Genome Project 2011), hinting at a potential regulatory role at this enhancer. Calmodulin (*Cam*), a calcium sensor, is a component of the Ca^2+^/Calmodulin signaling pathway (Martin and Bayley 2004) with an expression peak during embryonic nervous system development (Andruss *et al*. 1997). Calmodulin is involved in ovarian germline maintenance (Andruss *et al*. 1997), cell movement (Toth *et al*. 2005), and mitosis (Goshima *et al*. 2007). MYC binds to Calmodulin, and high levels of Calmodulin increase the transcription and oncogenic activity of MYC in cell cultures (Raffeiner *et al*. 2017). Calmodulin might be a potential regulator of *Myc* transcription.

Terribly reduced optic lobes (*trol*) encodes the extracellular matrix heparan sulfate proteoglycan Perlecan, which plays a crucial role during fate commitment and tissue patterning through the integration of multiple signaling cascades, including Hedgehog (Park *et al*. 2003b), Wingless/Wg, TGF-β (Lindner *et al*. 2007), FGFR (Park *et al*. 2003b; Dragojlovic-Munther and Martinez-Agosto 2013), and EGFR (Takemura and Nakato 2017). Trol plays a role in maintaining epithelial-cell polarity in ovarian follicle cells (Schneider *et al*. 2006) and epithelial morphogenesis in the egg chamber (Isabella and Horne-Badovinac 2016; Crest *et al*. 2017). For comparison, Dally-like, another heparan sulfate proteoglycan, activates the Dpp pathway to antagonize Wingless signaling and control *Myc* expression during the establishment of larval hematopoietic niche size and progenitor stem cell (PSC) differentiation (Pennetier *et al*. 2012). Trol might function similarly to control *Myc* expression during cell differentiation.

Spaghetti Squash (*sqh*), the regulatory light chain of nonmuscle type 2 myosin II (Wang *et al*. 2023), plays roles in the cell cycle, cell proliferation (Somma *et al*. 2002; Royou *et al*. 2004; Dean and Spudich 2006), morphogenesis (Simoes *et al*. 2006; Sharma *et al*. 2018), embryonic gastrulation, syncytial cellularization (Vasquez *et al*. 2014; He *et al*. 2016), and oogenesis (Fulga and Rorth 2002; Nicolas *et al*. 2009; Sharma *et al*. 2018; Doerflinger *et al*. 2022). Based on these functions, Spaghetti Squash may play an important role as a *Myc* regulator during cell growth, proliferation, and differentiation.

Lethal-(2)-denticleless (*l(2)dtl*) encodes the Cdt2 protein (*l(2)dtl/Cdt2*), an adapter for the Cul4-based E3 ubiquitin ligase crucial for modifying proteins during S phase and for DNA damage repair at the G2 checkpoint (GO Reference Genome Project 2011). Essential for embryogenesis, male fertility, and larval development (Sloan *et al*. 2012), Cdt2 was found associated exclusively with the P37/P38 oocyte element in our study. This specific association suggests Cdt2 might play a role in regulating *Myc* expression from this particular element during early development.

Abnormal wing disc (Awd/*awd*), a nucleoside-diphosphate kinase (Lascu *et al*. 1992) associated with P37/P38, participates in vesicle trafficking, receptor internalization, and GDP/GTP conversion (Morera *et al*. 1995; Ignesti *et al*. 2014); its human homolog *NmE1/2* is essential for tissue patterning (Dammai *et al*. 2003; Hsouna *et al*. 2010; Serafini *et al*. 2020). Highly expressed during oogenesis, Awd maintains follicular epithelial integrity and morphogenesis by modulating adherens junction components (Woolworth *et al*. 2009). Since key *Myc*-regulating pathways like Notch and EGFR depend on Awd/NmE1/2 activity (Dammai *et al*. 2003; Ignesti *et al*. 2014), Awd likely acts as an indirect regulator of *Myc*, particularly during oocyte development and subsequent cell differentiation.

Survival motor neuron (*Smn*), a component of the SMN (BORG *et al*. 2015) and SMN-Gemmin2 complexes (KROISS *et al*. 2008) and involved in a multiprotein interaction network (CAUCHI *et al*. 2008), is an RNA binding protein containing a Tudor domain. It is essential for oocyte maturation and viability, with higher protein levels required for neuromuscular development (CHAN *et al*. 2003). Research using Rat-1 cell cultures indicated that Smn induces apoptosis by activating the *c-MYC* gene, similar to the action of Bcl-2 and IAP-2 (VYAS *et al*. 2002). Coilin, an SMN interactor and member of the large SMN protein complex, localizes to the nucleoplasm (HEBERT AND MATERA 2000). These findings suggest that SMN might regulate *Myc* during oogenesis.

#### Post-transcriptional/-translational factors associated with the oocyte element

Cleavage and polyadenylation factor 5 (Cpsf5) and 160 (Cpsf160) enable transcription termination and mRNA processing. Both factors were associated with the oocyte element. Upstream of N-ras (*Unr*), characterized as an lncRNA binding RNA chaperone (Patalano *et al*. 2009; Militti *et al*. 2014), is involved in the posttranscriptional regulation of gene expression (Kakumani *et al*. 2020), including roles in translation (Abaza *et al*. 2006; Duncan *et al*. 2006) and dosage compensation (Patalano *et al*. 2009; Militti *et al*. 2014). *Unr* binds to the IRES sequence within the 5’-end of *c-MYC* mRNA and mediates its cap-independent translation (Mitchell *et al*. 2001). Additionally, MYC and Max suppress *Unr* expression in a developmental stage-dependent manner (Anderson and Catnaigh 2015). These observations suggest that Unr might play a role in the translational regulation of *Drosophila Myc*.

SBDS ribosome maturation factor (*Sbds*) is a nucleolar protein that enables cytosolic ribosome assembly, translation, protein biogenesis, and cell growth (Takai *et al*. 2020). Since MYC proteins play an essential role in regulating genes important for ribosome assembly, biogenesis, and translation during cell growth and proliferation (Gomez-Roman *et al*. 2003; Schmidt 2004; van Riggelen *et al*. 2010; Piazzi *et al*. 2019), SBDS might potentially regulate *Myc* expression.

The eukaryotic translation initiation factor 3 subunit j (*eIF3j*) is a conserved component of the translation initiation complex (Andersen and Leevers 2007). The larger translation initiation factor 3 (eIF3) complex, which includes eIF3j, is the largest subunit in mammalian translation initiation complexes and plays an essential role in the formation and initiation of translation in highly proliferative cells (Asano *et al*. 1997).

Nucleostemin 3 (*Ns3*), an evolutionarily conserved body size regulator with GTPase activity (Hartl *et al*. 2013), functions in the export of the 60S ribosomal subunit from the nucleus and serves as a global growth controller (Hartl *et al*. 2013). Given that the regulation of *Myc* mRNA translation rate and stability are key factors controlling *Myc* activity during development and adult life (Samuels *et al*. 2020; Zacarias-Fluck *et al*. 2024), Ns3’s role in growth control could be linked to *Myc* regulation.

Effete (*eff, UbcD1*) is an E2 ubiquitin-conjugating enzyme (Ryoo *et al*. 2002; Herman-Bachinsky *et al*. 2007; Asaoka *et al*. 2016) involved in multiple processes. These include the ubiquitination and degradation of mitotic cyclins to regulate female germline stem cell maintenance (Chen *et al*. 2009) and the ubiquitination and proteolysis of a telomeric polypeptide during mitosis and male meiosis to maintain chromosome structure (Cenci *et al*. 1997). Effete might therefore be involved in Myc degradation during early cell division cycles.

### XVII. Posttranscriptional regulation of *Myc*

A considerable number of the identified proteins belong to the factors that participate in RNA transport, ribosome biogenesis, translation, mRNA processing, posttranscriptional silencing, and protein metabolism. Examples of these factors as potential regulators of *Myc* expression will be described here (Figure 16; see also supporting information, Table 3 in File S3).

**Figure 16.**
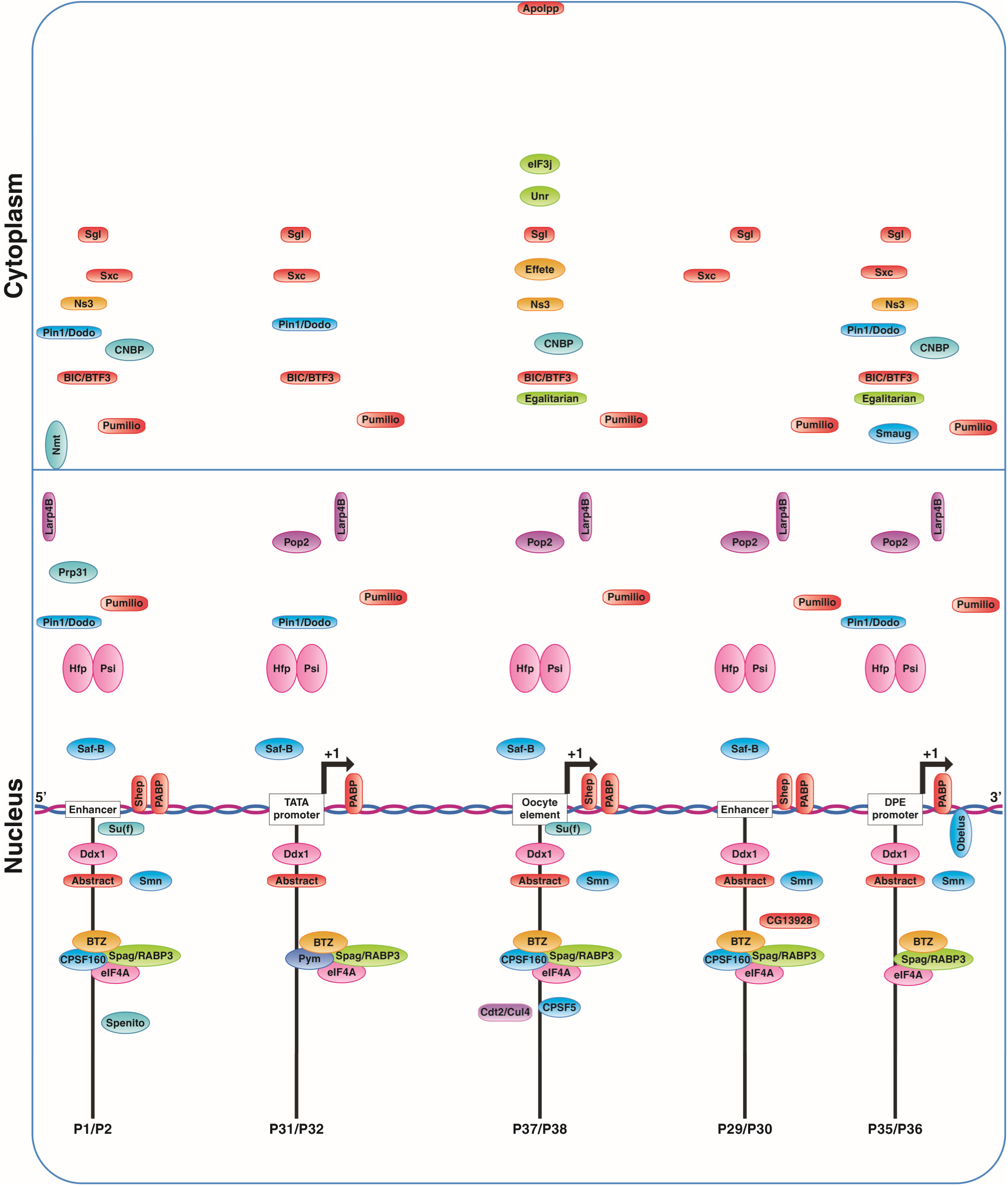
Examples of factors that might act as posttranscriptional regulators of the *Drosophila Myc* gene. We identified Upstream of N-ras (**Unr**), an RNA chaperone with a role in *Myc* mRNA translation, associated exclusively with the oocyte element P37/P38. The conserved small CCHC-type Zinc Finger protein **CNBP**, a regulator of Myc protein synthesis, was associated with the Myc-CRMs P1/P2, P37/P38, and P35/P36. Core components of the Exon Junction Complex (EJC) associated with Myc-CRMs included Barentsz (**BTZ**), eukaryotic initiation factor 4A (**eIF4A**) (an essential RNA helicase and cap-binding translation initiation factor), Dead-box-1 (**Ddx1**), Partner of Y14 and Mago (**Pym**), and Spaghetti (**Spag, RABP3**) (identified in association with the pol II transcription regulator complex and RNP protein folding complex). Among the cleavage and polyadenylation factors involved in nascent mRNA 3’-end formation, **CPSF160** was identified in association with the P1/P2, P29/P30, and P37/P38 targets, while **CPSF5** was associated exclusively with the oocyte element (P37/P38) initiation site. Poly(A) binding protein **PABP** was associated with all tested targets. Both Half pint (**Hfp**) and P-element somatic inhibitor (Psi), which have roles in transcription and RNA processing, and **Larp4B**, an RNA binding protein and translation inhibitor of *Myc* mRNA, were associated with all targets. For more information about the association of the illustrated factors with the Myc-CRMs and their abundance ratios, see supplementary materials (Table 3 in File S3, Table 7 in File S7, and Table S8).

#### Several enzymes identified in this study might regulate different aspects of Myc posttranscriptionally

O-GlcNAc transferase ‘Super sex combs’ Ogt/Sxc (*Ogt/Sxc*) adds acetylglucosamine moieties to cytoplasmic and nuclear proteins (Gambetta *et al*. 2009), including members of the Polycomb group, thereby playing a key role in regulating epigenetic gene silencing and developmental gene expression (Akan *et al*. 2016). Notably, MYC is a direct target of Ogt/Sxc, and O-GlcNAcylation stabilizes the MYC protein in humans during development and disease (Itkonen *et al*. 2013; Itkonen *et al*. 2019; Luanpitpong *et al*. 2021).

N-myristoyl transferase (*Nmt*) is an important protein contributing to the myristoylation of developmental proteins (Ntwasa *et al*. 1997) during processes such as cell shape changes and the completion of dorsal closure (Harden 2002). Notably, the MYC binding partner MAX is myristoylated posttranslationally prior to caspase cleavage during apoptosis, differentiation, and development (Martin *et al*. 2011), a modification that results in the downregulation of MYC.

Peptidylprolyl isomerase Pin1 (*Pin1*), the mammalian homolog of *Drosophila* Dodo (*dod*), facilitates rhomboid gene expression via degradation of chorion factor 2 (CF2) and induces dorsoventral patterning of the follicular epithelial cell layer within the egg chamber (Hsu *et al*. 2001). MYC is a known target of Pin1, which promotes its ubiquitination and proteasomal degradation, thereby tightly controlling MYC protein stability during cell cycle transitions; loss of this control contributes to oncogenesis in humans (Yeh *et al*. 2004; Yeh *et al*. 2006; Helander *et al*. 2015). Additionally, Pin1 directly associates with MYC and, through recruitment of p300 and indirect involvement of ABL protein, can acetylate the MYC protein, potentially facilitating its transcriptional activity (Sanchez-Arevalo Lobo *et al*. 2013).

Spaghetti/RNA pol II-associated protein 3 (*Spag/RABP3*) is a cochaperone containing a TPR domain that can associate with HSP70 and HSP90 (GO Reference Genome Project 2011). It mediates protein-protein interactions (Das *et al*. 1998a), leading to the formation of multiprotein complexes (D’Andrea and Regan 2003), including scaffolding RNA pol II and connecting it to regulatory protein complexes essential for pol II stabilization and transcriptional regulation (Rodriguez and Llorca 2020). In complex with ribonucleoproteins, Spag/RABP3 can also mediate protein folding immediately upon translation completion (Boulon *et al*. 2008; Eckert *et al*. 2010). Based on its role in transcriptional regulation, Spag/RABP3 might act as a potential regulator of *Myc*.

Large proline-rich protein CG7546 (*CG7546*, *clone 2.45*), part of the BAT3 complex, is the homolog of human BCL2-associated athanogene 6 (BAG6) (Broderick *et al*. 2010) (BAG cochaperone 6). BAG6 functions in the ubiquitin-dependent ERAD pathway, targeting proteins for degradation by the cytoplasmic proteasome (GO Reference Genome Project 2011), and is involved in regulating the DNA damage response and apoptosis (Krenciute *et al*. 2013). Given that MYC plays an important role in inducing cell death (Thompson 1998), BAG6 might participate in regulating *Myc* expression through protein degradation processes.

EGF-domain serine glucosyl-/xylol transferase Rumi (*rumi*) functions as a Notch modulator and an ER enzyme that contributes to O-linked protein modifications (Acar *et al*. 2008; Takeuchi *et al*. 2011; Sethi *et al*. 2012; Yu *et al*. 2016). Myc and Notch signaling interact during the induction of cell fate and lineage commitment, most probably in the G1 phase of the cell cycle, to produce progenitors for differentiation into various tissue types (Singh and Dalton 2009). The glucosylation of EGF repeats by Rumi is essential for the correct folding and cleavage of Notch, enabling the activation of its target genes (Yu *et al*. 2016). Through its modulation of Notch signaling, Rumi may indirectly influence Myc regulation.

Purity of essence Poe (*poe*) exhibits both calmodulin-binding and E3 ubiquitin ligase activity (Xu *et al*. 1998; Gene Ontology Curators 2002) and plays roles in glial cell growth (Yager *et al*. 2001) and sperm individualization (Fabrizio *et al*. 1998; Arama *et al*. 2003). Myc also plays key roles during the *Drosophila* sperm maturation process and in glial cell growth and differentiation (Hempel *et al*. 2006; Reddy and Irvine 2011; Ota and Kobayashi 2020; Shohayeb *et al*. 2020). Therefore, Poe might be involved in regulating *Myc* expression via degradation of the Myc protein during glial cell differentiation and sperm maturation.

#### The following factors might impact Myc mRNA processing and translation

Bicaudal (*bic/Btf3*) is the β subunit of the nascent polypeptide-associated complex (NAC), whose α and β subunits enable the activity of the ribosomal translational machinery (Markesich *et al*. 2000; Braat *et al*. 2004; Kogan *et al*. 2017; Kogan *et al*. 2022). Besides its chaperone activity, the NAC complex might function as a coactivator during developmental transcription and cell differentiation (Rospert *et al*. 2002; Liu *et al*. 2010). Quantitative proteomics approaches have revealed the coexpression of BTF3 with MYCN, *N-myc* downstream regulated gene 1 (NDRG1), and the *MYC* downstream target Ornithine Decarboxylase 1 (ODC1) in epithelial cancer cells (Bello-Fernandez *et al*. 1993; Tu *et al*. 2007; Hogarty *et al*. 2008; Symes *et al*. 2013), suggesting complex interactions between the MYC/Max network and other factors, including BTF3.

The conserved small CCHC-type zinc finger protein CNBP (*CNBP*) binds to ssDNA/RNA and plays a key role during development and disease (Antonucci *et al*. 2014). CNBP binds G-rich regions in mRNAs and stabilizes their structure by inhibiting G-quadruplex formation (InterPro Project Members 2004; GO Reference Genome Project 2011). CNBP promotes *Myc* protein synthesis through IRES-dependent translation of *Myc* mRNA, and the absence of CNBP causes embryonic lethality (Antonucci *et al*. 2014). Furthermore, CNBP is involved in *Drosophila* wing disc morphogenesis and the control of wing size by modulating Myc protein levels (Antonucci *et al*. 2014). Identified as a *Myc* regulator, CNBP was found associated with the upstream enhancer P1/P2, the proximal P37/P38 oocyte element, and the DPE promoter in this study.

The F-box-like/WD40 repeat-containing protein Ebi (*ebi*), the *Drosophila* homolog of mammalian *TBL1*, is a nuclear conserved repressor identified in association with transcription repressor complexes (Tsuda *et al*. 2006) and histone deacetylase complexes (GO Reference Genome Project 2011). Ebi integrates multiple developmental signaling pathways, including JNK (Lim *et al*. 2012), EGFR (Dong *et al*. 1999), Notch (Tsuda *et al*. 2002; Marygold *et al*. 2011; Nguyen *et al*. 2016), and Wingless (Matsuzawa and Reed 2001), to regulate cell cycle switches, gene expression, and the induction of differentiation and patterning. Ebi might therefore have a direct and/or indirect role in controlling *Myc* expression during development.

Pre-mRNA processing factor 31 (*Prp31*) (Figure 16; Table 3 in File S3) is a splicing factor involved in pre-mRNA processing/splicing (Mount and Salz 2000; Herold *et al*. 2009) and photoreceptor development (Ray *et al*. 2010). Low levels of the protein or loss of the Prp31 gene lead to embryonic lethality in mice (Bujakowska *et al*. 2009). Given that MYC plays a crucial role in controlling the expression of splicing machinery factors to maintain proper pre-mRNA splicing during development and disease (Koh *et al*. 2015; Abou Faycal *et al*. 2016; Phillips *et al*. 2020; Jablonowski *et al*. 2023), Prp31 might potentially be involved in the regulation of *Myc*.

Spenito (*nito*) (Figure 16), an RNA-binding protein (Lasko 2000), acts downstream of the Wnt signaling pathway and is required during larval development (Chang *et al*. 2008). In association with the WMM complex, Spenito mediates the m6A methylation of mRNAs, increasing the efficiency of alternative mRNA splicing for targets such as Sex-lethal (*Sxl*) during sex determination and dosage compensation (Yan and Perrimon 2015). Notably, *Myc*, *Sxl*, and transformer (*tra*) genes together control male and female body size dimorphism (Mathews *et al*. 2017). This suggests Spenito might have a role in regulating *Myc* expression, potentially during sex determination among other processes.

Suppressor of forked Su(f) was enriched at the P1/P2 and P37/P38 enhancer clusters. Su(f) enables mRNA processing (InterPro Project Members 2004; GO Reference Genome Project 2011), and high concentrations are found during mitosis in proliferating cells (Mitchelson *et al*. 1993). Su(f) negatively regulates the gypsy transposon and inhibits gypsy mRNA accumulation (Parkhurst and Corces 1986). Notably, the retrotransposon gypsy is inserted in the proximal 5’-end region of the *Myc* gene, within the first intron (Gallant *et al*. 1996) near the P37/P38 enhancer cluster; this location potentially underscores the significance of the observed association between Su(f) and the P37/P38 enhancer.

La-related protein 4B (*Larp4B*) is an RNA binding protein involved in the downregulation of cell growth via posttranscriptional inhibition of *Myc* mRNA translation. Overexpression of Larp4B results in reduced cell size and small organ size (Funakoshi *et al*. 2018).

Pumilio (*pum*), an RNA binding protein, is involved in modulating RNA stability, translation repression (Dean *et al*. 2002; Kim *et al*. 2012), and RNA degradation. Target mRNAs deadenylated by Pop2 are recognized by repression domains (RDs) of Pumilio for destruction during stem cell differentiation, neurogenesis, and embryogenesis (Arvola *et al*. 2020). Pumilio plays roles in regulating Hunchback mRNA in early embryos, germline stem cell proliferation and migration (Asaoka-Taguchi *et al*. 1999; Wickens *et al*. 2002), and fate specification in sensory organ precursor (SOP) cells via EGFR signaling (Kim *et al*. 2012). By integrating inputs from multiple signaling pathways, Pumilio and Pop2 might be involved in the negative regulation of *Myc* mRNA levels during processes such as bristle and wing vein development.

Dead-box-1 (*Ddx1*), an RNA helicase, plays roles in RNA trafficking, nucleic acid duplex unwinding (Gene Ontology Curators 2002), translation, and protein synthesis (Rafti *et al*. 1996). In the early embryo, Ddx1 positively regulates cell proliferation and controls body size and germ cell development (Rafti *et al*. 1996; Germain *et al*. 2015). The human homolog of *Ddx1* has been shown to be amplified in a series of MYCN-amplified tumors, such as retinoblastoma (Godbout and Squire 1993; Manohar *et al*. 1995; Squire *et al*. 1995; Godbout *et al*. 1998). Furthermore, in pediatric neuroblastoma, the MYC-binding partner Max showed strong interaction with the Ddx1 protein (Jin *et al*. 2021), suggesting that Ddx1 might be indirectly involved in *Myc* regulation through its interaction with Max.

Abstrakt (*abs*), a DEAD/H-Box RNA helicase, is involved in RNA splicing (Herold *et al*. 2009; GO Reference Genome Project 2011) and functions as a Notch regulator in certain sibling progenitor cells by controlling mRNA levels of Inscuteable (*insc*) during neurogenesis (Irion *et al*. 2004). Abstrakt function is also required for the regulation of epithelial cell polarity in the oocyte and during embryogenesis (Irion and Leptin 1999). Notably, DEAD Box helicases have been shown to be direct targets of MYC/Max heterodimers (Grandori *et al*. 1996), potentially linking Abstrakt to *Myc* regulatory networks.

*Drosophila* Barentsz (*btz*), whose vertebrate homologs include *CASC3* and *MLN51*, is a component of the Exon Junction Complex (EJC) pathway. The EJC marks intron removal sites on spliced mRNAs, a process essential for subsequent mRNA translation, localization, and decay of nonsense-mediated mRNAs. Core EJC components, including MLN15/Barentsz, eIF4A3, Ddx1, and Mago/Y14 (Le Hir and Andersen 2008; Chazal et al. 2013), are required during transcription termination and RNA processing (Ishigaki et al. 2014). High levels of EJC are present in the nuclei of cells during late oogenesis, and the EJC associates with pol II elongation and termination complexes in cycling and differentiating cells (Kiselev et al. 2017). Additionally, the EJC component Y14 was shown to colocalize with Myc in the nucleus during pre-/vitellogenic stages of oocytes (Kiselev et al. 2017). Although Mago was not significantly enriched in the original list (Table 3 in File S3), other key EJC subunits— Barentsz, Ddx1, and partner of Y14 and Mago (Pym) (Ghosh et al. 2014)—were enriched on the Myc-CRMs in this study. This suggests that the EJC might play an important role in terminating *Myc* transcription and processing *Myc* mRNAs.

Smaug (*smg*) is the founding member of the SMAUG family of sequence-specific RNA-binding proteins. Smaug functions as a translation repressor (Dean *et al*. 2002; Kim and Bowie 2003) during the maternal-to-zygotic transition in early embryos and induces the degradation of hundreds of maternal mRNAs by recruiting the CCR4-NOT deadenylase complex (Jeske *et al*. 2006; Zaessinger *et al*. 2006; Chartier *et al*. 2015). This activity thereby controls embryonic patterning (Smibert *et al*. 1999), the activation of zygotic transcription, and early embryonic cell cycles (Liang *et al*. 2008). These functions might indicate a role for Smaug in the regulation of *Myc* as an immediate early gene.

Obelus (*obe*), which belongs to the Ski2 family of DNA helicases (Vichas *et al*. 2015), is required for adherence junction arrangement and the alternative splicing of Crumbs (*crb*) mRNA, a transmembrane protein responsible for apico-basal cell polarity during embryogenesis and oogenesis (Vichas *et al*. 2015). Myc, a master regulator of growth and proliferation, must be tightly controlled to maintain the balance between cell polarity and proliferation rates and avoid malignant transformation (Grifoni *et al*. 2013). Here, we found the Obelus protein, a regulator of apico-basal cell polarity, associated with the DPE-containing Myc-CRM P35/P36. Obelus might therefore be a potential regulator of *Myc* mRNA and protein levels during early development.

How/Struthio (*how*, *struthio*), known as ‘held out wings’, is a conserved KH domain RNA binding protein (Baehrecke 1997) with a role in the alternative splicing of mRNAs (Volohonsky *et al*. 2007; GO Reference Genome Project 2011). Its isoform how(L) destabilizes RNA, whereas How(S) induces RNA stabilization (Volohonsky *et al*. 2007) during embryonic cell differentiation and morphogenesis (Walsh and Brown 1998). How/Struthio functions downstream of various signaling cascades, including EGFR (Martin-Bermudo 2000; Edenfeld *et al*. 2006), MAPK (Toledano-Katchalski *et al*. 2007), ERK (Nir *et al*. 2012), DPP/BMP (Israeli *et al*. 2007), Hedgehog (Lobbardi *et al*. 2011), Wnt-TCF (Riese *et al*. 1997; Marsden and DeSimone 2001), Insulin-like receptor, Ras, and Akt (Lasko 2003), to control cell cycle progression and induce differentiation. The mouse homolog of How/Struthio, Quaking, is a tumor suppressor RNA binding protein that contributes to controlling cell cycle progression and promoting differentiation pathways (Biedermann *et al*. 2010). These functions suggest How/Struthio might control *Myc* expression during cell cycle progression and the induction of fate specification. Scaffold attachment factor B (Saf-B) demonstrates versatility as both an RNA and DNA binding protein involved in regulating alternative mRNA splicing (PARK *et al*. 2004) and pol II transcription (GO REFERENCE GENOME PROJECT 2011). Structurally, *Drosophila* SAF-B links the nuclear matrix, chromosomes, and the transcription regulator complex, playing a role in maintaining nucleosome-chromosome architecture and overall nucleus organization (ALFONSO-PARRA AND MAGGERT 2010). Contextually relevant is the finding that the human *MYC* gene contains a nuclear matrix attachment region (MAR) within its 3’-UTR, bound by proteins like the transcription factor Sox2 (LEI *et al*. 2005). MARs generally function as boundary elements situated between genes, serving as protective shields during DNA replication and transcription (BODE *et al*. 1996; KOHWI-SHIGEMATSU *et al*. 1997; BODE *et al*. 2003). Considering its role in nuclear architecture and MAR association, Saf-B might provide an essential structural hub for ensuring nuclear integrity during *Myc* transcriptional regulation.

Highlighting the close connection between Myc and protein synthesis, our analysis revealed that numerous components of the ribosome and mRNA translation machinery associate with the Myc-CRMs. These included a wide array of proteins from both the large (60S) ribosomal subunit (RpL5, RpL8, RpL9, RpL13, RpL13A, RpL14, RpL19, RpL27A, RpL31, RpL32, RpL36, RpL38) and the small (40S) ribosomal subunit (RpS3, RpS3A, RpS6, RpS5a6, RpS10b, RpS13, RpS24, RpS26, RpS28b, RpS29), in addition to the mitochondrial ribosomal protein mRpL16. Furthermore, a significant number of key eukaryotic mRNA translation factors were found associated with Myc-CRMs, encompassing factors involved in initiation (eIF2α, eIF3B, eIF3C, eIF3J, eIF3I, eIF4A, eIF4B, eIF4E1, eIF4H1), elongation (eEF1α1), and termination (eRF1), alongside mitochondrial initiation (mIF2) and elongation (mEFTu1) factors.

Several of the aforementioned ribosomal proteins and the translation initiation factor eIF2α, as part of the eIF-2 complex, have been classified as ‘Minute’ proteins because heterozygous animals grow slowly and exhibit small body size and short bristles (Marygold *et al*. 2007). Initiation factor eIF3I, a component of the eIF-3 complex, stimulates the binding of mRNA to the 40S ribosome (Lasko 2000; GO Reference Genome Project 2011) to specifically target and initiate translation of a subset of mRNAs involved in cell proliferation (Asano *et al*. 1997). Translation initiation factor eIF4B plays a role in cap-dependent translation, cell survival, and proliferation (Hernandez *et al*. 2004). Translation initiation factor eIF4E1 is a component of the eIF4F cap-binding complex and plays a crucial role in the cap-dependent translation of mRNA (Lavoie *et al*. 1996).

Ribosomal genes are known *Myc* target genes (Orian *et al*. 2003; Orian *et al*. 2005). Notably, a high percentage of the Rps and translation factors identified here were associated with the tissue-specific enhancer cluster P1/P2 and are mostly involved in cell growth and tissue patterning. For instance, the X-linked mRpL16 (*mRpL16*) (Marygold *et al*. 2007), enriched exclusively on the P1/P2 enhancer, is expressed in the adult head, embryonic/larval midgut, larval muscle system, larval visceral muscle cells, and spermatozoon (Hamada *et al*. 2005; Deng and Meller 2006; Worringer and Panning 2007). Translation release factor eRF1, associated only with P1/P2, induces the release of the nascent peptide chain in response to termination codons UAA, UAG, and UGA. Mutations in the eRF1 gene result in the readthrough of stop codons, causing maternal-effect nonsense suppression and lethality in larval stages (Chao *et al*. 2003). We conclude that components of the ribosome machinery and translation protein complexes might regulate *Myc* mRNA and protein levels during cell and tissue growth, mainly involving the P1/P2 enhancer cluster.

## Discussion

This study used a newly developed, highly sensitive and selective Solid Surface Magnetic Enrichment Protocol (SSMEP) to purify proteins from *Drosophila* embryonic nuclear extracts and subsequently identified *Myc* regulators by mass spectrometry analysis. The goal of this project was to discover potential trans-regulators responsible for *Myc* regulation at the levels of transcription, post-transcriptional control, and post-translational protein stability and decay. We justify the chosen target sequences—P1/P2, P29/P30, P31/P32, P35/P36, and P37/P38—based on the prior identification of conserved Myc-CRMs through *in silico* phylogenetic alignment using *cis*-Decoder (Kharazmi *et al*. 2012) and by reporter expression patterns observed under the control of these *Myc cis*-elements (Figures 1 and 2; Table S1.3). The datasets obtained show that mitogenic stimuli trigger transcription through cascades of multiprotein complexes reaching the nucleus, leading to the assembly of pre-initiation complexes on *Myc* promoters accompanied by the binding of trans-acting factors to different sets of enhancers within the gene body.

To validate our experimental approach with the identified factors associated with Myc-CRMs, we used a Wnt *cis*-Regulatory Module (W-CRM) as a positive control that has been previously designed and tested in *Drosophila* and *in vitro* using expressed and purified proteins (Archbold *et al*. 2014). In our experiment, the W-CRM sequence (Figure 3) was tested with crude nuclear embryonic extracts, leading to the identification of potential *wingless/wg* regulators (Figure 11; Table 3 in File S3 and Table S8). In addition to components of the Wnt/Wg signaling pathway, factors from the Notch, Hedgehog, Dpp, EGFR, and MAPK pathways were also associated with the W-CRM, among others. Developmental collaboration between Wnt/Wg and other pathways is well-known; for instance, Wingless and Notch cooperate in wing pattern formation (Couso *et al*. 1995). We believe the tested W-CRM containing a TCF/LEF site can serve as a suitable positive control in similar studies.

### Developmental signals and cell cycle factors control Myc expression

Prominent among the identified *Myc* regulators are cofactors controlling cell cycle phases (Figure 13), highlighting the correlation between cell growth, proliferation, and *Myc* expression levels (Bretones *et al*. 2015). Following cell cycle factors, many of the identified proteins are targets of the Wingless, Notch, and EGFR signaling pathways, as well as components of the Dpp, Hedgehog, ecdysone, and miRNA pathways. The identification of known factors among our protein list corroborates previous findings on the control of *Myc* expression.

Advances in *Myc* regulation indicate that a variety of cofactors act as *Myc* regulators, functioning both transcriptionally and posttranscriptionally via positive and/or negative feedback loops. For example, the transcriptional coactivator Yorkie (*yki*), the *Drosophila* homolog of Yap in the Hippo pathway, stimulates *Myc* transcription, while high levels of Myc protein, in turn, repress *yki* (Neto-Silva *et al*. 2010).

The complexity of *MYC* regulation is multilayered, involving multiple initiation sites and differential regulation to orchestrate its early ubiquitous expression and later patterned expression during development (Liu *et al*. 2006). Analyses of the most relevant identified trans-acting factors reveal that the *Myc* gene is at the nexus of diverse signal transducers, allowing it to deliver messages and trigger appropriate promoter responses according to developmental stage and the physiological needs of cell types. Cdk5, Scribbler/Brakeless/Thickveins, and SAP18/HDAC1, found associated with Myc-CRMs in this study, exemplify the tight regulation of *Myc* in conjunction with cell cycle factors and cell signaling components. Cdk5 controls axonogenesis in *Drosophila* embryos through interaction with Rac GTPase and, by partnering with P35, induces the differentiation of post-mitotic neurons (Connell-Crowley *et al*. 2000). Given the induction of *MYC* by Cdk5 in tumor cells via inhibition of the *MYC* suppressor Bin1 (Ito *et al*. 2014; Zhang *et al*. 2019) and the regulation of CDKs by MYC (Bretones *et al*. 2015), we postulate that Cdk5 might regulate *Myc* expression during the onset of tissue patterning in *Drosophila*.

The transcriptional coregulator Scribbler, in association with the corepressor Atrophin/Grunge (an interactor of the histone deacetylase complex), represses transcription during early embryonic segmentation and later during axon projection from photoreceptor cells to the medulla (Haecker *et al*. 2007). Through modulation of Notch, Wingless, and Hedgehog signaling target gene expression, Scribbler determines many aspects of wing disc patterning (Dworkin and Gibson 2006). Scribbler requires binding to *cis*-elements of Kruppel to repress transcription (Haecker *et al*. 2007).

The chromatin repressive complex SAP18/HDAC1 potentially reflects the complexity of the signaling network relevant to *Myc* regulation. SAP18/HDAC1 functions in various settings during body patterning and, like Scribbler, requires tethering to Kruppel CRMs to recruit the enhancer-specific Sin3A/HDAC1 repressor complex (Matyash *et al*. 2009). This recruitment represses the transcription of certain genes, such as Bicoid during embryonic germband extension (Zhu *et al*. 2001). This suggests that Scribbler, Grunge, SAP18/HDAC1, and the signaling pathways linked to these proteins might form subsets of activating and repressing complexes to pattern *Myc* expression during cell and tissue growth.

To sum up, the transcription factor MYC, a master regulator of transcription, is under the tight control of cell signaling and cell cycle control programs; deregulation of MYC drives cells toward malignant transformation and metabolic adversities.

### The distal enhancer cluster P1/P2 controls Myc transcription during tissue patterning and DNA replication

A large subset of the P1/P2-associated proteins constitute components of conserved signaling pathways with activities in the induction of cell differentiation, morphology changes, and patterning during larval and pupal development. The P1/P2 enhancer cluster sequence 5’-GAAGCTTT**TTGTTG**T**TTTTGCAGCTGC**C**GTTT**TCGCCCTT-3’ (ChrX:3,369,776..3,369,815) contains an E-box (indicated by gray) within a sequence of conserved bases (bold letters). Correspondingly, Net (*Net, CG11450*), the E-box binding (GO Reference Genome Project 2011) basic helix-loop-helix (bHLH) transcription repressor, was pulled down by the P1/P2 sequence. Net acts as an antagonist of EGFR signaling and is part of the Wnt enhanceosome, with roles in proliferation, cell fate induction, and wing vein formation (Brentrup *et al*. 2000; Peyrefitte *et al*. 2001).

Factors found uniquely associated with P1/P2 encompassed the tissue-specific T-box transcription factor DOC3 (a cofactor for TGF-beta, Dpp and Wg signaling) (Reim and Frasch 2005; Hatton-Ellis *et al*. 2007; Lu *et al*. 2022), Spenito (Wg cofactor) (Chang *et al*. 2008), and N-myristoyl transferase Nmt (TGF-β/Dpp and Wg cofactor) (Harden 2002). Interestingly, MED24, a sensor for developmental signals essential for pupal growth (Wang *et al*. 2010), was also exclusively associated with P1/P2. One scenario could be that Wingless signaling forms a transcription inhibition complex with the transcription repressor Net and recruits further repressive factors to the E-box, then mediates this information to the pol II complex via MED24 to downregulate *Myc* during wing vein specification. A parallel activating complex might be formed involving the Wingless/Wg pathway, perhaps through the phosphorylation of ‘Enhancer of Rudimentary’ (ERH) (Gelsthorpe *et al*. 2006) (enriched exclusively on P1/P2 and the W-CRM) by CKII at either target site, which could counteract the repressive behavior of the Net complex and upregulate *Myc*. Similar regulatory switches are conceivable involving the binding of the trans-acting factor DOC3 to the P1/P2 *cis*-element for *Myc* control during cardiogenesis, neuronal development, or similar processes. We hypothesize that complex regulatory programs initiating from the distal enhancer P1/P2 govern the differential spatio-temporal expression of *Myc* during larval and pupal development.

The association of Topoisomerase II (Top2) and RecQ4 helicase with P1/P2 suggests a role for this *cis*-element in the removal of torsional stress generated by DNA replication and transcription, and in the maintenance of genome integrity. The DNA 3’-5’ RecQ4 helicase plays a pivotal role during S phase in maintaining genome stability during DNA strand separation and exchange. This raises the possibility that RecQ4 might control *Myc* levels, mainly during S phase and the G1/S checkpoint, to ensure healthy cell cycle progression.

Taken together, P1/P2 appears to function as an enhancer with dual roles: transmitting patterning signals to *Myc* promoters to establish differential *Myc* expression during tissue growth, and relaxing DNA topology through the removal of supercoils during transcription and replication.

### The *Myc* TATA-box containing P1 promoter might represent a major transcriptional bursting site

The Myc-CRM P31/P32 sequence, 5’-GA**GCGCGGC**AGTCTGGTACGATAG**AAATTTTATTTAA**GCCACAG-3’ (ChrX:3,373,105..3,373,148), derived from the P1 promoter region, retains the GC-box and the TATA-box (indicated by gray) (Kharazmi *et al*. 2012). Most of the proteins found associated with this target belong to the RNA pol II complex or are involved in different steps of transcription, chromatin remodeling, cell cycle regulation, and DNA replication/repair (Figure 14 and Figure 15; Table 3 in File S3).

### Basal transcription machinery at the Myc TATA promoter

We identified general transcription factors and components of RNA pol II indicative of the assembly of a functional pre-initiation complex (PIC) on TATA promoter. Literature has shown that purified TFIID incubated with the adenovirus major late promoter can form a stable PIC (Van Dyke *et al*. 1989). Formation of the PIC on a core promoter *in vivo* requires TFIID, TFIIA, TFIIB, TFIIE, TFIIF, TFIIH, and RNA pol II (Thomas and Chiang 2006). The binding of TBP/TFIID to the promoter nucleates PIC assembly (Smale and Kadonaga 2003), followed by the recruitment of TFIIA for stabilization of TBP/TFIID binding, and promoter contact with TFIIB, which recruits RNA pol II to the complex (Thomas and Chiang 2006).

The TFIID complex subunits TAF5, TAF6, TAF12, and dFUS/Cabeza were associated with the TATA promoter; TAF12 recruits TAF4, and TAF12 can also recruit TAF9 to the TBP-TFIID complex *in vivo* (Yokomori *et al*. 1993). A stable core TFIID complex consists of TAF4, TAF5, TAF6, TAF9, and TAF12 (Wright *et al*. 2006). Although TFIIA-S (*Drosophila* isoform of TFIIA) (Aoyagi and Wassarman 2000) appeared on the original list, it was not significantly enriched. Most of the RNA pol II subunits required for transcription were associated with the TATA promoter. Among the identified pol II subunits were RPB1, the largest subunit of RNA pol II (Greenleaf 1983; Brickey and Greenleaf 1995; InterPro Project Members 2004); RPB2, the second largest component of the pol II core complex (Falkenburg *et al*. 1987; Aoyagi and Wassarman 2000); and the evolutionarily conserved RPB7. RPB7 is essential in *Drosophila* and can recruit and heterodimerize with RPB4, both of which are vital for transcriptional and posttranscriptional events (Pankotai *et al*. 2010).

The activity of the TATA-bound PIC might be sufficient for a basal level of *Myc* transcription. However, achieving cell-type or tissue-specific *Myc* patterning likely requires the activity of additional GTFs and the Mediator complex, working independently or in collaboration, to transmit regulatory cues between upstream activators and/or repressors and the initiation complex. Further GTFs associated with the Myc-CRMs included TFIIEα, CycH-TFIIH, Ssl1-TFIIH, and topoisomerase II (Top2). TFIIE (α and β subunits) constitutes an important component of the initiation complex that interacts with bound pol II and contributes to promoter clearance. TFIIE can function independently or in collaboration with other GTFs and TFIIH. TFIIH could potentially contribute to many aspects of initiation and elongation via its helicase activity (involving p44/Ssl1 and the XPD subunit) for promoter opening, transcription-coupled nucleotide excision repair, phosphorylation of the pol II CTD, and E3 ubiquitin ligase activity. Top2 can instantly neutralize torsional stress upstream and downstream of the transcribing machinery. The dFUS/Cabeza subunit of TFIID may accommodate the pol II complex during initiation, termination, and mRNA processing.

The associated Mediator subunits consisted of MED4, MED11, MED13/Skuld, MED17, and MED25, which might suggest the presence of a Mediator complex containing both large and accessory subcomplexes capable of bridging enhancers to GTFs. The Mediator complex could transmit signaling stimuli from activators and repressors to the TATA promoter and initiation complex, enabling precisely patterned expression of *Myc*.

Finally, the association of Trf2 with the TATA promoter might offer an alternative mechanism for *Myc* transcriptional regulation in addition to TBP-dependent transcription. We hypothesize that both TAF-TBP and TAF-Trf2 initiation pathways might be possible from the TATA promoter for expressing *Myc* throughout development.

### The Myc DPE promoter and the downstream enhancer cluster P29/P30 participate in the formation of the basal transcription machinery on the DPE promoter

Analysis of the *Drosophila* promoter database indicates that approximately 40% of *Drosophila* promoters contain a DPE element (Kutach and Kadonaga 2000; Thomas and Chiang 2006). In previous work, we located an initiator (Inr) and a DPE element, the P2 promoter, within the *Myc* downstream intronic sequences (Kharazmi *et al*. 2012) (Figure 2A). Here, the factors associated with the DPE-containing target P35/P36 and the adjacent enhancer cluster P29/P30 show that these two elements constitute a functional initiation site within the downstream intronic sequences, capable of cooperatively initiating robust transcription from this site (Figure 14 and Figure 15). TAF6 and TFIIEα, found associated with the enhancer P29/P30, might complement the factors on the DPE to enable the formation of a stable PIC (Figure 15). The strength and choice of transcription factors, as well as the composition of the core promoter, play crucial roles in determining the robustness and magnitude of transcription initiation (Sloutskin *et al*. 2024). The *Myc* DPE promoter responds to mitogenic stimuli transduced by conserved developmental signaling pathways and to abiotic stress. The outcome of each context-dependent response to an inducer contributes to changes in *Myc* expression levels, affecting cell cycle progression, cell proliferation rate, and differentiation. DPE promoters, in general, have been reported to play pivotal roles during different developmental stages by translating input signals from evolutionarily conserved pathways into transcriptional regulatory programs that contribute to animal growth (Sloutskin *et al*. 2024). Precisely designed genetic and functional studies will be needed to elucidate the molecular mechanism of the regulatory network at the *Myc* DPE promoter.

### Our findings suggest the Myc enhancer cluster P37/P38 may function as a transcribing developmental super enhancer synthesizing eRNAs

Work by others has shown that microinjection of a plasmid containing a *MYC* promoter into Xenopus oocytes and embryos leads to high levels of *MYC* expression during oogenesis, followed by silencing in early fertilized embryos, and subsequent expression reactivation at the midblastula transition (MBT) (Prioleau *et al*. 1994). Although early embryos contain all the necessary GTFs for transcription complex assembly, high nuclear levels of histones hamper the DNA binding of transcription factors (Prioleau *et al*. 1994). To initiate transcription during the early stages of embryonic development, TBP-DNA binding must be established at promoter complexes to overcome histone inhibition (Prioleau *et al*. 1994).

In addition to the three previously reported P0, P1, and P2 promoters (Kharazmi *et al*. 2012), our proteomics data indicate that the enhancer cluster P37/P38 within the proximal 5’-UTR region may serve as a potential fourth initiation site at the *Myc* locus, with main activity during oogenesis and embryogenesis. Factors associated with P37/P38 primarily indicate the commencement of TAF-TFIID and/or TAF-TRF2 transcription (Figure 15; Table 3 in File S3). To supply cells with sufficient biomass during oocyte maturation and embryogenesis, transcriptional bursting from the oocyte element at the MBT transition and the start of zygotic gene expression is expected to be a crucial source.

*MYC* is thought to be transcribed into mRNAs of varying lengths, including enhancer RNAs (eRNAs). The mouse *MYC* gene generates transcripts of varying lengths throughout development (Spencer and Groudine 1991; Benassayag et al. 2005; Kubickova et al. 2023). Furthermore, in tumor cells experiencing replication stress, the production of transcripts can increase, yielding up to five shorter variants at the expense of full-length *MYC* mRNAs (Ibrahim et al. 2022).

Developmental enhancers can produce eRNAs independently of promoters (Catarino and Stark 2018; Henriques *et al*. 2018). Such eRNAs are initially short-lived molecules and can potentially be translated with or without undergoing polyadenylation (Koch *et al*. 2011). The association of key regulatory factors with the P37/P38 element—such as subunits of the pol II complex, subunits of TFIID and Mediator complexes, GTFs, chromatin remodeling and histone-modifying enzymes, TRF2/GAF, and TRL (Figure 14 and Figure 15)—strongly suggests that the oocyte element P37/P38 serves as a transcribing super enhancer, empowering *Myc* in its function as a master regulator of cell growth and proliferation and as a determinant of organ size. Proteins associated with the P37/P38 also includes other factors, such as Effete/UbcD1, eIF3j, the ribosome maturation factor SBDS, survival motor neuron Smn, and the Cdt2 protein (encoded by the *lethal-(2)-denticleless* gene), whose known functions point to roles in oocyte maturation, female germline cell maintenance, sex determination, early phases of the mitotic cell cycle, and the establishment of germ layers.

### Myc control involves a dynamic chromatin landscape and regulatory switches utilizing gene-specific factors

Competition between chromatin changes and transcription complex assembly is a key determinant in the regulation of *MYC* transcription during development (King *et al*. 1986; Vriz *et al*. 1989; Prioleau *et al*. 1994). We identified components of major chromatin assembly and histone modification complexes associated with Myc-CRMs, indicating vital roles for major remodeling complexes such as SAGA, SWI/SNF (Brahma), ATAC, NuRF, NURD, ACF, and CHRAC throughout fly development.

Certain components of GTFs, like TAF5 and TAF6, show interrelation with chromatin remodeling complexes for the modulation of cell-type-specific transcription. TAF5 and TAF6 (found associated with Myc-CRMs), subunits of TFIID, are also parts of the coactivator SAGA complex (Wang *et al*. 2020). SAGA exhibits both histone acetyltransferase (HAT) and ubiquitin protease activity (Guelman *et al*. 2006b). SAGA links TBP to transcription factors, ensures promoter opening, degrades unwanted transcription factors to initiate differentiation in certain cell types, and is enriched at the proximal 5’-ends of genes (Hansen and Tjian 1995; Wright *et al*. 2006; Weake *et al*. 2011). Furthermore, 5’-end paused SAGA may contribute to the immediate reinitiation of transcription for genes like *Myc* via TAF-TRF2/pol II and/or TAF-TFIID/pol II pause release in response to active enhancers (Weake *et al*. 2011).

In addition, the SAGA complex plays an essential role during pol II elongation events. The transcription elongation factor SPT6 is a component of the SAGA/TFTC complex and associates with both nucleosomes and actively transcribing pol II (Ardehali *et al*. 2009). SPT6 recruits other elongation factors such as SPT5 and PAF1 (SPT6 and PAF1/Antimeros were both found associated with Myc-CRMs) and contributes to chromatin remodeling and pol II elongation (Andrulis *et al*. 2002; Petruk *et al*. 2006; Hsu *et al*. 2008; Ardehali *et al*. 2009). Taken together, our data suggest that the SAGA complex might be actively involved in different steps of *Myc* transcriptional control.

SWI/SNF (Brahma), a known chromatin remodeling complex in *Drosophila* (MARENDA *et al*. 2004), might have an essential role in regulating *Myc*. For instance, the component Snr1/BAP45, found associated with Myc-CRMs, induces ATP-dependent chromatin remodeling linked to endosome-mediated signaling cascades; this activity negatively regulates pol II transcription during cell proliferation and induces differentiation (DINGWALL *et al*. 1995; KAL *et al*. 2000; MARENDA *et al*. 2003; MOHRMANN *et al*. 2004; CHALKLEY *et al*. 2008; BONNAY *et al*. 2014; BARISH *et al*. 2020; MASHTALIR *et al*. 2020). In addition, Snr1/BAP45 inhibits CycE during S phase and induces histone tail methylation during differentiation and patterning (BRUMBY *et al*. 2002; KOE *et al*. 2014). This raises the possibility that Snr1/BAP45 might directly inhibit *Myc* transcription during S phase to promote cell differentiation.

A further function of chromatin assembly complexes might involve associating with gene-specific factors that target the *Myc* promoter, or alternatively, transcription factors might themselves be parts of remodeling complexes and act as *Myc* regulators. For instance, Host cell factor (HCF), found associated with Myc-CRMs and known as a cell cycle regulator (MAHAJAN *et al*. 2003), is a component of the Ada2a complex (ATAC). Furthermore, the Ash2-Sin3A-HCF protein complex activates promoters of genes involved in growth and proliferation via histone modifications (GUELMAN *et al*. 2006a; SUGANUMA AND WORKMAN 2008; FURRER *et al*. 2010; SUGANUMA *et al*. 2012). In a similar manner, heterodimers such as HCF/MLF or MLF/DnaJ-1 might downregulate *Myc* transcription during tissue growth and organogenesis (JASPER AND BOHMANN 2002) and during hematopoiesis (BRAS *et al*. 2012; MILLER *et al*. 2017), potentially in response to signaling cascades such as Hedgehog (FOUIX *et al*. 2003).

### Myc mRNA and protein levels are under tight control during development

The vital role of Myc in controlling pol I, II, and III transcription to modulate various aspects of cell physiology, including ribosome biogenesis and DNA replication, has been evidenced by the work of others (Raha *et al*. 2010a; Mitchell *et al*. 2015). Two of the RNA polymerase subunits found associated with the Myc-CRMs, RPB5 and RPB8, exhibit polymerase activity from the promoters of pol I, pol II, and pol III. Polr2E/RPB5, which enables pol II transcription, is also involved in transcription elongation by pol I during rRNA synthesis and in tRNA synthesis by pol III (Aoyagi and Wassarman 2000; Gene Ontology Curators 2002; GO Reference Genome Project 2011). RNA pol I, II, and III subunit H, Polr2H/RPB8, enables transcription by all three polymerases (Aoyagi and Wassarman 2000; Gene Ontology Curators 2002; GO Reference Genome Project 2011).

Myc’s function in controlling target gene expression during cell growth and proliferation (Orian *et al*. 2007) might be linked to the control of *Myc* expression by its associated cofactors, which include components of RNA pol I, II, and III, as well as factors involved in ribosome biogenesis and mRNA translation. The identification of numerous ribosomal proteins and translation factors associated with the Myc-CRMs might point to the potential roles of these proteins in controlling *Myc* mRNA and protein levels during cell growth, proliferation, and differentiation. The posttranscriptional/-translational factors associated with Myc-CRMs include parts of the cytoplasmic 40S and 60S subunits of the eukaryotic 80S functional ribosomal apparatus, mitochondrial ribosomal proteins, and eukaryotic translation factors.

Point mutations in certain genes encoding ribosomal proteins (RPs) and mRNA translation factors, including RpL15, RpS5a6, and eIF2α, cause the ‘Minute’ condition due to allelic loss-of-function leading to genetic haploinsufficiency (Lambertsson 1998; Marygold *et al*. 2007). Animals with the ‘Minute’ syndrome show small body size, short and thin bristles, and reduced viability and fertility (Qian *et al*. 1987; Marygold *et al*. 2007). Ribosome biogenesis and protein synthesis are key factors sensed by cells for controlling DNA replication and the G1/S transition (Pardee 1989; Norbury and Nurse 1992). Myc is a fine-tuning regulator of DNA synthesis and ribosome biogenesis, thereby controlling cell and animal size (Pardee 1989; Norbury and Nurse 1992; Destefanis *et al*. 2020). The ‘Minute’ syndrome might therefore link the haploinsufficiency of ribosomal genes to defects in *Myc* mRNA and protein turnover. We propose that genes at ‘Minute’ loci might be main regulators of *Myc* in highly proliferative cells during early development.

DEAD-box RNA helicases constitute a large family of proteins that induce structural changes in RNA molecules required for their physiological function. Many genes encoding DEAD-box RNA helicases are targets of MYC/Max heterodimers (Grandori *et al*. 1996). Furthermore, DEAD-box RNA helicases associate with the target genes of MYC/Max heterodimers, exhibit nucleic acid duplex unwinding ability, and are essential for rRNA processing and ribosome biogenesis during cell growth and germline development (Hirling *et al*. 1989; Grandori *et al*. 1996; Lasko 2013; Kara *et al*. 2023).

Members of the DEAD-box RNA helicase family, including eIF4A (*eIF-4a*), Dead-box-1 (*Ddx1*), and Abstrakt (*abs*), were pulled down by the Myc-CRMs. eIF4A is dependent on eIF4B (also found associated with Myc-CRMs) for its helicase activity. Together with eIF4B and ribosomal protein S6 kinase (*S6k*), a component of the insulin receptor signaling pathway (Radimerski *et al*. 2002), eIF4A selectively regulates the translation of pro-survival mRNAs, including MYC and Bcl-2, to promote cell proliferation (Wolfe *et al*. 2014; Roux and Topisirovic 2018). Furthermore, eIF4A has been demonstrated to behave as an oncogene and to drive the overexpression of *MYC* and *Notch1* through mRNA translation in acute lymphatic leukemia (ALL) (Wolfe *et al*. 2014).

Dead-box-1 (Ddx1), with roles in nucleic acid duplex unwinding (Gene Ontology Curators 2002) and protein synthesis (Rafti *et al*. 1996), is a positive regulator of cell proliferation and germ cell development in early embryos (Rafti *et al*. 1996; Germain *et al*. 2015). The human homolog DDX1, which has high similarity to the *Drosophila* Ddx1, has been found to be overexpressed and amplified together with MYCN in subsets of retinoblastoma and neuroblastoma tumors, as well as some other cancer types like breast cancer (Squire *et al*. 1995; Godbout *et al*. 1998; Balko and Arteaga 2011; Germain *et al*. 2011). This suggests that Ddx1 plays a role in cell proliferation with potential connections to the regulation of growth regulators like *Myc*.

Abstrakt, a member of the DEAD/DEAH-box RNA helicases, is required for cell survival throughout the *Drosophila* life cycle, with essential impacts on cell morphology, mRNA processing and localization, and apoptosis (Irion and Leptin 1999). Abstrakt also functions as a Notch regulator by modulating Inscuteable (*insc*) mRNA levels in certain sibling progenitor cells during neurogenesis (Irion *et al*. 2004). The high enrichment of Abstrakt observed in all Myc-CRM samples might indicate a context-dependent role for Abstrakt in regulating *Myc* expression. In conclusion, based on our proteomics data, DEAD-box RNA helicases appear to have a significant influence on the direct or indirect control of *Myc* expression across different developmental stages and in disease.

## Concluding remarks

The Myc-CRMs tested in this study may contain overlapping binding sites that accommodate the binding of multiple transcription factors, either simultaneously or sequentially. Alternatively, factors might compete for binding to the same site(s) depending on the transcriptional context. The exact sequences of these binding sites, the specific binding mechanisms, relevant allosteric conditions, and the potential involvement of putative adaptor molecules and mediators remain to be investigated.

Considering the stringent filtering applied to the obtained data and the implementation of different quality control approaches, we could deliver credible datasets with minimal background noise. The dataset represents a wide range of factors regulating *Myc* expression at different levels of gene control mechanisms. Extensive *in vivo* follow-up analyses are necessary to provide sufficient proof for the functionality of the identified proteins and to validate these results further. These findings provide a solid foundation for the scientific community to plan and carry out numerous exploratory studies aimed at clarifying the molecular and cellular mechanisms of the genes of interest, as well as their linkages to the powerful network of the super-oncogene *MYC*.

## Supporting information

Highlights

## Acknowledgements

The authors thank C. Kerscher (Licor Biosciences) for analytical support with the Gel Shift Images; Prof. V. Schwämmle and Prof. A. Rogowska-Wrzesinska (University of Southern Denmark) for assistance with using the Software Tool ComplexBrowser; Prof R. Dubey and Prof M. Calcagni for access to research space and laboratory facilities of the University Hospital Zurich; C. Scavenius Sonne-Schmidt (Danish Technological Institute, DTI) for analytical support with the mass spectrometry data; and D. Ackermann and C. Winiger (Oligonucleotide services, Microsynth AG, Switzerland) for the synthesis of the novel oligonucleotides. The authors also thank Prof. H-P Lipp for a critical reading of the final pre-submission manuscript and the members of BroadPharm company (San Diego, CA) for the synthesis of the free-end PEG-linker ‘BP-25103 DBCO-PEG23-amine’.

## CRediT authorship contribution statement

**Jasmine Kharazmi**: Conceptualization, Data curation, Formal analysis, Funding acquisition, Investigation, Methodology, Project administration, Resources, Supervision, Validation, visualization, Writing - Original draft preparation, Writing - review and editing. **Cameron Moshfegh**: Data curation, Formal analysis, Funding acquisition, Methodology, Software, visualization, Review and editing. **Thomas Brody**: Formal analysis, Review and editing.

## References

1. Abaza, I., O. Coll, S. Patalano and F. Gebauer, 2006 Drosophila UNR is required for translational repression of male-specific lethal 2 mRNA during regulation of X-chromosome dosage compensation. Genes Dev 20: 380–389.

2. Abel, J., R. Eskeland, G. D. Raffa, E. Kremmer and A. Imhof, 2009 Drosophila HP1c is regulated by an auto-regulatory feedback loop through its binding partner Woc. PLoS One 4: e5089.

3. Aberger, F., and A. Ruiz i Altaba, 2014 Context-dependent signal integration by the GLI code: the oncogenic load, pathways, modifiers and implications for cancer therapy. Semin Cell Dev Biol 33: 93–104.

4. Abou Faycal, C., S. Gazzeri and B. Eymin, 2016 RNA splicing, cell signaling, and response to therapies. Curr Opin Oncol 28: 58–64.

5. Acar, M., H. Jafar-Nejad, H. Takeuchi, A. Rajan, D. Ibrani et al., 2008 Rumi is a CAP10 domain glycosyltransferase that modifies Notch and is required for Notch signaling. Cell 132: 247–258.

6. Adelman, K., W. Wei, M. B. Ardehali, J. Werner, B. Zhu et al., 2006 Drosophila Paf1 modulates chromatin structure at actively transcribed genes. Mol Cell Biol 26: 250–260.

7. Agianian, B., K. Leonard, E. Bonte, H. Van der Zandt, P. B. Becker et al., 1999 The glutamine-rich domain of the Drosophila GAGA factor is necessary for amyloid fibre formation in vitro, but not for chromatin remodelling. J Mol Biol 285: 527–544.

8. Akan, I., D. C. Love, K. R. Harwood, M. R. Bond and J. A. Hanover, 2016 Drosophila O-GlcNAcase Deletion Globally Perturbs Chromatin O-GlcNAcylation. J Biol Chem 291: 9906–9919.

9. Akhmetova, K., M. Balasov, R. P. Huijbregts and I. Chesnokov, 2015 Functional insight into the role of Orc6 in septin complex filament formation in Drosophila. Mol Biol Cell 26: 15–28.

10. Alfonso-Parra, C., and K. A. Maggert, 2010 Drosophila SAF-B links the nuclear matrix, chromosomes, and transcriptional activity. PLoS One 5: e10248.

11. Alland, L., R. Muhle, H. Hou, Jr., J. Potes, L. Chin et al., 1997 Role for N-CoR and histone deacetylase in Sin3-mediated transcriptional repression. Nature 387: 49–55.

12. Alpar, L., C. Bergantinos and L. A. Johnston, 2018 Spatially Restricted Regulation of Spatzle/Toll Signaling during Cell Competition. Dev Cell 46: 706–719 e705.

13. Anatskaya, O. V., and A. E. Vinogradov, 2022 Polyploidy and Myc Proto-Oncogenes Promote Stress Adaptation via Epigenetic Plasticity and Gene Regulatory Network Rewiring. Int J Mol Sci 23.

14. Andersen, D. S., and S. J. Leevers, 2007 The essential Drosophila ATP-binding cassette domain protein, pixie, binds the 40 S ribosome in an ATP-dependent manner and is required for translation initiation. J Biol Chem 282: 14752–14760.

15. Andersen, P. R., L. Tirian, M. Vunjak and J. Brennecke, 2017 A heterochromatin-dependent transcription machinery drives piRNA expression. Nature 549: 54–59.

16. Anderson, A. E., U. C. Karandikar, K. L. Pepple, Z. Chen, A. Bergmann et al., 2011 The enhancer of trithorax and polycomb gene Caf1/p55 is essential for cell survival and patterning in Drosophila development. Development 138: 1957–1966.

17. Anderson, E. C., and P. O. Catnaigh, 2015 Regulation of the expression and activity of Unr in mammalian cells. Biochem Soc Trans 43: 1241–1246.

18. Andersson, R., C. Gebhard, I. Miguel-Escalada, I. Hoof, J. Bornholdt et al., 2014 An atlas of active enhancers across human cell types and tissues. Nature 507: 455–461.

19. Andrulis, E. D., J. Werner, A. Nazarian, H. Erdjument-Bromage, P. Tempst et al., 2002 The RNA processing exosome is linked to elongating RNA polymerase II in Drosophila. Nature 420: 837–841.

20. Andruss, B. F., A. Q. Lu and K. Beckingham, 1997 Expression of calmodulin in Drosophila is highly regulated in a stage- and tissue-specific manner. Dev Genes Evol 206: 541–545.

21. Angulo, M., M. Corominas and F. Serras, 2004 Activation and repression activities of ash2 in Drosophila wing imaginal discs. Development 131: 4943–4953.

22. Antonucci, L., D. D’Amico, L. Di Magno, S. Coni, L. Di Marcotullio et al., 2014 CNBP regulates wing development in Drosophila melanogaster by promoting IRES-dependent translation of dMyc. Cell Cycle 13: 434–439.

23. Aoyagi, N., S. Matsuoka, A. Furunobu, A. Matsukage and K. Sakaguchi, 1994 Drosophila DNA polymerase delta. Purification and characterization. J Biol Chem 269: 6045–6050.

24. Aoyagi, N., and D. A. Wassarman, 2000 Genes encoding Drosophila melanogaster RNA polymerase II general transcription factors: diversity in TFIIA and TFIID components contributes to gene-specific transcriptional regulation. J Cell Biol 150: F45–50.

25. Arama, E., J. Agapite and H. Steller, 2003 Caspase activity and a specific cytochrome C are required for sperm differentiation in Drosophila. Dev Cell 4: 687–697.

26. Arce, L., K. T. Pate and M. L. Waterman, 2009 Groucho binds two conserved regions of LEF-1 for HDAC-dependent repression. BMC Cancer 9: 159.

27. Archbold, H. C., C. Broussard, M. V. Chang and K. M. Cadigan, 2014 Bipartite recognition of DNA by TCF/Pangolin is remarkably flexible and contributes to transcriptional responsiveness and tissue specificity of wingless signaling. PLoS Genet 10: e1004591.

28. Ardehali, M. B., J. Yao, K. Adelman, N. J. Fuda, S. J. Petesch et al., 2009 Spt6 enhances the elongation rate of RNA polymerase II in vivo. EMBO J 28: 1067–1077.

29. Armstrong, J. D., M. J. Texada, R. Munjaal, D. A. Baker and K. M. Beckingham, 2006 Gravitaxis in Drosophila melanogaster: a forward genetic screen. Genes Brain Behav 5: 222–239.

30. Arvola, R. M., C. T. Chang, J. P. Buytendorp, Y. Levdansky, E. Valkov et al., 2020 Unique repression domains of Pumilio utilize deadenylation and decapping factors to accelerate destruction of target mRNAs. Nucleic Acids Res 48: 1843–1871.

31. Asano, K., T. G. Kinzy, W. C. Merrick and J. W. Hershey, 1997 Conservation and diversity of eukaryotic translation initiation factor eIF3. J Biol Chem 272: 1101–1109.

32. Asaoka-Taguchi, M., M. Yamada, A. Nakamura, K. Hanyu and S. Kobayashi, 1999 Maternal Pumilio acts together with Nanos in germline development in Drosophila embryos. Nat Cell Biol 1: 431–437.

33. Asaoka, T., J. Almagro, C. Ehrhardt, I. Tsai, A. Schleiffer et al., 2016 Linear ubiquitination by LUBEL has a role in Drosophila heat stress response. EMBO Rep 17: 1624–1640.

34. Ashton-Beaucage, D., C. M. Udell, P. Gendron, M. Sahmi, M. Lefrancois et al., 2014 A functional screen reveals an extensive layer of transcriptional and splicing control underlying RAS/MAPK signaling in Drosophila. PLoS Biol 12: e1001809.

35. Azuma, Y., T. Tokuda, Y. Kushimura, I. Yamamoto, I. Mizuta et al., 2018 Hippo, Drosophila MST, is a novel modifier of motor neuron degeneration induced by knockdown of Caz, Drosophila FUS. Exp Cell Res 371: 311–321.

36. Azuma, Y., T. Tokuda, M. Shimamura, A. Kyotani, H. Sasayama et al., 2014 Identification of ter94, Drosophila VCP, as a strong modulator of motor neuron degeneration induced by knockdown of Caz, Drosophila FUS. Hum Mol Genet 23: 3467–3480.

37. Badenhorst, P., M. Voas, I. Rebay and C. Wu, 2002 Biological functions of the ISWI chromatin remodeling complex NURF. Genes Dev 16: 3186–3198.

38. Badenhorst, P., H. Xiao, L. Cherbas, S. Y. Kwon, M. Voas et al., 2005 The Drosophila nucleosome remodeling factor NURF is required for Ecdysteroid signaling and metamorphosis. Genes Dev 19: 2540–2545.

39. Baehrecke, E. H., 1997 who encodes a KH RNA binding protein that functions in muscle development. Development 124: 1323–1332.

40. Bag, I., Y. Chen, K. D’Orazio, P. Lopez, S. Wenzel et al., 2022 Isha is a su(Hw) mRNA-binding protein required for gypsy insulator function. G3 (Bethesda) 12.

41. Bajpai, R., K. Makhijani, P. R. Rao and L. S. Shashidhara, 2004 Drosophila Twins regulates Armadillo levels in response to Wg/Wnt signal. Development 131: 1007–1016.

42. Baker, D. A., B. Mille-Baker, S. M. Wainwright, D. Ish-Horowicz and N. J. Dibb, 2001 Mae mediates MAP kinase phosphorylation of Ets transcription factors in Drosophila. Nature 411: 330–334.

43. Bakkenist, C. J., and S. Cotterill, 1994 The 50-kDa primase subunit of Drosophila melanogaster DNA polymerase alpha. Molecular characterization of the gene and functional analysis of the overexpressed protein. J Biol Chem 269: 26759–26766.

44. Balko, J. M., and C. L. Arteaga, 2011 Dead-box or black-box: is DDX1 a potential biomarker in breast cancer? Breast Cancer Res Treat 127: 65–67.

45. Baluapuri, A., J. Hofstetter, N. Dudvarski Stankovic, T. Endres, P. Bhandare et al., 2019 MYC Recruits SPT5 to RNA Polymerase II to Promote Processive Transcription Elongation. Mol Cell 74: 674–687 e611.

46. Barbosa, V., M. Gatt, E. Rebollo, C. Gonzalez and D. M. Glover, 2003 Drosophila dd4 mutants reveal that gammaTuRC is required to maintain juxtaposed half spindles in spermatocytes. J Cell Sci 116: 929–941.

47. Bardin, A. J., R. Le Borgne and F. Schweisguth, 2004 Asymmetric localization and function of cell-fate determinants: a fly’s view. Curr Opin Neurobiol 14: 6–14.

48. Barish, S., T. S. Barakat, B. C. Michel, N. Mashtalir, J. B. Phillips et al., 2020 BICRA, a SWI/SNF Complex Member, Is Associated with BAF-Disorder Related Phenotypes in Humans and Model Organisms. Am J Hum Genet 107: 1096–1112.

49. Barker, N., P. J. Morin and H. Clevers, 2000 The Yin-Yang of TCF/beta-catenin signaling. Adv Cancer Res 77: 1–24.

50. Barral, A., and J. Dejardin, 2023 The chromatin signatures of enhancers and their dynamic regulation. Nucleus 14: 2160551.

51. Bashirullah, A., G. Lam, V. P. Yin and C. S. Thummel, 2007 dTrf2 is required for transcriptional and developmental responses to ecdysone during Drosophila metamorphosis. Dev Dyn 236: 3173–3179.

52. Bauer, A., S. Chauvet, O. Huber, F. Usseglio, U. Rothbacher et al., 2000 Pontin52 and reptin52 function as antagonistic regulators of beta-catenin signalling activity. EMBO J 19: 6121–6130.

53. Bejarano, F., C. M. Luque, H. Herranz, G. Sorrosal, N. Rafel et al., 2008 A gain-of-function suppressor screen for genes involved in dorsal-ventral boundary formation in the Drosophila wing. Genetics 178: 307–323.

54. Bello-Fernandez, C., G. Packham and J. L. Cleveland, 1993 The ornithine decarboxylase gene is a transcriptional target of c-Myc. Proc Natl Acad Sci U S A 90: 7804–7808.

55. Bellosta, P., and P. Gallant, 2010 Myc Function in Drosophila. Genes Cancer 1: 542–546.

56. Bellosta, P., T. Hulf, S. Balla Diop, F. Usseglio, J. Pradel et al., 2005 Myc interacts genetically with Tip48/Reptin and Tip49/Pontin to control growth and proliferation during Drosophila development. Proc Natl Acad Sci U S A 102: 11799–11804.

57. Beltran, S., M. Angulo, M. Pignatelli, F. Serras and M. Corominas, 2007 Functional dissection of the ash2 and ash1 transcriptomes provides insights into the transcriptional basis of wing phenotypes and reveals conserved protein interactions. Genome Biol 8: R67.

58. Benassayag, C., L. Montero, N. Colombie, P. Gallant, D. Cribbs et al., 2005 Human c-Myc isoforms differentially regulate cell growth and apoptosis in Drosophila melanogaster. Mol Cell Biol 25: 9897–9909.

59. Berman, B. P., B. D. Pfeiffer, T. R. Laverty, S. L. Salzberg, G. M. Rubin et al., 2004 Computational identification of developmental enhancers: conservation and function of transcription factor binding-site clusters in Drosophila melanogaster and Drosophila pseudoobscura. Genome Biol 5: R61.

60. Bertolotti, A., Y. Lutz, D. J. Heard, P. Chambon and L. Tora, 1996 hTAF(II)68, a novel RNA/ssDNA-binding protein with homology to the pro-oncoproteins TLS/FUS and EWS is associated with both TFIID and RNA polymerase II. EMBO J 15: 5022–5031.

61. Bhambhani, C., J. L. Chang, D. L. Akey and K. M. Cadigan, 2011 The oligomeric state of CtBP determines its role as a transcriptional co-activator and co-repressor of Wingless targets. EMBO J 30: 2031–2043.

62. Biedermann, B., H. R. Hotz and R. Ciosk, 2010 The Quaking family of RNA-binding proteins: coordinators of the cell cycle and differentiation. Cell Cycle 9: 1929–1933.

63. Biggs, W. H., 3rd, K. H. Zavitz, B. Dickson, A. van der Straten, D. Brunner et al., 1994 The Drosophila rolled locus encodes a MAP kinase required in the sevenless signal transduction pathway. EMBO J 13: 1628-1635.

64. Biryukova, I., and P. Heitzler, 2005 The Drosophila LIM-homeo domain protein Islet antagonizes pro-neural cell specification in the peripheral nervous system. Dev Biol 288: 559–570.

65. Blanco, J., J. C. Cooper and N. E. Baker, 2020 Roles of C/EBP class bZip proteins in the growth and cell competition of Rp (’Minute’) mutants in Drosophila. Elife 9.

66. Bode, J., S. Goetze, H. Heng, S. A. Krawetz and C. Benham, 2003 From DNA structure to gene expression: mediators of nuclear compartmentalization and dynamics. Chromosome Res 11: 435–445.

67. Bode, J., M. Stengert-Iber, V. Kay, T. Schlake and A. Dietz-Pfeilstetter, 1996 Scaffold/matrix-attached regions: topological switches with multiple regulatory functions. Crit Rev Eukaryot Gene Expr 6: 115–138.

68. Boi, D., E. Rubini, S. Breccia, G. Guarguaglini and A. Paiardini, 2023 When Just One Phosphate Is One Too Many: The Multifaceted Interplay between Myc and Kinases. Int J Mol Sci 24.

69. Bonchuk, A., S. Denisov, P. Georgiev and O. Maksimenko, 2011 Drosophila BTB/POZ domains of “ttk group” can form multimers and selectively interact with each other. J Mol Biol 412: 423–436.

70. Bonke, M., M. Turunen, M. Sokolova, A. Vaharautio, T. Kivioja et al., 2013 Transcriptional networks controlling the cell cycle. G3 (Bethesda) 3: 75-90.

71. Bonnay, F., X. H. Nguyen, E. Cohen-Berros, L. Troxler, E. Batsche et al., 2014 Akirin specifies NF-kappaB selectivity of Drosophila innate immune response via chromatin remodeling. EMBO J 33: 2349–2362.

72. Borg, R. M., R. Bordonne, N. Vassallo and R. J. Cauchi, 2015 Genetic Interactions between the Members of the SMN-Gemins Complex in Drosophila. PLoS One 10: e0130974.

73. Bose, A., B. Kahali, S. Zhang, J. M. Lin, R. Allada et al., 2006 Drosophila CK2 regulates lateral-inhibition during eye and bristle development. Mech Dev 123: 649–664.

74. Bosso, G., F. Cipressa, M. L. Moroni, R. Pennisi, J. Albanesi et al., 2019 NBS1 interacts with HP1 to ensure genome integrity. Cell Death Dis 10: 951.

75. Boube, M., C. Faucher, L. Joulia, D. L. Cribbs and H. M. Bourbon, 2000 Drosophila homologs of transcriptional mediator complex subunits are required for adult cell and segment identity specification. Genes Dev 14: 2906–2917.

76. Bouchard, C., K. Thieke, A. Maier, R. Saffrich, J. Hanley-Hyde et al., 1999 Direct induction of cyclin D2 by Myc contributes to cell cycle progression and sequestration of p27. EMBO J 18: 5321–5333.

77. Boulon, S., N. Marmier-Gourrier, B. Pradet-Balade, L. Wurth, C. Verheggen et al., 2008 The Hsp90 chaperone controls the biogenesis of L7Ae RNPs through conserved machinery. J Cell Biol 180: 579-595.

78. Braat, A. K., N. Yan, E. Arn, D. Harrison and P. M. Macdonald, 2004 Localization-dependent oskar protein accumulation; control after the initiation of translation. Dev Cell 7: 125–131.

79. Bras, S., S. Martin-Lanneree, V. Gobert, B. Auge, O. Breig et al., 2012 Myeloid leukemia factor is a conserved regulator of RUNX transcription factor activity involved in hematopoiesis. Proc Natl Acad Sci U S A 109: 4986–4991.

80. Brentrup, D., H. Lerch, H. Jackle and M. Noll, 2000 Regulation of Drosophila wing vein patterning: net encodes a bHLH protein repressing rhomboid and is repressed by rhomboid-dependent Egfr signalling. Development 127: 4729–4741.

81. Bretones, G., M. D. Delgado and J. Leon, 2015 Myc and cell cycle control. Biochim Biophys Acta 1849: 506–516.

82. Brewer-Jensen, P., C. B. Wilson, J. Abernethy, L. Mollison, S. Card et al., 2016 Suppressor of sable [Su(s)] and Wdr82 down-regulate RNA from heat-shock-inducible repetitive elements by a mechanism that involves transcription termination. RNA 22: 139–154.

83. Brickey, W. J., and A. L. Greenleaf, 1995 Functional studies of the carboxy-terminal repeat domain of Drosophila RNA polymerase II in vivo. Genetics 140: 599–613.

84. Broderick, P., D. Cunningham, J. Vijayakrishnan, R. Cooke, A. Ashworth et al., 2010 IRF4 polymorphism rs872071 and risk of Hodgkin lymphoma. Br J Haematol 148: 413–415.

85. Brody, T., W. Rasband, K. Baler, A. Kuzin, M. Kundu et al., 2007 cis-Decoder discovers constellations of conserved DNA sequences shared among tissue-specific enhancers. Genome Biol 8: R75.

86. Brou, C., F. Logeat, M. Lecourtois, J. Vandekerckhove, P. Kourilsky et al., 1994 Inhibition of the DNA-binding activity of Drosophila suppressor of hairless and of its human homolog, KBF2/RBP-J kappa, by direct protein-protein interaction with Drosophila hairless. Genes Dev 8: 2491-2503.

87. Brumby, A. M., C. B. Zraly, J. A. Horsfield, J. Secombe, R. Saint et al., 2002 Drosophila cyclin E interacts with components of the Brahma complex. EMBO J 21: 3377–3389.

88. Brun, S., S. Vidal, P. Spellman, K. Takahashi, H. Tricoire et al., 2006 The MAPKKK Mekk1 regulates the expression of Turandot stress genes in response to septic injury in Drosophila. Genes Cells 11: 397–407.

89. Brunner, D., K. Ducker, N. Oellers, E. Hafen, H. Scholz et al., 1994 The ETS domain protein pointed-P2 is a target of MAP kinase in the sevenless signal transduction pathway. Nature 370: 386–389.

90. Bujakowska, K., C. Maubaret, C. F. Chakarova, N. Tanimoto, S. C. Beck et al., 2009 Study of gene-targeted mouse models of splicing factor gene Prpf31 implicated in human autosomal dominant retinitis pigmentosa (RP). Invest Ophthalmol Vis Sci 50: 5927-5933.

91. Burgio, G., G. La Rocca, A. Sala, W. Arancio, D. Di Gesu et al., 2008 Genetic identification of a network of factors that functionally interact with the nucleosome remodeling ATPase ISWI. PLoS Genet 4: e1000089.

92. Burke, T. W., and J. T. Kadonaga, 1997 The downstream core promoter element, DPE, is conserved from Drosophila to humans and is recognized by TAFII60 of Drosophila. Genes Dev 11: 3020–3031.

93. Butler, J. E., and J. T. Kadonaga, 2001 Enhancer-promoter specificity mediated by DPE or TATA core promoter motifs. Genes Dev 15: 2515–2519.

94. Cadigan, K. M., 2012 TCFs and Wnt/beta-catenin signaling: more than one way to throw the switch. Curr Top Dev Biol 98: 1–34.

95. Caforio, M., C. Sorino, S. Iacovelli, M. Fanciulli, F. Locatelli et al., 2018 Recent advances in searching c-Myc transcriptional cofactors during tumorigenesis. J Exp Clin Cancer Res 37: 239.

96. Cakouros, D., K. Mills, D. Denton, A. Paterson, T. Daish et al., 2008 dLKR/SDH regulates hormone-mediated histone arginine methylation and transcription of cell death genes. J Cell Biol 182: 481–495.

97. Campbell, S. D., F. Sprenger, B. A. Edgar and P. H. O’Farrell, 1995 Drosophila Wee1 kinase rescues fission yeast from mitotic catastrophe and phosphorylates Drosophila Cdc2 in vitro. Mol Biol Cell 6: 1333–1347.

98. Capelson, M., Y. Liang, R. Schulte, W. Mair, U. Wagner et al., 2010 Chromatin-bound nuclear pore components regulate gene expression in higher eukaryotes. Cell 140: 372–383.

99. Capp, C., J. Wu and T. S. Hsieh, 2009 Drosophila RecQ4 has a 3’-5’ DNA helicase activity that is essential for viability. J Biol Chem 284: 30845–30852.

100. Cappadocia, L., and C. D. Lima, 2018 Ubiquitin-like Protein Conjugation: Structures, Chemistry, and Mechanism. Chem Rev 118: 889–918.

101. Carbonell, A., A. Mazo, F. Serras and M. Corominas, 2013 Ash2 acts as an ecdysone receptor coactivator by stabilizing the histone methyltransferase Trr. Mol Biol Cell 24: 361–372.

102. Carreno, S., I. Kouranti, E. S. Glusman, M. T. Fuller, A. Echard et al., 2008 Moesin and its activating kinase Slik are required for cortical stability and microtubule organization in mitotic cells. J Cell Biol 180: 739–746.

103. Carrera, I., F. Janody, N. Leeds, F. Duveau and J. E. Treisman, 2008 Pygopus activates Wingless target gene transcription through the mediator complex subunits Med12 and Med13. Proc Natl Acad Sci U S A 105: 6644–6649.

104. Carrera, P., Y. M. Moshkin, S. Gronke, H. H. Sillje, E. A. Nigg et al., 2003 Tousled-like kinase functions with the chromatin assembly pathway regulating nuclear divisions. Genes Dev 17: 2578–2590.

105. Casper, A. L., K. Baxter and M. Van Doren, 2011 no child left behind encodes a novel chromatin factor required for germline stem cell maintenance in males but not females. Development 138: 3357–3366.

106. Castro, B., S. Barolo, A. M. Bailey and J. W. Posakony, 2005 Lateral inhibition in proneural clusters: cis-regulatory logic and default repression by Suppressor of Hairless. Development 132: 3333–3344.

107. Catarino, R. R., C. Neumayr and A. Stark, 2017 Promoting transcription over long distances. Nat Genet 49: 972–973.

108. Catarino, R. R., and A. Stark, 2018 Assessing sufficiency and necessity of enhancer activities for gene expression and the mechanisms of transcription activation. Genes Dev 32: 202–223.

109. Cauchi, R. J., K. E. Davies and J. L. Liu, 2008 A motor function for the DEAD-box RNA helicase, Gemin3, in Drosophila. PLoS Genet 4: e1000265.

110. Cavallo, R. A., R. T. Cox, M. M. Moline, J. Roose, G. A. Polevoy et al., 1998 Drosophila Tcf and Groucho interact to repress Wingless signalling activity. Nature 395: 604–608.

111. Cenci, G., R. B. Rawson, G. Belloni, D. H. Castrillon, M. Tudor et al., 1997 UbcD1, a Drosophila ubiquitin-conjugating enzyme required for proper telomere behavior. Genes Dev 11: 863–875.

112. Cencioni, C., F. Scagnoli, F. Spallotta, S. Nasi and B. Illi, 2023 The “Superoncogene” Myc at the Crossroad between Metabolism and Gene Expression in Glioblastoma Multiforme. Int J Mol Sci 24.

113. Chalkley, G. E., Y. M. Moshkin, K. Langenberg, K. Bezstarosti, A. Blastyak et al., 2008 The transcriptional coactivator SAYP is a trithorax group signature subunit of the PBAP chromatin remodeling complex. Mol Cell Biol 28: 2920–2929.

114. Chambers, C., K. Cermakova, Y. S. Chan, K. Kurtz, K. Wohlan et al., 2023 SWI/SNF Blockade Disrupts PU.1-Directed Enhancer Programs in Normal Hematopoietic Cells and Acute Myeloid Leukemia. Cancer Res 83: 983-996.

115. Chan, Y. B., I. Miguel-Aliaga, C. Franks, N. Thomas, B. Trulzsch et al., 2003 Neuromuscular defects in a Drosophila survival motor neuron gene mutant. Hum Mol Genet 12: 1367–1376.

116. Chang, J. L., H. V. Lin, T. A. Blauwkamp and K. M. Cadigan, 2008 Spenito and Split ends act redundantly to promote Wingless signaling. Dev Biol 314: 100–111.

117. Chang, Y. C., J. W. Wu, Y. C. Hsieh, T. H. Huang, Z. M. Liao et al., 2018 Rap1 Negatively Regulates the Hippo Pathway to Polarize Directional Protrusions in Collective Cell Migration. Cell Rep 22: 2160–2175.

118. Chao, A. T., H. A. Dierick, T. M. Addy and A. Bejsovec, 2003 Mutations in eukaryotic release factors 1 and 3 act as general nonsense suppressors in Drosophila. Genetics 165: 601–612.

119. Chartier, A., P. Klein, S. Pierson, N. Barbezier, T. Gidaro et al., 2015 Mitochondrial dysfunction reveals the role of mRNA poly(A) tail regulation in oculopharyngeal muscular dystrophy pathogenesis. PLoS Genet 11: e1005092.

120. Chau, B. L., K. P. Ng, K. K. Li and K. A. Lee, 2016 RGG boxes within the TET/FET family of RNA-binding proteins are functionally distinct. Transcription 7: 141–151.

121. Chazal, P. E., E. Daguenet, C. Wendling, N. Ulryck, C. Tomasetto et al., 2013 EJC core component MLN51 interacts with eIF3 and activates translation. Proc Natl Acad Sci U S A 110: 5903–5908.

122. Chen, D., Q. Wang, H. Huang, L. Xia, X. Jiang et al., 2009 Effete-mediated degradation of Cyclin A is essential for the maintenance of germline stem cells in Drosophila. Development 136: 4133–4142.

123. Chen, F. X., A. R. Woodfin, A. Gardini, R. A. Rickels, S. A. Marshall et al., 2015 PAF1, a Molecular Regulator of Promoter-Proximal Pausing by RNA Polymerase II. Cell 162: 1003–1015.

124. Chen, X., and L. Xu, 2010 Specific nucleoporin requirement for Smad nuclear translocation. Mol Cell Biol 30: 4022–4034.

125. Cheng, S. Y., and J. M. Bishop, 2002 Suppressor of Fused represses Gli-mediated transcription by recruiting the SAP18-mSin3 corepressor complex. Proc Natl Acad Sci U S A 99: 5442–5447.

126. Chiang, C. S., P. G. Mitsis and I. R. Lehman, 1993 DNA polymerase delta from embryos of Drosophila melanogaster. Proc Natl Acad Sci U S A 90: 9105–9109.

127. Cho, I. K., C. L. Chang and Q. X. Li, 2013 Diet-induced over-expression of flightless-I protein and its relation to flightlessness in Mediterranean fruit fly, Ceratitis capitata. PLoS One 8: e81099.

128. Chodagam, S., A. Royou, W. Whitfield, R. Karess and J. W. Raff, 2005 The centrosomal protein CP190 regulates myosin function during early Drosophila development. Curr Biol 15: 1308–1313.

129. Churchill, M. E., D. N. Jones, T. Glaser, H. Hefner, M. A. Searles et al., 1995 HMG-D is an architecture-specific protein that preferentially binds to DNA containing the dinucleotide TG. EMBO J 14: 1264–1275.

130. Ciapponi, L., G. Cenci, J. Ducau, C. Flores, D. Johnson-Schlitz et al., 2004 The Drosophila Mre11/Rad50 complex is required to prevent both telomeric fusion and chromosome breakage. Curr Biol 14: 1360–1366.

131. Ciapponi, L., G. Cenci and M. Gatti, 2006 The Drosophila Nbs protein functions in multiple pathways for the maintenance of genome stability. Genetics 173: 1447–1454.

132. Ciribilli, Y., P. Singh, R. Spanel, A. Inga and J. Borlak, 2015 Decoding c-Myc networks of cell cycle and apoptosis regulated genes in a transgenic mouse model of papillary lung adenocarcinomas. Oncotarget 6: 31569–31592.

133. Clegg, N. J., D. M. Frost, M. K. Larkin, L. Subrahmanyan, Z. Bryant et al., 1997 maelstrom is required for an early step in the establishment of Drosophila oocyte polarity: posterior localization of grk mRNA. Development 124: 4661–4671.

134. Clemens, J. C., C. A. Worby, N. Simonson-Leff, M. Muda, T. Maehama et al., 2000 Use of double-stranded RNA interference in Drosophila cell lines to dissect signal transduction pathways. Proc Natl Acad Sci U S A 97: 6499–6503.

135. Clouse, K. N., S. B. Ferguson and T. Schupbach, 2008 Squid, Cup, and PABP55B function together to regulate gurken translation in Drosophila. Dev Biol 313: 713-724.

136. Collins, B. E., C. B. Greer, B. C. Coleman and J. D. Sweatt, 2019 Histone H3 lysine K4 methylation and its role in learning and memory. Epigenetics Chromatin 12: 7.

137. Collins, R. T., and J. E. Treisman, 2000 Osa-containing Brahma chromatin remodeling complexes are required for the repression of wingless target genes. Genes Dev 14: 3140–3152.

138. Conerly, M. L., S. S. Teves, D. Diolaiti, M. Ulrich, R. N. Eisenman et al., 2010 Changes in H2A.Z occupancy and DNA methylation during B-cell lymphomagenesis. Genome Res 20: 1383–1390.

139. Connell-Crowley, L., M. Le Gall, D. J. Vo and E. Giniger, 2000 The cyclin-dependent kinase Cdk5 controls multiple aspects of axon patterning in vivo. Curr Biol 10: 599–602.

140. Cordero, J. B., R. K. Stefanatos, K. Myant, M. Vidal and O. J. Sansom, 2012 Non-autonomous crosstalk between the Jak/Stat and Egfr pathways mediates Apc1-driven intestinal stem cell hyperplasia in the Drosophila adult midgut. Development 139: 4524–4535.

141. Corona, D. F., A. Eberharter, A. Budde, R. Deuring, S. Ferrari et al., 2000 Two histone fold proteins, CHRAC-14 and CHRAC-16, are developmentally regulated subunits of chromatin accessibility complex (CHRAC). EMBO J 19: 3049-3059.

142. Cottage, C. T., B. Bailey, K. M. Fischer, D. Avitabile, B. Collins et al., 2010 Cardiac progenitor cell cycling stimulated by pim-1 kinase. Circ Res 106: 891–901.

143. Cotterill, S., G. Chui and I. R. Lehman, 1987 DNA polymerase-primase from embryos of Drosophila melanogaster. DNA primase subunits. J Biol Chem 262: 16105–16108.

144. Couso, J. P., E. Knust and A. Martinez Arias, 1995 Serrate and wingless cooperate to induce vestigial gene expression and wing formation in Drosophila. Curr Biol 5: 1437–1448.

145. Crest, J., A. Diz-Munoz, D. Y. Chen, D. A. Fletcher and D. Bilder, 2017 Organ sculpting by patterned extracellular matrix stiffness. Elife 6.

146. Crevel, G., E. Mathe and S. Cotterill, 2005 The Drosophila Cdc6/18 protein has functions in both early and late S phase in S2 cells. J Cell Sci 118: 2451–2459.

147. Crona, F., B. Singla and M. Mannervik, 2015 Gene expression profiling of brakeless mutant Drosophila embryos. Data Brief 5: 134–137.

148. Crosby, M. A., C. Miller, T. Alon, K. L. Watson, C. P. Verrijzer et al., 1999 The trithorax group gene moira encodes a brahma-associated putative chromatin-remodeling factor in Drosophila melanogaster. Mol Cell Biol 19: 1159–1170.

149. Crucianelli, C., J. Jaiswal, A. Vijayakumar Maya, L. Nogay, A. Cosolo et al., 2022 Distinct signaling signatures drive compensatory proliferation via S-phase acceleration. PLoS Genet 18: e1010516.

150. Csiszar, A., E. Vogelsang, H. Beug and M. Leptin, 2010 A novel conserved phosphotyrosine motif in the Drosophila fibroblast growth factor signaling adaptor Dof with a redundant role in signal transmission. Mol Cell Biol 30: 2017–2027.

151. Cuartero, S., U. Fresan, O. Reina, E. Planet and M. L. Espinas, 2014 Ibf1 and Ibf2 are novel CP190-interacting proteins required for insulator function. EMBO J 33: 637–647.

152. Curtis, B. J., C. B. Zraly, D. R. Marenda and A. K. Dingwall, 2011 Histone lysine demethylases function as co-repressors of SWI/SNF remodeling activities during Drosophila wing development. Dev Biol 350: 534–547.

153. Curtis, C. D., and C. T. Griffin, 2012 The chromatin-remodeling enzymes BRG1 and CHD4 antagonistically regulate vascular Wnt signaling. Mol Cell Biol 32: 1312–1320.

154. D’Andrea, L. D., and L. Regan, 2003 TPR proteins: the versatile helix. Trends Biochem Sci 28: 655–662.

155. Dahmus, G. K., C. V. Glover, D. L. Brutlag and M. E. Dahmus, 1984 Similarities in structure and function of calf thymus and Drosophila casein kinase II. J Biol Chem 259: 9001–9006.

156. Dai, H., C. Hogan, B. Gopalakrishnan, J. Torres-Vazquez, M. Nguyen et al., 2000 The zinc finger protein schnurri acts as a Smad partner in mediating the transcriptional response to decapentaplegic. Dev Biol 227: 373–387.

157. Dammai, V., B. Adryan, K. R. Lavenburg and T. Hsu, 2003 Drosophila awd, the homolog of human nm23, regulates FGF receptor levels and functions synergistically with shi/dynamin during tracheal development. Genes Dev 17: 2812–2824.

158. Daneshvar, K., A. Khan and J. M. Goodliffe, 2011 Myc localizes to histone locus bodies during replication in Drosophila. PLoS One 6: e23928.

159. Dang, C. V., 2012 MYC on the path to cancer. Cell 149: 22–35.

160. Dansereau, D. A., and P. Lasko, 2008 RanBPM regulates cell shape, arrangement, and capacity of the female germline stem cell niche in Drosophila melanogaster. J Cell Biol 182: 963–977.

161. Dao, L. T. M., A. O. Galindo-Albarran, J. A. Castro-Mondragon, C. Andrieu-Soler, A. Medina-Rivera et al., 2017 Genome-wide characterization of mammalian promoters with distal enhancer functions. Nat Genet 49: 1073–1081.

162. Das, A. K., P. W. Cohen and D. Barford, 1998a The structure of the tetratricopeptide repeats of protein phosphatase 5: implications for TPR-mediated protein-protein interactions. EMBO J 17: 1192–1199.

163. Das, P., L. L. Maduzia, H. Wang, A. L. Finelli, S. H. Cho et al., 1998b The Drosophila gene Medea demonstrates the requirement for different classes of Smads in dpp signaling. Development 125: 1519–1528.

164. Das Thakur, M., Y. Feng, R. Jagannathan, M. J. Seppa, J. B. Skeath et al., 2010 Ajuba LIM proteins are negative regulators of the Hippo signaling pathway. Curr Biol 20: 657–662.

165. De Celis, J. F., 2003 Pattern formation in the Drosophila wing: The development of the veins. Bioessays 25: 443–451.

166. de la Cova, C., and L. A. Johnston, 2006 Myc in model organisms: a view from the flyroom. Semin Cancer Biol 16: 303–312.

167. de Pretis, S., T. R. Kress, M. J. Morelli, A. Sabo, C. Locarno et al., 2017 Integrative analysis of RNA polymerase II and transcriptional dynamics upon MYC activation. Genome Res 27: 1658–1664.

168. Dean, K. A., A. K. Aggarwal and R. P. Wharton, 2002 Translational repressors in Drosophila. Trends Genet 18: 572–577.

169. Dean, S. O., and J. A. Spudich, 2006 Rho kinase’s role in myosin recruitment to the equatorial cortex of mitotic Drosophila S2 cells is for myosin regulatory light chain phosphorylation. PLoS One 1: e131.

170. Delanoue, R., M. Slaidina and P. Leopold, 2010 The steroid hormone ecdysone controls systemic growth by repressing dMyc function in Drosophila fat cells. Dev Cell 18: 1012–1021.

171. Demontis, F., and N. Perrimon, 2009 Integration of Insulin receptor/Foxo signaling and dMyc activity during muscle growth regulates body size in Drosophila. Development 136: 983–993.

172. Deng, X., and V. H. Meller, 2006 roX RNAs are required for increased expression of X-linked genes in Drosophila melanogaster males. Genetics 174: 1859–1866.

173. Destefanis, F., V. Manara and P. Bellosta, 2020 Myc as a Regulator of Ribosome Biogenesis and Cell Competition: A Link to Cancer. Int J Mol Sci 21.

174. Dey, B., S. Thukral, S. Krishnan, M. Chakrobarty, S. Gupta et al., 2012 DNA-protein interactions: methods for detection and analysis. Mol Cell Biochem 365: 279–299.

175. Dhanasekaran, R., A. Deutzmann, W. D. Mahauad-Fernandez, A. S. Hansen, A. M. Gouw et al., 2022 The MYC oncogene - the grand orchestrator of cancer growth and immune evasion. Nat Rev Clin Oncol 19: 23–36.

176. Diaz-Benjumea, F. J., and E. Hafen, 1994 The sevenless signalling cassette mediates Drosophila EGF receptor function during epidermal development. Development 120: 569–578.

177. Dingwall, A. K., S. J. Beek, C. M. McCallum, J. W. Tamkun, G. V. Kalpana et al., 1995 The Drosophila snr1 and brm proteins are related to yeast SWI/SNF proteins and are components of a large protein complex. Mol Biol Cell 6: 777–791.

178. Dobi, K. C., M. S. Halfon and M. K. Baylies, 2014 Whole-genome analysis of muscle founder cells implicates the chromatin regulator Sin3A in muscle identity. Cell Rep 8: 858–870.

179. Doerflinger, H., V. Zimyanin and D. St Johnston, 2022 The Drosophila anterior-posterior axis is polarized by asymmetric myosin activation. Curr Biol 32: 374–385 e374.

180. Dong, X., L. Tsuda, K. H. Zavitz, M. Lin, S. Li et al., 1999 ebi regulates epidermal growth factor receptor signaling pathways in Drosophila. Genes Dev 13: 954–965.

181. Dorfman, R., L. Glazer, U. Weihe, M. F. Wernet and B. Z. Shilo, 2002 Elbow and Noc define a family of zinc finger proteins controlling morphogenesis of specific tracheal branches. Development 129: 3585–3596.

182. Douziech, M., M. Sahmi, G. Laberge and M. Therrien, 2006 A KSR/CNK complex mediated by HYP, a novel SAM domain-containing protein, regulates RAS-dependent RAF activation in Drosophila. Genes Dev 20: 807–819.

183. Doyen, C. M., G. E. Chalkley, O. Voets, K. Bezstarosti, J. A. Demmers et al., 2015 A Testis-Specific Chaperone and the Chromatin Remodeler ISWI Mediate Repackaging of the Paternal Genome. Cell Rep 13: 1310–1318.

184. Dragan, A. I., J. Klass, C. Read, M. E. Churchill, C. Crane-Robinson et al., 2003 DNA binding of a non-sequence-specific HMG-D protein is entropy driven with a substantial non-electrostatic contribution. J Mol Biol 331: 795–813.

185. Dragojlovic-Munther, M., and J. A. Martinez-Agosto, 2013 Extracellular matrix-modulated Heartless signaling in Drosophila blood progenitors regulates their differentiation via a Ras/ETS/FOG pathway and target of rapamycin function. Dev Biol 384: 313–330.

186. Dreos, R., A. Sloutskin, N. Malachi, D. Ideses, P. Bucher et al., 2021 Computational identification and experimental characterization of preferred downstream positions in human core promoters. PLoS Comput Biol 17: e1009256.

187. Du, L., A. Sohr, G. Yan and S. Roy, 2018 Feedback regulation of cytoneme-mediated transport shapes a tissue-specific FGF morphogen gradient. Elife 7.

188. Ducat, D., S. Kawaguchi, H. Liu, J. R. Yates, 3rd and Y.Zheng, 2008 Regulation of microtubule assembly and organization in mitosis by the AAA+ ATPase Pontin. Mol Biol Cell 19: 3097-3110.

189. Duman-Scheel, M., L. A. Johnston and W. Du, 2004 Repression of dMyc expression by Wingless promotes Rbf-induced G1 arrest in the presumptive Drosophila wing margin. Proc Natl Acad Sci U S A 101: 3857–3862.

190. Duncan, K., M. Grskovic, C. Strein, K. Beckmann, R. Niggeweg et al., 2006 Sex-lethal imparts a sex-specific function to UNR by recruiting it to the msl-2 mRNA 3’ UTR: translational repression for dosage compensation. Genes Dev 20: 368–379.

191. Duncan, R., L. Bazar, G. Michelotti, T. Tomonaga, H. Krutzsch et al., 1994 A sequence-specific, single-strand binding protein activates the far upstream element of c-myc and defines a new DNA-binding motif. Genes Dev 8: 465–480.

192. Dworkin, I., and G. Gibson, 2006 Epidermal growth factor receptor and transforming growth factor-beta signaling contributes to variation for wing shape in Drosophila melanogaster. Genetics 173: 1417–1431.

193. Dzhindzhev, N. S., S. L. Rogers, R. D. Vale and H. Ohkura, 2005 Distinct mechanisms govern the localisation of Drosophila CLIP-190 to unattached kinetochores and microtubule plus-ends. J Cell Sci 118: 3781–3790.

194. Eberharter, A., S. Ferrari, G. Langst, T. Straub, A. Imhof et al., 2001 Acf1, the largest subunit of CHRAC, regulates ISWI-induced nucleosome remodelling. EMBO J 20: 3781–3788.

195. Eckert, K., J. M. Saliou, L. Monlezun, A. Vigouroux, N. Atmane et al., 2010 The Pih1-Tah1 cochaperone complex inhibits Hsp90 molecular chaperone ATPase activity. J Biol Chem 285: 31304–31312.

196. Edenfeld, G., G. Volohonsky, K. Krukkert, E. Naffin, U. Lammel et al., 2006 The splicing factor crooked neck associates with the RNA-binding protein HOW to control glial cell maturation in Drosophila. Neuron 52: 969–980.

197. Edgar, B. A., J. Britton, A. F. de la Cruz, L. A. Johnston, D. Lehman et al., 2001 Pattern- and growth-linked cell cycles in Drosophila development. Novartis Found Symp 237: 3–12; discussion 12-18, 36-42.

198. Eeftens, J. M., J. van der Torre, D. R. Burnham and C. Dekker, 2015 Copper-free click chemistry for attachment of biomolecules in magnetic tweezers. BMC Biophys 8: 9.

199. Eggert, H., A. Gortchakov and H. Saumweber, 2004 Identification of the Drosophila interband-specific protein Z4 as a DNA-binding zinc-finger protein determining chromosomal structure. J Cell Sci 117: 4253–4264.

200. Eivers, E., L. C. Fuentealba, V. Sander, J. C. Clemens, L. Hartnett et al., 2009 Mad is required for wingless signaling in wing development and segment patterning in Drosophila. PLoS One 4: e6543.

201. Ekengren, S., and D. Hultmark, 2001 A family of Turandot-related genes in the humoral stress response of Drosophila. Biochem Biophys Res Commun 284: 998–1003.

202. Emelyanov, A. V., A. Y. Konev, E. Vershilova and D. V. Fyodorov, 2010 Protein complex of Drosophila ATRX/XNP and HP1a is required for the formation of pericentric beta-heterochromatin in vivo. J Biol Chem 285: 15027–15037.

203. Espinas, M. L., S. Canudas, L. Fanti, S. Pimpinelli, J. Casanova et al., 2000 The GAGA factor of Drosophila interacts with SAP18, a Sin3-associated polypeptide. EMBO Rep 1: 253–259.

204. Espinas, M. L., E. Jimenez-Garcia, A. Vaquero, S. Canudas, J. Bernues et al., 1999 The N-terminal POZ domain of GAGA mediates the formation of oligomers that bind DNA with high affinity and specificity. J Biol Chem 274: 16461–16469.

205. Fabrizio, J. J., G. Hime, S. K. Lemmon and C. Bazinet, 1998 Genetic dissection of sperm individualization in Drosophila melanogaster. Development 125: 1833–1843.

206. Fahey-Lozano, N., J. E. La Marca, M. Portela and H. E. Richardson, 2019 Drosophila Models of Cell Polarity and Cell Competition in Tumourigenesis. Adv Exp Med Biol 1167: 37–64.

207. Falkenburg, D., B. Dworniczak, D. M. Faust and E. K. Bautz, 1987 RNA polymerase II of Drosophila. Relation of its 140,000 Mr subunit to the beta subunit of Escherichia coli RNA polymerase. J Mol Biol 195: 929-937.

208. Fang, M., J. Li, T. Blauwkamp, C. Bhambhani, N. Campbell et al., 2006 C-terminal-binding protein directly activates and represses Wnt transcriptional targets in Drosophila. EMBO J 25: 2735–2745.

209. Farkas, G., J. Gausz, M. Galloni, G. Reuter, H. Gyurkovics et al., 1994 The Trithorax-like gene encodes the Drosophila GAGA factor. Nature 371: 806–808.

210. Feillet, C., P. Krusche, F. Tamanini, R. C. Janssens, M. J. Downey et al., 2014 Phase locking and multiple oscillating attractors for the coupled mammalian clock and cell cycle. Proc Natl Acad Sci U S A 111: 9828–9833.

211. Fiedler, M., M. Graeb, J. Mieszczanek, T. J. Rutherford, C. M. Johnson et al., 2015 An ancient Pygo-dependent Wnt enhanceosome integrated by Chip/LDB-SSDP. Elife 4.

212. Findley, S. D., M. Tamanaha, N. J. Clegg and H. Ruohola-Baker, 2003 Maelstrom, a Drosophila spindle-class gene, encodes a protein that colocalizes with Vasa and RDE1/AGO1 homolog, Aubergine, in nuage. Development 130: 859–871.

213. Fioresi, R., P. Demurtas and G. Perini, 2022 Deep learning for MYC binding site recognition. Front Bioinform 2: 1015993.

214. Firth, L. C., and N. E. Baker, 2005 Extracellular signals responsible for spatially regulated proliferation in the differentiating Drosophila eye. Dev Cell 8: 541–551.

215. Font-Burgada, J., D. Rossell, H. Auer and F. Azorin, 2008 Drosophila HP1c isoform interacts with the zinc-finger proteins WOC and Relative-of-WOC to regulate gene expression. Genes Dev 22: 3007–3023.

216. Fouix, S., S. Martin-Lanneree, M. Sanial, L. Morla, C. Lamour-Isnard et al., 2003 Over-expression of a novel nuclear interactor of Suppressor of fused, the Drosophila myelodysplasia/myeloid leukaemia factor, induces abnormal morphogenesis associated with increased apoptosis and DNA synthesis. Genes Cells 8: 897–911.

217. Franz, A., D. Shlyueva, E. Brunner, A. Stark and K. Basler, 2017 Probing the canonicity of the Wnt/Wingless signaling pathway. PLoS Genet 13: e1006700.

218. Fratini, L., M. G. S. Dalmolin, M. Sinigaglia, A. da Silveira Perla, C. B. de Farias et al., 2023 ZEB1 is a Subgroup-Specific Marker of Prognosis and Potential Drug Target in Medulloblastoma. Neuromolecular Med 25: 64–74.

219. Fulga, T. A., and P. Rorth, 2002 Invasive cell migration is initiated by guided growth of long cellular extensions. Nat Cell Biol 4: 715–719.

220. Funakoshi, M., M. Tsuda, K. Muramatsu, H. Hatsuda, S. Morishita et al., 2018 Overexpression of Larp4B downregulates dMyc and reduces cell and organ sizes in Drosophila. Biochem Biophys Res Commun 497: 762–768.

221. Furrer, M., M. Balbi, M. Albarca-Aguilera, M. Gallant, W. Herr et al., 2010 Drosophila Myc interacts with host cell factor (dHCF) to activate transcription and control growth. J Biol Chem 285: 39623–39636.

222. Gallant, P., 2013 Myc function in Drosophila. Cold Spring Harb Perspect Med 3: a014324.

223. Gallant, P., Y. Shiio, P. F. Cheng, S. M. Parkhurst and R. N. Eisenman, 1996 Myc and Max homologs in Drosophila. Science 274: 1523–1527.

224. Gambetta, M. C., K. Oktaba and J. Muller, 2009 Essential role of the glycosyltransferase sxc/Ogt in polycomb repression. Science 325: 93–96.

225. Gao, G., X. Bi, J. Chen, D. Srikanta and Y. S. Rong, 2009 Mre11-Rad50-Nbs complex is required to cap telomeres during Drosophila embryogenesis. Proc Natl Acad Sci U S A 106: 10728–10733.

226. Gaubatz, S., A. Imhof, R. Dosch, O. Werner, P. Mitchell et al., 1995 Transcriptional activation by Myc is under negative control by the transcription factor AP-2. EMBO J 14: 1508–1519.

227. Gaytan de Ayala Alonso, A., L. Gutierrez, C. Fritsch, B. Papp, D. Beuchle et al., 2007 A genetic screen identifies novel polycomb group genes in Drosophila. Genetics 176: 2099–2108.

228. Gelsthorpe, M. E., Z. Tan, A. Phillips, J. C. Eissenberg, A. Miller et al., 2006 Regulation of the Drosophila melanogaster protein, enhancer of rudimentary, by casein kinase II. Genetics 174: 265–270.

229. Gene Ontology Curators, -. 2002 Manual transfer of experimentally-verified manual GO annotation data to orthologs by curator judgment of sequence similarity.

230. Gennaro, V. J., H. Wedegaertner and S. B. McMahon, 2019 Interaction between the BAG1S isoform and HSP70 mediates the stability of anti-apoptotic proteins and the survival of osteosarcoma cells expressing oncogenic MYC. BMC Cancer 19: 258.

231. Gerasimova, T. I., D. A. Gdula, D. V. Gerasimov, O. Simonova and V. G. Corces, 1995 A Drosophila protein that imparts directionality on a chromatin insulator is an enhancer of position-effect variegation. Cell 82: 587–597.

232. Gerlach, J. M., M. Furrer, M. Gallant, D. Birkel, A. Baluapuri et al., 2017 PAF1 complex component Leo1 helps recruit Drosophila Myc to promoters. Proc Natl Acad Sci U S A 114: E9224–E9232.

233. Germain, D. R., K. Graham, D. D. Glubrecht, J. C. Hugh, J. R. Mackey et al., 2011 DEAD box 1: a novel and independent prognostic marker for early recurrence in breast cancer. Breast Cancer Res Treat 127: 53–63.

234. Germain, D. R., L. Li, M. R. Hildebrandt, A. J. Simmonds, S. C. Hughes et al., 2015 Loss of the Drosophila melanogaster DEAD box protein Ddx1 leads to reduced size and aberrant gametogenesis. Dev Biol 407: 232–245.

235. Ghiglione, C., N. Perrimon and L. A. Perkins, 1999 Quantitative variations in the level of MAPK activity control patterning of the embryonic termini in Drosophila. Dev Biol 205: 181–193.

236. Ghosh, S., A. Obrdlik, V. Marchand and A. Ephrussi, 2014 The EJC binding and dissociating activity of PYM is regulated in Drosophila. PLoS Genet 10: e1004455.

237. Giagtzoglou, N., P. Alifragis, K. A. Koumbanakis and C. Delidakis, 2003 Two modes of recruitment of E(spl) repressors onto target genes. Development 130: 259–270.

238. Gibson, M. C., and N. Perrimon, 2003 Apicobasal polarization: epithelial form and function. Curr Opin Cell Biol 15: 747–752.

239. Giraldez, A. J., and S. M. Cohen, 2003 Wingless and Notch signaling provide cell survival cues and control cell proliferation during wing development. Development 130: 6533–6543.

240. Glover-Cutter, K., S. Kim, J. Espinosa and D. L. Bentley, 2008 RNA polymerase II pauses and associates with pre-mRNA processing factors at both ends of genes. Nat Struct Mol Biol 15: 71–78.

241. Glover, C. V., E. R. Shelton and D. L. Brutlag, 1983 Purification and characterization of a type II casein kinase from Drosophila melanogaster. J Biol Chem 258: 3258–3265.

242. GO Reference Genome Project, -. 2011 Phylogenetic annotation using the Gene Ontology.

243. Godbout, R., M. Packer and W. Bie, 1998 Overexpression of a DEAD box protein (DDX1) in neuroblastoma and retinoblastoma cell lines. J Biol Chem 273: 21161–21168.

244. Godbout, R., and J. Squire, 1993 Amplification of a DEAD box protein gene in retinoblastoma cell lines. Proc Natl Acad Sci U S A 90: 7578–7582.

245. Goedhart, J., and M. S. Luijsterburg, 2020 VolcaNoseR is a web app for creating, exploring, labeling and sharing volcano plots. Sci Rep 10: 20560.

246. Goldberg, M. L., R. P. Lifton, G. R. Stark and J. G. Williams, 1979 Isolation of specific RNA’s using DNA covalently linked to diazobenzyloxymethyl cellulose or paper. Methods Enzymol 68: 206–220.

247. Gomez-Roman, N., C. Grandori, R. N. Eisenman and R. J. White, 2003 Direct activation of RNA polymerase III transcription by c-Myc. Nature 421: 290–294.

248. Gonzalez-Gaitan, M., and H. Jackle, 1997 Role of Drosophila alpha-adaptin in presynaptic vesicle recycling. Cell 88: 767–776.

249. Gonzalez-Prieto, R., S. A. Cuijpers, R. Kumar, I. A. Hendriks and A. C. Vertegaal, 2015 c-Myc is targeted to the proteasome for degradation in a SUMOylation-dependent manner, regulated by PIAS1, SENP7 and RNF4. Cell Cycle 14: 1859-1872.

250. Goodman, J. K., C. G. Zampronio, A. M. E. Jones and J. R. Hernandez-Fernaud, 2018 Updates of the In-Gel Digestion Method for Protein Analysis by Mass Spectrometry. Proteomics 18: e1800236.

251. Goodman, L. D., M. Prudencio, N. J. Kramer, L. F. Martinez-Ramirez, A. R. Srinivasan et al., 2019 Toxic expanded GGGGCC repeat transcription is mediated by the PAF1 complex in C9orf72-associated FTD. Nat Neurosci 22: 863–874.

252. Gorski, M. M., R. J. Romeijn, J. C. Eeken, A. W. de Jong, B. L. van Veen et al., 2004 Disruption of Drosophila Rad50 causes pupal lethality, the accumulation of DNA double-strand breaks and the induction of apoptosis in third instar larvae. DNA Repair (Amst) 3: 603–615.

253. Goshima, G., R. Wollman, S. S. Goodwin, N. Zhang, J. M. Scholey et al., 2007 Genes required for mitotic spindle assembly in Drosophila S2 cells. Science 316: 417–421.

254. Gouw, A. M., K. Margulis, N. S. Liu, S. J. Raman, A. Mancuso et al., 2019 The MYC Oncogene Cooperates with Sterol-Regulated Element-Binding Protein to Regulate Lipogenesis Essential for Neoplastic Growth. Cell Metab 30: 556–572 e555.

255. Goyal, Y., G. A. Jindal, J. L. Pelliccia, K. Yamaya, E. Yeung et al., 2017 Divergent effects of intrinsically active MEK variants on developmental Ras signaling. Nat Genet 49: 465–469.

256. Grandori, C., S. M. Cowley, L. P. James and R. N. Eisenman, 2000 The Myc/Max/Mad network and the transcriptional control of cell behavior. Annu Rev Cell Dev Biol 16: 653–699.

257. Grandori, C., J. Mac, F. Siebelt, D. E. Ayer and R. N. Eisenman, 1996 Myc-Max heterodimers activate a DEAD box gene and interact with multiple E box-related sites in vivo. EMBO J 15: 4344–4357.

258. Greenleaf, A. L., 1983 Amanitin-resistant RNA polymerase II mutations are in the enzyme’s largest subunit. J Biol Chem 258: 13403–13406.

259. Grewal, S. S., 2015 Why should cancer biologists care about tRNAs? tRNA synthesis, mRNA translation and the control of growth. Biochim Biophys Acta 1849: 898–907.

260. Grewal, S. S., L. Li, A. Orian, R. N. Eisenman and B. A. Edgar, 2005 Myc-dependent regulation of ribosomal RNA synthesis during Drosophila development. Nat Cell Biol 7: 295–302.

261. Grifoni, D., F. Froldi and A. Pession, 2013 Connecting epithelial polarity, proliferation and cancer in Drosophila: the many faces of lgl loss of function. Int J Dev Biol 57: 677–687.

262. Gryzik, T., and H. A. Muller, 2004 FGF8-like1 and FGF8-like2 encode putative ligands of the FGF receptor Htl and are required for mesoderm migration in the Drosophila gastrula. Curr Biol 14: 659–667.

263. Gu, J. Y., J. M. Park, E. J. Song, G. Mizuguchi, J. H. Yoon et al., 2002 Novel Mediator proteins of the small Mediator complex in Drosophila SL2 cells. J Biol Chem 277: 27154–27161.

264. Gu, Z., K. Inomata, K. Ishizawa and A. Horii, 2007 The FBXW7 beta-form is suppressed in human glioma cells. Biochem Biophys Res Commun 354: 992–998.

265. Guelman, S., T. Suganuma, L. Florens, S. K. Swanson, C. L. Kiesecker et al., 2006a Host cell factor and an uncharacterized SANT domain protein are stable components of ATAC, a novel dAda2A/dGcn5-containing histone acetyltransferase complex in Drosophila. Mol Cell Biol 26: 871–882.

266. Guelman, S., T. Suganuma, L. Florens, V. Weake, S. K. Swanson et al., 2006b The essential gene wda encodes a WD40 repeat subunit of Drosophila SAGA required for histone H3 acetylation. Mol Cell Biol 26: 7178–7189.

267. Guo, A., H. Huang, Z. Zhu, M. J. Chen, H. Shi et al., 2022 cBAF complex components and MYC cooperate early in CD8(+) T cell fate. Nature 607: 135–141.

268. Guo, L., O. Zaysteva, Z. Nie, N. C. Mitchell, J. E. Amanda Lee et al., 2016 Defining the essential function of FBP/KSRP proteins: Drosophila Psi interacts with the mediator complex to modulate MYC transcription and tissue growth. Nucleic Acids Res 44: 7646–7658.

269. Haecker, A., D. Qi, T. Lilja, B. Moussian, L. P. Andrioli et al., 2007 Drosophila brakeless interacts with atrophin and is required for tailless-mediated transcriptional repression in early embryos. PLoS Biol 5: e145.

270. Haines, N., and K. D. Irvine, 2003 Glycosylation regulates Notch signalling. Nat Rev Mol Cell Biol 4: 786–797.

271. Hallikas, O., K. Palin, N. Sinjushina, R. Rautiainen, J. Partanen et al., 2006 Genome-wide prediction of mammalian enhancers based on analysis of transcription-factor binding affinity. Cell 124: 47–59.

272. Hamada, F. N., P. J. Park, P. R. Gordadze and M. I. Kuroda, 2005 Global regulation of X chromosomal genes by the MSL complex in Drosophila melanogaster. Genes Dev 19: 2289–2294.

273. Hamiche, A., R. Sandaltzopoulos, D. A. Gdula and C. Wu, 1999 ATP-dependent histone octamer sliding mediated by the chromatin remodeling complex NURF. Cell 97: 833–842.

274. Hamilton, B. J., M. A. Mortin and A. L. Greenleaf, 1993 Reverse genetics of Drosophila RNA polymerase II: identification and characterization of RpII140, the genomic locus for the second-largest subunit. Genetics 134: 517–529.

275. Han, Y., Q. Shi and J. Jiang, 2015 Multisite interaction with Sufu regulates Ci/Gli activity through distinct mechanisms in Hh signal transduction. Proc Natl Acad Sci U S A 112: 6383–6388.

276. Han, Y., B. Wang, Y. S. Cho, J. Zhu, J. Wu et al., 2019 Phosphorylation of Ci/Gli by Fused Family Kinases Promotes Hedgehog Signaling. Dev Cell 50: 610–626 e614.

277. Hansen, S. K., and R. Tjian, 1995 TAFs and TFIIA mediate differential utilization of the tandem Adh promoters. Cell 82: 565–575.

278. Harden, N., 2002 Signaling pathways directing the movement and fusion of epithelial sheets: lessons from dorsal closure in Drosophila. Differentiation 70: 181–203.

279. Harper, S. E., Y. Qiu and P. A. Sharp, 1996 Sin3 corepressor function in Myc-induced transcription and transformation. Proc Natl Acad Sci U S A 93: 8536–8540.

280. Harris, D. A., K. Kim, K. Nakahara, C. Vasquez-Doorman and R. W. Carthew, 2011 Cargo sorting to lysosome-related organelles regulates siRNA-mediated gene silencing. J Cell Biol 194: 77–87.

281. Harris, L. R., M. A. Churchward, R. H. Butt and J. R. Coorssen, 2007 Assessing detection methods for gel-based proteomic analyses. J Proteome Res 6: 1418–1425.

282. Hartl, T. A., J. Ni, J. Cao, K. L. Suyama, S. Patchett et al., 2013 Regulation of ribosome biogenesis by nucleostemin 3 promotes local and systemic growth in Drosophila. Genetics 194: 101–115.

283. Harvey, A. J., A. P. Bidwai and L. K. Miller, 1997 Doom, a product of the Drosophila mod(mdg4) gene, induces apoptosis and binds to baculovirus inhibitor-of-apoptosis proteins. Mol Cell Biol 17: 2835–2843.

284. Hata, T., H. Rajabi, H. Takahashi, Y. Yasumizu, W. Li et al., 2019 MUC1-C Activates the NuRD Complex to Drive Dedifferentiation of Triple-Negative Breast Cancer Cells. Cancer Res 79: 5711–5722.

285. Hatton-Ellis, E., C. Ainsworth, Y. Sushama, S. Wan, K. VijayRaghavan et al., 2007 Genetic regulation of patterned tubular branching in Drosophila. Proc Natl Acad Sci U S A 104: 169–174.

286. Hayashi, R., S. M. Wainwright, S. J. Liddell, S. M. Pinchin, S. Horswell et al., 2014 A genetic screen based on in vivo RNA imaging reveals centrosome-independent mechanisms for localizing gurken transcripts in Drosophila. G3 (Bethesda) 4: 749-760.

287. He, B., A. Martin and E. Wieschaus, 2016 Flow-dependent myosin recruitment during Drosophila cellularization requires zygotic dunk activity. Development 143: 2417–2430.

288. He, L., J. M. Thomson, M. T. Hemann, E. Hernando-Monge, D. Mu et al., 2005 A microRNA polycistron as a potential human oncogene. Nature 435: 828–833.

289. Hebert, M. D., and A. G. Matera, 2000 Self-association of coilin reveals a common theme in nuclear body localization. Mol Biol Cell 11: 4159–4171.

290. Heck, B. W., B. Zhang, X. Tong, Z. Pan, W. M. Deng et al., 2012 The transcriptional corepressor SMRTER influences both Notch and ecdysone signaling during Drosophila development. Biol Open 1: 182–196.

291. Helander, S., M. Montecchio, R. Pilstal, Y. Su, J. Kuruvilla et al., 2015 Pre-Anchoring of Pin1 to Unphosphorylated c-Myc in a Fuzzy Complex Regulates c-Myc Activity. Structure 23: 2267–2279.

292. Helman, A., E. Cinnamon, S. Mezuman, Z. Hayouka, T. Von Ohlen et al., 2011 Phosphorylation of Groucho mediates RTK feedback inhibition and prolonged pathway target gene expression. Curr Biol 21: 1102–1110.

293. Hempel, L. U., C. Rathke, S. J. Raja and R. Renkawitz-Pohl, 2006 In Drosophila, don juan and don juan like encode proteins of the spermatid nucleus and the flagellum and both are regulated at the transcriptional level by the TAF II80 cannonball while translational repression is achieved by distinct elements. Dev Dyn 235: 1053–1064.

294. Henriques, T., B. S. Scruggs, M. O. Inouye, G. W. Muse, L. H. Williams et al., 2018 Widespread transcriptional pausing and elongation control at enhancers. Genes Dev 32: 26–41.

295. Herman-Bachinsky, Y., H. D. Ryoo, A. Ciechanover and H. Gonen, 2007 Regulation of the Drosophila ubiquitin ligase DIAP1 is mediated via several distinct ubiquitin system pathways. Cell Death Differ 14: 861–871.

296. Hernandez, G., P. Vazquez-Pianzola, A. Zurbriggen, M. Altmann, J. M. Sierra et al., 2004 Two functionally redundant isoforms of Drosophila melanogaster eukaryotic initiation factor 4B are involved in cap-dependent translation, cell survival, and proliferation. Eur J Biochem 271: 2923–2936.

297. Herold, N., C. L. Will, E. Wolf, B. Kastner, H. Urlaub et al., 2009 Conservation of the protein composition and electron microscopy structure of Drosophila melanogaster and human spliceosomal complexes. Mol Cell Biol 29: 281–301.

298. Herranz, H., L. Perez, F. A. Martin and M. Milan, 2008 A Wingless and Notch double-repression mechanism regulates G1-S transition in the Drosophila wing. EMBO J 27: 1633–1645.

299. Herrera, F. J., T. Yamaguchi, H. Roelink and R. Tjian, 2014 Core promoter factor TAF9B regulates neuronal gene expression. Elife 3: e02559.

300. Hirling, H., M. Scheffner, T. Restle and H. Stahl, 1989 RNA helicase activity associated with the human p68 protein. Nature 339: 562–564.

301. Hochheimer, A., S. Zhou, S. Zheng, M. C. Holmes and R. Tjian, 2002 TRF2 associates with DREF and directs promoter-selective gene expression in Drosophila. Nature 420: 439–445.

302. Hogarty, M. D., M. D. Norris, K. Davis, X. Liu, N. F. Evageliou et al., 2008 ODC1 is a critical determinant of MYCN oncogenesis and a therapeutic target in neuroblastoma. Cancer Res 68: 9735–9745.

303. Hsieh, A. L., X. Zheng, Z. Yue, Z. E. Stine, A. Mancuso et al., 2019 Misregulation of Drosophila Myc Disrupts Circadian Behavior and Metabolism. Cell Rep 29: 1778–1788 e1774.

304. Hsouna, A., G. Nallamothu, N. Kose, M. Guinea, V. Dammai et al., 2010 Drosophila von Hippel-Lindau tumor suppressor gene function in epithelial tubule morphogenesis. Mol Cell Biol 30: 3779–3794.

305. Hsu, J. Y., T. Juven-Gershon, M. T. Marr, 2nd, K. J. Wright, R. Tjian et al., 2008 TBP, Mot1, and NC2 establish a regulatory circuit that controls DPE-dependent versus TATA-dependent transcription. Genes Dev 22: 2353-2358.

306. Hsu, T., D. McRackan, T. S. Vincent and H. Gert de Couet, 2001 Drosophila Pin1 prolyl isomerase Dodo is a MAP kinase signal responder during oogenesis. Nat Cell Biol 3: 538–543.

307. Huang, C., F. Yang, Z. Zhang, J. Zhang, G. Cai et al., 2017 Mrg15 stimulates Ash1 H3K36 methyltransferase activity and facilitates Ash1 Trithorax group protein function in Drosophila. Nat Commun 8: 1649.

308. Huijbregts, R. P., A. Svitin, M. W. Stinnett, M. B. Renfrow and I. Chesnokov, 2009 Drosophila Orc6 facilitates GTPase activity and filament formation of the septin complex. Mol Biol Cell 20: 270–281.

309. Hyun, J., H. Jasper and D. Bohmann, 2005 DREF is required for efficient growth and cell cycle progression in Drosophila imaginal discs. Mol Cell Biol 25: 5590–5598.

310. Ibrahim, D., L. Prevaud, N. Faumont, D. Troutaud, J. Feuillard et al., 2022 Alternative c-MYC mRNA Transcripts as an Additional Tool for c-Myc2 and c-MycS Production in BL60 Tumors. Biomolecules 12.

311. Ichikawa, K., Y. Kubota, T. Nakamura, J. S. Weng, T. Tomida et al., 2015 MCRIP1, an ERK substrate, mediates ERK-induced gene silencing during epithelial-mesenchymal transition by regulating the co-repressor CtBP. Mol Cell 58: 35–46.

312. Ignesti, M., M. Barraco, G. Nallamothu, J. A. Woolworth, S. Duchi et al., 2014 Notch signaling during development requires the function of awd, the Drosophila homolog of human metastasis suppressor gene Nm23. BMC Biol 12: 12.

313. Ihry, R. J., and A. Bashirullah, 2014 Genetic control of specificity to steroid-triggered responses in Drosophila. Genetics 196: 767–780.

314. Ilyin, A. A., S. S. Ryazansky, S. A. Doronin, O. M. Olenkina, E. A. Mikhaleva et al., 2017 Piwi interacts with chromatin at nuclear pores and promiscuously binds nuclear transcripts in Drosophila ovarian somatic cells. Nucleic Acids Res 45: 7666–7680.

315. Immanuel, D., H. Zinszner and D. Ron, 1995 Association of SARFH (sarcoma-associated RNA-binding fly homolog) with regions of chromatin transcribed by RNA polymerase II. Mol Cell Biol 15: 4562–4571.

316. Inoue, H., T. Imamura, Y. Ishidou, M. Takase, Y. Udagawa et al., 1998 Interplay of signal mediators of decapentaplegic (Dpp): molecular characterization of mothers against dpp, Medea, and daughters against dpp. Mol Biol Cell 9: 2145–2156.

317. InterPro Project Members, -. 2004 Gene Ontology annotation through association of InterPro records with GO terms.

318. Irion, U., and M. Leptin, 1999 Developmental and cell biological functions of the Drosophila DEAD-box protein abstrakt. Curr Biol 9: 1373–1381.

319. Irion, U., M. Leptin, K. Siller, S. Fuerstenberg, Y. Cai et al., 2004 Abstrakt, a DEAD box protein, regulates Insc levels and asymmetric division of neural and mesodermal progenitors. Curr Biol 14: 138–144.

320. Isabella, A. J., and S. Horne-Badovinac, 2016 Rab10-Mediated Secretion Synergizes with Tissue Movement to Build a Polarized Basement Membrane Architecture for Organ Morphogenesis. Dev Cell 38: 47–60.

321. Ishigaki, Y., Y. Nakamura, T. Tatsuno, M. Hashimoto, K. Iwabuchi et al., 2014 RNA-binding protein RBM8A (Y14) and MAGOH localize to centrosome in human A549 cells. Histochem Cell Biol 141: 101–109.

322. Israeli, D., R. Nir and T. Volk, 2007 Dissection of the target specificity of the RNA-binding protein HOW reveals dpp mRNA as a novel HOW target. Development 134: 2107–2114.

323. Itkonen, H. M., S. Minner, I. J. Guldvik, M. J. Sandmann, M. C. Tsourlakis et al., 2013 O-GlcNAc transferase integrates metabolic pathways to regulate the stability of c-MYC in human prostate cancer cells. Cancer Res 73: 5277–5287.

324. Itkonen, H. M., A. Urbanucci, S. E. Martin, A. Khan, A. Mathelier et al., 2019 High OGT activity is essential for MYC-driven proliferation of prostate cancer cells. Theranostics 9: 2183–2197.

325. Ito, T., M. Bulger, M. J. Pazin, R. Kobayashi and J. T. Kadonaga, 1997 ACF, an ISWI-containing and ATP-utilizing chromatin assembly and remodeling factor. Cell 90: 145–155.

326. Ito, T., M. E. Levenstein, D. V. Fyodorov, A. K. Kutach, R. Kobayashi et al., 1999 ACF consists of two subunits, Acf1 and ISWI, that function cooperatively in the ATP-dependent catalysis of chromatin assembly. Genes Dev 13: 1529-1539.

327. Ito, Y., A. Asada, H. Kobayashi, T. Takano, G. Sharma et al., 2014 Preferential targeting of p39-activated Cdk5 to Rac1-induced lamellipodia. Mol Cell Neurosci 61: 34–45.

328. Jablonowski, C., W. Quarni, S. Singh, H. Tan, D. H. Bostanthirige et al., 2023 Metabolic reprogramming of cancer cells by JMJD6-mediated pre-mRNA splicing is associated with therapeutic response to splicing inhibitor. bioRxiv.

329. Jackstadt, R., S. Roh, J. Neumann, P. Jung, R. Hoffmann et al., 2013 AP4 is a mediator of epithelial-mesenchymal transition and metastasis in colorectal cancer. J Exp Med 210: 1331–1350.

330. Janody, F., Z. Martirosyan, A. Benlali and J. E. Treisman, 2003 Two subunits of the Drosophila mediator complex act together to control cell affinity. Development 130: 3691–3701.

331. Janssens, D. H., H. Komori, D. Grbac, K. Chen, C. T. Koe et al., 2014 Earmuff restricts progenitor cell potential by attenuating the competence to respond to self-renewal factors. Development 141: 1036–1046.

332. Jasper, H., V. Benes, A. Atzberger, S. Sauer, W. Ansorge et al., 2002 A genomic switch at the transition from cell proliferation to terminal differentiation in the Drosophila eye. Dev Cell 3: 511–521.

333. Jasper, H., and D. Bohmann, 2002 Drosophila innate immunity: a genomic view of pathogen defense. Mol Cell 10: 967–969.

334. JayaNandanan, N., R. Mathew and M. Leptin, 2014 Guidance of subcellular tubulogenesis by actin under the control of a synaptotagmin-like protein and Moesin. Nat Commun 5: 3036.

335. Jeske, M., S. Meyer, C. Temme, D. Freudenreich and E. Wahle, 2006 Rapid ATP-dependent deadenylation of nanos mRNA in a cell-free system from Drosophila embryos. J Biol Chem 281: 25124–25133.

336. Jewett, J. C., and C. R. Bertozzi, 2010 Cu-free click cycloaddition reactions in chemical biology. Chem Soc Rev 39: 1272–1279.

337. Ji, J., X. Tang, W. Hu, K. A. Maggert and Y. S. Rong, 2019 The processivity factor Pol32 mediates nuclear localization of DNA polymerase delta and prevents chromosomal fragile site formation in Drosophila development. PLoS Genet 15: e1008169.

338. Jia, D., M. Soylemez, G. Calvin, R. Bornmann, J. Bryant et al., 2015 A large-scale in vivo RNAi screen to identify genes involved in Notch-mediated follicle cell differentiation and cell cycle switches. Sci Rep 5: 12328.

339. Jia, H., Y. Liu, R. Xia, C. Tong, T. Yue et al., 2010 Casein kinase 2 promotes Hedgehog signaling by regulating both smoothened and Cubitus interruptus. J Biol Chem 285: 37218–37226.

340. Jimeno-Martin, A., E. Sousa, R. Brocal-Ruiz, N. Daroqui, M. Maicas et al., 2022 Joint actions of diverse transcription factor families establish neuron-type identities and promote enhancer selectivity. Genome Res 32: 459–473.

341. Jin, Y., J. Shi, H. Wang, J. Lu, C. Chen et al., 2021 MYC-associated protein X binding with the variant rs72780850 in RNA helicase DEAD box 1 for susceptibility to neuroblastoma. Sci China Life Sci 64: 991–999.

342. Johnston, L. A., D. A. Prober, B. A. Edgar, R. N. Eisenman and P. Gallant, 1999 Drosophila myc regulates cellular growth during development. Cell 98: 779–790.

343. Jung, P., A. Menssen, D. Mayr and H. Hermeking, 2008 AP4 encodes a c-MYC-inducible repressor of p21. Proc Natl Acad Sci U S A 105: 15046–15051.

344. Kadamb, R., S. Mittal, N. Bansal, H. Batra and D. Saluja, 2013 Sin3: insight into its transcription regulatory functions. Eur J Cell Biol 92: 237–246.

345. Kadonaga, J. T., 2002 The DPE, a core promoter element for transcription by RNA polymerase II. Exp Mol Med 34: 259–264.

346. Kadonaga, J. T., 2012 Perspectives on the RNA polymerase II core promoter. Wiley Interdiscip Rev Dev Biol 1: 40–51.

347. Kadrmas, J. L., M. A. Smith, K. A. Clark, S. M. Pronovost, N. Muster et al., 2004 The integrin effector PINCH regulates JNK activity and epithelial migration in concert with Ras suppressor 1. J Cell Biol 167: 1019–1024.

348. Kaguni, L. S., J. M. Rossignol, R. C. Conaway, G. R. Banks and I. R. Lehman, 1983 Association of DNA primase with the beta/gamma subunits of DNA polymerase alpha from Drosophila melanogaster embryos. J Biol Chem 258: 9037–9039.

349. Kakumani, P. K., L. M. Harvey, F. Houle, T. Guitart, F. Gebauer et al., 2020 CSDE1 controls gene expression through the miRNA-mediated decay machinery. Life Sci Alliance 3.

350. Kal, A. J., T. Mahmoudi, N. B. Zak and C. P. Verrijzer, 2000 The Drosophila brahma complex is an essential coactivator for the trithorax group protein zeste. Genes Dev 14: 1058–1071.

351. Kalverda, B., H. Pickersgill, V. V. Shloma and M. Fornerod, 2010 Nucleoporins directly stimulate expression of developmental and cell-cycle genes inside the nucleoplasm. Cell 140: 360–371.

352. Kaminker, J. S., J. Canon, I. Salecker and U. Banerjee, 2002 Control of photoreceptor axon target choice by transcriptional repression of Runt. Nat Neurosci 5: 746–750.

353. Kang, M. J., J. Chung and H. D. Ryoo, 2012 CDK5 and MEKK1 mediate pro-apoptotic signalling following endoplasmic reticulum stress in an autosomal dominant retinitis pigmentosa model. Nat Cell Biol 14: 409–415.

354. Kanoh, H., L. L. Tong, T. Kuraishi, Y. Suda, Y. Momiuchi et al., 2015 Genome-wide RNAi screening implicates the E3 ubiquitin ligase Sherpa in mediating innate immune signaling by Toll in Drosophila adults. Sci Signal 8: ra107.

355. Kara, E., A. McCambridge, M. Proffer, C. Dilts, B. Pumnea et al., 2023 Mutational analysis of the functional motifs of the DEAD-box RNA helicase Me31B/DDX6 in Drosophila germline development. FEBS Lett 597: 1848–1867.

356. Karandikar, U. C., R. L. Trott, J. Yin, C. P. Bishop and A. P. Bidwai, 2004 Drosophila CK2 regulates eye morphogenesis via phosphorylation of E(spl)M8. Mech Dev 121: 273–286.

357. Karim, F. D., H. C. Chang, M. Therrien, D. A. Wassarman, T. Laverty et al., 1996 A screen for genes that function downstream of Ras1 during Drosophila eye development. Genetics 143: 315–329.

358. Karim, F. D., and G. M. Rubin, 1999 PTP-ER, a novel tyrosine phosphatase, functions downstream of Ras1 to downregulate MAP kinase during Drosophila eye development. Mol Cell 3: 741–750.

359. Katoh, M., M. Igarashi, H. Fukuda, H. Nakagama and M. Katoh, 2013 Cancer genetics and genomics of human FOX family genes. Cancer Lett 328: 198–206.

360. Kedmi, A., Y. Zehavi, Y. Glick, Y. Orenstein, D. Ideses et al., 2014 Drosophila TRF2 is a preferential core promoter regulator. Genes Dev 28: 2163–2174.

361. Kelleher, F. C., A. Rao and A. Maguire, 2014 Circadian molecular clocks and cancer. Cancer Lett 342: 9–18.

362. Kenney, A. M., M. D. Cole and D. H. Rowitch, 2003 Nmyc upregulation by sonic hedgehog signaling promotes proliferation in developing cerebellar granule neuron precursors. Development 130: 15–28.

363. Kharazmi, J., and C. Moshfegh, 2013 Investigation of dmyc Promoter and Regulatory Regions. Gene Regul Syst Bio 7: 85–102.

364. Kharazmi, J., C. Moshfegh and T. Brody, 2012 Identification of cis-Regulatory Elements in the dmyc Gene of Drosophila Melanogaster. Gene Regul Syst Bio 6: 15–42.

365. Khoury, M. J., and D. Bilder, 2020 Distinct activities of Scrib module proteins organize epithelial polarity. Proc Natl Acad Sci U S A 117: 11531–11540.

366. Kiessling, A. A., R. Bletsa, B. Desmarais, C. Mara, K. Kallianidis et al., 2009 Evidence that human blastomere cleavage is under unique cell cycle control. J Assist Reprod Genet 26: 187–195.

367. Kim, C. A., and J. U. Bowie, 2003 SAM domains: uniform structure, diversity of function. Trends Biochem Sci 28: 625–628.

368. Kim, S. H., E. R. Park, E. Cho, W. H. Jung, J. Y. Jeon et al., 2017 Mael is essential for cancer cell survival and tumorigenesis through protection of genetic integrity. Oncotarget 8: 5026–5037.

369. Kim, S. N., A. Jeibmann, K. Halama, H. T. Witte, M. Walte et al., 2014 ECM stiffness regulates glial migration in Drosophila and mammalian glioma models. Development 141: 3233–3242.

370. Kim, S. Y., J. Y. Kim, S. Malik, W. Son, K. S. Kwon et al., 2012 Negative regulation of EGFR/MAPK pathway by Pumilio in Drosophila melanogaster. PLoS One 7: e34016.

371. Kim, Y. O., S. J. Park, R. S. Balaban, M. Nirenberg and Y. Kim, 2004 A functional genomic screen for cardiogenic genes using RNA interference in developing Drosophila embryos. Proc Natl Acad Sci U S A 101: 159–164.

372. King-Jones, K., G. Korge and M. Lehmann, 1999 The helix-loop-helix proteins dAP-4 and daughterless bind both in vitro and in vivo to SEBP3 sites required for transcriptional activation of the Drosophila gene Sgs-4. J Mol Biol 291: 71–82.

373. King, M. W., J. M. Roberts and R. N. Eisenman, 1986 Expression of the c-myc proto-oncogene during development of Xenopus laevis. Mol Cell Biol 6: 4499–4508.

374. Kirilly, D., E. P. Spana, N. Perrimon, R. W. Padgett and T. Xie, 2005 BMP signaling is required for controlling somatic stem cell self-renewal in the Drosophila ovary. Dev Cell 9: 651–662.

375. Kiselev, A. M., I. S. Stepanova, L. S. Adonin, F. M. Batalova, V. N. Parfenov et al., 2017 The exon junction complex factor Y14 is dynamic in the nucleus of the beetle Tribolium castaneum during late oogenesis. Mol Cytogenet 10: 41.

376. Kitadate, Y., S. Shigenobu, K. Arita and S. Kobayashi, 2007 Boss/Sev signaling from germline to soma restricts germline-stem-cell-niche formation in the anterior region of Drosophila male gonads. Dev Cell 13: 151–159.

377. Klymenko, T., B. Papp, W. Fischle, T. Kocher, M. Schelder et al., 2006 A Polycomb group protein complex with sequence-specific DNA-binding and selective methyl-lysine-binding activities. Genes Dev 20: 1110–1122.

378. Knoepfler, P. S., X. Y. Zhang, P. F. Cheng, P. R. Gafken, S. B. McMahon et al., 2006 Myc influences global chromatin structure. EMBO J 25: 2723–2734.

379. Koch, F., R. Fenouil, M. Gut, P. Cauchy, T. K. Albert et al., 2011 Transcription initiation platforms and GTF recruitment at tissue-specific enhancers and promoters. Nat Struct Mol Biol 18: 956–963.

380. Koe, C. T., S. Li, F. Rossi, J. J. Wong, Y. Wang et al., 2014 The Brm-HDAC3-Erm repressor complex suppresses dedifferentiation in Drosophila type II neuroblast lineages. Elife 3: e01906.

381. Koelzer, S., and T. Klein, 2006 Regulation of expression of Vg and establishment of the dorsoventral compartment boundary in the wing imaginal disc by Suppressor of Hairless. Dev Biol 289: 77–90.

382. Kogan, G. L., N. V. Akulenko, Y. A. Abramov, O. A. Sokolova, E. A. Fefelova et al., 2017 [Nascent Polypeptide-Associated Complex as Tissue-Specific Cofactor during Germinal Cell Differentiation in Drosophila Testes]. Mol Biol (Mosk) 51: 677–682.

383. Kogan, G. L., E. A. Mikhaleva, O. M. Olenkina, S. S. Ryazansky, O. V. Galzitskaya et al., 2022 Extended disordered regions of ribosome-associated NAC proteins paralogs belong only to the germline in Drosophila melanogaster. Sci Rep 12: 11191.

384. Koh, C. M., M. Bezzi, D. H. Low, W. X. Ang, S. X. Teo et al., 2015 MYC regulates the core pre-mRNA splicing machinery as an essential step in lymphomagenesis. Nature 523: 96–100.

385. Kohwi-Shigematsu, T., K. Maass and J. Bode, 1997 A thymocyte factor SATB1 suppresses transcription of stably integrated matrix-attachment region-linked reporter genes. Biochemistry 36: 12005–12010.

386. Koizumi, K., H. Higashida, S. Yoo, M. S. Islam, A. I. Ivanov et al., 2007 RNA interference screen to identify genes required for Drosophila embryonic nervous system development. Proc Natl Acad Sci U S A 104: 5626–5631.

387. Kon, C., K. M. Cadigan, S. L. da Silva and R. Nusse, 2005 Developmental roles of the Mi-2/NURD-associated protein p66 in Drosophila. Genetics 169: 2087–2100.

388. Konig, A., and H. R. Shcherbata, 2015 Soma influences GSC progeny differentiation via the cell adhesion-mediated steroid-let-7-Wingless signaling cascade that regulates chromatin dynamics. Biol Open 4: 285–300.

389. Kopecky, B., P. Santi, S. Johnson, H. Schmitz and B. Fritzsch, 2011 Conditional deletion of N-Myc disrupts neurosensory and non-sensory development of the ear. Dev Dyn 240: 1373–1390.

390. Koppen, M., B. G. Fernandez, L. Carvalho, A. Jacinto and C. P. Heisenberg, 2006 Coordinated cell-shape changes control epithelial movement in zebrafish and Drosophila. Development 133: 2671–2681.

391. Kozlova, T., and C. S. Thummel, 2000 Steroid regulation of postembryonic development and reproduction in Drosophila. Trends Endocrinol Metab 11: 276–280.

392. Krenciute, G., S. Liu, N. Yucer, Y. Shi, P. Ortiz et al., 2013 Nuclear BAG6-UBL4A-GET4 complex mediates DNA damage signaling and cell death. J Biol Chem 288: 20547–20557.

393. Kress, T. R., A. Sabo and B. Amati, 2015 MYC: connecting selective transcriptional control to global RNA production. Nat Rev Cancer 15: 593–607.

394. Kristo, I., C. Bajusz, B. N. Borsos, T. Pankotai, J. Dopie et al., 2017 The actin binding cytoskeletal protein Moesin is involved in nuclear mRNA export. Biochim Biophys Acta Mol Cell Res 1864: 1589–1604.

395. Kroiss, M., J. Schultz, J. Wiesner, A. Chari, A. Sickmann et al., 2008 Evolution of an RNP assembly system: a minimal SMN complex facilitates formation of UsnRNPs in Drosophila melanogaster. Proc Natl Acad Sci U S A 105: 10045–10050.

396. Kuan, Y. S., P. Brewer-Jensen, W. L. Bai, C. Hunter, C. B. Wilson et al., 2009 Drosophila suppressor of sable protein [Su(s)] promotes degradation of aberrant and transposon-derived RNAs. Mol Cell Biol 29: 5590–5603.

397. Kuan, Y. S., P. Brewer-Jensen and L. L. Searles, 2004 Suppressor of sable, a putative RNA-processing protein, functions at the level of transcription. Mol Cell Biol 24: 3734–3746.

398. Kubickova, A., J. B. De Sanctis and M. Hajduch, 2023 Isoform-Directed Control of c-Myc Functions: Understanding the Balance from Proliferation to Growth Arrest. Int J Mol Sci 24.

399. Kugler, S. J., E. M. Gehring, V. Wallkamm, V. Kruger and A. C. Nagel, 2011 The Putzig-NURF nucleosome remodeling complex is required for ecdysone receptor signaling and innate immunity in Drosophila melanogaster. Genetics 188: 127–139.

400. Kugler, S. J., and A. C. Nagel, 2007 putzig is required for cell proliferation and regulates notch activity in Drosophila. Mol Biol Cell 18: 3733–3740.

401. Kumar, J. P., M. Tio, F. Hsiung, S. Akopyan, L. Gabay et al., 1998 Dissecting the roles of the Drosophila EGF receptor in eye development and MAP kinase activation. Development 125: 3875–3885.

402. Kuroda, K., R. Kagiyama-Takahashi and T. Shinomiya, 1990 Immunoaffinity purification and properties of Drosophila melanogaster DNA polymerase alpha-primase complex. J Biochem 108: 926–933.

403. Kusch, T., L. Florens, W. H. Macdonald, S. K. Swanson, R. L. Glaser et al., 2004 Acetylation by Tip60 is required for selective histone variant exchange at DNA lesions. Science 306: 2084–2087.

404. Kutach, A. K., and J. T. Kadonaga, 2000 The downstream promoter element DPE appears to be as widely used as the TATA box in Drosophila core promoters. Mol Cell Biol 20: 4754–4764.

405. Lacoste, J., S. Codani-Simonart, M. Best-Belpomme and F. Peronnet, 1995 Characterization and cloning of p11, a transrepressor of Drosophila melanogaster retrotransposon 1731. Nucleic Acids Res 23: 5073–5079.

406. LaJeunesse, D., and A. Shearn, 1995 Trans-regulation of thoracic homeotic selector genes of the Antennapedia and bithorax complexes by the trithorax group genes: absent, small, and homeotic discs 1 and 2. Mech Dev 53: 123–139.

407. Lambertsson, A., 1998 The minute genes in Drosophila and their molecular functions. Adv Genet 38: 69–134.

408. Lancaster, O. M., C. F. Cullen and H. Ohkura, 2007 NHK-1 phosphorylates BAF to allow karyosome formation in the Drosophila oocyte nucleus. J Cell Biol 179: 817–824.

409. Langst, G., and P. B. Becker, 2001 ISWI induces nucleosome sliding on nicked DNA. Mol Cell 8: 1085–1092.

410. Laranjeiro, R., T. K. Tamai, E. Peyric, P. Krusche, S. Ott et al., 2013 Cyclin-dependent kinase inhibitor p20 controls circadian cell-cycle timing. Proc Natl Acad Sci U S A 110: 6835–6840.

411. Larochelle, S., J. Chen, R. Knights, J. Pandur, P. Morcillo et al., 2001 T-loop phosphorylation stabilizes the CDK7-cyclin H-MAT1 complex in vivo and regulates its CTD kinase activity. EMBO J 20: 3749–3759.

412. Lascu, I., A. Chaffotte, B. Limbourg-Bouchon and M. Veron, 1992 A Pro/Ser substitution in nucleoside diphosphate kinase of Drosophila melanogaster (mutation killer of prune) affects stability but not catalytic efficiency of the enzyme. J Biol Chem 267: 12775–12781.

413. Lasko, P., 2000 The drosophila melanogaster genome: translation factors and RNA binding proteins. J Cell Biol 150: F51–56.

414. Lasko, P., 2003 Ribosomes rule: translation, not transcription, is the primary target of two major intercellular signaling pathways. Dev Cell 5: 671–672.

415. Lasko, P., 2013 The DEAD-box helicase Vasa: evidence for a multiplicity of functions in RNA processes and developmental biology. Biochim Biophys Acta 1829: 810–816.

416. Lavoie, C. A., P. E. Lachance, N. Sonenberg and P. Lasko, 1996 Alternatively spliced transcripts from the Drosophila eIF4E gene produce two different Cap-binding proteins. J Biol Chem 271: 16393–16398.

417. Le Hir, H., and G. R. Andersen, 2008 Structural insights into the exon junction complex. Curr Opin Struct Biol 18: 112–119.

418. Lecuit, T., W. J. Brook, M. Ng, M. Calleja, H. Sun et al., 1996 Two distinct mechanisms for long-range patterning by Decapentaplegic in the Drosophila wing. Nature 381: 387–393.

419. Lee, J. E., N. J. Cranna, A. S. Chahal and L. M. Quinn, 2012 Genetic systems to investigate regulation of oncogenes and tumour suppressor genes in Drosophila. Cells 1: 1182–1196.

420. Lee, L. A., and T. L. Orr-Weaver, 2003 Regulation of cell cycles in Drosophila development: intrinsic and extrinsic cues. Annu Rev Genet 37: 545–578.

421. Lee, M. P., M. Sander and T. S. Hsieh, 1989 Single strand DNA cleavage reaction of duplex DNA by Drosophila topoisomerase II. J Biol Chem 264: 13510–13518.

422. Lehner, C. F., and P. H. O’Farrell, 1990 The roles of Drosophila cyclins A and B in mitotic control. Cell 61: 535–547.

423. Lei, J. X., Q. Y. Liu, C. Sodja, J. LeBlanc, M. Ribecco-Lutkiewicz et al., 2005 S/MAR-binding properties of Sox2 and its involvement in apoptosis of human NT2 neural precursors. Cell Death Differ 12: 1368–1377.

424. Leitner, A., T. Walzthoeni, A. Kahraman, F. Herzog, O. Rinner et al., 2010 Probing native protein structures by chemical cross-linking, mass spectrometry, and bioinformatics. Mol Cell Proteomics 9: 1634–1649.

425. Levanon, D., R. E. Goldstein, Y. Bernstein, H. Tang, D. Goldenberg et al., 1998 Transcriptional repression by AML1 and LEF-1 is mediated by the TLE/Groucho corepressors. Proc Natl Acad Sci U S A 95: 11590–11595.

426. Levine, J. D., C. I. Casey, D. D. Kalderon and F. R. Jackson, 1994 Altered circadian pacemaker functions and cyclic AMP rhythms in the Drosophila learning mutant dunce. Neuron 13: 967–974.

427. Levine, M., 2010 Transcriptional enhancers in animal development and evolution. Curr Biol 20: R754–763.

428. Li, C., J. Lv, G. Wumaier, Y. Zhao, L. Dong et al., 2023 NDRG1 promotes endothelial dysfunction and hypoxia-induced pulmonary hypertension by targeting TAF15. Precis Clin Med 6: pbad024.

429. Li, L., S. Anderson, J. Secombe and R. N. Eisenman, 2013 The Drosophila ubiquitin-specific protease Puffyeye regulates dMyc-mediated growth. Development 140: 4776–4787.

430. Li, L., P. Williams, Z. Gao and Y. Wang, 2020 VEZF1-guanine quadruplex DNA interaction regulates alternative polyadenylation and detyrosinase activity of VASH1. Nucleic Acids Res 48: 11994–12003.

431. Li, L. Y., J. F. Yang, F. Rong, Z. P. Luo, S. Hu et al., 2021 ZEB1 serves an oncogenic role in the tumourigenesis of HCC by promoting cell proliferation, migration, and inhibiting apoptosis via Wnt/beta-catenin signaling pathway. Acta Pharmacol Sin 42: 1676–1689.

432. Li, Q., S. Li, S. Mana-Capelli, R. J. Roth Flach, L. V. Danai et al., 2014 The conserved misshapen-warts-Yorkie pathway acts in enteroblasts to regulate intestinal stem cells in Drosophila. Dev Cell 31: 291–304.

433. Li, S., Y. S. Cho, T. Yue, Y. T. Ip and J. Jiang, 2015 Overlapping functions of the MAP4K family kinases Hppy and Msn in Hippo signaling. Cell Discov 1: 15038.

434. Li, W., E. M. Skoulakis, R. L. Davis and N. Perrimon, 1997 The Drosophila 14-3-3 protein Leonardo enhances Torso signaling through D-Raf in a Ras 1-dependent manner. Development 124: 4163–4171.

435. Liang, H. L., C. Y. Nien, H. Y. Liu, M. M. Metzstein, N. Kirov et al., 2008 The zinc-finger protein Zelda is a key activator of the early zygotic genome in Drosophila. Nature 456: 400–403.

436. Lim, A. K., and T. Kai, 2007 Unique germ-line organelle, nuage, functions to repress selfish genetic elements in Drosophila melanogaster. Proc Natl Acad Sci U S A 104: 6714–6719.

437. Lim, Y. M., S. Hayashi and L. Tsuda, 2012 Ebi/AP-1 suppresses pro-apoptotic genes expression and permits long-term survival of Drosophila sensory neurons. PLoS One 7: e37028.

438. Lim, Y. M., K. Nishizawa, Y. Nishi, L. Tsuda, Y. H. Inoue et al., 1999 Genetic analysis of rolled, which encodes a Drosophila mitogen-activated protein kinase. Genetics 153: 763–771.

439. Lin, Y. H., H. Currinn, S. M. Pocha, A. Rothnie, T. Wassmer et al., 2015 AP-2-complex-mediated endocytosis of Drosophila Crumbs regulates polarity by antagonizing Stardust. J Cell Sci 128: 4538–4549.

440. Lindhorst, D., and M. S. Halfon, 2023 Reporter gene assays and chromatin-level assays define substantially non-overlapping sets of enhancer sequences. BMC Genomics 24: 17.

441. Lindner, J. R., P. R. Hillman, A. L. Barrett, M. C. Jackson, T. L. Perry et al., 2007 The Drosophila Perlecan gene trol regulates multiple signaling pathways in different developmental contexts. BMC Dev Biol 7: 121.

442. Liu, H., X. Tang, A. Srivastava, T. Pecot, P. Daniel et al., 2015 Redeployment of Myc and E2f1-3 drives Rb-deficient cell cycles. Nat Cell Biol 17: 1036–1048.

443. Liu, J., F. Kouzine, Z. Nie, H. J. Chung, Z. Elisha-Feil et al., 2006 The FUSE/FBP/FIR/TFIIH system is a molecular machine programming a pulse of c-myc expression. EMBO J 25: 2119–2130.

444. Liu, N., T. Konuma, R. Sharma, D. Wang, N. Zhao et al., 2023a Histone H3 lysine 27 crotonylation mediates gene transcriptional repression in chromatin. Mol Cell 83: 2206–2221 e2211.

445. Liu, S., G. H. Baeg, Y. Yang, F. G. Goh, H. Bao et al., 2023b The Integrator complex desensitizes cellular response to TGF-beta/BMP signaling. Cell Rep 42: 112007.

446. Liu, X., and J. Secombe, 2015 The Histone Demethylase KDM5 Activates Gene Expression by Recognizing Chromatin Context through Its PHD Reader Motif. Cell Rep 13: 2219–2231.

447. Liu, Y., Y. Hu, X. Li, L. Niu and M. Teng, 2010 The crystal structure of the human nascent polypeptide-associated complex domain reveals a nucleic acid-binding region on the NACA subunit. Biochemistry 49: 2890–2896.

448. Liu, Y., J. Mattila and V. Hietakangas, 2020 Systematic Screen for Drosophila Transcriptional Regulators Phosphorylated in Response to Insulin/mTOR Pathway. G3 (Bethesda) 10: 2843-2849.

449. Liu, Y., J. Mattila, S. Ventela, L. Yadav, W. Zhang et al., 2017 PWP1 Mediates Nutrient-Dependent Growth Control through Nucleolar Regulation of Ribosomal Gene Expression. Dev Cell 43: 240–252 e245.

450. Liu, Y. I., M. V. Chang, H. E. Li, S. Barolo, J. L. Chang et al., 2008 The chromatin remodelers ISWI and ACF1 directly repress Wingless transcriptional targets. Dev Biol 323: 41–52.

451. Lloret-Llinares, M., C. Carre, A. Vaquero, N. de Olano and F. Azorin, 2008 Characterization of Drosophila melanogaster JmjC+N histone demethylases. Nucleic Acids Res 36: 2852–2863.

452. Lobbardi, R., G. Lambert, J. Zhao, R. Geisler, H. R. Kim et al., 2011 Fine-tuning of Hh signaling by the RNA-binding protein Quaking to control muscle development. Development 138: 1783–1794.

453. Loncle, N., M. Boube, L. Joulia, C. Boschiero, M. Werner et al., 2007 Distinct roles for Mediator Cdk8 module subunits in Drosophila development. EMBO J 26: 1045–1054.

454. Loo, L. W., J. Secombe, J. T. Little, L. S. Carlos, C. Yost et al., 2005 The transcriptional repressor dMnt is a regulator of growth in Drosophila melanogaster. Mol Cell Biol 25: 7078–7091.

455. Lu, J., Y. Wang, X. Wang, D. Wang, G. O. Pflugfelder et al., 2022 The Tbx6 Transcription Factor Dorsocross Mediates Dpp Signaling to Regulate Drosophila Thorax Closure. Int J Mol Sci 23.

456. Lu, Y., L. Yang, M. Shen, Z. Zhang, S. Wang et al., 2023 Tespa1 facilitates hematopoietic and leukemic stem cell maintenance by restricting c-Myc degradation. Leukemia 37: 1039–1047.

457. Luanpitpong, S., J. Poohadsuan, P. Klaihmon, X. Kang, K. Tangkiettrakul et al., 2021 Metabolic sensor O-GlcNAcylation regulates megakaryopoiesis and thrombopoiesis through c-Myc stabilization and integrin perturbation. Stem Cells 39: 787–802.

458. Luo, S., N. B. Wehr and R. L. Levine, 2006 Quantitation of protein on gels and blots by infrared fluorescence of Coomassie blue and Fast Green. Anal Biochem 350: 233–238.

459. Luo, W., M. S. Friedman, K. D. Hankenson and P. J. Woolf, 2011 Time series gene expression profiling and temporal regulatory pathway analysis of BMP6 induced osteoblast differentiation and mineralization. BMC Syst Biol 5: 82.

460. Luo, Y., S. Yang, X. Wu, S. Takahashi, L. Sun et al., 2021 Intestinal MYC modulates obesity-related metabolic dysfunction. Nat Metab 3: 923–939.

461. Lupo, R., A. Breiling, M. E. Bianchi and V. Orlando, 2001 Drosophila chromosome condensation proteins Topoisomerase II and Barren colocalize with Polycomb and maintain Fab-7 PRE silencing. Mol Cell 7: 127–136.

462. Luscher-Firzlaff, J., I. Gawlista, J. Vervoorts, K. Kapelle, T. Braunschweig et al., 2008 The human trithorax protein hASH2 functions as an oncoprotein. Cancer Res 68: 749–758.

463. Luscher, B., 2001 Function and regulation of the transcription factors of the Myc/Max/Mad network. Gene 277: 1–14.

464. Lyman, D. F., and B. Yedvobnick, 1995 Drosophila Notch receptor activity suppresses Hairless function during adult external sensory organ development. Genetics 141: 1491–1505.

465. Ma, J., H. Plesken, J. E. Treisman, I. Edelman-Novemsky and M. Ren, 2004 Lightoid and Claret: a rab GTPase and its putative guanine nucleotide exchange factor in biogenesis of Drosophila eye pigment granules. Proc Natl Acad Sci U S A 101: 11652–11657.

466. Mahajan, S. S., K. M. Johnson and A. C. Wilson, 2003 Molecular cloning of Drosophila HCF reveals proteolytic processing and self-association of the encoded protein. J Cell Physiol 194: 117–126.

467. Maier, D., P. Kurth, A. Schulz, A. Russell, Z. Yuan et al., 2011 Structural and functional analysis of the repressor complex in the Notch signaling pathway of Drosophila melanogaster. Mol Biol Cell 22: 3242–3252.

468. Maines, J. Z., L. M. Stevens, X. Tong and D. Stein, 2004 Drosophila dMyc is required for ovary cell growth and endoreplication. Development 131: 775–786.

469. Mallik, M., M. Catinozzi, C. B. Hug, L. Zhang, M. Wagner et al., 2018 Xrp1 genetically interacts with the ALS-associated FUS orthologue caz and mediates its toxicity. J Cell Biol 217: 3947–3964.

470. Manohar, C. F., H. R. Salwen, G. M. Brodeur and S. L. Cohn, 1995 Co-amplification and concomitant high levels of expression of a DEAD box gene with MYCN in human neuroblastoma. Genes Chromosomes Cancer 14: 196–203.

471. Marenda, D. R., C. B. Zraly and A. K. Dingwall, 2004 The Drosophila Brahma (SWI/SNF) chromatin remodeling complex exhibits cell-type specific activation and repression functions. Dev Biol 267: 279–293.

472. Marenda, D. R., C. B. Zraly, Y. Feng, S. Egan and A. K. Dingwall, 2003 The Drosophila SNR1 (SNF5/INI1) subunit directs essential developmental functions of the Brahma chromatin remodeling complex. Mol Cell Biol 23: 289–305.

473. Marinho, J., F. Casares and P. S. Pereira, 2011 The Drosophila Nol12 homologue viriato is a dMyc target that regulates nucleolar architecture and is required for dMyc-stimulated cell growth. Development 138: 349–357.

474. Marinkovic, T., and D. Marinkovic, 2021 Obscure Involvement of MYC in Neurodegenerative Diseases and Neuronal Repair. Mol Neurobiol 58: 4169–4177.

475. Markesich, D. C., K. M. Gajewski, M. E. Nazimiec and K. Beckingham, 2000 bicaudal encodes the Drosophila beta NAC homolog, a component of the ribosomal translational machinery*. Development 127: 559–572.

476. Marmorstein, R., 2003 Structure of SET domain proteins: a new twist on histone methylation. Trends Biochem Sci 28: 59–62.

477. Marquez, R. M., M. A. Singer, N. T. Takaesu, W. R. Waldrip, Y. Kraytsberg et al., 2001 Transgenic analysis of the Smad family of TGF-beta signal transducers in Drosophila melanogaster suggests new roles and new interactions between family members. Genetics 157: 1639–1648.

478. Marsden, M., and D. W. DeSimone, 2001 Regulation of cell polarity, radial intercalation and epiboly in Xenopus: novel roles for integrin and fibronectin. Development 128: 3635-3647.

479. Martin-Bermudo, M. D., 2000 Integrins modulate the Egfr signaling pathway to regulate tendon cell differentiation in the Drosophila embryo. Development 127: 2607–2615.

480. Martin, D. D., E. Beauchamp and L. G. Berthiaume, 2011 Post-translational myristoylation: Fat matters in cellular life and death. Biochimie 93: 18–31.

481. Martin, S. R., and P. M. Bayley, 2004 Calmodulin bridging of IQ motifs in myosin-V. FEBS Lett 567: 166–170.

482. Martins, T., F. Meghini, F. Florio and Y. Kimata, 2017 The APC/C Coordinates Retinal Differentiation with G1 Arrest through the Nek2-Dependent Modulation of Wingless Signaling. Dev Cell 40: 67–80.

483. Marygold, S. J., J. Roote, G. Reuter, A. Lambertsson, M. Ashburner et al., 2007 The ribosomal protein genes and Minute loci of Drosophila melanogaster. Genome Biol 8: R216.

484. Marygold, S. J., C. Walker, M. Orme and S. Leevers, 2011 Genetic characterization of ebi reveals its critical role in Drosophila wing growth. Fly (Austin) 5: 291–303.

485. Mashtalir, N., H. Suzuki, D. P. Farrell, A. Sankar, J. Luo et al., 2020 A Structural Model of the Endogenous Human BAF Complex Informs Disease Mechanisms. Cell 183: 802–817 e824.

486. Masri, S., M. Cervantes and P. Sassone-Corsi, 2013 The circadian clock and cell cycle: interconnected biological circuits. Curr Opin Cell Biol 25: 730–734.

487. Matharu, N. K., T. Hussain, R. Sankaranarayanan and R. K. Mishra, 2010 Vertebrate homologue of Drosophila GAGA factor. J Mol Biol 400: 434–447.

488. Mathews, K. W., M. Cavegn and M. Zwicky, 2017 Sexual Dimorphism of Body Size Is Controlled by Dosage of the X-Chromosomal Gene Myc and by the Sex-Determining Gene tra in Drosophila. Genetics 205: 1215–1228.

489. Matias-Barrios, V. M., and X. Dong, 2023 The Implication of Topoisomerase II Inhibitors in Synthetic Lethality for Cancer Therapy. Pharmaceuticals (Basel) 16.

490. Matsui, M., K. C. Sharma, C. Cooke, B. T. Wakimoto, M. Rasool et al., 2011 Nuclear structure and chromosome segregation in Drosophila male meiosis depend on the ubiquitin ligase dTopors. Genetics 189: 779–793.

491. Matsuo, T., S. Yamaguchi, S. Mitsui, A. Emi, F. Shimoda et al., 2003 Control mechanism of the circadian clock for timing of cell division in vivo. Science 302: 255–259.

492. Matsuzawa, S. I., and J. C. Reed, 2001 Siah-1, SIP, and Ebi collaborate in a novel pathway for beta-catenin degradation linked to p53 responses. Mol Cell 7: 915-926.

493. Matyash, A., N. Singh, S. D. Hanes, H. Urlaub and H. Jackle, 2009 SAP18 promotes Kruppel-dependent transcriptional repression by enhancer-specific histone deacetylation. J Biol Chem 284: 3012–3020.

494. Mazur, P. K., H. Einwachter, M. Lee, B. Sipos, H. Nakhai et al., 2010 Notch2 is required for progression of pancreatic intraepithelial neoplasia and development of pancreatic ductal adenocarcinoma. Proc Natl Acad Sci U S A 107: 13438–13443.

495. McGrail, M., and T. S. Hays, 1997 The microtubule motor cytoplasmic dynein is required for spindle orientation during germline cell divisions and oocyte differentiation in Drosophila. Development 124: 2409–2419.

496. Meng, Z., T. Moroishi, V. Mottier-Pavie, S. W. Plouffe, C. G. Hansen et al., 2015 MAP4K family kinases act in parallel to MST1/2 to activate LATS1/2 in the Hippo pathway. Nat Commun 6: 8357.

497. Methot, N., and K. Basler, 2000 Suppressor of fused opposes hedgehog signal transduction by impeding nuclear accumulation of the activator form of Cubitus interruptus. Development 127: 4001–4010.

498. Michalak, W., V. Tsiamis, V. Schwammle and A. Rogowska-Wrzesinska, 2019 ComplexBrowser: A Tool for Identification and Quantification of Protein Complexes in Large-scale Proteomics Datasets. Mol Cell Proteomics 18: 2324–2334.

499. Miles, W. O., E. Jaffray, S. G. Campbell, S. Takeda, L. J. Bayston et al., 2008 Medea SUMOylation restricts the signaling range of the Dpp morphogen in the Drosophila embryo. Genes Dev 22: 2578–2590.

500. Militti, C., S. Maenner, P. B. Becker and F. Gebauer, 2014 UNR facilitates the interaction of MLE with the lncRNA roX2 during Drosophila dosage compensation. Nat Commun 5: 4762.

501. Miller, C. L. W., J. L. Warner and F. Winston, 2023 Insights into Spt6: a histone chaperone that functions in transcription, DNA replication, and genome stability. Trends Genet 39: 858–872.

502. Miller, M., A. Chen, V. Gobert, B. Auge, M. Beau et al., 2017 Control of RUNX-induced repression of Notch signaling by MLF and its partner DnaJ-1 during Drosophila hematopoiesis. PLoS Genet 13: e1006932.

503. Mishra, A. K., N. Sachan, M. Mutsuddi and A. Mukherjee, 2015 Kinase active Misshapen regulates Notch signaling in Drosophila melanogaster. Exp Cell Res 339: 51–60.

504. Mitchell, N. C., E. B. Tchoubrieva, A. Chahal, S. Woods, A. Lee et al., 2015 S6 Kinase is essential for MYC-dependent rDNA transcription in Drosophila. Cell Signal 27: 2045–2053.

505. Mitchell, S. A., E. C. Brown, M. J. Coldwell, R. J. Jackson and A. E. Willis, 2001 Protein factor requirements of the Apaf-1 internal ribosome entry segment: roles of polypyrimidine tract binding protein and upstream of N-ras. Mol Cell Biol 21: 3364–3374.

506. Mitchelson, A., M. Simonelig, C. Williams and K. O’Hare, 1993 Homology with Saccharomyces cerevisiae RNA14 suggests that phenotypic suppression in Drosophila melanogaster by suppressor of forked occurs at the level of RNA stability. Genes Dev 7: 241–249.

507. Mizuguchi, G., A. Vassilev, T. Tsukiyama, Y. Nakatani and C. Wu, 2001 ATP-dependent nucleosome remodeling and histone hyperacetylation synergistically facilitate transcription of chromatin. J Biol Chem 276: 14773–14783.

508. Mohan, M., M. Bartkuhn, M. Herold, A. Philippen, N. Heinl et al., 2007 The Drosophila insulator proteins CTCF and CP190 link enhancer blocking to body patterning. EMBO J 26: 4203–4214.

509. Mohrmann, L., K. Langenberg, J. Krijgsveld, A. J. Kal, A. J. Heck et al., 2004 Differential targeting of two distinct SWI/SNF-related Drosophila chromatin-remodeling complexes. Mol Cell Biol 24: 3077–3088.

510. Moller, A., F. W. Avila, J. W. Erickson and H. Jackle, 2005 Drosophila BAP60 is an essential component of the Brahma complex, required for gene activation and repression. J Mol Biol 352: 329–337.

511. Molnar, C., and J. F. de Celis, 2013 Tay bridge is a negative regulator of EGFR signalling and interacts with Erk and Mkp3 in the Drosophila melanogaster wing. PLoS Genet 9: e1003982.

512. Moore, R., K. Vogt, A. E. Acosta-Martin, P. Shire, M. Zeidler et al., 2020 Integration of JAK/STAT receptor-ligand trafficking, signalling and gene expression in Drosophila melanogaster cells. J Cell Sci 133.

513. Morales-Mulia, S., and J. M. Scholey, 2005 Spindle pole organization in Drosophila S2 cells by dynein, abnormal spindle protein (Asp), and KLP10A. Mol Biol Cell 16: 3176–3186.

514. Morelli, E., M. Fulciniti, M. K. Samur, C. F. Ribeiro, L. Wert-Lamas et al., 2023 A MIR17HG-derived long noncoding RNA provides an essential chromatin scaffold for protein interaction and myeloma growth. Blood 141: 391–405.

515. Morera, S., M. Chiadmi, G. LeBras, I. Lascu and J. Janin, 1995 Mechanism of phosphate transfer by nucleoside diphosphate kinase: X-ray structures of the phosphohistidine intermediate of the enzymes from Drosophila and Dictyostelium. Biochemistry 34: 11062–11070.

516. Morgan, J. L., Y. Song and E. Barbar, 2011 Structural dynamics and multiregion interactions in dynein-dynactin recognition. J Biol Chem 286: 39349–39359.

517. Morillo Prado, J. R., S. Srinivasan and M. T. Fuller, 2013 The histone variant His2Av is required for adult stem cell maintenance in the Drosophila testis. PLoS Genet 9: e1003903.

518. Moser, M., A. Pscherer, C. Roth, J. Becker, G. Mucher et al., 1997 Enhanced apoptotic cell death of renal epithelial cells in mice lacking transcription factor AP-2beta. Genes Dev 11: 1938–1948.

519. Mount, S. M., and H. K. Salz, 2000 Pre-messenger RNA processing factors in the Drosophila genome. J Cell Biol 150: F37–44.

520. Muller, B., B. Hartmann, G. Pyrowolakis, M. Affolter and K. Basler, 2003 Conversion of an extracellular Dpp/BMP morphogen gradient into an inverse transcriptional gradient. Cell 113: 221–233.

521. Muller, D., S. J. Kugler, A. Preiss, D. Maier and A. C. Nagel, 2005 Genetic modifier screens on Hairless gain-of-function phenotypes reveal genes involved in cell differentiation, cell growth and apoptosis in Drosophila melanogaster. Genetics 171: 1137–1152.

522. Murphy, T. D., 2003 Drosophila skpA, a component of SCF ubiquitin ligases, regulates centrosome duplication independently of cyclin E accumulation. J Cell Sci 116: 2321–2332.

523. Myster, S. H., and M. Peifer, 2001 Wingless can’t fly so it hitches a ride with dynein. Bioessays 23: 869–872.

524. Nagel, A. C., A. Krejci, G. Tenin, A. Bravo-Patino, S. Bray et al., 2005 Hairless-mediated repression of notch target genes requires the combined activity of Groucho and CtBP corepressors. Mol Cell Biol 25: 10433–10441.

525. Nagel, A. C., D. Maier and A. Preiss, 2000 Su(H)-independent activity of hairless during mechano-sensory organ formation in Drosophila. Mech Dev 94: 3–12.

526. Nair, M., A. Teng, V. Bilanchone, A. Agrawal, B. Li et al., 2006 Ovol1 regulates the growth arrest of embryonic epidermal progenitor cells and represses c-myc transcription. J Cell Biol 173: 253–264.

527. Narayanan, R., and M. Ramaswami, 2001 Endocytosis in Drosophila: progress, possibilities, prognostications. Exp Cell Res 271: 28–35.

528. Neto-Silva, R. M., S. de Beco and L. A. Johnston, 2010 Evidence for a growth-stabilizing regulatory feedback mechanism between Myc and Yorkie, the Drosophila homolog of Yap. Dev Cell 19: 507–520.

529. Neves, A., and R. N. Eisenman, 2019 Distinct gene-selective roles for a network of core promoter factors in Drosophila neural stem cell identity. Biol Open 8.

530. Nguyen, M. B., L. T. Vuong and K. W. Choi, 2016 Ebi modulates wing growth by ubiquitin-dependent downregulation of Crumbs in Drosophila. Development 143: 3506–3513.

531. Nicholson, S. C., B. N. Nicolay, M. V. Frolov and K. H. Moberg, 2011 Notch-dependent expression of the archipelago ubiquitin ligase subunit in the Drosophila eye. Development 138: 251–260.

532. Nicolas, E., N. Chenouard, J. C. Olivo-Marin and A. Guichet, 2009 A dual role for actin and microtubule cytoskeleton in the transport of Golgi units from the nurse cells to the oocyte across ring canals. Mol Biol Cell 20: 556–568.

533. Nie, M., Y. Xie, J. A. Loo and A. J. Courey, 2009 Genetic and proteomic evidence for roles of Drosophila SUMO in cell cycle control, Ras signaling, and early pattern formation. PLoS One 4: e5905.

534. Nikalayevich, E., and H. Ohkura, 2015 The NuRD nucleosome remodelling complex and NHK-1 kinase are required for chromosome condensation in oocytes. J Cell Sci 128: 566–575.

535. Nilkanta, C., and A. Bagchi, 2018 Comparative analysis of prokaryotic and eukaryotic transcription factors using machine-learning techniques. Bioinformation 14: 315–326.

536. Nir, R., R. Grossman, Z. Paroush and T. Volk, 2012 Phosphorylation of the Drosophila melanogaster RNA-binding protein HOW by MAPK/ERK enhances its dimerization and activity. PLoS Genet 8: e1002632.

537. Nollet, F., G. Berx and F. van Roy, 1999 The role of the E-cadherin/catenin adhesion complex in the development and progression of cancer. Mol Cell Biol Res Commun 2: 77–85.

538. Norbury, C., and P. Nurse, 1992 Animal cell cycles and their control. Annu Rev Biochem 61: 441–470.

539. Noura, M., S. Tomita, T. Yasuda, S. Tsuzuki, H. Kiyoi et al., 2024 NUP98-BPTF promotes oncogenic transformation through PIM1 upregulation. Cancer Med 13: e7445.

540. Nowak, A. J., C. Alfieri, C. U. Stirnimann, V. Rybin, F. Baudin et al., 2011 Chromatin-modifying complex component Nurf55/p55 associates with histones H3 and H4 and polycomb repressive complex 2 subunit Su(z)12 through partially overlapping binding sites. J Biol Chem 286: 23388–23396.

541. Ntwasa, M., M. Egerton and N. J. Gay, 1997 Sequence and expression of Drosophila myristoyl-CoA: protein N-myristoyl transferase: evidence for proteolytic processing and membrane localisation. J Cell Sci 110 (Pt 2): 149–156.

542. Nurminsky, D. I., M. V. Nurminskaya, E. V. Benevolenskaya, Y. Y. Shevelyov, D. L. Hartl et al., 1998 Cytoplasmic dynein intermediate-chain isoforms with different targeting properties created by tissue-specific alternative splicing. Mol Cell Biol 18: 6816–6825.

543. O’Keefe, D. D., E. Gonzalez-Nino, M. Burnett, L. Dylla, S. M. Lambeth et al., 2009 Rap1 maintains adhesion between cells to affect Egfr signaling and planar cell polarity in Drosophila. Dev Biol 333: 143–160.

544. O’Neill, E. M., I. Rebay, R. Tjian and G. M. Rubin, 1994 The activities of two Ets-related transcription factors required for Drosophila eye development are modulated by the Ras/MAPK pathway. Cell 78: 137–147.

545. Oegema, K., C. Wiese, O. C. Martin, R. A. Milligan, A. Iwamatsu et al., 1999 Characterization of two related Drosophila gamma-tubulin complexes that differ in their ability to nucleate microtubules. J Cell Biol 144: 721–733.

546. Oellers, N., and E. Hafen, 1996 Biochemical characterization of rolledSem, an activated form of Drosophila mitogen-activated protein kinase. J Biol Chem 271: 24939–24944.

547. Ogirenko, A. A., D. A. Karagodin, N. V. Pavlova, S. A. Fedorova, M. A. Voloshina et al., 2008 [Molecular and genetic description of a new hypomorphic mutation of Trithorax-like gene and analysis of its effect on Drosophila melanogaster oogenesis]. Ontogenez 39: 134–142.

548. Ogiyama, Y., B. Schuettengruber, G. L. Papadopoulos, J. M. Chang and G. Cavalli, 2018 Polycomb-Dependent Chromatin Looping Contributes to Gene Silencing during Drosophila Development. Mol Cell 71: 73–88 e75.

549. Ogura, Y., F. L. Wen, M. M. Sami, T. Shibata and S. Hayashi, 2018 A Switch-like Activation Relay of EGFR-ERK Signaling Regulates a Wave of Cellular Contractility for Epithelial Invagination. Dev Cell 46: 162–172 e165.

550. Oh, S., M. Kato, C. Zhang, Y. Guo and P. A. Beachy, 2015 A Comparison of Ci/Gli Activity as Regulated by Sufu in Drosophila and Mammalian Hedgehog Response. PLoS One 10: e0135804.

551. Ohkuma, Y., H. Sumimoto, A. Hoffmann, S. Shimasaki, M. Horikoshi et al., 1991 Structural motifs and potential sigma homologies in the large subunit of human general transcription factor TFIIE. Nature 354: 398–401.

552. Ohler, U., G. C. Liao, H. Niemann and G. M. Rubin, 2002 Computational analysis of core promoters in the Drosophila genome. Genome Biol 3: RESEARCH0087.

553. Ohler, U., and D. A. Wassarman, 2010 Promoting developmental transcription. Development 137: 15–26.

554. Ohshiro, T., Y. Emori and K. Saigo, 2002 Ligand-dependent activation of breathless FGF receptor gene in Drosophila developing trachea. Mech Dev 114: 3–11.

555. Okada, M., and Y. B. Shi, 2018 The balance of two opposing factors Mad and Myc regulates cell fate during tissue remodeling. Cell Biosci 8: 51.

556. Okamoto, T., S. Yamamoto, Y. Watanabe, T. Ohta, F. Hanaoka et al., 1998 Analysis of the role of TFIIE in transcriptional regulation through structure-function studies of the TFIIEbeta subunit. J Biol Chem 273: 19866–19876.

557. On, K., G. Crevel, S. Cotterill, M. Itoh and Y. Kato, 2021 Drosophila telomere capping protein HOAP interacts with DSB sensor proteins Mre11 and Nbs. Genes Cells 26: 219–229.

558. Orian, A., J. J. Delrow, A. E. Rosales Nieves, M. Abed, D. Metzger et al., 2007 A Myc-Groucho complex integrates EGF and Notch signaling to regulate neural development. Proc Natl Acad Sci U S A 104: 15771–15776.

559. Orian, A., S. S. Grewal, P. S. Knoepfler, B. A. Edgar, S. M. Parkhurst et al., 2005 Genomic binding and transcriptional regulation by the Drosophila Myc and Mnt transcription factors. Cold Spring Harb Symp Quant Biol 70: 299–307.

560. Orian, A., B. van Steensel, J. Delrow, H. J. Bussemaker, L. Li et al., 2003 Genomic binding by the Drosophila Myc, Max, Mad/Mnt transcription factor network. Genes Dev 17: 1101–1114.

561. Oster, S. K., W. W. Marhin, C. Asker, L. M. Facchini, P. A. Dion et al., 2000 Myc is an essential negative regulator of platelet-derived growth factor beta receptor expression. Mol Cell Biol 20: 6768–6778.

562. Ota, R., and S. Kobayashi, 2020 Myc plays an important role in Drosophila P-M hybrid dysgenesis to eliminate germline cells with genetic damage. Commun Biol 3: 185.

563. Pai, C. Y., E. P. Lei, D. Ghosh and V. G. Corces, 2004 The centrosomal protein CP190 is a component of the gypsy chromatin insulator. Mol Cell 16: 737–748.

564. Pan, D., and G. M. Rubin, 1995 cAMP-dependent protein kinase and hedgehog act antagonistically in regulating decapentaplegic transcription in Drosophila imaginal discs. Cell 80: 543–552.

565. Panda, D., P. Pascual-Garcia, M. Dunagin, M. Tudor, K. C. Hopkins et al., 2014 Nup98 promotes antiviral gene expression to restrict RNA viral infection in Drosophila. Proc Natl Acad Sci U S A 111: E3890–3899.

566. Pankotai, T., Z. Ujfaludi, E. Vamos, K. Suri and I. M. Boros, 2010 The dissociable RPB4 subunit of RNA Pol II has vital functions in Drosophila. Mol Genet Genomics 283: 89–97.

567. Papoulas, O., G. Daubresse, J. A. Armstrong, J. Jin, M. P. Scott et al., 2001 The HMG-domain protein BAP111 is important for the function of the BRM chromatin-remodeling complex in vivo. Proc Natl Acad Sci U S A 98: 5728–5733.

568. Pardee, A. B., 1989 G1 events and regulation of cell proliferation. Science 246: 603–608.

569. Park, J. M., J. M. Kim, L. K. Kim, S. N. Kim, J. Kim-Ha et al., 2003a Signal-induced transcriptional activation by Dif requires the dTRAP80 mediator module. Mol Cell Biol 23: 1358–1367.

570. Park, J. W., K. Parisky, A. M. Celotto, R. A. Reenan and B. R. Graveley, 2004 Identification of alternative splicing regulators by RNA interference in Drosophila. Proc Natl Acad Sci U S A 101: 15974–15979.

571. Park, Y., C. Rangel, M. M. Reynolds, M. C. Caldwell, M. Johns et al., 2003b Drosophila perlecan modulates FGF and hedgehog signals to activate neural stem cell division. Dev Biol 253: 247–257.

572. Parker, D. S., Y. Y. Ni, J. L. Chang, J. Li and K. M. Cadigan, 2008 Wingless signaling induces widespread chromatin remodeling of target loci. Mol Cell Biol 28: 1815–1828.

573. Parkhurst, S. M., and V. G. Corces, 1986 Mutations at the suppressor of forked locus increase the accumulation of gypsy-encoded transcripts in Drosophila melanogaster. Mol Cell Biol 6: 2271–2274.

574. Parkinson, W. M., M. Dookwah, M. L. Dear, C. L. Gatto, K. Aoki et al., 2016 Synaptic roles for phosphomannomutase type 2 in a new Drosophila congenital disorder of glycosylation disease model. Dis Model Mech 9: 513–527.

575. Parrott, B. B., Y. Chiang, A. Hudson, A. Sarkar, A. Guichet et al., 2011 Nucleoporin98-96 function is required for transit amplification divisions in the germ line of Drosophila melanogaster. PLoS One 6: e25087.

576. Pascual-Garcia, P., J. Jeong and M. Capelson, 2014 Nucleoporin Nup98 associates with Trx/MLL and NSL histone-modifying complexes and regulates Hox gene expression. Cell Rep 9: 433–442.

577. Patalano, S., M. Mihailovich, Y. Belacortu, N. Paricio and F. Gebauer, 2009 Dual sex-specific functions of Drosophila Upstream of N-ras in the control of X chromosome dosage compensation. Development 136: 689–698.

578. Patel, J. H., Y. Du, P. G. Ard, C. Phillips, B. Carella et al., 2004 The c-MYC oncoprotein is a substrate of the acetyltransferases hGCN5/PCAF and TIP60. Mol Cell Biol 24: 10826–10834.

579. Pearson, J. C., and S. T. Crews, 2014 Enhancer diversity and the control of a simple pattern of Drosophila CNS midline cell expression. Dev Biol 392: 466–482.

580. Peck, V. M., E. W. Gerner and A. E. Cress, 1992 Delta-type DNA polymerase characterized from Drosophila melanogaster embryos. Nucleic Acids Res 20: 5779–5784.

581. Peissert, S., A. Schlosser, R. Kendel, J. Kuper and C. Kisker, 2020 Structural basis for CDK7 activation by MAT1 and Cyclin H. Proc Natl Acad Sci U S A 117: 26739–26748.

582. Pek, J. W., A. K. Lim and T. Kai, 2009 Drosophila maelstrom ensures proper germline stem cell lineage differentiation by repressing microRNA-7. Dev Cell 17: 417–424.

583. Pek, J. W., B. F. Ng and T. Kai, 2012 Polo-mediated phosphorylation of Maelstrom regulates oocyte determination during oogenesis in Drosophila. Development 139: 4505–4513.

584. Pennetier, D., J. Oyallon, I. Morin-Poulard, S. Dejean, A. Vincent et al., 2012 Size control of the Drosophila hematopoietic niche by bone morphogenetic protein signaling reveals parallels with mammals. Proc Natl Acad Sci U S A 109: 3389–3394.

585. Perez-Lluch, S., E. Blanco, A. Carbonell, D. Raha, M. Snyder et al., 2011 Genome-wide chromatin occupancy analysis reveals a role for ASH2 in transcriptional pausing. Nucleic Acids Res 39: 4628–4639.

586. Perez-Roger, I., S. H. Kim, B. Griffiths, A. Sewing and H. Land, 1999 Cyclins D1 and D2 mediate myc-induced proliferation via sequestration of p27(Kip1) and p21(Cip1). EMBO J 18: 5310–5320.

587. Perrin, L., C. Benassayag, D. Morello, J. Pradel and J. Montagne, 2003 Modulo is a target of Myc selectively required for growth of proliferative cells in Drosophila. Mech Dev 120: 645–655.

588. Perrin, L., O. Demakova, L. Fanti, S. Kallenbach, S. Saingery et al., 1998 Dynamics of the sub-nuclear distribution of Modulo and the regulation of position-effect variegation by nucleolus in Drosophila. J Cell Sci 111 (Pt 18): 2753–2761.

589. Perrin, L., P. Romby, P. Laurenti, H. Berenger, S. Kallenbach et al., 1999 The Drosophila modifier of variegation modulo gene product binds specific RNA sequences at the nucleolus and interacts with DNA and chromatin in a phosphorylation-dependent manner. J Biol Chem 274: 6315–6323.

590. Petit, V., U. Nussbaumer, C. Dossenbach and M. Affolter, 2004 Downstream-of-FGFR is a fibroblast growth factor-specific scaffolding protein and recruits Corkscrew upon receptor activation. Mol Cell Biol 24: 3769–3781.

591. Petruk, S., Y. Sedkov, K. M. Riley, J. Hodgson, F. Schweisguth et al., 2006 Transcription of bxd noncoding RNAs promoted by trithorax represses Ubx in cis by transcriptional interference. Cell 127: 1209–1221.

592. Peyrefitte, S., D. Kahn and M. Haenlin, 2001 New members of the Drosophila Myc transcription factor subfamily revealed by a genome-wide examination for basic helix-loop-helix genes. Mech Dev 104: 99–104.

593. Pflugfelder, G. O., F. Eichinger and J. Shen, 2017 T-Box Genes in Drosophila Limb Development. Curr Top Dev Biol 122: 313–354.

594. Phillips, J. W., Y. Pan, B. L. Tsai, Z. Xie, L. Demirdjian et al., 2020 Pathway-guided analysis identifies Myc-dependent alternative pre-mRNA splicing in aggressive prostate cancers. Proc Natl Acad Sci U S A 117: 5269–5279.

595. Phizicky, E. M., and S. Fields, 1995 Protein-protein interactions: methods for detection and analysis. Microbiol Rev 59: 94–123.

596. Piazzi, M., A. Bavelloni, A. Gallo, I. Faenza and W. L. Blalock, 2019 Signal Transduction in Ribosome Biogenesis: A Recipe to Avoid Disaster. Int J Mol Sci 20.

597. Pojer, J. M., S. A. Manning, B. Kroeger, S. Kondo and K. F. Harvey, 2021 The Hippo pathway uses different machinery to control cell fate and organ size. iScience 24: 102830.

598. Polesello, C., I. Delon, P. Valenti, P. Ferrer and F. Payre, 2002 Dmoesin controls actin-based cell shape and polarity during Drosophila melanogaster oogenesis. Nat Cell Biol 4: 782–789.

599. Port, F., M. Kuster, P. Herr, E. Furger, C. Banziger et al., 2008 Wingless secretion promotes and requires retromer-dependent cycling of Wntless. Nat Cell Biol 10: 178–185.

600. Prioleau, M. N., J. Huet, A. Sentenac and M. Mechali, 1994 Competition between chromatin and transcription complex assembly regulates gene expression during early development. Cell 77: 439–449.

601. Prober, D. A., and B. A. Edgar, 2002 Interactions between Ras1, dMyc, and dPI3K signaling in the developing Drosophila wing. Genes Dev 16: 2286-2299.

602. Prochownik, E. V., 2022 Regulation of Normal and Neoplastic Proliferation and Metabolism by the Extended Myc Network. Cells 11.

603. Qian, S., J. Y. Zhang, M. A. Kay and M. Jacobs-Lorena, 1987 Structural analysis of the Drosophila rpA1 gene, a member of the eucaryotic ‘A’ type ribosomal protein family. Nucleic Acids Res 15: 987–1003.

604. Qiao, F., B. Harada, H. Song, J. Whitelegge, A. J. Courey et al., 2006 Mae inhibits Pointed-P2 transcriptional activity by blocking its MAPK docking site. EMBO J 25: 70–79.

605. Qiu, H., C. Hu, N. A. Gaur and A. G. Hinnebusch, 2012 Pol II CTD kinases Bur1 and Kin28 promote Spt5 CTR-independent recruitment of Paf1 complex. EMBO J 31: 3494–3505.

606. Quinn, L. M., J. Secombe and G. R. Hime, 2013 Myc in stem cell behaviour: insights from Drosophila. Advances in experimental medicine and biology 786: 269–285.

607. Radimerski, T., J. Montagne, F. Rintelen, H. Stocker, J. van der Kaay et al., 2002 dS6K-regulated cell growth is dPKB/dPI(3)K-independent, but requires dPDK1. Nat Cell Biol 4: 251–255.

608. Raffeiner, P., A. Schraffl, T. Schwarz, R. Rock, K. Ledolter et al., 2017 Calcium-dependent binding of Myc to calmodulin. Oncotarget 8: 3327–3343.

609. Rafti, F., D. Scarvelis and P. F. Lasko, 1996 A Drosophila melanogaster homologue of the human DEAD-box gene DDX1. Gene 171: 225–229.

610. Raha, D., M. Hong and M. Snyder, 2010a ChIP-Seq: a method for global identification of regulatory elements in the genome. Curr Protoc Mol Biol Chapter 21: Unit 21 19 21-14.

611. Raha, D., Z. Wang, Z. Moqtaderi, L. Wu, G. Zhong et al., 2010b Close association of RNA polymerase II and many transcription factors with Pol III genes. Proc Natl Acad Sci U S A 107: 3639–3644.

612. Rajan, A., and N. Perrimon, 2010 Steroids make you bigger? Fat chance says Myc. Cell Metab 12: 7–9.

613. Ramanujam, P. L., S. Mehrotra, R. P. Kumar, S. Verma, G. Deshpande et al., 2021 Global chromatin organizer SATB1 acts as a context-dependent regulator of the Wnt/Wg target genes. Sci Rep 11: 3385.

614. Ramdas Nair, A., P. Singh, D. Salvador Garcia, D. Rodriguez-Crespo, B. Egger et al., 2016 The Microcephaly-Associated Protein Wdr62/CG7337 Is Required to Maintain Centrosome Asymmetry in Drosophila Neuroblasts. Cell Rep 14: 1100–1113.

615. Rauskolb, C., S. Sun, G. Sun, Y. Pan and K. D. Irvine, 2014 Cytoskeletal tension inhibits Hippo signaling through an Ajuba-Warts complex. Cell 158: 143–156.

616. Ravens, S., C. Yu, T. Ye, M. Stierle and L. Tora, 2015 Tip60 complex binds to active Pol II promoters and a subset of enhancers and co-regulates the c-Myc network in mouse embryonic stem cells. Epigenetics Chromatin 8: 45.

617. Ray-Jones, H., and M. Spivakov, 2021 Transcriptional enhancers and their communication with gene promoters. Cell Mol Life Sci 78: 6453–6485.

618. Ray, P., X. Luo, E. J. Rao, A. Basha, E. A. Woodruff, 3rd et al., 2010 The splicing factor Prp31 is essential for photoreceptor development in Drosophila. Protein Cell 1: 267-274.

619. Reaume, A. G., D. A. Knecht and A. Chovnick, 1991 The rosy locus in Drosophila melanogaster: xanthine dehydrogenase and eye pigments. Genetics 129: 1099–1109.

620. Reddy, B. V., and K. D. Irvine, 2011 Regulation of Drosophila glial cell proliferation by Merlin-Hippo signaling. Development 138: 5201–5212.

621. Reich, A., and B. Z. Shilo, 2002 Keren, a new ligand of the Drosophila epidermal growth factor receptor, undergoes two modes of cleavage. EMBO J 21: 4287–4296.

622. Reim, I., and M. Frasch, 2005 The Dorsocross T-box genes are key components of the regulatory network controlling early cardiogenesis in Drosophila. Development 132: 4911–4925.

623. Reiter, F., B. P. de Almeida and A. Stark, 2023 Enhancers display constrained sequence flexibility and context-specific modulation of motif function. Genome Res 33: 346–358.

624. Ren, F., Q. Shi, Y. Chen, A. Jiang, Y. T. Ip et al., 2013 Drosophila Myc integrates multiple signaling pathways to regulate intestinal stem cell proliferation during midgut regeneration. Cell Res 23: 1133–1146.

625. Ren, F., B. Wang, T. Yue, E. Y. Yun, Y. T. Ip et al., 2010 Hippo signaling regulates Drosophila intestine stem cell proliferation through multiple pathways. Proc Natl Acad Sci U S A 107: 21064–21069.

626. Ren, M., E. Coutavas, P. D’Eustachio and M. G. Rush, 1994 Effects of mutant Ran/TC4 proteins on cell cycle progression. Mol Cell Biol 14: 4216–4224.

627. Riese, J., X. Yu, A. Munnerlyn, S. Eresh, S. C. Hsu et al., 1997 LEF-1, a nuclear factor coordinating signaling inputs from wingless and decapentaplegic. Cell 88: 777–787.

628. Rincon-Limas, D. E., C. H. Lu, I. Canal and J. Botas, 2000 The level of DLDB/CHIP controls the activity of the LIM homeodomain protein apterous: evidence for a functional tetramer complex in vivo. EMBO J 19: 2602–2614.

629. Roberts, D. M., M. I. Pronobis, J. S. Poulton, E. G. Kane and M. Peifer, 2012 Regulation of Wnt signaling by the tumor suppressor adenomatous polyposis coli does not require the ability to enter the nucleus or a particular cytoplasmic localization. Mol Biol Cell 23: 2041–2056.

630. Rodriguez, C. F., and O. Llorca, 2020 RPAP3 C-Terminal Domain: A Conserved Domain for the Assembly of R2TP Co-Chaperone Complexes. Cells 9.

631. Roignant, J. Y., K. Legent, F. Janody and J. E. Treisman, 2010 The transcriptional co-factor Chip acts with LIM-homeodomain proteins to set the boundary of the eye field in Drosophila. Development 137: 273–281.

632. Rosales-Vega, M., D. Resendez-Perez, M. Zurita and M. Vazquez, 2023 TnaA, a trithorax group protein, modulates wingless expression in different regions of the Drosophila wing imaginal disc. Sci Rep 13: 15162.

633. Rospert, S., Y. Dubaquie and M. Gautschi, 2002 Nascent-polypeptide-associated complex. Cell Mol Life Sci 59: 1632–1639.

634. Rosselot, C., S. Baumel-Alterzon, Y. Li, G. Brill, L. Lambertini et al., 2021 The many lives of Myc in the pancreatic beta-cell. J Biol Chem 296: 100122.

635. Roux, P. P., and I. Topisirovic, 2018 Signaling Pathways Involved in the Regulation of mRNA Translation. Mol Cell Biol 38.

636. Roy, F., G. Laberge, M. Douziech, D. Ferland-McCollough and M. Therrien, 2002 KSR is a scaffold required for activation of the ERK/MAPK module. Genes Dev 16: 427–438.

637. Royou, A., C. Field, J. C. Sisson, W. Sullivan and R. Karess, 2004 Reassessing the role and dynamics of nonmuscle myosin II during furrow formation in early Drosophila embryos. Mol Biol Cell 15: 838–850.

638. Rust, K., M. D. Tiwari, V. K. Mishra, F. Grawe and A. Wodarz, 2018 Myc and the Tip60 chromatin remodeling complex control neuroblast maintenance and polarity in Drosophila. EMBO J 37.

639. Ryoo, H. D., A. Bergmann, H. Gonen, A. Ciechanover and H. Steller, 2002 Regulation of Drosophila IAP1 degradation and apoptosis by reaper and ubcD1. Nat Cell Biol 4: 432–438.

640. Sabino, D., N. H. Brown and R. Basto, 2011 Drosophila Ajuba is not an Aurora-A activator but is required to maintain Aurora-A at the centrosome. J Cell Sci 124: 1156–1166.

641. Samuels, T. J., A. I. Jarvelin, D. Ish-Horowicz and I. Davis, 2020 Imp/IGF2BP levels modulate individual neural stem cell growth and division through myc mRNA stability. Elife 9.

642. Sanchez-Arevalo Lobo, V. J., M. Doni, A. Verrecchia, S. Sanulli, G. Faga et al., 2013 Dual regulation of Myc by Abl. Oncogene 32: 5261–5271.

643. Sarikas, A., T. Hartmann and Z. Q. Pan, 2011 The cullin protein family. Genome Biol 12: 220.

644. Sato, M., and T. B. Kornberg, 2002 FGF is an essential mitogen and chemoattractant for the air sacs of the drosophila tracheal system. Dev Cell 3: 195–207.

645. Satou, A., T. Taira, S. M. Iguchi-Ariga and H. Ariga, 2001 A novel transrepression pathway of c-Myc. Recruitment of a transcriptional corepressor complex to c-Myc by MM-1, a c-Myc-binding protein. J Biol Chem 276: 46562-46567.

646. Savitsky, M., M. Kim, O. Kravchuk and Y. B. Schwartz, 2016 Distinct Roles of Chromatin Insulator Proteins in Control of the Drosophila Bithorax Complex. Genetics 202: 601–617.

647. Sazer, S., and M. Dasso, 2000 The ran decathlon: multiple roles of Ran. J Cell Sci 113 (Pt 7): 1111–1118.

648. Scantlebury, N., X. L. Zhao, V. G. Rodriguez Moncalvo, A. Camiletti, S. Zahanova et al., 2010 The Drosophila gene RanBPM functions in the mushroom body to regulate larval behavior. PLoS One 5: e10652.

649. Schaub, C., M. Rose and M. Frasch, 2019 Yorkie and JNK revert syncytial muscles into myoblasts during Org-1-dependent lineage reprogramming. J Cell Biol 218: 3572–3582.

650. Scheer, E., F. Delbac, L. Tora, D. Moras and C. Romier, 2012 TFIID TAF6-TAF9 complex formation involves the HEAT repeat-containing C-terminal domain of TAF6 and is modulated by TAF5 protein. J Biol Chem 287: 27580–27592.

651. Schmidt, E. V., 2004 The role of c-myc in regulation of translation initiation. Oncogene 23: 3217–3221.

652. Schneider, M., A. A. Khalil, J. Poulton, C. Castillejo-Lopez, D. Egger-Adam et al., 2006 Perlecan and Dystroglycan act at the basal side of the Drosophila follicular epithelium to maintain epithelial organization. Development 133: 3805–3815.

653. Schnepp, B., T. Donaldson, G. Grumbling, S. Ostrowski, R. Schweitzer et al., 1998 EGF domain swap converts a drosophila EGF receptor activator into an inhibitor. Genes Dev 12: 908–913.

654. Schweitzer, R., M. Shaharabany, R. Seger and B. Z. Shilo, 1995 Secreted Spitz triggers the DER signaling pathway and is a limiting component in embryonic ventral ectoderm determination. Genes Dev 9: 1518–1529.

655. Schweizer, L., D. Nellen and K. Basler, 2003 Requirement for Pangolin/dTCF in Drosophila Wingless signaling. Proc Natl Acad Sci U S A 100: 5846–5851.

656. Sears, R., G. Leone, J. DeGregori and J. R. Nevins, 1999 Ras enhances Myc protein stability. Mol Cell 3: 169–179.

657. Secombe, J., L. Li, L. Carlos and R. N. Eisenman, 2007 The Trithorax group protein Lid is a trimethyl histone H3K4 demethylase required for dMyc-induced cell growth. Genes Dev 21: 537–551.

658. Serafini, G., G. Giordani, L. Grillini, D. Andrenacci, G. Gargiulo et al., 2020 The Impact of Drosophila Awd/NME1/2 Levels on Notch and Wg Signaling Pathways. Int J Mol Sci 21.

659. Sethi, M. K., F. F. Buettner, A. Ashikov, V. B. Krylov, H. Takeuchi et al., 2012 Molecular cloning of a xylosyltransferase that transfers the second xylose to O-glucosylated epidermal growth factor repeats of notch. J Biol Chem 287: 2739–2748.

660. Shalaby, N. A., A. L. Parks, E. J. Morreale, M. C. Osswalt, K. M. Pfau et al., 2009 A screen for modifiers of notch signaling uncovers Amun, a protein with a critical role in sensory organ development. Genetics 182: 1061–1076.

661. Sharma, A., S. Halder, M. Felix, K. Nisaa, G. Deshpande et al., 2018 Insulin signaling modulates border cell movement in Drosophila oogenesis. Development 145.

662. Shchepinov, M. S., I. A. Udalova, A. J. Bridgman and E. M. Southern, 1997 Oligonucleotide dendrimers: synthesis and use as polylabelled DNA probes. Nucleic Acids Res 25: 4447–4454.

663. Shcherbata, H. R., C. Althauser, S. D. Findley and H. Ruohola-Baker, 2004 The mitotic-to-endocycle switch in Drosophila follicle cells is executed by Notch-dependent regulation of G1/S, G2/M and M/G1 cell-cycle transitions. Development 131: 3169-3181.

664. Shi, J., W. A. Whyte, C. J. Zepeda-Mendoza, J. P. Milazzo, C. Shen et al., 2013 Role of SWI/SNF in acute leukemia maintenance and enhancer-mediated Myc regulation. Genes Dev 27: 2648–2662.

665. Shi, Q., S. Li, S. Li, A. Jiang, Y. Chen et al., 2014 Hedgehog-induced phosphorylation by CK1 sustains the activity of Ci/Gli activator. Proc Natl Acad Sci U S A 111: E5651–5660.

666. Shidlovskii, Y. V., A. N. Krasnov, J. V. Nikolenko, L. A. Lebedeva, M. Kopantseva et al., 2005 A novel multidomain transcription coactivator SAYP can also repress transcription in heterochromatin. EMBO J 24: 97–107.

667. Shilatifard, A., 2012 The COMPASS family of histone H3K4 methylases: mechanisms of regulation in development and disease pathogenesis. Annu Rev Biochem 81: 65–95.

668. Shima, S., T. Aigaki, T. Nojima and D. Yamamoto, 2007 Identification of trf2 mutants of Drosophila with defects in anterior spiracle eversion. Arch Insect Biochem Physiol 64: 157–163.

669. Shimamura, M., A. Kyotani, Y. Azuma, H. Yoshida, T. Binh Nguyen et al., 2014 Genetic link between Cabeza, a Drosophila homologue of Fused in Sarcoma (FUS), and the EGFR signaling pathway. Exp Cell Res 326: 36–45.

670. Shir-Shapira, H., A. Sloutskin, O. Adato, A. Ovadia-Shochat, D. Ideses et al., 2019 Identification of evolutionarily conserved downstream core promoter elements required for the transcriptional regulation of Fushi tarazu target genes. PLoS One 14: e0215695.

671. Shohayeb, B., N. Mitchell, S. S. Millard, L. M. Quinn and D. C. H. Ng, 2020 Elevated levels of Drosophila Wdr62 promote glial cell growth and proliferation through AURKA signalling to AKT and MYC. Biochim Biophys Acta Mol Cell Res 1867: 118713.

672. Siddiqui-Jain, A., C. L. Grand, D. J. Bearss and L. H. Hurley, 2002 Direct evidence for a G-quadruplex in a promoter region and its targeting with a small molecule to repress c-MYC transcription. Proc Natl Acad Sci U S A 99: 11593–11598.

673. Siebel, C. W., R. Kanaar and D. C. Rio, 1994 Regulation of tissue-specific P-element pre-mRNA splicing requires the RNA-binding protein PSI. Genes Dev 8: 1713–1725.

674. Sienski, G., D. Donertas and J. Brennecke, 2012 Transcriptional silencing of transposons by Piwi and maelstrom and its impact on chromatin state and gene expression. Cell 151: 964–980.

675. Sim, D. Y., H. J. Lee, C. H. Ahn, J. Park, S. Y. Park et al., 2024 Negative Regulation of CPSF6 Suppresses the Warburg Effect and Angiogenesis Leading to Tumor Progression Via c-Myc Signaling Network: Potential Therapeutic Target for Liver Cancer Therapy. Int J Biol Sci 20: 3442–3460.

676. Simoes, S., B. Denholm, D. Azevedo, S. Sotillos, P. Martin et al., 2006 Compartmentalisation of Rho regulators directs cell invagination during tissue morphogenesis. Development 133: 4257–4267.

677. Sinenko, S. A., T. Hung, T. Moroz, Q. M. Tran, S. Sidhu et al., 2010 Genetic manipulation of AML1-ETO-induced expansion of hematopoietic precursors in a Drosophila model. Blood 116: 4612–4620.

678. Sinenko, S. A., L. Mandal, J. A. Martinez-Agosto and U. Banerjee, 2009 Dual role of wingless signaling in stem-like hematopoietic precursor maintenance in Drosophila. Dev Cell 16: 756–763.

679. Singh, A. M., and S. Dalton, 2009 The cell cycle and Myc intersect with mechanisms that regulate pluripotency and reprogramming. Cell Stem Cell 5: 141–149.

680. Siomi, M. C., K. Higashijima, A. Ishizuka and H. Siomi, 2002 Casein kinase II phosphorylates the fragile X mental retardation protein and modulates its biological properties. Mol Cell Biol 22: 8438–8447.

681. Slack, C., N. Alic, A. Foley, M. Cabecinha, M. P. Hoddinott et al., 2015 The Ras-Erk-ETS-Signaling Pathway Is a Drug Target for Longevity. Cell 162: 72–83.

682. Sloan, R. S., C. I. Swanson, L. Gavilano, K. N. Smith, P. Y. Malek et al., 2012 Characterization of null and hypomorphic alleles of the Drosophila l(2)dtl/cdt2 gene: Larval lethality and male fertility. Fly (Austin) 6: 173–183.

683. Sloutskin, A., D. Itzhak, G. Vogler, H. Pozeilov, D. Ideses et al., 2024 From promoter motif to cardiac function: a single DPE motif affects transcription regulation and organ function in vivo. Development 151.

684. Smale, S. T., and J. T. Kadonaga, 2003 The RNA polymerase II core promoter. Annu Rev Biochem 72: 449–479.

685. Small, S., and D. N. Arnosti, 2020 Transcriptional Enhancers in Drosophila. Genetics 216: 1–26.

686. Smelkinson, M. G., Q. Zhou and D. Kalderon, 2007 Regulation of Ci-SCFSlimb binding, Ci proteolysis, and hedgehog pathway activity by Ci phosphorylation. Dev Cell 13: 481–495.

687. Smibert, C. A., Y. S. Lie, W. Shillinglaw, W. J. Henzel and P. M. Macdonald, 1999 Smaug, a novel and conserved protein, contributes to repression of nanos mRNA translation in vitro. RNA 5: 1535–1547.

688. Smith-Bolton, R. K., M. I. Worley, H. Kanda and I. K. Hariharan, 2009 Regenerative growth in Drosophila imaginal discs is regulated by Wingless and Myc. Dev Cell 16: 797–809.

689. Smith, G. D., W. H. Ching, P. Cornejo-Paramo and E. S. Wong, 2023 Decoding enhancer complexity with machine learning and high-throughput discovery. Genome Biol 24: 116.

690. Smylla, T. K., M. Meier, A. Preiss and D. Maier, 2019 The Notch repressor complex in Drosophila: in vivo analysis of Hairless mutants using overexpression experiments. Dev Genes Evol 229: 13–24.

691. Solinet, S., K. Mahmud, S. F. Stewman, K. Ben El Kadhi, B. Decelle et al., 2013 The actin-binding ERM protein Moesin binds to and stabilizes microtubules at the cell cortex. J Cell Biol 202: 251–260.

692. Soltani-Bejnood, M., S. E. Thomas, L. Villeneuve, K. Schwartz, C. S. Hong et al., 2007 Role of the mod(mdg4) common region in homolog segregation in Drosophila male meiosis. Genetics 176: 161–180.

693. Somma, M. P., B. Fasulo, G. Cenci, E. Cundari and M. Gatti, 2002 Molecular dissection of cytokinesis by RNA interference in Drosophila cultured cells. Mol Biol Cell 13: 2448–2460.

694. Song, H., P. Hasson, Z. Paroush and A. J. Courey, 2004 Groucho oligomerization is required for repression in vivo. Mol Cell Biol 24: 4341–4350.

695. Song, H., C. Spichiger-Haeusermann and K. Basler, 2009 The ISWI-containing NURF complex regulates the output of the canonical Wingless pathway. EMBO Rep 10: 1140–1146.

696. Song, S., H. Herranz and S. M. Cohen, 2017 The chromatin remodeling BAP complex limits tumor-promoting activity of the Hippo pathway effector Yki to prevent neoplastic transformation in Drosophila epithelia. Dis Model Mech 10: 1201–1209.

697. Spain, M. M., J. A. Caruso, A. Swaminathan and L. A. Pile, 2010 Drosophila SIN3 isoforms interact with distinct proteins and have unique biological functions. J Biol Chem 285: 27457–27467.

698. Spencer, C. A., and M. Groudine, 1991 Control of c-myc regulation in normal and neoplastic cells. Adv Cancer Res 56: 1–48.

699. Squire, J. A., P. S. Thorner, S. Weitzman, J. D. Maggi, P. Dirks et al., 1995 Co-amplification of MYCN and a DEAD box gene (DDX1) in primary neuroblastoma. Oncogene 10: 1417–1422.

700. St Clair, S. L., H. Li, U. Ashraf, J. A. Karty and J. M. Tennessen, 2017 Metabolomic Analysis Reveals That the Drosophila melanogaster Gene lysine Influences Diverse Aspects of Metabolism. Genetics 207: 1255–1261.

701. Stern, B., G. Ried, N. J. Clegg, T. A. Grigliatti and C. F. Lehner, 1993 Genetic analysis of the Drosophila cdc2 homolog. Development 117: 219–232.

702. Strick, T. R., V. Croquette and D. Bensimon, 2000 Single-molecule analysis of DNA uncoiling by a type II topoisomerase. Nature 404: 901–904.

703. Stroebele, E., and A. Erives, 2016 Integration of Orthogonal Signaling by the Notch and Dpp Pathways in Drosophila. Genetics 203: 219–240.

704. Su, Y. C., J. E. Treisman and E. Y. Skolnik, 1998 The Drosophila Ste20-related kinase misshapen is required for embryonic dorsal closure and acts through a JNK MAPK module on an evolutionarily conserved signaling pathway. Genes Dev 12: 2371–2380.

705. Suganuma, T., A. Mushegian, S. K. Swanson, L. Florens, M. P. Washburn et al., 2012 A metazoan ATAC acetyltransferase subunit that regulates mitogen-activated protein kinase signaling is related to an ancient molybdopterin synthase component. Mol Cell Proteomics 11: 90–99.

706. Suganuma, T., and J. L. Workman, 2008 Crosstalk among Histone Modifications. Cell 135: 604–607.

707. Swarup, S., and E. M. Verheyen, 2011 Drosophila homeodomain-interacting protein kinase inhibits the Skp1-Cul1-F-box E3 ligase complex to dually promote Wingless and Hedgehog signaling. Proc Natl Acad Sci U S A 108: 9887–9892.

708. Symes, A. J., M. Eilertsen, M. Millar, J. Nariculam, A. Freeman et al., 2013 Quantitative analysis of BTF3, HINT1, NDRG1 and ODC1 protein over-expression in human prostate cancer tissue. PLoS One 8: e84295.

709. Szentirmay, M. N., and M. Sawadogo, 1991 Transcription factor requirement for multiple rounds of initiation by human RNA polymerase II. Proc Natl Acad Sci U S A 88: 10691–10695.

710. Takahashi, S., and I. Takada, 2023 Recent advances in prostate cancer: WNT signaling, chromatin regulation, and transcriptional coregulators. Asian J Androl 25: 158–165.

711. Takai, A., T. Chiyonobu, I. Ueoka, R. Tanaka, T. Tozawa et al., 2020 A novel Drosophila model for neurodevelopmental disorders associated with Shwachman-Diamond syndrome. Neurosci Lett 739: 135449.

712. Takayama, M., T. Taira, S. M. Iguchi-Ariga and H. Ariga, 2000 CDC6 interacts with c-Myc to inhibit E-box-dependent transcription by abrogating c-Myc/Max complex. FEBS Lett 477: 43–48.

713. Takemura, M., and H. Nakato, 2017 Drosophila Sulf1 is required for the termination of intestinal stem cell division during regeneration. J Cell Sci 130: 332–343.

714. Takeuchi, H., R. C. Fernandez-Valdivia, D. S. Caswell, A. Nita-Lazar, N. A. Rana et al., 2011 Rumi functions as both a protein O-glucosyltransferase and a protein O-xylosyltransferase. Proc Natl Acad Sci U S A 108: 16600–16605.

715. Taliaferro, J. M., D. Marwha, J. L. Aspden, D. Mavrici, N. E. Cheng et al., 2013 The Drosophila splicing factor PSI is phosphorylated by casein kinase II and tousled-like kinase. PLoS One 8: e56401.

716. Tang, R., W. Huang, J. Guan, Q. Liu, B. T. Beerntsen et al., 2021 Drosophila H2Av negatively regulates the activity of the IMD pathway via facilitating Relish SUMOylation. PLoS Genet 17: e1009718.

717. Taniguchi, K., S. Hozumi, R. Maeda, T. Okumura and K. Matsuno, 2007 Roles of type I myosins in Drosophila handedness. Fly (Austin) 1: 287–290.

718. Tarayrah, L., Y. Li, Q. Gan and X. Chen, 2015 Epigenetic regulator Lid maintains germline stem cells through regulating JAK-STAT signaling pathway activity. Biol Open 4: 1518–1527.

719. Tariq, A., L. Green, J. C. G. Jeynes, C. Soeller and J. G. Wakefield, 2020 In vitro reconstitution of branching microtubule nucleation. Elife 9.

720. Tavares, L., A. Correia, M. A. Santos, J. B. Relvas and P. S. Pereira, 2017 dMyc is required in retinal progenitors to prevent JNK-mediated retinal glial activation. PLoS Genet 13: e1006647.

721. Taylor, J. P., R. H. Brown, Jr. and D. W. Cleveland, 2016 Decoding ALS: from genes to mechanism. Nature 539: 197–206.

722. Terriente-Felix, A., and J. F. de Celis, 2009 Osa, a subunit of the BAP chromatin-remodelling complex, participates in the regulation of gene expression in response to EGFR signalling in the Drosophila wing. Dev Biol 329: 350–361.

723. Teufel, A., T. Maass, S. Strand, S. Kanzler, T. Galante et al., 2010 Liver-specific Ldb1 deletion results in enhanced liver cancer development. J Hepatol 53: 1078–1084.

724. Thao, D. T., H. Seto and M. Yamaguchi, 2008 Drosophila Myc is required for normal DREF gene expression. Exp Cell Res 314: 184–192.

725. Thevenon, D., I. Seffouh, C. Pillet, X. Crespo-Yanez, M. O. Fauvarque et al., 2020 A Nucleolar Isoform of the Drosophila Ubiquitin Specific Protease dUSP36 Regulates MYC-Dependent Cell Growth. Front Cell Dev Biol 8: 506.

726. Thomas, M. C., and C. M. Chiang, 2006 The general transcription machinery and general cofactors. Crit Rev Biochem Mol Biol 41: 105–178.

727. Thompson, E. B., 1998 The many roles of c-Myc in apoptosis. Annu Rev Physiol 60: 575–600.

728. Tokamov, S. A., T. Su, A. Ullyot and R. G. Fehon, 2021 Negative feedback couples Hippo pathway activation with Kibra degradation independent of Yorkie-mediated transcription. Elife 10.

729. Toledano-Katchalski, H., R. Nir, G. Volohonsky and T. Volk, 2007 Post-transcriptional repression of the Drosophila midkine and pleiotrophin homolog miple by HOW is essential for correct mesoderm spreading. Development 134: 3473–3481.

730. Toth, J., M. Kovacs, F. Wang, L. Nyitray and J. R. Sellers, 2005 Myosin V from Drosophila reveals diversity of motor mechanisms within the myosin V family. J Biol Chem 280: 30594–30603.

731. Tran, D. D., S. Saran, A. J. Williamson, A. Pierce, O. Dittrich-Breiholz et al., 2014 THOC5 controls 3’end-processing of immediate early genes via interaction with polyadenylation specific factor 100 (CPSF100). Nucleic Acids Res 42: 12249–12260.

732. Treiber, T., N. Treiber, U. Plessmann, S. Harlander, J. L. Daiss et al., 2017 A Compendium of RNA-Binding Proteins that Regulate MicroRNA Biogenesis. Mol Cell 66: 270–284 e213.

733. Tritto, P., V. Palumbo, L. Micale, M. Marzulli, M. P. Bozzetti et al., 2015 Loss of Pol32 in Drosophila melanogaster causes chromosome instability and suppresses variegation. PLoS One 10: e0120859.

734. Tsai, C. C., H. Y. Kao, A. Mitzutani, E. Banayo, H. Rajan et al., 2004 Ataxin 1, a SCA1 neurodegenerative disorder protein, is functionally linked to the silencing mediator of retinoid and thyroid hormone receptors. Proc Natl Acad Sci U S A 101: 4047–4052.

735. Tsuda, L., M. Kaido, Y. M. Lim, K. Kato, T. Aigaki et al., 2006 An NRSF/REST-like repressor downstream of Ebi/SMRTER/Su(H) regulates eye development in Drosophila. EMBO J 25: 3191–3202.

736. Tsuda, L., R. Nagaraj, S. L. Zipursky and U. Banerjee, 2002 An EGFR/Ebi/Sno pathway promotes delta expression by inactivating Su(H)/SMRTER repression during inductive notch signaling. Cell 110: 625–637.

737. Tu, L. C., X. Yan, L. Hood and B. Lin, 2007 Proteomics analysis of the interactome of N-myc downstream regulated gene 1 and its interactions with the androgen response program in prostate cancer cells. Mol Cell Proteomics 6: 575–588.

738. Turnage, M. A., P. Brewer-Jensen, W. L. Bai and L. L. Searles, 2000 Arginine-rich regions mediate the RNA binding and regulatory activities of the protein encoded by the Drosophila melanogaster suppressor of sable gene. Mol Cell Biol 20: 8198–8208.

739. Tyler, J. K., M. Bulger, R. T. Kamakaka, R. Kobayashi and J. T. Kadonaga, 1996 The p55 subunit of Drosophila chromatin assembly factor 1 is homologous to a histone deacetylase-associated protein. Mol Cell Biol 16: 6149–6159.

740. Uemura, T., K. Shiomi, S. Togashi and M. Takeichi, 1993 Mutation of twins encoding a regulator of protein phosphatase 2A leads to pattern duplication in Drosophila imaginal discs. Genes Dev 7: 429–440.

741. Ukken, F. P., I. Aprill, N. JayaNandanan and M. Leptin, 2014 Slik and the receptor tyrosine kinase Breathless mediate localized activation of Moesin in terminal tracheal cells. PLoS One 9: e103323.

742. Valanne, S., H. Myllymaki, J. Kallio, M. R. Schmid, A. Kleino et al., 2010 Genome-wide RNA interference in Drosophila cells identifies G protein-coupled receptor kinase 2 as a conserved regulator of NF-kappaB signaling. J Immunol 184: 6188–6198.

743. Van Dyke, M. W., M. Sawadogo and R. G. Roeder, 1989 Stability of transcription complexes on class II genes. Mol Cell Biol 9: 342–344.

744. van Meyel, D. J., D. D. O’Keefe, S. Thor, L. W. Jurata, G. N. Gill et al., 2000 Chip is an essential cofactor for apterous in the regulation of axon guidance in Drosophila. Development 127: 1823–1831.

745. van Riggelen, J., A. Yetil and D. W. Felsher, 2010 MYC as a regulator of ribosome biogenesis and protein synthesis. Nat Rev Cancer 10: 301–309.

746. van Tienen, L. M., J. Mieszczanek, M. Fiedler, T. J. Rutherford and M. Bienz, 2017 Constitutive scaffolding of multiple Wnt enhanceosome components by Legless/BCL9. Elife 6.

747. Varga-Weisz, P. D., M. Wilm, E. Bonte, K. Dumas, M. Mann et al., 1997 Chromatin-remodelling factor CHRAC contains the ATPases ISWI and topoisomerase II. Nature 388: 598–602.

748. Vasquez, C. G., M. Tworoger and A. C. Martin, 2014 Dynamic myosin phosphorylation regulates contractile pulses and tissue integrity during epithelial morphogenesis. J Cell Biol 206: 435–450.

749. Verni, F., R. Gandhi, M. L. Goldberg and M. Gatti, 2000 Genetic and molecular analysis of wings apart-like (wapl), a gene controlling heterochromatin organization in Drosophila melanogaster. Genetics 154: 1693–1710.

750. Vichas, A., M. T. Laurie and J. A. Zallen, 2015 The Ski2-family helicase Obelus regulates Crumbs alternative splicing and cell polarity. J Cell Biol 211: 1011–1024.

751. Villani, G., B. Sauer and I. R. Lehman, 1980 DNA polymerase alpha from Drosophila melanogaster embryos. Subunit structure. J Biol Chem 255: 9479–9483.

752. Vinayagam, A., Y. Hu, M. Kulkarni, C. Roesel, R. Sopko et al., 2013 Protein complex-based analysis framework for high-throughput data sets. Sci Signal 6: rs5.

753. Voelker, R. A., W. Gibson, J. P. Graves, J. F. Sterling and M. T. Eisenberg, 1991 The Drosophila suppressor of sable gene encodes a polypeptide with regions similar to those of RNA-binding proteins. Mol Cell Biol 11: 894–905.

754. Volohonsky, G., G. Edenfeld, C. Klambt and T. Volk, 2007 Muscle-dependent maturation of tendon cells is induced by post-transcriptional regulation of stripeA. Development 134: 347–356.

755. Vorobyeva, N. E., N. V. Soshnikova, J. L. Kuzmina, M. R. Kopantseva, J. V. Nikolenko et al., 2009 The novel regulator of metazoan development SAYP organizes a nuclear coactivator supercomplex. Cell Cycle 8: 2152–2156.

756. Vriz, S., M. Taylor and M. Mechali, 1989 Differential expression of two Xenopus c-myc proto-oncogenes during development. EMBO J 8: 4091–4097.

757. Vuilleumier, R., M. Miao, S. Medina-Giro, C. M. Ell, S. Flibotte et al., 2022 Dichotomous cis-regulatory motifs mediate the maturation of the neuromuscular junction by retrograde BMP signaling. Nucleic Acids Res 50: 9748–9764.

758. Vyas, S., C. Bechade, B. Riveau, J. Downward and A. Triller, 2002 Involvement of survival motor neuron (SMN) protein in cell death. Hum Mol Genet 11: 2751–2764.

759. Wagner, C. R., K. Hamana and S. C. Elgin, 1992 A high-mobility-group protein and its cDNAs from Drosophila melanogaster. Mol Cell Biol 12: 1915–1923.

760. Wakefield, L. M., and C. Stuelten, 2007 Keeping order in the neighborhood: new roles for TGFbeta in maintaining epithelial homeostasis. Cancer Cell 12: 293–295.

761. Walsh, E. P., and N. H. Brown, 1998 A screen to identify Drosophila genes required for integrin-mediated adhesion. Genetics 150: 791–805.

762. Wan, M., J. Liang, Y. Xiong, F. Shi, Y. Zhang et al., 2013 The trithorax group protein Ash2l is essential for pluripotency and maintaining open chromatin in embryonic stem cells. J Biol Chem 288: 5039–5048.

763. Wang, C., Z. Liu and X. Huang, 2012 Rab32 is important for autophagy and lipid storage in Drosophila. PLoS One 7: e32086.

764. Wang, G., K. Amanai, B. Wang and J. Jiang, 2000 Interactions with Costal2 and suppressor of fused regulate nuclear translocation and activity of cubitus interruptus. Genes Dev 14: 2893–2905.

765. Wang, H., C. Dienemann, A. Stutzer, H. Urlaub, A. C. M. Cheung et al., 2020 Structure of the transcription coactivator SAGA. Nature 577: 717–720.

766. Wang, J., M. Michel, L. Bialas, G. Pierini and C. Dahmann, 2023 Preferential recruitment and stabilization of Myosin II at compartment boundaries in Drosophila. J Cell Sci 136.

767. Wang, J. W., J. R. Brent, A. Tomlinson, N. A. Shneider and B. D. McCabe, 2011 The ALS-associated proteins FUS and TDP-43 function together to affect Drosophila locomotion and life span. J Clin Invest 121: 4118–4126.

768. Wang, L., G. Lam and C. S. Thummel, 2010 Med24 and Mdh2 are required for Drosophila larval salivary gland cell death. Dev Dyn 239: 954–964.

769. Wang, M., M. Ly, A. Lugowski, J. D. Laver, H. D. Lipshitz et al., 2017 ME31B globally represses maternal mRNAs by two distinct mechanisms during the Drosophila maternal-to-zygotic transition. Elife 6.

770. Wang, X., S. K. Hansen, R. Ratts, S. Zhou, A. J. Snook et al., 1997 Drosophila TFIIE: purification, cloning, and functional reconstitution. Proc Natl Acad Sci U S A 94: 433–438.

771. Wang, X., X. Huang, C. Wu and L. Xue, 2018a Pontin/Tip49 acts as a novel regulator of JNK pathway. Cell Death Dis 9: 978.

772. Wang, X., X. Huang, C. Wu and L. Xue, 2018b Pontin/Tip49 negatively regulates JNK-mediated cell death in Drosophila. Cell Death Discov 4: 8.

773. Wang, Y. L., S. H. Duttke, K. Chen, J. Johnston, G. A. Kassavetis et al., 2014 TRF2, but not TBP, mediates the transcription of ribosomal protein genes. Genes Dev 28: 1550–1555.

774. Wang, Z., S. Buratowski, J. Q. Svejstrup, W. J. Feaver, X. Wu et al., 1995 The yeast TFB1 and SSL1 genes, which encode subunits of transcription factor IIH, are required for nucleotide excision repair and RNA polymerase II transcription. Mol Cell Biol 15: 2288–2293.

775. Wang, Z. H., Y. Liu, V. Chaitankar, M. Pirooznia and H. Xu, 2019 Electron transport chain biogenesis activated by a JNK-insulin-Myc relay primes mitochondrial inheritance in Drosophila. Elife 8.

776. Weake, V. M., J. O. Dyer, C. Seidel, A. Box, S. K. Swanson et al., 2011 Post-transcription initiation function of the ubiquitous SAGA complex in tissue-specific gene activation. Genes Dev 25: 1499–1509.

777. Weake, V. M., S. K. Swanson, A. Mushegian, L. Florens, M. P. Washburn et al., 2009 A novel histone fold domain-containing protein that replaces TAF6 in Drosophila SAGA is required for SAGA-dependent gene expression. Genes Dev 23: 2818–2823.

778. Wei, Y., D. Resetca, Z. Li, I. Johansson-Akhe, A. Ahlner et al., 2019 Multiple direct interactions of TBP with the MYC oncoprotein. Nat Struct Mol Biol 26: 1035–1043.

779. Weiss, A., E. Charbonnier, E. Ellertsdottir, A. Tsirigos, C. Wolf et al., 2010 A conserved activation element in BMP signaling during Drosophila development. Nat Struct Mol Biol 17: 69–76.

780. Wickens, M., D. S. Bernstein, J. Kimble and R. Parker, 2002 A PUF family portrait: 3’UTR regulation as a way of life. Trends Genet 18: 150–157.

781. Wiersdorff, V., T. Lecuit, S. M. Cohen and M. Mlodzik, 1996 Mad acts downstream of Dpp receptors, revealing a differential requirement for dpp signaling in initiation and propagation of morphogenesis in the Drosophila eye. Development 122: 2153–2162.

782. Willemsen, M. H., B. Nijhof, M. Fenckova, W. M. Nillesen, E. M. Bongers et al., 2013 GATAD2B loss-of-function mutations cause a recognisable syndrome with intellectual disability and are associated with learning deficits and synaptic undergrowth in Drosophila. J Med Genet 50: 507–514.

783. Willert, K., M. Brink, A. Wodarz, H. Varmus and R. Nusse, 1997 Casein kinase 2 associates with and phosphorylates dishevelled. EMBO J 16: 3089–3096.

784. Willy, P. J., R. Kobayashi and J. T. Kadonaga, 2000 A basal transcription factor that activates or represses transcription. Science 290: 982–985.

785. Windler, S. L., and D. Bilder, 2010 Endocytic internalization routes required for delta/notch signaling. Curr Biol 20: 538–543.

786. Wolf, D., T. K. Smylla, J. Reichmuth, P. Hoffmeister, L. Kober et al., 2019 Nucleo-cytoplasmic shuttling of Drosophila Hairless/Su(H) heterodimer as a means of regulating Notch dependent transcription. Biochim Biophys Acta Mol Cell Res 1866: 1520–1532.

787. Wolfe, A. L., K. Singh, Y. Zhong, P. Drewe, V. K. Rajasekhar et al., 2014 RNA G-quadruplexes cause eIF4A-dependent oncogene translation in cancer. Nature 513: 65–70.

788. Wolfe, S. A., L. Nekludova and C. O. Pabo, 2000 DNA recognition by Cys2His2 zinc finger proteins. Annu Rev Biophys Biomol Struct 29: 183–212.

789. Wong, J. J., S. Li, E. K. Lim, Y. Wang, C. Wang et al., 2013 A Cullin1-based SCF E3 ubiquitin ligase targets the InR/PI3K/TOR pathway to regulate neuronal pruning. PLoS Biol 11: e1001657.

790. Wong, K. H., Y. Jin and K. Struhl, 2014 TFIIH phosphorylation of the Pol II CTD stimulates mediator dissociation from the preinitiation complex and promoter escape. Mol Cell 54: 601–612.

791. Wong, L. C., A. Costa, I. McLeod, A. Sarkeshik, J. Yates, 3rd et al., 2011 The functioning of the Drosophila CPEB protein Orb is regulated by phosphorylation and requires casein kinase 2 activity. PLoS One 6: e24355.

792. Woolworth, J. A., G. Nallamothu and T. Hsu, 2009 The Drosophila metastasis suppressor gene Nm23 homolog, awd, regulates epithelial integrity during oogenesis. Mol Cell Biol 29: 4679–4690.

793. Worringer, K. A., and B. Panning, 2007 Zinc finger protein Zn72D promotes productive splicing of the maleless transcript. Mol Cell Biol 27: 8760–8769.

794. Wright, K. J., M. T. Marr, 2nd and R. Tjian, 2006 TAF4 nucleates a core subcomplex of TFIID and mediates activated transcription from a TATA-less promoter. Proc Natl Acad Sci U S A 103: 12347-12352.

795. Wu, D., L. Wu, H. An, H. Bao, P. Guo et al., 2019 RanGAP-mediated nucleocytoplasmic transport of Prospero regulates neural stem cell lifespan in Drosophila larval central brain. Aging Cell 18: e12854.

796. Wu, D. C., and L. A. Johnston, 2010 Control of wing size and proportions by Drosophila myc. Genetics 184: 199–211.

797. Wu, J., C. Capp, L. Feng and T. S. Hsieh, 2008 Drosophila homologue of the Rothmund-Thomson syndrome gene: essential function in DNA replication during development. Dev Biol 323: 130–142.

798. Xia, P., S. Wang, G. Huang, P. Zhu, M. Li et al., 2014 WASH is required for the differentiation commitment of hematopoietic stem cells in a c-Myc-dependent manner. J Exp Med 211: 2119–2134.

799. Xia, R., Y. Liu, L. Yang, J. Gal, H. Zhu et al., 2012 Motor neuron apoptosis and neuromuscular junction perturbation are prominent features in a Drosophila model of Fus-mediated ALS. Mol Neurodegener 7: 10.

800. Xu, J., K. Hopkins, L. Sabin, A. Yasunaga, H. Subramanian et al., 2013 ERK signaling couples nutrient status to antiviral defense in the insect gut. Proc Natl Acad Sci U S A 110: 15025–15030.

801. Xu, X. Z., P. D. Wes, H. Chen, H. S. Li, M. Yu et al., 1998 Retinal targets for calmodulin include proteins implicated in synaptic transmission. J Biol Chem 273: 31297–31307.

802. Xu, Y., Z. Lei, H. Huang, W. Dui, X. Liang et al., 2009 dRecQ4 is required for DNA synthesis and essential for cell proliferation in Drosophila. PLoS One 4: e6107.

803. Xue, C., D. M. Yu, S. Gherardi, J. Koach, G. Milazzo et al., 2016 MYCN promotes neuroblastoma malignancy by establishing a regulatory circuit with transcription factor AP4. Oncotarget 7: 54937–54951.

804. Yager, J., S. Richards, D. S. Hekmat-Scafe, D. D. Hurd, V. Sundaresan et al., 2001 Control of Drosophila perineurial glial growth by interacting neurotransmitter-mediated signaling pathways. Proc Natl Acad Sci U S A 98: 10445–10450.

805. Yamashita, T., A. D. Agulnick, N. G. Copeland, D. J. Gilbert, N. A. Jenkins et al., 1998 Genomic structure and chromosomal localization of the mouse LIM domain-binding protein 1 gene, Ldb1. Genomics 48: 87-92.

806. Yan, D., and N. Perrimon, 2015 spenito is required for sex determination in Drosophila melanogaster. Proc Natl Acad Sci U S A 112: 11606–11611.

807. Yang, L., S. Paul, K. G. Trieu, L. G. Dent, F. Froldi et al., 2016 Minibrain and Wings apart control organ growth and tissue patterning through down-regulation of Capicua. Proc Natl Acad Sci U S A 113: 10583–10588.

808. Yang, Y., D. A. Primrose, A. C. Leung, R. B. Fitzsimmons, M. C. McDermand et al., 2012 The PP1 phosphatase flapwing regulates the activity of Merlin and Moesin in Drosophila. Dev Biol 361: 412–426.

809. Yano, M., S. Nakamuta, X. Wu, Y. Okumura and H. Kido, 2006 A novel function of 14-3-3 protein: 14-3-3zeta is a heat-shock-related molecular chaperone that dissolves thermal-aggregated proteins. Mol Biol Cell 17: 4769–4779.

810. Yao, B., Y. Li, Z. Wang, L. Chen, M. Poidevin et al., 2018a Active N(6)-Methyladenine Demethylation by DMAD Regulates Gene Expression by Coordinating with Polycomb Protein in Neurons. Mol Cell 71: 848–857 e846.

811. Yao, J., M. B. Ardehali, C. J. Fecko, W. W. Webb and J. T. Lis, 2007 Intranuclear distribution and local dynamics of RNA polymerase II during transcription activation. Mol Cell 28: 978–990.

812. Yao, L. C., I. L. Blitz, D. A. Peiffer, S. Phin, Y. Wang et al., 2006 Schnurri transcription factors from Drosophila and vertebrates can mediate Bmp signaling through a phylogenetically conserved mechanism. Development 133: 4025–4034.

813. Yao, W. D., and C. F. Wu, 2001 Distinct roles of CaMKII and PKA in regulation of firing patterns and K(+) currents in Drosophila neurons. J Neurophysiol 85: 1384–1394.

814. Yao, Y., X. Li, W. Wang, Z. Liu, J. Chen et al., 2018b MRT, Functioning with NURF Complex, Regulates Lipid Droplet Size. Cell Rep 24: 2972–2984.

815. Yau, T. Y., O. Molina and A. J. Courey, 2020 SUMOylation in development and neurodegeneration. Development 147.

816. Yavatkar, A. S., Y. Lin, J. Ross, Y. Fann, T. Brody et al., 2008 Rapid detection and curation of conserved DNA via enhanced-BLAT and EvoPrinterHD analysis. BMC Genomics 9: 106.

817. Yeh, E., M. Cunningham, H. Arnold, D. Chasse, T. Monteith et al., 2004 A signalling pathway controlling c-Myc degradation that impacts oncogenic transformation of human cells. Nat Cell Biol 6: 308–318.

818. Yeh, E. S., B. O. Lew and A. R. Means, 2006 The loss of PIN1 deregulates cyclin E and sensitizes mouse embryo fibroblasts to genomic instability. J Biol Chem 281: 241–251.

819. Yeh, T. H., S. Y. Huang, W. Y. Lan, G. J. Liaw and J. Y. Yu, 2015 Modulation of cell morphogenesis by tousled-like kinase in the Drosophila follicle cell. Dev Dyn 244: 852–865.

820. Yeom, E., S. T. Hong and K. W. Choi, 2015 Crumbs interacts with Xpd for nuclear division control in Drosophila. Oncogene 34: 2777–2789.

821. Yokomori, K., J. L. Chen, A. Admon, S. Zhou and R. Tjian, 1993 Molecular cloning and characterization of dTAFII30 alpha and dTAFII30 beta: two small subunits of Drosophila TFIID. Genes Dev 7: 2587–2597.

822. Yokoshi, M., K. Kawasaki, M. Cambon and T. Fukaya, 2022 Dynamic modulation of enhancer responsiveness by core promoter elements in living Drosophila embryos. Nucleic Acids Res 50: 92–107.

823. Yoon, J. W., M. Gallant, M. L. Lamm, S. Iannaccone, K. F. Vieux et al., 2013 Noncanonical regulation of the Hedgehog mediator GLI1 by c-MYC in Burkitt lymphoma. Mol Cancer Res 11: 604–615.

824. Yu, H., H. Takeuchi, M. Takeuchi, Q. Liu, J. Kantharia et al., 2016 Structural analysis of Notch-regulating Rumi reveals basis for pathogenic mutations. Nat Chem Biol 12: 735–740.

825. Yuen, H. F., V. K. Gunasekharan, K. K. Chan, S. D. Zhang, A. Platt-Higgins et al., 2013 RanGTPase: a candidate for Myc-mediated cancer progression. J Natl Cancer Inst 105: 475–488.

826. Zacarias-Fluck, M. F., L. Soucek and J. R. Whitfield, 2024 MYC: there is more to it than cancer. Front Cell Dev Biol 12: 1342872.

827. Zaessinger, S., I. Busseau and M. Simonelig, 2006 Oskar allows nanos mRNA translation in Drosophila embryos by preventing its deadenylation by Smaug/CCR4. Development 133: 4573–4583.

828. Zaman, S., K. Sukhodolets, P. Wang, J. Qin, D. Levens et al., 2014 FBP1 Is an Interacting Partner of Menin. Int J Endocrinol 2014: 535401.

829. Zanet, J., S. Pibre, C. Jacquet, A. Ramirez, I. M. de Alboran et al., 2005 Endogenous Myc controls mammalian epidermal cell size, hyperproliferation, endoreplication and stem cell amplification. J Cell Sci 118: 1693–1704.

830. Zavortink, M., L. N. Rutt, S. Dzitoyeva, J. C. Henriksen, C. Barrington et al., 2020 The E2 Marie Kondo and the CTLH E3 ligase clear deposited RNA binding proteins during the maternal-to-zygotic transition. Elife 9.

831. Zaytseva, O., and L. M. Quinn, 2017 Controlling the Master: Chromatin Dynamics at the MYC Promoter Integrate Developmental Signaling. Genes (Basel) 8.

832. Zehring, W. A., J. M. Lee, J. R. Weeks, R. S. Jokerst and A. L. Greenleaf, 1988 The C-terminal repeat domain of RNA polymerase II largest subunit is essential in vivo but is not required for accurate transcription initiation in vitro. Proc Natl Acad Sci U S A 85: 3698–3702.

833. Zeitler, J., C. P. Hsu, H. Dionne and D. Bilder, 2004 Domains controlling cell polarity and proliferation in the Drosophila tumor suppressor Scribble. J Cell Biol 167: 1137–1146.

834. Zeke, A., M. Misheva, A. Remenyi and M. A. Bogoyevitch, 2016 JNK Signaling: Regulation and Functions Based on Complex Protein-Protein Partnerships. Microbiol Mol Biol Rev 80: 793–835.

835. Zeng, X., S. R. Singh, D. Hou and S. X. Hou, 2010 Tumor suppressors Sav/Scrib and oncogene Ras regulate stem-cell transformation in adult Drosophila malpighian tubules. J Cell Physiol 224: 766–774.

836. Zenvirt, S., Y. Nevo-Caspi, S. Rencus-Lazar and D. Segal, 2008 Drosophila LIM-only is a positive regulator of transcription during thoracic bristle development. Genetics 179: 1989–1999.

837. Zhan, W., X. Liao, Y. Wang, L. Li, J. Li et al., 2019 circCTIC1 promotes the self-renewal of colon TICs through BPTF-dependent c-Myc expression. Carcinogenesis 40: 560–568.

838. Zhang, J., Y. Liu, K. Jiang and J. Jia, 2017 SUMO regulates the activity of Smoothened and Costal-2 in Drosophila Hedgehog signaling. Sci Rep 7: 42749.

839. Zhang, P., Y. Wu, T. Y. Belenkaya and X. Lin, 2011 SNX3 controls Wingless/Wnt secretion through regulating retromer-dependent recycling of Wntless. Cell Res 21: 1677–1690.

840. Zhang, X., J. Wang, Y. Jia, T. Liu, M. Wang et al., 2019 CDK5 neutralizes the tumor suppressing effect of BIN1 via mediating phosphorylation of c-MYC at Ser-62 site in NSCLC. Cancer Cell Int 19: 226.

841. Zhang, Y., R. Cai, R. Zhou, Y. Li and L. Liu, 2016 Tousled-like kinase mediated a new type of cell death pathway in Drosophila. Cell Death Differ 23: 146–157.

842. Zhang, Y. W., and D. N. Arnosti, 2011 Conserved catalytic and C-terminal regulatory domains of the C-terminal binding protein corepressor fine-tune the transcriptional response in development. Mol Cell Biol 31: 375–384.

843. Zhao, L. J., P. M. Loewenstein and M. Green, 2018 Identification of a panel of MYC and Tip60 co-regulated genes functioning primarily in cell cycle and DNA replication. Genes Cancer 9: 101–113.

844. Zheng, Y., J. Wildonger, B. Ye, Y. Zhang, A. Kita et al., 2008 Dynein is required for polarized dendritic transport and uniform microtubule orientation in axons. Nat Cell Biol 10: 1172–1180.

845. Zhou, H., I. Grubisic, K. Zheng, Y. He, P. J. Wang et al., 2013 Taf7l cooperates with Trf2 to regulate spermiogenesis. Proc Natl Acad Sci U S A 110: 16886–16891.

846. Zhou, Z. Q., and P. J. Hurlin, 2001 The interplay between Mad and Myc in proliferation and differentiation. Trends Cell Biol 11: S10–14.

847. Zhu, J. Y., J. van de Leemput and Z. Han, 2024 Distinct roles of COMPASS subunits to Drosophila heart development. Biol Open 13.

848. Zhu, W., M. Foehr, J. B. Jaynes and S. D. Hanes, 2001 Drosophila SAP18, a member of the Sin3/Rpd3 histone deacetylase complex, interacts with Bicoid and inhibits its activity. Dev Genes Evol 211: 109–117.

849. Zoranovic, T., L. Grmai and E. A. Bach, 2013 Regulation of proliferation, cell competition, and cellular growth by the Drosophila JAK-STAT pathway. JAKSTAT 2: e25408.

