## Supplemental Figures for "Targeting Regulatory Factors Associated with the *Drosophila Myc cis*-Elements by Reporter Expression, Gel Shift Assay, and Mass Spectrometric Protein Identification"

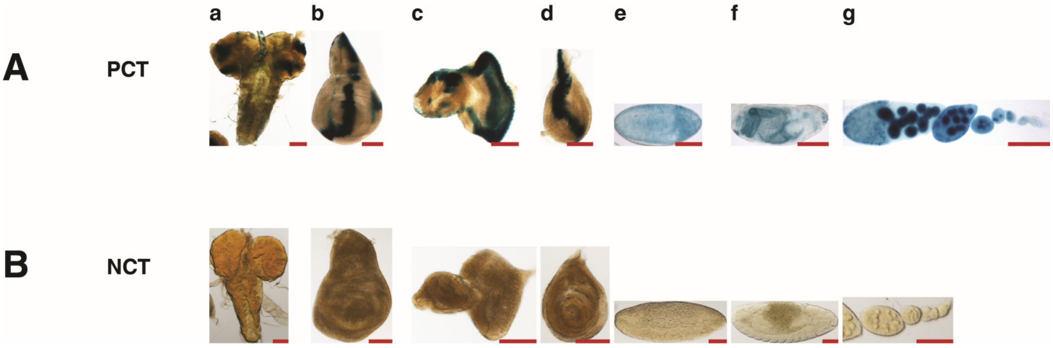


**Figure S1**. **β-Galactosidase (β-Gal) staining of positive and negative controls**. Each staining of transgenic tissues (Figures 1 and 2) was accompanied by positive and negative control staining. Tissues taken from 3rd instar larvae and adult females of the dpp-lacZ fly strain were used as positive control (A, PCT), and the "*y[1] w[1118]*" flies served as negative control (B, NCT). **Abbreviations**: **a**, brain; **b**, wing disc; **c**, eye-antennal disc, **d**, leg disc; **e** and **f**, embryos, **g**; ovary. Scale bar in (**a–g**) indicates 100 μm.

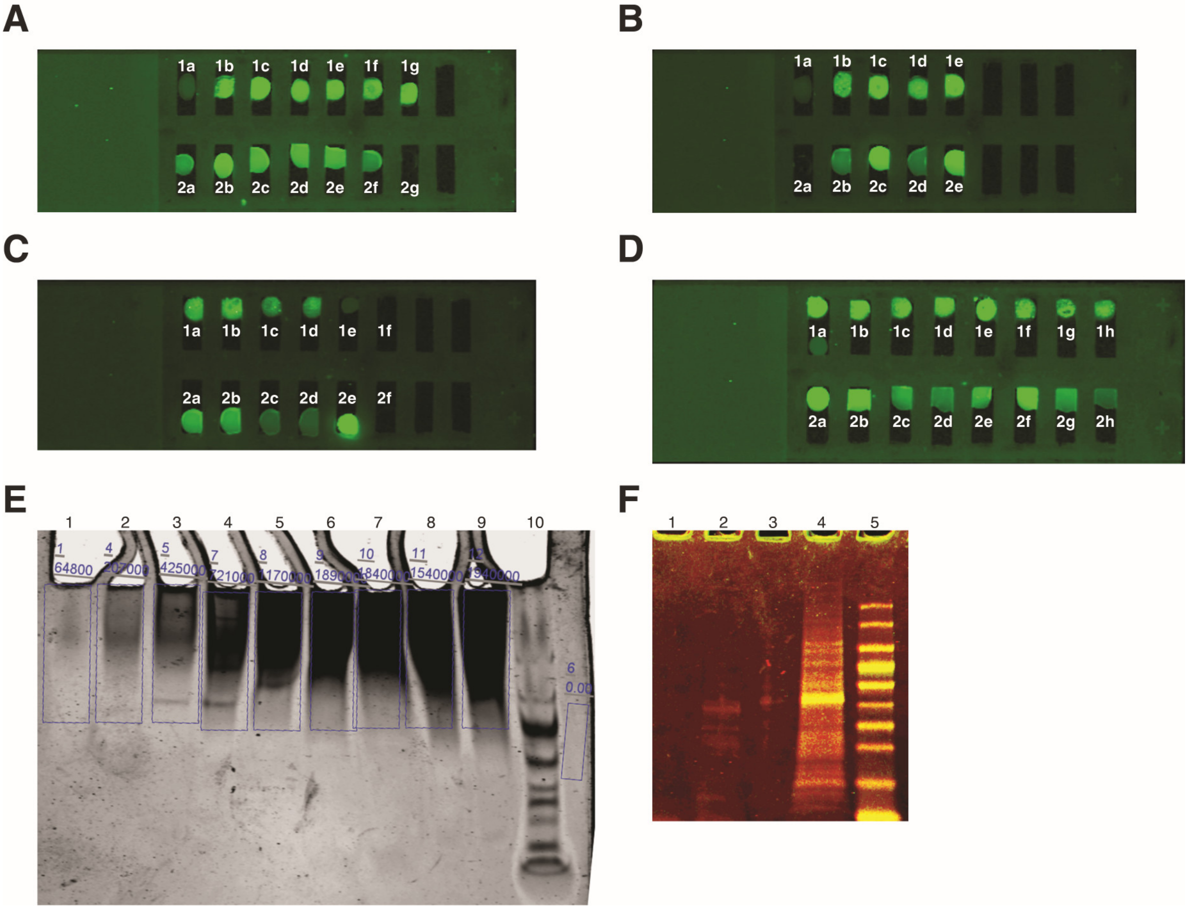


**Figure S2**. **Magnetic Beads (blank beads), Bead-Peg23-DNA constructs, and Soluble Nuclear Fraction (SNF) were tested before sample preparation for protein identification**. **(A-D): Enrichment constructs Bead-Peg23-DNA (IR-labeled) on glass slide before protein binding, A: 1a**: Raw beads 30 μL, **1b/1c**: Bead-Peg23-IR-Pos1/Pos2, **1d/1g**: Bead-Peg23-IR-P1/P2, **2a**: Supernatant of 1b, **2b**: Supernatant of 1c, **2c**: Supernatant of 1d, **2d**: Supernatant of 1e, **2e**: Supernatant of 1f, **2f**: Supernatant of 1g, **2g**: Binding buffer 30 μL (lanes **1b-1g**: each 3 μL of 0.3 pmol input/300 μL, supernatants **2a-2f**: each 30 μL of 0.3 pmol input/300 μL); **B: 1a**: Raw beads only 30 μL, **1b-1e**: Bead-Peg23-IR-Scm1/Scm2, **2a**: Binding buffer only: 30 μL, **2b**: Supernatant of 1b, **2c**: Supernatant of 1c, **2d**: Supernatant of 1d, **2e**: Supernatant of 1e (lanes **1b-1e**: each 3 μL of 0.3 pmol input/300 μL, supernatants **2b-2e**: each 30 μL of 0.3 pmol input/300 μL); **C: 1a-1d**: Bead-Peg23-IR-P29/P30, **1e**: Raw beads only 30 μL, **1f**: Binding buffer 30 μL, **2a**: Supernatant of 1a, **2b**: Supernatant of 1b, **2c**: Supernatant of 1c, **2d**: Supernatant of 1d, **2e**: IR5-P29/P30-IR3 only, **2f**: Storage buffer only 30 μL (lanes **1a-1d**: each 3 μL of 0.3 pmol input/300 μL, supernatants **2a-2d**: each 30 μL of 0.3 pmol input/300 μL, **2e:** 0.3 pmol); **D: 1a top & 1b-1d**: Bead-Peg-IR-P31/P32, **1a**

**bottom**: Raw beads only 30 µL, **1e-1h**: Bead-Peg-IR-P35/P36, **2a-2h**: supernatants of **1a top-1h** accordingly (lanes **1a top-1h**: each 3 µL of 0.3 pmol input/300 µL, supernatants of **1a top-1h**: each 30 µL of 0.3 pmol input/300 µL); **E**: Quality & quantity test of SNF with increasing concentrations in non-denaturing conditions (low salt 0.5% TBE running buffer & non-denaturing 5% Precast PAGE gel; SNF concentration: **1**: 10µg; **2**: 15µg; **3**: 30µg, **4**: 50µg; **5**: 60µg; **6**: 70µg; **7**: 80µg **8**: 90µg **9**: 100µg; **10**: 5µl of PageRuler Protein Ladder Thermo Fisher Cat# 26616). **F:** Magnetic Beads contamination test, **1**: Beads: Laemmli treatment without urea addition, **2**: Beads: Laemmli & 8M urea treatment, **3**: carry over of lane4, **4**: 50µg SNF, **5**: 5µl of PageRuler Protein Ladder Thermo Fisher Cat# 26616), (**1 & 2** 100µg Beads). Electrophoresis conditions: run on 5% denaturing SDS PAGE gel for 50 minutes at voltage 90. **Note**: Artifactual bands of ~15 - 40 kDa seen in lane 2 might have leeched off magnetic beads through boiling and urea treatment. However, the bands are very weak (Bovine Serum Albumin (BSA) "fraction V") contains 583 residues with a Molecular Weight of 66.5 kDa; after protein identification the only contaminant unique to the blank beads turned out to be Collagen alpha I(III) chain, Accession P04258, 1466 residues and MW 138.4 kDa.

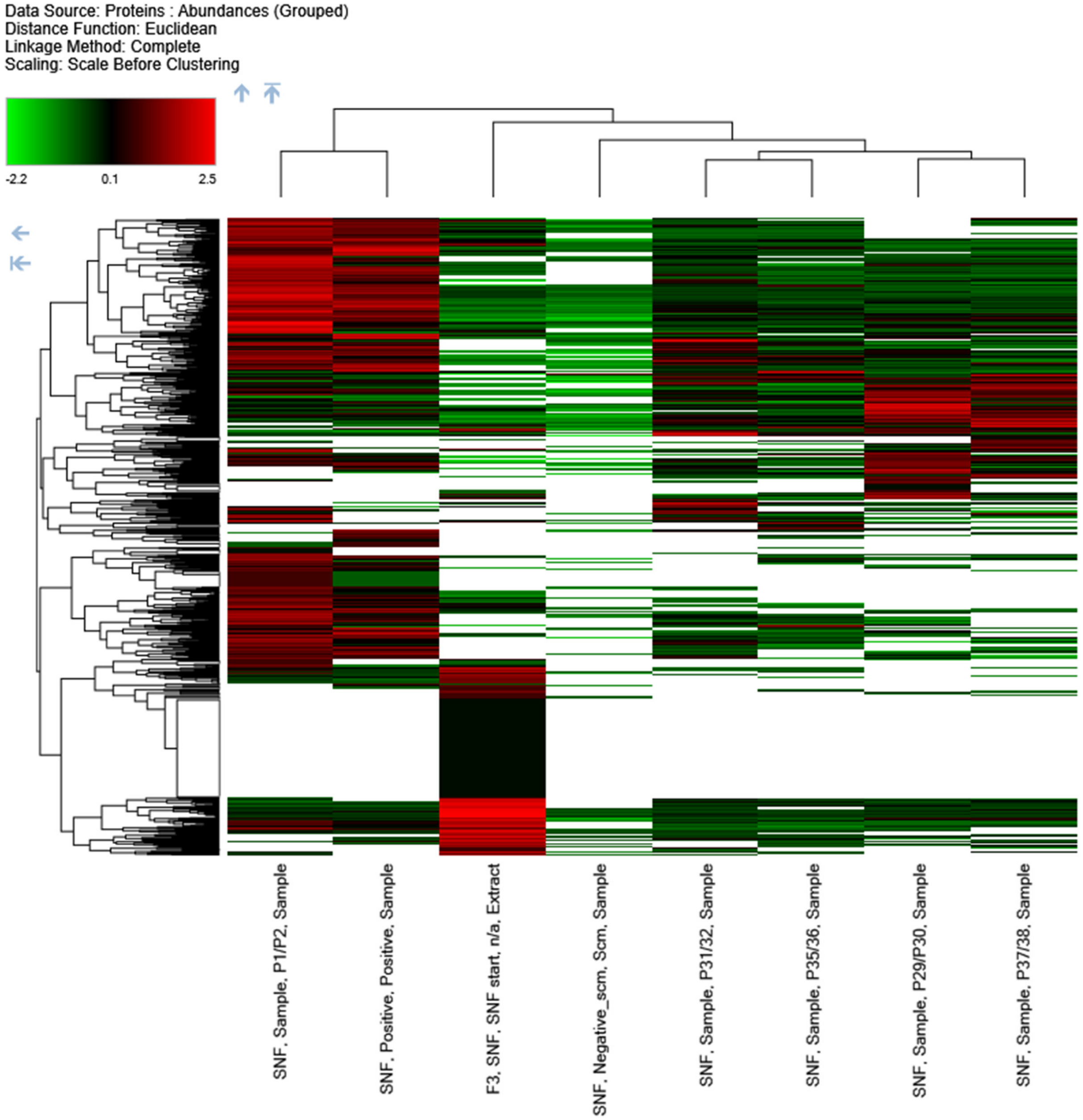


**Figure S3**. **Heat Map of identified proteins using hierarchical clustering log2 Label-Free identification ratios of 1001 quality filtered proteins**. The identified proteins originate from *Drosophila* embryonic nuclear extracts (SNF), 0-72 hours After Egg laying (AEL). The final applied quality filter was"SNF_10x greater than Scm" for all samples and the positive control and displayed are only the hierarchically clustered proteins of each sample group based on Euclidean neighbor joining distance similarity (KEY 2012) The applied filter included "the mass over charge ratio for the sample is 10 times higher than for the Scrambled negative control" (Scm) to minimize False Detection Rate (FDR) and optimize the confidence of identified values. Each sample and control included 2 replicates; in one double-stranded oligo pair the "top" strand and in the other double-stranded pair the "bottom" strand was attached to the Bead-PEG23 solid surface. This strategy helped to retain the proteins associated with either strand (attached or not attached to the bead) through the washing processes. In this Heat Map the oligo pairs are grouped. For example, the Bead-PEG23-IR-P1/P2 is grouped with the Bead-PEG23-IR-P2/P1. Graphically detected overall patterns in the data show P1/P2 & Pos1/Pos2, P31/P32 & P35/P36 and P29/P30 & P37/P38 as relative and similar groups which indicates each two samples in one relative group exhibits a similar expression pattern and biological function. For example, P1/P2 & Pos1/Pos2 might function mainly as tissue-specific late enhancers, P31/P32 & P35/P36 as promoters and P29/P30 & P37/P38 as developmental super enhancers during development.

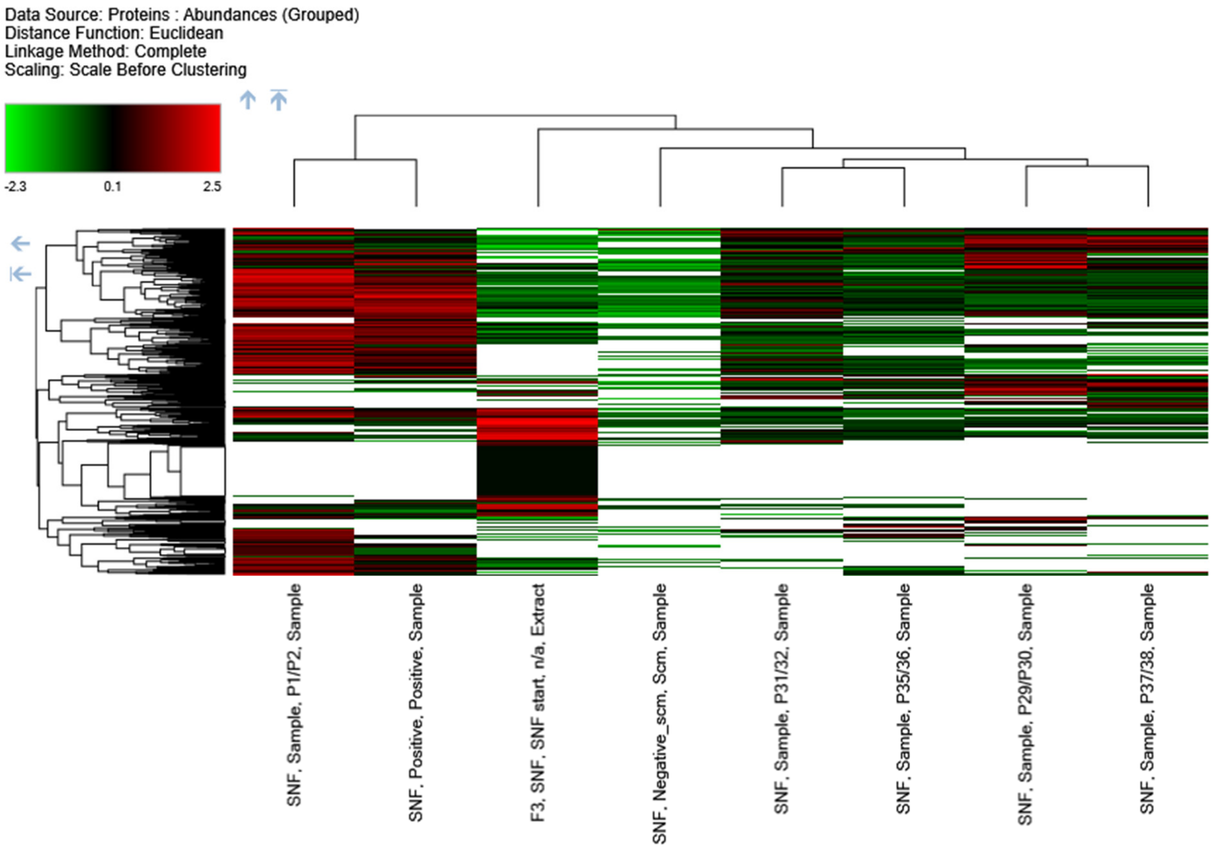


**Figure S4**. **Heat Map of the proteomics data after application of the "SNF_2x greater than Scm" quality filter to Myc samples, positive and negative controls, and the crude nuclear extract:** In this Heat Map "the mass over charge ratio for samples is 2 times higher than for the Scrambled negative control". With this quality filter functionally similar and related groups are in vicinity to each other, and the sample pairs are grouped as is the case in Figure S3.

Key, M., 2012 A tutorial in displaying mass spectrometry-based proteomic data using heat maps. BMC Bioinformatics 13 Suppl 16: S10.
