## Supplemental Tables for "Targeting Regulatory Factors Associated with the *Drosophila Myc cis*-Elements by Reporter Expression, Gel Shift Assay, and Mass Spectrometric Protein Identification": Image 2.pdf

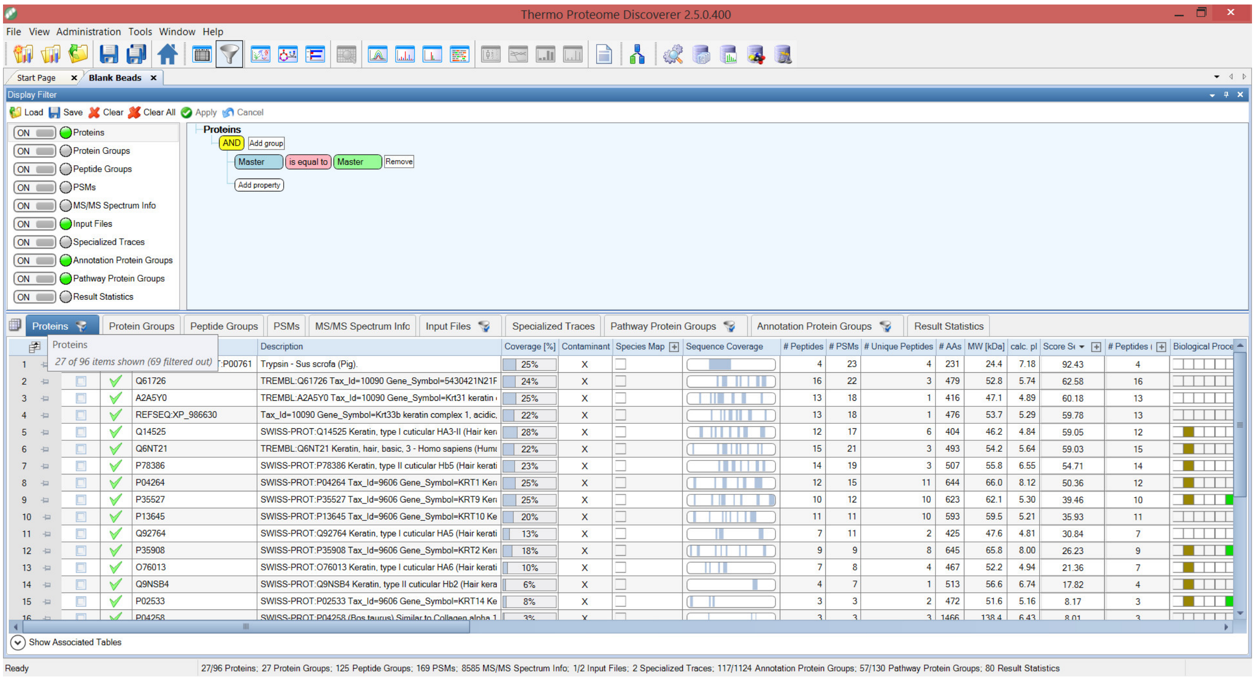

**Image 2. Analysis of the blank (raw) magnetic beads.** Depicted is a Print Screen of the filter used for the raw beads. We applied the same quality control filter 'Master is equal to Master' to the proteomics result obtained for the Dynabeads® MyOne™ Carboxylic Acid (Invitrogen by Thermo Fisher cat. #65001) as we did for the other samples. We identified 26 proteins annotated to *D melanogaster* that were associated with the raw beads. For the complete list of the proteins see Table 2 in File S2.
