## Supplemental Tables for "Targeting Regulatory Factors Associated with the *Drosophila Myc cis*-Elements by Reporter Expression, Gel Shift Assay, and Mass Spectrometric Protein Identification": Image 3.pdf

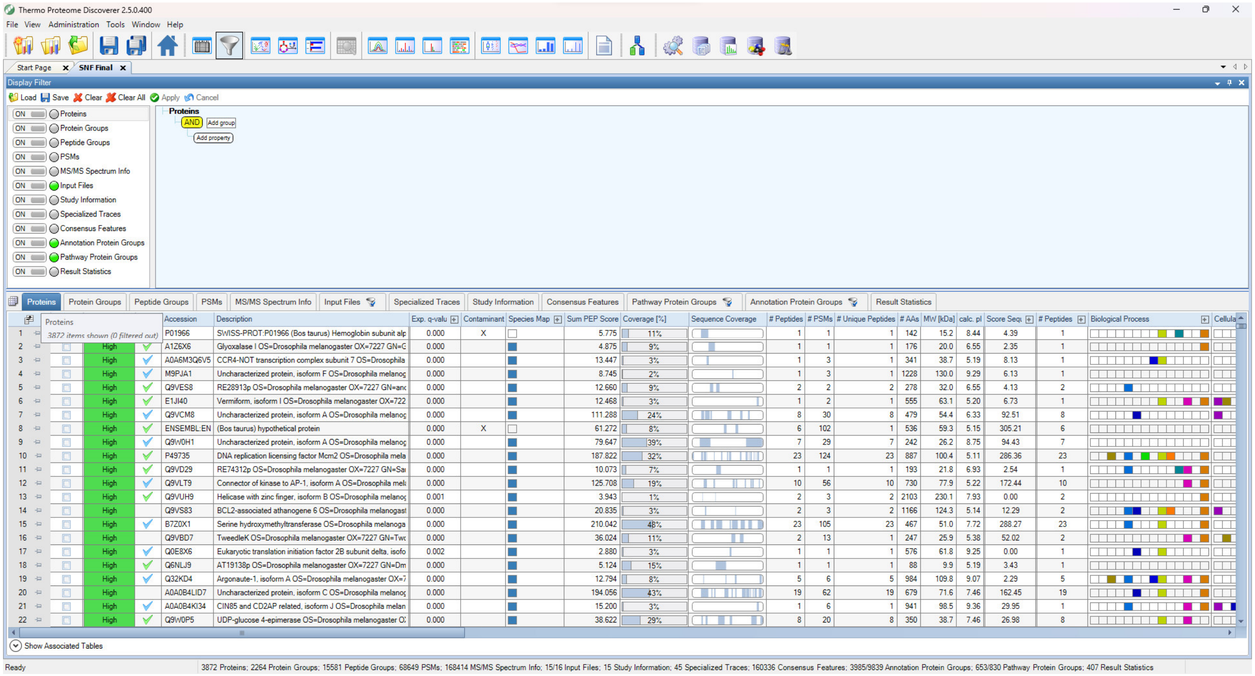


**Image S3. No filter was applied to the original dataset.** The screenshot displays the original unbiased data file with no filter applied. The original Excel file with the replicates ungrouped contained 3872 proteins. The contaminants have not been excluded. For the complete list of the proteins see Table 3 in File S3.
