## Supplemental Tables for "Targeting Regulatory Factors Associated with the *Drosophila Myc cis*-Elements by Reporter Expression, Gel Shift Assay, and Mass Spectrometric Protein Identification": Image 4.pdf

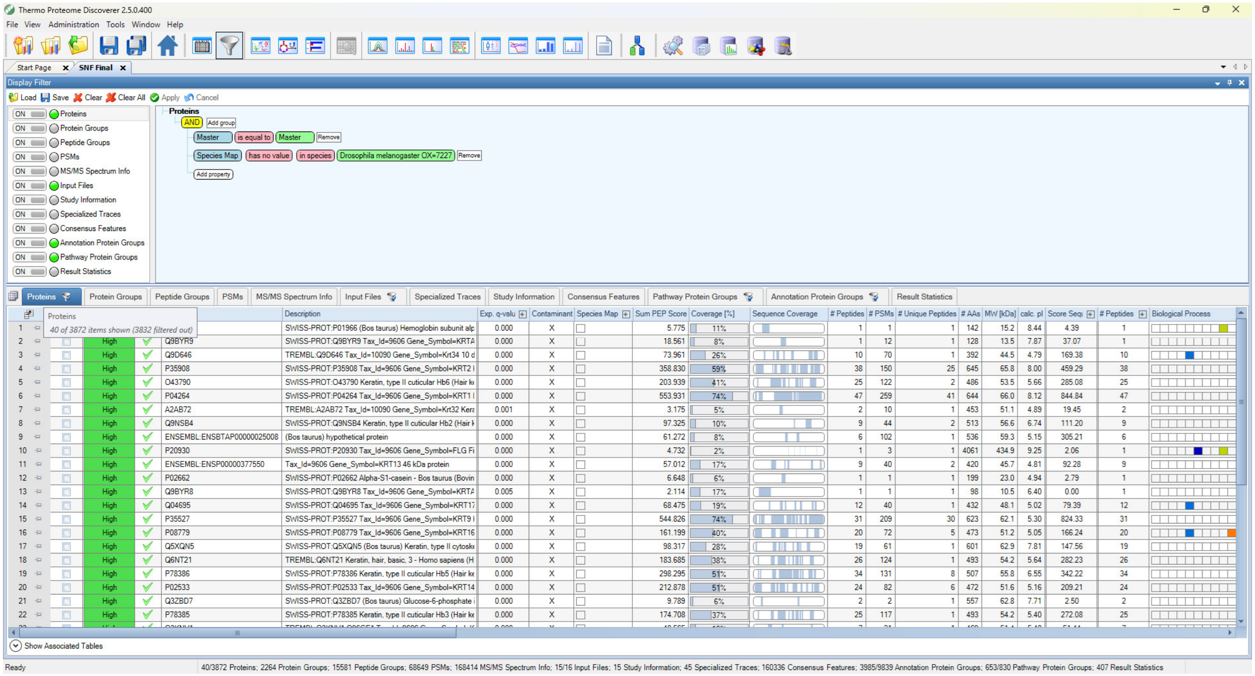


**Image S4. The display shows the filter application for the exclusion of contaminants.** This filter helped to remove 40 factors that did not belong to the species *D. melanogaster*. For the list of the contaminants see Table 4 in File S4.
