## Supplemental Tables for "Targeting Regulatory Factors Associated with the *Drosophila Myc cis*-Elements by Reporter Expression, Gel Shift Assay, and Mass Spectrometric Protein Identification": Image 5.pdf

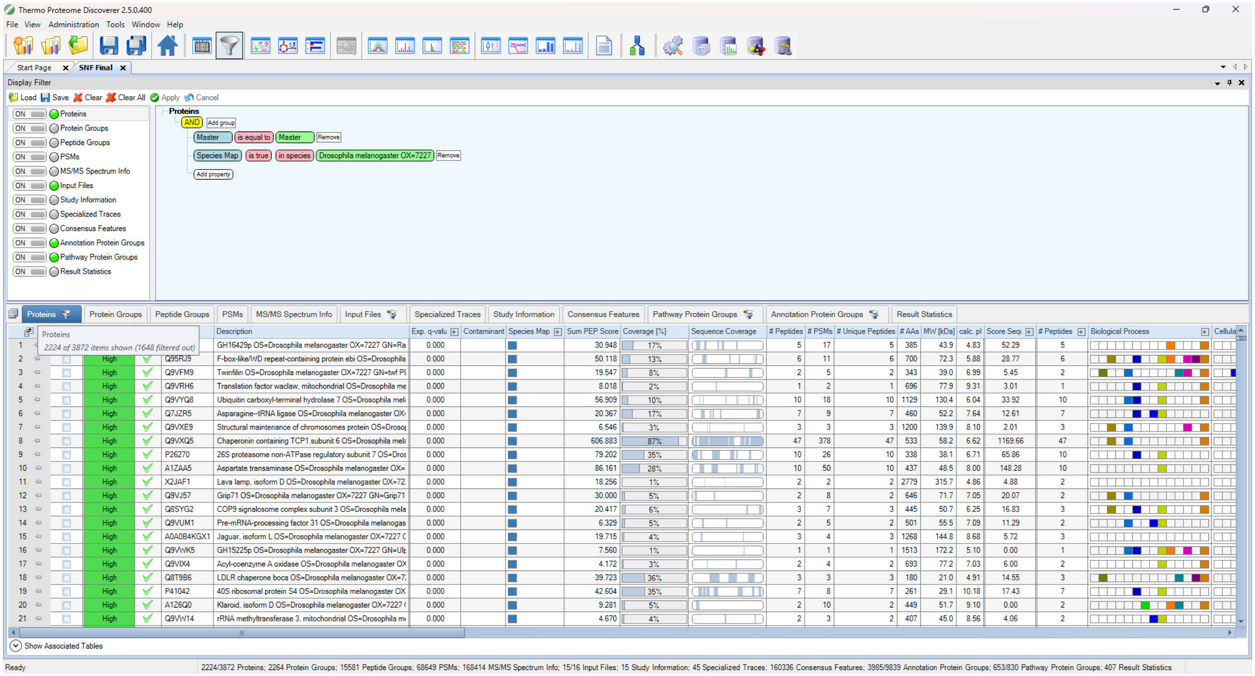


**Image S5. Replicates ungrouped.** After removal of the contaminants and inclusion of the species *D. melanogaster*, with the ungrouped replicates 2224 proteins were obtained. For the complete list of the proteins see Table 5 in File S5.
