## Supplemental Tables for "Targeting Regulatory Factors Associated with the *Drosophila Myc cis*-Elements by Reporter Expression, Gel Shift Assay, and Mass Spectrometric Protein Identification": Image 6.pdf

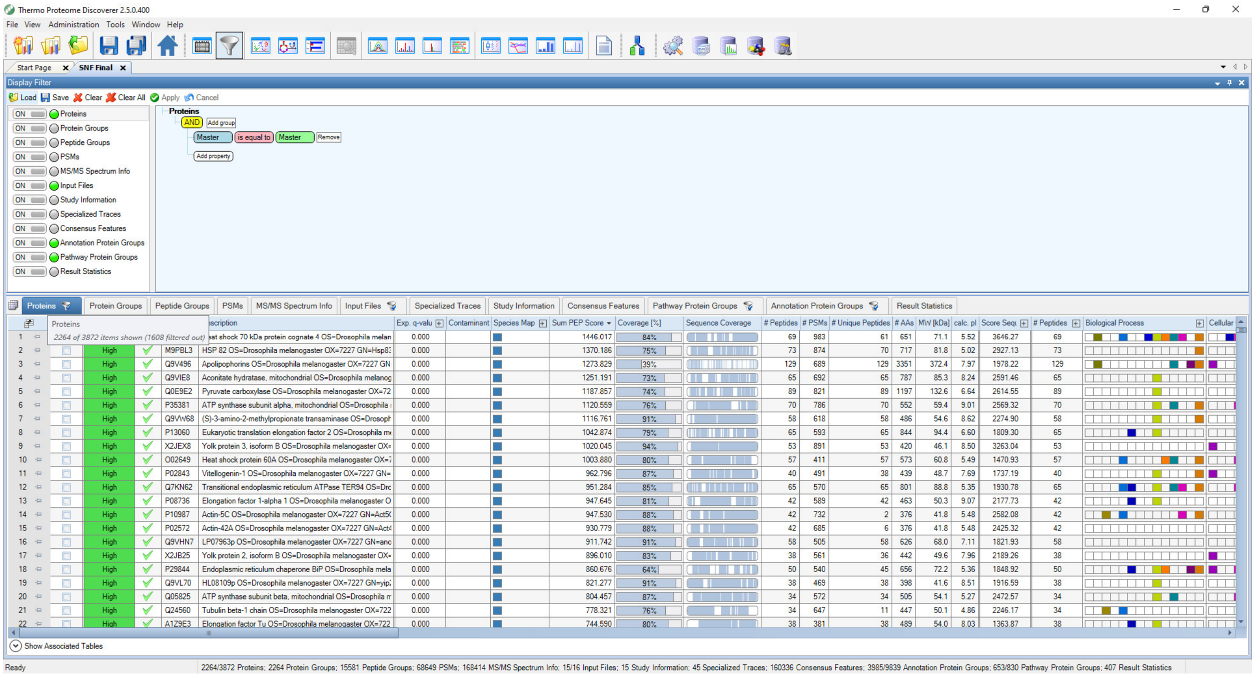


**Image S6. Replicates grouped.** In the unbiased proteome result with the grouped replicates 2264 proteins were obtained. In the following analyses this list served as working material with the application of further filters such as removal of the contaminants and inclusion of the species *D. melanogaster* species. For the complete list of the proteins see Table 6 in File S6.
