## Supplemental Tables for "Targeting Regulatory Factors Associated with the *Drosophila Myc cis*-Elements by Reporter Expression, Gel Shift Assay, and Mass Spectrometric Protein Identification": Image 7.pdf

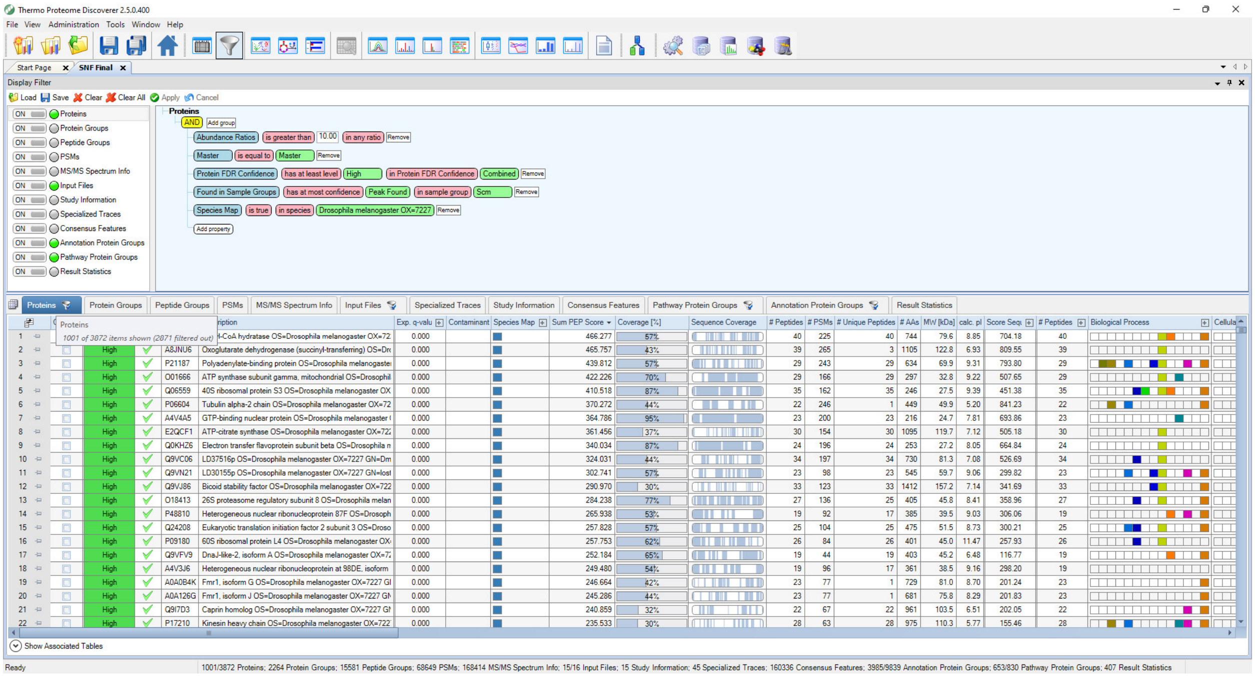


**Image S7. Stringent quality control and filtering of the original data.** The 1001 factors resulting from the original identified dataset that passed stringent quality control were further computationally and manually analyzed to identify *Myc* regulators. For the complete list of factors see Table 7 in File S7.
