## Supplemental Tables for "Targeting Regulatory Factors Associated with the *Drosophila Myc cis*-Elements by Reporter Expression, Gel Shift Assay, and Mass Spectrometric Protein Identification": Table B.pdf

| Regulatory Factors | Abbreviations |
|---|---|
| Activator (Transcriptional) | **ACT** |
| Co-activator (Transcriptional) | **COACT** |
| Co-activator Binding (Transcriptional) | **COACTB** |
| Co-Regulator (Transcriptional) | **COREG** |
| Corepressor (Transcriptional) | **COREP** |
| Repressor/Silencer (Transcriptional) | **REP** |
| Suppressor (Transcription) | **SUP** |
| Chromatin Silencing/Repression/Epigenetic Silencing | **CHRSILEN** |
| Posttranscriptional/Posttranslational Silencing/Repression | **PSILEN** |
| DNA Binding, POLI Transcription | **DBTR1** |
| DNA Binding, POLII Transcription | **DBTR2** |
| DNA Binding, POLIII Transcription | **DBTR3** |
| DNA Binding, Replication/Repair/Recombination | **DBRR** |
| DNA binding Mitochondrial Transcription | **DBMTTR** |
| DNA binding Mitochondrial Replication/Repair/Recombination | **DBMTRR** |
| DNA Helicase, POLI Transcription | **DNAHTR1** |
| DNA Helicase, POLII Transcription | **DNAHTR2** |
| DNA Helicase, POLIII Transcription | **DNAHTR3** |
| DNA Helicase, Replication/Repair/Recombination | **DNAHRR** |
| RNA Helicase, POLI Transcription | **RNAHTR1** |
| RNA Helicase, POLII Transcription | **RNAHTR2** |
| RNA Helicase, POLIII Transcription | **RNAHTR3** |
| RNA Helicase, Translation | **RNAHTN** |
| RNA Helicase, RNA Processing | **RNAHPR** |
| RNA Helicase, Posttranscriptional/Posttranslational | **RNAHPT** |
| RNA Helicase, Replication/Repair/Recombination | **RNAHRR** |
| RNA Binding, POLI Transcription | **RNABTR1** |
| RNA Binding, POLII Transcription | **RNABTR2** |
| RNA Binding, POLIII Transcription | **RNABTR3** |
| RNA Binding, Translation | **RNABTN** |
| RNA Binding, RNA Processing | **RNABPR** |
| RNA Binding, Posttranscriptional/Posttranslational Regulation | **RNABPT** |
| RNA Binding, (RRR) | **RNABRR** |
| DNA primase 1, DNA polymerase α | **PRIM1** |
| DNA primase large subunit, PRIM2 | **PRIM2** |
| DNA polymerase alpha catalytic subunit (Dmel\PolA1) | **DNAPOLA1** |
| DNA polymerase delta subunit 2 (Dmel\PolD2) | **DNAPOLD2** |
| DNA polymerase delta catalytic subunit (Dmel\PolD1) | **DNAPOLD1** |
| DNA polymerase delta subunit 3 (Dmel\PolD3) | **DNAPOLD3** |
| DNA polymerase epsilon subunit 1 (Dmel\PolE1) | **DNAPOLE1** |
| Polymerase (DNA-directed), delta interacting protein 2 (POLDIP2) | **POLDIP2** |
| Chromatin/Nucleosome Remodeling/Organization POLI Transcription | **CRTR1** |
| Chromatin/Nucleosome Remodeling/Organization POLII Transcription | **CRTR2** |
| Chromatin/Nucleosome Remodeling/Organization POLIII Transcription | **CRTR3** |
| Chromatin/Nucleosome Remodeling/Organization, (RRR) | **CRR** |
| Chromosome Maintenance/stability/organization/Condensation | **CHRM** |
| (Chromatin) Insulator | **INSUL** |
| (Chromatin) Insulator Sequence Binding | **INSULB** |
| Chromatin/Hetero-/-DNA Binding, POLI TR | **CBTR1** |
| Chromatin/Hetero-/-DNA Binding, POLII TR | **CBTR2** |
| Chromatin/Hetero-/-DNA Binding, POLIII TR | **CBTR3** |
| Chromatin/Hetero-/-DNA Binding, Translation | **CBTN** |
| Chromatin/Hetero-/-DNA Binding, RNA processing | **CBPR** |
| Chromatin/Hetero-/-DNA Binding, Post-TR/Post-Translational | **CBPT** |
| Chromatin/Hetero-/-DNA Binding, Replication/Repair/Recombination | **CBRR** |
| Enhancer Blocking | **ENBLOC** |
| Histone Modification (other than Acetylation/Deacetylation) | **HMOD** |

| | |
|---|---|
| Histone Acetylation (Acetyl Transferase) | **HAT** |
| Histone Deacetylation | **HDAC** |
| Heterochromatin Formation/Organization | **HF** |
| Mediator Subunit | **MS** |
| Mismatched DNA Binding | **MISDB** |
| Integrator Complex | **INTCOM** |
| Promoter Recognition Factor (Trf2) | **PRF** |
| Polyadenylation Factor | **PAFAC** |
| Poly(A) Binding Protein | **PAB** |
| RNA POL I & III subunit C (Dmel\Polr1C) | **RPI1C** |
| RNA POL I & III subunit C (Dmel\Polr1C) | **RPIII1C** |
| RNA Pol II subunits (Dmel\Polr2B, RPBII140, RPB2) | **RPIIB2** |
| RNA Pol II subunits (Dme\Polr2A, RpII215, RPB1) | **RPIIB1** |
| RNA Pol II subunits ( Dmel\Polr2C, l(2)34Dg, RpII33, RPB3) | **RPIIB3** |
| RNA POL I, II & III subunit E (Dmel\Polr2E, Rpb5, DmRPB5) | **RPIB5** |
| RNA POL I, II & III subunit E (Dmel\Polr2E, Rpb5, DmRPB5) | **RPIIB5** |
| RNA POL I, II & III subunit E (Dmel\Polr2E, Rpb5, DmRPB5) | **RPIIIB5** |
| RNA POL II subunits (Dmel\Polr2G, RPB7) | **RPIIB7** |
| RNA POL I, II, & III subunit RPABC3 (Dmel\Polr2H, RPB8) | **RPIB8** |
| RNA POL I, II, & III subunit RPABC3 (Dmel\Polr2H, RPB8) | **RPIIB8** |
| RNA POL I, II, & III subunit RPABC3 (Dmel\Polr2H, RPB8) | **RPIIIB8** |
| Regulation of Gene Expression, POLI Transcription | **RGEXPTR1** |
| Regulation of Gene Expression, POLII Transcription | **RGEXPTR2** |
| Regulation of Gene Expression, POLIII Transcription | **RGEXPTR3** |
| Regulation of Gene Expression, Posttranscriptional | **RGEXPPT** |
| Regulation of Gene Expression, RNA Processing | **RGEXPPR** |
| Regulation of Gene Expression, Translation | **RGEXPTN** |
| Regulation of Gene Expression, Replication/Repair | **RGEXPRR** |
| Regulation of Gene Expression, Mitochondrial | **RGEXPMT** |
| Replication/Repair/Recombination | **RRR** |
| Mitochondrial Replication/Repair/Recombination | **MTRR** |
| RNA Pol I Transcription | **POLITR** |
| RNA Pol II Transcription | **POLIITR** |
| RNA Pol III Transcription | **POLIIITR** |
| Mitochondrial Transcription Factor | **MTTF** |
| Mitochondrial transcription factor A | **TFAM** |
| Mitochondrial transcription factor B2 | **TFB2M** |
| Mitochondrial Transcription | **MTTR** |
| Mitochondrial Transcription Termination | **MTTT** |
| Mitochondrial RNA binding processing | **MTRNABPR** |
| Transcription Factor, Gene-specific | **TF** |
| TBP &TFIID-associated factors, TAFs (Pre-initiation/initiation) (TPI) | **TPI** |
| Transcription Initiation | **TI** |
| Transcription Elongation | **TE** |
| Transcription Termination | **TT** |
| Transcription Export Complex | **TREX** |
| (DNA-Binding) Transcription Factor Binding | **DBTFB** |
| Telomere Capping/Maintenance Mechanism | **TMM** |
| Translation | **TN** |
| Translation Factor | **TNFAC** |
| Mitochondrial Translation | **MTTN** |
| Mitochondrial Translation Factor | **MTTNF** |
| Uncharacterized | **U** |

**Table B. Subdivisions of regulatory factors associated with the *Myc* target P1/P2 (plotted in Figure 12)**

| Gene name | Transcriptional Category | Activity/Function |
|---|---|---|
| Su(f) | **PAFAC; RNABPR** | **Suppressor of forked**, polyadenylation factor; mRNA 3'-end processing (the nuclear cleavage/polyadenylation reaction) & alternative poly(A) site utilization; accumulation of Suppressor of forked in dividing cells required for mitosis progression (InterPro Project Members, 2004-); histone pre-mRNA cleavage complex. |
| IntS14 | **TT; RNABPR; RGEXPPR; INTCOM** | **Integrator 14**, component of the Integrator complex, a complex with a role in the transcription of small nuclear RNAs (snRNA) and their 3'-box-dependent processing. Involved in the 3'-end processing of the U7 snRNA, and also the spliceosomal snRNAs U1 and U5 (Chen et al., 2012). |
| Tailor | **POLIIITR; RGEXPTR3; RNABTR3; RNABPR** | **Tailor**, cytoplasmic RNA-specific terminal uridylyltransferase (Cheng et al., 2019; Lin et al., 2017; Reimão-Pinto et al., 2015), part of the terminal RNA uridylation-mediated processing (TRUMP) complex; involved in 3'-to-5' exoribonucleolytic decay of RNA species by Dis3l2 (Lin et al., 2017); regulation of microRNA biogenesis by targeting precursor-microRNAs (predominantly mirtron hairpins) & targets unprocessed RNA polymerase III transcripts for degradation in a cytoplasmic RNA surveillance pathway (Reimão-Pinto et al., 2015); Uridyltransferases. |
| Sirt4 | **HDAC; MTTR** | **Sirtuin4**, Sole mitochondrial sirtuin in *Drosophila*, NAD-dependent protein deacylase. Catalyzes the NAD-dependent hydrolysis of acyl groups from lysine residues (Feller et al., 2015); Transcriptional activation of mitochondrial biogenesis, mitochondrial regulator of life span & metabolism, cellular response to starvation, Sirt4 knockout causes short lifespan, increased sensitivity to starvation, decreased fertility & activity (Wood et al., 2018). |
| Ski6 (dSki6) | **RGEXPPT** | **Ski6**, involved in gene expression regulation at the RNA level (RNA Exosome Complex) (Kiss and Andrulis, 2010). |
| DCP1 | **RNABPT; RGEXPPT** | **Decapping protein 1 (DCP1)**, encodes a subunit of the mRNA decapping holoenzyme (Nishihara et al., 2013); involved in oskar mRNA localization (Lee et al., 2020; Lin et al., 2006) and miRNA-mediated gene silencing; mRNA binding (Rehwinkel et al., 2005). |
| Rga | **TN; RNAHTN; RNAHPR; RNAHPT; RGEXPPT; PSILEN** | **Regena, isoform C**, bulk mRNA degradation, miRNA-mediated repression, translational repression (Bawankar et al., 2013); component of CCR4-NOT complex (one of the major cellular mRNA deadenylases); mRNA catabolic process (Bawankar et al., 2013). |

**Table B. Subdivisions of regulatory factors associated with the *Myc* target P1/P2 (plotted in Figure 12)**

| Gene name | Transcriptional Category | Activity/Function |
|---|---|---|
| me31B | **TN; RNAHTN; RNAHPT; RGEXPPT; PSILEN** | **(maternal expression at 31B)**, Dead-Box RNA Helicase, TP-dependent RNA helicase which is a core component of a variety of ribonucleoprotein complexes (RNPs) that play critical roles in translational repression and mRNA decapping during embryogenesis, oogenesis, neurogenesis and neurotransmission; translational repressor activity (Ruscica et al., 2019); miRNA-mediated gene silencing by inhibition of translation (Hillebrand et al., 2010). |
| Su(var)2-10 | **COACT; POLIITR; CHRM** | **Suppressor of variegation 2-10, isoform L**, a member of the PIAS protein family that regulates chromosome structure and function; chromosome organization & condensation (Hari et al., 2001); transcription coregulatory activity, regulation of transcription by RNA pol II (GO Reference Genome Project, 2011-); JAK/STAT pathway regulator, contributes to eye formation & eye determination (Betz et al., 2001); negative regulation of JAK/STAT signaling (Muller et al., 2005; Betz et al., 2001); positive regulators of Hedgehog signaling (Ma et al., 2016; Zhang et al., 2017); negative regulators of Imd signaling pathway (Tang et al., 2021; Cronin et al., 2009). |
| Chi | **COACT; DBTR2; POLIITR** | **Chip, isoform B**, encodes a transcriptional coactivator, LIM domain-binding protein 2; LIM domain-binding protein/SEUSS; LIM interaction domain; imaginal disc-derived wing morphogenesis (Rincon-Limas et al., 2000) / leg development & axon guidance (van Meyel et al., 2000); positive regulation of transcription by RNA pol II (Werner et al., 2017; van Meyel et al., 2000). |
| Dhx15 | **RNAHPR; RNABPT; RGEXPPT** | **DEAH-box helicase 15**, RNA helicase activity; involved in mRNA processing (Herold et al., 2009; Mount and Salz, 2000). |
| Cbp20 | **RNABPR; PSILEN; RGEXPPR** | **20 kDa nuclear cap-binding protein**, mRNA binding (Lasko, 2000); RNA binding; RNA cap binding (InterPro Project Members, 2004-); siRNA processing (Sabin et al., 2009); ncRNA-mediated post-transcriptional gene silencing; NCBP2 RNA Recognition Motif. |
| CNBP | **DBTR2; RNABTN; TN** | **CCHC-type zinc finger nucleic acid binding protein**, single-stranded DNA/RNA-binding protein with sequence specificity; CNBP/MYC axis, involved in the control of wing size by regulating *Myc* levels; IRES trans-acting factor (ITAF), promote IRES-dependent *Myc* mRNA translation (Antonucci and Ganettieri, cc-2014). |

**Table B. Subdivisions of regulatory factors associated with the *Myc* target P1/P2 (plotted in Figure 12)**

| Gene name | Transcriptional Category | Activity/Function |
|---|---|---|
| Doc3 | **TF; POLIITR; DBTR2** | **Dorsocross**, one of the three tissue-specific T-box transcription factors encoded by the Dorsocross cluster, RNA pol II *cis*-regulatory region sequence-specific DNA binding (GO Reference Genome Project, 2011-; InterPro Project Members, 2004-); regulation of Wnt/Wg and TGF-beta pathways (Hatton-Ellis et al., 2007); cardiac induction in *Drosophila* based on combinatorial activities of Dpp and Wg signaling derived from the ectoderm (Reim and Frasch, 2005). |
| Dmel\lola | **DBTR2; POLIITR; TF** | **Longitudinals lacking protein, isoforms H/M/V**, involved in Notch signaling; cell death; neurogenesis; regulation of retrotransposons; oogenesis; spermatogenesis; wiring, eye development & a variety of behaviors (Tripathy, R. et al., 2014; Iyer, E.P. et al., 2013; Bass, B.P. et al., 2007; Girard, F. et al., 2006; Crowner, D. et al., 2002; Giniger, E. et al., 1994); positive regulators of immune deficiency (Imd) pathway based on the activity of NF-κB-like transcription factor (*Rel*) in the nucleus; DNA binding (Dinges et al., 2017); positive regulation of peptidoglycan recognition protein signaling pathway (Kleino et al., 2005); DNA binding transcription factor activity (Giniger et al., 1994); C2H2 Zinc Finger Transcription Factors. |
| ash2 | **DBTR2; TI; COACT; POLIITR; HMOD; CRTR2** | **(absent, small, or homeotic discs 2)**, a component of the histone methyltransferase complex, specifically methylates lysine 4 of histone H3 & a member of the trithorax family; mutants show homeotic transformations & a variety of pattern-formation defects; DNA binding (Pérez-Lluch et al., 2011); imaginal disc-derived wing vein specification (Angulo et al., 2004); transcription coactivator activity (Carbonell et al., 2013); chromatin remodeling (Beltran et al., 2007); transcription initiation coupled chromatin remodeling (LaJeunesse and Shearn, 1995). |
| fon (CG1582) | **RNABTN; RNAHTN; TN** | **Fondue, isoform B**, RNA helicase activity; translation initiation activity (Linsalata et al., 2019; Lasko, 2000); RNA binding activity (GO Reference Genome Project, 2011-); DEAH-Box RNA Helicases. |
| ebi | **REP; POLIITR; COREP; DBTFB** | **Ebi**, F-box-like/WD repeat-containing protein, conserved repressor/silencer; JNK signaling: Ebi/ AP-1 complex represses pro-/anti-apoptotic genes; suppresses basal transcription levels of apoptotic genes protecting sensory neurons degeneration (Lim et al., 2012); regulates EGFR (Dong et al., 1999) and Notch signaling pathways (Nguyen et al., 2016; Marygold et al., 2011; Tsuda et al., 2002); negative regulation of transcription (Lim et al., 2012); DNA binding transcription factor binding (Qi et al., 2008). |

**Table B. Subdivisions of regulatory factors associated with the *Myc* target P1/P2 (plotted in Figure 12)**

| Gene name | Transcriptional Category | Activity/Function |
|---|---|---|
| Trf2 | **PRF; CBTR2; DBTR2; TF; POLIITR; TPI; TI** | **TATA box-binding protein-like 1**, a core promoter recognition factor (PRF), mediates gene transcription; Ecdysone signaling (Bashirullah et al., 2007); ecdysteroid signaling (Shima et al., 2007); chromatin binding (Wang et al., 2014); TFIIA-class transcription factor complex (Andersen et al., 2017); RNA pol II initiation factor (Hochheimer et al., 2002); RNA pol II core promoter sequence-specific DNA binding (Kedmi et al., 2014); TAT binding protein & TBP-Related Factor. |
| Su(fu) | **DBTFB; REP; POLIITR** | **Suppressor of fused**, (Ser/Thr-protein kinase; negative regulator of Hedgehog signaling by forming a complex with the transcription factor Cubitus interruptus (Ci); (Han et al., 2019; Han et al., 2015; Smelkinson et al., 2007; Zhang et al., 2011; Fukumoto et al., 2001; Methot and Basler, 2000); negative regulator of Dpp pathway *(*Jia et al., 2002-nature); DNA binding transcription factor binding (Han et al., 2015); molecular sequestering activity (Han et al., 2019; Han et al., 2015; Oh et al., 2015; Shi et al., 2014; Wang et al., 2000). |
| nito | **RGEXPPT; RGEXPPR** | **RNA-binding protein Spenito**, a component of the WMM complex, mediation of N6-methyladenosine (m6A) methylation of mRNAs (Guo et al., 2018; Knuckles et al., 2018; Lence et al., 2016);positive regulator of canonical Wg-TCF signaling during wing disk & eye development (Chang et al., 2008; Jemc and Rebay, 2006); positive regulation of gene expression (Dewald et al., 2014); regulation of mRNA splicing, via spliceosome (Lence et al., 2016). |
| Rcc1 | **CBRR; CHRM** | **Regulator of chromosome condensation**, nuclear import & export of beta-Catenin (Koyama et at., 2017); regulation of mitotic cell cycle, apoptosis pathway; chromatin binding (Trieselmann and Wilde, 2002). |
| Doa | **RNABPR; RGEXPPR** | **Darkener of apricot**, 105 kDa Doa protein kinase (non-specific Ser/Thr protein kinase), positive regulators of Toll NF-κB signaling pathway (Kanoh et al., 2015); MAPK cascade (Yun et al., 2000); TORC1 signaling inhibits CDK8 & DOA kinases, which directly phosphorylate CPSF6, a component of the CPA complex, and regulate CPSF6 localization, RNA binding, & starvation-induced alternative RNA processing of transcripts involved in autophagy (Tang et al., 2018); negative regulation of male germ cell proliferation (Zhao et al., 2013); regulation of RNA splicing (Park et al., 2004). |

**Table B. Subdivisions of regulatory factors associated with the *Myc* target P1/P2 (plotted in Figure 12)**

| Gene name | Transcriptional Category | Activity/Function |
|---|---|---|
| ca | **RGEXPPT; PSILEN** | **Claret**, negative regulation of post-transcriptional gene silencing by regulatory ncRNA (Harris et al., 2011) acts as a guanine nucleotide exchange factor (GEF) for Lightoid/Rab-RP1 in an adaptor protein 3-independent vesicular trafficking pathway of pigment granule biogenesis (Ma et al., 2004); Rab32 GEF/Claret involved in autophagy, affecting lipid storage (Wang et al., 2012). |
| how | **RNABPT; RGEXPPT** | **(held out wings)**, RNA-binding protein (Volohonsky et al., 2007; Di Fruscio et al., 1998) highly expressed in mesoderm & tendon cells; two isoforms: how(L) & how(S). how(L) induces RNA destabilization, How(S) stabilizes the RNA targets; integrin signaling, apposition of dorsal/ventral imaginal disc-derived wing surfaces (Walsh and Brown, 1998); cell adhesion (Lo and Frasch, 1997); EGFR signaling: Glial cell migration; axon ensheatment (Edenfeld et al., 2006; Lasko, 2003); Dpp (Israeli & Volk, 2007). |
| pum | **RNABTN; RNABPT; RNABPR; TN; RGEXPPT; PSILEN** | **Pumilio, isoform G**, a founding member of the PUF family of RNA binding proteins (Chen et al., 2008); mRNA regulatory element binding translation repressor activity (Kim et al., 2012); sequence-specific mRNA binding (Arvola et al., 2020); post-transcriptional gene silencing; nuclear-transcribed mRNA catabolic process (Miles et al., 2012); negative regulation of DNA-templated transcription (Leatherman and Jongens, 2003). |
| Fmr1 | **RNABPT; RNAHPT; RGEXPPT; PSILEN** | **Fragile X messenger ribonucleoprotein 1**, RNA & channel binding protein; DEAD/H-Box RNA helicase binding (Barbee et al., 2006); mRNA regulatory element binding repressor activity (Fajner et al., 2021); post-transcriptional regulation of gene expression (Luhur et al., 2017). |
| His2Av | **DBRR; RRR; CBRR; RGEXPRR; HF** | **Histone H2A variant**, the only histone H2A variant (H2AV), a chimera of H2AZ & H2AX in *Drosophila*; required for larval hematopoiesis (Grigorian et al., 2017); DNA binding; structural constituent of chromatin (InterPro Project Members, 2004-); element of a class of active promoter structure; gene regulation (Baldi and Becker, 2013); phosphorylated in response to DNA damage in the less conserved C-terminal tail; protein-containing complex binding (Kusch et al., 2004); cell division & kinetochore driven microtubule formation (Vernì and Cenci, 2015). |

**Table B. Subdivisions of regulatory factors associated with the *Myc* target P1/P2 (plotted in Figure 12)**

| Gene name | Transcriptional Category | Activity/Function |
|---|---|---|
| (puf) | **RRR; RGEXPRR** | **Puffyeye**, Ubiquitin-Specific Protease (USP), an essential deubiquitinating enzyme, acts as a ubiquitin-specific protease, removes ubiquitin polypeptide chains from Myc & CycE, stabilizes/increases their abundance to influence cell growth & proliferation; post-translational protein modification (Li et al., 2013); positive regulation of double-strand break repair via homologous recombination (Páhi et al., 2022); TCF dependent signaling in response to WNT. |
| IntS11 | **RNABTR2; INTCOM; POLIITR; RNABPR** | **Integrator complex subunit 11**, involved in the transcription of small nuclear RNAs (snRNA) & their 3'-box-dependent processing; pol II transcription (Baillat et al., 2005); Notch signaling (Shersher et.al., 2021); Epidermal Growth Factor Receptor signaling pathway (F. C.Tilley & G. Mollet, 2021). |
| sxc | **CHRSILEN; RGEXPPT** | **Super sex combs**, a Polycomb group gene, encodes a O-GlcNAc transferase, involved in epigenetic gene silencing; regulation of gene expression (Akan et al., 2016); positive regulation of Fibroblast Growth Factor Receptor (FGFR) signaling pathway (Mariappa et al., 2011). |
| SkpA | **RRR; CHRM** | **SKP1-related A**, a subunit of Skp, Cullin, F-box (SCF)-containing ubiquitin ligase complexes, regulates centrosome duplication, chromatin condensation (Murphy, 2003); cell cycle progression, cell polarity, dendrite pruning & endoreduplication.; mitotic cell cycle (Ducat et al., 2008). |
| Cdk1 | **RRR** | **Cyclin-dependent kinase 1**, regulation of DNA replication initiation; embryonic syncytial blastoderm development (Seller and O'Farrell, 2018); Cyclin-dependent protein Ser/Thr kinase activity; regulation of the G1/S & G2/M transition phases of mitotic cell cycle (Lehner and O'Farrell, 1990); Protein binding (Jacobs et al., 1998). |
| l(3)72Ab (Brr2) | **RNAHPR; RNABPR** | **U5 small nuclear ribonucleoprotein 200 kDa helicase**, mitotic cell cycle (Ducat et al., 2008); steroid hormone ecdysone signaling (Claudius et al., 2014); RNA Helicase activity (Lasko, 2000); mRNA splicing, via spliceosome (Monedero Cobeta et al., 2018). |
| Hrb87F | **RNABPR; RGEXPPR; DBRR; TMM** | **Heterogeneous nuclear ribonucleoprotein at 87F**, hnRNP-A family RNA-binding protein; involved in gene expression & RNA processing (Lasko, 2000); an essential component of the nucleoplasmic omega speckles; necessary for telomere maintenance; sequence-specific DNA binding (Borah et al., 2009); regulation of alternative mRNA splicing, via spliceosome (Borah et al., 2009; Park et al., 2004); Wnt/Wingless and JNK signaling (Yadav & Tapadia, 2016). |

**Table B. Subdivisions of regulatory factors associated with the *Myc* target P1/P2 (plotted in Figure 12)**

| Gene name | Transcriptional Category | Activity/Function |
|---|---|---|
| 14-3-3ε | **COACTB; POLIITR; RRR** | **14-3-3 protein epsilon**; an acidic protein, preferentially heterodimerizes with other members of the family (Messaritou et al., 2010), also can homodimerize; functions in multiple signaling pathways, most prominently in the Ras/MAPK cascade; regulation of nuclear mitotic cell division (Ashton-Beaucage et al., 2014; Su et al., 2001); interacts with kinases such as PKC or Raf-1; transcription coactivator binding (Ren et al., 2010); in plants associated with a complex that binds to the G-box promoter elements; Ras/MAPK (Ashton-Beaucage et al., 2014); DNA damage checkpoint signaling (Su et al., 2001; Brodsky et al., 2000); contains 14-3-3 domain. |
| BAP155 moira (mor) | **POLIITR; CRTR2; CBTR2; DBTR2** | **Brahma associated protein 155 kDa**, a member of the trithorax group of homeotic gene regulators; chromatin remodeling as the Swi3 component of the Brahma complex (Crosby et al., 1999); regulation of transcription by RNA pol II (Bonnay et al., 2014); imaginal disc-derived wing margin morphogenesis, regulation of EGFR signaling pathway (Terriente-Felix & de Celis, 2009); negative regulation of G1/S transition of mitotic cell cycle (Brumby et al., 2002); regulation of innate immune response via NF-κB signaling pathway (Bonnay et al., 2014). |
| Mlf | **TF; POLIITR** | **Myelodysplasia/myeloid leukemia factor**, interacts with various factors involved in transcriptional regulation, regulation of cell proliferation during eye morphogenesis, Lozenge activity during hematopoietic development and assembly of the COP9 signalosome complex; (Miller et al., 2017); Regulation of DNA-template transcription (Bras et al., 2012). |
| Bap111 | **DBTR2; CRTR2; POLIITR; TF** | **Brahma associated protein 111kD**, extensive homology of the Brahma (BRM) complex to SWI/ SNF; DNA binding; chromatin remodeling (Papoulas et al., 2001); Polybromo-Containing Proteins Complex; High Mobility Group Box Transcription Factors. |
| glu (CG11397) | **CHRM; CBRR** | **Gluon**, part of the multiprotein complex condensin, mitotic cell cycle: required for prometaphase chromosome condensation (Nikalayevich and Ohkura, 2015); sister chromatid segregation; nervous system development; glucose metabolism; expressed in dividing cells throughout embryogenesis—pole cells, neuroblasts in the CNS & the PNS; chromosome condensation (Bivik et al., 2015); chromatin binding (Gene Ontology Curators, 2002-); Structural Maintenance of Chromosomes Family. |

**Table B. Subdivisions of regulatory factors associated with the *Myc* target P1/P2 (plotted in Figure 12)**

| Gene name | Transcriptional Category | Activity/Function |
|---|---|---|
| Helz | **RNAHPT; PSILEN** | **Helicase with zinc finger**, RNA helicase activity (Hanet et al., 2019); involved in ncRNA-mediated post-transcriptional gene silencing (GO Reference Genome Project, 2011-). |
| Hmg-2 (HMGB2) | **TF; POLIITR; DBTR2; CRTR2; HDAC** | **High mobility group protein 2**, together with the Wnt signaling regulate chondrocyte hypertrophy by mediating Runt-related transcription factor 2 expression (Taniguchi et al., 2018); Wnt signaling & HMGB2 regulate articular cartilage surface maintenance (Taniguchi et al., 2009); association with chromatin, ubiquitous distribution in nucleus, and non-specific DNA minor groove binding; DNA bending / unwinding and promotion of DNA flexibility (McCauley et al., 2005; Lorenz et al., 1999); regulation of gene expression (GO Reference Genome Project, 2011-). |
| (mod) | **POLIITR; DBTR2; CRTR2; TF** | **Modulo**, the *Drosophila* homologue of nucleolin (Mikhaylova et al., 2006); MYC pathway (target of MYC selectively required for the growth of proliferative cells); association with the proto-oncogene MYC (Perinn et al., 2003); required for meiosis & spermatid differentiation in male germ line; sequence-specific DNA binding (Mikhaylova et al., 2006); involved in chromatin packaging; dominant suppressor of variegation (Bantignies et al., 2002). |
| psi | **POLIITR; DBTR2; RNABPR** | **P-element somatic inhibitor, isoform C**, far upstream element binding with dual roles in RNA processing (Labourier et al., 2002; Siebel et al., 1994) & transcriptional regulation; required for activating MYC transcription. KSRP (KHSRP) binding & destabilization of mRNA; mRNA binding (Siebel et al., 1994); regulation of DNA-templated transcription (InterPro Project Members, 2004-). |
| mask | **RNABTR2; POLIITR** | **Multiple ankyrin repeats single KH domain**, RNA binding (InterPro Project Members, 2004-); positive regulation of transcription by RNA pol II (Sansores-Garcia et al., 2013; Sidor et al., 2013). |
| bel | **RNAHPT; RNABPT; RGEXPPT** | **Belle**, Dead-Box RNA Helicase; regulatory ncRNA-mediated post-transcriptional gene silencing (Ulvila et al., 2006); mitotic cell cycle (Pek and Kai, 2011). |
| Ddx1 | **RNABPR; RNAHPT; CBTR2; COREG; DBTR2** | **Dead-box-1**, a member of the DEAD box family of RNA helicases that bind and unwind double-stranded RNA-RNA; DNA-RNA helicase activity, chromatin binding; transcription coregulatory activity (Gene Ontology Curators, 2002-); tRNA processing (Schmidt et al., 2019); DEAD-Box 1 is a novel & independent prognostic marker for early recurrence in breast cancer (Germain et al., 2011). |

**Table B. Subdivisions of regulatory factors associated with the *Myc* target P1/P2 (plotted in Figure 12)**

| Gene name | Transcriptional Category | Activity/Function |
|---|---|---|
| lin-28 | **RNABPT; RGEXPPT** | **Protein lin-28 homolog**, cold shock and RNA-binding protein (Sreejith et al., 2019); regulator of developmental timing; regulator of microRNA maturation; post-transcriptional regulation of gene expression (Luhur et al., 2017; Chen et al., 2015); positive regulation of receptor signaling pathway via JAK/STAT (Sreejith et al., 2019). |
| Cul4 | **RRR; RGEXPRR** | **Cullin 4, isoform A**, molecular scaffold for the CRL4 E3 ubiquitin ligase complex, catalyzes the ubiquitylation (Reynolds et al., 2008) & subsequent destruction of proteins functioning in cell growth, proliferation, transcription, replication & repair of the genome; SCF-dependent proteasomal ubiquitin-dependent protein catabolic process; cellular response to DNA damage stimulus (GO Reference Genome Project, 2011-); negative regulation of Smoothened signaling pathway (Li et al., 2018); protein ubiquitination (Li et al., 2018; Reynolds et al., 2008). |
| asun | **POLIITR; INTCOM** | **Protein asunder**, mitotic cell cycle (Lee et al., 2005); component of the Integrator complex involved in the transcription of small nuclear RNAs (snRNA) by RNA pol II-transcribed snRNAs & their 3'-box-dependent processing (Chen et al., 2012). |
| HDAC1 | **HDAC; RGEXPPT; CHRSILEN; CRTR2; POLIITR; CHRM** | **Histone deacetylase 1**, deacetylation of lysine residues on the N-terminal part of the core histones (H2A, H2B, H3 and H4); negative regulation of gene expression, epigenetics (Janssens et al., 2017); Hedgehog signaling (Zhang et al., 2013); transcription corepressor activity (Miotto et al., 2006); chromosome condensation (Nikalayevich and Ohkura, 2015); NAD-independent histone deacetylation (Feller et al., 2015); regulation of DNA-templated transcription (Cho et al., 2005). |
| CG6664 | **CHRM** | Establishment of meiotic spindle orientation; spindle pole, condensed chromosomes (Gene Ontology Curators, 2002-). |
| gw | **RGEXPPT; PSILEN; RNABPR; RNABTN; TN** | **Gawky**, RNAi pathway (miRNA-mediated gene silencing pathway) (Chekulaeva et al., 2010; Chekulaeva et al., 2009; Eulalio et al., 2009; Eulalio et al., 2009; Zekri et al., 2009; Eulalio et al., 2008); mRNA catabolic process (Behm-Ansmant et al., 2006); miRNA induced silencing complex (RISC) (Behm-Ansmant et al., 2006); negative regulation of gene expression, RNA binding (Sienski et al., 2015); miRNA-mediated gene silencing by inhibition of translation (InterPro Project Members, 2004-). |

**Table B. Subdivisions of regulatory factors associated with the *Myc* target P1/P2 (plotted in Figure 12)**

| Gene name | Pathway | Activity/Function |
|---|---|---|
| Nurf-38 | **CBTR2; CRR; POLIITR; RRR** | **Nucleosome remodeling factor - 38kD**, component of NURF (nucleosome remodeling factor), ATP-dependent nucleosome sliding facilitates transcription (Gdula et al., 1998); chromatin remodeling (Mizuguchi et al., 2001; Gdula et al., 1998); positive regulation of DNA-templated transcription (Mizuguchi et al., 2001); ecdysone receptor-mediated signaling pathway (Badenhorst et al., 2005). |
| Elp4 (CG6907) | **POLIITR; TN; RRR; RNABTN** | **Elongator complex protein 4**, establishment of mitotic spindle asymmetry; regulation of translation through targeted transfer-RNA (tRNA) modification (Planelles-Herrero et al., 2022); phosphorylase kinase regulator activity; regulation of transcription by RNA pol II (Gene Ontology Curators, 2002-); Elongator Complex. |
| woc | **POLIITR; TF; CBTR2; DBTR2; TMM** | **Without children, isoform B, ecdysone**, involved in biosynthetic process (Wismar et al., 2000); regulation of transcription by RNA pol II (Abel et al., 2009; Font-Burgada et al., 2008); chromatin-binding factor (Font-Burgada et al., 2008) related to the mammalian MYM-type family of transcription factors; involved in telomere capping. |
| shep | **REP; DBTR2; CBTR2; POLIITR** | **Protein alan shepard**, an evolutionarily conserved RNA/DNA binding protein; regulation of alternative splicing & gypsy insulator activities (chromatin insulator associated) (Chen et al., 2021). |
| btz | **RNABTN; TN; RNABPR; RGEXPPT** | **Barentsz (CASC3 in mammals) (other name MLN51)**, a component of Exon Junction Complex pathway (EXJC), recruited to spliced mRNAs to mark introns removal sites; translation activator linking the EJC & the translation machinery, direct role in protein synthesis (Chazal et al., 2013); BTZ domain found on CASC3 (cancer susceptibility candidate gene 3 protein, also known as **MLN51= M**etastatic **L**ymph **N**ode **51)** also known as Barentsz (Btz); CASC3, component of EJC involved in post-transcriptional regulation of mRNA in metazoa; complex formed by association of 4 proteins (eIF4AIII, Barentsz, Mago, & Y14), mRNA, & ATP; BTZ wraps around eIF4AIII & stacks against the 5' nucleotide (Bono et al., 2006; Palacios et al., 2004); Barentsz (MLN51) overexpressed in breast cancer (Degot et al., 2002-); repression of MNL51 by brain-specific miR-128 (Wilkinson et al., 2011); mRNA binding; RNA binding (InterPro Project Members, 2004-). |

**Table B. Subdivisions of regulatory factors associated with the *Myc* target P1/P2 (plotted in Figure 12)**

| Gene name | Transcriptional Category | Activity/Function |
|---|---|---|
| MRG15 | **HAT; CRTR2; CBTR2; CRR; CBRR; RGEXPPT; POLIITR; CHRM; RRR; HF** | **MORF-related gene 15**, histone acetylation; chromatin binding; positive regulation of gene expression (Huang et al., 2017); chromatin organization; chromosome separation (Smith et al., 2013); DNA repair-dependent chromatin remodeling (Kusch et al., 2004); heterochromatin formation (Qi et al., 2006); regulation of DNA-templated transcription (InterPro Project Members, 2004-). |
| wds | **POLIITR; CRTR2** | **Will die slowly, isoform B**, (Hollmann et al, 2002), essential gene codes for a WD-repeat protein with seven repeats; WD40 repeat implicated in transcription regulation; positions the N-terminus of histone H3 for efficient trimethylation at 'Lys-4' (Pascual-Garcia et al., 2014); chromatin remodeling (Suganuma et al., 2008); Mad interacts with the Trithorax-related complex protein Wds to maintain active transcription by dynamically demethylating intragenic 6mA; maintenance of transcriptional activation for specific sets of genes (yao et al., 2018); ATAC Complex, TRR complex, COMPASS Complex, TRX Complex. |
| kis | **CBTR2; CRTR2; DBTR2; TF; POLIITR; RGEXPPT** | **Kismet, isoform F**, a conserved chromodomain containing ATP-dependent transcription factor (Terriente-Félix et al., 2011); gene control through epigenetic mechanisms; chromatin binding; ATP-dependent chromatin remodeling; DNA binding; regulation of gene expression (GO Reference Genome Project, 2011-); negative regulation of DNA-templated transcription (Thompson et al., 2008). |
| CG17202 | **POLIITR; COACT; DBTR2** | **c-Myc-binding protein homolog,** transcription coactivator activity; regulation of transcription, DNA-templated (InterPro Project Members, 2004-); stimulation of *c-Myc* transcription (Taira et al., 1998); associate of Myc 1; orthologous to human MYCBP (MYC binding protein). |
| Mi-2 | **CBTR2; CRTR2; DBTR2; POLIITR; CHRM** | **Mi-2**, a nuclear ATP-dependent nucleosome (chromatin) remodeling activity (Kunert et al., 2009; Murawska et al., 2008); chromatin binding (Kunert et al., 2009); required for repression of cell type-specific genes; DNA binding (InterPro Project Members, 2004-); chromosome condensation (Nikalayevich and Ohkura, 2015); regulation of transcription by RNA pol II (Li et al., 2010); Notch signaling (Zacharioudaki E, Falo Sanjuan J, Bray S., Elife. 2019); Wingless and ecdysone signaling (Kon &Nusse; 2005); Nucleosome Remodeling Deacetylase Activity; SNF2-Like Chromatin Remodelers. |

**Table B. Subdivisions of regulatory factors associated with the *Myc* target P1/P2 (plotted in Figure 12)**

| Gene name | Transcriptional Category | Activity/Function |
|---|---|---|
| barr | **CBRR; CRR; RRR** | **Barren**, chromatin binding protein, chromatin condensation, DNA topoisomerase binding (Lupo et al., 2001) regulates Malpighian tubule development & epithelial morphogenesis; Tube development (Liu et al., 1999): Hedgehog, required for the completion of bud evagination (Hoch and Pankratz 1996), Wingless, required for cell division & morphogenesis in the tubules (Skaer and Martinez Arias 1992; Harbecke and Lengyel 1995), Notch receptor, required to define the single tip cell at the end of each tubule (Hoch et al. 1994), which leads out the elongation of the tubule (Skaer, 1989), EGFR (EGF-like) ligand required for the proliferation of the distal cells of the tubule (Baumann and Skaer 1993; Kerber et al. 1998); Dpp expressed in foregut/hindgut, required for morphogenesis of both structures (Pankratz and Hoch 1995; Hoch and Pankratz 1996); involved in cell cycle interphase (Lupo et al., 2001). |
| msi | **RNABTN; TN; PSILEN** | M**usashi**, RNA binding protein, binds to the 3' UTR region of target mRNAs; negatively regulates the Hypoxia Inducible Factor (HIF) pathway, contributes to cell fate determination, as well as cellular response to normoxic/hypoxic conditions; negative regulation of translation, asymmetric cell fate determination by Notch signaling; mRNA regulatory element binding translation repressor activity (Bardin et al., 2004). |
| Cdc6 | **DBRR; RRR** | **Cell division cycle 6**, encodes an essential component of the pre-Replication Complex (preRC) together with the origin recognition complex, the product of (*dup*) & MCM2-7 proteins (InterPro Project Members, 2004-). |
| fs(1)h | **POLIITR; TE; DBTR2; TF; CBTR2** | **Homeotic protein female sterile** (*Drosophila* female sterile (1) homeotic), non-specific serine/threonine protein kinase, Ras signaling: a multifunctional transcriptional regulator modulated by Ras signaling (Florence & Faller, 2008); Hedgehog (Hh) signaling; (Xiangdong et al., 2018; Bagley et al., 2014); a Bromodomain & Extra-Terminal domain (BET) family of protein, characterized by the presence of two tandem bromodomains & an extra-terminal (ET) domain. BET proteins can bind acetylated lysine residues in histones and regulate transcription. Homeotic protein female sterile & human BRD2 possess protein kinase activity; chromatin binding (Gaub et al., 2020); DNA binding transcription activator activity (Chang et al., 2007); unclassified DNA binding domain transcription factors; atypical protein kinases. |

**Table B. Subdivisions of regulatory factors associated with the *Myc* target P1/P2 (plotted in Figure 12)**

| Gene name | Transcriptional Category | Activity/Function |
|---|---|---|
| BRWD3 | **CRTR2; RGEXPPT; POLIITR; DBTR2** | **BRWD3**, positive regulator of JAK/STAT signaling (Muller et al., 2005); RHOBTB2 GTPase cycle; Interleukin-7 signaling; chromatin modifying enzymes; member of bromodomain & WD repeat containing protein (BRWD) family; regulation of gene expression; regulation of ecdysone & JAK/STAT signaling (Chen et al., 2015); regulation of transcription by RNA pol II (GO Reference Genome Project, 2011-). |
| atms | **POLIITR; TE; CBTR2; DBTR2; HMOD** | **Antimeros**, a component of the PAF1 complex; physically interacts with components of the basal transcription machinery and sequence-specific transcription factors to control histone modifications and pause release, positive regulation of DNA-templated transcription (Adelman et al., 2006); chromatin binding; RNA pol II binding (GO Reference Genome Project, 2011-); alternative mRNA splicing pathway (Lence et al., 2016; Burnette et al., 1999); transcription elongation by RNA pol II; histone modification (InterPro Project Members, 2004-); E3 ubiquitin ligases ubiquitinate target proteins; Notch signaling (Lence et al., 2016; Hongay & Orr-Weaver, 2011); RNA Polymerase II-Associated Factor 1 Homolog. |
| aub | **RGEXPPT; RNABPT** | **Aubergine**, piRNA binding (Huang et al., 2021; Webster et al., 2015; Nagao et al., 2010); global gene silencing by mRNA cleavage (Kennerdell et al., 2002); ncRNA-mediated post-transcriptional gene silencing (Tomari et al., 2004); RNA-mediated gene silencing (Bozzetti et al., 2015); JAK/STAT signaling pathway controls host defense in the gut by regulating stem cell proliferation & epithelial cell homeostasis (Cronin et al., 2009); RNA binding (Brennecke et al., 2007). |
| e(r) | **RRR; POLIITR; REP** | **Enhancer of rudimentary (ERH)**, evolutionarily highly conserved, implicated in the regulation of pyrimidine biosynthesis, DNA replication, transcription, mRNA splicing, cellular proliferation, & tumorigenesis (Gelsthorpe et al., 1997); cell-type specific transcriptional repressor activity (Pogge von Strandmann et al., 2001) Notch signaling pathway (Tsubota et.al., 2011); Fission yeast homologue of ERH is implicated in meiotic mRNA elimination during vegetative growth (Hazra et al., 2020; Shichino et al., 2018; Sugiyama et al., 2016); involvement of ERH-SRPK1-SAFB proteins in mitotic phosphorylation of Lamin B receptor (LBR) at the nuclear matrix (Drakouli et al., 2017; Nikolakaki et al., 1997). |

**Table B. Subdivisions of regulatory factors associated with the *Myc* target P1/P2 (plotted in Figure 12)**

| Gene name | Transcriptional Category | Activity/Function |
|---|---|---|
| HmgD | **TF; POLIITR; DBTR2; CRTR2** | **High mobility group protein D, isoform C & High mobility group protein C, isoform D**, EGFR signaling (Anan Ragab & Travers, 2006); ecdysone/ecdysteroid signaling pathway (Chen et al., 2008); Dpp pathway: HmgD binding to the Dpp-responsive enhancer of (*tinman*) as well as to the Tinman protein during *Drosophila* cardiogenesis (Zaffran, 2002); Wnt/TCF signaling (Archbold et al., 2014); a highly abundant chromosomal protein involved in DNA binding, bending and chromatin organization (Dragan et al., 2003); minor groove of Adenine-Thymine-rich DNA binding (Churchill et al., 1995); High Mobility Group Box Transcription Factors. |
| CtBP, isoform G | **COREP; DBTFB; COACT; CRTR2; POLIITR** | **C-terminal binding protein, isoform G**, corepressor targeting diverse transcription regulators; Hairy (*hry*)-interacting protein; positive regulator of Wnt/TCF pathway (Bhambhani et al., 2011; Zhang and Arnosti, 2011; Fang et al., 2006); negative regulation of canonical Wnt signaling pathway (Bhambhani et al., 2011; Fang et al., 2006); Wingless signaling pathway involved in embryo segmentation (Chan et al., 2008); chromatin remodeling (Emelyanov et al., 2012); DNA binding transcription factor binding (Qi et al., 2008); transcription coactivator & corepressor (Fang et al., 2006); regulation of transcription by RNA pol II (Zhang and Arnosti, 2011); TORC Remodeling Complex. |
| REPTOR | **TF; POLIITR; DBTR2** | **Repressed by TOR**, a transcription factor, shuttles between cytoplasm & nucleus depending on the state of the activity of mechanistic Target of rapamycin (*mTor*). Together with its binding partner REPTOR-BP, it mediates much of the transcriptional response observed upon Tor complex 1 inhibition; response to starvation; TORC1 signaling; positive regulation of transcription (Tiebe et al., 2015); DNA binding transcription factor activity (GO Reference Genome Project, 2011-; InterPro Project Members, 2004-); Basic Leucine Zipper Transcription Factors. |
| Tlk | **CRR; CHRM** | **Tousled-like kinase**, conserved anti-silencing function protein 1 (ASF1)/Tousled-like kinase (TLK), coordination of cell cycle phases via chromatin organization (Carrera et al., 2003); cell migration/apoptosis pathway (Zhang Y, Cai R, Zhou R, Li Y, Liu L. 2016); association of Tousled-like kinase with the protein complex of Wingless signaling regulators (Milan et al. 1998); chromosome segregation (Li et al., 2009). |

**Table B. Subdivisions of regulatory factors associated with the *Myc* target P1/P2 (plotted in Figure 12)**

| Gene name | Transcriptional Category | Activity/Function |
|---|---|---|
| 14-3-3zeta | **POLIITR; COACTB; CHRM** | **14-3-3 protein zeta,** Ras/Raf/MAPK signaling-dependent photoreceptor development (Kockel et al., 1997); regulation of Yorkie nuclear localization by Hippo (Hpo) signaling pathway; transcription coactivator binding (Ren et al., 2010); chromosome segregation (Su et al., 2001). |
| piwi | **DBTR2; CHRSILEN; PSILEN; REP; HF; RGEXPPT** | **P-element induced wimpy testis, isoform B**, repression of transposable elements during meiosis by complexes of piRNAs/Piwi containing proteins, methylation & repression of transposons (Sienski et al., 2015); chromatin silencing (Brower-Toland et al., 2007; Grimaud et al., 2006; Pal-Bhadra et al., 2004; Pal-Bhadra et al., 2002); gene silencing by RNA, ncRNA mediated gene silencing (Le Thomas et al., 2013); heterochromatin organization involved in chromatin silencing (Sienski et al., 2015; Sienski et al., 2012); heterochromatin formation (Brower-Toland et al., 2007; Grimaud et al., 2006; Pal-Bhadra et al., 2004; Pal-Bhadra et al., 2002); DNA methylation-dependent heterochromatin formation (Yu et al., 2015); Argonaut Endoribonucleases. |
| SMC3 | **CBRR; DBRR; CHRM; RRR** | **Structural maintenance of chromosomes 3**, a subunit of the cohesin complex, involved in planar cell polarity by regulating the membrane enrichment of the transmembrane cadherin encoded by "starry night" (*stan*); mitotic sister chromatid cohesion; cell division (GO Reference Genome Project, 2011-); Notch Wnt/Frizzled-PCP signaling: establishment of imaginal disc-derived wing hair orientation; imaginal disc-derived wing morphogenesis (Mouri et al., 2012); chromatin binding, double-stranded DNA binding (Gene Ontology Curators, 2002-); chromosome organization and DNA condensation (InterPro Project Members, 2004-). |
| Trl | **TF; CRTR2; DBTR2; POLIITR; HF** | **Trithorax-like**, a GAGA transcription factor involved in chromatin modification; chromatin binding (Chopra et al., 2008); chromatin remodeling; core promoter sequence-specific DNA binding (Agianian et al., 1999); RNA pol II-specific transcription activator activity (Rieder et al., 2017); DNA-binding activity (Read et al., 2000); POZ domain binding (Bonchuk et al., 2011); heterochromatin formation (Muolland et al., 2003); positive regulation of transcription by RNA pol II (Tsai et al., 2016; Chopra et al., 2008); negative regulation of DNA-templated transcription (Chen et al., 2009); dosage compensation and gametogenesis; Dpp & EGFR signaling, imaginal disc-derived wing morphogenesis (Dworkin and Gibson, 2006); syncytial mitotic cell cycle (Bhat et al., 1996). |

**Table B. Subdivisions of regulatory factors associated with the *Myc* target P1/P2 (plotted in Figure 12)**

| Gene name | Transcriptional Category | Activity/Function |
|---|---|---|
| Taf5 | **POLIITR; DBTR2; COACT; TPI; TI** | **TBP-associated factor 5**, part of the multisubunit basal transcription factor TFIID & important for its assembly or stability; RNA pol II general transcription initiation factor activity; transcription factor TFIID complex (Wright et al., 2006; Hansen and Tjian, 1995). |
| CG34159 (LD45253) | **COREP; POLIITR; CRTR2** | **MAPK-regulated corepressor-interacting protein**, orthologous to human MCRIP2 (MAPK regulated corepressor interacting protein 2) (Gelbart and Emmert, 2013); MCRIP1 binds to and inhibits the transcriptional co-repressor CtBP; MCRIP1 is an ERK substrate; when phosphorylated by ERK, MCRIP1 releases CtBP to induce chromatin modifications (Ichikawa et al., 2015). |
| Rpt4 | **POLITR; DBTR1** | **Regulatory particle triple-A ATPase 4**, 19S proteasomal ATPase, a component of the 26S proteosome complex, nucleolar protein & regulator of rRNA transcription; physical interaction with the tumor suppressor protein "Birt Hogg Dube" (BHD), RNA pol I transcription, regulatory region sequence-specific DNA binding (association of Rpt4 with the rDNA locus) (GO Reference Genome Project, 2011-); Notch-mediated follicle cell differentiation and cell cycle switches, insulin-PI3K pathway (Jia et al., 2015). |
| Chro | **CBTR2; CRTR2; POLIITR; CHRM** | **Chromator**, a chromodomain protein, chromatin binding; chromatin organization; negative regulation of transcription by RNA pol II (GO Reference Genome Project, 2011-); chromosome organization (Ding et al., 2009; Rath et al., 2006); microtubule spindle formation, normal cell cycle progression, functioning as a spatial regulator of cell cycle factors (GO Reference Genome Project, 2011-); metamorphosis (Wasser et al., 2007). |
| CG6683 | **POLIITR** | Involved in regulation of transcription by RNA pol II; part of transcription regulator complex (GO Reference Genome Project, 2011-), active in nucleus; MAD-BESS domain Transcription Regulators. |
| SMC5 | **DBRR; CHRM; RRR** | **Structural maintenance of chromosomes protein 2**, a component of the SMC5/6 protein complex, critical to genome stability & required for homologous DNA recombination-based processes; single-stranded DNA binding (GO Reference Genome Project, 2011-); damaged DNA binding (Gene Ontology Curators, 2002-); DNA damage response (Li et al., 2013). |

**Table B. Subdivisions of regulatory factors associated with the *Myc* target P1/P2 (plotted in Figure 12)**

| Gene name | Transcriptional Category | Activity/Function |
|---|---|---|
| gro | **COREP; POLIITR** | **Groucho, isoform F**, a global developmental corepressor in collaboration with DNA-binding repressor partner proteins & tethering to target promoters (Ajuria et al., 2011; Giagtzoglou et al., 2003; Jimenez et al., 2000; Goldstein et al., 1999; Valentine et al., 1998); downstream effector of signaling pathways such as Wg/Wnt & Dpp/TGF-beta; phosphorylation & attenuation of Groucho repressor activity in response to MAPK activation; "context dependent regulatory domain binding" (CRD), a domain of about 130aa, the most divergent region among the LEF/TCF proteins (Arce et al., 2009). |
| Polr2A (RpII215, RPB1) | **RPIIB1; DBTR2; POLIITR; DBRR; TI; RRR** | **RNA polymerase II subunit A (RPB1)**, DNA-directed 5'-3' polymerase activity (Brickey and Greenleaf, 1995; Zehring et al., 1988); RNA pol II, core complex (Gu et al., 2002; Greenleaf, 1983); largest and catalytic component of RNA pol II, synthesizes mRNA precursors and many functional non-coding RNAs; forms the polymerase active center together with the second largest subunit; central component of the basal RNA polymerase II transcription machinery. |
| BEAF-32 | **INSUL; POLIITR; DBTR2; RGEXPTR2; CBTR2; CRTR2** | **Boundary element-associated factor of 32kD**, a DNA-binding protein with binding sites near transcription start sites; roles include chromatin domain insulator function, chromatin insulator sequence binding (Cuvier et al., 1998; Zhao et al., 1995); chromatin organization (Emberly et al., 2008; Roy et al., 2007); gene regulation and genome organization, DNA binding (Ogiyama et al., 2018; Hart et al., 1997); regulation of transcription by RNA pol II (Zhao et al., Zhao et al., 1995 1995). |
| mre11 | **DBRR; RRR; TMM; CHRM** | **Meiotic recombination 11**, Double-strand break repair protein; G2/M DNA damage checkpoint signaling (Bi et al., 2006); chromosome organization (Bi et al., 2004; Ciapponi et al., 2004); telomere capping/maintenance (Gao et al., 2009; Bi et al., 2005; Bi et al., 2004; Ciapponi et al., 2004). |
| PolE1 (DNApol-epsilon, DNApol-ε255) | **DNAPOLE1; RRR; DBRR** | **DNA polymerase epsilon subunit 1** (DNA Pol-epsilon, DNA Pol-ε, DNA Pol-ε255), the large subunit of DNA pol epsilon, an essential DNA polymerase participating with DNA pol alpha & DNA pol delta in cellular DNA replication. The C-terminal and N-terminal regions have differential requirements in mitotic and endo-replicating cells; 3'-5' DNA exonuclease activity (Oshige et al., 2004; Aoyagi et al., 1997); DNA replication proofreading (Oshige et al., 2004); endomitotic cells (Suyari et al., 2012). |

**Table B. Subdivisions of regulatory factors associated with the *Myc* target P1/P2 (plotted in Figure 12)**

| Gene name | Transcriptional Category | Activity/Function |
|---|---|---|
| Polr1C | **DBTR1; POLITR; DBTR3; POLIIITR** | **RNA polymerase I and III subunit C**, DNA-directed 5'-3' RNA polymerase activity and protein dimerization activity; tRNA transcription by RNA polymerase III; transcription by RNA polymerase I (Gene Ontology Curators, 2002-); part of RNA polymerase I complex and RNA polymerase III complex. |
| caz, dFUS, Sarcoma-associated RNA-binding fly homolog, P19, SARFH | **CBTR2; RNABPR; COACT; TI; POLIITR; RGEXPPR; TPI** | **Cabeza** ('cabeza' means 'head' in Spanish), a chromatin binding protein (Mallik et al., 2018); RNA-binding protein, mRNA processing (Lasko, 2000); a single ortholog of human FUS in *Drosophila*; mRNA splicing (Herold et al., 2009); transcription initiation by RNA pol II promoter (Aoyagi and Wassarman, 2000); transcription coregulatory activity (GO Reference Genome Project, 2011-); synaptic assembly at the neuromuscular junction (Azuma et al., 2014); Cabeza shares homology domains with EWS & TLS, two human genes involved in chromosomal translocations with sarcoma formation; enriched in *Drosophila* head (David T. Stolow and Susan R. Haynes, 1995); Transcription Factor TFIID complex. |
| Mcm3 | **DBRR; RRR; DNAHRR** | **Minichromosome maintenance 3**, a component of the MCM2-7 hexamer, forms part of the CMG complex, together with the product of CDC45L & GINS proteins. The CMG complex is the main DNA helicase (Moyer et al., 2006), functions during DNA replication; DNA binding; ssDNA binding; duplex DNA unwinding; DNA replication initiation (InterPro Project Members, 2004-). |
| Rfc37 (CG8142) | **RRR; DBRR** | **Replication factor C37**, ATP binding activity; ATP hydrolysis activity; DNA binding activity (InterPro Project Members, 2004-): Predicted to contribute to DNA clamp loader activity (GO Reference Genome Project, 2011-). Predicted to be involved in DNA repair and DNA-templated DNA replication. Part of Elg1 RFC-like complex. |
| Fkbp39 | **CRR; CBRR; RRR** | **FK506-binding protein 39kD**, acts as an inhibitor of autophagy in larval fat body; juvenile hormone response element binding ("JHRE binding") (Li et al., 2007); FK506-binding (Theopold et al., 1995); nucleoplasmin-like NPL domain signature, present in *Drosophila* FKBP39 and a large number of chromatin-related proteins/chromatin binding & chromatin regulation: histone chaperone related to DNA activity during transcription, replication and repair (Edlich-Muth et al., 2015). |

**Table B. Subdivisions of regulatory factors associated with the *Myc* target P1/P2 (plotted in Figure 12)**

| Gene name | Transcriptional Category | Activity/Function |
|---|---|---|
| mtSSB | **MTRR; DBMTRR** | **Mitochondrial single stranded DNA-binding protein**, replication of mitochondrial DNA (Farr et al., 2004); single-stranded DNA binding (Ciesielski et al., 2015; Farr et al., 2004; Stroumbakis et al., 1994). |
| Su(var)205 | **CBTR2; DBTR2; POLIITR; REP** | **Suppressor of variegation 205**, structural component of heterochromatin, involved in gene repression and the modification of position-effect-variegation; recognizes & binds histone H3 tails methylated at 'Lys-9', leading to epigenetic repression; chromatin binding (Sienski et al., 2015; Johansson et al., 2007); pol II transcription (Liu et al., 2005; Kellum, 2003); Heterochromatin I family. |
| BcDNA:RE18748 (CG15390) | **DBMTTR; MTTR; MTTT** | Double-stranded DNA binding activity; involved in regulation of DNA-templated transcription (InterPro Project Members, 2004-); orthologous to human MTERF4 (mitochondrial transcription termination factor 4); mitochondrial/chloroplastic transcription termination; mitochondrial transcription termination (TTM). |
| PolD2 (Pol31) | **DNAPOLD2; RRR; DBRR** | **DNA polymerase delta subunit 2**, **DNA polymerase delta subunit 2**, a component of DNA pol delta complex; DNA strand elongation involved in DNA replication; mitotic DNA-templated DNA replication (Ji et al., 2019); a component of DNA pol zeta complex (Gene Ontology Curators, 2002-); high fidelity genome replication, including lagging strand synthesis, DNA recombination & repair; promotes the function of the DNA pol-delta complex accessory subunit PolD3 in both embryonic and post-embryonic somatic cells (Ji et al., 2019); DNA binding; DNA replication (InterPro Project Members, 2004-); DNA strand elongation involved in DNA replication (GO Reference Genome Project, 2011-). |
| lid (Kdm5) | **DBTR2; ACT; POLIITR; TF; CRTR2** | **Lysine-specific demethylase 5 (lid)**, activator of transcription (JAK/STAT) signaling (Tarayrah et al., 2015); regulation of cell growth, circadian rhythm, stress resistance, hematopoiesis & fertility; male germline stem cell population maintenance (Tarayrah et al., 2015; Lloret-Llinares et al., 2008; Secombe et al., 2007); histone H4R3 demethylase activity, histone reader activity (Liu and Secombe, 2015); pol II transcription (Secombe et al., 2007) H3K4m / H3K4m2 / H3K4M3 demethylase activity (Tarayrah et al., 2015; Lloret-Llinares et al., 2008; Secombe et al., 2007); chromatin remodeling (GO Reference Genome Project, 2011-); DNA binding (InterPro Project Members, 2004-). |

**Table B. Subdivisions of regulatory factors associated with the *Myc* target P1/P2 (plotted in Figure 12)**

| Gene name | Transcriptional Category | Activity/Function |
|---|---|---|
| La (CG10922) | **TT; POLIIITR; DBTR3; RNABTR3** | **La autoantigen-like**, involved in transcription termination by RNA polymerase III. Binds RNA & DNA. Binds to precursors of RNA polymerase III transcripts. May play a specialized role during fly development |
| Orc6 | **RRR; DBRR** | **Origin recognition complex subunit 6**, a subunit of the origin recognition complex (ORC), which is essential for the initiation of DNA replication in eukaryotic cells (Chesnokov et al., 1999); mitotic cell cycle (Balasov et al., 2009). |
| mtTFB2 | **MTTF; TFB2M; MTTR; DBMTTR; TI; MTRR; DBMTRR** | **Mitochondrial Transcription Factor B2**, dimethylation of mitochondrial 12S rRNA at the conserved stem loop; required for basal transcription of mitochondrial DNA (Gene Ontology Curators, 2002-); regulation of mitochondrial DNA copy number; stimulation of transcription independent of the methyltransferase activity; mitochondrial transcription (Adan et al., 2008). |
| Prim2 | **PRIM2; RRR; DBRR** | **DNA primase subunit 2**, a regulatory subunit of the DNA polymerase alpha-primase complex, synthesis of short RNA-DNA primers (Kuroda et al., 1990; Cotterill et al., 1987; Kaguni et al., 1983) on the lagging strand during DNA synthesis; mitotic cell cycle (Chen et al., 2000); DNA polymerase Alpha Primase. |
| vas | **RNABTN; RNAHTN; TN; RGEXPTN** | **Vasa**, a DEAD-Box RNA helicase protein (Mahowald, 2001; van Eeden and St. Johnston, 1999; Cooperstock and Lipshitz, 1997), promotes translation of (*grk*) & (*mei-26*) mRNAs (Liu et al., 2009); also functions in piRNA biogenesis as a component of an amplifier complex. maternal transcripts required for oogenesis, transposon silencing in the female germ line, A-P embryonic patterning, & germ cell specification; RNA binding (Sengoku et al., 2006); DEAD Box RNA Helicases. |
| His3 | **DBRR; DBTR1; DBTR2; DBTR3; POLITR; POLIITR; POLIIITR; RRR; CHRM** | **Histone 3**, core component of nucleosome, nucleosomes wrap and compact DNA into chromatin, limiting DNA accessibility to the cellular machineries requiring DNA as a template; histones play a central role in transcription regulation, DNA repair, DNA replication & chromosomal stability; regulation of DNA accessibility via a complex set of post-translational modifications of histones, also called histone code, & nucleosome remodeling; DNA binding (InterPro Project Members, 2004-); nucleosome DNA binding (Gene Ontology Curators, 2002-). |

**Table B. Subdivisions of regulatory factors associated with the *Myc* target P1/P2 (plotted in Figure 12)**

| Gene name | Transcriptional Category | Activity/Function |
|---|---|---|
| XRCC1 | **RRR; DBRR** | **X-ray repair cross complementing 1**, damaged DNA binding activity; involved in base-excision repair; active in nucleus; single strand break repair; double-strand break repair via nonhomologous end joining (InterPro Project Members, 2004-). |
| Top2 | **POLITR; POLIITR; POLIIITR; RRR; DBRR; DBTR1; DBTR2; DBTR3; CBRR** | **Topoisomerase 2**, alteration of the topology of DNA by creating a transient DNA double strand breakage; essential for removing supercoils generated via DNA replication & transcription; decatenating replicated sister chromosomes during cell division (Ducat et al., 2008); chromatin binding; DNA binding; rDNA binding; satellite DNA binding (Blattes et al., 2006); sister chromatid segregation (Chang et al., 2003; Coelho et al., 2003). |
| Bin1 (SAP18) | **COREP; HDAC; HF; CBTR2; POLIITR; RNABTR2** | **Histone deacetylase complex subunit SAP18 (Bicoid interacting protein 1 Bin 1)**, regulation of hedgehog (Hh) signaling pathway by transcription factor Gli in mammals, repression of Gli-mediated transcription by Su(fu) in cooperation with SAP18 for the recruitment of the SAP18-mSin3 complex to promoters containing the Gli-binding element (Yan Cheng and Bisho, 2002); transcription corepressor activity; heterochromatin formation (Matyash et al., 2009); negative regulation of DNA-templated transcription (Matyash et al., 2009; Zhu et al., 2001). |
| POLDIP2 | **POLDIP2; DBRR; RRR** | **Polymerase (DNA-directed), delta interacting protein 2**, DNA binding activity (InterPro Project Members, 2004-); involved in error-free translesion synthesis (GO Reference Genome Project, 2011-); active in mitochondrial nucleoid and nucleus. |
| Dp1 | **TN; RGEXPTN; CBTR2; DBTR2** | **Dodeca-satellite-binding protein 1**, translation enhancer; ssDND binding (Cortes and Azorin, 2000; Cortes et al., 1999); mRNA 3'-UTR binding (Nelson et al., 2007); satellite DNA binding (Cortes and Azorin, 2000); chromatin condensation (Huertas et al., 2004); chromatin formation (Wang et al., 2005). |
| Prim1 | **RRR; PRIM1; DBRR** | **DNA primase subunit 1, DNA polymerase α**, a subunit of the DNA-polymerase alpha complex, together with the subunit Primase2 responsible for primase activity (synthesis of short RNA primers) (Bakkenist and Cotterill, 1994) required during DNA replication to prime DNA synthesis at origins, also on the lagging strand during Okazaki fragment synthesis (Bakkenist and Cotterill, 1994; Kuroda et al., 1990; Cotterill et al., 1987; Kaguni et al., 1983). |

**Table B. Subdivisions of regulatory factors associated with the *Myc* target P1/P2 (plotted in Figure 12)**

| Gene name | Transcriptional Category | Activity/Function |
|---|---|---|
| Ciz1 | **RRR; DBRR** | **Ciz1, Cip1 Interacting Zinc Finger Protein 1**, nuclear protein; negative regulation of cell cycle, regulation of the subcellular localization of p21(CIP/WAF1); consensus DNA sequence, ARYSR(0–2)YYAC, recognized by Ciz1 (Warder and Keherly, 2003); no evidence found in transcriptional regulation (Mitsui et al., 1999); DNA replication factor (Lukasik et al., 2008). |
| hang | **DBTR2; POLIITR; TF; RNABTR2** | **Hangover**, a nuclear zinc finger protein, DNA binding transcription factor activity, pol II-specific (GO Reference Genome Project, 2011-); mRNA binding (Ruppert et al., 2017); C2H2 Zinc Finger Transcription Factor. |
| row | **CBTR2; DBTR2; POLIITR; TF** | **Relative of woc**, a zinc-finger transcription factor involved in transcription regulation, required for the (HP1) to bind chromatin (Font-Burgada et al., 2008); sequence-specific DNA binding; pol II transcription (GO Reference Genome Project, 2011-); regulation of transcription by RNA pol II (Abel et al., 2009; Font-Burgada et al., 2008); Zinc Finger C2H2 TYPE. |
| Ndf | **DBTR1; DBTR2; DBTR3; POLITR; POLIITR; POLIIITR; CBTR2; TE; CRTR2** | **Nucleosome-destabilizing factor**, a putative H3K36me3 binding protein, which binds active gene bodies, DNA binding, nucleosome binding, chromatin binding, pol II transcription elongation-coupled chromatin remodeling (Fei et al., 2018). |
| dpa | **DBRR; RRR; DNAHRR** | **disc proliferation abnormal**, 3'-5' DNA helicase activity (Moyer et al., 2006); DNA binding (InterPro Project Members, 2004-); DNA replication (Pflumm and Botchan, 2001); mitotic DNA replication initiation (GO Reference Genome Project, 2011-); essential for 'once per cell cycle' DNA replication initiation and elongation in eukaryotic cells; required for DNA replication and cell proliferation. Essential role in mitotic DNA replication but not in endoreplication; MCM2-7 complex. |
| Nse4 | **RRR** | **Non-SMC element 4** (non-structural maintenance of chromosomes element 4), a member of the EID (EP300-interacting inhibitor of differentiation) family of proteins, is the kleisin component of the SMC5/6 protein complex; critical role in genome stability; required for homologous DNA recombination-based processes; DNA repair (GO Reference Genome Project, 2011-). |

**Table B. Subdivisions of regulatory factors associated with the *Myc* target P1/P2 (plotted in Figure 12)**

| Gene name | Transcriptional Category | Activity/Function |
|---|---|---|
| Ssl1 | **POLIITR; TI; DBTR2; RRR; TPI** | **Suppressor of stem-loop mutation**, zinc ion binding activity, involved in transcription by RNA polymerase II (Fregoso et al., 2007); part of transcription factor TFIIH holo complex; orthologous to human GTF2H2C (GTF2H2 family member C); GTF2H2C_2 (GTF2H2 family member C, copy 2); & GTF2H2 (general transcription factor IIH subunit 2); DNA repair (InterPro Project Members, 2004-); nucleosome excision repair; transcription initiation at RNA pol II promoter and transcription open complex formation (Aoyagi and Wassarman, 2000). |
| Polr2H | **RPIIB8; DBTR1; POLITR; DBTR2; POLIITR; DBTR3; POLIIITR** | **RNA polymerase II, I and III subunit H**, RNA pol I, RNA pol II (GO Reference Genome Project, 2011-; Aoyagi and Wassarman, 2000), and RNA pol III activity (GO Reference Genome Project, 2011-); transcription by RNA pol I, pol II and pol III; part of RNA polymerase I complex; RNA polymerase II, core complex; and RNA polymerase III complex. |
| vig2 | **CHRSILEN; CRTR2; HF; RNABPT** | **Vig2**, heterochromatin organization (formation), histone H3-K9 methylation & chromatin silencing regulation; Vig2 & its homolog are components of RISC complex (Gracheva et al., 2009); Hyaluronan/mRNA-binding protein, chromatin remodeling, regulation of transcription (Nery et al., 2006; Nery et al., 2004). |
| vig | **HF; RGEXPPT; RNABPT** | V**asa intronic gene**, RNA interference & heterochromatin organization; Vig and its homolog are components of RISC complex (Gracheva et al., 2009); ncRNA-mediated post-transcriptional gene silencing, ncRNA-mediated post-transcriptional gene silencing (Caudy et al., 2002); RNA binding (GO Reference Genome Project, 2011-). |
| rad50 | **CHRM; RRR; TMM; DBRR** | **RAD50**, SMC family & RAD50 subfamily required for double-strand break repair; double-stranded DNA binding (Gorski et al., 2004) & telomere maintenance via recombination (Ciapponi et al., 2004); chromosome organization, DNA repair (Ciapponi et al., 2004); G-quadruplex DNA binding, single stranded & double stranded telomeric DNA binding (GO Reference Genome Project, 2011-). |
| koi | **RRR; DBRR** | **Klaroid**, double-strand break repair via homologous recombination (Ryu et al., 2015); nuclear migration (Kracklauer et al., 2007); nuclear organization (Tan et al., 2018; Elhanany-Tamir et al., 2012). |

**Table B. Subdivisions of regulatory factors associated with the *Myc* target P1/P2 (plotted in Figure 12)**

| Gene name | Transcriptional Category | Activity/Function |
|---|---|---|
| Mcm7 | **DBRR; RRR; DNAHRR** | **Minichromosome maintenance 7**, a component of the MCM2-7 hexamer, forms part of the CMG complex, together with CDC45L & GINS proteins. CMG complex is the main DNA helicase, functions during DNA replication; DNA binding; DNA helicase activity; single-stranded DNA binding; duplex DNA binding (InterPro Project Members, 2004-); DNA replication initiation (GO Reference Genome Project, 2011-). |
| PolD1 (DNApol-delta, DNApol-δ) | **RRR; DBRR; DNAPOLD1** | **DNA polymerase delta catalytic subunit**, 3'-5' exonuclease activity and DNA polymerase activity, DNA replication proofreading (Aoyagi et al., 1994); catalytic component of the DNA polymerase delta complex, plays a crucial role in high fidelity genome replication, including lagging strand synthesis; DNA recombination & repair (Aoyagi et al., 1994; Chiang et al., 1993; Peck et al., 1992); required at the nucleus of rapidly dividing embryonic cells to activate genome replication during the earliest cell division (Ji et al., 2019); base-excision repair gap-filling (GO Reference Genome Project, 2011-); delta DNA polymerase complex (Ji et al., 2019; Peck et al., 1992). |
| RecQ4 | **RRR; DBRR** | **RecQ4 helicase**, ATP-dependent hydrolytic DNA strand separation and exchange, playing a critical role in replication and repair; 3'-5' helicase activity (Capp et al., 2009); four-way junction helicase activity (GO Reference Genome Project, 2011-); nucleic acid binding (InterPro Project Members, 2004-; (Xu et al., 2009). |
| TFAM | **MTTF; TFAM; DBMTTR; MTTR; MTRR; DBMTRR** | **Mitochondrial transcription factor A**, essential for mtDNA transcription & replication; binds to mtDNA nonspecifically; plays a role in mtDNA maintenance by packaging mtDNA; mitochondrial promoter sequence-specific DNA binding (Gene Ontology Curators, 2002-); transcription *cis*-regulatory region binding (GO Reference Genome Project, 2011-);High Mobility Group Box Transcription Factors. |
| Cpsf160 | **TT; PAFAC; RNABPR** | **Cleavage and polyadenylation specificity factor 160**, a key role in pre-mRNA 3'-end formation, recognizing the AAUAAA signal sequence and interacting with poly(A) polymerase and other factors to bring about cleavage and poly(A) addition (Salinas et al., 1998); mRNA polyadenylation (Mount and Salz, 2000); polyadenylation factors. |

**Table B. Subdivisions of regulatory factors associated with the *Myc* target P1/P2 (plotted in Figure 12)**

| Gene name | Transcriptional Category | Activity/Function |
|---|---|---|
| CG11164 | **RGEXPPR** | **Nucleoplasmic protein**, involved in RNA catabolic process (part of ribonuclease H2 complex) (GO Reference Genome Project, 2011-). |
| Msh6 (CG7003) | **RRR; MISDB** | **Msh6 (DNA mismatch repair protein Msh6)**, detection of base-base mismatches & small insertion/deletion loops; recruitment of the rest of the mismatch repair machinery; mismatched DNA binding; ATP-dependent DNA damage sensor activity (InterPro Project Members, 2004-); meiotic mismatch repair; MutSalpha complex (GO Reference Genome Project, 2011-). |
| Mad | **DBTR2; POLIITR; TF; REP; ACT; COACT** | **Mothers against decapentaplegic homolog (dpp)**, the primary transcription factor, BMP signaling pathway core component (Vuilleumier et al., 2022; Guo et al., 2013; Weiss et al., 2010; Kamiya et al., 2008; Yao et al., 2006; Muller et al., 2003; Dai et al., 2000; Das et al., 1998; Inoue et al., 1998); wing development by EGFR & BMP signaling (Dworkin and Gibson, 2006; Lecuit et al., 1996); Wg/Wingless signaling (Bradley et al., 2001); DNA binding transcription activator activity, pol II specific (Vuilleumier et al., 2022; Weiss et al., 2010; Saller and Bienz, 2001); DNA binding transcription repressor activity, RNA pol II specific (Yao et al., 2006; Muller et al., 2003); RNA pol II *cis*-regulatory region sequence-specific DNA binding (Xu et al., 1998); transcription co-activator activity (Dai et al., 2000). |
| Cp190 | **TF; DBTR2; CBTR2; INSULB** | **Centrosomal protein 190kD**, C2H2 zinc finger transcription factor, binds to most boundaries of contact domains; defined by enhanced internal contact frequencies; forms most contact domain boundaries distal to a transcribed promoter; recruited to DNA by proteins such as TCTF and Su(Hw) & associates with them (Sabirov et at., 2021; Maksimenko et al., 215); essential for early development, prevents regulatory cross-talk between specific gene loci that pattern the embryo; chromatin binding (Whitfield et al., 1995); chromatin insulator sequence binding (Erokhin et al., 2010; Oliver et al., 2010; Mohan et al., 2007); DNA binding (Pai et al., 2004); POZ domain binding (Bonchuk et al., 2011). |
| Taf6 | **POLIITR; TF; TI; COACT; TPI** | **TBP-associated factor 6**; part of the multisubunit basal transcription factor TFIID; pol II general transcription initiation factor (Wright et al., 2006; Hansen and Tjian, 1995); transcription co-activator activity (GO Reference Genome Project, 2011-); transcription by RNA pol II (Hansen and Tjian, 1995); Transcription Factor TFIID. |

**Table B. Subdivisions of regulatory factors associated with the *Myc* target P1/P2 (plotted in Figure 12)**

| Gene name | Transcriptional Category | Activity/Function |
|---|---|---|
| Dmel\CG5543 | **U** | **Gastrulation defective protein 1 homolog**, Active in nucleus, located at the site of double-strand break. |
| pds5 | **CHRM; ACT; REP** | **(precocious dissociation of sisters 5)**, sister chromatid separation at mitosis by removing the cohesin ring complex from chromosomes (Gause et al., 2010; Dorsett et al., 2005); influences gene activation & silencing through interactions with cohesin; required to initiate and/or maintain sister chromatid cohesion; TGF-alpha (ATR/Mei-41 kinase) (Barbosa et al., 2007); Armadillo-Like helical. |
| tplus3a | **POLIITR; DBTR2** | **(testis-specific Plus3 domain a)**, RNA pol II CTD phosphoserine binding; part of Cdc73/Paf1 complex; DNA binding (InterPro Project Members, 2004-); RNA pol II C-terminal domain phosphoserine binding (GO Reference Genome Project, 2011-). |
| Sas-6 | **RRR** | **Spindle assembly abnormal 6**, a centriole protein, essential for centriole assembly, homo-oligomerizes to form a 9-fold symmetric "cartwheel" structure, plays an important part in setting the 9-fold symmetry of the assembling centriole (Jana et al., 2018); centriole replication (Stevens et al., 2010; Dobbelaere et al., 2008; Peel et al., 2007); centriole duplication (Gene Ontology Curators, 2002-). |
| Taf12 | **POLIITR; DBTR2; COACT; TPI; TI** | **TBP-associated factor 12**, part of the multisubunit basal transcription factor TFIID; pol II general transcription initiation factor (Yokomori et al., 1993) forms a histone-like pair with Taf4; Taf12 also an integral component of the *Drosophila* SAGA histone acetyltransferase complex; pol II preinitiation complex assembly (GO Reference Genome Project, 2011-); SAGA complex; TFIIFD complex. |
| tplus3a | **POLIITR; DBTR2** | **(testis-specific Plus3 domain a)**, RNA pol II C-terminal domain phosphoserine binding activity; part of Cdc73/Paf1 complex; DNA binding (InterPro Project Members, 2004-); RNA pol II C-terminal domain phosphoserine binding (GO Reference Genome Project, 2011-). |
| nonA-l | **POLIITR; RNABTR2** | **nonA-like**, nuclear protein, mRNA binding activity (Lasko, 2000); regulation of DNA-templated transcription (GO Reference Genome Project, 2011-); C-terminal to the RNA recognition motifs (RRMs). |
| CG6418 (DmRH27) | **RNAHPR; RNABPR** | RNA helicase, nuclear RNA binding; mRNA splicing (InterPro Project Members, 2004-; Gene Ontology Curators, 2002-); DEAD-BOX RNA HELICASES. |

**Table B. Subdivisions of regulatory factors associated with the *Myc* target P1/P2 (plotted in Figure 12)**

| Gene name | Transcriptional Category | Activity/Function |
|---|---|---|
| CG12134 | **U** | Uncharacterized, unknown; WD40/YVTN Repeat-like Containing Domain Superfamily. |
| su(sable), su(s) | **POLIITR; TT; SUP; RGEXPPR** | **suppressor of sable**, su(sable) & Wdr82, part of a transcription termination checkpoint; promotes transcription termination of RNAs & their subsequent degradation by the nuclear exosome, negative regulation of transcription termination (Brewer-Jensen et al., 2016); negative regulation of transcription (suppressor) (Kuan et al., 2004). |
| nclb | **POLITR; POLIIITR; TE; CBTR1; CBTR3; TPI** | **no child left behind**, chromatin DNA binding (Casper et al., 2011); pol I & poI III transcription; regulator of the pol I-elongation factor TFIIH (Liu et al., 2017); chromatin-associated factor (Caf), regulates transcription. |
| B52 | **POLIITR; TI; CRTR2** | **B52**, regulation of gene expression; regulation of transcription start site selection at RNA pol II promoter (Bradley et al., 2015); condensation or decondensation of chromatin; associated with boundaries of transcriptionally active chromatin (Champlin et al., 1991). |
| tst | **RNABPR; RNAHPR; RGEXPPR** | (**twister**) RNA helicase, helicase activity (Lasko, 2000); RNA helicase activity; RNA catabolic process (InterPro Project Members, 2004-); nuclear-transcribed mRNA catabolic process, 3'-5' exonucleolytic nonsense-mediated decay (GO Reference Genome Project, 2011-); regulation of mRNA alternative splicing (Park et al., 2004); SKI2-like RNA helicases. |
| hfp | **POLIITR; RGEXPPR; REP** | (**half pint**), a single stranded nucleic acid binding protein with roles in transcription & RNA splicing; functions in *Myc* transcriptional repression & cell growth control, via interaction with the transcription factor Haywire (*hay*). |
| Polr2B (RpII140) | **RPIIB2; POLIITR; TI; TE; DBTR2** | **RNA polymerase II subunit B**, present in all RNA pol II complexes; DNA-directed 5'-3' RNA polymerase activity (Falkenburg et al., 1987); RNA pol II activity (Aoyagi and Wassarman, 2000). |
| CAP-D2 | **CBRR; CHRM; DBRR; RRR** | **CAP-D2 condensin subunit**, chromatin binding; mitotic chromosome condensation (Savvidou et al., 2005). |
| Rm62 | **RNAHPR; POLIITR; RGEXPPR** | **Rm62**, RNA helicase; pre-mRNA splicing, alternative splicing, rRNA processing, miRNA processing, and transcription regulation (Ishizuka et al., 2002); DEAD-BOX RNA HELICASES. |

**Table B. Subdivisions of regulatory factors associated with the *Myc* target P1/P2 (plotted in Figure 12)**

| Gene name | Transcriptional Category | Activity/Function |
|---|---|---|
| X16 | **POLIITR; RGEXPPT; RNABPR; DBTR2** | **x16 splicing factor**, involved in mRNA splicing and RNA metabolism regulation; regulation of transcription start site selection at RNA pol II promoter; plays role in alternative promoter choices, polyadenylation site selection & overall transcript levels; regulation of gene expression (Bradley et al., 2015). |
| Pep (CG6143) | **DBTR2; POLIITR; RNABPR** | **Zinc finger protein on ecdysone puffs**, may play a role in the process of early and late gene activation, or possibly in RNA processing, for a defined set of developmentally regulated loci; DNA binding; single-stranded RNA binding (Hamann and Stratling, 1998); single-stranded DNA binding (Amero et al., 1993); Cip1-Interacting Zinc Finger Protein (Zinc Finger C2H2-type: most common DNA-binding motifs found in eukaryotic transcription factors). |
| Saf-B | **DBTR2; POLIITR; RNABPR; RGEXPPT** | **Scaffold attachment factor B**, regulation of mRNA processing; regulation of transcription by RNA pol II; sequence-specific DNA binding (GO Reference Genome Project, 2011-); RNA binding (InterPro Project Members, 2004-); RNA Recognition Motif Domain. |
| ssx | **RGEXPPT; RNABPR; RNABPT** | **(sister-of-Sex-lethal)**, RNA binding activity; involved in post-transcriptional gene silencing; regulation of RNA splicing (Moschall et al., 2019). |
| Lrpprc2 | **RNABTN; MTTR; TN** | **Leucine-rich pentatricopeptide repeat containing 2**, regulation of mitochondrial transcription; coordination of mitochondrial translation (Baggio et al., 2014); RNA-binding protein; mRNA 3'-UTR binding (GO Reference Genome Project, 2011-); regulation of opsin-mediated signaling pathway (Jaiswal et al., 2015). |
| Hsc70-4 | **RGEXPPT; PSILEN** | **Heat shock protein 70 cognate 4**, regulatory ncRNA-mediated post-transcriptional gene silencing (Dorner et al., 2006). |
| Hyls1 | **U** | **Hyls1** centriolar and ciliogenesis, involved in cilium assembly; located in centriole & ciliary basal body. |
| Mcm5 | **RRR; DBRR; CHRM** | **Minichromosome maintenance 5**, involved in mitotic DNA replication; required to resolve meiotic double-strand breaks into crossovers; 3'-5' helicase activity (Moyer et al., 2006); DNA binding; DNA replication origin binding; DNA duplex unwinding (InterPro Project Members, 2004-); chromosome condensation/organization (Christensen and Tye, 2003); DNA endoreplication (Park and Asano, 2008). |

**Table B. Subdivisions of regulatory factors associated with the *Myc* target P1/P2 (plotted in Figure 12)**

| Gene name | Transcriptional Category | Activity/Function |
|---|---|---|
| Tudor-SN | **RNABTN; TN; RGEXPPT** | **Tudor staphylococcal nuclease**, shows activity towards both DNA and RNA substrates; translation regulation through its association with the RNA-induced silencing complex (RISC) (Ku et al., 2016; Caudy et al., 2003); regulatory ncRNA-mediated post-transcriptional gene silencing (InterPro Project Members, 2004-); RNA binding (GO Reference Genome Project, 2011-). |
| RnrL | **RRR** | **Ribonucleoside diphosphate reductase large subunit**, preparation of precursors necessary for DNA synthesis; catalysis of the biosynthesis of deoxyribonucleotides from the corresponding ribonucleotides (Gene Ontology Curators, 2002-); DNA replication (InterPro Project Members, 2004-). |
| RPA2 | **DBRR; RRR** | **Replication protein A2**, nuclear protein, DNA binding activity, involved in DNA recombination; DNA repair; and DNA replication (InterPro Project Members, 2004-). |
| MED4 | **COACT; MS; POLIITR; TPI** | **Mediator complex subunit 4**, component of the Mediator complex transcription coactivator activity (Gu et al., 2002); transcription coregulator activity (Park et al., 2001); RNA pol II transcription; serves as a scaffold for the assembly of a functional preinitiation complex with RNA pol II and the general transcription factors (Gu et al., 2002; Park et al., 2001); Mediator Complex. |
| Hcf | **COACT; CBTR2; CRTR2; POLIITR** | **Host cell factor**, chromatin binding (Guelman et al., 2006); chromatin remodeling (Suganuma et al., 2008); transcription coactivator activity (Mahajan et al., 2003); positive regulation of DNA-templated transcription (Furrer et al., 2010); ATAC Complex, COMPASS Complex, TRX Complex, SWRI Complex. |
| Nasp (CG8223) | **CRR; RRR** | **Nuclear autoantigenic sperm protein**, histone chaperone activity (Tirgar et al., 2023); histone binding; CENP-A containing chromatin assembly; DNA replication-dependent chromatin assembly (GO Reference Genome Project, 2011-); Belongs to the NASP family. |
| MED11 | **COACT; MS; POLIITR; TPI** | **Mediator complex subunit 11**, component of the Mediator complex, a coactivator involved in the regulated transcription of nearly all RNA pol II-dependent genes; regulation of transcription by RNA pol II (InterPro Project Members, 2004-); Mediator Complex. |

**Table B. Subdivisions of regulatory factors associated with the *Myc* target P1/P2 (plotted in Figure 12)**

| Gene name | Transcriptional Category | Activity/Function |
|---|---|---|
| MED17 | **COACT; MS; POLIITR; TPI** | **Mediator complex subunit 17**, component of the Mediator complex transcription coactivator activity (Gu et al., 2002); RNA pol II transcription; serves as a scaffold for the assembly of a functional preinitiation complex with RNA pol II and the general transcription factors; regulation of transcription by RNA pol II (Gu et al., 2002; Park et al., 2001); DNA-binding transcription factor binding (*Park et al., 2003*); Mediator Complex. |
| MED24 | **COACT; MS; POLIITR; TPI** | **Mediator complex subunit 24**, component of the Mediator complex transcription coactivator activity (Gu et al., 2002); RNA Pol II transcription; serves as a scaffold for the assembly of a functional preinitiation complex with RNA pol II and the general transcription factors; regulation of transcription by RNA pol II (Gu et al., 2002; Park et al., 2001); DNA-binding transcription factor binding (Park et al., 2001); transcription coregulator activity (Gu et al., 2002; Boube et al., 2000); Mediator Complex. |
| skd | **COACT; MS; POLIITR; TI** | **Skuld/MED13**, a subunit of the kinase module of the mediator complex; not required for all the transcriptional functions of mediator; links Mediator to a set of transcription factors involved in developmental signaling; transcription coregulator activity (Janody et al., 2003; Boube et al., 2000); regulation of transcription by RNA pol II (Janody et al., 2003); transcription initiation at RNA pol II promoter (Boube et al., 2000); Mediator Complex. |
| Parg | **HF; RRR** | **Poly(ADP-ribose) glycohydrolase**, DNA damage response (Fontana et al., 2023); heterochromatin formation (Tulin et al., 2006). |
| E(bx) | **POLIITR; DBTR2; CRTR2; TF** | **Enhancer of bithorax**, lysine-acetylated histone binding; methylated histone binding (Kwon et al., 2009); RNA pol II *cis*-regulatory region sequence-specific DNA binding (GO Reference Genome Project, 2011-); chromatin remodeling (Bai et al., 2007; Badenhorst et al., 2002; Mizuguchi et al., 2001; Xiao et al., 2001); negative regulation of DNA-templated transcription (Yao et al., 2018); nucleosome organization (Hamiche et al., 1999); regulation of DNA-templated transcription (Badenhorst et al., 2002); Nucleosome Remodeling Complex; Unclassified DNA Binding Domain Transcription Factors. |

**Table B. Subdivisions of regulatory factors associated with the *Myc* target P1/P2 (plotted in Figure 12)**

| Gene name | Transcriptional Category | Activity/Function |
|---|---|---|
| Iswi | **CRTR2; CHRM; CBTR2; TF; DBTR2; POLIITR; DBTFB** | **Imitation SWI**, energy-transducing component of the chromatin-remodeling complexes NURF (nucleosome-remodeling factor), ACF (ATP-utilizing chromatin assembly and remodeling factor), & CHRAC (chromatin accessibility complex (Corona et al., 2007; Eberharter et al., 2001); DNA-binding transcription factor binding (Vanolst et al., 2005); nucleosome array spacer activity (Ito et al., 1997; Varga-Weisz et al., 1997); nucleosome binding (InterPro Project Members, 2004-); chromatin organization (Corona et al., 2007; Mizuguchi et al., 1997); regulation of pol II transcription (Li et al., 2010); sperm DNA condensation (Doyen et al., 2015). |
| rept | **DBTR1; DBTR2; CBRR; DNAHRR; DNAHTR1; DNAHTR2; RRR; CRTR2; HF; RGEXPPT; POLITR; COACT; POLIITR; HAT** | **RuvB-like helicase**, DNA repair-dependent chromatin remodeling; NuA4 histone acetyltransferase complex (Kusch et al., 2004); negative regulation of gene expression (Diop et al., 2008); heterochromatin formation (Qi et al., 2006); positive regulation of transcription of nucleolar large rRNA by RNA pol I (Vinayagam et al., 2016); regulation of transcription by RNA pol II (GO Reference Genome Project, 2011-); core component of chromatin remodeling Ino80 complex; chromatin remodeling (Klymenko et al., 2006); transcriptional coactivator in Wg pathway by altered arm signaling; Pontin/Reptin antagonistically interfere with the nuclear Arm signaling; an essential cofactor for the normal function of Myc during cell growth & proliferation (Bellosta et al., 2005); ecdysone-mediated salivary gland cell autophagy cell death (Ihry and Bashirullah, 2014); negative regulation of Wnt signaling (Bauer et al., 2000); TIP60 Complex; SWR1 Complex. |
| Nup107 | **CBRR; RRR** | **Nucleoporin 107kD**, nuclear pore complex (NPC) assembly & maintenance; chromatin binding; (Gozalo et al., 2020); double strand break repair via homologous recombination (Ryu et al., 2015); post-transcriptional tethering of RNA pol II gene DNA to nuclear periphery (GO Reference Genome Project, 2011-). |
| Art4 | **HMOD; CRTR2; POLIITR** | **Arginine methyltransferase 4**, methylation of histone H3 at 'Arg-17' (Gene Ontology Curators, 2002-); histone methyltransferase activity (Boulanger et al., 2004; Cakouros et al., 2004); protein arginine omega N-asymmetric methyltransferase activity (Boulanger et al., 2004) activation of transcription by RNA pol II via chromatin remodeling (Cakouros et al., 2008). |

**Table B. Subdivisions of regulatory factors associated with the *Myc* target P1/P2 (plotted in Figure 12)**

| Gene name | Transcriptional Category | Activity/Function |
|---|---|---|
| Spt20 | **POLIITR; HAT; COREG** | **Spt20, isoform C,** Spt-Ada-Gcn5-acetyltransferase (SAGA) complex; involved in histone acetylation (Gene Ontology Curators, 2002-); regulation of transcription by RNA pol II (GO Reference Genome Project, 2011-); transcription coregulator activity (InterPro Project Members, 2004-); SAF6 (SAGA factor-like TAF6), a histone fold domain-containing protein can replace Taf6 in *Drosophila* SAGA complex & is required for SAGA-dependent gene expression; the gene CG17689 encodes an ortholog of Spt20/p38IP, CG9866 encodes a potential ortholog of Sgf73 & CG3883 encodes a novel histone fold domain (HFD)-containing protein SAF6 (Weake et al., 2009); SAGA (Spt-Ada-Gcn5 Acetyltransferase) Complex; Transcription Factor Spt20. |
| pont | **DBRR; DBTR2; TF; RRR; CRTR2; CBRR; POLIITR; DNAHRR; DNAHTR2; RGEXPPT** | **Pontin**, involved in transcriptional regulation by RNA pol II (GO Reference Genome Project, 2011-), DNA repair-dependent chromatin remodeling (Kusch et al., 2004); ribonucleoprotein assembly; transcriptional coregulator activity (Bellosta et al., 2005; Bauer et al., 2000); DNA helicase activity (GO Reference Genome Project, 2011-); positive regulation of gene expression (Diop et al., 2008). |
| Spt6 | **CBTR2; CRTR2; POLIITR; DBTR2; TE** | **Spt6**, transcription elongation factor (other elongation factor Spt5, Paf1); binds to histone H3 & enhances transcription elongation by RNA pol II (RNAPII) (InterPro Project Members, 2004-); chromatin binding (Petruk et al., 2006); RNA pol II complex binding (Saunders et al., 2003); DNA binding (InterPro Project Members, 2004-); positive regulation of transcription by RNA pol II (Ardehali et al., 2009; Petruk et al., 2006). |
| Mtor (Mgtor) | **CBTR2; CRTR2; POLIITR** | **Megator**, transcriptional attenuator of X chromosome gene expression in male X chromosome dosage compensation; histone acetyltransferase binding; negative regulation of transcription by RNA pol II (Aleman et al., 2021); chromatin DNA binding; chromatin organization; chromatin remodeling; positive regulation of transcription by RNA pol II (Vaquerizas et al., 2010); Nuclear Pore Complex. |
| spag | **POLIITR** | **Spaghetti**, homolog of RNA pol II-associated protein 3-like C-terminal (RPAP3, contains TPR-repeats towards the N terminus); negative regulation of motor neuron apoptotic process (Means et al., 2015); plays role in the assembly of RNA pol II complex (Benbahouche et al., 2014). |

**Table B. Subdivisions of regulatory factors associated with the *Myc* target P1/P2 (plotted in Figure 12)**

| Gene name | Transcriptional Category | Activity/Function |
|---|---|---|
| HDAC6 | **HDAC; RGEXPPT; CRTR2; POLIITR** | **Histone Deacetylase 6, isoform G**, cytosolic deacetylase (Miskiewicz et al., 2014), key modulator of proteostasis by mediating ubiquitin-proteasomal & lysosomal degradation of native and/or misfolded proteins; histone deacetylase activity (Feller et al., 2015; Cho et al., 2005; Barlow et al., 2001); regulation of DNA-templated transcription (Cho et al., 2005); negative regulation of transcription by RNA pol II (GO Reference Genome Project, 2011-) chromatin organization (InterPro Project Members, 2004-); RPD3/HDA1 Lysine Deacetylases. |
| D1 | **TF; DBTR2; REP** | **D1 chromosomal protein**, a multi-AT-hook chromosomal protein, associates with AT-rich satellites, including the SAT-III repeats of the X-chromosome (Blattes et al., 2006); binds to the minor-groove of the DNA (Levinger and Varshavsky, 1982); favors heterochromatin-mediated gene repression involving its interaction with topoisomerase II; Unclassified DNA Binding Domain Transcription Factors. |
| Rae1 | **RNABPT; RGEXPPT; TREX; RRR** | **Rae1**, a nucleoporin member of the WD40-repeat β propeller protein super family with pleiotropic functions including poly(A)+ mRNA export, cell cycle regulation, male meiosis control & male germ cell post-meiotic differentiation; RNA binding; transcription-dependent tethering of RNA pol II gene DNA at nuclear periphery (GO Reference Genome Project, 2011-); male meiotic I (Volpi et al., 2013); positive regulation of gene expression (Tian et al., 2011); Nuclear Pore Complex. |
| Map60 (CP-60) | **POLIITR; DBTR2** | **Microtubule-associated protein**, microtubule binding; localized to the centrosome in a cell cycle-dependent manner (Kellogg et al., 1995); regulation of transcription by RNA pol II (GO Reference Genome Project, 2011-); a component of a Centrosomal complex; MADF Domain. |
| Lon | **DBMTTR; DBMTRR; RGEXPRR** | **Lon protease**, a conserved nuclear ATP-stimulated serine protease; ssDNA binding (GO Reference Genome Project, 2011-), targeted to the mitochondrial matrix for protein turnover; serine hydrolase activity (Kumar et al., 2021); regulation of mitochondrial DNA copy number & transcription by degradation of mitochondrial TFAM (Matsushima et al., 2010). |
| Nlp | **CBTR2; CRTR2; CHRM; RNABTR2** | **Nucleoplasmin**, chromatin binding; RNA binding (GO Reference Genome Project, 2011-); chromatin remodeling; sperm DNA condensation (Emelyanov et al., 2014). |

**Table B. Subdivisions of regulatory factors associated with the *Myc* target P1/P2 (plotted in Figure 12)**

| Gene name | Transcriptional Category | Activity/Function |
|---|---|---|
| Nup98-96 | **CBTR2; RNABTR2; TI; POLIITR; TMM** | **Nucleoporin 98-96kD**, precursor of Nup98 and Nup96 proteins, two integral parts of the nuclear pore; male germ line cells differentiation; Chromatin DNA binding (Ilyin et al., 2017; Pascual-Garcia et al., 2017; Kalverda et al., 2010); promoter-enhancer loop anchoring (Pascual-Garcia et al., 2017); promoter-specific chromatin binding (Pascual-Garcia et al., 2017; Pascual-Garcia et al., 2014); RNA binding (GO Reference Genome Project, 2011-); heat shock-mediated polytene chromosome puffing (Kalverda et al., 2010); positive regulation of transcription by RNA pol II (Panda et al., 2014; Pascual-Garcia et al., 2014; Capelson et al., 2010; Kalverda et al., 2010); positive regulation of transcription initiation by RNA pol II (Capelson et al., 2010); post-transcriptional tethering of RNA pol II gene DNA to nuclear periphery; telomere tethering to nuclear periphery (GO Reference Genome Project, 2011-); Nuclear Pore Complex. |
| tex | **TREX** | (**tex**), part of THO complex part of transcription export complex; mRNA export from nucleus (GO Reference Genome Project, 2011-; InterPro Project Members, 2004-); Transcription Export Complex. |
| Nup93-1 | **CBPT; RGEXPPT** | **Nucleoporin 93kD-1**, chromatin binding; negative regulation of gene expression, epigenetic; chromatin DNA binding (Gozalo et al., 2020). |
| sqd | **RNABTN; TN; RGEXPPT** | **Squid, isoform E,** Dpp signaling, squid & Hephaestus (but not Hrb27C) are necessary for proper bone morpho-genetic protein (BMP) signaling in GSCs; novel roles for RNA binding proteins Squid, Hephaestus, & Hrb27C during *Drosophila* oogenesis (Finger et al., 2022); negative regulation of RNA splicing (Ji and Tulin, 2009); negative regulation of translation (Cooperstock and Lipshitz, 1997); regulation of gene expression (GO Reference Genome Project, 2011-). |
| CG30291 | **POLIITR** | Probable regulator of cell proliferation. May regulate CDK5, NF κB-mediated gene transcription & p53/TP53 activation; regulation of mitotic cell cycle; regulation of cyclin-dependent protein Ser/Thr kinase activity (GO Reference Genome Project, 2011-); CDK5 regulatory subunit-associated protein 3. |
| CG13850 | **RGEXPMT; MTRNABPR** | RNA binding; mitochondrial RNA processing; regulation of mitochondrial mRNA stability (GO Reference Genome Project, 2011-). |

**Table B. Subdivisions of regulatory factors associated with the *Myc* target P1/P2 (plotted in Figure 12)**

| Gene name | Transcriptional Category | Activity/Function |
|---|---|---|
| l(2)10685 | **RNABPT; RGEXPPT** | **Lethal (2) 10685**, methyltransferase activity (GO Reference Genome Project, 2011-); S-adenosylmethionine-dependent methyltransferase activity (InterPro Project Members, 2004-); orthologous to human NSUN4 (NOP2/Sun RNA methyltransferase 4); Unclassified Methyltransferases. |
| lig | **RNABPT; RGEXPPT** | **Lingerer**, RNA binding protein; regulation of gene expression (Baumgartner et al., 2013). UBA-like Superfamily. |
| ncd | **CHRM** | **non-claret disjunctional**, a minus-end-directed kinesin microtubule motor protein and the sole member of the kinesin-14 motor family; chromosome segregation (Hallen et al., 2008); distributive segregation (Whyte et al., 1993); mRNA transport (Fahmy et al., 2014). |
| Lam (CG6944) | **CBRR; HF** | **Lamin**, chromatin binding (Verboon et al., 2015); Heterochromatin formation (Verboon et al., 2015; Dialynas et al., 2010; Shevelyov et al., 2009); regulation of meiosis I cytokinesis (Hayashi et al., 2016); Lamins. |
| tral | **RNAHPT; RNABPT; RGEXPPT** | **Trailer hitch, isoform D**, DEAD/H-Box RNA helicase binding (Barbee et al., 2006; Wilhelm et al., 2005); mRNA binding (GO Reference Genome Project, 2011-); RNA binding (InterPro Project Members, 2004-); piRNA pathway; retrotransposon silencing (Liu et al., 2011); required for oocyte dorsoventral patterning via actin and microtubule cytoskeleton organization. |
| abs | **RNAHPR; RNABPR; RGEXPPR** | **Abstrakt**, DEAD/DEAH-Box RNA helicase (Lasko, 2000), RNA binding (GO Reference Genome Project, 2011-); RNA splicing, via spliceosome (Herold et al., 2009); regulates cell polarity in oocytes & embryos; downregulation of Notch signaling in asymmetric cell division in ganglion mother cell (GMC2-4a) in collaboration with Inscuteable (*Insc*) (Irion et al., 2004). |
| Nup50 | **CBTR2; POLIITR** | **Nuclear pore complex protein Nup50 (Nucleoporin 50kD)**, chromatin DNA binding; positive regulation of transcription by RNA pol II (Kalverda et al., 2010); an important protein for TGF-beta signal transduction by mediating the nuclear translocation of the Mothers against dpp protein (*Mad*); Association of 400 genes interacting with Nup50 Nucleoporin, among which transcriptionally active genes inside the nucleoplasm are predominantly involved in development & cell cycle regulation (Kalverda et al., 2010); Nuclear Pore Complex. |

**Table B. Subdivisions of regulatory factors associated with the *Myc* target P1/P2 (plotted in Figure 12)**

| Gene name | Transcriptional Category | Activity/Function |
|---|---|---|
| CG8064 | **RNABPT; RGEXPPT** | SnoRNA binding; involved in maturation of SSU-rRNA (GO Reference Genome Project, 2011-); ); human ortholog(s) implicated in papillary thyroid carcinoma; orthologous to human WDR3 (WD repeat domain 3). |
| Patr-1 | **RNABPR; RGEXPPR** | **Protein associated with topo II related – 1**, a P body component involved in deadenylation-dependent decapping of nuclear transcribed-mRNA (Nishihara et al., 2013; Braun et al., 2010) & regulation of synaptic growth at neuromuscular junctions (Pradhan et al., 2012); RNA binding (GO Reference Genome Project, 2011-); positive regulation of mRNA catabolic process (Nish ihara et al., 2013). |
| AGO2 | **RNABPT; RGEXPPT; PSILEN** | **Argonaut 2**, interaction with small interfering RNAs (siRNAs) to form RNA-induced silencing complexes (RISCs), siRNA binding (Goh and Okamura, 2019; Kawamura et al., 2008; Tomari et al., 2007; Rand et al., 2005; Lingel et al., 2003); miRNA-mediated gene silencing (Besnard-Guérin et al., 2015); single-stranded RNA binding (Goh and Okamura, 2019); siRNA binding (Goh and Okamura, 2019; Kawamura et al., 2008; Tomari et al., 2007; Rand et al., 2005; Lingel et al., 2003). |
| Edc3 | **RNABPR; RGEXPPT** | **Enhancer of decapping 3**, during mRNA degradation may play a role in mRNA decapping; mRNA binding; deadenylation-independent decapping of nuclear-transcribed mRNAs (GO Reference Genome Project, 2011-); RNA binding (InterPro Project Members, 2004-); mRNA catabolic process (Tritschler et al., 2007). |
| CG44270 | **U** | Expressed in several structures, including anterior ectoderm; ectoderm anlage; embryonic central brain neurons; extended germ band embryo; and somatic precursor cell; Domain of Unknown Function; DUF4799. |
| bsf | **RNABPT; PAFAC; RGEXPPT; MTTR** | **(bicoid stability factor)**, a member of the family of proteins containing the pentatricopeptide motif, an RNA binding domain; mitochondrial mRNA stability (Pajak et al., 2019; Jaiswal et al., 2015); post-transcriptional control of gene expression in mitochondria, where it has multiple roles in gene expression; required for progression through oogenesis and viability; mRNA 3'-UTR binding (Mancebo et al., 2001); mitochondrial mRNA polyadenylation (Bratic et al., 2011); regulation of mitochondrial gene expression (Matsushima et al., 2017); regulation of mitochondrial transcription (Bratic et al., 2011). |

**Table B. Subdivisions of regulatory factors associated with the *Myc* target P1/P2 (plotted in Figure 12)**

| Gene name | Transcriptional Category | Activity/Function |
|---|---|---|
| CG44270 | **U** | Expressed in several structures, including anterior ectoderm; ectoderm anlage; embryonic central brain neurons; extended germ band embryo; and somatic precursor cell; Domain of Unknown Function; DUF4799. |
| CG43813 | **U** | Uncharacterized; Unknown. |
| CG8187 | **U** | Uncharacterized; Putative Adhesion Molecule |
| CG7406 | **U** | Uncharacterized; Domain of Unknown Function; DUF4766 |
| CG7173 | **U** | expressed in embryonic dorsal epidermis; embryonic head epidermis; embryonic ventral epidermis; and spermatozoon; contains KAZAL Domain, KAZAL Domain Superfamily. |
| CG15731 | **U** | Expressed in several structures, including anterior endoderm anlage; embryonic epidermis; and germ layer; Domain of Unknown Function; DUF4766 |
| Nin (Bsg25D) | **RRR** | **Ninein**, a microtubule-anchoring protein in humans; microtubule binding (Kowanda et al., 2016); microtubule anchoring at centrosome (GO Reference Genome Project, 2011-); microtubule organizing center attachment site organization (Zheng et al., 2016). |
| CG15239 | **U** | expressed in embryonic dorsal epidermis; embryonic esophagus; embryonic head epidermis; embryonic ventral epidermis; and embryonic/larval salivary gland; Domain of Unknown Function; DUF4773. |
| CG8003 | **RGEXPPT** | Expressed in adult head and organism; orthologous to human ANKMY2 (ankyrin repeat and MYND domain containing 2). |
| Ars2 | **RGEXPPT; PSILEN; POLIITR** | **Arsenic resistance protein 2**, binds the cap binding complex; plays roles in small RNA biogenesis (Garcia et al., 2016); primary miRNA processing to pre-miRNA required for siRNA-mediated silencing; regulation of transcription by RNA pol II (Speth et al., 2018); Serrate/Ars2 heterodimers. |
| pAbp | **PAB; RNABTN; TN; RGEXPTN** | **Poly(A) binding protein**, Poly(A) binding (Mount and Salz, 2000); post-transcriptional regulation of gene expression; mRNA binding (Lasko, 2000); mRNA translation (Herold et al., 2009). |

**Table B. Subdivisions of regulatory factors associated with the *Myc* target P1/P2 (plotted in Figure 12)**

| Gene name | Transcriptional Category | Activity/Function |
|---|---|---|
| scny | **COREP; HF; POLIITR** | **Scrawny**, ubiquitinyl hydrolase 1; hydrolase & deubiquitination activity deubiquitinating Imd and prevention of constitutive activation of Imd/NF-κB cascade (Engel et al., 2014; Buszczak et al 2009; Thevenon et al 2009); functions as a transcriptional repressor by continually deubiquitinating histone H2B at the promoters of genes critical for cellular differentiation, thereby preventing histone H3 'Lys-4' trimethylation (H3K4me3); heterochromatin formation (Buszczak et al., 2009); Ubiquitin Specific Protease (USP) Deubiquitinases. |
| lost | **RNABPR; RGEXPPR** | **lost**, interaction with the RNA-binding protein Rumpelstiltskin (*rump*) for posterior localization of mRNAs by diffusion/entrapment during late stages of oogenesis; mRNA splicing, via spliceosome (Herold et al., 2009); localization to various RNP complexes with a broad role in RNA metabolism. |
| Srp54 | **RNABPR; RGEXPPR; DBTR2; POLIITR** | **Splicing regulatory protein 54**, regulation of mRNA alternative splicing; mRNA binding; regulation of gene expression; regulation of the transcriptional start site selection at RNA pol II promoter (Bradley et al., 2015); poly-pyrimidine tract binding (Kennedy et al., 1998); RNA binding (InterPro Project Members, 2004-). |
| Smn | **RNABPR; CRR; RGEXPPR** | **Survival motor neuron**, eponymous member of the SMN complex, functions as an assembly chaperone for Sm-class small nuclear ribonucleoproteins; RNA binding; mRNA processing (InterPro Project Members, 2004-); chromatin organization (Lee et al., 2009); protein-RNA-complex assembly (Shpargel et al., 2009); stem cell differentiation, division, and proliferation (Grice and Liu, 2011); Survival Motor Neuron Complex-GEM4A Variant. |
| Larp4B | **PSILEN; RNABPT; TN; RGEXPPT** | **La-related protein Larp4B**, RNA-binding protein; mRNA 3'-UTR binding (Gene Ontology Curators, 2002-); negative regulation of cell growth and translation; post-transcriptional inhibition of MYC protein (Funakoshi et al., 2018). |
| SF2 | **DBTR2; RNABPR; RGEXPPR** | **Splicing factor 2**, DNA binding (Lynch and Maniatis, 1996); mRNA binding; regulation of gene expression; regulation of transcriptional start site selection at RNA pol II promoter (Bradley et al., 2015; Lasko, 2000); RNA binding (GO Reference Genome Project, 2011-); Canonical Ser/Arg Rich Splice Factors. |

**Table B. Subdivisions of regulatory factors associated with the *Myc* target P1/P2 (plotted in Figure 12)**

| Gene name | Transcriptional Category | Activity/Function |
|---|---|---|
| Hrb98DE | **RNABTN; TN; PSILEN; RGEXPPT** | **Heterogeneous nuclear ribonucleoprotein at 98DE**, nuclear RNA-binding protein; hnRNA stability, splicing, IRES-dependent translation, & translational repression; one of the main targets of the poly(ADP-ribosyl)ation pathway; tissue polarity patterning & germ-line stem cell fate; negative regulation of RNA splicing; mRNA 5'-UTR binding (Ji et al., 2009); RNA binding (InterPro Project Members, 2004-); mRNA binding (Lasko, 2000); post-translational modification of hnRNPs, such as poly(ADP-ribosyl)ation, regulation of gene expression during development, such as eye patterning (Ji et al., 2009); Spliceosome Comple A. |
| Ge-1 | **PSILEN; RNABPR; RGEXPPT** | **Ge-1**, RNAi pathway (miRNA-mediated gene silencing) (Eulalio et al., 2009); signaling pathways that activate NF-κB , Toll & Imd pathways (Jin et al., 2008; De Gregorio et al., 2002); mRNA degradation, plays role in mRNA decapping (GO Reference Genome Project, 2011-; Eulalio et al., 2009); Enhancer of mRNA-Decapping Protein 4 Homolog. |
| HnRNP-K | **DBTR2; DBTFB; POLIITR; CRTR2; RNABTN; TN; RGEXPTN** | **Heterogeneous nuclear ribonucleoprotein K**, localizes in nucleus, cytoplasm & mitochondria; involved in gene regulation; post-transcriptional RNA processing; RNA transport; mRNA binding; regulation of transcription by RNA pol II (GO Reference Genome Project, 2011-); RNA binding (InterPro Project Members, 2004-); transcription factor binding; DNA binding; involved in chromatin remodeling, transcription, splicing & translation; docking platform for the integration of signaling cascades (Bomsztyk et al., 2004). |
| NfI | **TF; DBTR2; POLIITR** | **Nuclear factor I, isoform B**, **CCAAT box-binding transcription factor** (CTF) (Mermod et al., 1989); regulation of transcription by RNA pol II; PNA pol II *cis*-regulatory region sequence-specific DNA binding (GO Reference Genome Project, 2011-); regulation of DNA-templated transcription (InterPro Project Members, 2004-); MAD Homology Domain Transcription Factors. |
| LKRSDH | **COREP; POLIITR** | **Lysine ketoglutarate reductase/saccharopine dehydrogenase**, a bifunctional enzyme in the lysine degradation pathway, and transcriptional corepressor in ecdysone signaling; histone binding; nuclear receptor binding; negative regulation of transcription by RNA pol II (Cakouros et al., 2008); CH-NH Oxidoreductases, NAD Or NADP As Acceptor. |

**Table B. Subdivisions of regulatory factors associated with the *Myc* target P1/P2 (plotted in Figure 12)**

| Gene name | Transcriptional Category | Activity/Function |
|---|---|---|
| Alsin2 (CG7564) | **RNABPR** | **Alsin2**, mRNA binding; mRNA splice site recognition (InterPro Project Members, 2004-); mRNA splicing, via spliceosome; precatalytic spliceosome complex (Herold et al., 2009; Mount and Salz, 2000). |
| wapl | **CRR; ACT; REP; CHRM; CHRSILEN** | **wings apart-like**, (*wapl*) & precocious dissociation of sisters 5 (*pds5*) proteins form the releasin complex for separation of sister chromatid at mitosis by removing the cohesin ring complex from chromosomes; gene activation & silencing via interaction with cohesin (Gause et al., 2010; Dorsett et al., 2005); chromatin organization via sister chromatid cohesin (Verni et al., 2000); regulation of Wnt/Wingless signaling pathway to determine the wing size in the Mediterranean fruit fly (medfly) (Cho et al., 2013). |
| tex | **TREX** | **tex**, involved in mRNA export from nucleus; part of THO complex part of transcription export complex (GO Reference Genome Project, 2011-; InterPro Project Members, 2004-); Transcription Export Complex. |
| net (CG11450) | **DBTR2; TF; REP; POLIITR** | **Net**, Basic Helix-Loop-Helix protein, acts as a transcriptional repressor; during wing vein formation it is expressed in all interveins territories; negative regulation of transcription by RNA pol II (Brentrup et al., 2000); antagonist of the product of EGFR; DNA binding transcription factor activity; regulation of transcription by RNA pol II (Peyrefitte et al., 2001); E-Box binding; positive regulation of transcription by RNA pol II (GO Reference Genome Project, 2011-). |
| mri | **POLIITR** | **mrityu**, a BTB/POZ domain containing protein (Many BTB proteins are transcriptional regulators) (Zollman et al., 1994); positive regulation of phosphorylation (GO Reference Genome Project, 2011-); involved in retinal apoptosis via its regulation by transcription factor Klumpfuss (Rusconi and Challa); Pseudo-Glycerol Kinase. |
| CG2962 | **POLIITR; COACT** | Transcription coactivator activity; positive regulation of transcription by RNA pol II (GO Reference Genome Project, 2011-). |
| nonA | **RNABPR; POLIITR** | **no on or off transient A**, mRNA binding (Greenspan and Ferveur, 2000; Lasko, 2000); RNA binding; regulation of DNA-templated transcription (GO Reference Genome Project, 2011-); mRNA splicing, via spliceosome (Herold et al., 2009); RNA Recognition Motif Domain. |

**Table B. Subdivisions of regulatory factors associated with the *Myc* target P1/P2 (plotted in Figure 12)**

| Gene name | Transcriptional Category | Activity/Function |
|---|---|---|
| rump | **RNABPR; RGEXPPR** | **Rumpelstiltskin**, the *Drosophila* hnRNP M homolog, binds to (*nanos*) & (*oskar*) mRNAs & localizes them to the germ plasm by diffusion/entrapment during late stages of oogenesis; splicing factor, binds to exonic splicing enhancers on pre-mRNA targets; mRNA 3'-UTR binding (Sinsimer et al., 2011; Jain and Gavis, 2008); mRNA binding (Lasko, 2000); RNA binding (InterPro Project Members, 2004-); mitotic cell cycle (Ducat et al., 2008). |
| rin (CG9412) | **RNABPR; RNAHPR; RGEXPPR** | **Rasputin**, evolutionarily conserved RNA-binding protein, RNA helicase activity (Gene Ontology Curators, 2002-); functions as a link between Ras signaling & RNA metabolism (Costa et al., 2013; Pazman et al., 2000); Rho-mediated signaling (Pazman et al., 2000); mRNA binding (GO Reference Genome Project, 2011-); positive regulation of gene expression (Costa et al., 2013); response to amino acid starvation (Aguilera-Gomez et al., 2017); catalytic step 2 spliceosome (Herold et al, 2009); Unclassified RNA Helicases. |
| Usp10 | **POLIITR; TE; RGEXPPT** | **Ubiquitin specific protease 10**, cysteine-type deubiquitinase activity; ubiquitin-dependent protein catabolic process (Gene Ontology Curators, 2002-); negative regulation of transcription elongation (GO Reference Genome Project, 2011-); positive regulation of Notch signaling (Zhang et al., 2012). |
| Mtr4 | **RNAHPR; RNABPR; RGEXPPR** | **Mtr4 helicase**, interferon immune signaling, viral defense response (Molleston et al., 2016); RNA binding; RNA catabolic process (InterPro Project Members, 2004-); RNA helicase activity (GO Reference Genome Project, 2011-); SKI2-Like RNA Helicases. |
| tsu | **RNABPR; RGEXPPR; POLIITR** | **Tsunagi**, RNA binding; mRNA binding; RNA processing (InterPro Project Members, 2004-); regulation of MAPK levels by the Exon Junction Complex (EJC) (Ashton-Beaucage & Therrien, 2010); disassembly of EJC by interaction between Mago/Tsunagi heterodimer with the EJC key regulator Pym in cytoplasm (Ghosh et al., 2014). |
| san | **CHRM; RRR** | **separation anxiety**, mitotic sister chromatid cohesin; couples the processes of cohesion and DNA replication (Ribeiro et al., 2016; Williams et al., 2003). |
| CG4849 | **RNABTN; TN; RGEXPPT** | U5 snRNA binding (GO Reference Genome Project, 2011-); translation elongation (Lasko, 2000); positive regulation of gene expression (Ashton-Beaucage et al., 2014). |

**Table B. Subdivisions of regulatory factors associated with the *Myc* target P1/P2 (plotted in Figure 12)**

| Gene name | Transcriptional Category | Activity/Function |
|---|---|---|
| eIF4A | **RNABTN; RNAHTN; TNFAC; TN; RGEXPTN** | **Eukaryotic translation initiation factor 4A**, an essential DEAD-Box RNA helicase protein (Lasko, 2000); RNA binding (InterPro Project Members, 2004-); a canonical translation initiation factor; a component of the eIF4F cap-binding complex, essential for cap-dependent translation of mRNA; SMAD binding; negative regulation of BMP signaling pathway (Li and Li, 2006); mitotic cell cycle (Ducat et al., 2008). |
| Cbp80 | **RNABPR; RGEXPPT; PSILEN** | **cap binding protein 80**, component of the cap-binding complex (CBC), which binds cotranscriptionally to the 5'-cap of pre-mRNAs and is involved in various processes such as pre-mRNA splicing and RNA-mediated gene silencing (RNAi); RNA binding; RNA cap binding; mRNA export from nucleus (InterPro Project Members, 2004-); mRNA binding (GO Reference Genome Project, 2011-); primary-miRNA processing; regulatory ncRNA-mediated post-transcriptional gene silencing; RNA-mediated gene silencing (RNAi); miRNA-mediated RNA interference via its interaction with Ars2, required for primary microRNAs (miRNAs) processing (Sabin et al., 2009). |
| LSm7 | **RNABPR; RGEXPPR** | **Like Sm 7**, RNA binding activity; nuclear-transcribed mRNA catabolic process (InterPro Project Members, 2004-). |
| Sf3a1 | **RNABPR; RGEXPPR** | **Splicing factor 3a subunit 1**, RNA binding (GO Reference Genome Project, 2011-); RNA processing (InterPro Project Members, 2004-). |
| thoc6 | **POLIITR; TE; TT; TREX** | **THO complex subunit 6**, linking transcription elongation to several cellular processes including mitotic recombination, co-transcriptional mRNP assembly, & nuclear mRNA export (Rehwinkel et al., 2004; Aguilera, 2002); Tho Complex; Transcription Export Complex. |
| CG9641 | **U** | Uncharacterized; Unknown; Domain of Unknown Function DUF4780. |
| Fkbp39 (FK506-bp1) | **DBTR2; POLIITR; CBTR2** | **FK506-bp1 binding protein**, FK506-bp1 binding; DNA binding, Juvenil Hormone Response Element binding (JHRE); chromatin binding; transcription (Theopold et al., 1995). |
| Usp7 | **HF; RRR** | **Ubiquitin-specific protease 7**, positive regulation of heterochromatin formation (van der Knaap et al., 2005); Nucleotide Excision Repair (NER) (Higa et al., 2018). |

**Table B. Subdivisions of regulatory factors associated with the *Myc* target P1/P2 (plotted in Figure 12)**

| Gene name | Transcriptional Category | Activity/Function |
|---|---|---|
| RpS3 | **DBRR; RRR; RNABPR** | **Ribosomal protein S3**, haploinsufficient - heterozygous mutants display the 'Minute' phenotype, characterized by a slower developmental rate and small adult bristles DNA binding; DNA repair; oxidized purine nuclease lesion DNA N-glycosylase activity (Yacoub et al., 1996); damaged DNA binding; oxidized purine DNA binding (Hegde et al., 2004); RNA binding (InterPro Project Members, 2004-); translation initiation complex formation; class I DNA-(apurinic or apyrimidinic site) endonuclease activity (Yacoub et al., 1996, Wilson et al., 1994). |
| Pdcd4 | **POLIITR** | **Programmed cell death 4**, negative regulation of DNA-templated transcription (InterPro Project Members, 2004-); Armadillo-type fold. |
| bic (l(2)49Da, l(2)k10712, Btf, E(Bic)) | **TF; POLIITR** | **Bicaudal**, β subunit of the nascent polypeptide-associated complex (NAC), a heterodimeric cytosolic protein complex (GO Reference Genome Project, 2011-); transcription regulation & mitochondrial translocation (Liu et al., 2010; Rospert et al., 2002); Transcription Factor BTF3. |
| Lst8 | **POLIITR** | **Lst8**, a conserved TOR-binding protein, CREB-regulated transcription coactivator (Crtc)-dependent regulation of cell growth based on genetic evidence (Ctrc, a highly conserved transcriptional coactivator of cAMP-response element-binding protein (CREB)); positive regulation of transcription by transcription factor localization (Kuo et al., 2015); TORC2 Complex. |
| Nup62 | **DBTR2; CBTR2; TREX; POLIITR** | **Nup62kD**, binds to transcriptionally active genes; chromatin DNA binding (Kalverda et al., 2010); structural constituent of nuclear pore (InterPro Project Members, 2004-); chromosome attachment to the nuclear pore (Breuer and Ohkura, 2015); RNA export from nucleus (GO Reference Genome Project, 2011-); Nuclear Pore Complex. |
| CG7518 | **U** | Uncharacterized; FAM 193 Family |
| dod | **DBTFB; POLIITR; RGEXPPT** | **Dodo**, peptidylprolyl isomerase; transcription factor binding; positive regulation of protein ubiquitination (Hsu et al., 2001); PARVULINS. |
| Rolled (rl) | **DBTFB; TF; POLIITR** | **Mitogen-activated protein kinase ERK A**, downstream effector of the RAS/MAPK pathway; mitotic cell cycle (Marenda et al., 2006); DNA-binding transcription factor binding (Astigarraga et al., 2007). |

**Table B. Subdivisions of regulatory factors associated with the *Myc* target P1/P2 (plotted in Figure 12)**

| Gene name | Transcriptional Category | Activity/Function |
|---|---|---|
| CG32486 | **DBTR2; POLIITR** | Nuclear zinc ion binding (InterPro Project Members, 2004-); DNA binding, transcription in response to metabolic gene expression (Ling et al., 2010); orthologous to human ZFTRAF1 (zinc finger TRAF-type containing 1); Zinc finger, RING/FYVE/PHD-type. |
| Nopp140 | **RNABTR1; POLITR** | **Nopp140**, nucleolar and Cajal body protein; rRNA 2'-O-methylation (He et al., 2015); interaction with RNA polymerase I (Tantos et al., 2013); Srp40; C-terminal. |
| DnaJ-1 | **POLIITR; DBTFB** | **DnaJ-like-1**, heat-shock protein co-factor, regulates & interacts with larger heat shock proteins providing client specificity; regulates the folding of proteins whose misfolding leads to age-dependent neurodegeneration; DNA binding transcription factor binding; protein binding (Miller et al., 2017). |
| Polr1B (Rpl135) | **DBTR1; POLITR; TI** | **RNA polymerase I subunit B**, DNA binding; DNA-directed 5'-3' RNA polymerase activity (InterPro Project Members, 2004-); RNA Pol I activity (GO Reference Genome Project, 2011-); RNA Polymerase I Complex. |
| Hpf1(CG1218) | **RRR** | **Histone PARylation factor 1**, zinc ion binding (Isogai et al., 2010); histone binding (GO Reference Genome Project, 2011-); DNA damage response (Fontana et al., 2023). |
| heph | **RNABTN; TN; RGEXPTN** | **Hephaestus**, nucleo-cytoplasmic shuttling protein, mRNA regulatory element binding translation repressor activity (Besse et al., 2009); regulates Oskar mRNA translation; spermatid individualization (Robida et al., 2010; Robida and Singh, 2003); regulation of Notch signaling (Dansereau et al., 2002); Dpp signaling, Squid & Hephaestus (but not Hrb27C) are necessary for proper bone morpho-genetic protein (BMP) signaling in GSCs; novel roles for RNA binding proteins Squid, Hephaestus, & Hrb27C in *Drosophila* oogenesis (Finger & Ables, 2022); RNA binding (InterPro Project Members, 2004-); mRNA binding (Lasko, 2000). |
| CG10077(BcDNA:HL07910) | **RNAHPR; RNABPR** | RNA helicase, RNA binding activity; alternative mRNA splicing, via spliceosome; part of ribonucleoprotein complex (GO Reference Genome Project, 2011-); RNA helicase activity DEAD-Box RNA Helicases (Lasko, 2000). |
| tho2 | **POLIITR; RNABTR2; TREX** | **(tho2), component of THO Complex**, nuclear export of mRNA (Rehwinkel et al., 2004); mRNA binding (GO Reference Genome Project, 2011-); Transcription Export Complex. |

**Table B. Subdivisions of regulatory factors associated with the *Myc* target P1/P2 (plotted in Figure 12)**

| Gene name | Transcriptional Category | Activity/Function |
|---|---|---|
| Prim2 | **DBRR; RRR; PRIM2** | **DNA primase large subunit, PRIM2**; synthesis of short RNA-DNA primers on the lagging strand during DNA synthesis (Kuroda et al., 1990; Cotterill et al., 1987; Kaguni et al., 1983); compound eye morphogenesis (Chen et al., 2000); alpha DNA polymerase: primase complex (Kuroda et al., 1990; Cotterill et al., 1987; Kaguni et al., 1983; Kaguni et al., 1983; Villani et al., 1980). |
| mEFG1 | **MTTNF; MTTN** | **Mitochondrial translation elongation factor G1**, developmental signal from mitochondria to nucleus for slow proliferation in the case of low mitochondrial energy level (Trivigno and Haerry, 2011); GTP binding; catalysis of mRNAs and tRNAs translocation along the ribosomes via GTP hydrolysis (InterPro Project Members, 2004-). |
| Hrb27C | **RNABTN; TN; DBTR2; RGEXPTN** | **Heterogeneous nuclear ribonucleoprotein at 27C**, component of ribonucleosomes; mRNA 3'-UTR binding (Nelson et al., 2007); ssDNA binding (Matunis et al., 1992); translation repressor activity (Szostak et al., 2018); regulation of border cell migration (Mathieu et al., 2007); the *Drosophila* PDGF/VEGF Receptor (PVR) pathway. |
| sti | **CHRSILEN; CRR** | **Sticky**, a member of the AGC family of kinases, functions to regulate both actin-myosin-mediated cytokinesis & epigenetic gene silencing (histone H3-K9 methylation, HP1 localization, & heterochromatin-mediated gene silencing); chromatin organization (Sweeney et al., 2008). |
| SmF | **RNABPR** | **Small ribonucleoprotein particle protein SmF**, RNA binding (GO Reference Genome Project, 2011-). |
| Klp10A | **RRR** | **Kinesin-like protein at 10A** (current: **Plus-end-directed kinesin ATPase**), establishment of mitotic spindle asymmetry; centriole assembly; non-motile cilium assembly (Gottardo et al., 2016); centrosome duplication (*Pavlova et al., 2019*); establishment of meiotic spindle orientation; spindle assembly involved in female meiotic I (Zou et al., 2008); meiotic spindle organization (Radford et al., 2012); microtubule depolymerization; mitotic chromosome movement towards spindle pole (Goshima and Vale, 2005); mitotic spindle organization (Pavlova et al., 2019; Goshima and Vale, 2005); spindle organization (Morales-Mulia and Scholey, 2005); kinesin complex (GO Reference Genome Project, 2011-). |

**Table B. Subdivisions of regulatory factors associated with the *Myc* target P1/P2 (plotted in Figure 12)**

| Gene name | Transcriptional Category | Activity/Function |
|---|---|---|
| wmd | **RNABPR** | **wing morphogenesis defect**; RNA binding (GO Reference Genome Project, 2011-); an essential WD-repeat protein required for wing development and motor behavior; imaginal disc-derived wing morphogenesis with involvement TGF-beta and Epidermal Growth Factor Receptor signaling pathways (Dworkin and Gibson, 2006); SMN complex (Survival Motor Neuron), involved in spliceosomal snRNP assembly in the cytoplasm and in pre-mRNA splicing in the nucleus (Carissimi et al., 2006). |
| PolD3 (pol32) | **DNAPOLD3; RRR; DBRR** | **DNA polymerase delta subunit 3**, non-essential subunit of the multi-subunit DNA polymerases delta and zeta; involved in embryonic DNA replication and promotes several types of DNA repair (Tritto et al., 2015), including homologous recombination repair of double-strand breaks; DNA polymerase processivity factor; repair via homologous recombination (Kane et al., 2012); DNA polymerase delta DNA polymerase zeta. |
| CRIF | **PSILEN** | **CR6-interacting factor**, positive regulation of post-transcriptional gene silencing by RNA, positive regulation of siRNA production, RNAi pathway, (Lim et al., 2014); Growth arrest/DNA-damage-inducible protein-interacting protein 1. |
| Polr2E | **RPIB5; RPIIB5; RPIIIB5; DBTR1; DBTR2; DBTR3; POLITR; POLIITR; POLIIITR** | **RNA polymerase II, I and III subunit E**, DNA binding (InterPro Project Members, 2004-); DNA-directed 5'-3' RNA polymerase activity; RNA pol II activity (Aoyagi and Wassarman, 2000); RNA pol I activity; RNA pol III activity (GO Reference Genome Project, 2011-); RNA Pol I Complex; RNA Pol II Complex; RNA pol III Complex. |
| (CG7583) CtBP(G) A0A0B4KHB2; CtBP(F) A0A0B4KGG9 | **COREP; DBTFB; COACT; CRTR2; POLIITR** | **C-terminal binding protein, isoform G**, corepressor targeting diverse transcription regulators; Hairy (*hry*)-interacting protein; positive regulator of Wnt/TCF pathway (Bhambhani et al., 2011; Zhang and Arnosti, 2011; Fang et al., 2006); negative regulation of canonical Wnt signaling pathway (Bhambhani et al., 2011; Fang et al., 2006); Wingless signaling pathway involved in embryo segmentation (Chan et al., 2008); chromatin remodeling (Emelyanov et al., 2012); DNA binding Transcription Factor Binding (Qi et al., 2008); transcription coactivator & corepressor (Fang et al., 2006); regulation of transcription by RNA pol II (Zhang and Arnosti, 2011); TORC Remodeling Complex. |

**Table B. Subdivisions of regulatory factors associated with the *Myc* target P1/P2 (plotted in Figure 12)**

| Gene name | Transcriptional Category | Activity/Function |
|---|---|---|
| fl(2)d | **RNABPR; RGEXPPR** | **female lethal d**, associated component of the WMM complex, mediates N6-methyladenosine (m6A) methylation of mRNAs, a modification that plays a role in the efficiency of mRNA splicing & is required for sex determination; mRNA binding (Lence et al., 2016); regulation of mRNA splicing, via spliceosome (Bawankar et al., 2021); RNA splicing via transesterification (Burnette et al., 1999). |
| SMC2 | **CBRR; CHRM** | **Structural maintenance of chromosomes protein 2**, condensation of prometaphase chromosomes, chromatin binding (GO Reference Genome Project, 2011-); condensation of prophase chromosomes; mitotic cell cycle (Ducat et al., 2008); regulation of microtubule assembly & organization in mitosis by the AAA+ ATPase Pontin, neurogenesis, stem cell differentiation (Liu et al., 2016). |
| Bap55 (Arp4) | **CBRR; CRR; RRR; POLIITR** | **Brahma associated protein 55kD**, member of two chromatin remodeling complexes; chromatin binding (GO Reference Genome Project, 2011-); DNA repair-dependent chromatin remodeling (Kusch et al., 2004); regulation of transcription by RNA pol II (Bonnay et al., 2014); chromatin remodeling (Kal et al., 2000); Notch signaling (Pillidge & Bray, 2019); response to EGFR signaling in the *Drosophila* wing (Terriente-Félix & de Celis, 2009); Brahma Associated Proteins Complex; TIP60 Complex; Non-Canonical Brahma Associated Proteins Complex; Polybromo-Containing Proteins Complex; SWRI Complex. |
| mbo, (Nup88) | **CBTR2; POLIITR** | **members only**, encodes a nucleoporin (nuclear pore complex protein Nup88), chromatin binding (Capelson et al., 2010); involved in immune response transduction by mediating the nuclear translocation of Mad. Toll-NF-κB, antimicrobial humoral response (Uv et al., 2000). |
| Capr | **RNABPT; TN; RGEXPPT** | **Caprin**, a cytoplasmic RNA granules protein (neuronal & stress granules); RNA-binding; regulation of embryonic mitotic cell cycle; cellularization; regulation of translation; ribonucleoprotein complex (Papoulas et al., 2010); translational regulator, Caprin-1 Dimerization Domain. |
| Nup358 | **RGEXPPT; TREX; PSILEN** | **Nucleoporin 358kD**, positive regulation of gene expression (He et al., 2017); positive regulation of RNA export from nucleus (Forler et al., 2004); SUMO transferase activity (Gene Ontology Curators, 2002-); protein translocation into nucleus, including Smad & phosphorylated Mad (Chen and Xu, 2010); transposon silencing & piRNA biogenesis (Parikh et al., 2018); Nuclear Pore Complex. |

**Table B. Subdivisions of regulatory factors associated with the *Myc* target P1/P2 (plotted in Figure 12)**

| Gene name | Transcriptional Category | Activity/Function |
|---|---|---|
| P32 | **CRTR2; CHRM** | **P32**, evolutionarily conserved mitochondrial glycoprotein, functions in presynaptic calcium signaling & neurotransmitter release as well as chromatin metabolism, histone binding; chromatin remodeling; nucleosome assembly; histone binding; sperm DNA binding (Emelyanov et al., 2014). |
| Caf1-55 | **POLIITR; CBTR2; DBTR2; CRTR2; HF; HDAC; RRR; DBRR** | **Chromatin assembly factor 1, p55 subunit**, a subunit of the MuvB core complex, MuvB core binds to the oncoprotein Myb & Rbf-E2f2-Dp tumor suppressor complex, thereby controlling the expression of many genes, including critical regulators of the cell cycle; histone binding (Nowak et al., 2011); histone deacetylase binding (Tyler et al., 1996); nucleosome binding (Nekrasov et al., 2005); chromatin remodeling (Mizuguchi et al., 2001); chromatin organization (Martinez-Balbas et al., 1998; Bulger et al., 1995); DNA replication-dependent chromatin assembly (Tyler et al., 1999; Ito et al., 1997; Tyler et al., 1996); positive regulation of transcription by RNA pol II (Mizuguchi et al., 2001); negative regulation of transcription by RNA pol II; transcription *cis*-regulatory region binding (Yao et al., 2018); nucleosome organization (Hamiche et al., 1999); Nucleosome Remodeling Factor; Polycomb Repressive Complex 2, PCL Variant; Chromatin Assembly Factor; Nucleosome Remodeling Deacetylase Complex; Polycomb Repressive Complex 2, JARID2-JING Variant. |
