## Supplemental Tables for "Targeting Regulatory Factors Associated with the *Drosophila Myc cis*-Elements by Reporter Expression, Gel Shift Assay, and Mass Spectrometric Protein Identification": Table C.pdf

**Table C. Subdivisions of regulatory factors associated with the Myc target P29/P30 (plotted in Figure 12)**

| Gene name | Pathway | Activity-Function |
|---|---|---|
| MED4 | **COACT; MS; POLIITR; TPI** | **Mediator of RNA pol II transcription subunit 4**, component of the Mediator complex transcription coactivator activity (Gu et al., 2002); transcription coregulator activity (Park et al., 2001); RNA pol II transcription; serves as a scaffold for the assembly of a functional preinitiation complex with RNA pol II and the general transcription factors (Gu et al., 2002; Park et al., 2001); Mediator Complex. |
| MED17 | **COACT; MS; POLIITR; TPI** | **Mediator of RNA pol II transcription subunit 17**, component of the Mediator complex, transcription coactivator activity (Gu et al., 2002); RNA pol II transcription; serves as a scaffold for the assembly of a functional preinitiation complex with RNA pol II and the general transcription factors; regulation of transcription by RNA pol II (Gu et al., 2002; Park et al., 2001); DNA binding transcription factor binding (Park et al., 2003); Mediator Complex. |
| MED25 | **COACT; POLIITR; TPI** | **Mediator of RNA poly II transcription subunit 25**, transcription coregulatory activity (Gene Ontology Curators, 2002-); regulation of transcription by RNA pol II (Boube et al., 2000); regulation of Imd and NF-κB Signaling pathways; positive regulation of antibacterial peptide biosynthetic process (Valanne et al., 2010); Mediator Complex. |
| pont | **DBRR; DBTR2; TF; RRR; CRTR2; CBRR; POLIITR; DNAHRR; DNAHTR2; RGEXPPT** | **Pontin**, involved in transcriptional regulation by RNA pol II (GO Reference Genome Project, 2011-), chromatin remodeling, DNA repair-dependent chromatin remodeling (Kusch et al., 2004); ribonucleoprotein assembly; transcriptional coregulator activity (Bellosta et al., 2005; Bauer et al., 2000); DNA helicase activity (GO Reference Genome Project, 2011-); positive regulation of gene expression (Diop et al., 2008). |
| E(bx) | **POLIITR; DBTR2; CRTR2; TF** | **Enhancer of bithorax**, lysine-acetylated histone binding; methylated histone binding (Kwon et al., 2009); RNA pol II *cis*-regulatory region sequence-specific DNA binding (GO Reference Genome Project, 2011-); chromatin remodeling (Bai et al., 2007; Badenhorst et al., 2002; Mizuguchi et al., 2001; Xiao et al., 2001); negative regulation of DNA-templated transcription (Yao et al., 2018); nucleosome organization (Hamiche et al., 1999); regulation of transcription (Badenhorst et al., 2002); Nucleosome Remodeling Complex; Unclassified DNA Binding Domain Transcription Factors. |

**Table C. Subdivisions of regulatory factors associated with the Myc target P29/P30 (plotted in Figure 12)**

| Gene name | Pathway | Activity-Function |
|---|---|---|
| lid (Kdm5) | **DBTR2; ACT; POLIITR; TF; CRR** | **Lysine-specific demethylase 5 (lid)**, activator of transcription (JAK/STAT) signaling (Tarayrah et al., 2015); regulation of cell growth, circadian rhythm, stress resistance, hematopoiesis & fertility; male germline stem cell population maintenance (Tarayrah et al., 2015; Lloret-Llinares et al., 2008; Secombe et al., 2007); histone H4R3 demethylase activity, histone reader activity (Liu and Secombe, 2015); pol II transcription (Secombe et al., 2007); H3K4m/H3K4m2/H3K4M3 demethylase activity (Tarayrah et al., 2015; Lloret-Llinares et al., 2008; Secombe et al., 2007); chromatin remodeling (GO Reference Genome Project, 2011-). |
| pds5 | **CHRM; ACT; REP** | **precocious dissociation of sisters 5**, sister chromatid separation at mitosis by removing the cohesin ring complex from chromosomes (Gause et al., 2010; Dorsett et al., 2005); influences gene activation & silencing through interactions with cohesin; required to initiate and/or maintain sister chromatid cohesion; TGF-alpha (ATR/Mei-41 kinase) (Barbosa et al., 2007); Armadillo-Like helical. |
| pzg | **CBRR; CRR; RRR; DBTR2; POLIITR; TF** | **Putzig**, involved in chromatin activation of replication related genes & regulation of signaling pathways including Notch, Ecdysone & JAK/STAT; chromatin binding (Kugler and Nagel, 2010); transcription *cis*-regulatory region binding; negative regulation of DNA-templated transcription (Yao et al., 2018); chromatin organization (Kugler and Nagel, 2007; Eggert et al., 2004); positive regulation of transcription by RNA pol II (GO Reference Genome Project, 2011-); regulation of growth, cell death and various developmental processes (Kugler et al., 2011); C2H2 Zinc Finger Transcription Factors. |
| Dbp80 | **RNAHTR2; RNABTR2; TREX** | **Dead box protein 80**, poly(A) + mRNA export from nucleus; RNA binding; RNA helicase activity (GO Reference Genome Project, 2011-); DEAD-Box RNA Helicases. |
| sbb | **COREG; COREP; POLIITR** | **Scribbler**, transcriptional co-regulator, acts mainly as a co-repressor (Haecker et al., 2007); negative regulation of transcription by RNA pol II (Kaminker et al., 2002); interactor of Grunge (*Gug*) & is used by the repressor Tailless (*tll*) in early embryos; Hedgehog signaling; Notch signaling wing disc D/V pattern formation (Bejarano et al., 2008); (Bejarano et al., 2008); Zinc Finger C2H2-Type. |

**Table C. Subdivisions of regulatory factors associated with the Myc target P29/P30 (plotted in Figure 12)**

| Gene name | Pathway | Activity-Function |
|---|---|---|
| Cpsf160 | **TT; PAFAC; RNABPR** | **Cleavage and polyadenylation specificity factor 160**, a key role in pre-mRNA 3'-end formation, recognizing the AAUAAA signal sequence and interacting with poly(A) polymerase and other factors to bring about cleavage and poly(A) addition (Salinas et al., 1998); mRNA polyadenylation (Mount and Salz, 2000); polyadenylation factors. |
| Trf2 | **PRF; CBTR2; DBTR2; TF; POLIITR; TPI; TI** | **TATA box-binding protein-like 1**, a core promoter recognition factor, mediates gene transcription; ecdysone signaling (Bashirullah et al., 2007); ecdysteroid signaling (Shima et al., 2007); chromatin binding (Wang et al., 2014); TFIIA-class transcription factor complex (Andersen et al., 2017); RNA pol II initiation factor (Hochheimer et al., 2002); RNA pol II core promoter sequence-specific DNA binding (Kedmi et al., 2014); TATA binding protein & TBP-Related Factor. |
| e(y)3 (SAYP) | **CBTR2; CRTR2; TPI; POLIITR; COREP; COACT** | **Enhancer of yellow 3, isoform D**, a nuclear protein; participates in gene activation in euchromatin as a component of both the SWI/SNF chromatin remodeling complex (Chalkley et al., 2008) & the TFIID transcription coactivator; contributes to gene silencing in pericentric heterochromatin; chromatin binding; transcription coactivator activity transcription corepressor activity; negative and positive regulation of transcription; coactivator of JAK/STAT signaling pathway (Shidlovskii et al., 2005); required for embryogenesis and oogenesis (Vorobyeva et al., 2009); Polybromo Containing Brahma Associated Proteins Complex. |
| Dhx15 | **RNAHPR; RNABPT; RGEXPPT** | **DEAH-box helicase 15**, RNA helicase activity; involved in mRNA processing (GO Reference Genome Project, 2011-). |
| Su(var)2-10 | **COACT; POLIITR; CHRM** | **Suppressor of variegation 2-10, isoform L**, a member of the PIAS protein family that regulates chromosome structure and function; chromosome organization & condensation (Hari et al., 2001); transcription coregulatory activity, regulation of transcription by RNA pol II (GO Reference Genome Project, 2011-); JAK/STAT pathway regulator, contributes to eye formation & eye determination (Betz et al., 2001); negative regulation of JAK/STAT signaling (Muller et al., 2005; Betz et al., 2001); positive regulators of Hedgehog signaling (Ma et al., 2016; Zhang et al., 2017); negative regulators of Imd signaling pathway (Tang et al., 2021; Cronin et al., 2009). |

**Table C. Subdivisions of regulatory factors associated with the Myc target P29/P30 (plotted in Figure 12)**

| Gene name | Pathway | Activity-Function |
|---|---|---|
| asun | **POLIITR; INTCOM** | **Protein asunder**, mitotic cell cycle (Lee et al., 2005); component of the Integrator complex involved in the transcription of small nuclear RNAs (snRNA) by RNA polymerase II-transcribed snRNAs & their 3'-box-dependent processing (Chen et al., 2012); cell cycle regulator Mat89Bb. |
| swm | **RNABPR; RGEXPPT; PSILEN** | **second mitotic wave missing**, regulates neural-specific glycosylation by binding to FucTA mRNA and facilitating its nuclear export in neural cells; negatively regulates Hedgehog (Hh) protein signal in wing development (Casso et al., 2008) mRNA binding; negative regulation of mRNA polyadenylation; positive regulation of gene expression (Yamamoto-Hino et al., 2010); RNA binding (GO Reference Genome Project, 2011-); mRNA processing (InterPro Project Members, 2004-). |
| CG10077(BcDNA:HL07910) | **RNAHPR; RNABPR** | RNA helicase, RNA binding activity; alternative mRNA splicing, via spliceosome; part of ribonucleoprotein complex (GO Reference Genome Project, 2011-); RNA helicase activity DEAD-Box RNA Helicases (Lasko, 2000). |
| spag | **POLIITR** | **Spaghetti**, homolog of RNA pol II-associated protein 3-like C-terminal (RPAP3, contains TPR-repeats towards the N-terminus); plays role in the assembly of RNA pol II complex (Benbahouche et al., 2014); negative regulation of motor neuron apoptotic process (Means et al., 2015). |
| smash | **U** | **Smallish, isoform H**, the *Drosophila* homolog of human LIM domain only 7 (LMO7); Domain of Unknown Function DUF 4757; Zinc Finger, Lim Type. |
| Mad | **DBTR2; POLIITR; TF; REP; ACT; COACT** | **Mothers against decapentaplegic**, the primary transcription factor, BMP signaling pathway core components (Vuilleumier et al., 2022; Guo et al., 2013; Weiss et al., 2010; Kamiya et al., 2008; Yao et al., 2006; Muller et al., 2003; Dai et al., 2000; Das et al., 1998; Inoue et al., 1998); wing development via EGFR & BMP pathways (Dworkin and Gibson, 2006; Lecuit et al., 1996); Wg/Wingless signaling (Bradley et al., 2001); DNA binding transcription activator activity, RNA pol II specific (Vuilleumier et al., 2022; Weiss et al., 2010; Saller and Bienz, 2001); DNA binding transcription repressor activity, RNA pol II specific (Yao et al., 2006; Muller et al., 2003); RNA pol II *cis*-regulatory region sequence-specific binding (Xu et al., 1998); transcription co-activator activity (Dai et al., 2000). |

**Table C. Subdivisions of regulatory factors associated with the Myc target P29/P30 (plotted in Figure 12)**

| Gene name | Pathway | Activity-Function |
|---|---|---|
| wmd | **RNABPR** | **wing morphogenesis defect**; RNA binding (GO Reference Genome Project, 2011-); an essential WD-repeat protein required for wing development and motor behavior; imaginal disc-derived wing morphogenesis with involvement of TGF-beta and Epidermal Growth Factor Receptor signaling pathways (Dworkin and Gibson, 2006). |
| His2Av | **DBRR; RRR; CBRR; RGEXPRR; HF** | **Histone H2A variant**, the only histone H2A variant (H2AV), a chimera of H2AZ & H2AX in *Drosophila*; required for larval hematopoiesis (Grigorian et al., 2017); DNA binding; structural constituent of chromatin (InterPro Project Members, 2004-); element of a class of active promoter structure proteins; gene regulation (Baldi and Becker, 2013); phosphorylated in response to DNA damage in the less conserved C-terminal tail; protein-containing complex binding (Kusch et al., 2004); cell division & kinetochore driven microtubule formation (Vernì and Cenci, 2015). |
| Btk29A | **RRR** | **Bruton tyrosine kinase** (non-specific protein-tyrosine kinase), DNA endoreplication (Chandrasekaran and Beckendorf, 2005); plays role in processes such as cellularization, morphogenesis, and germ cell proliferation (Hamada-Kawaguchi et al., 2015; Tsikala et al., 2014; Chandrasekaran and Beckendorf, 2005; Roulier et al., 1998) with the involvement of different cascades including JNK and TGF-β signaling pathways (Harden, 2002); Non-Receptor Tyrosine Kinases. |
| Ote | **DBTFB; COREP; POLIITR** | **Otefin**, a nuclear membrane-associated protein, acts in concert with BMP/DPP signaling to mediate "bag of marbles" (*bam*) transcriptional silencing; positive regulation of BMP/DPP signaling; DNA binding transcription factor binding; transcription corepressor activity; negative regulation of transcription; interacts with Medea/Smad4 at the (*bam*) silencer element to regulate germline stem cell (GSC) fate (Jiang et al., 2008). |
| mod | **POLIITR; DBTR2; CRR; TF** | **Modulo**, the *Drosophila* homologue of Nucleolin (Mikhaylova et al., 2006); association with the proto-oncogene MYC (perinn et al., 2003) (target of MYC selectively required for growth & proliferation); required for meiosis & spermatid differentiation in male germline cells; sequence-specific DNA binding (Mikhaylova et al., 2006); involved in chromatin packaging; dominant suppressor of variegation (Bantignies et al., 2002). |

**Table C. Subdivisions of regulatory factors associated with the Myc target P29/P30 (plotted in Figure 12)**

| Gene name | Pathway | Activity-Function |
|---|---|---|
| H | **DBTR2; COREP; SUP; REP; POLIITR** | **Hairless**, major antagonist of Notch during imaginal disc development; silencer of Notch targets by assembling a transcriptional repressor complex including transcription factor Suppressor of Hairless (*Su(H)*) & general co-repressors like Groucho (*gro*) & CtBP (*CtBP*); transcription corepressor activity (Maier et al., 2011; Brou et al., 1994); negative regulation of transcription by RNA pol II (Maier et al., 2011); negative regulator of Notch signaling (Smylla et al., 2019; Wolf et al., 2019; Berndt et al., 2017; Maier et al., 2013; Troost and Klein, 2012; Kurth et al., 2011; Maier et al., 2011; Lee et al., 2009; Nagel et al., 2005; Barolo et al., 2002; Go et al., 1998; Lyman and Yedvobnick, 1995); EGFR signaling; (Protzer et al., 2008); negative regulator of Wnt/TCF pathway (Bhambhani et al., 2011; Fang et al., 2006); positive regulator of Wnt/TCF pathway (Zhang and Arnosti, 2011). |
| caz, dFUS, Sarcoma-associated RNA-binding fly homolog, P19, SARFH | **CBTR2; RNABPR; COACT; TI; POLIITR; RGEXPPR; TPI** | **Cabeza** ('cabeza' means 'head' in Spanish), a chromatin binding protein (Mallik et al., 2018), RNA-binding protein, mRNA processing (Lasko, 2000), a single ortholog of human FUS in *Drosophila*; mRNA splicing (Herold et al., 2009); transcription initiation by RNA pol II promoter (Aoyagi and Wassarman, 2000); transcription coregulatory activity (GO Reference Genome Project, 2011-); synaptic assembly at the neuromuscular junction (Azuma et al., 2014); Cabeza shares homology domains with EWS & TLS, two human genes involved in chromosomal translocations with sarcoma formation; enriched in *Drosophila* head (David T. Stolow and Susan R. Haynes, 1995); Transcription Factor TFIID complex. |
| Rm62 | **RNAHPR; POLIITR; RGEXPPR** | **Rm62**, RNA helicase; pre-mRNA splicing, alternative splicing, rRNA processing, miRNA processing & transcription regulation (Ishizuka et al., 2002); DEAD-BOX RNA HELICASES. |
| Hmg-2 (HMGB2) | **TF; POLIITR; DBTR2; CRTR2; HDAC** | **High mobility group protein 2**, together with the Wnt signaling regulate chondrocyte hypertrophy by mediating Runt-related transcription factor 2 expression (Taniguchi et al., 2018); Wnt signaling & HMGB2 regulate articular cartilage surface maintenance (Taniguchi et al., 2009); association with chromatin, ubiquitous distribution in nucleus, and non-specific DNA minor groove binding; DNA bending / unwinding and promotion of DNA flexibility (McCauley et al., 2005; Lorenz et al., 1999); regulation of gene expression (GO Reference Genome Project, 2011-). |

**Table C. Subdivisions of regulatory factors associated with the Myc target P29/P30 (plotted in Figure 12)**

| Gene name | Pathway | Activity-Function |
|---|---|---|
| Bin1 (SAP18) | **COREP; HDAC; HF; CBTR2; POLIITR; RNABTR2** | **Bicoid interacting protein 1 Bin 1 (Histone deacetylase complex subunit SAP18)**, regulation of hedgehog (Hh) signaling pathway by transcription factor Gli in mammals, repression of Gli-mediated transcription by Su(fu) in cooperation with SAP18 for the recruitment of the SAP18-mSin3 complex to promoters containing the Gli-binding element (Yan Cheng and Bisho, 2002); transcription corepressor activity; heterochromatin formation (Matyash et al., 2009); negative regulation of DNA-templated transcription (Matyash et al., 2009; Zhu et al., 2001). |
| psi | **POLIITR; DBTR2; RNABPR** | **P-element somatic inhibitor, isoform C**, dual roles in RNA processing (Labourier et al., 2002; Siebel et al., 1994) & transcriptional regulation; required for activating MYC transcription. KSRP (KHSRP) binding & destabilization of mRNA; mRNA binding (Siebel et al., 1994); regulation of DNA-templated transcription (InterPro Project Members, 2004-); far upstream element-binding. |
| vas | **RNABTN; RNAHTN; TN; RGEXPTN** | **Vasa**, a DEAD-Box RNA helicase protein (Mahowald, 2001; van Eeden and St. Johnston, 1999; Cooperstock and Lipshitz, 1997), promotes translation of (*grk*) & (*mei-P26*) mRNAs (Liu et al., 2009); functions in piRNA biogenesis as a component of an amplifier complex; maternal transcripts required for oogenesis, transposon silencing in the female germ line, A-P embryonic patterning, & germ cell specification; RNA binding (Sengoku et al., 2006); DEAD Box RNA Helicases. |
| fon (CG1582) | **RNABTN; RNAHTN; TN** | **Fondue, isoform B**, RNA helicase activity; translation initiation activity (Linsalata et al., 2019; Lasko, 2000); RNA binding activity (GO Reference Genome Project, 2011-); DEAH-Box RNA Helicases. |
| AGO2 | **RNABPT; RGEXPPT; PSILEN** | **Argonaut 2**, interaction with small interfering RNAs (siRNAs) for the formation of RNA-induced silencing complexes (RISCs), siRNA binding (Goh and Okamura, 2019; Kawamura et al., 2008; Tomari et al., 2007; Rand et al., 2005; Lingel et al., 2003); mRNA-mediated gene silencing (Besnard-Guérin et al., 2015); single-stranded RNA binding (Goh and Okamura, 2019); siRNA binding (Goh and Okamura, 2019; Kawamura et al., 2008; Tomari et al., 2007; Rand et al., 2005; Lingel et al., 2003). |

**Table C. Subdivisions of regulatory factors associated with the Myc target P29/P30 (plotted in Figure 12)**

| Gene name | Pathway | Activity-Function |
|---|---|---|
| (CG7583) CtBP(G) A0A0B4KHB2; CtBP(F) A0A0B4KGG9 | **COREP; DBTFB; COACT; CRTR2; POLIITR** | **C-terminal binding protein, isoform G**, corepressor targeting diverse transcription regulators; Hairy (*hry*)-interacting protein; positive regulator of Wnt/TCF pathway (Bhambhani et al., 2011; Zhang and Arnosti, 2011; Fang et al., 2006); negative regulation of canonical Wnt signaling pathway (Bhambhani et al., 2011; Fang et al., 2006); embryo segmentation (Chan et al., 2008); chromatin remodeling (Emelyanov et al., 2012); DNA binding transcription factor binding (Qi et al., 2008); transcription coactivator & corepressor (Fang et al., 2006); regulation of transcription by RNA pol II (Zhang and Arnosti, 2011); TORC Remodeling Complex. |
| sqd | **RNABTN; TN; RGEXPPT** | **Squid, isoform E**, Dpp signaling, squid & Hephaestus (but not Hrb27C) are necessary for proper functioning of bone morpho-genetic protein (BMP) signaling in GSCs; novel roles for RNA binding proteins Squid, Hephaestus & Hrb27C in *Drosophila* oogenesis (Finger et al., 2022); negative regulation of RNA splicing (Ji and Tulin, 2009); negative regulation of translation (Cooperstock and Lipshitz, 1997); regulation of gene expression (GO Reference Genome Project, 2011-). |
| Bre1 | **POLIITR; CRTR2** | **E3 ubiquitin protein ligase**, required for Notch signaling and histone modification; chromatin organization (Bray et al., 2005), E3 ubiquitin ligase, interacts with E2 ubiquitin ligase Ubc6, ubiquitination of H2B on lysine 120 at most RNA pol II transcripts; negative regulation of heterochromatin formation; positive regulation of transcription; Toll-dependent Bre1/Rad6-cact feedback loop in controlling host innate immune response (Cai et al., 2022); Unclassified Ring Domain Ubiquitin Ligases. |
| CG9705 | **TF; RNABPT; RGEXPPT** | **Cold shock domain-containing protein CG9705,** regulation of mRNA stability; mRNA 3'UTR binding (GO Reference Genome Project, 2011-); transcription factor, involved in the regulation of gene expression, dendrite morphogenesis (Iyer et al., 2013); Cold-shock (CSD) Domain. |
| larp | **DBMTRR; RNABTN; MTRR; RGEXPTN; TN** | **La related protein, isoform F**, poly(A) binding protein; 3'-UTR binding; poly-pyrimidine tract binding (Martin et al., 2022); male meiotic nuclear division (Blagden et al., 2009; Ichihara et al., 2007); Ras/MAPK signaling (Blagden et al., 2009); positive regulation of mitochondrial DNA replication (Zhang et al., 2016); mRNA stabilization (InterPro Project Members, 2004-); RNA binding; positive regulation of translation (GO Reference Genome Project, 2011-). |

**Table C. Subdivisions of regulatory factors associated with the Myc target P29/P30 (plotted in Figure 12)**

| Gene name | Pathway | Activity-Function |
|---|---|---|
| btz | **RNABTN; TN; RNABPR; RGEXPPT** | **Barentsz (CASC3, mammalian homolog) (other name MLN51)**, a component of Exon Junction Complex pathway (EXJC), recruited to spliced mRNAs to mark introns removal site; translation activator linking the EJC & the translation machinery, direct role in protein synthesis (Chazal et al., 2013); BTZ domain found on CASC3 (cancer susceptibility candidate gene 3 protein, also known as **MLN51= M**etastatic **L**ymph **N**ode **51)** also known as Barentsz (Btz); CASC3, component of EJC involved in post-transcriptional regulation of mRNA in metazoa; complex formed by association of 4 proteins (eIF4AIII, Barentsz, Mago, & Y14), mRNA, & ATP; BTZ wraps around eIF4AIII & stacks against the 5' nucleotide (Bono et al., 2006; Palacios et al., 2004); Barentsz (MLN51) overexpressed in breast cancer (Degot et al., 2002); repression of MNL51 by brain-specific miR-128 (Wilkinson et al., 2011); mRNA binding; RNA binding (InterPro Project Members, 2004-). |
| lin-28 | **RNABPT; RGEXPPT** | **Lin-28** protein, cold shock and RNA-binding protein; positive regulation of receptor signaling pathway via JAK/STAT (Sreejith et al., 2019); regulator of developmental timing; regulator of microRNA maturation; post-transcriptional regulation of gene expression (Luhur et al., 2017; Chen et al., 2015). |
| mask | **RNABTR2; POLIITR** | **Multiple ankyrin repeats single KH domain**, RNA binding (InterPro Project Members, 2004-); positive regulation of transcription by RNA pol II (Sansores-Garcia et al., 2013; Sidor et al., 2013). |
| Mi-2 | **CBTR2; CRTR2; DBTR2; POLIITR; CHRM** | **Mi-2**, a nuclear ATP-dependent nucleosome (chromatin) remodeler activity (Kunert et al., 2009; Murawska et al., 2008); chromatin binding (Kunert et al., 2009); required for repression of cell type-specific genes; DNA binding (GO Reference Genome Project, 2011-); chromosome condensation (Nikalayevich and Ohkura, 2015); regulation of transcription by RNA pol II (Li et al., 2010); Notch signaling (Zacharioudaki E, Falo Sanjuan J, Bray S., Elife. 2019); involved in Wingless and ecdysone signaling pathways (Kon et al., 2005); Nucleosome Remodeling Deacetylase Activity; SNF2-Like Chromatin Remodelers. |

**Table C. Subdivisions of regulatory factors associated with the Myc target P29/P30 (plotted in Figure 12)**

| Gene name | Pathway | Activity/Function |
|---|---|---|
| glu (CG11397) | **CHRM; CBRR** | **Gluon**, a subunit of the multiprotein complex Condensin, mitotic cell cycle: required for prometaphase chromosome condensation (Nikalayevich and Ohkura, 2015); sister chromatid segregation; nervous system development; glucose metabolism; expressed in dividing cells throughout embryogenesis—pole cells, neuroblasts in the CNS & the PNS; chromosome condensation (Bivik et al., 2015); chromatin binding (Gene Ontology Curators, 2002-); Structural Maintenance of Chromosomes Family. |
| sm (CG9218) | **RNABPR; RGEXPPR** | **Smooth**, heterogeneous nuclear ribonucleoprotein, *Drosophila* smooth (sm) closest to human hnRNP (zur Lage et al. 1997); mRNA processing; RNA binding; regulation of RNA splicing (GO Reference Genome Project, 2011-); RNA binding (InterPro Project Members, 2004-); ecdysteroid signaling & insulin-like peptide in context of determination of adult lifespan & in conjunction with ImpL2 (ecdysone-inducible gene L2) (Paik et al., 2012); RNA Recognition Motif. |
| Ddx1 | **RNABPR; RNAHPT; CBTR2; COREG; DBTR2** | **Dead-box-1**, a member of the DEAD box family of RNA helicases that bind and unwind double-stranded RNA; DNA/RNA helicase activity, chromatin binding; transcription coregulatory activity (Gene Ontology Curators, 2002-); DEAD-Box 1 is a novel & independent prognostic marker for early recurrence in breast cancer (Germain et al., 2011). |
| Pop2 | **TN; PSILEN; RGEXPPR** | **poly(A)-specific ribonuclease**; involved in translation inhibition (Ruscica et al., 2019; Braun et al., 2011); miRNA-mediated mRNA degradation (Braun et al., 2011); poly(A)-specific ribonuclease activity (Temme et al., 2004); nuclear-transcribed mRNA poly(A) tail shortening (Temme et al., 2010; Bönisch et al., 2007). |
| Sirt4 | **HDAC; MTTR** | **Sirtuin4,** Sole mitochondrial Sirtuin in *Drosophila*, NAD-dependent protein deacylase; catalyzes the NAD-dependent hydrolysis of acyl groups from lysine residues (Feller et al., 2015);transcriptional activation of mitochondrial biogenesis, mitochondrial regulator of life span & metabolism, cellular response to starvation, Sirt4 knockout causes short lifespan, increased sensitivity to starvation, decreased fertility, & activity (Wood et al., 2018). |

**Table C. Subdivisions of regulatory factors associated with the Myc target P29/P30 (plotted in Figure 12)**

| Gene name | Pathway | Activity-Function |
|---|---|---|
| Bap111 | **DBTR2; CRTR2; POLIITR; TF** | **Brahma associated protein 111kD**, extensive homology of the Brahma (BRM) complex to SWI/ SNF; DNA binding; chromatin remodeling (Papoulas et al., 2001); transcription induction (Kal et al., 2000); Brahma Associated Proteins Complex; Polybromo-Containing Proteins Complex; High Mobility Group Box Transcription Factors. |
| mod(mdg4) | **DBTR2; CHRM; POLIITR; ENBLOC; CBTR2** | **modifier of mdg4, isoform AD**, a nuclear protein, specifically interacts with various DNA binding proteins; involved in meiotic chromosome condensation (Matsui et al., 2011); male meiotic chromosome segregation (Soltani-Bejnood et al., 2007); regulation of transcription & enhancer blocking; chromatin binding (Bag et al., 2019; Gerasimova et al., 1995) DNA binding (Ogiyama et al., 2018; Bonchuk et al., 2011); POZ (Pox virus and Zinc finger) domain binding (Harvey et al., 1997); regulation of transcription by RNA pol II (GO Reference Genome Project, 2011-); SKP1/BTB/POZ Domain Superfamily. |
| IntS11 | **RNABTR2; INTCOM; POLIITR** | **Integrator complex subunit 11,** involved in the transcription of small nuclear RNAs (snRNA) & their 3'-box-dependent processing; pol II transcription (Baillat et al., 2005); Notch signaling (Shersher et.al., 2021); Epidermal Growth Factor Receptor pathway (Tilley and Mollet, 2021). |
| Sxc | **CHRSILEN; RGEXPPT** | **Super sex combs**, a Polycomb group gene, a O-GlcNAc transferase, involved in epigenetic gene silencing; regulation of gene expression (Akan et al., 2016); positive regulation of Fibroblast Growth Factor Receptor (FGFR) signaling pathway (Mariappa et al., 2011). |
| Hrb98DE | **RNABTN; TN; PSILEN; RGEXPPT** | **Heterogeneous nuclear ribonucleoprotein at 98DE**, nuclear RNA-binding protein, controls hnRNA stability, splicing, IRES-dependent translation, & translational repression; represents one of the main targets of the poly(ADP-ribosyl)ation pathway; also regulates tissue polarity patterning & germ-line stem cell fate; negative regulation of RNA splicing; mRNA 5'-UTR binding (Ji et al., 2009); RNA binding (InterPro Project Members, 2004-); mRNA binding (Lasko, 2000); post-translational modification of hnRNPs, such as poly(ADP-ribosyl)ation, is an important mechanism in the regulation of gene expression during development, such as eye pattern formation (Ji et al., 2013); Spliceosome Complex A. |

**Table C. Subdivisions of regulatory factors associated with the Myc target P29/P30 (plotted in Figure 12)**

| Gene name | Pathway | Activity-Function |
|---|---|---|
| Elp4 (CG6907) | **POLIITR; TN; RRR; RNABTN** | **Elongator complex protein 4**, establishment of mitotic spindle asymmetry; regulation of translation through targeted transfer-RNA (tRNA) modification (Planelles-Herrero et al., 2022); phosphorylase kinase regulator activity; regulation of transcription by RNA pol II (Gene Ontology Curators, 2002-); Elongation Complex. |
| Tudor-SN | **RNABTN; TN; RGEXPPT** | **Tudor staphylococcal nuclease**, shows activity towards both DNA and RNA substrates; translation regulation through its association with the RNA-induced silencing complex (RISC) (Ku et al., 2016; Caudy et al., 2003); regulatory ncRNA-mediated post-transcriptional gene silencing (InterPro Project Members, 2004-); (GO Reference Genome Project, 2011-). |
| rump | **RNABPR; RGEXPPR** | **Rumpelstiltskin**, the *Drosophila* hnRNP M homolog. It binds to nanos & oskar mRNAs, plays a role in mRNA localization to the germ plasm by diffusion/entrapment during late stages of oogenesis; splicing factor, binds to exonic splicing enhancers on pre-mRNA targets; mRNA 3'-UTR binding (Sinsimer et al., 2011; Jain and Gavis, 2008); mRNA binding (Lasko, 2000); RNA binding (InterPro Project Members, 2004-); mitotic cell cycle (Ducat et al., 2008). |
| abs | **RNAHPR; RNABPR; RGEXPPR** | **Abstrakt**, DEAD/DEAH-Box RNA helicase (Lasko, 2000), RNA binding (GO Reference Genome Project, 2011-); RNA splicing, via spliceosome (Herold et al., 2009); regulates cell polarity in oocytes & embryos; downregulation of Notch signaling in asymmetric cell division in ganglion mother cell (GMC2-4a) in interaction with Inscuteable (*Insc*) (Irion et al., 2004). |
| Ndf | **DBTR1; DBTR2; DBTR3; POLITR; POLIITR; POLIIITR; CBTR2; TE; CRTR2** | **Nucleosome-destabilizing factor**, H3K36me3 binding in active gene bodies, DNA binding, nucleosome binding, chromatin binding, transcription elongation-coupled chromatin remodeling (Fei et al., 2018). |
| Tailor | **POLIIITR; RGEXPTR3; RNABTR3; RNABPR** | **Tailor**, part of the terminal RNA uridylation-mediated processing (TRUMP) complex; involved in 3'-to-5' exoribonucleolytic decay of RNA species by Dis3l2 (Lin et al., 2017); regulation of microRNA biogenesis by targeting precursor-microRNAs (predominantly mirtron hairpins) & targets unprocessed RNA pol III transcripts for degradation in a cytoplasmic RNA surveillance pathway (Reimão-Pinto et al., 2015); cytoplasmic RNA-specific terminal uridylyltransferase (Cheng et al., 2019; Lin et al., 2017; Reimão-Pinto et al., 2015), Uridyltransferases. |

**Table C. Subdivisions of regulatory factors associated with the Myc target P29/P30 (plotted in Figure 12)**

| Gene name | Pathway | Activity-Function |
|---|---|---|
| aub | **RGEXPPT; RNABPT** | **Aubergine**, piRNA binding (Huang et al., 2021; Webster et al., 2015; Nagao et al., 2010); global gene silencing by mRNA cleavage (Kennerdell et al., 2002) ncRNA-mediated post-transcriptional gene silencing (Tomari et al., 2004); RNA-mediated gene silencing (Bozzetti et al., 2015); JAK/STAT signaling pathway controls host defense in the gut by regulating stem cell proliferation & epithelial cell homeostasis (Cronin et al., 2009); RNA binding (Brennecke et al., 2007). |
| mre11 | **DBRR; RRR; TMM; CHRM** | **meiotic recombination 11**, Double-strand break repair protein; G2/M DNA damage checkpoint signaling; (Bi et al., 2006); chromosome organization (Bi et al., 2006; Ciapponi et al., 2004); telomere capping/maintenance (Gao et al., 2009; Bi et al., 2005; Bi et al., 2004; Ciapponi et al., 2004). |
| Rcc1 | **CBRR; CHRM** | **Regulator of chromosome condensation**, nuclear import & export of β-Catenin (Koyama et at., 2017); regulation of mitotic cell cycle, apoptosis pathway (Trieselmann and Wilde, 2002); regulation of chromatin condensation. |
| CG8003 | **RGEXPPT** | Expressed in adult head and organism; orthologous to human ANKMY2 (ankyrin repeat and MYND domain containing 2). |
| Nup50 | **DBTR2; POLIITR** | **Nuclear pore complex protein Nup50** (Nucleoporin 50kD), chromatin DNA binding; positive regulation of transcription by RNA pol II (Kalverda et al., 2010); an important protein for TGF-β signal transduction by mediating the nuclear translocation of the Mothers against dpp protein (Mad); association of 400 genes interacting with Nup50 Nucleoporin, among which transcriptionally active genes inside the nucleoplasm are predominantly involved in development & cell cycle regulation (Kalverda et al., 2010); Nuclear Pore Complex. |
| Sf3a1 | **RNABPR; RGEXPPR** | **Splicing factor 3a subunit 1**, RNA binding (GO Reference Genome Project, 2011-); RNA processing (InterPro Project Members, 2004-). |
| CG12134 | **U** | Uncharacterized, unknown; WD40/YVTN Repeat-like Containing Domain Superfamily. |
| shep | **REP; DBTR2; CBTR2; POLIITR** | **Protein Alan shepard**, an evolutionarily conserved RNA/DNA binding protein; regulation of alternative splicing & gypsy insulator activities (chromatin insulator associated) (Chen et al., 2021). |

**Table C. Subdivisions of regulatory factors associated with the Myc target P29/P30 (plotted in Figure 12)**

| Gene name | Pathway | Activity/Function |
|---|---|---|
| wapl | **CRR; ACT; REP; CHRM; CHRSILEN** | **wings apart-like**, (*wapl*) & precocious dissociation of sisters 5 (*pds5*) proteins form the releasin complex for separation of sister chromatid at mitosis by removing the cohesin ring complex from chromosomes; gene activation & silencing via interaction with cohesin (Gause et al., 2010; Dorsett et al., 2005); chromatin organization; sister chromatid cohesin (Verni et al., 2000); regulation of Wnt/Wingless signaling pathway to determine the wing size in the mediterranean fruit fly (medfly) (Cho et al., 2013). |
| Mlf | **TF; POLIITR** | **Myelodysplasia/myeloid leukemia factor**, interacts with various factors involved in transcriptional regulation, regulates cell proliferation during eye morphogenesis, Lozenge activity during hematopoietic development and assembly of the COP9 signalosome complex (Miller et al., 2017); regulation of transcription (Bras et al., 2012). |
| rin (CG9412) | **RNABPR; RNAHPR; RGEXPPR** | **Rasputin**, evolutionarily conserved RNA-binding protein, RNA helicase activity (Gene Ontology Curators, 2002-); functions as a link between Ras signaling pathway & RNA metabolism (Costa et al., 2013; Pazman et al., 2000); Rho-mediated signaling pathway (Pazman et al., 2000); mRNA binding (GO Reference Genome Project, 2011-); positive regulation of gene expression (Costa et al., 2013); response to amino acid starvation (Aguilera-Gomez et al., 2017); Unclassified RNA Helicases. |
| Fmr1 | **RNABPT; RNAHPT; RGEXPPT; PSILEN** | **Fragile X messenger ribonucleoprotein 1**, channel binding protein; DEAD/H-Box RNA helicase binding (Barbee et al., 2006); mRNA regulatory element binding repressor activity (Fajner et al., 2021); post-transcriptional regulation of gene expression (Luhur et al., 2017). |
| CG4849 | **RNABTN; RGEXPPT; TN** | U5 snRNA binding activity (GO Reference Genome Project, 2011-); translation elongation (Lasko, 2000); positive regulation of gene expression (Ashton-Beaucage et al., 2014). |
| crp | **TF; POLIITR; DBTR2** | **Activator protein 4 (Cropped)**, a downstream target of MYC, plays role in cell growth, organ size & survival; myosin binding (Liu et al., 2008); transcription factor; sequence-specific DNA binding (King-Jones et al., 1999); RNA pol II *cis*-regulatory region, sequence-specific DNA binding (GO Reference Genome Project, 2011-); protein dimerization activity (InterPro Project Members, 2004-); larval somatic muscle development (Dobi et al., 2014); Basic Helix-Loop-Helix Transcription Factors. |

**Table C. Subdivisions of regulatory factors associated with the Myc target P29/P30 (plotted in Figure 12)**

| Gene name | Pathway | Activity/Function |
|---|---|---|
| rept | **DBTR1; DBTR2; CBRR; DNAHRR; DNAHTR1; DNAHTR2; RRR; CRTR2; HF; RGEXPPT; POLITR; COACT; POLIITR; HAT** | **Reptin, RuvB-like helicase**, involved in transcriptional regulation; DNA repair-dependent chromatin remodeling; NuA4 histone acetyltransferase complex (Kusch et al., 2004); heterochromatin formation (Qi et al., 2006); negative regulation of gene expression (Diop et al., 2008); positive regulation of transcription of nucleolar large rRNA by RNA polymerase I (Vinayagam et al., 2016); regulation of transcription by RNA pol II (GO Reference Genome Project, 2011-); core component of the chromatin remodeling Ino80 complex; chromatin remodeling (Klymenko et al., 2006); transcriptional coactivator in Wg signaling caused by altered Armadillo (*arm*) signaling; Pontin/Reptin antagonistically interfere with the nuclear Armadillo signaling; an essential cofactor for the normal function of Myc; required for cellular proliferation and growth; ecdysone-mediated salivary gland cell autophagy cell death (Ihry and Bashirullah, 2014); negative regulation of canonical Wnt signaling (Bauer et al., 2000); regulation of cell population proliferation (Bellosta et al., 2005); INO80 Complex; TIP60 Complex; SWR1 Complex; RUVB-Like DNA Helicases. |
| Usp10 | **POLIITR; TE; RGEXPPT** | **Ubiquitin specific protease 10**, cysteine-type deubiquitinase activity; ubiquitin-dependent protein catabolic process (Gene Ontology Curators, 2002-); negative regulation of transcription elongation (GO Reference Genome Project, 2011-); positive regulation of Notch signaling (Zhang et al., 2012). |
| Chro | **CBTR2; CRTR2; POLIITR; CHRM** | **Chromator**, a chromodomain protein, chromatin binding; chromatin organization; negative regulation of transcription by RNA pol II (GO Reference Genome Project, 2011-); chromosome organization (Ding et al., 2009; Rath et al., 2006); metamorphosis (Wasser et al., 2007); microtubule spindle formation, cell cycle progression, functions as a spatial regulator of cell cycle factors (Ding et al., 2009). |
| Usp7 | **HF; RRR** | **Ubiquitin-specific protease 7**, positive regulation of heterochromatin formation (van der Knaap et al., 2005); Nucleotide Excision Repair (NER) (Higa et al., 2018). |
| Nurf-38 | **CBTR2; CRR; POLIITR; RRR** | **Nucleosome remodeling factor - 38kD**, component of NURF (Nucleosome Remodeling Factor), ATP-dependent nucleosome sliding & facilitates transcription of chromatin (Gdula et al., 1998); chromatin remodeling (Mizuguchi et al., 2001; Gdula et al., 1998); positive regulation of transcription (Mizuguchi et al., 2001); ecdysone receptor-mediated signaling pathway (Badenhorst et al., 2005). |

**Table C. Subdivisions of regulatory factors associated with the Myc target P29/P30 (plotted in Figure 12)**

| Gene name | Pathway | Activity/Function |
|---|---|---|
| Rpt4 | **POLITR; DBTR1** | **Regulatory particle triple-A ATPase 4**, 19S proteasomal ATPase, a component of the 26S proteasome complex, nucleolar protein & regulator of rRNA transcription; physical interaction with the tumor suppressor protein "Birt Hogg Dube" (BHD), RNA pol I transcription, regulatory region sequence-specific DNA binding (association of Rpt4 with the rDNA locus) (Gaur et al., 2013); Notch-mediated follicle cell differentiation and cell cycle switches, regulation of insulin-PI3K signaling pathway (Jia et al., 2015). |
| Snr1 | **CRTR2; COACT; POLIITR; TF; DBTR2** | **Snf5-related 1**, (ATP-dependent chromatin remodeler; a counterpart of yeast SWI/SNF) (Swi3 component of the Brahma Associated Proteins Complex), SET Domain Binding (Marenda et al., 2003); cell differentiation; Tat protein binding (Koe et al., 2014); transcription coactivator activity (GO Reference Genome Project, 2011-); chromatin remodeling (Dingwall et al., 1995); regulation of transcription by RNA pol II (Bonnay et al., 2014); positive regulation of DNA-templated transcription (Dingwall et al., 1995; Kal et al., 2000). |
| CDK2AP1 | **HDAC; CRTR2; REP; POLIITR** | **CDK2-associated protein 1**, DNA polymerase binding; positive regulation of protein phosphorylation (Gene Ontology Curators, 2002-); association with Nucleosome Remodeling Deacetylase (NURD) Complex; tumor suppressor gene and *Drosophila* homolog of mammalian DOC1 (Deleted in Oral Cancer) (Reddy et al., 2010); Transcription repression (Mohd-Sarip, 2017). |
| MRG15 | **HAT; CRTR2; CBTR2; CRR; CBRR; RGEXPPT; POLIITR; CHRM; RRR; HF** | **MORF-related gene 15**, histone acetylation; chromatin binding; positive regulation of gene expression (Huang et al., 2017); chromatin organization; chromosome separation (Smith et al., 2013); DNA repair-dependent chromatin remodeling (Kusch et al., 2004); heterochromatin formation (Qi et al., 2006); regulation of DNA-templated transcription (InterPro Project Members, 2004-). |
| Smr | **COREP; POLIITR; CBTR2** | **Smrter, SMRT-related** ecdysone receptor-interacting protein; transcriptional corepressor (Kozlova and Thummel, 2000) mediating transcriptional silencing of the ecdysone receptor; chromatin binding (Tsuda et al., 2006); negative regulation of transcription by RNA pol II (Heck et al., 2012); Insulin-PI3K signaling pathway (Jia et al., 2015); Notch signaling pathway (Jia et al., 2015; Heck et al., 2012); ecdysteroid signaling (Heck et al., 2012); SANT-MYB Domain Transcription Regulators. |

**Table C. Subdivisions of regulatory factors associated with the Myc target P29/P30 (plotted in Figure 12)**

| Gene name | Pathway | Activity/Function |
|---|---|---|
| Hrb27C | **RNABTN; TN; DBTR2; RGEXPTN** | **Heterogeneous nuclear ribonucleoprotein at 27C**, component of ribonucleosomes; mRNA 3'-UTR binding (Nelson et al., 2007); ssDNA binding (Matunis et al., 1992); translation repressor activity (Szostak et al., 2018); regulation of border cell migration; (Mathieu et al., 2007); PVR (the *Drosophila* PDGF/VEGF receptor). |
| PolD3 (pol32) | **DNAPOLD3; RRR; DBRR** | **DNA polymerase delta subunit 3**, non-essential subunit of the multi-subunit DNA polymerases delta and zeta; involved in embryonic DNA replication and promotes several types of DNA repair (Tritto et al., 2015), including homologous recombination repair of double-strand breaks; DNA polymerase processivity factor; DNA repair via homologous recombination (Kane et al., 2012); DNA polymerase delta DNA polymerase zeta. |
| NfI (Q86P06) | **TF; DBTR2; POLIITR** | **Nuclear factor I, isoform B**, CCAAT box-binding transcription factor (CTF) (Mermod et al., 1989); STAT3 signaling (Chen et al, 2017; Stringer et al, 2016); RNA pol II *cis*-regulatory region sequence-specific DNA binding; regulation of transcription by RNA pol II (GO Reference Genome Project, 2011-); canonical Wnt signaling (High-mobility group AT-Hook 1 mediates the role of nuclear factor I/X in osteogenic differentiation through activating canonical Wnt signaling (Wu et al, 2021); MAD Homology Domain Transcription Factors. |
| CG17493 | **RRR** | Centriole replication (GO Reference Genome Project, 2011-); active in centriole and centrosome. |
| tral | **RNAHPT; RNABPT; RGEXPPT** | **Trailer hitch, isoform D**, DEAD/H-Box RNA helicase binding (Barbee et al., 2006; Wilhelm et al., 2005); mRNA binding; (GO Reference Genome Project, 2011-); RNA binding (InterPro Project Members, 2004-); piRNA pathway; retrotransposon silencing (Liu et al., 2011); required for oocyte dorsoventral patterning via actin and microtubule cytoskeleton organization. |
| TfIIEalpha | **TPI; TI; POLIITR; TF; DBTR2** | **Transcription factor IIEα**, the largest subunit of the RNA pol II general transcription factor TFIIE; TfIIEα & TfIIEβ essential for transcription initiation in vitro acting with RNA pol II & the other general transcription factors; mRNA transcription by RNA pol II (Wang et al., 1997); transcription initiation at RNA pol II promoter (Aoyagi and Wassarman, 2000); transcription open complex formation at RNA pol II promoter (GO Reference Genome Project, 2011-); Transcription Factor II E. |

**Table C. Subdivisions of regulatory factors associated with the Myc target P29/P30 (plotted in Figure 12)**

| Gene name | Pathway | Activity/Function |
|---|---|---|
| Cdc6 | **DBRR; RRR** | **Cell division cycle 6**, encodes an essential component of the pre-replication complex (preRC) together with the origin recognition complex, the 'Double parked' (*dup*) & MCM2-7 proteins (InterPro Project Members, 2004-). |
| CG17002 | **COREP; POLIITR; HDAC** | Transcription coregulator activity; regulation of transcription by RNA pol II (GO Reference Genome Project, 2011-); a component of the Nuclear Receptor Corepressor Histone Deacetylase, N-CoR-HDAC3 (Zhang et al., 2002) and SMRT corepressor complexes (essential for estrogen alpha-mediated transcriptional regulation) (Cheng and Kao, 2009). |
| D1 | **TF; DBTR2; REP** | **D1 chromosomal protein**, a multi-AT-hook chromosomal protein, associates with AT-rich satellites, including the SAT-III repeats of the X-chromosome (Blattes et al., 2006); binds to the minor-groove of the DNA (Levinger and Varshavsky, 1982); favors heterochromatin-mediated gene repression involving its interaction with topoisomerase II; Unclassified DNA Binding Domain Transcription Factors. |
| Polr2G (RPB7) | **RPIIB7; DBTR2; RNABTR2; TI; POLIITR** | **RNA pol II subunit G**, ssDNA binding activity; transcription initiation activity; RNA pol II nuclear-transcribed mRNA catabolic process, exonucleolytic; positive regulation of nuclear-transcribed mRNA poly(A) tail shortening (GO Reference Genome Project, 2011-); transcription by RNA pol II (Aoyagi and Wassarman, 2000). |
| pds5 | **CHRM; ACT; REP** | **precocious dissociation of sisters 5**, sister chromatid separation at mitosis by removing the cohesin ring complex from chromosomes (Gause et al., 2010; Dorsett et al., 2005); influences gene activation & silencing through interactions with cohesin; required to initiate and/or maintain sister chromatid cohesion; TGF-alpha (ATR/Mei-41 kinase) (Barbosa et al., 2007); Armadillo-Like helical. |
| Bap55 | **CBRR; CRR; RRR; POLIITR** | **Brahma associated protein 55kDa**, chromatin binding (GO Reference Genome Project, 2011-); DNA repair-dependent chromatin remodeling (Kusch et al., 2004); regulation of transcription by RNA pol II (Bonnay et al., 2014); chromatin remodeling (Kal et al., 2000); Notch signaling (Pillidge & Bray, 2019); response to EGFR signaling in the *Drosophila* wing (Terriente-Félix & de Celis, 2009); Brahma Associated Proteins Complex; TP60 Complex; Non-Canonical Brahma Associated Proteins Complex; Polybromo-Containing Proteins Complex; SWRI Complex. |

**Table C. Subdivisions of regulatory factors associated with the Myc target P29/P30 (plotted in Figure 12)**

| Gene name | Pathway | Activity/Function |
|---|---|---|
| CG2962 | **POLIITR; COACT** | Transcription coactivator activity; positive regulation of transcription by RNA pol II (GO Reference Genome Project, 2011-). |
| Polr2B (RpII140) | **RPIIB2; POLIITR; TI; TE; DBTR2** | **RNA polymerase II subunit B**, present in all RNA pol II complexes; DNA-directed 5'-3' RNA polymerase activity (Falkenburg et al., 1987); RNA polymerase II activity (Aoyagi and Wassarman, 2000). |
| Sas-6 | **RRR** | **Spindle assembly abnormal 6**, a centriole protein, essential for centriole assembly, homo-oligomerizes to form a 9-fold symmetric "cartwheel" structure, plays an important part in setting the 9-fold symmetry of the assembling centriole (Jana et al., 2018); centriole replication (Stevens et al., 2010; Dobbelaere et al., 2008; Peel et al., 2007); centriole duplication (Gene Ontology Curators, 2002-). |
| Isha (cg4266) | **POLIITR; TT; CBTR2; RNABTR2; INSUL** | **Insulator su(Hw) mRNA adaptor**, chromatin binding activity; mRNA binding activity (Bag et al., 2022; Lasko, 2000); negative regulation of transcription by RNA pol II; required for gypsy insulator function (Bag et al., 2022). |
| REPTOR | **TF; POLIITR; DBTR2** | **Repressed by TOR**, a transcription factor, shuttles between cytoplasm & nucleus based on the activity of mechanistic target of rapamycin (*mTor*); together with its binding partner REPTOR-BP, it mediates much of the transcriptional response upon Tor complex 1 inhibition; response to starvation; TORC1 signaling; positive regulation of transcription (Tiebe et al., 2015); DNA binding transcription factor activity (GO Reference Genome Project, 2011-; InterPro Project Members, 2004-); Basic Leucine Zipper Transcription Factors. |
| Taf6 | **POLIITR; TF; TI; COACT; TPI** | **TBP-associated factor 6**; part of the multisubunit basal transcription factor TFIID; pol II general transcription initiation factor (Wright et al., 2006; Hansen and Tjian, 1995); transcription co-activator activity (GO Reference Genome Project, 2011-); transcription by RNA pol II (Hansen and Tjian, 1995); TFIID Complex. |
| Taf12 | **POLIITR; DBTR2; COACT; TPI; TI** | **TBP-associated factor 12**, part of the multisubunit basal transcription factor TFIID; pol II general transcription initiation factor (Yokomori et al., 1993) forms a histone-like pair with Taf4; Taf12 also an integral component of the *Drosophila* SAGA histone acetyltransferase complex; pol II preinitiation complex assembly (GO Reference Genome Project, 2011-); SAGA complex; TFIIFD Complex. |

**Table C. Subdivisions of regulatory factors associated with the Myc target P29/P30 (plotted in Figure 12)**

| Gene name | Pathway | Activity/Function |
|---|---|---|
| su(sable) su(s) | **POLIITR; TT; SUP; RGEXPPR** | **suppressor of sable**, su(sable) & Wdr82 (Wdr82 turned up on the protein identification list but not enriched), part of a transcription termination checkpoint; promotes transcription termination of RNAs & their subsequent degradation by the nuclear exosome; promotes transcription termination of aberrant RNAs, transcripts from genes containing a transposon inserted at their very 5' end or RNAs from heat-shock-inducible repetitive elements (Brewer-Jensen et al., 2016; Kuan et al., 2009; Voelker et al., 1991); nuclear RNA surveillance (Brewer-Jensen et al., 2016; Kuan et al., 2009); negative regulation of transcription termination (Brewer-Jensen et al., 2016; negative regulation of transcription (suppressor) (Kuan et al., 2004); protein Suppressor of sable; Zinc finger, CCCH-type. |
| Cp190 | **TF; DBTR2; CBTR2; INSULB** | **Centrosomal protein 190kD**, a C2H2 zinc finger transcription factor, binds to most boundaries of contact domains; defined by enhanced internal contact frequencies; required to form most contact domain boundaries distal to a transcribed promoter; essential for early development, prevention of regulatory cross-talk between specific gene loci that pattern the embryo; chromatin binding (Whitfield et al., 1995); chromatin insulator binding (Erokhin et al., 2010; Oliver et al., 2010; Mohan et al., 2007); DNA binding (Pai et al., 2004); POZ domain binding (Bonchuk et al., 2011). |
| Cdk1 | **RRR** | **Cyclin-dependent kinase 1**, regulation of DNA replication initiation (Seller and O'Farrell, 2018). |
| CG6683 | **POLIITR** | Involved in regulation of transcription by RNA polymerase II; part of transcription regulator complex (GO Reference Genome Project, 2011-); active in nucleus; MAD-BESS Domain Transcription Regulators. |
| ash2 | **DBTR2; TI; COACT; POLIITR; HMOD; CRTR2** | **absent, small, or homeotic discs 2**, a component of the histone methyltransferase complex, specifically methylates lysine 4 of histone H3 & a member of the trithorax family; mutants show homeotic transformations & a variety of pattern-formation defects; DNA binding (Pérez-Lluch et al., 2011); imaginal disc-derived wing vein specification (Angulo et al., 2004); transcription coactivator activity (Carbonell et al., 2013); chromatin remodeling (Beltran et al., 2007); transcription initiation coupled chromatin remodeling (La Jeunesse and Shearn, 1995); histone H3-H4 methylation (InterPro Project Members, 2004-). |

**Table C. Subdivisions of regulatory factors associated with the Myc target P29/P30 (plotted in Figure 12)**

| Gene name | Pathway | Activity/Function |
|---|---|---|
| HDAC6 | **HDAC; RGEXPPT; CRTR2; POLIITR** | **Histone Deacetylase 6, isoform G**, cytosolic deacetylase (Miskiewicz et al., 2014), key modulator of proteostasis by mediating ubiquitin-proteasomal & lysosomal degradation of native and/or misfolded proteins; histone deacetylase activity (Feller et al., 2015; Cho et al., 2005; Barlow et al., 2001); regulation of DNA-templated transcription (Cho et al., 2005) negative regulation of transcription by RNA pol II (GO Reference Genome Project, 2011-); chromatin organization (InterPro Project Members, 2004-); RPD3/HDA1 Lysine Deacetylases. |
| Ssb-c31a | **DBTR2; REP; ACT; POLIITR** | **Single stranded-binding protein c31A**, characterized as a nuclear protein binds specifically to NssBF element, a 26 nucleotides sequence in the long terminal repeat (LTR) of the *Drosophila* melanogaster 1731 retrotransposon to repress promoter activity; negative regulation of DNA-templated transcription (Lacoste et al., 1995); DNA binding; positive regulation of transcription by RNA pol II (InterPro Project Members, 2004-); RNA pol II transcription Coactivator Sub 1/Tcp4-like; Transcription Coactivator p15 (Pc4), C-terminal. |
| Rae1 | **RNABPT; RGEXPPT; TREX; RRR** | **Rae1**, a nucleoporin member of the WD40-repeat β propeller protein super family with pleiotropic functions including poly(A)+ mRNA export, cell cycle regulation, male meiosis control & male germ cell post-meiotic differentiation; RNA binding; transcription-dependent tethering of RNA pol II gene DNA at nuclear periphery (GO Reference Genome Project, 2011-); male meiotic I (Volpi et al., 2013); positive regulation of gene expression (Tian et al., 2011); Nuclear Pore Complex. |
| ebi | **REP; POLIITR; COREP; DBTFB** | **Ebi**, **F-box-like/WD repeat-containing protein**, evolutionarily conserved repressor/silencer; JNK signaling: Ebi/ AP-1 complex (activator protein 1) represses pro-/anti-apoptotic genes; suppresses basal transcription levels of apoptotic genes protecting sensory neurons degeneration (Lim et al., 2012); regulation of EGFR (Dong et al., 1999) and Notch pathways (Nguyen et al., 2016; Marygold et al., 2011; Tsuda et al., 2002); negative regulation of transcription (Lim et al., 2012); DNA binding transcription factor binding (Qi et al., 2008). |
| CG5543 | **U** | **Gastrulation defective protein 1 homolog**, Active in nucleus, located at the site of DNA double-strand break. |

**Table C. Subdivisions of regulatory factors associated with the Myc target P29/P30 (plotted in Figure 12)**

| Gene name | Pathway | Activity/Function |
|---|---|---|
| RfC4 | **DBRR; RRR** | **Replication factor C subunit 4**, DNA binding; DNA replication (Krause et al., 2001); DNA clamp loader activity (Gene Ontology Curators, 2002-); DNA strand elongation involved in DNA replication; DNA damage check point signaling; DNA replication check point signaling; DNA repair (Mossi et al., 1997); sister chromatid cohesion; leading strand elongation (Gene Ontology Curators, 2002-); ELG1 Complex; RFC Complex ATPases. |
| Nup98-96 | **CBTR2; RNABTR2; TI; POLIITR; TMM** | **Nucleoporin 98-96kDa**, precursor of Nup98 and Nup96 proteins, two integral parts of the nuclear pore; Nup98-96 loss of function results in premature differentiation of germ line cells at the expense of proliferation; chromatin DNA binding (Ilyin et al., 2017; Pascual-Garcia et al., 2017; Kalverda et al., 2010); promoter-enhancer loop anchoring (Pascual-Garcia et al., 2017); promoter-specific chromatin binding (Pascual-Garcia et al., 2017; Pascual-Garcia et al., 2014); RNA binding (GO Reference Genome Project, 2011-); heat shock-mediated polytene chromosome puffing (Kalverda et al., 2010); positive regulation of transcription by RNA pol II (Panda et al., 2014; Pascual-Garcia et al., 2014; Capelson et al., 2010; Kalverda et al., 2010); positive regulation of transcription initiation by RNA pol II (Capelson et al., 2010); post-transcriptional tethering of RNA pol II gene DNA to nuclear periphery (GO Reference Genome Project, 2011-); telomere tethering to nuclear periphery (GO Reference Genome Project, 2011-); Nuclear Pore Complex. |
| XRCC1 | **RRR; DBRR** | **X-ray repair cross complementing 1**, damaged DNA binding activity; involved in base-excision repair; active in nucleus; single strand break repair; double-strand break repair via nonhomologous end joining (InterPro Project Members, 2004-). |
| jub | **COREP; POLIITR; RRR** | **Ajuba LIM protein, isoform D**, LIM domain containing protein and regulator of Hippo signaling, bind to & inhibits activation of the Hippo pathway kinase Warts (Huang et al., 2016; Fletcher et al., 2015; Rauskolb et al., 2014; Sun and Irvine, 2013; Rauskolb et al., 2011; Das Thakur et al., 2010); transcription corepressor activity; positive regulation of miRNA-mediated gene silencing; regulation of DNA-templated transcription; transcription regulator complex (GO Reference Genome Project, 2011-); mitotic cell cycle/DNA replication (Sabino et al., 2011). |

**Table C. Subdivisions of regulatory factors associated with the Myc target P29/P30 (plotted in Figure 12)**

| Gene name | Pathway | Activity/Function |
|---|---|---|
| RnrL | **RRR** | **Ribonucleoside diphosphate reductase large subunit**, preparation of precursors necessary for DNA synthesis; catalysis of the biosynthesis of deoxyribonucleotides from the corresponding ribonucleotides (Gene Ontology Curators, 2002-); DNA replication (InterPro Project Members, 2004-). |
| dpa | **DBRR; RRR; DNAHRR** | **disc proliferation abnormal**, 3'-5' DNA helicase activity (Moyer et al., 2006); DNA binding (InterPro Project Members, 2004-); DNA replication (Pflumm and Botchan, 2001); mitotic DNA replication initiation (GO Reference Genome Project, 2011-); essential for 'once per cell cycle' DNA replication initiation and elongation in eukaryotic cells; required for DNA replication and cell proliferation. essential role in mitotic DNA replication but not in endoreplication; MCM2-7 complex. |
| Mcm3 | **DBRR; RRR; DNAHRR** | **Minichromosome maintenance 3**, a component of the MCM2-7 hexamer, forms part of the CMG complex, together with the CDC45L & GINS proteins resulting in the main DNA helicase complex (Moyer et al., 2006), functions during DNA replication; DNA binding; ssDNA binding; duplex DNA unwinding; DNA replication initiation (InterPro Project Members, 2004-). |
| Rfc37 (CG8142) | **RRR; DBRR** | **Replication factor C37**, ATP binding activity; ATP hydrolysis activity; DNA binding activity (InterPro Project Members, 2004-);contributes to DNA clamp loader activity (GO Reference Genome Project, 2011-); involved in DNA repair and DNA-templated DNA replication; part of Elg1 RFC-like complex. |
| Orc6 | **RRR; DBRR** | **Origin recognition complex subunit 6**, a subunit of the origin recognition complex (ORC), which is essential for the initiation of DNA replication in eukaryotic cells (Chesnokov et al., 1999); mitotic cell cycle (Balasov et al., 2009). |
| cutlet | **CHRM; DBRR** | **Cutlet**, chromosome cohesion factor, (chromosome transmission fidelity protein 18 homolog) cell cycle; involved in sister chromatid cohesion and fidelity of chromosome transmission; DNA clamp loader activity; positive regulation of DNA-directed DNA polymerase activity (Gene Ontology Curators, 2002-); CTF18-replication factor C (CTF18-RFC); DNA binding (GO Reference Genome Project, 2011-); RFC Complex ATPases. |

**Table C. Subdivisions of regulatory factors associated with the Myc target P29/P30 (plotted in Figure 12)**

| Gene name | Pathway | Activity/Function |
|---|---|---|
| La (CG10922) | **TT; POLIIITR; DBTR3; RNABTR3** | **La autoantigen-like**, involved in transcription termination by RNA pol III; binds RNA, DNA, and precursors of RNA pol III transcripts; exhibits specialized role during fly development. |
| PolE1 (DNApol-epsilon, DNApol-ε255) | **DNAPOLE1; RRR; DBRR** | **DNA polymerase epsilon subunit 1** (DNA pol-epsilon, DNA pol-ε, DNA pol-ε255, DNA polymerase ε, l(3)pl10), the large subunit of DNA pol epsilon, an essential DNA polymerase participating with DNA pol alpha & DNA pol delta in cellular DNA replication. The C-terminal and N-terminal regions have differential requirements in mitotic and endo-replicating cells; 3'-5' DNA exonuclease activity (Oshige et al., 2004; Aoyagi et al., 1997); DNA replication proofreading (Oshige et al., 2004); endomitotic cells (Suyari et al., 2012). |
| Prim2 | **DBRR; RRR; PRIM2** | **DNA primase large subunit**, synthesis of short RNA-DNA primers on the lagging strand during DNA synthesis (Kuroda et al., 1990; Cotterill et al., 1987; Kaguni et al., 1983); compound eye morphogenesis (Chen et al., 2000); alpha DNA polymerase: primase complex (Kuroda et al., 1990; Cotterill et al., 1987; Kaguni et al., 1983; Kaguni et al., 1983; Villani et al., 1980). |
| PolD2 (Pol31) | **RRR; DBRR** | **DNA polymerase delta subunit 2**, accessory component of both the DNA polymerase delta complex and possibly the DNA polymerase zeta complex; as a component of the delta complex, participates in high fidelity genome replication, including lagging strand synthesis, DNA recombination & repair; promotes the function of the DNA pol-delta complex accessory subunit PolD3 in both embryonic & postembryonic somatic cells (Ji et al., 2019); DNA binding; DNA replication (InterPro Project Members, 2004-); mitotic DNA templated DNA replication (Ji et al., 2019); DNA strand elongation involved in DNA replication (GO Reference Genome Project, 2011-). |
| Nup93-1 | **CBPT; RGEXPPT** | **Nucleoporin 93kD-1**, direct interactor of Nup154 chromatin binding; negative regulation of gene expression, epigenetic; chromatin DNA binding (Gozalo et al., 2020). |
| tex | **TREX** | **Transcription export complex**, part of THO complex part of transcription export complex; mRNA export from nucleus (GO Reference Genome Project, 2011-; InterPro Project Members, 2004-); Transcription Export Complex. |

**Table C. Subdivisions of regulatory factors associated with the Myc target P29/P30 (plotted in Figure 12)**

| **Gene name** | **Pathway** | **Activity/Function** |
|---|---|---|
| Ciz1 | **RRR; DBRR** | **Ciz1, Cip1 Interacting Zinc Finger Protein 1**, nuclear protein; negative regulation of cell cycle, regulation of the subcellular localization of p21(CIP/WAF1); consensus DNA sequence, ARYSR(0–2)YYAC, recognized by Ciz1 (Warder and Keherly, 2003); no evidence found in transcriptional regulation (Mitsui et al., 1999); DNA replication factor (Lukasik et al., 2008). |
| SMC3 | **CBRR; DBRR; CHRM; RRR** | **Structural maintenance of chromosomes 3**, a subunit of the cohesin complex, involved in planar cell polarity by regulating the membrane enrichment of the transmembrane cadherin encoded by "starry night" (*stan*); mitotic sister chromatid cohesion; cell division (GO Reference Genome Project, 2011-); involved in Notch and Wnt/Frizzled-PCP signaling pathways for the establishment of imaginal disc-derived wing hair orientation; imaginal disc-derived wing morphogenesis (Mouri et al., 2012); chromatin binding, double-stranded DNA binding (Gene Ontology Curators, 2002-); chromosome organization and DNA condensation (InterPro Project Members, 2004-). |
| Mcm5 | **RRR; DBRR; CHRM** | **Minichromosome maintenance 5**, involved in mitotic DNA replication; required to resolve meiotic double-strand breaks into crossovers; 3'-5' helicase activity (Moyer et al., 2006); DNA binding; DNA replication origin binding (InterPro Project Members, 2004-); DNA duplex unwinding; chromosome organization/DNA condensation (Christensen and Tye, 2003); DNA endoreplication (Park and Asano, 2008). |
| Rfc38 | **RRR; DBRR** | **Replication factor C 38kD subunit**; DNA clamp loader activity; involved in DNA repair (GO Reference Genome Project, 2011-); leading strand elongation; sister chromatid cohesion (Gene Ontology Curators, 2002-); part of Elg1 RFC-like complex; DNA binding; DNA replication (InterPro Project Members, 2004-). |
| Rfc37 (CG8142) | **RRR; DBRR** | ATP binding activity; ATP hydrolysis activity; DNA binding activity (InterPro Project Members, 2004-); contributes to DNA clamp loader activity (GO Reference Genome Project, 2011-); involved in DNA repair and DNA-templated DNA replication; part of Elg1 RFC-like complex. |
| Helz | **RNAHPT; PSILEN** | **Helicase with zinc finger**, RNA helicase activity (Hanet et al., 2019); involved in ncRNA-mediated post-transcriptional gene silencing (GO Reference Genome Project, 2011-). |

**Table C. Subdivisions of regulatory factors associated with the Myc target P29/P30 (plotted in Figure 12)**

| Gene name | Pathway | Activity/Function |
|---|---|---|
| PolD1 | **RRR; DBRR** | **DNA polymerase delta subunit 1 (**DNA-directed DNA polymerase**)**, catalytic component of the DNA polymerase delta complex, crucial role in high fidelity genome replication, including lagging strand synthesis (Aoyagi et al., 1994); DNA recombination & repair; DNA polymerase & 3'-5'-exonuclease activities (Aoyagi et al., 1994; Chiang et al., 1993; Peck et al., 1992); required at the nucleus of rapidly dividing embryonic cells to activate genome replication during the earliest cell proliferation (Ji et al., 2019); DNA-directed DNA polymerase activity (InterPro Project Members, 2004-);base-excision repair gap filling (GO Reference Genome Project, 2011-). |
| CG30291 | **POLIITR** | Probable regulator of cell proliferation, may regulate CDK5, NF-kappa-B-mediated gene transcription & p53/TP53 activation; regulation of mitotic cell cycle (GO Reference Genome Project, 2011-); regulation of cyclin-dependent protein Ser/Thr kinase activity (Gene Ontology Curators, 2002-); CDK5 regulatory subunit-associated protein 3. |
| CG13850 | **MTRNABPR; RGEXPMT** | RNA binding; mitochondrial RNA processing; regulation of mitochondrial mRNA stability (GO Reference Genome Project, 2011-). |
| Lam (CG6944) | **CBRR; HF** | **Lamin**, chromatin binding (Verboon et al., 2015); Heterochromatin formation (Verboon et al., 2015; Dialynas et al., 2010; Shevelyov et al., 2009); regulation of meiosis I cytokinesis (Hayashi et al., 2016); Lamins. |
| POLDIP2 | **POLDIP2; DBRR; RRR** | **Polymerase (DNA-directed) delta interacting protein 2**, DNA binding activity (InterPro Project Members, 2004-); involved in error-free translesion synthesis (GO Reference Genome Project, 2011-); active in mitochondrial nucleoid and nucleus. |
| CCG31673 | **U** | Glycerate dehydrogenase, glyoxylate reductase (NADP(+)), oxidoreductase activity, actin on CH-OH group of donors NAD or NADP as acceptor (InterPro Project Members, 2004-); The catalytic domain contains a number of conserved charged residues which may play a role in the catalytic mechanism (Dengler et al., 1997); Unclassified CH-OH Oxidoreductases, NAD or NADP as Acceptor. |
| l(2)10685 | **RNABPT; RGEXPPT** | **Lethal (2) 10685**, methyltransferase activity (GO Reference Genome Project, 2011-); S-adenosylmethionine-dependent methyltransferase activity (InterPro Project Members, 2004-); orthologous to human NSUN4 (NOP2/Sun RNA methyltransferase 4); Unclassified Methyltransferases. |

**Table C. Subdivisions of regulatory factors associated with the Myc target P29/P30 (plotted in Figure 12)**

| Gene name | Pathway | Activity/Function |
|---|---|---|
| CG10077 (BcDNA:HL07910) | **RNAHPR; RNABPR** | RNA helicase, RNA binding activity & RNA helicase activity (GO Reference Genome Project, 2011-); DEAD-Box RNA Helicases. |
| DCP1 | **RNABPT; RGEXPPT** | **Decapping protein 1**, a subunit of the mRNA decapping holoenzyme (Nishihara et al., 2013); involved in oskar mRNA localization (Lee et al., 2020; Lin et al., 2006) and miRNA-mediated gene silencing; mRNA binding (GO Reference Genome Project, 2011-). |
| ncd | **CHRM** | **non-claret disjunctional**, a minus-end-directed kinesin microtubule motor protein and the sole member of the kinesin-14 motor family; chromosome segregation (Hallen et al., 2008); distributive segregation (Whyte et al., 1993); mRNA transport (Fahmy et al., 2014). |
| CG12384 (BcDNA:RH17411) | **POLIITR** | Negative regulation of DNA-templated transcription (Gene Ontology Curators, 2002-); apoptotic signaling pathway (GO Reference Genome Project, 2011-); cellular response to amino acid starvation; Death-Associated Protein1 (DAP1/DAPL1). |
| SMC5 | **DBRR; CHRM; RRR** | **Structural maintenance of chromosomes protein 2**, a component of the SMC5/6 protein complex, critical to genome stability & required for homologous DNA recombination-based processes; single-stranded DNA binding (GO Reference Genome Project, 2011-); damaged DNA binding (Gene Ontology Curators, 2002-); DNA damage response (Li et al., 2013). |
| IntS14 | **TT; RNABPR; RGEXPPR; INTCOM** | **Integrator 14**, component of the Integrator complex, a complex involved in the transcription of small nuclear RNAs (snRNA) and their 3'-box-dependent processing; involved in the 3'-end processing of the U7 snRNA, and also the spliceosomal snRNAs U1 and U5 (Chen et al., 2012). |
| CG8064 | **RNABPT; RGEXPPT** | SnoRNA binding; involved in maturation of SSU-rRNA (GO Reference Genome Project, 2011-); human ortholog(s) implicated in papillary thyroid carcinoma; orthologous to human WD repeat domain 3 (WDR3). |
| CG8149 | **TREX; TT** | Orthologous to human SARNP (SAP domain containing ribonucleoprotein), poly(A) + mRNA export from nucleus (GO Reference Genome Project, 2011-); SAP Domain/DNA binding domain containing protein (Bejarano et al., 2008). |

**Table C. Subdivisions of regulatory factors associated with the Myc target P29/P30 (plotted in Figure 12)**

| Gene name | Pathway | Activity/Function |
|---|---|---|
| Not3 | **RNABPR; RNABTN; TN; RGEXPPT; POLIITR** | **CCR4-NOT transcription complex subunit 3**, a poly(A)-specific ribonuclease involved in translation inhibition (Bawankar et al., 2013); nuclear transcribed mRNA catabolic process, deadenylation-dependent decay (Arvola et al., 2020); nuclear transcribed mRNA poly(A) tail shortening (Temme et al., 2010); regulation of DNA-templated transcription (InterPro Project Members, 2004-). |
| Patr-1 | **RNABPR; RGEXPPR** | **Protein associated with topo II related – 1**, a P body component involved in deadenylation-dependent decapping of nuclear transcribed-mRNA (Nishihara et al., 2013; Braun et al., 2010) & regulation of synaptic growth at neuromuscular junctions (Pradhan et al., 2012); RNA binding (GO Reference Genome Project, 2011-); positive regulation of mRNA catabolic process (Nish ihara et al., 2013). |
| Ge-1 | **PSILEN; RNABPR; RGEXPPT** | **Ge-1**, RNAi pathway (miRNA-mediated gene silencing) (Eulalio et al., 2009); signaling pathways that activate NF-κB, Toll & Imd pathways (Jin et al., 2008; De Gregorio et al., 2002); mRNA degradation with role in mRNA decapping (GO Reference Genome Project, 2011-; Eulalio et al., 2009); Enhancer of mRNA-Decapping Protein 4. |
| Atu | **TI; TE; POLIITR** | **Another transcription unit**, RNA pol II C-terminal domain phosphoserine binding; positive regulation of transcription by RNA pol II; positive regulation of transcription elongation by RNA pol II (GO Reference Genome Project, 2011-); histone modification (InterPro Project Members, 2004-); RNA Polymerase-Associated Factor 1 Complex; part of Cdc73/Paf1 complex. |
| bsf | **RNABPT; PAFAC; RGEXPPT; MTTR** | **bicoid stability factor**, a member of the family of proteins containing the pentatricopeptide motif, RNA binding domain; functions in mitochondrial mRNA stability (Pajak et al., 2019; Jaiswal et al., 2015) & post-transcriptional control of gene expression; multiple roles in mitochondrial gene expression; required for progression through oogenesis and viability; mRNA 3'-UTR binding (Mancebo et al., 2001); mitochondrial mRNA polyadenylation (Bratic et al., 2011); regulation of mitochondrial gene expression (Matsushima et al., 2017); regulation of mitochondrial transcription (Bratic et al., 2011). |
| Edc3 | **RNABPR; RGEXPPT** | **Enhancer of decapping 3**, deadenylation-independent decapping of nuclear-transcribed mRNAs (GO Reference Genome Project, 2011-); RNA binding (InterPro Project Members, 2004-); mRNA catabolic process (Tritschler et al., 2007). |

**Table C. Subdivisions of regulatory factors associated with the Myc target P29/P30 (plotted in Figure 12)**

| Gene name | Pathway | Activity/Function |
|---|---|---|
| Dis3 | **RNABPR; RGEXPPR** | **Dis3**, RNA endo-/exonuclease activity (Snee et al., 2016); RNA binding (InterPro Project Members, 2004-); nuclear RNA surveillance (Kuan et al., 2009); regulation of gene expression (Kiss and Andrulis, 2010); rRNA catabolic process (GO Reference Genome Project, 2011-); RNA Exosome Complex. |
| CG44270 | **U** | Expressed in several structures, including anterior ectoderm, ectoderm anlage, embryonic central brain neurons, extended germ band embryo, and somatic precursor cell; Domain of Unknown Function, DUF4799. |
| CG12708 | **U** | Uncharacterized; Unknown; Domain of Unknown Function, DUF 4799. |
| CG43813 | **U** | Uncharacterized; Unknown. |
| CG8187 | **U** | Uncharacterized; Putative Adhesion. |
| CG7406 | **U** | Uncharacterized; Domain of Unknown Function, DUF4766 |
| CG7173 | **U** | Expressed in embryonic dorsal epidermis, embryonic head epidermis, embryonic ventral epidermis, and spermatozoon; contains KAZAL Domain, KAZAL Domain Superfamily. |
| CG15731 | **U** | Expressed in several structures, including anterior endoderm anlage, embryonic epidermis, and germ layer; Domain of Unknown Function, DUF4766. |
| Nin (Bsg25D) | **RRR** | **Ninein**, a microtubule-anchoring protein in humans; microtubule binding (Kowanda et al., 2016); microtubule anchoring at centrosome (GO Reference Genome Project, 2011-); microtubule organizing center attachment site organization (Zheng et al., 2016). |
| CG15239 | **U** | Expressed in embryonic dorsal epidermis; embryonic esophagus; embryonic head epidermis; embryonic ventral epidermis; and embryonic/larval salivary gland; Domain of Unknown Function, DUF4773. |
| Hyls1 | **U** | **Hyls1**, centriolar and ciliogenesis, involved in cilium assembly; located in centriole & ciliary basal body. |
| CG15506 | **U** | Uncharacterized, Unknown. |

**Table C. Subdivisions of regulatory factors associated with the Myc target P29/P30 (plotted in Figure 12)**

| Gene name | Pathway | Activity/Function |
|---|---|---|
| CG1648 | **U** | Molecular Function: Uncharacterized, Unknown. |
| fit | **U** | **Female-specific independent of transformer**, expressed in adult fat body, adult head, and fat cell; Molecular Function: Uncharacterized, Unknown. |
| scny | **COREP; HF; POLIITR** | **Scrawny**, ubiquitinyl hydrolase 1; hydrolase & deubiquitination activity deubiquitinating Imd and prevention of constitutive activation of Imd/NF-κB cascade (Engel et al., 2014; Buszczak et al 2009; Thevenon et al 2009); functions as a transcriptional repressor by continually deubiquitinating histone H2B at the promoters of genes critical for cellular differentiation, thereby preventing histone H3 'Lys-4' trimethylation (H3K4me3); heterochromatin formation (Buszczak et al., 2009); Ubiquitin Specific Protease (USP) Deubiquitinases. |
| Dmac2 (CG4042) | **U** | Mitochondrial respiratory chain complex I assembly factor (Gene Ontology Curators, 2002-); cellular component: mitochondrion (Murari et al., 2021); orthologous to human DMAC2 (distal membrane arm assembly component 2). |
| Hsc70-4 | **RGEXPPT; PSILEN** | **Heat shock protein 70 cognate 4**, regulatory ncRNA-mediated post-transcriptional gene silencing (Dorner et al., 2006) |
| gro | **COREP; POLIITR** | **Groucho, isoform F**, a global developmental co-repressor in collaboration with DNA-binding repressor partner proteins & tethering to target promoters (Ajuria et al., 2011; Giagtzoglou et al., 2003; Jimenez et al., 2000; Goldstein et al., 1999; Valentine et al., 1998); downstream effector of signaling pathways such as Wg/Wnt & Dpp/TGF-beta; phosphorylation & attenuation of Groucho repressor activity in response to MAPK activation; "context dependent regulatory domain binding" (CRD), a domain of about 130aa, the most divergent region among the LEF/TCF proteins (Arce et al., 2009). |
| CG7518 | **U** | Uncharacterized, FAM 193 Family |
| CG10543 | **TF; POLIITR; DBTR2** | Nuclear protein, DNA binding transcription factor activity, RNA polymerase II-specific; RNA polymerase II transcription regulatory region sequence-specific DNA binding activity; zinc ion binding activity (GO Reference Genome Project, 2011-); involved in gravitaxis (Armstrong et al., 2006); C2H2 Zinc Finger Transcription Factors. |

**Table C. Subdivisions of regulatory factors associated with the Myc target P29/P30 (plotted in Figure 12)**

| Gene name | Pathway | Activity/Function |
|---|---|---|
| lost | **RNABPR; RGEXPPR** | (**lost**), interacts with the RNA-binding protein rumpelstiltskin (*rump*) for posterior localization of mRNAs by diffusion/entrapment during late stages of oogenesis; mRNA splicing, via spliceosome (Herold et al., 2009); localization to various RNP complexes with a broad role in RNA metabolism. |
| Ars2 | **RGEXPPT; PSILEN; POLIITR** | **Arsenic resistance protein 2**, binds the cap binding complex; plays roles in small RNA biogenesis (Garcia et al., 2016); facilitates miRNA processing from primary miRNAs to pre-miRNAs; required for siRNA-mediated silencing; regulation of transcription by RNA pol II (Speth et al., 2018) Serrate/Ars2. |
| Hrb87F | **RNABPR; RGEXPPR; DBRR; TMM** | **Heterogeneous nuclear ribonucleoprotein at 87F**, hnRNP-A family RNA-binding protein; involved in gene expression & RNA processing (Lasko, 2000); an essential component of the nucleoplasmic omega speckles; necessary for telomere maintenance; sequence-specific DNA binding (Borah et al., 2009); regulation of alternative mRNA splicing, via spliceosome (Borah et al., 2009; Park et al., 2004); Wnt/Wingless and JNK signaling pathways (Yadav & Tapadia, 2016). |
| glo (CG6946) | **RNABPR; RNABTN; TN; CBPR; CRTR2; RGEXPPR** | **glorund**, mRNA binding (Lasko, 2000); regulation of RNA splicing (GO Reference Genome Project, 2011-); chromosome organization; intracellular mRNA localization involved in pattern specification process; translational repressor, splicing regulator; regulation of translation (Kalifa et al., 2009); hnRNP F/H family RNA-binding protein; RNA Recognition Motive Domain¸ part of a complex containing the hnRNP protein Hrp48 & the splicing factor Half-pint. |
| Saf-B | **DBTR2; POLIITR; RNABPR; RGEXPPT** | **Scaffold attachment factor B**, regulation of mRNA processing; regulation of transcription by RNA pol II; sequence-specific DNA binding (GO Reference Genome Project, 2011-); RNA binding (InterPro Project Members, 2004-); RNA Recognition Motif Domain. |
| SmF | **RNABPR** | **Small ribonucleoprotein particle protein SmF**, RNA binding (GO Reference Genome Project, 2011-). |
| X16 | **POLIITR; RGEXPPT; RNABPR; DBTR2** | **x16 splicing factor**, involved in mRNA splicing and RNA metabolism regulation; regulation of transcription start site selection at RNA pol II promoter; play role in alternative promoter choices, polyadenylation site selection & overall transcript levels; regulation of gene expression (Bradley et al., 2015). |

**Table C. Subdivisions of regulatory factors associated with the Myc target P29/P30 (plotted in Figure 12)**

| Gene name | Pathway | Activity/Function |
|---|---|---|
| Pep (CG6143) | **DBTR2; POLIITR; RNABPR** | **Zinc finger protein on ecdysone puffs**, plays role in the process of early and late gene activation, and in RNA processing, for a defined set of developmentally regulated loci; DNA binding; single-stranded RNA binding (Hamann and Stratling, 1998); single-stranded DNA binding (Amero et al., 1993); Cip1-Interacting Zinc Finger Protein (Zinc Finger C2H2-type: most common DNA-binding motifs found in eukaryotic transcription factors). |
| Srp54 | **RNABPR; RGEXPPR; DBTR2; POLIITR** | **Splicing regulatory protein 54**, regulation of mRNA alternative splicing; mRNA binding; regulation of gene expression; regulation of the transcriptional start site selection at RNA pol II promoter (Bradley et al., 2015); poly-pyrimidine tract binding (Kennedy et al., 1998); RNA binding (InterPro Project Members, 2004-). |
| Smn | **RNABPR; CRR; RGEXPPR** | **Survival motor neuron**, eponymous member of the SMN complex, functions as an assembly chaperone for Sm-class small nuclear ribonucleoproteins; RNA binding; mRNA processing (InterPro Project Members, 2004-); chromatin organization (Lee et al., 2009); protein-RNA-complex assembly (Shpargel et al., 2009); Survival Motor Neuron Complex-GEM4A Variant. |
| CG7564 | **RNABPR** | **Alsin2**, mRNA binding & splice site recognition (InterPro Project Members, 2004-); mRNA splicing via spliceosome (Herold et al., 2009; Mount and Salz, 2000). |
| pum | **RNABTN; RNABPT; RNABPR; TN; RGEXPPT; PSILEN** | **Pumilio, isoform G**, a member of the PUF family of RNA binding proteins (Chen et al., 2008); mRNA regulatory element binding translation repressor activity (Kim et al., 2012); sequence-specific mRNA binding (Arvola et al., 2020); post-transcriptional gene silencing (Miles et al., 2012); nuclear-transcribed mRNA catabolic process (Miles et al., 2012); negative regulation of DNA-templated transcription (Leatherman and Jongens, 2003). |
| CG13928 | **TT; RGEXPPR; TN; RNABTN** | Negative regulation of translation and positive regulation of nuclear-transcribed mRNA poly(A) tail shortening (Khan et al., 2015). |
| SF2 | **DBTR2; RNABPR; RGEXPPR** | **Splicing factor 2**, DNA binding (Lynch and Maniatis, 1996); mRNA binding; regulation of gene expression; regulation of transcriptional start site selection at RNA pol II promoter (Bradley et al., 2015; Lasko, 2000); RNA binding (GO Reference Genome Project, 2011-); regulation of gene expression ((Bradley et al., 2015; Canonical Ser/Arg Rich Splice Factors. |

**Table C. Subdivisions of regulatory factors associated with the Myc target P29/P30 (plotted in Figure 12)**

| Gene name | Pathway | Activity/Function |
|---|---|---|
| Larp4B | **PSILEN; RNABPT; TN; RGEXPPT** | **La-related protein Larp4B**, RNA-binding; inhibition of MYC protein; negative regulation of cell growth & translation (Funakoshi et al., 2018); mRNA 3'-UTR binding (Gene Ontology Curators, 2002-). |
| sti | **CHRSILEN; CRR** | **Sticky**, a member of the AGC family of kinases, functions to regulate both actin-myosin-mediated cytokinesis & epigenetic gene silencing (histone H3-K9 methylation, HP1 localization, & heterochromatin-mediated gene silencing); chromatin organization (Sweeney et al., 2008). |
| gw | **RGEXPPT; PSILEN; RNABPR; RNABTN; TN** | **Gawky**, RNAi pathway (gene silencing pathway) (Chekulaeva et al., 2010; Chekulaeva et al., 2009; Eulalio et al., 2009; Eulalio et al., 2009; Zekri et al., 2009; Eulalio et al., 2008); mRNA catabolic process; miRNA induced silencing complex (RISC) (Behm-Ansmant et al., 2006); negative regulation of gene expression, RNA binding (Sienski et al., 2015); miRNA-mediated gene silencing by inhibition of translation (InterPro Project Members, 2004-). |
| lola | **DBTR2; POLIITR; TF** | **Longitudinals lacking protein, isoforms H/M/V** , involved in Notch signaling, cell death, regulation of retrotransposons & expression of axon & dendrite patterning genes; oogenesis, spermatogenesis, neural wiring, eye development & a variety of behaviors; positive regulation of peptidoglycan recognition protein signaling pathway (Kleino et al., 2005); positive regulators of immune deficiency (Imd) pathway with activity of the NF-κB-like transcription factor Rel in the nucleus; DNA binding (Dinges et al., 2017); DNA binding transcription factor activity (Giniger et al., 1994); C2H2 Zinc Finger Transcription Factors. |
| hang | **DBTR2; POLIITR; TF; RNABTR2** | **Hangover**, a nuclear zinc finger protein, DNA binding transcription factor, pol II-specific (GO Reference Genome Project, 2011-); mRNA binding (Ruppert et al., 2017); C2H2 Zinc Finger Transcription Factor. |
| HnRNP-K | **DBTR2; DBTFB; POLIITR; CRTR2; RNABTN; TN; RGEXPTN** | **Heterogeneous nuclear ribonucleoprotein K**, localizes in nucleus, cytoplasm & mitochondria; involved in gene regulation; post-transcriptional RNA processing; RNA transport; mRNA binding; regulation of transcription by RNA pol II (GO Reference Genome Project, 2011-); RNA binding (InterPro Project Members, 2004-); transcription factor binding; DNA binding; involved in chromatin remodeling, transcription, splicing & translation; docking platform for the integration of signaling cascades (Bomsztyk et al., 2004). |

**Table C. Subdivisions of regulatory factors associated with the Myc target P29/P30 (plotted in Figure 12)**

| Gene name | Pathway | Activity/Function |
|---|---|---|
| net (CG11450) | **DBTR2; TF; POLIITR; DBTFB** | **Net, isoform B**; E-box binding, positive regulation of transcription by RNA pol II (GO Reference Genome Project, 2011-), Basic Helix-Loop-Helix protein, transcriptional repressor; during wing vein formation, expressed in all interveins territories; negative regulation of transcription by RNA pol II (Brentrup et al., 2000); EGFR antagonist; DNA binding transcription factor activity; involved in the regulation of Dpp, Hedgehog, & Wingless signaling pathways (De Celis, 2003); regulation of transcription by RNA pol II (Peyrefitte et al., 2001). |
| CG6418 (DmRH27) | **RNAHPR; RNABPR** | Nuclear RNA binding activity & RNA helicase activity; mRNA splicing (InterPro Project Members, 2004-; Gene Ontology Curators, 2002-); DEAD-BOX RNA Helicases. |
| Mtr4 | **RNAHPR; RNABPR; RGEXPPR** | **Mtr4 helicase**, interferon immune signaling, RNA helicase (GO Reference Genome Project, 2011-) involved in the viral defense response (Molleston et al., 2016); RNA binding; RNA catabolic process (InterPro Project Members, 2004-); SKI2-Like RNA Helicases. |
| san | **CHRM; RRR** | **(separation anxiety)**, mitotic sister chromatid cohesin; couples the processes of cohesion and DNA replication (Ribeiro et al., 2016; Williams et al., 2003). |
| DCG10077(B cDNA:HL079 10) | **RNAHPR; RNABPR** | RNA helicase, RNA binding (GO Reference Genome Project, 2011-); DEAD-Box RNA Helicases. |
| tsu | **POLIITR; RNABPR; RGEXPPR** | **Tsunagi**, RNA binding & processing (InterPro Project Members, 2004-); regulation of MAPK levels by the Exon Junction Complex (EJC) (Ashton-Beaucage & Therrien, 2010); disassembly of EJC by interaction between Mago/Tsunagi with the EJC key regulator Pym in cytoplasm (Ghosh et al., 2014). |
| LSm7 | **RNABPR; RGEXPPR** | **Like Sm 7**, RNA binding activity; nuclear-transcribed mRNA catabolic process (InterPro Project Members, 2004-). |
| Lig | **RNABPT; RGEXPPT** | **Lingerer**, RNA binding protein; regulation of gene expression (Baumgartner et al., 2013). UBA-like superfamily. |
| nonA | **RNABTR2; POLIITR** | **(no on or off transient A)**, mRNA binding (Greenspan and Ferveur, 2000; Lasko, 2000); RNA binding; regulation of transcription (GO Reference Genome Project, 2011-); contains RNA Recognition Motif (RRM) Domain. |

**Table C. Subdivisions of regulatory factors associated with the Myc target P29/P30 (plotted in Figure 12)**

| Gene name | Pathway | Activity/Function |
|---|---|---|
| Mtor (Mgtor) | **CBTR2; CRTR2; POLIITR** | **Megator**, plays role as a transcriptional attenuator of X chromosome gene expression in male X chromosome dosage compensation; histone acetyltransferase binding (Aleman et al., 2021); chromatin DNA binding; chromatin organization; chromatin remodeling; positive regulation of transcription by RNA pol II (Vaquerizas et al., 2010); negative regulation of transcription by RNA pol II (Aleman et al., 2021); Nuclear Pore Complex. |
| Nasp (CG8223) | **CRR; RRR** | **Nuclear autoantigenic sperm protein**, histone chaperone activity (Tirgar et al., 2023); histone binding; CENP-A containing chromatin assembly; DNA replication-dependent chromatin assembly (GO Reference Genome Project, 2011-); Belongs to the NASP family. |
| PolD3 (Pol32) | **DBRR; RRR; DNAPOLD3** | **DNA polymerase delta subunit 3**, non-essential subunit of the multi-subunit DNA polymerases delta and zeta; embryonic DNA replication; DNA repair (Tritto et al., 2015); homologous recombination repair of double-strand breaks (Kane et al., 2012); delta DNA polymerase complex (Ji et al., 2019); zeta DNA polymerase complex (Gene Ontology Curators, 2002-). |
| mEFG1 | **MTTN; MTTNF** | **Mitochondrial translation elongation factor G1**, developmental signal from mitochondria to nucleus for slow proliferation in the case of low mitochondrial energy level (Trivigno and Haerry, 2011); GTP binding; catalysis of mRNAs and tRNAs translocation along the ribosomes via GTP hydrolysis (InterPro Project Members, 2004-). |
| pAbp | **PAB; RNABTN; TN; RGEXPTN** | **Poly(A) binding protein**, poly(A) binding (Mount and Salz, 2000); post-transcriptional regulation of gene expression; mRNA binding (Lasko, 2000); mRNA translation (Herold et al., 2009). |
| Polr2H | **RPIIB8; DBTR1; POLITR; DBTR2; POLIITR; DBTR3; POLIIITR** | **RNA polymerase II, I and III subunit H,** RNA pol I & RNA pol III activity (GO Reference Genome Project, 2011-); RNA pol II activity (GO Reference Genome Project, 2011-; Aoyagi and Wassarman, 2000); transcription by RNA pol I, RNA pol II and RNA pol III; part of RNA polymerase I complex; RNA polymerase II, core complex; and RNA polymerase III complex. |
| mbo (Nup88) | **CBTR2** | **members only**, chromatin binding (Capelson et al., 2010); involved in immune response transduction by mediating the nuclear translocation of Mad; Toll-NF-κB, antimicrobial humoral response (Uv et al., 2000). |

**Table C. Subdivisions of regulatory factors associated with the Myc target P29/P30 (plotted in Figure 12)**

| Gene name | Pathway | Activity/Function |
|---|---|---|
| Dp1 | **TN; RGEXPTN; CBTR2; DBTR2** | **Dodeca-satellite-binding protein 1**, translation enhancer; ssDNA binding (Cortes and Azorin, 2000; Cortes et al., 1999); mRNA 3'-UTR binding (Nelson et al., 2007); satellite DNA binding (Cortes and Azorin, 2000); chromatin condensation (Huertas et al., 2004); chromatin formation (Wang et al., 2005). |
| Caf1-55 | **POLIITR; CBTR2; DBTR2; CRTR2; HF; HDAC; RRR; DBRR** | **Chromatin assembly factor 1, p55 subunit**, a subunit of the MuvB core complex, MuvB core binds to the oncoprotein Myb & Rbf-E2f2-Dp tumor suppressor complex, thereby controlling the expression of many genes, including critical regulators of the cell cycle; histone binding (Nowak et al., 2011); histone deacetylase binding (Tyler et al., 1996); nucleosome binding (Nekrasov et al., 2005); chromatin remodeling (Mizuguchi et al., 2001); chromatin organization (Martinez-Balbas et al., 1998; Bulger et al., 1995); DNA replication-dependent chromatin assembly (Tyler et al., 1999; Ito et al., 1997; Tyler et al., 1996); positive regulation of transcription by RNA pol II (Mizuguchi et al., 2001); negative regulation of transcription by RNA pol II; transcription *cis*-regulatory region binding (Yao et al., 2018); nucleosome organization (Hamiche et al., 1999); Nucleosome Remodeling Factor (NURF); Polycomb Repressive Complex 2, PCL Variant; Chromatin Assembly Factor; Nucleosome Remodeling Deacetylase (NURD) Complex; Polycomb Repressive Complex 2, JARID2-JING Variant. |
| Rga | **TN; RNAHTN; RNAHPR; RNAHPT; RGEXPTN; PSILEN** | **Regena, isoform C**, bulk mRNA degradation, miRNA-mediated repression, translational repression (Bawankar et al., 2013); component of CCR4-NOT complex (one of the major cellular mRNA deadenylases) (Frolov et al., 1998); mRNA catabolic process (Bawankar et al., 2013). |
| Rat1 | **RNABPR; RGEXPPR; POLIITR; TT** | **Rat1 5'-3' exoribonuclease**, nuclear protein, 5'-3'-exonuclease activity (Chen et al., 2016); termination of RNA pol II transcription (Gene Ontology Curators, 2002-); RNA binding; nuclear-transcribed mRNA catabolic process (GO Reference Genome Project, 2011-); 5'-3'-Exoribonucleases. |
| Taf5 | **POLIITR; DBTR2; COACT; TPI; TI** | **TBP-associated factor 5**, part of the multisubunit basal transcription factor TFIID & important for its assembly or stability; RNA pol II general transcription initiation factor activity; transcription factor TFIID complex (Wright et al., 2006; Hansen and Tjian, 1995). |
