## Supplemental Tables for "Targeting Regulatory Factors Associated with the *Drosophila Myc cis*-Elements by Reporter Expression, Gel Shift Assay, and Mass Spectrometric Protein Identification": Table D.pdf

**Table D. Subdivisions of regulatory factors associated with the Myc target P31/P32 (plotted in Figure 12)**

| Gene name | Pathway | Activity-Function |
|---|---|---|
| ncd | **CHRM** | **non-claret disjunctional**, a minus-end-directed kinesin microtubule motor protein and the sole member of the kinesin-14 motor family; chromosome segregation (Hallen et al., 2008); distributive segregation (Whyte et al., 1993); mRNA transport (Fahmy et al., 2014). |
| Sirt4 | **HDAC; MTTR** | **Sirtuin4,** Sole mitochondrial Sirtuin in *Drosophila*, NAD-dependent protein deacylase; catalyzes the NAD-dependent hydrolysis of acyl groups from lysine residues (Feller et al., 2015); transcriptional activation of mitochondrial biogenesis, mitochondrial regulator of life span & metabolism, cellular response to starvation, Sirt4 knockout causes short lifespan, increased sensitivity to starvation, and decreased fertility and activity (Wood et al., 2018). |
| tral | **RNAHPT; RNABPT; RGEXPPT** | **Trailer hitch, isoform D**, DEAD/H-Box RNA helicase binding (Barbee et al., 2006; Wilhelm et al., 2005); mRNA binding (GO Reference Genome Project, 2011-); RNA binding (InterPro Project Members, 2004-); piRNA pathway; retrotransposon silencing (Liu et al., 2011); required for oocyte dorsoventral patterning via actin and microtubule cytoskeleton organization. |
| Elp4 (CG6907) | **POLIITR; TN; RRR; RNABTN** | **Elongator complex protein 4**, establishment of mitotic spindle asymmetry; regulation of translation through targeted transfer-RNA (tRNA) modification (Planelles-Herrero et al., 2022); phosphorylase kinase regulator activity; regulation of transcription by RNA pol II (Gene Ontology Curators, 2002-); Elongator Complex. |
| CG7406 | **U** | Uncharacterized; Domain of Unknown Function; DUF4766 |
| CG15506 | **U** | Uncharacterized; Unknown. |
| e(y)3 (SAYP) | **CBTR2; CRTR2; TPI; POLIITR; COREP; COACT** | **Enhancer of yellow 3, isoform D**, nuclear protein, plays role in gene activation in euchromatin as a component of SWI/SNF chromatin remodeling complex (Chalkley et al., 2008) & transcription factor TFIID complex; gene silencing in pericentric heterochromatin; chromatin binding; transcription coactivator and corepressor activity; negative regulation of transcription; positive regulation of transcription; coactivator of JAK/STAT pathway (Shidlovskii et al., 2005); required for embryogenesis and oogenesis (Vorobyeva et al., 2009); Polybromo Containing Brahma Associated Proteins Complex. |

**Table D. Subdivisions of regulatory factors associated with the Myc target P31/P32 (plotted in Figure 12)**

| Gene name | Pathway | Activity-Function |
|---|---|---|
| Gug | **COREP; DBTFB; POLIITR** | **Grunge, isoform C** (related to human Atrophin-like proteins & is the only Atrophin-like protein in *Drosophila*), a nuclear repressor binding protein, responds to EGFR signaling to control cell behavior for normal developmental patterning; DNA binding transcription factor binding (Zhang et al., 2002); transcription corepressor activity (Wang et al., 2006; Zhang et al., 2002); activation of Frizzled, Notch, Dpp, & JNK signaling pathways (Zheng et al., 1995; Artavanis-Tsakonas et al., 1999; Zeitlinger & Bohmann, 1999); eye development; negative regulation of Epidermal Growth Factor Receptor (EGFR) signaling pathway (Charroux et al., 2006); negative regulation of Hedgehog signaling pathway (Zhang et al., 2013); SANT-MYB Domain Transcription Factors. |
| Map60 (CP-60) | **POLIITR; DBTR2** | **Microtubule-associated protein**, microtubule binding protein; localized to the centrosome in a cell cycle-dependent manner (Kellogg et al., 1995); a component of a Centrosomal complex; regulation of transcription by RNA pol II (GO Reference Genome Project, 2011-); MADF Domain. |
| cutlet | **CHRM; DBRR** | **Chromosome cohesion factor**, (chromosome transmission fidelity protein 18 homolog) cell cycle; DNA clamp loader activity; positive regulation of DNA-directed DNA polymerase activity (Gene Ontology Curators, 2002-); CTF18-replication factor C (CTF18-RFC); DNA binding (GO Reference Genome Project, 2011-); RFC Complex ATPases |
| pAbp | **PAB; RNABTN; TN; RGEXPTN** | **Poly(A) binding protein**, Poly(A) binding (Mount and Salz, 2000); post-transcriptional regulation; mRNA binding (Lasko, 2000); mRNA translation (Herold et al., 2009). |
| Cbp80 | **RNABPR; RGEXPPT; PSILEN** | **cap binding protein 80**, component of the cap-binding complex (CBC), which binds cotranscriptionally to the 5'-cap of pre-mRNAs and is involved in various processes such as pre-mRNA splicing and RNA-mediated gene silencing (RNAi); RNA binding; RNA cap binding; mRNA export from nucleus (InterPro Project Members, 2004-); mRNA binding (GO Reference Genome Project, 2011-); primary-miRNA processing; regulatory ncRNA-mediated post-transcriptional gene silencing; RNA-mediated gene silencing (RNAi); miRNA-mediated RNA interference via its interaction with Ars2, required for primary microRNAs (miRNAs) processing (Sabin et al., 2009). |

**Table D. Subdivisions of regulatory factors associated with the Myc target P31/P32 (plotted in Figure 12)**

| Gene name | Pathway | Activity-Function |
|---|---|---|
| LKRSDH | **COREP; POLIITR** | **Lysine ketoglutarate reductase/saccharopine dehydrogenase**, a bifunctional enzyme in the lysine degradation pathway, Independently of its enzymatic activity, also a transcriptional corepressor in ecdysone signaling; histone binding; nuclear receptor binding; negative regulation of transcription by RNA pol II (Cakouros et al., 2008); CH-NH Oxidoreductases, NAD Or NADP As Acceptor. |
| vig2 | **CHRSILEN; CRTR2; HF; RNABPT** | **Vig2**, involved in heterochromatin organization (formation), histone H3-K9 methylation & chromatin silencing regulation (Gracheva et al., 2009); Hyaluronan/mRNA-binding protein, involved in nuclear functions such as the remodeling of chromatin & the regulation of transcription (Nery et al., 2006; Nery et al., 2004). |
| Taf6 | **POLIITR; TF; TI; COACT; TPI** | **TBP-associated factor 6**; part of the multisubunit basal transcription factor TFIID; pol II general transcription initiation factor (Wright et al., 2006; Hansen and Tjian, 1995); transcription co-activator activity (GO Reference Genome Project, 2011-); transcription by RNA pol II (Hansen and Tjian, 1995); Transcription Factor TFIID. |
| mael | **TF; DBTR2; REP; POLIITR; HF; RNABPT; RGEXPPT** | **Maelstrom**, involved both in the piRNA & miRNA metabolic processes (Findley et al., 2003); a component of the meiotic nuage, plays a central role during oogenesis by repressing transposable elements and preventing their mobilization, which is essential for the germline integrity. Repression of transposable elements is mediated via the piRNA metabolic process; sequence-specific DNA binding; transcription *cis*-regulatory region binding; negative regulation of DNA-templated transcription (Pek et al., 2009); intracellular mRNA localization; signaling oocyte polarity; posterior localization of *grk* mRNA (EGFR ligand) (Clegg et al., 1997); piRNA-mediated retrotransposon silencing by heterochromatin formation (Sienski et al., 2012); regulatory nc-RNA-mediated gene silencing (Lim and Kai, 2007); RNA binding (Genzor et al., 2015); High Mobility Group Box Domain Superfamily; Maelstrom Domain. |
| CG6683 | **POLIITR** | Involved in regulation of transcription by RNA polymerase II. Predicted to be part of transcription regulator complex (GO Reference Genome Project, 2011-); active in nucleus; MAD-BESS domain Transcription Regulators. |

**Table D. Subdivisions of regulatory factors associated with the Myc target P31/P32 (plotted in Figure 12)**

| Gene name | Pathway | Activity-Function |
|---|---|---|
| row | **CBTR2; DBTR2; POLIITR; TF** | **Relative of woc**, a zinc-finger protein involved in transcription regulation, required for the (HP1c) to bind chromatin (Font-Burgada et al., 2008); sequence-specific DNA binding, pol II transcription (GO Reference Genome Project, 2011-); regulation of transcription by RNA pol II (Abel et al., 2009; Font-Burgada et al., 2008); Zinc Finger C2H2 TYPE. |
| Polr2C | **RPIIB3; DBTR2; POLIITR** | **RNA polymerase II subunit C, DNA-directed RNA polymerase** (associated with TATA, DPE & Positive Control), protein dimerization activity; DNA binding activity; DNA-templated transcription (InterPro Project Members, 2004-); DNA-directed 5'-3' RNA pol II activity (Aoyagi and Wassarman, 2000); involved in cellular response to heat (Yao et al., 2007); located in cytoplasm; nucleus; & polytene chromosome puff. Part of RNA pol II, core complex; RNA pol Catalytic Subunit. |
| Polr2A (RpII215) | **RPIIB1; DBTR2; POLIITR; DBRR; TI; RRR** | **RNA polymerase II subunit A**, DNA-directed 5'-3' polymerase activity (Brickey and Greenleaf, 1995; Zehring et al., 1988); RNA pol II, core complex (Gu et al., 2002; Greenleaf, 1983); Largest and catalytic component of RNA pol II which synthesizes mRNA precursors and many functional non-coding RNAs. Forms the polymerase active center together with the second largest subunit. pol II is the central component of the basal RNA pol II transcription machinery (RNA pol II subunit RPB1). |
| CG2962 | **POLIITR; COACT** | Transcription coactivator activity; positive regulation of transcription by RNA pol II (GO Reference Genome Project, 2011-). |
| Rfc37 (CG8142) | **RRR; DBRR** | **Replication factor C37**, ATP binding activity; ATP hydrolysis activity; DNA binding activity (InterPro Project Members, 2004-); predicted to contribute to DNA clamp loader activity (GO Reference Genome Project, 2011-); involved in DNA repair and DNA-templated DNA replication; part of Elg1 RFC-like complex. |
| La (CG10922) | **TT; POLIIITR; DBTR3; RNABTR3** | **La autoantigen-like**, involved in transcription termination by RNA pol III; binds RNA & DNA. Binds to precursors of RNA pol III transcripts; plays role during fly development. |
| Nup93-1 | **CBPT; RGEXPPT** | **Nucleoporin 93kD-1**, chromatin binding; negative regulation of gene expression, epigenetic; chromatin DNA binding (Gozalo et al., 2020). |

**Table D. Subdivisions of regulatory factors associated with the Myc target P31/P32 (plotted in Figure 12)**

| Gene name | Pathway | Activity-Function |
|---|---|---|
| Nup98-96 | **CBTR2; RNABTR2; TI; POLIITR; TMM** | **Nucleoporin 98-96kD**, precursor of Nup98 and Nup96 proteins, integral parts of the nuclear pore; male germline cell differentiation; Chromatin DNA binding (Ilyin et al., 2017; Pascual-Garcia et al., 2017; Kalverda et al., 2010); promoter-enhancer loop anchoring (Pascual-Garcia et al., 2017); promoter-specific chromatin binding (Pascual-Garcia et al., 2017; Pascual-Garcia et al., 2014); RNA binding (GO Reference Genome Project, 2011-); heat shock-mediated polytene chromosome puffing (Kalverda et al., 2010); positive regulation of transcription by RNA pol II (Panda et al., 2014; Pascual-Garcia et al., 2014; Capelson et al., 2010; Kalverda et al., 2010); positive regulation of transcription initiation by RNA pol II (Capelson et al., 2010); post-transcriptional tethering of RNA pol II gene DNA to nuclear periphery; telomere tethering to nuclear periphery (GO Reference Genome Project, 2011-); Nuclear Pore Complex. |
| Chi | **COACT; POLIITR** | **Chip, isoform B**, transcription co-activator, LIM domain-binding protein 2; LIM domain-binding protein/SEUSS; Imaginal disc-derived wing morphogenesis (Capelson et al., 2010) / leg development & axon guidance (van Meyel et al., 2000); positive regulation of transcription by RNA pol II (Werner et al., 2017; van Meyel et al., 2000). |
| Isha (cg4266) | **POLIITR; TT; CBTR2; RNABTR2; INSUL** | **Insulator su(Hw) mRNA adaptor**, chromatin binding activity; mRNA binding activity (Bag et al., 2022; Lasko, 2000); negative regulation of transcription by RNA pol II; required for gypsy insulator function (Bag et al., 2022). |
| lost | **RNABPR; RGEXPPR** | (**lost**), interacts with the RNA-binding protein rumpelstiltskin (*rump*) for posterior localization of mRNAs by diffusion/entrapment during late stages of oogenesis; mRNA splicing, via spliceosome (Herold et al., 2009); localization to various RNP complexes with a broad role in RNA metabolism. |
| Hsc70-4 | **RGEXPPT; PSILEN** | **Heat shock protein 70 cognate 4**, regulatory ncRNA-mediated post-transcriptional gene silencing (Dorner et al., 2006). |
| koi | **RRR; DBRR** | **Klaroid**, involved in double-strand break repair via homologous recombination (Ryu et al., 2015); nuclear migration (Kracklauer et al., 2007); nucleolus organization (Tan et al., 2018; Elhanany-Tamir et al., 2012). |

**Table D. Subdivisions of regulatory factors associated with the Myc target P31/P32 (plotted in Figure 12)**

| Gene name | Pathway | Activity-Function |
|---|---|---|
| PolD1 | **RRR; DBRR** | **DNA polymerase delta subunit 1 (**DNA-directed DNA polymerase**)**, catalytic component of the DNA polymerase delta complex, plays a crucial role in high fidelity genome replication, including lagging strand synthesis (Aoyagi et al., 1994); DNA recombination & repair; DNA polymerase & 3'-5'-exonuclease activities (Aoyagi et al., 1994; Chiang et al., 1993; Peck et al., 1992); required at the nucleus of rapidly dividing embryonic cells to activate genome replication during the earliest cell (Ji et al., 2019); DNA polymerase activity (InterPro Project Members, 2004-); base-excision repair, gap filling (Gene Ontology Curators, 2002-). |
| TFAM | **MTTF; TFAM; DBMTTR; MTTR; MTRR; DBMTRR** | **Mitochondrial transcription factor A**, essential for mtDNA transcription & replication; binds to mtDNA nonspecifically; plays a role in mtDNA maintenance by packaging mtDNA; mitochondrial promoter sequence-specific DNA binding (Gene Ontology Curators, 2002-);transcription *cis*-regulatory region binding (GO Reference Genome Project, 2011-); High Mobility Group Box Transcription Factors. |
| net (CG11450) | **DBTR2; TF; POLIITR; DBTFB** | **Net, isoform B**; E-box binding, positive regulation of transcription by RNA pol II (GO Reference Genome Project, 2011-), Basic Helix-Loop-Helix protein, transcriptional repressor; during wing vein formation, expressed in all interveins territories; negative regulation of transcription by RNA pol II (Brentrup et al., 2000); EGFR antagonist; DNA binding transcription factor activity; Dpp, Hedgehog & Wingless signaling (De Celis, 2003); regulation of transcription by RNA pol II (Peyrefitte et al., 2001). |
| RnrL | **RRR** | **Ribonucleoside diphosphate reductase large subunit**, preparation of precursors necessary for DNA synthesis; catalysis of the biosynthesis of deoxyribonucleotides from the corresponding ribonucleotides (Gene Ontology Curators, 2002-); DNA replication DNA helicase activity; single-stranded DNA binding; duplex DNA binding. |
| CG9411 | **U** | Uncharacterized, unknown. |
| dsx-c73A | **TREX** | **doublesex cognate 73A**, a secreted cuticle protein present in the epidermis, salivary duct and trachea; expressed under control of (ovo); structural constituent of nuclear pores; mRNA export from nucleus; regulation of mitotic cell cycle (GO Reference Genome Project, 2011-); Unclassified Cuticle Proteins. |

**Table D. Subdivisions of regulatory factors associated with the Myc target P31/P32 (plotted in Figure 12)**

| Gene name | Pathway | Activity-Function |
|---|---|---|
| Dbp80 | **RNAHTR2; RNABTR2; TREX** | Dead box protein 80, poly(A) + mRNA export from nucleus; RNA binding; RNA helicase activity (GO Reference Genome Project, 2011-); DEAD-Box RNA Helicases |
| l(2)10685 | **RNABPT; RGEXPPT** | **Lethal (2) 10685**, methyltransferase activity (GO Reference Genome Project, 2011-); S-adenosylmethionine-dependent methyltransferase activity (InterPro Project Members, 2004-); orthologous to human NSUN4 (NOP2/Sun RNA methyltransferase 4); Unclassified Methyltransferases. |
| CG6000 | **U** | Orthologous to human TSTD1 (thiosulfate sulfurtransferase like domain containing 1), Rhodanese-like domain. |
| Pym | **RNABPR; RGEXPPR; TREX; TN** | **Partner of Y14 and Mago**, RNA binding (Bono et al., 2004); regulator of the exon junction complex (EJC), a multiprotein complex, associates upstream of the exon-exon junction on mRNAs, serves as a positional landmark for the intron exon structure of genes (Ghosh et al., 2014); directs post-transcriptional processes in the cytoplasm such as mRNA export, nonsense-mediated mRNA decay (NMD) or translation; acts as an EJC disassembly factor by disrupting mature EJC from spliced mRNAs (GO Reference Genome Project, 2011-); required for normal localization of osk mRNA to the posterior pole of the developing oocyte; no interaction with the small ribosomal unit or components of the translation initiation complex; not involved in cap-dependent translation regulation. |
| SF2 | **DBTR2; RNABPR; RGEXPPR** | **Splicing factor 2**, DNA binding (Lynch and Maniatis, 1996); mRNA binding; regulation of gene expression; ; regulation of transcriptional start site selection at RNA pol II promoter (Bradley et al., 2015; Lasko, 2000); RNA binding (GO Reference Genome Project, 2011-); regulation of gene expression (Bradley et al., 2015); Canonical Ser/Arg Rich Splice Factors. |
| asun | **POLIITR; INTCOM** | **Protein asunder**, Mitotic cell cycle (Lee et al., 2005); Component of the Integrator complex involved in the transcription of small nuclear RNAs (snRNA) by RNA pol II-transcribed snRNAs & their 3'-box-dependent processing (Chen et al., 2012); Cell Cycle Regulated Mat89Bb. |
| LSm7 | **RNABPR; RGEXPPR** | **Like Sm 7**, RNA binding activity; nuclear-transcribed mRNA catabolic process (InterPro Project Members, 2004-). |

**Table D. Subdivisions of regulatory factors associated with the Myc target P31/P32 (plotted in Figure 12)**

| Gene name | Pathway | Activity-Function |
|---|---|---|
| Dmac2 (CG4042) | **U** | Mitochondrial respiratory chain complex I assembly factor (Gene Ontology Curators, 2002-); cellular component: mitochondrion (Murari et al., 2021); orthologous to human DMAC2 (distal membrane arm assembly component 2). |
| PolA1 | **RRR; DNAPOLA1; DBRR** | **DNA polymerase alpha subunit 1**, DNA-directed polymerase activity; Initial synthesis on the leading strand and on each Okazaki fragment of the lagging strand (Peck et al., 1992; Kuroda et al., 1990; Cotterill et al., 1987; Kaguni et al., 1984; Kaguni et al., 1983; Villani et al., 1980). |
| PolD2 (Pol31) | **RRR; DBRR** | **DNA polymerase delta subunit 2**, Accessory component of both the DNA polymerase delta complex and possibly the DNA polymerase zeta complex; as a component of the delta complex, participates in high fidelity genome replication, including lagging strand synthesis, DNA recombination & repair; promotes the function of the DNA pol-delta complex accessory subunit PolD3 in both embryonic & postembryonic somatic cells; mitotic DNA templated DNA replication (Ji et al., 2019); DNA binding; DNA replication (InterPro Project Members, 2004-); DNA strand elongation involved in DNA replication (GO Reference Genome Project, 2011-). |
| Prim2 | **DBRR; RRR; PRIM2** | **DNA primase large subunit, PRIM2**, synthesis of short RNA-DNA primers on the lagging strand during DNA synthesis (Kuroda et al., 1990; Cotterill et al., 1987; Kaguni et al., 1983); compound eye morphogenesis (Chen et al., 2000); alpha DNA polymerase: primase complex (Kuroda et al., 1990; Cotterill et al., 1987; Kaguni et al., 1984; Kaguni et al., 1983; Villani et al., 1980). |
| mEFG1 | **MTTN; MTTNF** | **Mitochondrial translation elongation factor G1**, developmental signal from mitochondria to nucleus for slow proliferation in the case of low mitochondrial energy level (Trivigno and Haerry, 2011); GTP binding; catalysis of mRNAs and tRNAs translocation along the ribosomes via GTP hydrolysis (InterPro Project Members, 2004-). |
| Rga | **TN; RNAHTN; RNAHPR; RNAHPT; RGEXPPT; PSILEN** | **Regena, isoform C**, bulk mRNA degradation, miRNA-mediated repression, translational repression (Bawankar et al., 2013); Component of CCR4-NOT Complex (one of the major cellular mRNA deadenylases) (Frolov et al., 1998); mRNA catabolic process (Bawankar et al., 2013). |

**Table D. Subdivisions of regulatory factors associated with the Myc target P31/P32 (plotted in Figure 12)**

| **Gene name** | **Pathway** | **Activity-Function** |
|---|---|---|
| RanBP3 | **CHRM** | **chromosome region maintenance 1 (CRM1)**, cofactor involved in the export of proteins from the nucleus; negatively regulates Wnt signaling (Hendriksen et al., 2005). |
| Orc6 | **RRR; DBRR** | **Origin recognition complex subunit 6**, a subunit of the origin recognition complex (ORC), which is essential for the initiation of DNA replication in eukaryotic cells (Chesnokov et al., 1999); mitotic cell cycle (Balasov et al., 2009). |
| abs | **RNAHPR; RNABPR; RGEXPPR** | **Abstrakt**, DEAD/DEAH-Box RNA helicase (Lasko, 2000), RNA binding (GO Reference Genome Project, 2011-); RNA splicing, via spliceosome (Herold et al., 2009); regulates cell polarity in oocytes & embryos; downregulation of Notch signaling in asymmetric cell division in ganglion mother cell (GMC2-4a) in collaboration with Inscuteable (*Insc*) (Irion et al., 2004). |
| Dp1 | **TN; RGEXPTN; CBTR2; DBTR2** | **Dodeca-satellite-binding protein 1**, translation enhancer; ssDND binding (Cortes and Azorin, 2000; Cortes et al., 1999); mRNA 3'-UTR binding (Nelson et al., 2007); satellite DNA binding (Cortes and Azorin, 2000); chromatin condensation (*Huertas et al., 2004*); chromatin formation (Wang et al., 2005). |
| CG6418 (DmRH27) | **RNAHPR; RNABPR** | Nuclear RNA binding activity; RNA helicase activity; mRNA splicing (InterPro Project Members, 2004-; Gene Ontology Curators, 2002-); DEAD-BOX RNA HELICASES. |
| Capr | **RNABPT; TN; RGEXPPT** | **Caprin**, a cytoplasmic RNA granules protein (neuronal granules & stress granules); RNA-binding; regulation of embryonic mitotic cell cycle; cellularization; regulation of translation; ribonucleoprotein complex (Papoulas et al., 2010); Caprin-1 Dimerization Domain. |
