## Supplemental Tables for "Targeting Regulatory Factors Associated with the *Drosophila Myc cis*-Elements by Reporter Expression, Gel Shift Assay, and Mass Spectrometric Protein Identification": Table E.pdf

**Table E. Subdivisions of regulatory factors associated with the Myc target P35/P36 (plotted in Figure 12)**

| Gene name | Pathway | Activity-Function |
|---|---|---|
| Rpt4 | **POLITR; DBTR1** | **Regulatory particle triple-A ATPase 4**, 19S proteasomal ATPase, a component of the 26S proteosome complex, nucleolar protein & regulator of rRNA transcription; physical interaction with the tumor suppressor protein “Birt Hogg Dube” (BHD), RNA pol I transcription, regulatory region sequence-specific DNA binding (association of Rpt4 with the rDNA locus) (GO Reference Genome Project, 2011-); Notch-mediated follicle cell differentiation and cell cycle switches, insulin-PI3K pathway (Jia et al., 2015). |
| LKRSDH | **COREP; POLIITR** | **Lysine ketoglutarate reductase/saccharopine dehydrogenase**, a bifunctional enzyme in the lysine degradation pathway, Independently of its enzymatic activity, also a transcriptional corepressor in ecdysone signaling; histone binding; nuclear receptor binding; negative regulation of transcription by RNA pol II (Cakouros et al., 2008); CH-NH Oxidoreductases, NAD Or NADP As Acceptor. |
| Cdk5 | **POLIITR; RRR** | **Cyclin-dependent kinase 5**, regulation of transcription involved in G1/S transition of mitotic cell cycle (GO Reference Genome Project, 2011-); cyclin-dependent protein, serine/threonine kinase activity (Connell-Crowley et al., 2000); proapoptotic signaling by CDK5 & MEKK1 (*Ryoo HD., 2018-cc*); centrosome localization (Jauffred et al., 2013); protein kinase 5 complex (Connell-Crowley et al., 2007). |
| barr | **CBRR; CRR; RRR** | **Barren**, chromatin binding protein, chromatin condensation, DNA topoisomerase binding (Lupo et al., 2001) regulates Malpighian tubule development & epithelial morphogenesis; Tube development (Liu et al., 1999): Hedgehog, completion of bud evagination (Hoch and Pankratz 1996), Wingless is required for cell division & morphogenesis in the tubules (Skaer and Martinez Arias 1992; Harbecke and Lengyel 1995), Notch receptor is required to define the single tip cell at the end of each tubule (Hoch et al. 1994), which leads out the elongation of the tubule (Skaer, 1989), EGFR (EGF-like) ligand required for the proliferation of the distal cells of the tubule (Baumann and Skaer 1993; Kerber et al. 1998); Dpp expressed in foregut/hindgut, required for morphogenesis of both structures (Pankratz and Hoch 1995; Hoch and Pankratz 1996); Cell Cycle interphase (Lupo et al., 2001). |

**Table E. Subdivisions of regulatory factors associated with the Myc target P35/P36 (plotted in Figure 12)**

| Gene name | Pathway | Activity-Function |
|---|---|---|
| asun | **POLIITR; INTCOM** | **Protein asunder**, Mitotic cell cycle (Lee et al., 2005); Component of the Integrator complex involved in the transcription of small nuclear RNAs (snRNA) by RNA polymerase II-transcribed snRNAs & their 3'-box-dependent processing (Chen et al., 2012). |
| pum | **RNABTN; RNABPT; RNABPR; TN; RGEXPPT; PSILEN** | **Pumilio, isoform G**, a founding member of the PUF family of RNA binding proteins (Chen et al., 2008); mRNA regulatory element binding translation repressor activity (Kim et al., 2012); sequence-specific mRNA binding (Arvola et al., 2020); post-transcriptional gene silencing (Miles et al., 2012); nuclear-transcribed mRNA catabolic process (Miles et al., 2012); negative regulation of DNA-templated transcription (Leatherman and Jongens, 2003). |
| Pop2 | **PSILEN; RGEXPPT** | **poly(A)-specific ribonuclease**; involved in translation inhibition (Ruscica et al., 2019; Braun et al., 2011); miRNA-mediated mRNA degradation (Braun et al., 2011); poly(A)-specific ribonuclease activity (Temme et al., 2004); nuclear-transcribed mRNA poly(A) tail shortening (Temme et al., 2010; Bönisch et al., 2007). |
| Mi-2 | **CBTR2; CRTR2; DBTR2; POLIITR; CHRM** | **Mi-2**, a nuclear ATP-dependent nucleosome (chromatin) remodeler activity (Kunert et al., 2009; Murawska et al., 2008); chromatin binding (Kunert et al., 2009); required for repression of cell type-specific genes; DNA binding (InterPro Project Members, 2004-); chromosome condensation (Nikalayevich and Ohkura, 2015); regulation of transcription by RNA pol II (Li et al., 2010); Notch signaling (Zacharioudaki E, Falo Sanjuan J, Bray S., Elife. 2019); wingless, ecdysone signaling (Kon &Nusse; 2005); Nucleosome Remodeling Deacetylase Activity; SNF2-Like Chromatin Remodelers. |
| fon (CG1582) | **RNABTN; RNAHTN; TN** | **Fondue, isoform B**, RNA helicase activity; translation initiation activity (Linsalata et al., 2019; Lasko, 2000); RNA binding activity (GO Reference Genome Project, 2011-); DEAH-Box RNA Helicases. |
| obe | **DNAHTR2; RGEXPPR** | **Obelus**, a Ski2 helicase regulates alternative mRNA splicing of crumb (*crb*); 3'-5' DNA helicase activity (GO Reference Genome Project, 2011-); regulation of alternative mRNA splicing, via spliceosome (Vichas et al., 2015); Wingless signaling alters the levels, subcellular distribution and dynamics of Armadillo and E-cadherin in third instar larval wing imaginal discs; (Somorjai&Martinez-Arias, 2008). |

**Table E. Subdivisions of regulatory factors associated with the Myc target P35/P36 (plotted in Figure 12)**

| Gene name | Pathway | Activity-Function |
|---|---|---|
| aub | **RGEXPPT; RNABPT** | **Aubergine**, piRNA binding (Huang et al., 2021; Webster et al., 2015; Nagao et al., 2010); global gene silencing by mRNA cleavage (Kennerdell et al., 2002) ncRNA-mediated post-transcriptional gene silencing (Tomari et al., 2004); RNA-mediated gene silencing (Bozzetti et al., 2015); JAK-STAT signaling pathway controls host defense in the gut by regulating stem cell proliferation & epithelial cell homeostasis (Cronin et al., 2009); RNA binding (Brennecke et al., 2007). |
| msi | **RNABTN; TN; PSILEN** | M**usashi** RNA binding protein, binds to the 3' UTR region of target mRNAs; Negatively regulates the Hypoxia Inducible Factor (HIF) pathway, contributes to cell fate determination, as well as cellular response to normoxic/hypoxic conditions; negative regulation of translation, asymmetric cell fate determination by Notch signaling; mRNA regulatory element binding translation repressor activity (Bardin et al., 2004). |
| Sirt4 | **HDAC; MTTR** | **Sirtuin4**, Sole mitochondrial sirtuin in *Drosophila*, NAD-dependent protein deacylase. Catalyzes the NAD-dependent hydrolysis of acyl groups from lysine residues (Feller et al., 2015); Transcriptional activation of mitochondrial biogenesis, mitochondrial regulator of life span & metabolism, cellular response to starvation, Sirt4 knockout causes short lifespan, increased sensitivity to starvation, decreased fertility & activity (Wood et al., 2018). |
| smg | **POLIITR; RNABTN; TN; PSILEN; RGEXPTN** | **Smaug**, founding member of the SMAUG family of sequence-specific RNA binding proteins; Translation repressor activity (Dahanukar et al., 1999); mRNA regulatory element binding repressor activity (Dean et al., 2002); positive regulation of nuclear-transcribed mRNA poly(A) tail shortening (Chartier et al., 2015); regulation of DNA-templated transcription; regulation of mRNA stability (InterPro Project Members, 2004-); mRNA 3'-UTR binding (Kim and Bowie, 2004; Dean et al., 2002; Johnstone and Lasko, 2001); cell cycle (Tadros et al., 2007). |
| tral | **RNAHPT; RNABPT; RGEXPPT** | **Trailer hitch, isoform D**, DEAD/H-Box RNA helicase binding (Barbee et al., 2006; Wilhelm et al., 2005); mRNA binding; (GO Reference Genome Project, 2011-); RNA binding (InterPro Project Members, 2004-); piRNA pathway; ); retrotransposon silencing (Liu et al., 2011); required for oocyte dorsoventral patterning via actin and microtubule cytoskeleton organization. |

**Table E. Subdivisions of regulatory factors associated with the Myc target P35/P36 (plotted in Figure 12)**

| Gene name | Pathway | Activity-Function |
|---|---|---|
| mod(mdg4) | **DBTR2; CHRM; POLIITR; ENBLOC; CBTR2** | **modifier of mdg4, isoform AD**, a nuclear protein, specifically interacts with various DNA binding proteins; involved in meiotic chromosome condensation (Matsui et al., 2011); male meiotic chromosome segregation (Soltani-Bejnood et al., 2007); regulation of transcription & enhancer blocking; chromatin binding (Bag et al., 2019; Gerasimova et al., 1995) DNA binding (Ogiyama et al., 2018; Bonchuk et al., 2011); POZ (Pox virus and Zinc finger) domain binding (Harvey et al., 1997); regulation of transcription by RNA pol II (GO Reference Genome Project, 2011-); SKP1/BTB/POZ Domain Superfamily. |
| CG1943 | **U** | Expressed in embryonic brain & organism; used to study congenital heart disease; orthologous to human JPT1 (Jupiter microtubule associated homolog; wing disc dorsal/ventral pattern formation (Bejarano et al., 2008). |
| CG6664 | **CHRM** | Establishment of meiotic spindle orientation; spindle pole, condensed chromosomes (Gene Ontology Curators, 2002-). |
| Tailor | **POLIIITR; RGEXPTR3; RNABTR3; RNABPR** | **Tailor**, cytoplasmic RNA-specific terminal uridylyl-transferase (Cheng et al., 2019; Lin et al., 2017; Reimão-Pinto et al., 2015), part of the terminal RNA uridylation-mediated processing (TRUMP) complex; involved in 3'-to-5' exoribonucleolytic decay of RNA species by Dis3l2 (Lin et al., 2017); regulation of microRNA biogenesis by targeting precursor-microRNAs (predominantly mirtron hairpins) & targets unprocessed RNA polymerase III transcripts for degradation in a cytoplasmic RNA surveillance pathway (Reimão-Pinto et al., 2015); Uridyltransferases. |
| IntS14 | **TT; RNABPR; RGEXPPR; INTCOM** | **Integrator 14**, component of the Integrator complex; involved in the transcription of small nuclear RNAs (snRNA) and their 3'-box-dependent processing & the 3'-end processing of the U7 snRNA, and spliceosomal snRNAs U1 and U5 (Chen et al., 2012). |
| Etl1 | **DBTR2; CBTR2; CRTR2; POLIITR; HAT** | **Etl1**, subfamily of the Snf2 family of helicase-related proteins, SWI/SNF-related matrix-associated actin-dependent regulator of chromatin subfamily A containing DEAD/H box 1; ATP-dependent chromatin remodeling activity (InterPro Project Members, 2004-); ATP-dependent activity, acting on DNA; chromatin binding; DNA binding (GO Reference Genome Project, 2011-); H2A acetylation resulting in regulation of transcription (Doiguchi et al., 2016); SNF2-Like Chromatin Remodelers. |

**Table E. Subdivisions of regulatory factors associated with the Myc target P35/P36 (plotted in Figure 12)**

| Gene name | Pathway | Activity-Function |
|---|---|---|
| P32 | **CRTR2; CHRM** | **P32**, evolutionarily conserved mitochondrial glycoprotein, functions in presynaptic calcium signaling & neurotransmitter release as well as chromatin metabolism, histone binding; chromatin remodeling; nucleosome assembly; histone binding; sperm DNA binding (Emelyanov et al., 2014). |
| Hyls1 | **U** | **Hyls1** centriolar and ciliogenesis, involved in cilium assembly; located in centriole & ciliary basal body. |
| mask | **RNABTR2; POLIITR** | **Multiple ankyrin repeats single KH domain**, RNA binding (InterPro Project Members, 2004-); positive regulation of transcription by RNA pol II (Sansores-Garcia et al., 2013; Sidor et al., 2013). |
| Map60 (CP-60) | **POLIITR; DBTR2** | **Microtubule-associated protein**, microtubule binding protein; localized to the centrosome in a cell cycle-dependent manner (Kellogg et al., 1995); a component of a Centrosomal complex; regulation of transcription by RNA pol II (GO Reference Genome Project, 2011-); MADF Domain. |
| CG33490 (CG6110) | **U** | Orthologous to human EFHB (EF-hand domain family member B), regulation of calcineurin-NFAT signaling. |
| Nup50 | **CBTR2; POLIITR** | **Nucleoporin 50kDa**, nuclear pore complex protein, chromatin DNA binding; positive regulation of transcription by RNA pol II (Kalverda et al., 2010); important for TGF-beta signal transduction by mediating the nuclear translocation of the Mothers against dpp protein (*Mad*); Association of 400 genes interacting with Nup50 Nucleoporin, including transcriptionally active genes inside the nucleoplasm are involved in development & cell cycle regulation (Kalverda et al., 2010); Nuclear Pore Complex. |
| mod | **POLIITR; DBTR2; CRR; TF** | **Modulo**, the *Drosophila* homologue of nucleolin (Mikhaylova et al., 2006); MYC pathway (target of MYC selectively required for the growth of proliferative cells); association with the proto-oncogene MYC (Perinn et al., 2003); required for meiosis & spermatid differentiation in male germ line; sequence-specific DNA binding (Mikhaylova et al., 2006); involved in chromatin packaging; dominant suppressor of variegation (Bantignies et al., 2002). |

**Table E. Subdivisions of regulatory factors associated with the Myc target P35/P36 (plotted in Figure 12)**

| Gene name | Pathway | Activity-Function |
|---|---|---|
| AGO2 | **RNABPT; RGEXPPT; PSILEN** | **Argonaut 2**, interaction with small interfering RNAs (siRNAs) to form RNA-induced silencing complexes (RISCs), siRNA binding (Goh and Okamura, 2019; Kawamura et al., 2008; Tomari et al., 2007; Rand et al., 2005; Lingel et al., 2003); miRNA-mediated gene silencing (Besnard-Guérin et al., 2015); single-stranded RNA binding (Goh and Okamura, 2019); siRNA binding (Goh and Okamura, 2019; Kawamura et al., 2008; Tomari et al., 2007; Rand et al., 2005; Lingel et al., 2003). |
| Not3 | **RNABPR; RNABTN; TN; RGEXPPT; POLIITR** | **CCR4-NOT transcription complex subunit 3**, a poly(A)-specific ribonuclease involved in translation inhibition (Bawankar et al., 2013); nuclear transcribed mRNA catabolic process, deadenylation-dependent decay (Arvola et al., 2020); nuclear transcribed mRNA poly(A) tail shortening (Temme et al., 2010); regulation of DNA-templated transcription (InterPro Project Members, 2004-). |
| ncd | **CHRM** | **non-claret disjunctional**, a minus-end-directed kinesin microtubule motor protein and the sole member of the kinesin-14 motor family; chromosome segregation (Hallen et al., 2008); distributive segregation (Whyte et al., 1993); mRNA transport (Fahmy et al., 2014). |
| dsx-c73A | **TREX** | **doublesex cognate 73A**, a secreted cuticle protein present in the epidermis, salivary duct and trachea; expressed under control of (*ovo*); structural constituent of nuclear pores; mRNA export from nucleus; regulation of mitotic cell cycle (GO Reference Genome Project, 2011-); Unclassified Cuticle Proteins. |
| Nup93-1 | **CBPT; RGEXPPT** | **Nucleoporin 93kD-1**, chromatin binding; negative regulation of gene expression, epigenetic; chromatin DNA binding (Gozalo et al., 2020). |
| wupA | **CHRM; RRR** | **Wings up A**, a nuclear protein (Sahota et al., 2009) also found in striated muscle thin filament (Sarov et al., 2016); involved in calcium-dependent regulation of muscle contraction, development of the embryonic heart development (Wolf et al., 2006), adult somatic muscle development (Barthmaier and Fyrberg, 1995), & flight muscle; also contributes to non-muscle functions such as apico-basal polarity formation, nuclear division; maintenance of nuclear integrity (Sahota et al., 2009); actin binding; nervous system development; Troponin Complex (Barbas et al., 1991); Troponins I Isoforms (TPNI). |

**Table E. Subdivisions of regulatory factors associated with the Myc target P35/P36 (plotted in Figure 12)**

| Gene name | Pathway | Activity-Function |
|---|---|---|
| gro | **COREP; POLIITR** | **Groucho, isoform F**, a global developmental co-repressor in collaboration with DNA-binding repressor partner proteins & tethering to target promoters (Ajuria et al., 2011; Giagtzoglou et al., 2003; Jimenez et al., 2000; Goldstein et al., 1999; Valentine et al., 1998); downstream effector of signaling pathways such as Wg/Wnt & Dpp/TGF-beta; Phosphorylation & attenuation of Groucho repressor activity in response to MAPK activation; "context dependent regulatory domain binding" (CRD), a domain of about 130aa, the most divergent region among the LEF/TCF proteins (Arce et al., 2009). |
| rept | **DBTR1; DBTR2; CBRR; DNAHRR; DNAHTR1; DNAHTR2; RRR; CRTR2; HF; RGEXPPT; POLITR; COACT; POLIITR; HAT** | **Reptin**, RuvB-like helicase, involved in transcriptional regulation; DNA repair-dependent chromatin remodeling; NuA4 histone acetyltransferase complex (Kusch et al., 2004); heterochromatin formation (Qi et al., 2006); negative regulation of gene expression (Diop et al., 2008); positive regulation of transcription of nucleolar large rRNA by RNA polymerase I (Vinayagam et al., 2016); regulation of transcription by RNA pol II (GO Reference Genome Project, 2011-); core component of the chromatin remodeling Ino80 complex; chromatin remodeling (Klymenko et al., 2006); transcriptional coactivator in Wg signaling caused by altered arm signaling; Pont/rept antagonistically interfere with the nuclear Arm signaling ; an essential cofactor for the normal function of Myc; required for cellular proliferation and growth; ecdysone-mediated salivary gland cell autophagy cell death (Ihry and Bashirullah, 2014); negative regulation of canonical Wnt signaling (Bauer et al., 2000); regulation of cell population proliferation (Bellosta et al., 2005); INO80 Complex; TIP60 Complex; SWR1 Complex; RUVB-Like DNA Helicases. |
| CG33490 (CG6110) | **U** | Orthologous to human EFHB (EF-hand domain family member B), regulation of calcineurin-NFAT signaling cascade; Uncharacterized. |
| baf | **DBRR; CRR; CHRM; RGEXPRR** | **Barrier to autointegration factor**, highly conserved in metazoan evolution, plays fundamental roles in nuclear assembly, chromatin organization, gene expression & gonad development, may potently compress chromatin structure & be involved in membrane recruitment & chromatin decondensation during nuclear assembly;, required in both M phase & interphase of the cell cycle (Nikalayevich and Ohkura, 2015; Furukawa et al., 2003); DNA binding (InterPro Project Members, 2004-). |

**Table E. Subdivisions of regulatory factors associated with the Myc target P35/P36 (plotted in Figure 12)**

| Gene name | Pathway | Activity-Function |
|---|---|---|
| Larp4B | **PSILEN; RNABPT; TN; RGEXPPT** | **La-related protein Larp4B**, RNA-binding protein; post-transcriptionally inhibition of MYC protein; negative regulation of cell growth & translation (Funakoshi et al., 2018); mRNA 3'-UTR binding (Gene Ontology Curators, 2002-). |
