## Supplemental Tables for "Targeting Regulatory Factors Associated with the *Drosophila Myc cis*-Elements by Reporter Expression, Gel Shift Assay, and Mass Spectrometric Protein Identification": Table F.pdf

**Table F. Subdivisions of regulatory factors associated with the Myc target P37/P38 (plotted in Figure 12)**

| Gene name | Pathway | Activity-Function |
|---|---|---|
| Dmel\CG9641 | **U** | Uncharacterized; Unknown; Domain of Unknown Function DUF4780. |
| Cdk5 | **POLIITR; RRR** | **Cyclin-dependent kinase 5**, regulation of transcription involved in G1/S transition of mitotic cell cycle (GO Reference Genome Project, 2011-); cyclin-dependent protein, serine/threonine kinase activity (Connell-Crowley et al., 2000); proapoptotic signaling by CDK5 & MEKK1 (Ryoo H.D., 2018); centrosome localization (Jauffred et al., 2013); protein kinase 5 complex (Connell-Crowley et al., 2007). |
| (baf) | **DBRR; CRR; CHRM; RGEXPRR** | **Barrier to autointegration factor**, highly conserved in metazoan evolution, plays fundamental roles in nuclear assembly, chromatin organization, gene expression & gonad development, may potently compress chromatin structure & be involved in membrane recruitment & chromatin decondensation during nuclear assembly;, required in both M phase & interphase of the cell cycle (Nikalayevich and Ohkura, 2015; Furukawa et al., 2003); DNA binding (InterPro Project Members, 2004-). |
| (eff) | **CHRM; CRR; RRR; RGEXPPT** | **Effete**, E2 ubiquitin-conjugating enzyme, nuclear protein, protein ubiquitination and degradation pathway (Treier et al., 1992); roles include female germline stem cell maintenance, eye development, apoptosis regulation & chromatin organization; chromosome organization; mitotic cell cycle (Cenci et al., 1997); negative regulation of smoothened signaling pathway (Pan et al., 2017); ubiquitin ligase complex (GO Reference Genome Project, 2011-). |
| Dmel\Su(fu) | **DBTFB; REP; POLIITR** | **Suppressor of fused**, (Ser/Thr-protein kinase; negative regulator of Hedgehog signaling by forming a complex with the transcription factor Cubitus interruptus (Ci); (Han et al., 2019; Han et al., 2015; Smelkinson et al., 2007; Zhang et al., 2011; Fukumoto et al., 2001; Methot and Basler, 2000); negative regulator of Dpp pathway *(*Jia et al., 2002-nature); DNA binding transcription factor binding (Han et al., 2015); molecular sequestering activity (Han et al., 2019; Han et al., 2015; Oh et al., 2015; Shi et al., 2014; Wang et al., 2000). |
| Dmel\Elp4 (CG6907) | **POLIITR; TN; RRR; RNABTN** | **Elongator complex protein 4**, establishment of mitotic spindle asymmetry; regulation of translation through targeted transfer-RNA (tRNA) modification (Planelles-Herrero et al., 2022); phosphorylase kinase regulator activity; regulation of transcription by RNA pol II (Gene Ontology Curators, 2002-); Elongator Complex. |

**Table F. Subdivisions of regulatory factors associated with the Myc target P37/P38 (plotted in Figure 12)**

| Gene name | Pathway | Activity/Function |
|---|---|---|
| Fmr1 | **RNABPT; RNAHPT; RGEXPPT; PSILEN** | **Fragile X messenger ribonucleoprotein 1**, RNA & channel binding protein; DEAD/H-Box RNA helicase binding (Barbee et al., 2006); mRNA regulatory element binding repressor activity (Fajner et al., 2021); post-transcriptional regulation of gene expression (Luhur et al., 2017). |
| Tailor | **POLIIITR; RGEXPTR3; RNABTR3; RNABPR** | **Tailor**, cytoplasmic RNA-specific terminal uridylyltransferase (Cheng et al., 2019; Lin et al., 2017; Reimão-Pinto et al., 2015), part of the terminal RNA uridylation-mediated processing (TRUMP) complex; involved in 3'-to-5' exoribonucleolytic decay of RNA species by Dis3l2 (Lin et al., 2017); regulation of microRNA biogenesis by targeting precursor-microRNAs (predominantly mirtron hairpins) & targets unprocessed RNA polymerase III transcripts for degradation in a cytoplasmic RNA surveillance pathway (Reimão-Pinto et al., 2015); Uridyltransferases |
| Dmel\Lam (CG6944) | **CBRR; HF** | **Lamin**, chromatin binding (Verboon et al., 2015); Heterochromatin formation (Verboon et al., 2015; Dialynas et al., 2010; Shevelyov et al., 2009); regulation of meiosis I cytokinesis (Hayashi et al., 2016); Lamins. |
| Rcd-1 | **RGEXPPR; TN; RGEXPTN** | **Regulator of cell death-1,** regulation of mRNA catabolic process; negative regulation of translation (Bawankar et al., 2013); programmed cell death (Daskalov et al., 2019); CCR4-NOT Complex (Temme et al., 2010); Armadillo-like-helical. |
| (abs) | **RNAHPR; RNABPR; RGEXPPR** | **Abstrakt**, DEAD/DEAH-Box RNA helicase (Lasko, 2000), RNA binding (GO Reference Genome Project, 2011-); RNA splicing, via spliceosome (Herold et al., 2009); regulates cell polarity in oocytes & embryos; downregulation of Notch signaling in asymmetric cell division in ganglion mother cell (GMC2-4a) in collaboration with Inscuteable (*Insc*) (Irion et al., 2004). |
| Dmel\CG8064 | **RNABPT; RGEXPPT** | SnoRNA binding; involved in maturation of SSU-rRNA (GO Reference Genome Project, 2011-); ); human ortholog(s) implicated in papillary thyroid carcinoma; orthologous to human WDR3 (WD repeat domain 3). |
| IntS11 | **RNABTR2; INTCOM; POLIITR; RNABPR** | **Integrator complex subunit 11**, involved in the transcription of small nuclear RNAs (snRNA) & their 3'-box-dependent processing; pol II transcription (Baillat et al., 2005); Notch signaling (Elena Shersher et.al., 2021-Cell Commun Signal); Epidermal growth factor pathway (F. C.Tilley & G. Mollet, 2021). |

**Table F. Subdivisions of regulatory factors associated with the Myc target P37/P38 (plotted in Figure 12)**

| Gene name | Pathway | Activity-Function |
|---|---|---|
| Rcc1 | **CBRR; CHRM** | **Regulator of chromosome condensation**, nuclear import & export of beta-Catenin (Koyama et at., 2017); regulation of mitotic cell cycle, apoptosis pathway; chromatin binding (Trieselmann and Wilde, 2002). |
| Usp10 | **POLIITR; TE; RGEXPPT** | **Ubiquitin specific protease 10**, cysteine-type deubiquitinase activity; ubiquitin-dependent protein catabolic process (Gene Ontology Curators, 2002-); negative regulation of transcription elongation (GO Reference Genome Project, 2011-); positive regulation of Notch signaling (Zhang et al., 2012). |
| Dmel\shep | **REP; DBTR2; CBTR2; POLIITR** | **Protein alan shepard**, an evolutionarily conserved RNA/DNA binding protein; regulation of alternative splicing & gypsy insulator activities (chromatin insulator associated) (Chen et al., 2021). |
| Dmel\Polr2E | **RPIB5; RPIIB5; RPIIIB5; DBTR1; DBTR2; DBTR3; POLITR; POLIITR; POLIIITR** | **RNA polymerase II, I and III subunit E**, DNA binding (InterPro Project Members, 2004-); DNA-directed 5'-3' RNA polymerase activity; RNA polymerase II activity (Aoyagi and Wassarman, 2000); RNA polymerase I activity; RNA polymerase III activity (GO Reference Genome Project, 2011-); RNA Polymerase I Complex; RNA Polymerase II Complex; RNA polymerase III Complex. |
| Dmel\hoip | **RNABPR; RNABTN; RGEXPPR; TN; RGEXPTN** | **(hoi-polloi),** RNA binding (GO Reference Genome Project, 2011-); positive regulation of translation (Williams et al., 2015); positive regulation of gene expression (Johnson et al., 2013); Spliceosome Complex B. |
| CG6418 (DmRH27) | **RNAHPR; RNABPR** | RNA helicase , nuclear RNA binding; mRNA splicing (InterPro Project Members, 2004-; Gene Ontology Curators, 2002-); DEAD-BOX RNA HELICASES. |
| Dmel\Sf3a1 | **RNABPR; RGEXPPR** | **Splicing factor 3a subunit 1**, RNA binding (GO Reference Genome Project, 2011-); RNA processing (InterPro Project Members, 2004-). |
| Dmel\Ote | **DBTFB; COREP; POLIITR** | **Otefin**, a nuclear membrane-associated protein, acts in concert with BMP/DPP signaling to mediate "bag of marbles" (*bam*) transcriptional silencing; positive regulation of BMP/DPP signaling; DNA binding transcription factor binding; transcription corepressor activity; negative regulation of DNA-templated transcription; interacts with Medea/Smad4 at the bam silencer element to regulate germline stem cell (GSC) fate (Jiang et al., 2008). |

**Table F. Subdivisions of regulatory factors associated with the Myc target P37/P38 (plotted in Figure 12)**

| Gene name | Pathway | Activity-Function |
|---|---|---|
| Dmel\crp | **TF; POLIITR; DBTR2** | **Activator protein 4 (Cropped)**, a downstream target of MYC, plays role in cell growth, organ size & survival; myosin binding (Liu et al., 2008); transcription factor; sequence-specific DNA binding (King-Jones et al., 1999); RNA pol II *cis*-regulatory region, sequence-specific DNA binding (GO Reference Genome Project, 2011-); protein dimerization activity (InterPro Project Members, 2004-); larval somatic muscle development (Dobi et al., 2014); Basic Helix-Loop-Helix Transcription Factors. |
| kis | **CBTR2; CRTR2; DBTR2; TF; POLIITR; RGEXPPT** | **Kismet, isoform F**, a conserved chromodomain containing ATP-dependent transcription factor (Terriente-Félix et al., 2011); gene control through epigenetic mechanisms; chromatin binding; ATP-dependent chromatin remodeling; DNA binding; regulation of gene expression (GO Reference Genome Project, 2011-); negative regulation of DNA-templated transcription (Thompson et al., 2008). |
| (gro) | **COREP; POLIITR** | **Groucho, isoform F**, a global developmental co-repressor in collaboration with DNA-binding repressor partner proteins & tethering to target promoters (Ajuria et al., 2011; Giagtzoglou et al., 2003; Jimenez et al., 2000; Goldstein et al., 1999; Valentine et al., 1998); downstream effector of signaling pathways such as Wg/Wnt & Dpp/TGF-beta; Phosphorylation & attenuation of Groucho repressor activity in response to MAPK activation; "context dependent regulatory domain binding" (CRD), a domain of about 130aa, the most divergent region among the LEF/TCF proteins (Arce et al., 2009). |
| (simj) | **REP; POLIITR; CRTR2; HDAC** | **Simjang, isoform D**, (mammalian GATAD2B ortholog), a component of the chromatin remodeling (deacetylation) NURD complex, couples chromatin remodeling & histone deacetylation to mediate transcriptional repression; negative regulation of transcription by RNA pol II (Kon et al., 2005); recruited by methyl-CpG-DNA binding proteins (Kim et al., 2004); modifier (repressor) of Wnt signaling (NURD complex in Wnt signaling): positive regulation of synaptic assembly at neuromuscular junction (Willemsen et al., 2013; Kon et al., 2005); Nucleosome Remodeling Deacetylase Complex; Transcriptional Repressor p66; coiled-coil MBD2 interaction domain: MBD2-NURD: complex recognition of methylated DNA & silencing the expression of associated genes through histone deacetylase & nucleosome remodeling (Walavalkar et al., 2013; Gnanapragasam et al., 2011). |

**Table F. Subdivisions of regulatory factors associated with the Myc target P37/P38 (plotted in Figure 12)**

| Gene name | Pathway | Activity-Function |
|---|---|---|
| Dmel\CG6000 | **U** | Orthologous to human TSTD1 (thiosulfate sulfurtransferase like domain containing 1), Rhodanese-like domain. |
| (Tlk) | **CRR; CHRM** | **Tousled-like kinase**, conserved anti-silencing function protein 1 (ASF1)/Tousled-like kinase (TLK), coordination of cell cycle phases via chromatin organization (Carrera et al., 2003); Cell migration/Apoptosis pathway (Zhang Y, Cai R, Zhou R, Li Y, Liu L. 2016); association of Tousled-like kinase with the protein complex of wingless signaling regulators (Milan et al. 1998); chromosome segregation (Li et al., 2009). |
| Dmel\CG12384(BcDNA:RH17411) | **POLIITR** | Negative regulation of DNA-templated transcription (Gene Ontology Curators, 2002-); apoptotic signaling pathway (GO Reference Genome Project, 2011-); cellular response to amino acid starvation; Death-Associated Protein 1: DAP1/DAPL1. |
| dsx-c73A | **TREX** | **(doublesex cognate 73A)**, a secreted cuticle protein present in the epidermis, salivary duct and trachea; expressed under control of (ovo); structural constituent of nuclear pores; mRNA export from nucleus; regulation of mitotic cell cycle (GO Reference Genome Project, 2011-); Unclassified Cuticle Proteins. |
| Hyls1 | **U** | **Hyls1**, centriolar and ciliogenesis, involved in cilium assembly; located in centriole & ciliary basal body. |
| koi | **RRR; DBRR** | **Klaroid**, double-strand break repair via homologous recombination (Ryu et al., 2015); nuclear migration (Kracklauer et al., 2007); nuclear organization (Tan et al., 2018; Elhanany-Tamir et al., 2012). |
| MRG15 | **HAT; CRTR2; CBTR2; CRR; CBRR; RGEXPPT; POLIITR; CHRM; RRR; HF** | **MORF-related gene 15**, histone acetylation; chromatin binding; positive regulation of gene expression (Huang et al., 2017); chromatin organization; chromosome separation (Smith et al., 2013); DNA repair-dependent chromatin remodeling (Kusch et al., 2004); heterochromatin formation (Qi et al., 2006); regulation of DNA-templated transcription (InterPro Project Members, 2004-). |
| Dmel\CG6026 | **COACT; POLIITR** | Nuclear protein, transcription coregulator activity; chromatin remodeling; positive regulation of pseudohyphal growth by positive regulation of transcription from RNA polymerase II promoter (GO Reference Genome Project, 2011-). |
