## Supplemental Tables for "Targeting Regulatory Factors Associated with the *Drosophila Myc cis*-Elements by Reporter Expression, Gel Shift Assay, and Mass Spectrometric Protein Identification": Table G.pdf

**Table G. Subdivisions of regulatory factors associated with the pos. control W-CRM (plotted in Figure 12)**

| Gene name | Pathway | Activity/Function |
|---|---|---|
| Bap111 | **DBTR2; CRTR2; POLIITR; TF** | **Brahma associated protein 111kD**, Extensive homology of the Brahma (BRM) complex to SWI/ SNF; DNA binding; chromatin remodeling (Papoulas et al., 2001); associated Proteins Complex; Polybromo-Containing Proteins Complex; High Mobility Group Box Transcription Factors. |
| Iswi | **CRTR2; CHRM; CBTR2; TF; DBTR2; POLIITR** | **Imitation SWI**, energy-transducing component of the chromatin-remodeling complexes NURF (nucleosome-remodeling factor), ACF (ATP-utilizing chromatin assembly and remodeling factor), & CHRAC (chromatin accessibility complex (Corona et al., 2007; Eberharter et al., 2001); DNA-binding transcription factor binding (Vanolst et al., 2005); nucleosome array spacer activity (Ito et al., 1997; Varga-Weisz et al., 1997); nucleosome binding (InterPro Project Members, 2004-); chromatin organization (Corona et al., 2007; Mizuguchi et al., 1997); regulation of pol II transcription (Li et al., 2010); sperm DNA condensation (Doyen et al., 2015). |
| Snr1 | **CRTR2; COACT; POLIITR; TF; DBTR2** | **Snf5-related 1**, (ATP-dependent chromatin remodeler; a counterpart of yeast SWI/SNF) (Swi3 component of the Brahma Associated Proteins complex), SET Domain Binding (Marenda et al., 2003); cell differentiation; Tat protein binding (Koe et al., 2014); transcription coactivator activity (GO Reference Genome Project, 2011-); chromatin remodeling (Dingwall et al., 1995); regulation of transcription by RNA pol II (Bonnay et al., 2014); positive regulation of DNA-templated transcription (Dingwall et al., 1995; Kal et al., 2000). |
| Dmel\Spt5 | **POLIITR; TE; CBTR2; CRTR2; RNABTR2** | Spt5, component of the DRB sensitivity-inducing factor complex (DSIF complex, DSIF enhances transcriptional pausing at sites proximal to the promoter, may facilitate the assembly of an elongation competent RNA pol II complex. DSIF may also promote transcriptional elongation within coding regions. DSIF is required for the transcriptional induction of heat shock response genes & regulation of genes which control A-P patterning during embryonic development; chromatin binding; mRNA binding; involved in promoter-proximal pausing (Qiu and Gilmour, 2017); protein-containing complex binding (Saunders et al., 2003); RNA pol II complex binding (Qiu and Gilmour, 2017; Saunders et al., 2003); negative regulation of DNA-templated transcription, elongation (Wu et al., 2003); transcription elongation-coupled chromatin remodeling (InterPro Project Members, 2004-). |

**Table G. Subdivisions of regulatory factors associated with the pos. control W-CRM (plotted in Figure 12)**

| Gene name | Pathway | Activity/Function |
|---|---|---|
| Spt20 | **POLIITR; HAT; COREG** | **Spt20, isoform C,** Spt-Ada-Gcn5-acetyltransferase **(SAGA) complex; i**nvolved in histone acetylation (Gene Ontology Curators, 2002-); regulation of transcription by RNA polymerase II (GO Reference Genome Project, 2011-); transcription coregulator activity (InterPro Project Members, 2004-); SAF6 (SAGA factor-like TAF6), a histone fold domain-containing containing protein can replace Taf6 in *Drosophila* SAGA complex & is required for SAGA-dependent gene expression; the gene CG17689 encodes an ortholog of Spt20/p38IP, CG9866 encodes a potential ortholog of Sgf73 & CG3883 encodes a novel histone fold domain (HFD)-containing protein SAF6; (Weake et al., 2009); SAGA (Spt-Ada-Gcn5 Acetyltransferase) Complex; Transcription Factor Spt20. |
| Mapmodulin | **RGEXPPT** | **Mapmodulin (MAPM), isoform D**, suppression of transformation (tumor suppressor), regulation of mRNA trafficking & stability; histone binding (GO Reference Genome Project, 2011-); inhibition of acetyltransferases as part of the inhibitor of histone acetyltransferases (INHAT) complex; nucleocytoplasmic transport (Gene Ontology Curators, 2002-); Acidic Leucine-rich Nuclear Phosphoprotein 32. |
| Su(var)2-10 | **COACT; POLIITR; CHRM** | **Suppressor of variegation 2-10, isoform L**, a member of the PIAS protein family that regulates chromosome structure and function; chromosome organization & condensation (Hari et al., 2001); transcription coregulatory activity, regulation of transcription by RNA pol II (GO Reference Genome Project, 2011-); JAK/STAT pathway regulator, contributes to eye formation & eye determination (Betz et al., 2001); negative regulation of JAK-STAT signaling (Muller et al., 2005; Betz et al., 2001); positive regulators of Hedgehog signaling (Ma et al., 2016; Zhang et al., 2017); negative regulators of Imd signaling pathway (Tang et al., 2021; Cronin et al., 2009). |
| HDAC1 | **HDAC; RGEXPPT; CHRSILEN; CRTR2; POLIITR; CHRM** | **Histone deacetylase 1**, deacetylation of lysine residues on the N-terminal part of the core histones (H2A, H2B, H3 and H4); negative regulation of gene expression, epigenetics (Janssens et al., 2017); Hedgehog signaling (Zhang et al., 2013); transcription corepressor activity (Miotto et al., 2006); chromosome condensation (Nikalayevich and Ohkura, 2015); NAD-independent histone deacetylation (Feller et al., 2015); regulation of DNA-templated transcription (Cho et al., 2005). |

**Table G. Subdivisions of regulatory factors associated with the pos. control W-CRM (plotted in Figure 12)**

| Gene name | Pathway | Activity-Function |
|---|---|---|
| Dmel\shep | **REP; DBTR2; CBTR2; POLIITR** | **Protein alan shepard**, an evolutionarily conserved RNA/DNA binding protein; regulation of alternative splicing & gypsy insulator activities (chromatin insulator associated) (Chen et al., 2021). |
| Dmel\Dbp80 | **RNAHTR2; RNABTR2; TREX** | Dead box protein 80, poly(A) + mRNA export from nucleus; RNA binding; RNA helicase activity (GO Reference Genome Project, 2011-); DEAD-Box RNA Helicases |
| l(3)72Ab (Dmel\Brr2) | **RNAHPR; RNABPR** | **U5 small nuclear ribonucleoprotein 200 kDa helicase**, mitotisc cell cycle (Ducat et al., 2008); steroid hormone ecdysone signaling (Claudius et al., 2014); RNA Helicase activity (Lasko, 2000); mRNA splicing, via spliceosome (Monedero Cobeta et al., 2018). |
| B52 | **POLIITR; TI; CRTR2** | **B52**, regulation of gene expression; regulation of transcription start site selection at RNA pol II promoter (Bradley et al., 2015); condensation or decondensation of chromatin; associated with boundaries of transcriptionally active chromatin (Champlin et al., 1991). |
| Dmel\Hpr1 | **TREX; TE** | **Hpr1,** orthologous to human THOC1 (THO complex subunit 1); mRNA export from nucleus in response to heat stress (Rehwinkel et al., 2004); regulation of DNA-templated transcription elongation (GO Reference Genome Project, 2011-); Transcription Export Complex. |
| (aub) | **RGEXPPT; RNABPT** | **Aubergine**, piRNA binding (Huang et al., 2021; Webster et al., 2015; Nagao et al., 2010); global gene silencing by mRNA cleavage (Kennerdell et al., 2002); ncRNA-mediated post-transcriptional gene silencing (Tomari et al., 2004); RNA-mediated gene silencing (Bozzetti et al., 2015); JAK-STAT signaling pathway controls host defense in the gut by regulating stem cell proliferation & epithelial cell homeostasis (Cronin et al., 2009); RNA binding (Brennecke et al., 2007). |
| **Dmel/how** | **RNABPT; RGEXPPT** | **held out wings**, RNA-binding protein (Volohonsky et al., 2007; Di Fruscio et al., 1998) highly expression: mesoderm & tendon cells; two isoforms: how(L) & how(S). how(L) induces RNA destabilization, How(S) stabilizes the RNA targets; integrin signaling, apposition of dorsal/ventral imaginal disc-derived wing surfaces (Walsh and Brown, 1998); cell adhesion (Lo and Frasch, 1997); EGFR signaling: Glial cell migration; axon ensheatment (Edenfeld et al., 2006; Lasko, 2003); Dpp (Israeli & Volk, 2007). |

**Table G. Subdivisions of regulatory factors associated with the pos. control W-CRM (plotted in Figure 12)**

| Gene name | Pathway | Activity/Function |
|---|---|---|
| MED4 | **COACT; MS; POLIITR; TPI** | **Mediator complex subunit 4**, component of the Mediator complex transcription coactivator activity (Gu et al., 2002); transcription coregulator activity (Park et al., 2001); RNA pol II transcription; serves as a scaffold for the assembly of a functional preinitiation complex with RNA polymerase II and the general transcription factors (Gu et al., 2002; Park et al., 2001); Mediator Complex. |
| (woc) | **POLIITR; TF; CBTR2; DBTR2; TMM** | **Without children, isoform B**, **ecdysone**, biosynthetic process (Wismar et al., 2000); regulation of transcription by RNA pol II (Abel et al., 2009; Font-Burgada et al., 2008); chromatin-binding factor (Font-Burgada et al., 2008) related to the mammalian MYM-type family of transcription factors; involved in telomere capping. |
| Hrb87F | **RNABPR; RGEXPPR; DBRR; TMM** | **Heterogeneous nuclear ribonucleoprotein at 87F**, hnRNP-A family RNA-binding protein; involved in gene expression & RNA processing (Lasko, 2000); an essential component of the nucleoplasmic omega speckles; necessary for telomere maintenance; sequence-specific DNA binding (Borah et al., 2009); regulation of alternative mRNA splicing, via spliceosome (Borah et al., 2009; Park et al., 2004); Wnt/wingless and JNK signaling (Yadav & Tapadia, 2016). |
| Dmel\lds | **TE; POLIITR; CRTR2; RRR** | **Lodestar**, Transcription termination factor 2, ATP-dependent activity, acting on DNA; DNA-templated transcription termination (Xie and Price, 1998); ATP-dependent chromatin remodeler activity (InterPro Project Members, 2004-); DNA repair (GO Reference Genome Project, 2011-); microtubule anchoring at centrosome, acts during the meiotic & cleavage divisions (Szalontai et al., 2009); SNF2-Like Chromatin Remodelers; part of spindle check point complex. |
| MRG15 | **HAT; CRTR2; CBTR2; CRR; CBRR; RGEXPPT; POLIITR; CHRM; RRR; HF** | **MORF-related gene 15**, histone acetylation; chromatin binding; positive regulation of gene expression (Huang et al., 2017); chromatin organization; chromosome separation (Smith et al., 2013); DNA repair-dependent chromatin remodeling (Kusch et al., 2004); heterochromatin formation (Qi et al., 2006); regulation of DNA-templated transcription (InterPro Project Members, 2004-). |
| XRCC1 | **RRR; DBRR** | **X-ray repair cross complementing 1**, damaged DNA binding activity; involved in base-excision repair; active in nucleus; single strand break repair; double-strand break repair via nonhomologous end joining (InterPro Project Members, 2004-). |

**Table G. Subdivisions of regulatory factors associated with the pos. control W-CRM (plotted in Figure 12)**

| Gene name | Pathway | Activity/Function |
|---|---|---|
| Dmel\CG9411 | **U** | Uncharacterized, unknown. |
| Dmel\CG10591 | **U** | Uncharacterized, unknown. |
| Dmel\CG15239 | **U** | Expressed in embryonic dorsal epidermis; embryonic esophagus; embryonic head epidermis; embryonic ventral epidermis; and embryonic/larval salivary gland; Domain of Unknown Function; DUF4773. |
| Dmel\CG42353 | **U** | Uncharacterized, unknown. |
| tst | **RNABPR; RNAHPR; RGEXPPR** | (**twister**) RNA helicase, helicase activity (Lasko, 2000); RNA helicase activity; RNA catabolic process (InterPro Project Members, 2004-); Nuclear-transcribed mRNA catabolic process, 3'-5' exonucleolytic nonsense-mediated decay (GO Reference Genome Project, 2011-); regulation of mRNA alternative splicing (Park et al., 2004); SKI2-like RNA helicases. |
| Cbp20 | **RNABPR; PSILEN; RGEXPPR** | **20 kDa nuclear cap-binding protein**, mRNA binding (Lasko, 2000); RNA binding; RNA cap binding (InterPro Project Members, 2004-); siRNA processing (Sabin et al., 2009); ncRNA-mediated post-transcriptional gene silencing; NCBP2 RNA Recognition Motif. |
| Dmel\nonA | **POLIITR; RNABTR2** | **no on or off transient A**, mRNA binding (Greenspan and Ferveur, 2000; Lasko, 2000); RNA binding; regulation of DNA-templated transcription (GO Reference Genome Project, 2011-); mRNA splicing, via spliceosome (Herold et al., 2009); RNA Recognition Motif Domain. |
| Dmel\Nup133 | **POLIITR; TREX** | **Nucleoporin 133kD**, component of the nuclear pore complex (NPC) (Probable). Plays a role in NPC assembly and/or maintenance; poly(A) + mRNA export from nucleus; transcription-dependent tethering of RNA pol II gene DNA at nuclear periphery (GO Reference Genome Project, 2011-); mRNA export from nucleus (Gene Ontology Curators, 2002-); Nuclear Pore Complex. |
| Cpsf160 | **TT; PAFAC; RNABPR** | **Cleavage and polyadenylation specificity factor 160**, a key role in pre-mRNA 3'-end formation, recognizing the AAUAAA signal sequence and interacting with poly(A) polymerase and other factors to bring about cleavage and poly(A) addition (Salinas et al., 1998); mRNA polyadenylation (Mount and Salz, 2000); polyadenylation factors. |
