## Supplemental Tables for "Targeting Regulatory Factors Associated with the *Drosophila Myc cis*-Elements by Reporter Expression, Gel Shift Assay, and Mass Spectrometric Protein Identification": Table A.pdf

| Signaling Pathways | Abbreviations |
|---|---|
| Apoptosis | APOP |
| Amphiphysin-Rho1-Dia/DAAM-Rok | ARDAMM |
| Autophagy (AUTO) | AUTO |
| Cell Cycle | CECY |
| CoA biosynthesis | COAS |
| DPP/BMP (TGF-β) | DPPBMP |
| Ecdysteroid | ECDY |
| Epidermal Growth Factor Receptor | EGFR |
| Eiger/ Wengen (EIWE) | EIWE |
| Ubiquitin-dependent ERAD | ERAD |
| ERK1/2 | ERK |
| Exon Junction Complex | EXJC |
| Fibroblast Growth Factor Receptor | FGFR |
| G-protein-coupled receptor Sun/Mth | GPCR |
| Heme-biosynthesis pathway | HEBI |
| Hepatocyte Growth Factor (HGF) | HGFS |
| Hedgehog/Smoothened | HH |
| Hypoxia Inducible Factor (HIF) | HIF |
| Hippo | HIPP |
| Immune deficiency (IMD) | IMD |
| Insulin-like Receptor | INR |
| interferon Immune | INTERIM |
| JAK/STAT | JAKSTAT |
| Jasmonate (JA) & Auxin | JASAU |
| Jun Kinase | JNK |
| Kennedy pathway | KENN |
| MAPK | MAPK |
| (Mitochondrial) Ca2+-mediated | MCAM |
| miRNA | miRNA |
| MYC | MYC |
| NF-κB | NFKB |
| Notch | NOTC |
| Poly(ADP-RIbosyl)ation | PADPRI |
| Platelet-derived Growth Factor Receptor | PDGFR |
| piRNA | piRNA |
| PVR (PDGFR/VEGFR) | PVR |
| Rac GTPase Pak kinase | RACP |
| RAS GTPase | RAS |
| Receptor Tyrosine Kinase (RTK) | RTK |
| Sema1,3A/Plexin-A | SAPA |
| Sevenless | SEVE |
| Sterol Regulatory-Element Binding protein | SREBP |
| TGFalpha (ATR/Mei-41 kinase) | TGFA |
| TNFα/Eiger-Mediated (TNFA) | TNFA |
| Toll | TOLL |
| TORC1,2 | TORC |
| Torso | TORS |
| Vascular Endothelial Growth Factor Receptor | VEGFR |
| Vesicular Trafficking | VTRA |
| Wingless (Wnt/β-Catenin/TCF) | WNBT |
| Wingless (Ca2+-mediated) | WNCA |
| With No Lysine kinase (WNK) | WNK |
| Wingless (Ca2+-mediated) | WNPCP |

**Table A. Supplement to Figure 13: Signaling pathways linked to the Myc-CRM P1/P2**

| Gene name | Signaling Pathway | Activity/Function |
|---|---|---|
| MED17 | **JASAU; RAS; MAPK** | **Mediator of RNA pol II transcription subunit 17**, sex comb development; required for adult cell & segment identity specification; SP1 transcription activation; thyroid hormone receptor complex component, VitD receptor interactor; jasmonate (JA) & Auxin signaling component (Boube et al., 2000); Ras/MAK pathway (Singh and Han 1995). |
| MED24 | **APOP; ECDY** | **Mediator of RNA pol II transcription subunit 24**, positive regulation of apoptosis pathway (Wang et al., 2008); ecdysone-mediated induction of salivary gland cell autophagic cell death (Ihry and Bashirullah, 2014). |
| skd | **WNBT; EGFR; HH; DPPBMP; NOTC** | **Skuld/MED13**, Wnt-TCF signaling pathway (Carrera et al., 2008); EGFR signaling; Hedgehog & Ras/MAPK signaling (Lim & Choi, 2007); Decapentaplegic (Dpp), and Notch (N) signaling (Borod and Heberlein, 1998; Fu and Baker, 2003; Greenwood and Struhl, 1999; Sun et al., 1998). |
| Hcf (dHCF, dHcf1, Hcf1) | **CECY; WNBT** | **Host cell factor**, phosphoprotein, involved in control of cell cycle (Guelman et al., 2006), & several processes, including chromatin remodeling, histone acetylation, & positive regulation of DNA-templated transcription; implicated in both activation & repression of transcription; associates with Wingless enhanceosome as a regulator of canonical Wingless signaling pathway during wing imaginal disc vein development (Rodriguez-Jato et al., 2011). |
| rept | **WNBT; MYC; APOP; CECY** | **RuvB-like helicase Reptin**, transcriptional coactivator in Wg signaling caused by altered Arm signaling; antagonistic interference of Pontin/Reptin with the nuclear Arm signaling; essential cofactor for the normal function of Myc; required for cellular proliferation and growth; ecdysone-mediated salivary gland cell autophagy cell death (Ihry and Bashirullah, 2014); negative regulation of canonical Wnt signaling pathway (Bauer et al., 2000); negative regulation of cell population proliferation (Bellosta et al., 2005). |
| Rpt4 | **NOTC; INR; CECY** | **Regulatory particle triple-A ATPase 4**, autophagy via hypoxia signaling (Lőw et al., 2013); Notch-mediated follicle cell differentiation and cell cycle switches, insulin-PI3K pathway (Jia et al., 2015). |
| Iswi | **ECDY** | **Imitation SWI**, chromatin-remodeling complex ATPase chain Iswi, ecdysteroid signaling and metamorphosis (Badenhorst et al., 2005; Tsukiyama & Wu, 1995). |

**Table A. Supplement to Figure 13: Signaling pathways linked to the Myc-CRM P1/P2**

| Gene name | Pathway | Activity/Function |
|---|---|---|
| E(bx) | **JAKSTAT** | **Enhancer of bithorax**, Component of Nucleosome Remodeling Factor NURF; negative regulation of receptor signaling pathway via JAK-STAT Signaling; spermatid differentiation; negative regulation in innate immune response; (Kwon et al., 2009); nuclear receptor binding (Badenhorst et al., 2005). |
| pont | **JNK; TNFA; WNBT** | **RuvB-like helicase 1 (Pontin),** canonical Wingless/Wnt pathway (Torres & Inestrosa, 2017), interaction with HMG box transcription factors of the lymphoid-enhancing factor-1 (LEF-1/T-cell factor (TCF) family, positive regulation of canonical Wnt signaling pathway (Bauer et al., 2000); JNK, negative regulation of tumor necrosis factor-mediated (TNFα-Eiger) signaling pathway (Wang et al., 2008). |
| Su(var)2-10 | **JAKSTAT; HH; IMD** | **Suppressor of variegation 2-10, isoform L**, JAK/STAT pathway regulator, contributes to eye formation & eye determination (Betz et al., 2001); negative regulators of JAK/STAT signaling (Muller et al., 2005; Betz et al., 2001); positive regulators of Smoothened signaling (Zhang et al., 2017; Ma et al., 2016); negative regulators of Imd signaling pathway (Tang et al., 2021; Cronin et al., 2009). |
| Mlf | **~~WNBT~~; HH; CECY** | **Myelodysplasia/myeloid leukemia factor**, regulation of cell proliferation involved in compound eye morphogenesis (Jasper et al., 2002); negative regulator of cell cycle progression functioning upstream of the tumor suppressor p53 (Yoneda-Kato et al, 2005); indirect regulator of Hedgehog & Wnt by binding Su(fu) (Bras et al, 2012), the negative regulator of both Hedgehog (Monnier et al., 1998) & Wnt (Meng X, et al. 2001) pathways. |
| Wdr62 | **CECY** | **WD repeat domain 62, isoform D**, regulation of mitotic cell cycle (Ramdas Nair et al., 2016). |
| caz | **HIPP** | **Chromatin and RNA-binding protein Cabeza** (Mallik et al., 2018; InterPro Project Members, 2004-), ('cabeza' means 'head' in Spanish), a single ortholog of human FUS in *Drosophila*; genetic link exists between Caz & Hippo signaling pathway: Hippo (Hpo) the *Drosophila* ortholog of human mammalian sterile 20-like kinase (MST) 1 rescues Cabezas knockdown-induced eye phenotype- and neuron-specific defects (Azuma et al., 2018; Shimamura et al., 2014; Xia et al., 2012); transcription coregulatory activity (GO Reference Genome Project, 2011-); transcription Factor TFIID. |

**Table A. Supplement to Figure 13: Signaling pathways linked to the Myc-CRM P1/P2**

| Gene name | Pathway | Activity/Function |
|---|---|---|
| Doc3 | **WNBT; DPPBMP** | **Dorsocross3**, Wnt signaling; TGF-beta pathway (Hatton-Ellis et al., 2007); cardiac induction in *Drosophila* relies on combinatorial Dpp and Wg signaling activities that are derived from the ectoderm (Reim and Frasch, 2005). |
| lola | **NOTC; APOP; IMD; NFKB** | **Longitudinals lacking protein, isoforms H/M/V**, involved in Notch signaling; cell death; expression of axon & dendrite patterning genes; regulation of retrotransposons; oogenesis; spermatogenesis; neural wiring; eye patterning & a variety of behaviors; positive regulation of peptidoglycan recognition protein signaling pathway; positive regulation of biosynthetic process of antibacterial peptides active against Gram-negative bacteria (Kleino et al., 2005); positive regulators of immune deficiency Imd signaling pathway (results in increased activity of the NF-κB-like transcription factor Rel in the nucleus). |
| ash2 | **EGFR; ECDY** | **absent, small, or homeotic discs 2**, imaginal disc-derived wing vein specification (Angulo et al., 2004); response to ecdysone (Carbonell et al., 2013). |
| Mad | **DPPBMP; WNBT; EGFR** | **Mothers against decapentaplegic**, BMP signaling pathway core components (Vuilleumier et al., 2022; Guo et al., 2013; Weiss et al., 2010; Kamiya et al., 2008; Yao et al., 2006; Muller et al., 2003; Dai et al., 2000; Das et al., 1998; Inoue et al., 1998); involved in wing development via modulation of EGFR and BMP signaling pathways (Dworkin and Gibson, 2006; Lecuit et al., 1996); negative regulation of salivary gland boundary formation with the involvement of Wnt/Wg signaling pathway (Bradley, P.L., Haberman, A.S., Andrew, D.J. 2001). |
| ebi | **JNK; EGFR; WNBT; NOTC** | **F-box-like/WD repeat-containing protein ebi-like**, evolutionarily conserved repressor/silencer; JNK signaling: Ebi/ AP-1 complex (Activator Protein 1) represses pro-/anti-apoptotic genes; suppresses basal transcription levels of apoptotic genes protecting sensory neurons degeneration (Lim et al., 2012); regulates EGFR (Dong et al., 1999), Notch (Nguyen et al., 2016; Marygold et al., 2011; Tsuda et al., 2002); Wg signaling; contributes to multiple processes including wing growth, eye development, regulation of transcription & innate immune response. |
| nclb | **TORC; MYC** | **no child left behind**; chromatin DNA-binding (Casper et al., 2011); pol I & pol III transcription (Liu et al., 2017). |

**Table A. Supplement to Figure 13: Signaling pathways linked to the Myc-CRM P1/P2**

| Gene name | Pathway | Activity/Function |
|---|---|---|
| PpD3 | **JNK; MAPK; CECY** | **Protein phosphatase D3**, conserved protein, involved in JNK and MAPK pathways (Miskei et al., 2011; Morrison et al., 2000); required for mitosis (G2/M progression). |
| Su(fu) | **HH; DPPBMP** | **Suppressor of fused**, fused (serine/threonine-protein kinase; negative regulator of Hedgehog signaling by forming a complex with the transcription factor Cubitus interruptus (Ci) (Han et al., 2019; Han et al., 2015; Smelkinson et al., 2007; Zhang et al., 2011; Fukumoto et al., 2001; Methot and Basler, 2000); negative regulator of Dpp pathway (Jia et al., 2002). |
| Ced-12 | **MAPK; TORC; JNK; PVR; RTK** | **Ced-12 (engulfment and cell motility)**, Pvr signaling: PDGF/VEGF (Platelet-Derived Growth Factor/Vascular Endothelial Growth Factor)-receptor related (Pvr), RTK activated by the binding of PDGF- & VEGF-related factors (Pvf1, Pvf2 or Pvf3), activates canonical Ras/Raf/MAPK (ERK) cascade, PI3K kinase pathway, TORC1 (FBrf0222697), RTK: Rho family small GTPases (Sopko & Perrimon, 2013; Ishimaru et al., 2004); JNK cascade (Sopko & Perrimon, 2013; Tran et al., 2013; Ishimaru et al., 2004). |
| CNBP | **MYC** | **CNBP/MYC** axis; Involved in the control of wing size by regulating Myc levels (Antonucci & Ganettieri, cc-2014). |
| rb | **NOTC; VTRA** | **Ruby**, *Drosophila* HOPS & AP-3 complex genes are required for a Deltex-regulated activation of Notch in the endosomal trafficking pathway, Notch receptor processing (Wilkin et al., 2008). |
| Larp4B | **MYC; CECY** | **La-related protein Larp4B**, negative regulation of cell growth and translation (Funakoshi et al., 2018). |
| Dci | **EIWE; APOP; IMD** | **Dodecenoyl-CoA delta-isomerase**, Eiger/Wengen signaling pathway, IMD signaling, apoptosis signaling (Berkey et al., 2009). |
| Doa | **TOLL; NFKB; MAPK; TORC; CECY; AUTO** | **Darkener of apricot (105 kDa Doa protein kinase)**, positive regulators of Toll NF-κB signaling pathway (Kanoh et al., 2015); MAPK cascade (Yun et al., 2000); TORC1 signaling inhibits CDK8 & DOA kinases, which directly phosphorylate CPSF6, a component of the CPA complex. These phosphorylation events regulate CPSF6 localization, RNA binding, & starvation-induced alternative RNA processing of transcripts involved in autophagy (Tang et al., 2018); negative regulation of male germ cell proliferation (Zhao et al., 2013). |

**Table A. Supplement to Figure 13: Signaling pathways linked to the Myc-CRM P1/P2**

| Gene name | Pathway | Activity/Function |
|---|---|---|
| Dhx15 | **MAPK** | **DEAH-box helicase 15**, MAPK/p38 MAPK cascade (Mosallanejad et al., 2014). |
| ca | **VTRA** | **Claret**, acts as a guanine nucleotide exchange factor (GEF) for Lightoid/Rab-RP1 in an adaptor protein 3-independent vesicular trafficking pathway of pigment granule biogenesis (Ma et al., 2004); Rab32 GEF/Claret involved in autophagy, affecting lipid storage (Wang et al., 2012). |
| how | **EGFR**; **MAPK; ERK; DPPBMP**; **HH; INR; WNBT; NOTC; RAS** | **held out wings**, integrin signaling, apposition of dorsal/ventral imaginal disc-derived wing surfaces (Walsh and Brown, 1998); cell adhesion (Lo and Frasch, 1997); EGFR signaling: glial cell migration; axon ensheatment (Edenfeld et al., 2006); MAPK signaling: germline stem cell population maintenance (Monk et al., 2010), mesoderm spreading via suppression of alternate target gene (*miple1*) (Toledano-Katchalski et al., 2007); ERK1/2 (Nir et al., 2012) muscle development, Hedgehog signaling (Lobbardi et al.,2011); RNA binding, intercellular signal transduction & activation of RNA (STAR) proteins; Insulin receptor (INR), Notch, Ras, and Rho signaling pathways (Lasko, 2003); Dpp signaling pathway (Israeli & Volk, 2007). |
| nito | **WNBT** | positive regulator of canonical Wg-TCF signaling during wing disk & eye development (Jemc and Rebay, 2006; Chang et al, 2008). |
| CG30291 | **NFKB**; **ERK** | NF-κB pathway; ERK1/2 pathways; (Utreras & Kulkarni; 2011). |
| pum | **EGFR** | **Pumilio, isoform G**, negative regulation of epidermal growth factor receptor (EGFR) signaling pathway (Kim et al., 2012). |
| Fmr1 | **miRNA; piRNA; ERK; WNBT; INR** | **Fragile X messenger ribonucleoprotein 1**, miRNA pathway (Specchia et al., 2017); MAPK/ERK (Leahy et al., 2023); Wg/Wnt pathway; Alk/Jeb pathway (The Jelly belly (Jeb)/ anterograde pathway involves presynaptic secretion of Jeb ligand to activate postsynaptic Anaplastic lymphoma kinase (Alk) receptors & drive phosphorylation of ERK (dpERK) resulting in modulation of both synaptic structure and function (Alk) (Friedman et al., 2013); negative regulation of insulin receptor signaling pathway (Luhur et al., 2017; Monyak et al., 2017; Callan et al., 2012). |

**Table A. Supplement to Figure 13: Signaling pathways linked to the Myc-CRM P1/P2**

| Gene name | Pathway | Activity/Function |
|---|---|---|
| IntS11 | **NOTC; EGFR; MYC** | **Integrator complex subunit 11**, Notch signaling (Shersher et.al., 2021); Epidermal growth factor pathway (Tilley & Mollet, 2021). |
| puf | **WNBT; MYC; TOLL; NFKB; IMD** | **Puffyeye**, Ubiquitin-Specific Protease (USP), an essential deubiquitinating enzyme, acts as a ubiquitin-specific protease, removes ubiquitin polypeptide chains from Myc & CycE, leading to stabilization and increases of their abundance during cell growth & proliferation (Li et al., 2013); TCF dependent signaling in response to WNT. |
| wap | **HIPP; ERK** | **wings apart**, negative regulation of Hippo Signaling (Degoutin et al., 2013); (ERK) pathway: involved in the control of organ growth and tissue patterning (Liu & Veraksa, 2016). |
| Gprk1 | **HH** | **G protein-coupled receptor kinase 1**, Hedgehog (Hh) signaling pathway: positive regulation of Smoothened signaling pathway (Cheng et al., 2010). |
| sti | **HIPP; DPPBMP; NOTC; TOLL; WNBT; EGFR; CECY** | **Sticky**, negative regulation of Hippo signaling pathway (Tran et al., 2019); involved in the regulating TGFb/BMP, Notch, Wingless, and EGFR pathways (Bivik et al., 2015); mitotic cytokinesis (Echard et al., 2004; Naim et al., 2004; Shandala et al., 2004; Rogers et al., 2003). |
| Mo25 | **WNK; TORC** | **Mo25, isoform B**, involved in neuroblast asymmetric cell division (Borkowsky et al., 2023); interactions between chloride & Mo25 regulate WNK [with no lysine (K)] kinases signaling in a transporting renal epithelium; WNK-SPAK/OSR1 (Rodan, 2018). |
| Vps26 | **WNBT; HH; DPPBMP** | **Vacuolar protein sorting 26**, regulation of Wnt, BMP, and Smoothened signaling (Port et al., 2008). |
| Sep1 | **APOP; DPPBMP; EGFR; NOTC** | **Septin-1**, positive regulation of apoptotic pathway (Bae et al., 2007); imaginal disc-derived wing morphogenesis (Sun et al., 2021); (TGF-b), epidermal growth factor (EGF), & Notch ligands govern vein patterning & differentiation (Sun et al., 2021; De Celis, 2003). |
| Idgf3 | **INR** | **Imaginal disc growth factor 3, isoform D**, member of chitinase-like protein family; involved in insulin signaling in imaginal disc cell lines (Kawamura et al., 1999). |

**Table A. Supplement to Figure 13: Signaling pathways linked to the Myc-CRM P1/P2**

| Gene name | Pathway | Activity/Function |
|---|---|---|
| sxc | **FGFR** | **Super sex combs**, belongs to Polycomb group, encodes a O-GlcNAc transferase, involved in epigenetic gene silencing; positive regulation of fibroblast growth factor receptor signaling pathway FGFR (Mariappa et al., 2011). |
| Rap1 | **HIPP; RAS; ERK; SEVE; HH; TORS; JNK** | **Ras-related protein Rap1**, negative regulation of hippo signaling (Chang et al., 2018); positive regulation of ERK1/2 signaling; regulation of Torso signaling (Mishra et al., 2005); positive regulation of sevenless signaling (Baril et al., 2014; Mavromatakis and Tomlinson, 2012; Marada et al., 2016); positive regulation of Smoothened signaling; JNK signaling (Boettner et al., 2003). |
| Septin5 (Sep5) | **CECY; APOP; DPPBMP; EGFR; NOTC** | **Septin5**, a member of the septin family of GTP-binding proteins; involved in cytokinesis; cell polarity & membrane rigidity; GTPase activity (GO Reference Genome Project, 2011-); imaginal disc-derived wing morphogenesis (Sun et al., 2021); involved in vein patterning & differentiation via modulation of TGF-β, Epidermal Growth Factor (EGF), & Notch ligands (Sun et al., 2021; De Celis, 2003). positive regulation of apoptosis pathway (Bae et al., 2007); regulation of cell cycle (O'Neill and Clark, 2013). |
| Sara | **NOTC** | A component of Notch signaling (Montagne and Gonzalez-Gaitan, 2014). |
| gukh | **RAS** | **GUK-holder**, Wiskott-Aldrich syndrome protein family member, Ras-mediated signaling pathways (Mathew et al., 2002; Chen et al., 1998). |
| dlg1 | **EGFR; JNK; ECDY** | **discs large 1**, regulation of epidermal growth factor receptor signaling pathway; JNK pathway: establishment or maintenance of polarity of larval imaginal disc epithelium (Bunker et al., 2015); Ecdysone Signaling: negative regulation of peptidoglycan recognition protein signaling pathway (Xiong et al., 2016). |
| Pak3 | **MAPK; RACP; RAS; ERK** | **p21-activated kinase-1 (Pak1)**, (non-specific serine/threonine protein kinase); an effector of Rho family GTPase Rac & Cdc42; Pak1 can influence signaling through the Ras/Erk pathway (Rac GTPase Pak kinase signaling: RACP) (Arias-Romero et al., 2010). |
| Mpcp2 | **NOTC** | **Phosphate carrier protein, mitochondrial**, Notch signaling wing disc D/V pattern formation (Bejarano et al., 2008). |

**Table A. Supplement to Figure 13: Signaling pathways linked to the Myc-CRM P1/P2**

| Gene name | Pathway | Activity/Function |
|---|---|---|
| CG11523 | **WNBT** | Kinase regulator activity and protein kinase A (PKA) binding activity; orthologous to human GSK3B interacting protein (GSKIP); regulation of canonical Wnt signaling pathway (InterPro Project Members, 2004-). |
| Amun | **NOTC** | **Amun, isoform A**, contains a putative DNA glycosylase domain; chaeta, compound eye & wing disc development (Shalaby et al., 2009). |
| wmd | **WNBT; DPPBMP; EGFR** | **wing morphogenesis defect**; imaginal disc-derived wing morphogenesis: TGF-beta, Epidermal Growth Factor Receptor (Dworkin and Gibson, 2006). |
| CG9231 | **APOP** | Cellular response to hypoxia; positive regulation of apoptosis; positive regulation of release of cytochrome c from mitochondria (Gene Ontology Curators, 2002-). |
| SkpA | **APOP; WNBT; HIPP; INR; JNK; IMD; CECY** | **SKP1-related A**, negative regulation of apoptosis (Fereres et al., 2013); negative regulation of Wnt-TCF signaling (Roberts et al., 2012); negative regulation of Hippo signaling (Tokamov et al., 2021; Zhang et al., 2015); negative regulation of Insulin receptor signaling pathway (Wong et al., 2013); negative regulation of JNK cascade signaling (Brace et al., 2014); Imd signaling: immune deficiency signaling cascade (Khush et al., 2002); mitotic cell cycle (Ducat et al., 2008). |
| His2Av | **IMD** | **Histone H2A variant**, Imd signaling negative regulation of peptidoglycan recognition protein signaling pathway (Tang et al., 2021). |
| LanB1 | **IMD; EGFR; JNK** | **Laminin B1**, Imd signaling Positive regulation of innate immune response (Cronin et al., 2009); Apoptosis pathway (Sun et al., 2021); JNK, imaginal disc eversion (Pastor-Pareja et al., 2004). |
| Cka | **RAS; MAPK; JNK; HIPP** | **Connector of kinase to AP-1**, Hippo (Neal et al., 2022; Neal et al., 2020; Pojer et al., 2021; Gil-Ranedo et al., 2019; Zheng et al., 2017; Liu et al., 2016; Ribeiro et al., 2010), JNK (La Marca et al., 2019; Ashton-Beaucage et al., 2014; Chen et al., 2002), and Ras/MAPK (Ashton-Beaucage et al., 2014) signaling; (Part of FAR/SIN/STRIPAK complex: Striatin-Interacting Phosphatase & Kinase Complex). |
| CG1943 | **NOTC** | Wing disc dorsal/ventral pattern formation via Notch signaling (Bejarano et al., 2008); orthologous to human JPT1 (Jupiter microtubule associated homolog 1). |

**Table A. Supplement to Figure 13: Signaling pathways linked to the Myc-CRM P1/P2**

| Gene name | Pathway | Activity/Function |
|---|---|---|
| kst | **EGFR; JNK; VTRA** | **Karst, isoform F**, EGFR signaling endosome transport via multivesicular body sorting pathway (Tjota et al., 2011); JNK signaling cascade in wound healing & epithelial sheath (Campos et al., 2010). |
| msk | **HH; DPPBMP; MAPK; NOTC; EGFR** | **Moleskin/Importin-7**, (Vrailas et al., 2006); Notch, EGFR, (Pepple et al., 2007), MAPK, Decapentaplegic (Dpp), & Hedgehog signaling pathways (Marenda et al., 2006). |
| Cdk1 | **HH; NOTC; HIPP; JAKSTAT; EGFR; WNBT; JNK; INR; CECY** | **Cyclin-dependent kinase 1**, G1/S transition of mitotic cell cycle (Lehner and O'Farrell, 1990); G2/S transition of mitotic cell cycle (Stern et al., 1993; Lehner and O'Farrell, 1990); follicle cell of egg chamber development, Notch, Hedgehog, EGFR, Wingless, JAK/STAT, Hippo, & JNK pathways; insulin-PI3K signaling pathway (Jia et al., 2015). |
| Gp150 | **NOTC** | **Glycoprotein 150, isoform E,** Transmembrane glycoprotein regulates Notch signaling, involved in compound eye development (Fetchko et al., 2002); transmembrane receptor protein tyrosine phosphatase signaling pathway (Tian and Zinn, 1994). |
| Cdc16 | **WNBT; CECY** | **Cell division cycle 16**, negative regulators of Wnt-TCF signaling; compound eye photoreceptor cell differentiation (Martins et al., 2017). |
| spag | **APOP** | **Spaghetti**, negative regulation of motor neuron apoptotic process (Means et al., 2015). |
| NfI (CG2380) | **JAKSTAT; WNBT** | **Nuclear factor I, isoform B**, CCAAT box-binding transcription factor (CTF) (Mermod et al., 1989); STAT3 signaling (Chen et al, 2017; Stringer et al, 2016); canonical Wnt signaling (High-mobility group AT-Hook 1 mediates the role of nuclear factor I/X in osteogenic differentiation via activating canonical Wnt signaling (Wu et al, 2021). |
| cype | **DPPBMP; HH** | **Cyclope**, encodes a cytochrome c oxidase subunit VIc homolog acting as an enhancer of dpp pathway phenotypes, involved in hair and cell growth, and in ommatidia development; regulation of Hedgehog signaling (Chang et al., 2001); significant alteration of *cype* gene product has been observed during the progression of prostate cancer (Herrmann, P. C., Petricoin III, E. F. et al., 2003). |
| Bap55 | **NOTC; EGFR** | **Brahma associated protein 55kD**, Notch signaling (Pillidge & Bray, 2019); response to EGFR signaling in the *Drosophila* wing (Terriente-Félix & de Celis, 2009). |

**Table A. Supplement to Figure 13: Signaling pathways linked to the Myc-CRM P1/P2**

| Gene name | Pathway | Activity/Function |
|---|---|---|
| 14-3-3ε | **HIPP; CECY; RAS; MAPK** | **14-3-3 protein epsilon**; key role in signal transduction pathways & cell cycle (DNA damage checkpoint signaling); interacts with kinases such as PKC or Raf-1; in plants associated with a complex that binds to the G-box promoter elements; positive regulators of Hippo signaling pathway (Pojer et al., 2021; Ren et al., 2010); Ras/MAPK (Ashton-Beaucage et al., 2014); DNA damage checkpoint signaling (Su et al., 2001; Brodsky et al., 2000). |
| glu | **CECY** | **Gluon**, a subunit of the multiprotein complex Condensin, mitotic cell cycle: prometaphase chromosome condensation (Bivik et al., 2015) & sister chromatid segregation; contributes to nervous system development & glucose metabolism; ubiquitous early, expressed in dividing cells throughout embryogenesis—pole cells, neuroblasts in the CNS & the PNS. |
| mod | **MYC; CECY** | **Modulo**, the *Drosophila* homologue of nucleolin; required for meiosis & spermatid differentiation in male germ line (Mikhaylova et al., 2006); MYC pathway (target of MYC selectively required for the growth of proliferative cells) (perinn et al., 2003); involved in chromatin packaging; dominant suppressor of variegation (Bantignies et al., 2002). |
| Hmg-2 (HMGB2) | **WNBT** | **High mobility group protein 2,** regulates chondrocyte hypertrophy by mediating Runt-related transcription factor 2 expression and Wnt signaling (Taniguchi et al., 2018); Wnt signaling and HMGB2 regulate articular cartilage surface maintenance (Taniguchi et al., 2009). |
| CRIF | **miRNA** | **CR6-interacting factor**, positive regulation of post-transcriptional gene silencing by RNA, positive regulation of siRNA production, RNAi pathway (Lim et al., 2014); growth arrest/DNA-damage-induced protein-interacting protein 1. |
| Wee1 | **CECY** | **Wee1-like protein kinase**, negative regulation of G2/M transition of mitotic cell cycle (Campbell et al., 1995). |
| Bin1 (SAP18) | **HH** | **Histone deacetylase complex subunit SAP18**, regulation of Hedgehog (Hh) signaling pathway by transcription factor Gli in mammals, repression of Gli-mediated transcription by Su(fu) for the recruitment of the SAP18- mSin3 complex to promoters containing the Gli-binding element (Yan Cheng and Bisho, 2002). |

**Table A. Supplement to Figure 13: Signaling pathways linked to the Myc-CRM P1/P2**

| Gene name | Pathway | Activity/Function |
|---|---|---|
| Rae1 | **HIPP; CECY** | **Rae1**, a nucleoporin member of the WD40-repeat β propeller protein super family; roles include poly(A)+ mRNA export, cell cycle regulation, male meiosis control & male germ cell post-meiotic differentiation; positive regulation of G1/S transition of mitotic cell cycle; Hippo signaling (Jahanshahi et al., 2016). |
| tsu | **MAPK; RAS** | **P-element somatic inhibitor, isoform C**, far upstream binding-element protein 1/2, C-terminal (FUBP1/2) (Davis-Smyth et al., 1996; Ni et al., 2020); dual roles in RNA processing (Labourier et al., 2002; Siebel et al., 1994) and transcriptional regulation; FUBP1required for the activation of MYC transcription by binding to a single-stranded-far upstream sequence element of MYC promoter (Duan et al., 2017; Duncan et al., 1994). |
| (psi) CG8912 | **MYC** | **P-element somatic inhibitor, isoform C**, dual roles in RNA processing (Labourier et al., 2002; Siebel et al., 1994) and transcriptional regulation. It is required for activating MYC transcription. KSRP (KHSRP) binds and destabilizes mRNA (Siebel et al., 1994). |
| Vig | **miRNA** | **vasa intronic gene**, involved in RNA interference; ncRNA-mediated post-transcriptional gene silencing; Fragile X-related protein and VIG associate with the RNA interference machinery (Caudy et al., 2002). |
| AGO2 | **miRNA** | **Argonaute 2**, interacts with small interfering RNAs (siRNAs) to form RNA-induced silencing complexes (RISCs), siRNA binding (Goh and Okamura, 2019, Kawamura et al., 2008; Tomari et al., 2007; Rand et al., 2005; Lingel et al., 2003); miRNA-mediated gene silencing (Besnard-Guérin et al., 2015). |
| l(3)72Ab (Brr2) | **DPPBMP; CECY; ECDY** | **Brr2 U5 snRNP complex subunit**, U5 small nuclear ribonucleoprotein 200 kDa helicase, brr2 and Prp8 control expression of FMRFa neuropeptide specifically in six neurons of the VNC (Tv4 neurons); brr2 control is executed by two independent mechanisms, both required for the activation of the BMP retrograde signaling pathway in Tv4 neurons: (1) proper axonal pathfinding to target tissue to receive BMP ligand. (2) proper RNA splicing of 2 genes in BMP pathway: thickveins (tkv) gene, encoding a BMP receptor subunit, & Medea gene (a co-Smad) (Ignacio Monedero Cobeta Benito-Sipos, 2018); mitotic cell cycle (Ducat et al., 2008); steroid hormone ecdysone signaling (Claudius et al., 2014). |

**Table A. Supplement to Figure 13: Signaling pathways linked to the Myc-CRM P1/P2**

| Gene name | Pathway | Activity/Function |
|---|---|---|
| Helz | **miRNA** | **Helicase with zinc finger**, involved in ncRNA-mediated post-transcriptional gene silencing (GO Reference Genome Project, 2011-). |
| Mtr4 | **INTERIM** | **Mtr4 helicase**, Interferon Immune signaling, RNA helicase involved in the viral defense response (Molleston et al., 2016). |
| Ddx1 | **MYC** | **DEAD box 1**, *Ddx1* depletion associated with small size & aberrant gametogenesis, possibly through alternative splicing of Sirup RNA transcript, oogenesis (Germain et al., 2015); MYC-associated protein X (Max) binds with stronger affinity to rs72780850, a genetic regulatory variant of DDX1 in neuroblastoma sample (Jin Y, Shi J, Wang H, Lu J, Chen C, Yu Y, Wang Y, Yang Y, Ren D, Zeng Q, Ni X, Guo Y. 2021); DEAD box 1: a novel and independent prognostic marker for early recurrence in breast cancer (Germain et al., 2011). |
| heph | **NOTC; DPPBMP** | **Hephaestus**, nucleo-cytoplasmic shuttling protein, regulates Oskar mRNA translation; spermatid individualization (Robida et al., 2010); regulation of Notch signaling (Dansereau et al., 2002); Dpp signaling, Squid & Hephaestus (but not Hrb27C) are necessary for proper bone morpho-genetic protein (BMP) signaling in GSCs; novel roles for RNA binding proteins Squid, Hephaestus, & Hrb27C in *Drosophila* oogenesis (Finger & Ables, 2022). |
| pod1 | **HIPP** | **pod1 coronin**, positive regulators of Hippo signaling pathway (Park et al., 2021). |
| unc-45 | **JAKSTAT** | **uncoordinated mutant number 45**, JAK/STAT signaling pathway controls host defense in the gut by regulating stem cell proliferation and thus epithelial cell homeostasis; negative regulation of innate immune response; defense response to Gram-negative bacterium (Cronin et al., 2009); involved in JAK/STAT pathway activity in a gradient-dependent manner during patterning of the anterior-posterior axis of the follicular epithelium (Xi et al., 2003). |
| gw | **miRNA** | **Gawky**, RNAi pathway (miRNA-mediated gene silencing pathway) (Chekulaeva et al., 2010; Chekulaeva et al., 2009; Eulalio et al., 2009; Eulalio et al., 2009; Zekri et al., 2009; Eulalio et al., 2008); miRNA induced silencing complex (Behm-Ansmant et al., 2006; Rehwinkel et al., 2005). |

**Table A. Supplement to Figure 13: Signaling pathways linked to the Myc-CRM P1/P2**

| Gene name | Pathway | Activity/Function |
|---|---|---|
| Ge-1 | **miRNA; NFKB; IMD; TOLL** | **Ge-1**, Enhancer of mRNA-decapping protein 4 homolog, RNAi pathway (miRNA-mediated gene silencing) (Eulalio et al., 2009); signaling pathways that activate NF-κB, Toll & Imd pathways (Jin et al., 2008; De Gregorio et al., 2002). |
| iPLA2-VIA | **KENN** | **Calcium-independent phospholipase A2 VIA**, Kennedy pathway (de novo synthesis pathway of Phospholipids for the rescue of ER stress via membrane remodeling), membrane remodeling supported by the regulation of neuronal functions & α-synuclein stability through Parkinson's disease-associated iPLA2-VIA/PLA2G6 (Mori & Hattoria, 2019). |
| awd | **FGFR; WNBT** | **abnormal wing discs**, *Drosophila* homolog of human gene (*Nm23*), crucial for follicle cells during oogenesis, loss-of-function (*awd*) mutant cells result in the accumulation & spreading of adherens junction components, such as E-cadherin, beta-Catenin/Armadillo, & Alpha-spectrin, & disruption of epithelial integrity (Woolworth et al., 2009); regulation of tracheal cell motility by modulating the FGFR levels (Dammai et al., 2003). |
| Mbo (Nup88) | **TOLL; NFKB** | **members only**, a nucleoporin involved in immune response transduction by mediating the nuclear translocation of Mad; Toll-NF-κB, antimicrobial humoral response (Uv et al., 2000). |
| Kcmf1 | **RAS; MAPK; ERK** | **Potassium channel modulatory factor 1**, negative regulation of the Ras/MAPK signaling pathway in the wing by acting with the E2 enzyme Unc6 and the putative E3 ligases Poe and Ufd4 to mediate the ubiquitination and proteasomal degradation of Rl/MAPK; negative regulation of ERK1 & ERK2 cascade (Ashton-Beaucage et al., 2016). |
| Cno | **HH**; **RAS**; **JNK** | **Canoe**, scaffold protein in adherens junctions, involved in morphogenesis in a variety of tissues; positive regulators of Hedgehog signaling pathway (Marada et al., 2016); Ras-association (RA) domains directly bind RanGTP & both the Canoe (RA) domains and RanGTP are required to recruit Mud to the cortex & activate the Pins/Mud/dynein spindle orientation pathway (Wee et al., 2011); Regulation of JNK cascade (Boettner et al., 2003). |
| Pi3K21B | **INR** | **Pi3K21B**, Insulin receptor signaling pathway (Breitkopf et al., 2016; Britton et al., 2002). |

**Table A. Supplement to Figure 13: Signaling pathways linked to the Myc-CRM P1/P2**

| Gene name | Pathway | Activity/Function |
|---|---|---|
| Rok, Drok | **WNPCP; CECY; RTK** | **Rho kinase**, *Drosophila* Rho kinase (Drok) (non-specific Ser/Thr protein kinase), activation by Rho1-dependent GTP, phosphorylation & modulation of cytoskeletal protein, particularly myosin II; dynamic regulation in subcellular locals influence cell polarization, movement, & cell shape (Dawes-Hoang et al., 2005) during interphase & mitosis; Fz/Dsh signaling (Fz/Dsh planar cell polarity pathway, cell adhesion & cell planar polarity (PCP) (Winter et al., 2001); mitotic cell cycle; mitotic spindle elongation (Hickson et al., 2006); mitotic cytokinesis (Tsankova et al., 2017; Dean and Spudich, 2006; Hickson et al., 2006; Dean et al., 2005); Rho protein signal transduction (GO Reference Genome Project, 2011-; Mizuno et al., 1999). |
| Dhc64C | **CECY** | **Dynein heavy chain 64C, isoform G**, centrosome-independent (acentrosomal) mitotic spindle assembly pathway; mitotic cell cycle (Maiato et al., 2004). |
| mask | **MAPK; JAKSTAT; PVR; SEVE; RTK** | **Multiple ankyrin repeats single KH domain**, mediator of RTK signaling & either downstream of MAPK or signal transducer through a parallel branch of the RTK; EGFR pathways, Sevenless signaling (Smith et al., 2002); positive regulation of receptor signaling via JAK/STAT (Fisher et al., 2018); vascular endothelial growth factor receptor signaling pathway (PVR signaling) (Tsai et al., 2022). |
| Sumo | **CECY; DPPBMP; RAS; MAPK; TOLL; NFKB; HH** | **Small ubiquitin-related modifier**, sole family member protein in *Drosophila*, required for embryonic patterning & mitosis (syncytial blastoderm mitotic cell cycle), also roles in wing patterning via Dpp (Miles et al., 2008), & Ras/MAPK signaling (Nie et al., 2009); localizes to the nucleus during interphase & to kinetochores & midbodies during mitosis; positive regulation of Smoothened signaling pathway (Zhang et al., 2017); regulation of Toll-NF-κB signaling pathway (Kanoh et al., 2015). |
| lin-28 | **INR; JAKSTAT** | **Protein lin-28 homolog**, cold shock and RNA-binding protein; regulator of developmental timing; regulator of microRNA maturation; INR (insulin/IGF signaling (IIS) pathway) (Luhur et al., 2017; Chen et al., 2015); positive regulation of receptor signaling pathway via JAK/STAT signaling (Sreejith et al., 2019). |

**Table A. Supplement to Figure 13: Signaling pathways linked to the Myc-CRM P1/P2**

| Gene name | Pathway | Activity/Function |
|---|---|---|
| btz | **miRNA; EXJC** | **Barentsz (CASC3 in mammals) (other name MLN51)**, a component of Exon Junction Complex pathway (EXJC), recruited to spliced mRNAs to mark introns removal site; translation activator linking the EJC & the translation machinery, direct role in protein synthesis (Chazal et al., 2013); BTZ domain found on CASC3 (cancer susceptibility candidate gene 3 protein, also known as **MLN51= Metastatic Lymph Node 51)** also known as Barentsz (Btz); CASC3, component of EJC involved in post-transcriptional regulation of mRNA in metazoa; complex formed by association of 4 proteins (eIF4AIII, Barentsz, Mago, & Y14), mRNA, & ATP; BTZ wraps around eIF4AIII & stacks against the 5' nucleotide (Bono et al., 2006; Palacios et al., 2004); Barentsz (MLN51) overexpressed in breast cancer (Degot et al., 2002-); repression of MNL51 by brain-specific miR-128 (Wilkinson et al., 2011); mRNA binding; RNA binding (InterPro Project Members, 2004-). |
| Amph | **ARDAMM** | **Amphiphysin**, Amphiphysin-Rho1-Dia/DAAM-Rok pathway: an actomyosin clamp assembled by the Amphiphysin-Rho1-Dia/DAAM-Rok pathway reinforces somatic cell membrane folded around spermatid heads (Kapoor et al., 2021); (DAAM: Diaphanous and Dishevelled Associated Activator of Morphogenesis). |
| spdi | **WNBT** | **split discs**, integrin signaling pathway in cell-migration & adhesion (Saadi et al., 2011). |
| l(2)gl | **NOTC; WNPCP** | **Lethal(2) giant larvae**, tumor suppressor, regulation of cell polarity & asymmetric cell division, acting on basolateral side of epithelial cells; antagonist of the activity of apical complex proteins Bazooka (Par-3), Par-6 & protein kinase C (aPKC); negative regulation of Notch (Johnson et al., 2016; Atwood and Prehoda, 2009); positive regulation of Hippo signaling (Parsons et al., 2014; Grzeschik et al., 2010); non-canonical Wnt signaling (planar cell polarity via Fz/Dsh); association with Dsh & regulation of adherence junction location, a process regulated by Wnt–Fz–Dsh signaling in vertebrates (Dollar et al., 2005; Na et al., 2007; Yamanaka and Nishida, 2007). |
| tws | **WNBT**; **HH; NFKB; CECY** | **Twins**, a regulatory subunit of protein phosphatase 2A (PP2A), involved in many developmental processes & signaling pathways (Bajpai et al., 2004); Smoothened signaling pathway (Su et al., 2011); positive regulator of Toll-NF-κB signaling pathway (Kanoh et al., 2021); mitotic cell cycle (Chen et al., 2007). |

**Table A. Supplement to Figure 13: Signaling pathways linked to the Myc-CRM P1/P2**

| Gene name | Pathway | Activity/Function |
|---|---|---|
| Idgf2 | **APOP; INR; IMD; JAKSTAT** | **Imaginal disc growth factor 2**, negative regulation of apoptotic process (Broz et al., 2017); insulin receptor pathway; (Varela et al., 2002); Imd and JAK/STAT pathways: The Imaginal Disc Growth Factors 2 and 3 participate in *Drosophila* immune response to nematode infection (Shruti, 2018). |
| Lig | **JAKSTAT; HIPP** | **Lingerer**, negative regulation of receptor signaling pathway JAK/STAT (Baumgartner et al., 2013); positive regulation of Hippo signaling pathway (Dong et al., 2015). |
| (sw) (CG18000) (E8NH77) | **CECY** | **short wing**, non-catalytic intermediate chain subunit cytoplasmic dynein motor complex; contributes to wing, eye, and oocyte development & polarity; mitotic cell division (Dzhindzhev et al., 2005); neuronal transport & neurogenesis; *Drosophila* Ser/Thr (MAST) kinase Drop interacts genetically with components of the dynein/dynactin complex (Hain et al., 2014). |
| Zip (Q99323) | **JNK; NOTC; WNPCP; CECY** | **Zipper**, Myosin heavy chain, non-muscle (myosin ATPase), JNK signaling, right-side-specific rearrangement of asymmetric morphogenetic cell, left/right axis specification (Okumura et al., 2010); Notch-dependent epithelial fold determines boundary formation between developmental fields in the *Drosophila* antenna. (Ku and Sun, 2017); *Drosophila* Rho-associated kinase (Drok) links Frizzled-mediated planar cell polarity signaling to the actin cytoskeleton (Wnt signaling via Fz/Dsh signaling) (Winter et al., 2001); cytoskeleton-dependent cytokinesis (Sechi et al., 2020). |
| Nurf-38 | **ECDY** | **Nucleosome remodeling factor - 38kD**, ecdysone receptor-mediated signaling pathway (Badenhorst et al., 2005). |
| Cul4 | **HH** | **Cullin 4, isoform A**, molecular scaffold for the CRL4 E3 ubiquitin ligase complex, catalyzes the ubiquitylation & subsequent destruction of proteins functioning in cell growth, proliferation, transcription, replication & repair of the genome; SCF-dependent proteasomal ubiquitin-dependent protein catabolic process; cellular response to DNA damage stimulus (GO Reference Genome Project, 2011-); negative regulation of Smoothened signaling pathway (Li et al., 2018); protein ubiquitination (GO Reference Genome Project, 2011-; Li et al., 2018; Reynolds et al., 2008). |

**Table A. Supplement to Figure 13: Signaling pathways linked to the Myc-CRM P1/P2**

| Gene name | Pathway | Activity/Function |
|---|---|---|
| chchd3 | **HIPP; JNK** | **Coiled-coil-helix-coiled-coil-helix domain containing 3**, Loss of (*chchd3*) leads to inactivation of Hippo activity, decreased tissue growth, cell proliferation defects, oxidative stress & JNK pathway activation (Deng et al., 2016). |
| HDAC1 | **HH** | **Histone deacetylase 1**, Hedgehog signaling (Zhang et al., 2013). |
| Mi-2 | **NOTC; WNBT; ECDY** | **Mi-2**, Notch signaling (Zacharioudaki et al., 2019); wingless, ecdysone signaling (Kon & Nusse; 2005). |
| udt | **JNK; JAKSTAT** | **Undicht**, wound healing, JNK signaling cascade, transduced by JUN/FOS transcriptional complexes (Ramet et al. 2002; Li et al. 2003; Ting et al. 2003, 2005a,b; Galko and Krasnow 2004; Mace et al. 2005; Campos et al., 2010); involvement of the JAK/STAT signaling cascade in this regenerative process (Mesilaty-Gross et al. 1999). |
| wds | **NOTC; CECY; APOP** | **Will die slowly, isoform B**, Trithorax-related complex protein (Hollmann et al, 2002), *Drosophila* essential gene codes for a WD-repeat protein with seven repeats; WD40 repeat implicated in a variety of functions ranging from signal transduction & transcription regulation to cell cycle control & apoptosis; positions the N-terminus of histone H3 for efficient trimethylation at 'Lys-4' (Pascual-Garcia et al., 2014); Mad interacts with the Wds to maintain active transcription by dynamically demethylating intragenic 6mA; maintenance of transcriptional activation for specific sets of genes in neurons together with DNA N6-methyl adenine demethylase Tet (yao et al., 2018); ATAC Complex, TRR complex, COMPASS Complex, TRX Complex. |
| Usp7 | **HIPP; HH** | **Ubiquitin-specific protease 7**, negative regulation of Hippo signaling pathway (Sun et al., 2019); positive regulation of Smoothened signaling pathway (Zhou et al., 2015). |
| Bap111 | **WNBT**; **HH**; **DPPBMP**; **NOTC** | **Brahma associated protein 111kD**, extensive homology of the Brahma (BRM) complex to SWI/ SNF; Osa/Eyelid (*osa*) shows a strong genetic interaction with (*brm*), suggesting close cooperation with the BRM complex (Treisman et al. 1997; Vazquez et al. 1999); (Wnt/TCF); Hh; Dpp; Notch signaling: Eyelid antagonizes wingless signaling during *Drosophila* development and has homology to the Bright family of DNA-binding proteins (Treisman et al., 1997; Kal et al., 2000). |

**Table A. Supplement to Figure 13: Signaling pathways linked to the Myc-CRM P1/P2**

| Gene name | Pathway | Activity/Function |
|---|---|---|
| barr | **CECY; HH; WNBT; NOTC; DPPBMP** | **Barren**, chromatin binding protein, involved in chromatin condensation; Malpighian tubule development & epithelial morphogenesis; Tube development (Liu et al., 1999): Hedgehog, completion of bud evagination (Hoch and Pankratz 1996), Wingless is required for cell division & morphogenesis in the tubules (Skaer and Martinez Arias 1992; Harbecke and Lengyel 1995); Notch receptor is required to define the single tip cell at the end of each tubule (Hoch et al. 1994), and elongation of the tubule (Skaer 1989), EGF (EGF-like) ligand required for the proliferation of the distal cells of the tubule (Baumann and Skaer 1993; Kerber et al. 1998); Dpp expressed in foregut & hindgut, required for morphogenesis of both structures (Pankratz and Hoch 1995*;* Hoch and Pankratz 1996); Cell Cycle interphase (Lupo et al., 2001). |
| LKRSDH | **ECDY** | **Lysine ketoglutarate reductase/saccharopine dehydrogenase**, negative regulation of ecdysone receptor signaling pathway (Cakouros et al., 2008). |
| kis | **HH; NOTC; RAS** | **Kismet, isoform F**, a conserved function of the chromatin ATPase Kismet is in the regulation of Hedgehog expression (Terriente-Félix et al., 2011); Notch signaling pathway (Go and Artavanis-Tsakonas, 1998; Verheyen et al., 1996);( *kis*) mutations have been identified in genetic screens as modifiers of the Ras and Notch signal transduction pathways (Go and Artavanis-Tsakonas, 1998; Therrien.et al., 2000; Melicharek et al., 2010). |
| CG17202 | **MYC** | **c-Myc-binding protein homolog,** transcription coactivator activity; regulation of transcription, DNA-templated (GO Reference Genome Project, 2011-) stimulation of *c-Myc* transcription (Taira et al., 1998); associate of Myc 1; orthologous to human MYCBP (MYC binding protein). |
| rump | **CECY** | **Rumpelstiltskin**, mitotic cell cycle (Ducat et al., 2008). |
| Grip71 | **CECY** | **Grip71**, mitotic cell cycle (Ducat et al., 2008; Verollet et al., 2006). |
| CG6664 | **CECY** | Establishment of meiotic spindle orientation; spindle pole, condensed chromosomes (Gene Ontology Curators, 2002-). |
| bel | **ECDY; miRNA; APOP** | **Belle**, Ecdysone-mediated induction of salivary gland cell autophagy cell death (Ihry and Bashirullah, 2014); ncRNA-mediated post-transcriptional gene silencing (Ulvila et al., 2006); Mitotic cell cycle (Pek and Kai, 2011). |

**Table A. Supplement to Figure 13: Signaling pathways linked to the Myc-CRM P1/P2**

| Gene name | Pathway | Activity/Function |
|---|---|---|
| asun | **CECY** | **Protein asunder**, mitotic cell cycle (Lee et al., 2005). |
| fs(1)h | **RAS**; **HH** | **Homeotic protein female sterile** (**female sterile (1) homeotic**), Ras signaling: a multifunctional transcriptional regulator modulated by Ras (Florence & Faller, 2008) and Hedgehog (Hh) signaling pathways (Xiangdong et al., 2018; Bagley et al., 2014). |
| msi | **HIF; NOTC** | **Musashi**, negatively regulates the Hypoxia Inducible Factor (HIF) pathway, contributes to cell fate determination, as well as cellular response to normoxic/hypoxic conditions; negative regulation of translation, asymmetric cell fate determination by Notch signaling (Bardin et al., 2004). |
| CaBP1 | **APOP** | **Calcium-binding protein 1**, (Protein disulfide-isomerase A6 homolog), apoptotic pathway (Okada et al., 2012); response to endoplasmic reticulum stress (GO Reference Genome Project, 2011-); protein disulfide isomerase activity (InterPro Project Members, 2004-). |
| slf | **HEBI** | **Schlaff**, chitin binding protein involved in wing formation, substrate for Transglutaminase (Tg); a putative C-type lectin needed for the adhesion between the horizontal cuticle layers (Zuber&Moussian, 2019); cooperation with the heme-biosynthesis pathway to stabilize the distribution of the cuticle tyrosinated proteins, exemplified by Resilin, the tyrosinated proteins network needed for correct contact between chitin laminae within the procuticle & the epicuticle (Zuber & Moussian, 2019). |
| RanBPM | **WNBT; JAKSTAT; SAPA** | **Ran-binding protein M, isoform F**, interacts with Armadillo (beta-catenin) (Dansereau and Lasko, 2008); JAK/STAT signaling (Baeg et al., 2005); Sema3A/Plexin-A signaling (Togashi et al., 2006). |
| lark | **CECY** | **Lark**, an essential RNA-binding protein of the RNA recognition motif (RRM); mitotic cell cycle (Ducat et al., 2008). |
| pds5 | **CECY**; **TGFA** | **precocious dissociation of sisters 5**, interacts with wings apart-like (*wapl*) to form the releasin complex, enables sister chromatid separation at mitosis by removing the cohesin ring complex from chromosomes (Gause et al., 2010; Dorsett et al., 2005); influences gene activation & silencing through interactions with cohesin; also required to initiate and/or maintain sister chromatid cohesion; TGF-alpha (ATR/Mei-41 kinase) (Barbosa et al., 2007). |

**Table A. Supplement to Figure 13: Signaling pathways linked to the Myc-CRM P1/P2**

| Gene name | Pathway | Activity/Function |
|---|---|---|
| CG34417 | **TOLL; DPPBMP; HH** | Mesoderm development; involved in Toll, Dpp, and Hedgehog signaling pathways (Furlong et al., 2001). |
| Idgf4 | **INR** | **Imaginal disc growth factor 4**, stimulation of insulin-like peptides for proliferation, polarization & motility of imaginal disk cells; stabilizing the binding of insulin-like peptides to its receptor (Kawamura et al., 1999) through a simultaneous interaction with both molecules to form a multiprotein signaling complex |
| (zip) (A0A0B4JD95) | **CECY; NOTC; JNK; WNPCP** | **Zipper, isoform H**, microtubule-binding protein involved in cytoskeleton-dependent intracellular transport. Its knockout or knockdown does not affect overall oocyte growth during mid-oogenesis, but the oocyte marker (*orb*) appears to be less evenly localized in (*zip*) loss-of-function mutant egg chambers. mitotic cytokinesis (Dean et al., 2005; Rogers et al., 2003); Notch signaling-dependent eye-antennal disc morphogenesis, epithelial fold boundary formation determination between developmental fields in the *Drosophila* antenna (Ku and Sun, 2017); left/right axis specification via JNK signaling (Okumura et al., 2010); Fz/Dsh signaling (a newly defined Fz/Dsh cytoskeletal signaling pathway with the involvement of fly myosin VIIA) (Winter et al., 2001). The seven-pass transmembrane receptor frizzled (Vinson et; al., 1989) requires the downstream signaling protein Dishevelled (Dsh) (Kling-ensmith et al., 1994; Theisen et al., 1994; Krasnow et al., 1995). Dsh & Fz also participate in the Wingless (Wg)/Wnt signaling to regulate developmental events including cell proliferation and cell fate specification (re- viewed in Wodarz and Nusse, 1998). However, Wg & planar cell polarity (PCP) distinct pathways downstream of Dsh (Axelrod et al., 1998; Boutros et al., 1998). |
| Hrb27C | **JNK**; **PVR** | **Heterogeneous nuclear ribonucleoprotein at 27C**, positive regulation of border follicle cell migration (Mathieu et al., 2007); PVR (the *Drosophila* PDGF/VEGF receptor) signaling pathway. |
| gbb | **DPPBMP** | **(glass bottom boat)**, a BMP ligand in the TGF-beta/BMP family of dimeric signaling molecules, binds to a receptor complex to transduce signal through phosphorylation of Mad; maintenance of stem cell populations, control of cell fate specification, proliferation, synapse growth, & neuropeptide release (Anderson and Wharton, 2017; Jensen et al., 2009; Shimmi et al., 2005). |

**Table A. Supplement to Figure 13: Signaling pathways linked to the Myc-CRM P1/P2**

| Gene name | Pathway | Activity/Function |
|---|---|---|
| Vps4 | **CECY; EGFR** | **Vacuolar protein sorting 4**, cell cycle; EGFR: *Drosophila* Vps4 promotes Epidermal Growth Factor Receptor signaling independent of its role in receptor degradation (Legent et al., 2015); JNK signaling: Disruption of Vps4 and JNK function in *Drosophila* causes tumor growth (Rodahl et al., 2009). |
| Ns3 | **INR** | **Nucleostemin 3**, GTPase (Hartl et al., 2013); nuclear export of 60S ribosomal subunit by mediating the release of Nmd3 from the 60S ribosomal subunit after export into the cytoplasm (Hartl et al., 2013); regulator of body size; acts in serotonergic neurons to regulate insulin signaling receptor pathway & thus exerts global growth control (Kaplan et al., 2008). |
| Snx6 | **WNBT; RTK** | **Sorting nexin 6**, phosphatidylinositol-3-phosphate binding activity (Gene Ontology Curators, 2002-); positive regulation of Wnt protein secretion; Wnt/Wg signaling (Zhang et al., 2011); phagosome-lysosome fusion involved in apoptotic cell clearance (GO Reference Genome Project, 2011-). |
| pins | **RAS; CECY** | **partner of inscuteable**, Ras-association (RA) domains directly bind RanGTP & both the Canoe (RA) domains and RanGTP are required to recruit Mud to the cortex & activate the Pins/Mud/dynein spindle orientation pathway (Wee et al., 2011); mitotic spindle orientation (Siegrist and Doe, 2005); asymmetric cell division (Januschke and Gonzalez, 2010; Yu et al., 2003; Parmentier et al., 2000). |
| Ppat-Dpck | **COAS** | **Bifunctional Phosphopantetheine adenylyl-transferase - Dephospho-CoA kinase**: unclassified kinases; conserved CoA biosynthesis pathway; imaginal disc-derived wing morphogenesis; triglyceride homeostasis (Bosveld et al., 2008). |
| cindr | **CECY** | **CIN85 and CD2AP related: adaptor protein**, links cell surface junctions & adhesion proteins with multiple components of the actin cytoskeleton; regulates cytoskeletal dynamics, eye patterning & endocytosis, cooperates with Scraps (*scra*) to promote intercellular bridge stability during cytokinesis; Cindr/dCortactin promote endocytosis (Quinones et al., 2010); Cindr:ArfGAP:dArf6 regulatory complex conserved across species, both during development & to maintain homeostasis (Johnson et al., 2011); Cindr/Anillin control cytokinesis in cleavage furrow (Haglund et al., 2010). |

**Table A. Supplement to Figure 13: Signaling pathways linked to the Myc-CRM P1/P2**

| Gene name | Pathway | Activity/Function |
|---|---|---|
| Idgf6 | **ECDY; INR** | **Imaginal disk growth factor 6**, stimulation of proliferation, polarization & motility of imaginal disk cells, may act by stabilizing the binding of insulin-like peptides to its receptor through a simultaneous interaction with both molecules to form a multiprotein signaling complex (InterPro Project Members, 2004-); ecdysis, chitin-based cuticle (Pesch et al., 2016). |
| Lst8 | **TORC; INR** | **Lst8**, conserved TOR-binding protein; required for "CREB-regulated transcription coactivator2 (Crtc)-dependent regulation of cell growth based on genetic evidence; TORC2 signaling (Kuo et al., 2015; Wang et al., 2012); TORC1 signaling (Wang et al., 2012); Positive regulation of Insulin-like receptor signaling (Yang et al., 2006); Target of Rapamycin Complex 2 regulates cell growth via Myc in *Drosophila* (Kuo et al., 2015). |
| eIF4A | **DPPBMP; CECY** | **Eukaryotic translation initiation factor 4A**, an essential DEAD box RNA helicase protein & a canonical translation initiation factor; a component of the eIF4F cap-binding complex, essential for cap-dependent translation of mRNA; SMAD binding; negative regulation of BMP signaling pathway (Li and Li, 2006); mitotic cell cycle (Ducat et al., 2008). |
| Elp4C (G6907) | **CECY** | **Elongator complex protein 4**, establishment of mitotic spindle asymmetry (Planelles-Herrero et al., 2022); phosphorylase kinase regulator activity (Gene Ontology Curators, 2002-). |
| e(r) | **NOTC; CECY** | **Protein enhancer of rudimentary (ERH)**, evolutionarily highly conserved; regulation of pyrimidine biosynthesis, DNA replication, transcription, mRNA splicing, cellular proliferation, tumorigenesis, & the Notch signaling pathway (Tsubota et.al., 2011); Fission yeast homologue of ERH is implicated in meiotic mRNA elimination during vegetative growth (Hazra et al., 2020; Shichino et al., 2018; Sugiyama et al., 2016); involvement of ERH-SRPK1-SAFB proteins in mitotic phosphorylation of Lamin B receptor (LBR) at the nuclear matrix (Hazra et al., 2020; Shichino et al., 2018; Sugiyama et al., 2016); (SAFB, scaffold attachment factor-B; SRPK1, SR protein kinase 1; SR protein, conserved splicing protein containing domains with long repeats of Serine & Arginine residues). |

**Table A. Supplement to Figure 13: Signaling pathways linked to the Myc-CRM P1/P2**

| Gene name | Pathway | Activity/Function |
|---|---|---|
| Tailor | **miRNA** | **Tailor**, terminal uridylyl-transferase Tailor, RNA-mediated gene silencing: positive regulation of miRNA catabolic process (Lin et al., 2017); pre-miRNA processing (Reimao-Pinto et al., 2015). |
| abs | **NOTC; CECY** | **Abstrakt**, DEAD-box protein, regulates cell polarity in oocytes & embryos; downregulation of Notch signaling in asymmetric cell division in ganglion mother cell (GMC2-4a) in collaboration with Inscuteable (*Insc*) (Irion et al., 2004). |
| apolpp | **WNBT; HH** | **Apolipophorin**, signal transduction activity, required for glycophosphatidylinositol-linked morphogens Wingless (Wg) & Hedgehog (HH) function by acting as vehicles for the movement of (Wg) & (HH) (Pana´kova et al., 2005). |
| BAP155<br>moira (mor) | **EGFR**; **NFKB; CECY** | **Brahma associated protein 155 kDa**, imaginal disc-derived wing margin morphogenesis, EGFR (Terriente-Felix & de Celis, 2009); negative regulation of G1/S transition of mitotic cell cycle (Brumby et al., 2002); regulation of innate immune response via NF κB signaling (Bonnay et al., 2014). |
| Slik | **CECY; NOTC; EGFR** | **Sterile20-like kinase, isoform A**, mitotic spindle midzone (Carreno et al., 2008); protein serine/threonine kinase activity; coordinated regulation of epithelial morphology & proliferation (Hughes and Fehon, 2006); regulation of Notch and EGF receptor signaling pathways (Hughes and Fehon, 2006; Maitra et al., 2006). |
| 'fandango'<br>(fand) | **FGFR; WNBT; HH; NOTC; DPPBMP** | **Pre-mRNA-splicing factor syf1 homolog**, tracheal development, specifically splicing (*bnl*) transcripts resulting in the activation of the BNL-FGF pathway; Wnt/Wg, Hedgehog, and Notch/Delta (N/Dl) signaling pathways (Hartenstein et al., 1992); TGF-b family (Dpp in *Drosophila*, BMP-2/4 in vertebrates signaling pathways (Fishman and Chien, 1997; Harvey, 1996; Bodmer, 1995). |
| (A4V4A5)<br>Ran | **RAS; CECY; WNBT** | **GTP-binding nuclear protein Ran**, a member of the Ras superfamily (Koizumi et al., 2001); localizes around the microtubule spindle in vivo during mitosis in *Drosophila* embryos (Trieselmann & Wilde, 2002); nuclear import of β-Catenin into nucleus (Yokoya et al., 1999). |

**Table A. Supplement to Figure 13: Signaling pathways linked to the Myc-CRM P1/P2**

| Gene name | Pathway | Activity/Function |
|---|---|---|
| Cen | **CECY** | **Centrocortin**, asymmetrical Centrosomal localization during mitosis on spindles (Kao and Megraw, 2009); involved in regulation of embryonic cleavage furrow (Kao and Megraw, 2009). |
| Lis-1 | **CECY; DPPBMP** | **Lissencephaly-1 homolog**, required during several dynein- and microtubule-dependent processes such as nuclear migration during cell division, mitotic spindle formation and the removal of mitotic checkpoint proteins from kinetochores at the metaphase to anaphase transition (Siller et al., 2006; Liu & Steward, 1999); positive regulation of BMP signaling pathway (Chen et al., 2010). |
| BRWD3 | **JAKSTAT; ECDY** | **Bromodomain & WD repeat containing protein 3**, positive regulator of JAK/STAT signaling (Muller et al., 2005); RHOBTB2 GTPase cycle; Interleukin-7 signaling; chromatin modifying enzymes; member of bromodomain & WD repeat containing protein (BRWD) family; regulation of ecdysone & JAK/STAT signaling (Chen et al., 2015). |
| Map60 (CP-60) | **CECY** | **Microtubule-associated protein 60**, centrosome-localized in a cell cycle-dependent manner (Kellogg et al., 1995). |
| CtBP (isoform G) | **NOTC; EGFR; WNBT; TORC** | **C-terminal binding protein, isoform G**, corepressor targeting diverse transcription regulators; Hairy (*hry*)-interacting protein; Hairless (*H*) major antagonist of Notch during imaginal tissue development; silences targets of (N) by assembling a transcriptional repressor complex containing transcription factor Su(H) and general corepressors such as Groucho (*gro*) and CtBP (*CtBP*); negative regulator of Notch signaling (Smylla et al., 2019; Wolf et al., 2019; Berndt et al., 2017; Maier et al., 2013; Troost and Klein, 2012; Kurth et al., 2011; Maier et al., 2011; Lee et al., 2009; Nagel et al., 2005; Barolo et al., 2002; Go et al., 1998; Lyman and Yedvobnick, 1995); plays role in the downregulation of EGFR signaling and induction of cell death by hairless (Protzer et al., 2008); CtBP corepressor directly activates & represses Wingless/Wnt transcriptional targets in *Drosophila* (Bhambhani et al., 2011; Fang et al., 2006); positive regulator of Wnt/TCF pathway (Bhambhani et al., 2011; Zhang and Arnosti, 2011; Fang et al., 2006); negative regulation of canonical Wnt signaling pathway (Bhambhani et al., 2011; Fang et al., 2006); plays role in Wingless signaling pathway during the process of embryo segmentation (Chan et al., 2008); part of TORC remodeling complex; belongs to other CH-OH oxidoreductases, NAD or NADP acceptor. |

**Table A. Supplement to Figure 13: Signaling pathways linked to the Myc-CRM P1/P2**

| Gene name | Pathway | Activity/Function |
|---|---|---|
| woc | **ECDY** | **Without children, isoform B**, ecdysone biosynthetic process (Wismar et al., 2000); regulation of transcription by RNA pol II (Font-Burgada et al., 2008). |
| HmgD | **EGFR; ECDY; DPPBMP; WNBT** | **High mobility group protein D, isoform C & High mobility group protein C, isoform D**, EGFR signaling (Anan Ragab & Travers, 2006); ecdysone/ecdysteroid (Chen et al., 2008); Dpp signaling: HMGD binding to the Dpp-responsive enhancer of (*tinman*) as well as to the Tinman protein during *Drosophila* cardiogenesis (Zaffran, 2002); Wnt/TCF signaling (Archbold et al., 2014). |
| shep | **MAPK; ERK; VTRA** | **Protein Alan shepard**, MAPK/ERK pathway phosphorylated ERK (PERK) (Clement et al., 2013); Vesicular Trafficking: neuropeptide Sorting through regulated secretory pathway (dense core vesicles) (Chen et al., 2014). |
| Spt20 | **JNK**; **NOTC** | **SPT20, isoform C,** Spt-Ada-Gcn5-acetyltransferase (SAGA) complex; JNK and Notch signaling pathway (Weake et al., 2009). |
| REPTOR | **TORC** | **Repressed by TOR**, response to starvation; TORC1 signaling (Tiebe et al., 2015). |
| sun | **GPCR** | **Stunted,** activation of G-protein coupled receptor Methuselah (*mth*) in vitro, leading to increased intracellular Calcium ion levels; positive regulation of G protein-coupled receptor signaling pathway involved in regulating ageing; associated with longevity in *Drosophila* (Ja et al., 2009; Cvejic et al., 2004). |
| sqh | **CECY; EGFR** | **Spaghetti squash**, Calcium ion binding (InterPro Project Members, 2004-); required for cytokinesis (Karess et al., 1991); EGFR signaling (Foronda et al., 2012). |
| mri | **EGFR; RAS; APOP** | **Mrityu, isoform E**, involved in Ras/EGFR and apoptotic pathway during retinal development (Rusconi & Challa, 2007). |
| atms/Paf1 | **NOTC** | **Antimeros, RNA polymerase II-associated factor 1 (PAF1)**, alternative mRNA splicing pathway (Lence et al., 2016; Burnette et al., 1999); formation of RNA Pol II elongation complex; RNA pol II pre-transcription events; E3 ubiquitin ligases ubiquitinate target proteins; Notch signaling (Lence et al., 2016; Hongay & Orr-Weaver, 2011). |

**Table A. Supplement to Figure 13: Signaling pathways linked to the Myc-CRM P1/P2**

| Gene name | Pathway | Activity/Function |
|---|---|---|
| net (CG11450) | **EGFR; DPPBMP; HH; WNBT** | **Net, isoform B**; E-box binding (GO Reference Genome Project, 2011-); imaginal disc-derived wing vein specification: EGFR signaling (Brentrup et al., 2000); imaginal disc-derived wing vein morphogenesis (De Celis, 2003); Notch signaling: the expression of E(spl)mβ and intervein-promoting genes *net*, *plexus*, and *bs* occur in the same domains as with the *iro* and *kni* during provein initiation; regulation of Dpp, Hedgehog, and Wingless signaling pathways (De Celis, 2003). |
| 14-3-3zeta | **RAS; MAPK; HIPP; CECY** | **14-3-3 protein zeta**, Ras/Raf/MAPK signaling-dependent photoreceptor development (Kockel et al., 1997); regulation of Yorkie nuclear localization by Hippo (Hpo) signaling pathway (Zhang & Jiang, 2010). |
| Cbp80 | **miRNA** | **Cap binding protein 80**, pre-mRNA splicing and RNA-mediated gene silencing (RNAi); miRNA-mediated RNA interference via its interaction with Ars2, required for primary microRNAs (miRNAs) processing (Sabin et al., 2009). |
| aub | **miRNA**; **JAKSTAT** | **Aubergine**, piRNA binding (Huang et al., 2021; Webster et al., 2015; Nagao et al., 2010); global gene silencing by mRNA cleavage (Kennerdell et al., 2002); ncRNA-mediated post-transcriptional gene silencing (Tomari et al., 2004); RNA-mediated gene silencing (Bozzetti et al., 2015); JAK/STAT signaling pathway controls host defense in the gut by regulating stem cell proliferation & epithelial cell homeostasis (Cronin et al., 2009). |
| Pka-C1 | **HH; HIPP; APOP; ERK; MAPK** | **cAMP-dependent protein kinase catalytic subunit 1**, negative regulation of Smoothened signaling pathway (Ranieri et al., 2014; Jia et al., 2002); positive regulation of Hippo signaling pathway (Yu et al., 2013); regulation of apoptotic pathway (Christiansen et al., 2013); plays role in neuronal plasticity via ERK/MAP kinase signaling (impey et al., 1999). |
| Vps29 | **WNBT** | **Vacuolar protein sorting 29**, Wnt ligand biogenesis & trafficking, retrograde transport endosome to Golgi (Franch-Marro et al., 2008; InterPro Project Members, 2004-). |
| mre11 | **CECY** | (**meiotic recombination 11**) double-strand break repair protein, G2/M DNA damage checkpoint signaling (Bi et al., 2006); telomere maintenance (Gao et al., 2009; Bi et al., 2005; Bi et al., 2004; Ciapponi et al., 2004). |

**Table A. Supplement to Figure 13: Signaling pathways linked to the Myc-CRM P1/P2**

| Gene name | Pathway | Activity/Function |
|---|---|---|
| shg | **PVR; EGFR; RTK; WNBT; JAKSTAT** | **Shotgun**, DE-cadherin (a major homophilic cell-cell adhesion molecule, inhibits motility of individual cells on matrix; functions downstream of the two chemoattractant receptors, PVR & EGFR to promote BC adhesion between the leader cells of the migrating cluster & the surrounding nurse cells; Receptor Tyrosine Kinase (Rac1/Rho-GTPase) activity (Cai et al., 2014); β-Catenin/Armadillo complex (cadherin-catenin cell-adhesion complex) (Menzel at al., 2008); β-Catenin binding (Huber et al., 2001); Wingless signaling & the control of cell shape in *Drosophila* wing imaginal discs (Widmer & Dahmann, 2009); JAK/STAT signaling cascade regulator & activator of molecules of the actin cytoskeleton, cell adhesion, & cell polarity (Lovegrove et al., 2006). |
| Cul1 | **CECY; INR; RTK; TORC; APOP; HH; IMD; HIPP; WNBT** | **Cullin homolog 1**, core component of multiple SCF (SKP1-CUL1-F-box protein) E3 ubiquitin-protein ligase complexes; ubiquitination of proteins involved in cell cycle progression (Wang et al., 2014), signal transduction & transcription; regulation of neuronal pruning through INR/PI3K/TOR pathway (Wong et al., 2013); apoptotic pathway during sperm differentiation (Bader et al., 2010); Regulation of Ci-SCF-Slimb binding, Ci proteolysis, & Hedgehog pathway activity by Ci phosphorylation (Smelkinson et al., 2007); repression of Imd immune response pathway by the ubiquitin-proteasome system (Kush et al., 2002); negative regulation of Hippo signaling (Tokamov et al., 2021; Zhang et al., 2015); negative regulation of canonical Wnt signaling (Roberts et al., 2012; Swarup & Verheyen, 2011). |
| thoc6 | **CECY** | **THO complex subunit 6**, linking transcription elongation with several cellular processes including mitotic recombination, co-transcriptional nuclear mRNA export, and export of heat shock mRNAs (Rehwinkel et al., 2004; Aguilera, 2002). |
| Galphas | **MAPK; WNBT; GPCR; WNPCP** | **G protein alpha s subunit, isoform E**, initiation of cAMP signaling; MAPK signaling; Wingless signaling; G-Protein coupled Receptor (GPCR) signaling cascade; non-canonical Frizzled signaling in the development of the fly pupal wing (Nichols et al., 2013). |

**Table A. Supplement to Figure 13: Signaling pathways linked to the Myc-CRM P1/P2**

| Gene name | Pathway | Activity/Function |
|---|---|---|
| Ank | **CECY; RTK; MAPK; MCAM** | **Ankyrin, isoform B**, recruited to the plasma membrane by its association with β-Spectrin (*β-Spec*) (Dubreuil & Yu, 1994), also binds to the adhesion protein Neuraglian (*Nrg*) leading to accumulation of Ankyrin & β-Spectrin at sites of cell-cell adhesion; asymmetric germline cell division; Receptor Tyrosine Kinase signaling pathway (Bennett & Baines, 2001); MAPK pathway (Whittard et al, 2006); (Higham et al., 2019; Cavaliere et al., 2013). |
| Vap33 | **EIWE; APOP; IMD** | **VAMP-associated protein 33kDa, isoform A**, plays a conserved role in synaptic homeostasis; defense response to Gram-negative bacterium: Eiger/ Wengen signaling pathway, Imd signaling, apoptosis signaling (Berkey et al., 2009); VAPB selectively triggers death of motoneurons through a Ca2+-dependent ER-associated pathway (Forrest et al., 2013; Langou et al., 2010); conserved phosphoinositide phosphatase Sac1 *Drosophila* VAP (DVAP)-binding partner, DVAP required to maintain normal levels of phosphoinositides; Vesicle-associated membrane protein (VAMP)-Associated Protein B (VAPB) in turn depends on phosphoinositide levels for its function. |
| CG8003 | **NOTC; WNBT** | Expressed in adult head and organism; orthologous to human ANKMY2 (ankyrin repeat and MYND domain containing 2). |
| Sirt4 | **INR** | **NAD-dependent protein deacylase Sirt4**, insulin signaling (Wood et al., 2018). |
| CG34159 | **MAPK** | MAPK-regulated corepressor-interacting protein; orthologous to human MCRIP2 (MAPK regulated corepressor interacting protein 2) (Gelbart and Emmert, 2013). |
| Chi | **NOTC; WNBT** | **Chip, isoform B**, LIM domain-binding protein 2; LIM domain-binding protein/SEUSS (LIM interaction domain); wing disc morphogenesis & leg development (Rincon-Limas et al., 2000); axon guidance (van Meyel et al., 2000). |
| Hrb87F | **WNBT; JNK** | **Heterogeneous nuclear ribonucleoprotein at 87F**, Wnt/wingless and JNK signaling (Yadav & Tapadia, 2016). |
| P32 | **MCAM** | **P32**, evolutionarily conserved mitochondrial protein, functions in presynaptic Calcium signaling & neurotransmitter release as well as chromatin metabolism, regulation of $Ca^{2+}$-mediated signaling (Lutas et al., 2012). |

**Table A. Supplement to Figure 13: Signaling pathways linked to the Myc-CRM P1/P2**

| Gene name | Pathway | Activity/Function |
|---|---|---|
| msn | **JNK; MAPK; NOTC; APOP; HIPP; TNFA** | **Misshapen, isoform A**, JNK activity (Martin and Wood, 2002); MAPK activity (InterPro Project Members, 2004-); negative regulation of Notch signaling pathway (Mishra et al., 2015); positive regulation of cell death; tumor necrosis factorα-mediated signaling pathway (Igaki et al., 2002; Moreno et al., 2002); positive regulation of Hippo signaling pathway (Li et al., 2015; Meng et al., 2015; Li et al., 2014). |
| Tm1 | **JNK; ERK; NFKB; DPPBMP; RAS; ERK** | **Tropomyosin**, in association with the troponin complex, plays a central role in the calcium dependent regulation of muscle contraction; response to hyperoxia (Zhao et al., 2010); JNK, ERK, NF-kappa B, and TGF-beta pathways play a role in the genesis of hyperoxia injury (Zhao et al., 2010); Ras signaling pathway (Zare et al., 2012); ERK pathway (Houle et al., 2007). |
| AP-2α | **NOTC; CECY; VTRA; JAKSTAT; WNBT** | **Adaptor Protein complex 2 α subunit**, Notch signaling, apico-basal polarity & neurogenesis; asymmetric cell division (Bardin et al., 2004); synaptic vesicle transport (Narayanan & Ramaswami, 2001; Gonzalez-Gaitan and Jackle, 1997); mitotic cleavage furrow ingression (Rodrigues et al., 2016); positive regulation of signal transduction by receptor internalization; WNT5A-dependent internalization of FZD2, FZD4**,** FZD5, and ROR2; integration of JAK/STAT receptor-ligand trafficking, signaling & gene expression in *Drosophila* melanogaster cells (Moore et al., 2020). |
| wapl | **CECY; WNBT** | **Protein wings apart-like**, together with the precocious dissociation of sisters 5 (pds5) protein form the releasing complex for separation of sister chromatid at mitosis by removing the cohesin ring complex from chromosomes; gene activation & silencing via interaction with Cohesin (Gause et al., 2010; Dorsett et al., 2005); regulation of Wnt/Wingless signaling pathway to determine the wing size in the Mediterranean fruit fly (medfly) (Cho et al., 2013). |
| Nup50 | **DPPBMP; CECY** | **Nuclear pore complex protein Nup50 (Nucleoporin 50kD)**, an important protein for TGF-beta signal transduction by mediating the nuclear translocation of Mothers against dpp protein (Mad); Association of 400 genes interacting with Nup50 Nucleoporin, among which transcriptionally active genes inside the nucleoplasm are predominantly involved in development & cell cycle regulation (Kalverda et al., 2010). |

**Table A. Supplement to Figure 13: Signaling pathways linked to the Myc-CRM P1/P2**

| **Gene name** | **Pathway** | **Activity/Function** |
|---|---|---|
| Rolled (rl) | **MAPK; RAS; EGFR; ERK; FGFR; INR; WNBT; CECY; SEVE; TORS; VEGFR; PVR** | **Mitogen-activated protein kinase ERK A**, core component of the RAS/MAPK pathway, is inactivated by the protein tyrosine phosphatase-ERK/Enhancer of Ras1 (PTP-ER) & Mitogen-activated protein kinase phosphatase 3 (Mkp3); phosphorylation of downstream cytoplasmic & nuclear effectors during cell fate decisions in a wide array of tissues; MAP kinase activity (Qiao et al., 2006; Baker et al., 2001; Karim and Rubin, 1999; Oellers and Hafen, 1996; Schweitzer et al., 1995; Brunner et al., 1994; O'Neill et al., 1994); EGFR signaling pathway (Ogura et al., 2018; Molnar and de Celis, 2013; Reich and Shilo, 2002; Yang and Baker, 2001; Kumar et al., 1998; Diaz-Benjumea and Hafen, 1994; Schnepp et al., 1998; Oellers and Hafen, 1996; Schweitzer et al., 1995); ERK1 & ERK2 cascade (Goyal et al., 2017; Ashton-Beaucage et al., 2014; Douziech et al., 2006; Roy et al., 2002; Clemens et al., 2000; Oellers and Hafen, 1996) FGFR signaling pathway (Du et al., 2018; Ukken et al., 2014; Csiszar et al., 2010; Gryzik and Muller, 2004; Petit et al., 2004; Ohshiro et al., 2002; Sato and Kornberg, 2002; Imam et al., 1999); insulin receptor signaling pathway (Slack et al., 2015; Xu et al., 2013; Kim et al., 2004; Clemens et al., 2000); lymph gland crystal cell differentiation, with the involvement of Wingless signaling (Dragojlovic-Munther & Martinez-Agosto, 2013; Sinenko et al., 2009); mitotic cell cycle (Marenda et al., 2006); Torso signaling pathway (Goyal et al., 2017; Lim et al., 1999; Ghiglione et al., 1999; Brunner et al., 1994); vascular endothelial growth factor receptor signaling pathway (Song et al., 2019; Sopko et al., 2015; Tsuzuki et al., 2014; Tran et al., 2013; Sims et al., 2009; Learte et al., 2008; Ishimaru et al., 2004; Cho et al., 2002; Duchek et al., 2001); PVR Signaling Pathway Core Components. |
| tral | **piRNA** | **Trailer hitch, isoform D**, piRNA pathway (Liu et al., 2011). |
| Tlk | **CECY**; **APOP; WNBT** | **Tousled-like kinase**, conserved anti-silencing function protein 1 (ASF1)/Tousled-like kinase (TLK), coordination of cell cycle phases via chromatin organization (Carrera et al., 2003); cell migration/apoptosis pathway (Zhang Y, Cai R, Zhou R, Li Y, Liu L.; 2016); association of Tousled-like kinase with the protein complex of wingless signaling regulators (Milan et al. 1998). |

**Table A. Supplement to Figure 13: Signaling pathways linked to the Myc-CRM P1/P2**

| Gene name | Pathway | Activity/Function |
|---|---|---|
| SMC2 | **CECY** | **Structural maintenance of chromosomes protein 2**, condensation of prometaphase chromosomes (GO Reference Genome Project, 2011-); condensation of prophase chromosomes; mitotic cell cycle (Ducat et al., 2008); regulation of microtubule assembly & organization in mitosis by the AAA+ ATPase Pontin, neurogenesis, stem cell differentiation (Liu et al., 2016). |
| CG10565 | **WNBT** | **DnaJ homolog subfamily C member 2**, chaperone J-domain superfamily; DnaJ domain; Homeobox-like domain superfamily; J-protein Zuotin/DnaJC2; Myb domain; ribosome-associated complex head domain superfamily; SANT/MYB domain; [Zuotin-related factor 1 (Zrf1) is necessary for chromatin displacement of the Polycomb-repressive complex 1 (PRC1); required for neural progenitor cells (NPC) specification from ESCs; promotes the expression of NPC markers, including the key regulator Pax6; essential to establish & maintain Wnt ligand expression levels, necessary for NPC self-renewal (Aloia et al., 2013). |
| piwi | **piRNA; CECY** | **P-element induced wimpy testis, isoform B**, piRNA-mediated metabolic process, repression of transposable elements during meiosis by complexes of piRNAs/Piwi containing proteins, methylation & subsequent repression of transposons; piRNA binding (Sienski et al., 2015); chromatin silencing (Brower-Toland et al., 2007; Grimaud et al., 2006; Pal-Bhadra et al., 2004; Pal-Bhadra et al., 2002); female germline stem cell asymmetric division; germline stem cell population maintenance; male germline stem cell asymmetric division (*Cox et al., 1998*); gene silencing by RNA (Le Thomas et al., 2013); heterochromatin organization involved in chromatin silencing (Sienski et al., 2015; Sienski et al., 2012). |
| bcn-3A | **VTRA; NOTC; WNBT** | **Rabconnectin-3A**, part of a regulatory subunit of the vacuolar H[+] ATPase (v-ATPase) required for acidification of intracellular vesicles & the lysosome; involved in endocytic trafficking & lysosome function (GO Reference Genome Project, 2011-; Yan et al., 2009); regulation of the Notch signaling pathway, mutations block Notch signaling in follicle cells & imaginal disc cells (Sethi et al., 2010; Yan et al., 2009); control of Wnt signaling during neural crest cells (NC) migration (Tuttle et al., 2014). |

**Table A. Supplement to Figure 13: Signaling pathways linked to the Myc-CRM P1/P2**

| Gene name | Pathway | Activity/Function |
|---|---|---|
| hop | **JAKSTAT; piRNA** | **Tyrosine-protein kinase hopscotch** (non-specific protein-tyrosine kinase) Tyrosine kinase of the non-receptor type; in combination with the Hsp83 & Piwi, mediates canalization, also known as developmental robustness, likely via epigenetic silencing of existing genetic variants & suppression of transposon-induced new genetic variations (Gangaraju et al., 2011; Binari and Perrimon, 1994); JAK/STAT signaling pathway (Bhaskar et al., 2022; Shen et al., 2022; Bailetti et al., 2019; Kallio et al., 2010; Bach et al., 2007; Muller et al., 2005; Wawersik et al., 2005; Johansen et al., 2003; Zeidler et al., 1999; Luo et al., 1997; Hou et al., 1996); tyrosine phosphorylation of STAT protein (Agaisse and Perrimon, 2004; Luo and Dearolf, 2001). |
| Pmm2 | WNBT | **Phosphomannomutase type 2**, synthesis of the GDP-mannose & dolichol-phosphate-mannose required for several critical Mannosyl transfer reactions (Parkinson et al., 2016); required for maintaining N-linked glycoprotein glycosylation at the neuromuscular junction (NMJ) synaptomatrix, & thus acts in multiple pathways that prevent NMJ structural overgrowth, restrict synaptic bouton differentiation, & limit NMJ neurotransmission strength, in order to maintain viability, coordinate movement, & in adults ensure correct wing positioning (Parkinson et al., 2016); acts in the NMJ trans-synaptic Wg pathway via glycosylation of synaptic Mmp2 which enables Dlp/Wg signaling during development (Parkinson et al., 2016). |
| Cp190 | **ECDY**; **WNBT** | **Centrosomal protein 190kD**, involved in the formation of most contact domain boundaries distal to a transcribed promoter; prevention of regulatory cross-talk between specific gene loci patterning the embryo; essential function during early development; response to ecdysone (Pascual-Garcia et al., 2017); Wingless signaling pathway: Cp190 found in complex with dTCF/Pan (pangolin), Su(Hw) and Pita (Marat Sabirov & Artem Bonchuk, 2021). |
| mbt | **WNBT; MAPK** | **mushroom body tiny**, Serine/threonine-protein kinase PAK (P21-activated kinase), association with Wingless pathway by modulating DE-cadherin mediated cell adhesion via phosphorylation of Armadillo (Menzel et al., 2008); regulation of MAP kinase cascade; protein phosphorylation (GO Reference Genome Project, 2011-; Schneeberger and Raabe, 2003); mushroom body development (Melzig et al., 1998). |

**Table A. Supplement to Figure 13: Signaling pathways linked to the Myc-CRM P1/P2**

| Gene name | Pathway | Activity/Function |
|---|---|---|
| SMC3 | **CECY; NOTC; WNPCP** | **Structural maintenance of chromosomes 3**, a subunit of the cohesin complex, involved in planar cell polarity by regulating the membrane enrichment of the transmembrane cadherin encoded by “starry night” (*stan*); mitotic sister chromatid cohesion; cell division (GO Reference Genome Project, 2011-); (; Notch and Wnt/Frizzled-PCP signaling during establishment of imaginal disc-derived wing hair orientation and imaginal disc-derived wing morphogenesis (Mouri et al., 2012). |
| Trl | **EGFR; DPPBMP** | **Trithorax-like**, a GAGA transcription factor involved in chromatin modification. It contributes to cell division, dosage compensation, and gametogenesis; Dpp & EGFR signaling, imaginal disc-derived wing morphogenesis (Dworkin and Gibson, 2006); ecdysteroid signaling (Pascual-Garcia et al., 2017); syncytial mitotic cell cycle (Bhat et al., 1996). |
| Hrb98DE | **PADPRI** | **Heterogeneous nuclear ribonucleoprotein at 98DE**, nuclear RNA-binding protein, controls hnRNA stability, splicing, IRES-dependent translation, & translational repression; represents one of the main targets of the poly(ADP-ribosyl)ation pathway; also regulates tissue polarity patterning & germ-line stem cell fate; negative regulation of RNA splicing (Ji and Tulin, 2009); post-translational modification of hnRNPs, such as poly(ADP-ribosyl)ation, is an important mechanism in the regulation of gene expression during development, such as eye pattern formation (Ji & Tulin, 2013). |
| Rcc1 | **CECY; WNBT; APOP** | **Regulator of chromosome condensation 1**, nuclear import & export of beta-Catenin (Koyama et at., 2017); regulation of mitotic cell cycle; apoptosis pathway (Trieselmann and Wilde, 2002). |
| Chro | **CECY; ECDY** | **Chromator**, a chromodomain protein, required for proper microtubule spindle formation, important for normal cell cycle progression, functioning as a spatial regulator of cell cycle factors (Ding et al., 2009); Metamorphosis (Wasser et al., 2007). |
| lid (Kdm5) | **JAKSTAT; CECY** | **Lysine-specific demethylase 5 (lid)**, activator of transcription (JAK/STAT) signaling; regulation of cell growth, circadian rhythm, stress resistance, hematopoiesis & fertility; male germline stem cell population maintenance (Tarayrah et al., 2015). |

**Table A. Supplement to Figure 13: Signaling pathways linked to the Myc-CRM P1/P2**

| Gene name | Pathway | Activity/Function |
|---|---|---|
| rin (CG9412) | **RAS; RTK** | **Rasputin**, evolutionarily conserved RNA-binding protein functions as a link between Ras signaling and RNA metabolism (Costa et al., 2013; Pazman et al., 2000); positive regulator of (*orb*) during *Drosophila* oogenesis; Rasputin (*Rin*), the *Drosophila* homologue of Ras-GAP SH3 Binding Protein (G3BP), associates with Orb in a messenger ribonucleoprotein complex (mRNP); Rho-mediated signaling (Pazman et al., 2000). |
| scny | **IMD; NFKB; RAS; NOTC** | **Scrawny**, Ubiquitin-specific protease 36; RAS processing (Ubiquitin carboxyl-terminal hydrolase 36); negative regulator of Imd/NF-kappa-B and (Imd) signaling; (Buszczak et al., 2009; Thevenon et al., 2009); somatic stem cell population maintenance; inhibition of H2B ubiquitylation within stem cells for the repression of premature expression of key differentiation genes, including Notch targets (Buszczak et al., 2009). |
| dgt4 | **CECY** | **dim γ-tubulin 4 (Augmin complex subunit dgt4)**, mitotic cell cycle, plays role in centrosome-independent generation of spindle microtubules (Goshima et al., 2008); mitotic spindle assembly (Goshima et.al., 2007). |
| rush | **VTRA** | **rush hour**, Pleckstrin homology domain-containing family F, an endosome-associated protein, directly binds phosphatidylinositol 3-phosphate (PI3P) & the Rab GDP dissociation inhibitor encoded by (*Gdi*); regulates endosomal trafficking by modulating the activity of Rab proteins (Gailite et al., 2012). |
| Klp10A | **CECY** | **Kinesin-like protein at 10A (Plus-end-directed kinesin ATPase)**, establishment of mitotic spindle asymmetry; centriole assembly; non-motile cilium assembly (Gottardo et al., 2016); centrosome duplication (Pavlova et al., 2019); establishment of meiotic spindle orientation; spindle assembly involved in female meiosis I (Zou et al., 2008); meiotic spindle organization (Radford et al., 2012); microtubule depolymerization; mitotic chromosome movement towards spindle pole (Goshima and Vale, 2005); mitotic spindle organization (Pavlova et al., 2019; Goshima and Vale, 2005); spindle organization (Morales-Mulia and Scholey, 2005). |
| Arpc3A | **RTK** | **Actin-related protein 2/3 complex, subunit 3A**, structural constituent of cytoskeleton, contributes to actin filament binding activity, involved in Arp2/3 complex-mediated actin nucleation; Receptor Tyrosine Kinases (RTK)-N-WASP/SCAR-Rho-GTPase-Arpc2,3 (Benesh et al., 2002). |

**Table A. Supplement to Figure 13: Signaling pathways linked to the Myc-CRM P1/P2**

| Gene name | Pathway | Activity/Function |
|---|---|---|
| CHOp24 | **WNBT** | Required for secretion of Wnt ligands (Port et al., 2011). |
| Orc6 | **CECY** | **Origin recognition complex subunit 6**, mitotic cell cycle (Balasov et al., 2009). |
| Poc1 | **CECY** | **Proteome of centrioles 1**, centrosome cycle (centrosome duplication and separation) (Blachon et al., 2009). |
| enc | **CECY** | **Encore** regulation of germline mitoses (Hawkins et al., 1996). |
| Trf2 | **ECDY** | **TATA box-binding protein-like 1**, ecdysone signaling (Bashirullah et al., 2007); ecdysteroid signaling (Shima et al., 2007). |
| sqd | **DPPBMP** | **Squid, isoform E**, Dpp signaling; Squid & Hephaestus (but not Hrb27C) are necessary for proper bone morphogenetic protein (BMP) signaling in GSCs. Novel roles for RNA binding proteins Squid, Hephaestus, and Hrb27C during *Drosophila* oogenesis (Finger et al., 2022). |
