## Supplemental Tables for "Targeting Regulatory Factors Associated with the *Drosophila Myc cis*-Elements by Reporter Expression, Gel Shift Assay, and Mass Spectrometric Protein Identification": Table B.pdf

| Signaling Pathways | Abbreviations |
|---|---|
| Apoptosis | APOP |
| Amphiphysin-Rho1-Dia/DAAM-Rok | ARDAMM |
| Autophagy (AUTO) | AUTO |
| Cell Cycle | CECY |
| CoA biosynthesis | COAS |
| DPP/BMP (TGF-β) | DPPBMP |
| Ecdysteroid | ECDY |
| Epidermal Growth Factor Receptor | EGFR |
| Eiger/ Wengen (EIWE) | EIWE |
| Ubiquitin-dependent ERAD | ERAD |
| ERK1/2 | ERK |
| Exon Junction Complex | EXJC |
| Fibroblast Growth Factor Receptor | FGFR |
| G-protein-coupled receptor Sun/Mth | GPCR |
| Heme-biosynthesis pathway | HEBI |
| Hepatocyte Growth Factor (HGF) | HGFS |
| Hedgehog/Smoothened | HH |
| Hypoxia Inducible Factor (HIF) | HIF |
| Hippo | HIPP |
| Immune deficiency (IMD) | IMD |
| Insulin-like Receptor | INR |
| interferon Immune | INTERIM |
| JAK/STAT | JAKSTAT |
| Jasmonate (JA) & Auxin | JASAU |
| Jun Kinase | JNK |
| Kennedy pathway | KENN |
| MAPK | MAPK |
| (Mitochondrial) Ca2+-mediated | MCAM |
| miRNA | miRNA |
| MYC | MYC |
| NF-κB | NFKB |
| Notch | NOTC |
| Poly(ADP-RIbosyl)ation | PADPRI |
| Platelet-derived Growth Factor Receptor | PDGFR |
| piRNA | piRNA |
| PVR (PDGFR/VEGFR) | PVR |
| Rac GTPase Pak kinase | RACP |
| RAS GTPase | RAS |
| Receptor Tyrosine Kinase (RTK) | RTK |
| Sema1,3A/Plexin-A | SAPA |
| Sevenless | SEVE |
| Sterol Regulatory-Element Binding protein | SREBP |
| TGFalpha (ATR/Mei-41 kinase) | TGFA |
| TNFα/Eiger-Mediated (TNFA) | TNFA |
| Toll | TOLL |
| TORC1,2 | TORC |
| Torso | TORS |
| Vascular Endothelial Growth Factor Receptor | VEGFR |
| Vesicular Trafficking | VTRA |
| Wingless (Wnt/β-Catenin/TCF) | WNBT |
| Wingless (Ca2+-mediated) | WNCA |
| With No Lysine kinase (WNK) | WNK |
| Wingless (Ca2+-mediated) | WNPCP |

**Table B. Supplement to Figure 13: Signaling pathways linked to the Myc-CRM P29/P30**

| Gene name | Pathway | Activity-Function |
|---|---|---|
| MED17 | **JASAU; RAS; MAPK** | **Mediator of RNA pol II transcription subunit 17**, sex comb development; required for adult cell & segment identity specification; SP1 transcription activation; thyroid hormone receptor complex component, VitD receptor interactor; jasmonate (JA) & Auxin signaling component (Boube et al., 2000); Ras/MAK pathway (Singh and Han 1995). |
| MED25 | **IMD; NFKB** | **Mediator of RNA poly II transcription subunit 25**, Imd pathway, NF-κB Signaling; positive regulation of antibacterial peptide biosynthetic process (Valanne et al., 2010). |
| pont | **JNK; TNFA; WNBT** | **RuvB-like helicase 1 (Pontin),** canonical Wingless/Wnt pathway (Torres & Inestrosa, 2017), interaction with HMG box transcription factors of the lymphoid-enhancing factor-1 (LEF-1/T-cell factor (TCF) family, positive regulation of canonical Wnt signaling pathway (Bauer et al., 2000); JNK pathway; negative regulation of tumor necrosis factor-mediated (TNFα-Eiger) signaling pathway (Wang et al., 2008). |
| E(bx) | **JAKSTAT** | **Enhancer of bithorax**, Component of Nucleosome Remodeling Factor NURF; negative regulation of receptor signaling pathway via JAK-STAT Signaling; spermatid differentiation; negative regulation in innate immune response; (Kwon et al., 2009); nuclear receptor binding (Badenhorst et al., 2005). |
| Snr1 | **EGFR; NOTC; DPPBMP** | **Snf5-related 1**, (ATP-dependent chromatin remodeler; a counterpart of yeast SWI/SNF) (Swi3 component of the Brahma complex), Notch signaling maintains neuroblast identity (Koe & Wang, 2014; Weng et al., 2010; Bowman et al., 2008; Wang et al., 2006); imaginal disc morphogenesis; EGFR signaling (Terriente-Félix and de Celis, 2009); Notch, EGFR, and DPP signaling important for wing development (Curtis et al., 2011). |
| Smr | **ECDY; INR; NOTC** | **Smrter**, Insulin-PI3K pathway (Jia et al., 2015); Notch pathway (Jia et al., 2015; Heck et al., 2012); ecdysteroid signaling (Heck et al., 2012). |
| CG17002 | **MAPK; RAS; JNK** | **G protein pathway suppressor 2**, suppresses a RAS- & MAPK-mediated signal and interferes with JNK activity (Cheng & Kao, 2009), suggesting that the function of this protein may be signal repression (pain BH, Bowdish KS, Pacal AR, Staub SF, Koo D, Chang CY, Xie W, Colicelli J., 1996). |

**Table B. Supplement to Figure 13: Signaling pathways linked to the Myc-CRM P29/P30**

| Gene name | Pathway | Activity-Function |
|---|---|---|
| Hrb27C | **JNK**; **PVR** | **Heterogeneous nuclear ribonucleoprotein at 27C**, positive regulation of border follicle cell migration (Mathieu et al., 2007); PVR (the *Drosophila* PDGF/VEGF receptor) signaling pathway. |
| CG8149 | **NOTC; PVR** | Notch signaling wing disc D/V patterning (Bejarano et al., 2008); PVR (*Drosophila* PDGF/VEGF receptor) signaling; larval lymph gland hemopoiesis (Mondal et al., 2014). |
| lid (Kdm5) | **JAKSTAT; CECY** | **Lysine-specific demethylase 5 (lid)**, activator of transcription (JAK/STAT) signaling; regulation of cell growth, circadian rhythm, stress resistance, hematopoiesis & fertility; male germline stem cell population maintenance (Tarayrah et al., 2015). |
| Su(fu) | **HH; DPPBMP** | **Suppressor of fused**, fused (serine/threonine-protein kinase; negative regulator of Hedgehog signaling by forming a complex with the transcription factor Cubitus interruptus (Ci); (Han et al., 2019; Han et al., 2015; Smelkinson et al., 2007; Zhang et al., 2011; Fukumoto et al., 2001; Methot and Basler, 2000); negative regulator of Dpp pathway (Jia et al., 2002). |
| pds5 | **CECY; TGFA** | **precocious dissociation of sisters 5**, interacts with wings apart-like (*wapl*) to form the releasin complex, enables sister chromatid separation at mitosis by removing the cohesin ring complex from chromosomes (Gause et al., 2010; Dorsett et al., 2005); influences gene activation / silencing through interactions with Cohesin; also required to initiate and/or maintain sister chromatid cohesion; TGF-alpha (TGFA) (ATR/Mei-41 kinase) (Barbosa et al., 2007). |
| sbb | **NOTC; HH** | **Scribbler**, transcriptional co-regulator, acts mainly as a co-repressor, interacts physically & genetically with Grunge (*Gug*) & is used by the repressor Tailless (*tll*) in early embryos, regulates larval behavior, photoreceptor axon target choice, and functions downstream of Hedgehog signaling; wing disc D/V pattern formation via Notch signaling (Bejarano et al., 2008). |
| Idgf2 | **APOP**; **INR**; **IMD**; **JAKSTAT** | **Imaginal disc growth factor 2**, negative regulation of apoptotic process (Broz et al., 2017); insulin receptor pathway; (Varela et al., 2002); Imd and JAK/STAT pathways: The Imaginal Disc Growth Factors 2 and 3 participate in the *Drosophila* response to nematode infection (Shruti, 2018). |

**Table B. Supplement to Figure 13: Signaling pathways linked to the Myc-CRM P29/P30**

| Gene name | Pathway | Activity-Function |
|---|---|---|
| Cpsf160 | **TORC** | **Cleavage and polyadenylation specificity factor 160**, TORC1 signaling, positive regulation of autophagy (Tang et al., 2018). |
| pzg | **NOTC; ECDY; JAKSTAT** | **Putzig**, involved in chromatin activation of replication related genes & signaling pathways including Notch, Ecdysone, & JAK/STAT; regulation of growth, cell death, and various developmental processes (Kugler et al., 2011). |
| Trf2 | **ECDY** | **TATA box-binding protein-like 1**, ecdysone signaling (Bashirullah et al., 2007); ecdysteroid signaling (Shima et al., 2007). |
| Su(var)2-10 | **JAKSTAT; HH; IMD** | **Suppressor of variegation 2-10, isoform L**, JAK/STAT pathway regulator, contributes to eye formation & eye determination (Betz et al., 2001); negative regulators of JAK/STAT signaling (Muller et al., 2005; Betz et al., 2001); positive regulators of Hedgehog signaling (Ma et al., 2016; Zhang et al., 2017); negative regulators of Imd signaling pathway (Tang et al., 2021; Cronin et al., 2009). |
| Rae1 | **HIPP; CECY** | **Rae1**, a nucleoporin member of the WD40-repeat β propeller protein super family; roles include poly(A)+ mRNA export, cell cycle regulation, male meiosis control, & male germ cell post-meiotic differentiation; positive regulation of G1/S transition of mitotic cell cycle, Hippo signaling (Jahanshahi et al., 2016). |
| ebi | **JNK; EGFR; WNBT; NOTC** | **F-box-like/WD repeat-containing protein ebi-like**, evolutionarily conserved repressor/silencer; JNK signaling: Ebi/ AP-1 complex (activator protein 1) represses pro-/anti-apoptotic genes; suppresses basal transcription levels of apoptotic genes protecting sensory neurons degeneration (Lim et al., 2012); regulation of EGFR pathway (Dong et al., 1999), Notch (Nguyen et al., 2016; Marygold et al., 2011; Tsuda et al., 2002); negative regulation of transcription (Lim et al., 2012); DNA binding transcription factor binding (Qi et al., 2008). |
| REPTOR | **TORC** | **Repressed by TOR**, response to starvation; TORC1 signaling (Tiebe et al., 2015). |
| jub | **HIPP** | **Ajuba LIM protein, isoform D**, Hippo signaling (Huang et al., 2016; Fletcher et al., 2015; Rauskolb et al., 2014; Sun and Irvine, 2013; Rauskolb et al., 2011; Das Thakur et al., 2010). |

**Table B. Supplement to Figure 13: Signaling pathways linked to the Myc-CRM P29/P30**

| Gene name | Pathway | Activity-Function |
|---|---|---|
| e(y)3 (SAYP) | **JAKSTAT** | **Enhancer of yellow 3, isoform D**, coactivator of JAK/STAT pathway (Shidlovskii et al., 2005); required for embryogenesis and oogenesis (Vorobyeva et al., 2009). |
| Rox8 | **HIPP** | Hippo signaling (Guo et al., 2020). |
| Dhx15 | **MAPK** | **DEAH-box helicase 15**, MAPK/p38 MAPK cascade (Mosallanejad et al., 2014). |
| asun | **CECY** | **Protein asunder**, mitotic cell cycle (Lee et al., 2005). |
| fs(1)h | **RAS**; **HH** | **Homeotic protein female sterile (female sterile (1) homeotic**), Ras signaling: a multifunctional transcriptional regulator modulated by Ras (Florence & Faller, 2008) and Hedgehog (Hh) signaling pathways (Xiangdong et al., 2018; Bagley et al., 2014). |
| Mad | **DPPBMP**; **WNBT; EGFR** | **Mothers against decapentaplegic**, BMP signaling pathway core components (Vuilleumier et al., 2022; Guo et al., 2013; Weiss et al., 2010; Kamiya et al., 2008; Yao et al., 2006; Muller et al., 2003; Dai et al., 2000; Das et al., 1998; Inoue et al., 1998); involved in wing development via modulation of EGFR and BMP signaling pathways (Dworkin and Gibson, 2006; Lecuit et al., 1996); negative regulation of salivary gland boundary formation with the involvement of Wnt/Wg signaling pathway (Bradley, P.L., Haberman, A.S., Andrew, D.J. 2001). |
| pum | **EGFR** | **Pumilio, isoform G**, negative regulation of epidermal growth factor receptor (EGFR) signaling pathway (Kim et al., 2012). |
| sti | **HIPP; DPPBMP; NOTC; TOLL; WNBT; EGFR; CECY** | **Sticky**, negative regulation of Hippo signaling (Tran et al., 2019); TGFb/BMP, Notch; Wingless; and EGFR signaling pathways (Bivik et al., 2015); mitotic cytokinesis (Echard et al., 2004; Naim et al., 2004; Shandala et al., 2004; Rogers et al., 2003). |
| Tlk | **CECY**; **APOP**; **WNBT** | **Tousled-like kinase**, conserved anti-silencing function protein 1 (ASF1)/Tousled-like kinase (TLK), coordination of cell cycle phases via chromatin organization (Carrera et al., 2003); cell migration/apoptosis pathway (Zhang Y, Cai R, Zhou R, Li Y, Liu L.; 2016); association of Tousled-like kinase with the protein complex of Wingless signaling regulators (Milan et al. 1998). |

**Table B. Supplement to Figure 13: Signaling pathways linked to the Myc-CRM P29/P30**

| Gene name | Pathway | Activity-Function |
|---|---|---|
| swm | **HH** | **(second mitotic wave missing)**, negatively regulates Hedgehog (Hh) protein signal in wing development (Casso & Karnberg, 2008). |
| IntS11 | **NOTC**; **EGFR**; **MYC** | **Integrator complex subunit 11**, Notch signaling (Shersher et.al., 2021); Epidermal Growth Factor Receptor signaling pathway (Tilley & Mollet, 2021). |
| Cdk1 | **HH; NOTC; HIPP; JAKSTAT; EGFR; WNBT; JNK; INR; CECY** | **Cyclin-dependent kinase 1**, G1/S transition of mitotic cell cycle (Lehner and O'Farrell, 1990); G2/S transition of mitotic cell cycle (Stern et al., 1993; Lehner and O'Farrell, 1990); follicle cell of egg chamber development, Notch, Hedgehog, EGFR, Wingless, JAK/STAT, Hippo, & JNK pathways; insulin-PI3K signaling pathway (Jia et al., 2015). |
| janA | **JAKSTAT** | **Janus A** (phosphohistidine phosphatase), sex regulated protein janus, JAK/STAT pathway (Myllym & Rämet, 2014); sex differentiation (Yanicostas et al., 1989). |
| Ulp1 | **HH; TOLL; NFKB** | **Ulp1**, negative regulators of Hedgehog signaling pathway (Zhang et al., 2017; Ma et al., 2016); negative regulators of Toll-NF-κB signaling pathway (Anjum et al., 2013). |
| awd | **FGFR; WNBT** | **(abnormal wing discs)**, *Drosophila* homolog of human Nm23, crucial for follicle cells during oogenesis, loss-of-function (awd) mutant cells result in the accumulation & spreading of adherens junction components, such as E-cadherin, beta-catenin/Armadillo, & alpha-spectrin, & disruption of epithelial integrity (Woolworth et al., 2009); awd regulates tracheal cell motility by modulating the FGFR level (Dammai et al., 2003). |
| for | **TOLL; NFKB** | **Foraging**, cGMP-dependent protein kinase, Toll-NF-κB Signaling (Kanoh et al., 2021); Protein Kinase G Family. |
| Pak3 | **MAPK; RACP; RAS; ERK** | **p21-activated kinase-1 (Pak1)**, (Non-specific serine/threonine protein kinase); an effector of Rho family GTPase Rac & Cdc42; Pak1 can influence signaling through the Ras/Erk pathway (Rac GTPase Pak kinase signaling, RACP) (Arias-Romero et al., 2010). |
| CG4115 | **IMD; NFKB** | Imd pathway (NF-kB signaling), carbohydrate binding (Tanji et al., 2006). |
| spag | **APOP** | **Spaghetti**, negative regulation of motor neuron apoptotic process (Means et al., 2015). |
| CG12384 | **APOP** | Apoptotic signaling pathway (GO Reference Genome Project, 2011-). |

**Table B. Supplement to Figure 13: Signaling pathways linked to the Myc-CRM P29/P30**

| Gene name | Pathway | Activity-Function |
|---|---|---|
| Amun | **NOTC** | **Amun, isoform A**, contains a putative DNA glycosylase domain; chaeta, compound eye, & wing disc development Shalaby et al., 2009). |
| wmd | **WNBT**; **DPPBMP; EGFR** | **(wing morphogenesis defect)**; imaginal disc-derived wing morphogenesis: TGF-beta, Epidermal Growth Factor Receptor (Dworkin and Gibson, 2006). |
| CG6607 | **VTRA** | **Coiled-coil domain-containing protein 128**, orthologous to human PPP1R21 (protein phosphatase 1 regulatory subunit 21); Rab protein signal transduction, early endosome, small GTPase binding (Gillingham et al., 2014). |
| Past1 | **NOTC** | **Putative Achaete Scute Target 1**, imaginal disc-derived wing vein specification (Olswang-Kutz et al., 2009). |
| Rpn13 | **HH** | **Regulatory particle non-ATPase 13**, positive regulation of Smoothened signaling pathway (Zhou et al., 2018). |
| SCAR | **RTK** | **SCAR**/WAVE complex pathway (Michael et al., 2013) lamellipodium and cell migration (Georgiou and Baum, 2010); actin filament organization (Sander et al., 2013). |
| scrib | **VTRA; EGFR; JAKSTAT; NOTC** | **Scribble**, scaffolding protein, part of conserved machinery regulating apicobasal polarity, interacts with "Discs large" (*dlg1*) & "lethal(2)giant larvae" (*l(2)gl*) to distinguish the basolateral domain of epithelial cells & neuroblasts via reciprocal antagonistic interactions with the aPKC/par-6 complex that impacts vesicle trafficking; EGFR, JAK, and Notch signaling pathways (Li et al., 2009). |
| sgl | **WNBT; DPPBMP; FGFR** | **Sugarless**, Wingless/Wnt signaling; Dpp signaling (Toyoda et al., 2000); Fibroblast Growth Factor Receptor Signaling pathway (Lin et al., 1999). |
| SkpA | **APOP**; **WNBT; HIPP; INR; JNK; IMD; CECY** | **SKP1-related A**, negative regulation of apoptosis (Fereres et al., 2013); negative regulation of Wnt-TCF signaling (Roberts et al., 2012); negative regulation of Hippo signaling (Tokamov et al., 2021; Zhang et al., 2015); negative regulation of Insulin receptor signaling pathway (Wong et al., 2013); negative regulation of JNK cascade signaling (Brace et al., 2014); Imd signaling: immune deficiency signaling cascade (Khush et al., 2002); mitotic cell cycle (Ducat et al., 2008). |

**Table B. Supplement to Figure 13: Signaling pathways linked to the Myc-CRM P29/P30**

| Gene name | Pathway | Activity-Function |
|---|---|---|
| Mpcp2 | **NOTC** | **Mitochondrial phosphate carrier protein 2**, Notch signaling involved in wing disc D/V pattern formation (Bejarano et al., 2008). |
| CadN | **WNBT** | **Cadherin-N**, roles of Armadillo, a *Drosophila* Catenin, during central nervous system development (Loureiro & Peifer, 1998). |
| clone 2.45 | **ERAD; APOP** | Gene: ***BCL2-associated athanogene 6***, Protein: **Large proline-rich protein BAG6**, ubiquitin-dependent ERAD pathway (GO Reference Genome Project, 2011-); negative regulation of apoptotic process (Gene Ontology Curators, 2002-); extracellular release via exosomes, is a ligand of the natural killer/NK cells receptor NCR3, stimulates NK cells cytotoxicity, may thereby trigger NK cells cytotoxicity against neighboring tumor cells & immature myeloid dendritic cells (DC) (UniProt Automatic Annotation); part of BAT3 complex. |
| Dap160 | **NOTC** | **Dynamin associated protein 160**; adaptor protein, contributes to endocytosis, regulates the Notch pathway, & mediates the asymmetric accumulation of a number or proteins, including aPKC during neuroblast division; negative regulation of Notch pathway (Tang et al., 2005). |
| Sep2 | **CECY** | **Septin 2**, regulation of cell cycle (O'Neill and Clark, 2013). |
| His2Av | **IMD** | **Histone H2A variant**, Imd signaling; negative regulation of peptidoglycan recognition protein signaling pathway (Tang et al., 2021). |
| Btk29A | **JNK; DPPBMP** | **Bruton tyrosine kinase 29A**, JNK signaling; TGF-beta signaling pathway (Harden, 2002). |
| LanB1 | **IMD; EGFR; JNK** | **Laminin B1**, Imd signaling; positive regulation of innate immune response (Cronin et al., 2009); apoptosis pathway (Sun et al., 2021); JNK, imaginal disc eversion (Pastor-Pareja et al., 2004). |
| Amph | **ARDAMM** | **Amphiphysin**, Amphiphysin-Rho1-Dia/DAAM-Rok pathway: an actomyosin clamp assembled by the Amphiphysin-Rho1-Dia/DAAM-Rok pathway reinforces somatic cell membrane folded around spermatid heads (Kapoor et al., 2021); (DAAM: Diaphanous & Dishevelled Associated Activator of Morphogenesis). |
| Ote | **DPPBMP** | **Otefin**, positive regulators of BMP (gbb/BMP) signaling pathway (Jiang et al., 2008). |

**Table B. Supplement to Figure 13: Signaling pathways linked to the Myc-CRM P29/P30**

| Gene name | Pathway | Activity-Function |
|---|---|---|
| Cortactin | **EGFR; PDGFR; VEGFR; PVR** | **Cortactin**, a cytoskeletal component, interacts with F-action, promotes actin polymerization via interaction with the Arp2/3 complex (O'Connell et al., 2019); Epidermal Growth Factor Receptor (EGFR) & the PDGF/VEGF-like (PVR) Receptor with involvement in cell migration and ring canal formation during *Drosophila* oogenesis (Quinones et al., 2010; Somogyi and Rorth, 2004); Cindr/dCortactin promote endocytosis (Quinones et al., 2010); invasive epithelial border cell migration is guided by *Drosophila* EGFR and PDGF/VEGF receptor (PVR) (Somogyi and Rorth, 2004). |
| H | **NOTC; EGFR; WNBT; APOP** | **Hairless**, major antagonist of Notch during imaginal disc development; silencer of Notch targets by assembling a transcriptional repressor complex including transcription factor Su(H) & general corepressors like Grouch (*gro*) & CtBP (*CtBP*); negative regulator of Notch signaling (Smylla et al., 2019; Wolf et al., 2019; Berndt et al., 2017; Maier et al., 2013; Troost and Klein, 2012; Kurth et al., 2011; Maier et al., 2011; Lee et al., 2009; Nagel et al., 2005; Barolo et al., 2002; Go et al., 1998; Lyman and Yedvobnick, 1995); EGFR signaling; (Protzer et al., 2008) cell death induction by downregulation of EGFR signaling; negative regulator of Wnt/TCF pathway (Bhambhani et al., 2011; Fang et al., 2006); positive regulator of Wnt/TCF pathway (Zhang and Arnosti, 2011). |
| CG1943 | **NOTC** | Wing disc dorsal/ventral pattern formation via Notch signaling (Bejarano et al., 2008); orthologous to human JPT1 (Jupiter microtubule associated homolog 1). |
| scny | **IMD; NFKB; RAS; NOTC** | **Scrawny**, ubiquitinyl hydrolase 1; RAS processing (Ubiquitin carboxyl-terminal hydrolase 36), negative regulator of Imd/ NF-kB, (Imd) signaling; (Buszczak et al., 2009; Thevenon et al., 2009); somatic stem cell population maintenance; inhibition of H2B ubiquitylation within stem cells for the repression of premature expression of key differentiation genes including Notch targets (Buszczak et al., 2009). |
| CG34417 | **TOLL; DPPBMP; HH** | Mesoderm development: Toll, Dpp, Hedgehog signaling pathways (Furlong et al., 2001). |
| smash | **WNBT; NOTC** | **Smallish, isoform H**, the *Drosophila* homologue of human LIM domain only 7 (LMO7); Role during eye development (Tanaka et al., 2019; Beati et al., 2018); contains the LIM, PDZ, & Calponin Homology (CH) domains. |

**Table B. Supplement to Figure 13: Signaling pathways linked to the Myc-CRM P29/P30**

| Gene name | Pathway | Activity-Function |
|---|---|---|
| Bin1 (SAP18) | **HH** | **Bin1**, Histone deacetylase complex subunit **SAP18**, regulation of Hedgehog signaling pathway by transcription factor Gli in mammals, repression of Gli-mediated transcription by Su(fu) for the recruitment of the SAP18-mSin3 complex to promoters containing the Gli-binding element (Yan Cheng & Bisho, 2002). |
| (psi) CG8912 | **MYC** | **P-element somatic inhibitor, isoform C**, far upstream binding-element protein 1/2, C-terminal (FUBP1/2) (Davis-Smyth et al., 1996; Ni et al., 2020); dual roles in RNA processing (Labourier et al., 2002; Siebel et al., 1994) and transcriptional regulation; FUBP1required for the activation of MYC transcription by binding to a single-stranded-far upstream sequence element of MYC promoter (Duan et al., 2017; Duncan et al., 1994). |
| tsu | **MAPK; RAS** | **Tsunagi**, rolled/MAPK pathway (Roignant & Treisman, 2010); RAS/MAPK signaling depends on the regulation of MAPK levels by the Exon Junction Complex (EJC) (Ashton-Beaucage & Therrien, 2010); The Mago-Tsunagi heterodimer interacts with the EJC key regulator Pym leading to EJC disassembly in the cytoplasm (Ghosh et al., 2014). |
| AGO2 | **miRNA** | **Argonaute 2**, interacts with small interfering RNAs (siRNAs) to form RNA-induced silencing complexes (RISCs), siRNA binding (Goh and Okamura, 2019, Kawamura et al., 2008; Tomari et al., 2007; Rand et al., 2005; Lingel et al., 2003); miRNA-mediated gene silencing (Besnard-Guérin et al., 2015). |
| sti | **HIPP; DPPBMP; NOTC; TOLL; WNBT; EGFR; CECY** | **Sticky**, negative regulation of Hippo signaling (Tran et al., 2019); TGFb/BMP, Notch, Wingless, and EGFR signaling pathways (Bivik et al., 2015); mitotic cytokinesis (Echard et al., 2004; Naim et al., 2004; Shandala et al., 2004; Rogers et al., 2003). |
| Ge-1 | **miRNA; NFKB; IMD; TOLL** | **Ge-1**, Enhancer of mRNA-decapping protein 4 homolog, RNAi pathway (miRNA-mediated gene silencing) (Eulalio et al., 2009); signaling pathways that activate NF-κB, Toll & Imd pathways (Jin et al., 2008; De Gregorio et al., 2002). |
| Helz | **miRNA** | **Helicase with zinc finger**, involved in ncRNA-mediated post-transcriptional gene silencing (GO Reference Genome Project, 2011-). |

**Table B. Supplement to Figure 13: Signaling pathways linked to the Myc-CRM P29/P30**

| Gene name | Pathway | Activity-Function |
|---|---|---|
| pod1 | **HIPP** | **pod1 coronin**, positive regulators of Hippo signaling pathway (Park et al., 2021). |
| lin-28 | **INR; JAKSTAT** | **Protein lin-28 homolog**, cold shock and RNA-binding protein - regulator of developmental timing - regulator of microRNA maturation; INR (insulin/IGF signaling (IIS) pathway) (Luhur et al., 2017; Chen et al., 2015); positive regulation of receptor signaling pathway via JAK/STAT (Sreejith et al., 2019). |
| gukh | **RAS** | **(GUK-holder)**, Wiskott-Aldrich syndrome protein family member, Ras-mediated signaling pathways (Mathew et al., 2002; Chen et al., 1998). |
| mask | **MAPK; JAKSTAT; PVR; SEVE; RTK** | **Multiple ankyrin repeats single KH domain**, mediator of RTK signaling either downstream of MAPK or signal transducer through a parallel branch of the RTK; EGFR pathways, Sevenless signaling (Smith et al., 2002); positive regulation of receptor signaling via JAK/STAT (Fisher et al., 2018); vascular endothelial growth factor receptor signaling pathway (PVR signaling) (Tsai et al., 2022). |
| Chchd3 | **HIPP; JNK** | **Coiled-coil-helix-coiled-coil-helix domain containing 3**, Loss of (*chchd3*) leads to inactivation of Hippo activity, decreased tissue growth, cell proliferation defects, oxidative stress & JNK pathway activation (Deng et al., 2016). |
| Lig | **JAKSTAT; HIPP** | **Lingerer**, negative regulation of receptor signaling pathway JAK/STAT (Baumgartner et al., 2013); positive regulation of Hippo signaling pathway (Dong et al., 2015). |
| (sw)<br>(CG18000)<br>(E8NH77) | **CECY** | **(short wing)**, non-catalytic intermediate chain subunit cytoplasmic dynein motor complex; contributes to wing, eye, oocyte development & polarity; mitotic cell division (Dzhindzhev et al., 2005); neuronal transport & neurogenesis; *Drosophila* Ser/Thr (MAST) kinase Drop interacts genetically with components of the dynein/dynactin complex (Hain et al., 2014). |
| twf | **TOLL** | **Twinfilin**, highly conserved ubiquitously expressed, actin monomer binding and inhibition of actin filament assembly (GO Reference Genome Project, 2011-); roles in bristle & neuronal development via Toll pathway (Cai et al., 2022). |
| Mi-2 | **NOTC; WNBT; ECDY** | **Mi-2**, Notch signaling (Zacharioudaki et al., 2019); Wingless, ecdysone signaling (Kon & Nusse; 2005). |

**Table B. Supplement to Figure 13: Signaling pathways linked to the Myc-CRM P29/P30**

| Gene name | Pathway | Activity-Function |
|---|---|---|
| Sara | **NOTC** | **Smad anchor for receptor activation**, Notch signaling (Montagne and Gonzalez-Gaitan, 2014). |
| CG9231 | **APOP** | Cellular response to hypoxia; positive regulation of apoptotic pathway; positive regulation of release of cytochrome c from mitochondria (Gene Ontology Curators, 2002-). |
| Bap111 | **WNBT**; **HH**; **DPPBMP**; **NOTC** | **Brahma associated protein 111kD**, extensive homology of the Brahma (BRM) complex to SWI/ SNF; Osa/Eyelid (*osa*) shows a strong genetic interaction with (*brm*), suggesting close cooperation with the BRM complex (Treisman et al. 1997; Vazquez et al. 1999); (Wnt/TCF); Hh; Dpp; and Notch signaling pathways: Eyelid antagonizes Wingless signaling during *Drosophila* development and has homology to the Bright family of DNA-binding proteins (Treisman et al., 1997; Kal et al., 2000). |
| udt | **JNK; JAKSTAT** | **Undicht**, wound healing, JNK signaling cascade, transduced by JUN/FOS transcriptional complexes (Ramet et al. 2002; Li et al. 2003; Ting et al. 2003, 2005a,b; Galko and Krasnow 2004; Mace et al. 2005; Campos et al., 2010); involvement of the JAK/STAT signaling cascade in this regenerative process (Mesilaty-Gross et al. 1999). |
| Hrb98DE | **PADPRI** | **Heterogeneous nuclear ribonucleoprotein at 98DE**, nuclear RNA-binding protein, controls hnRNA stability, splicing, IRES-dependent translation, & translational repression; represents one of the main targets of the poly(ADP-ribosyl)ation pathway; also regulates tissue polarity patterning & germline stem cell fate; negative regulation of RNA splicing (Ji and Tulin, 2009); post-translational modification of hnRNPs, such as poly(ADP-ribosyl)ation, is an important mechanism in the regulation of gene expression during development, such as eye pattern formation (Ji & Tulin, 2013). |
| glu | **CECY** | **Gluon**, a subunit of the multiprotein complex Condensin, mitotic cell cycle: prometaphase chromosome condensation (Bivik et al., 2015) & sister chromatid segregation, contributes to nervous system development & glucose metabolism; ubiquitous early, expressed in dividing cells throughout embryogenesis—pole cells, neuroblasts in the CNS & the PNS. |
| Chro | **CECY** | **Chromator**, mitotic spindle assembly check point signaling (Ding et al., 2009). |

**Table B. Supplement to Figure 13: Signaling pathways linked to the Myc-CRM P29/P30**

| Gene name | Pathway | Activity/Function |
|---|---|---|
| tral | **piRNA** | **Trailer hitch, isoform D**, piRNA pathway (Liu et al., 2011). |
| (sm) CG9218 | **ECDY** | **Small ubiquitin-related modifier (Smooth)**, sole family member protein in *Drosophila*, and closest to human heterogeneous nuclear ribonucleoprotein L (hnRNP L) (zur Lage et al. 1997); required for embryonic patterning & mitosis (syncytial blastoderm mitotic cell cycle), also has roles in wing patterning, Dpp (Miles et al., 2008) & Ras/MAPK signaling (Nie et al., 2009); localizes to the nucleus during interphase, & to kinetochores & midbodies during mitosis; positive regulation of Smoothened signaling pathway (Zhang et al., 2017); Toll-NF-κB signaling pathway (Kanoh et al., 2015). |
| rump | **CECY** | **Rumpelstiltskin**, mitotic cell cycle (Ducat et al., 2008). |
| Grip71 | **CECY** | **Grip71**, mitotic cell cycle (Ducat et al., 2008; Verollet et al., 2006). |
| enc | **CECY** | **Encore** regulation of germline mitosis (Hawkins et al., 1996). |
| Pop2 | **miRNA** | **Pop2**, poly(A)-specific ribonuclease; involved in translation inhibition (Ruscica et al., 2019; Braun et al., 2011); miRNA-mediated mRNA degradation (Braun et al., 2011). |
| CaBP1 | **APOP** | **Calcium-binding protein 1**, (protein disulfide-isomerase A6 homolog), apoptotic pathway (Okada et al., 2012); response to endoplasmic reticulum stress (GO Reference Genome Project, 2011-); protein disulfide isomerase activity (InterPro Project Members, 2004-). |
| Sirt4 | **INR** | **NAD-dependent protein deacylase Sirt4**, insulin signaling (Wood et al., 2018). |
| Ddx1 | **MYC** | **DEAD box 1**, *Ddx1* depletion associated with small size & aberrant gametogenesis, possibly through alternative splicing of Sirup RNA transcript, oogenesis (Germain et al., 2015); MYC-associated protein X (Max) binds with stronger affinity to rs72780850, a genetic regulatory variant of DDX1 in neuroblastoma sample (Jin Y, Shi J, Wang H, Lu J, Chen C, Yu Y, Wang Y, Yang Y, Ren D, Zeng Q, Ni X, Guo Y. 2021); DEAD box 1: a novel and independent prognostic marker for early recurrence in breast cancer (Germain et al., 2011). |

**Table B. Supplement to Figure 13: Signaling pathways linked to the Myc-CRM P29/P30**

| Gene name | Pathway | Activity/Function |
|---|---|---|
| mod(mdg4) | **APOP** | **(modifier of mdg4, isoform AD)**, apoptotic process (Harvey et al., 1997). |
| SPARC | **DPPBMP** | **SPARC/Osteonectin**, acidic cysteine-rich secreted protein, a small $Ca^{2+}$ & growth factor-binding secreted glycoprotein, enriched in basement membranes; during cell competition expressed in "loser" cells to avoid apoptosis mediated by Flower (*fwe*) & Ahuizotl (*azot*); hemocyte-secreted type IV collagen enhances BMP signaling to guide renal tubule morphogenesis in *Drosophila* (Bunt et al., 2010); cell adhesion (Hynes and Zhao, 2000). |
| wrd | **INR; TORC; HIPP** | **(well-rounded)**, one of the two regulatory B' subunits of the protein phosphatase PP2A; influences metabolism and growth via negative regulation of the INR/TORC (Insulin-like Receptor) signaling network (Van Hoof and Goris, 2003). |
| slf | **HEBI** | **Schlaff**, chitin binding protein involved in wing formation, substrate for Transglutaminase (Tg); a putative C-type lectin needed for the adhesion between the horizontal cuticle layers (Zuber&Moussian, 2019); cooperation with the heme-biosynthesis pathway to stabilize the distribution of the cuticle tyrosinated proteins, exemplified by Resilin, the tyrosinated proteins network needed for correct contact between chitin laminae within the procuticle & the epicuticle (Zuber & Moussian, 2019). |
| gbb | **DPPBMP** | **(glass bottom boat)**, a BMP ligand in the TGF-beta/BMP family of dimeric signaling molecules, binds to a receptor complex to transduce signal through phosphorylation of Mad; maintenance of stem cell populations, control of cell fate specification, proliferation, synapse growth, & neuropeptide release (Anderson and Wharton, 2017; Jensen et al., 2009; Shimmi et al., 2005). |
| Vps4 | **CECY; EGFR** | **Vacuolar protein sorting 4**, cell cycle; EGFR: *Drosophila* Vps4 promotes Epidermal Growth Factor Receptor signaling independent of its role in receptor degradation (Legent et al., 2015); JNK signaling: disruption of Vps4 and JNK function in *Drosophila* causes tumor growth (Rodahl et al., 2009). |
| sxc | **FGFR** | **Super sex combs**, a Polycomb group, encodes a O-GlcNAc transferase, involved in epigenetic gene silencing; positive regulation of fibroblast growth factor receptor signaling pathway FGFR (Mariappa et al., 2011). |

**Table B. Supplement to Figure 13: Signaling pathways linked to the Myc-CRM P29/P30**

| Gene name | Pathway | Activity-Function |
|---|---|---|
| pins | **RAS; CECY** | **partner of inscuteable**, Ras-association (RA) domains directly bind RanGTP & both the Canoe (RA) domains and RanGTP are required to recruit Mud to the cortex & activate the Pins/Mud/Dynein spindle orientation pathway (Wee et al., 2011); mitotic spindle orientation (Siegrist and Doe, 2005); asymmetric cell division (Januschke and Gonzalez, 2010; Yu et al., 2003; Parmentier et al., 2000). |
| Elp4C (G6907) | **CECY** | **Elongator complex protein 4**, establishment of mitotic spindle asymmetry (Planelles-Herrero et al., 2022); phosphorylase kinase regulator activity (Gene Ontology Curators, 2002-). |
| Orc6 | **CECY** | **Origin recognition complex subunit 6**, mitotic cell cycle (Balasov et al., 2009). |
| Tailor | **miRNA** | **Tailor**, terminal uridylyl-transferase Tailor, RNA-mediated gene silencing: positive regulation of miRNA catabolic process (Lin et al., 2017); pre-miRNA processing (Reimao-Pinto et al., 2015). |
| Cen | **CECY** | **Centrocortin**, asymmetrical Centrosomal localization during mitosis on spindles (Kao and Megraw, 2009); involved in regulation of embryonic cleavage furrow (Kao and Megraw, 2009). |
| apolpp | **WNBT; HH** | **Apolipophorin**, signal transduction activity, required for glycophosphatidylinositol-linked morphogens Wingless (Wg) & Hedgehog (Hh) function, by acting as vehicles for the movement of (Wg) & (Hh) (Pana´kova et al., 2005). |
| abs | **NOTC; CECY** | **Abstrakt**, DEAD-box protein, regulation of cell polarity in oocyte/embryos; downregulation of Notch signaling in asymmetric cell division in ganglion mother cell (GMC2-4a) together with Inscuteable (*Insc*) (Irion et al., 2004). |
| Manf | **EGFR; APOP; MAPK** | **Mesencephalic astrocyte-derived neurotrophic factor**, an invertebrate neurotrophic factor supporting dopaminergic neurons, cell-cell signaling between glial cells & neurons during development, EGFR signaling; apoptosis pathway; MAPK signaling (Palgi et al., 2009). |
| sun | **GPCR** | **Stunted,** activation of G-protein coupled receptor Methuselah (*mth*) in vitro, leading to increased intracellular Calcium ion levels; positive regulation of G protein-coupled receptor signaling pathway involved in regulating ageing; associated with longevity in *Drosophila* (Ja et al., 2009; Cvejic et al., 2004). |

**Table B. Supplement to Figure 13: Signaling pathways linked to the Myc-CRM P29/P30**

| Gene name | Pathway | Activity-Function |
|---|---|---|
| BAP155 moira (mor) | **EGFR; NFKB; CECY; NOTC; DPPBMP** | **Brahma associated protein 155 kDa**, imaginal disc-derived wing margin morphogenesis, EGFR (Terriente-Felix & de Celis, 2009); negative regulation of G1/S transition of mitotic cell cycle (Brumby et al., 2002); regulation of innate immune response via NF κB signaling (Bonnay et al., 2014). |
| Map60 (CP-60) | **CECY** | **Microtubule-associated protein 60**, localized to the centrosome in a cell cycle-dependent manner (Kellogg et al., 1995). |
| cutlet | **CECY** | **Chromosome transmission fidelity protein 18 homolog**, cell cycle: chromosome cohesion factor involved in sister chromatid cohesion, fidelity of chromosome transmission, and clamp loading activity; part of Ctf18 RFC-like complex (Gene Ontology Curators, 2002-). |
| 14-3-3zeta | **RAS; MAPK; HIPP; CECY** | **14-3-3 protein zeta**, Ras/Raf/MAPK signaling-dependent photoreceptor development (Kockel et al., 1997); regulation of Yorkie nuclear localization by Hippo (Hpo) signaling pathway (Zhang & Jiang, 2010). |
| aub | **miRNA; JAKSTAT** | **Aubergine**, piRNA binding (Huang et al., 2021; Webster et al., 2015; Nagao et al., 2010); global gene silencing by mRNA cleavage (Kennerdell et al., 2002); ncRNA-mediated post-transcriptional gene silencing (Tomari et al., 2004); RNA-mediated gene silencing (Bozzetti et al., 2015); JAK/STAT signaling pathway controls host defense in the gut by regulating stem cell proliferation & epithelial cell homeostasis (Cronin et al., 2009). |
| msn | **JNK; MAPK; NOTC; APOP; HIPP; TNFA** | **Misshapen, isoform A**, JNK activity (Martin and Wood, 2002); MAPK activity (InterPro Project Members, 2004-); negative regulation of Notch signaling pathway (Mishra et al., 2015); positive regulation of cell death; tumor necrosis factor α-mediated signaling pathway (Igaki et al., 2002; Moreno et al., 2002); positive regulation of Hippo signaling pathway (Li et al., 2015; Meng et al., 2015; Li et al., 2014). |
| mre11 | **CECY** | (**meiotic recombination 11**) double-strand break repair protein, G2/M DNA damage checkpoint signaling (Bi et al., 2006); telomere maintenance (Gao et al., 2009; Bi et al., 2005; Bi et al., 2004; Ciapponi et al., 2004). |
| shep | **MAPK; ERK; VTRA** | **Protein alan shepard**, MAPK/ERK pathway phosphorylated ERK (PERK) (Clement et al., 2013); vesicular trafficking: neuropeptide sorting through regulated secretory pathway (dense core vesicles) (Chen et al., 2014). |

**Table B. Supplement to Figure 13: Signaling pathways linked to the Myc-CRM P29/P30**

| Gene name | Pathway | Activity/Function |
|---|---|---|
| Ced-12 | **JNK; PVR; PDGFR; VEGFR** | **Ced-12 (engulfment and cell motility)**, PVR signaling: PDGF/VEGF (Platelet-Derived Growth Factor/Vascular Endothelial Growth Factor)-Receptor related (PVR), RTK activated by the binding of PDGF- & VEGF-related factors (Pvf1, Pvf2 or Pvf3), activates canonical Ras/Raf/MAPK (ERK) cascade, PI3K kinase pathway, TORC1 (FBrf0222697), RTK: Rho family small GTPases (Sopko & Perrimon, 2013; Ishimaru et al., 2004); JNK cascade (Sopko & Perrimon, 2013; Tran et al., 2013; Ishimaru et al., 2004). |
| crp (cropped) | **MYC** | **Activator protein 4**, **Cropped**, a downstream target of MYC, plays role in cell growth, organ size & survival; myosin binding (Liu et al., 2008); sequence-specific DNA binding (King-Jones et al., 1999); RNA pol II *cis*-regulatory region, sequence-specific DNA binding (GO Reference Genome Project, 2011-); protein dimerization activity (InterPro Project Members, 2004-); larval somatic muscle development (Dobi et al., 2014). |
| Septin5 (Sep5) | **CECY; APOP; DPPBMP; EGFR; NOTC** | **Septin5**, a member of the septin family of GTP-binding proteins; involved in cytokinesis; cell polarity & membrane rigidity; GTPase activity (GO Reference Genome Project, 2011-); imaginal disc-derived wing morphogenesis (Sun et al., 2021); involved in vein patterning & differentiation via modulation of TGF-β, Epidermal Growth Factor (EGF), & Notch ligands (Sun et al., 2021; De Celis, 2003). positive regulation of apoptosis pathway (Bae et al., 2007); regulation of cell cycle (O'Neill and Clark, 2013). |
| CG17202 | **MYC** | **c-Myc-binding protein homolog,** transcription coactivator activity; regulation of transcription, DNA-templated (GO Reference Genome Project, 2011-) stimulation of *c-Myc* transcription (Taira et al., 1998); associate of Myc 1; orthologous to human MYCBP (MYC binding protein). |
| Rpt4 | **NOTC; INR; CECY** | **Regulatory particle triple-A ATPase 4**, autophagy via hypoxia signaling (Lőw et al., 2013); Notch-mediated follicle cell differentiation and cell cycle switches, insulin-PI3K pathway (Jia et al., 2015). |
| ImpE1 | **ECDY** | **Ecdysone-inducible gene E1, isoform A**, This protein is similar to low-density lipoprotein LDL receptor; might play a role in the cell rearrangements associated with morphogenesis of the disc epithelium; imaginal disc eversion (Andres et al., 1993); (*ImpE1*) transcription is upregulated by 20-hydroxyecdysone. |

**Table B. Supplement to Figure 13: Signaling pathways linked to the Myc-CRM P29/P30**

| Gene name | Pathway | Activity/Function |
|---|---|---|
| mbt | **WNBT; MAPK** | **(mushroom body tiny)**, Serine/threonine-protein kinase PAK (P21-activated kinase), association with Wingless pathway by modulating DE-cadherin mediated cell adhesion via phosphorylation of Armadillo (Menzel et al., 2008); regulation of MAP kinase cascade; protein phosphorylation (GO Reference Genome Project, 2011-; Schneeberger and Raabe, 2003); mushroom body development (Melzig et al., 1998). |
| unc-45 | **JAKSTAT** | **(uncoordinated mutant number 45)**, JAK/STAT signaling pathway controls host defense in the gut by regulating stem cell proliferation and thus epithelial cell homeostasis; negative regulation of innate immune response; defense response to Gram-negative bacterium (Cronin et al., 2009); involved in JAK/STAT pathway activity in a gradient-dependent manner during patterning of the anterior-posterior axis of the follicular epithelium (Xi et al., 2003). |
| cib | **ERK; VEGFR; HIF** | **Ciboulot, isoform A**, (belongs to beta-thymosin family: thymosin beta-4, -10, -15) actin binding protein involved in brain development; remodeling of larval central nervous system (Boquet et al., 2000); anti-apoptotic response to an external stress (Sosne et al., 2004); HIF-1alpha stabilization via Erk activation (Hypoxia-inducible transcription factor (HIF)-1 alpha stabilization by actin-sequestering protein, thymosin beta-4 (TB4) in Hela cervical tumor cells); TB4P-increased HIF-1a stabilization & increase of the activity of hypoxia target gene, vascular endothelial growth factor (VEGF) transcription & elevated ERK phosphorylation (Oh et al., 2008). |
| Cp190 | **ECDY; WNBT** | **Centrosomal protein 190kD**, involved in the formation of most contact domain boundaries distal to a transcribed promoter; prevention of regulatory cross-talk between specific gene loci patterning the embryo; essential function during early development; response to ecdysone (Pascual-Garcia et al., 2017); Wingless signaling pathway: Cp190 found in complex with dTCF/Pan (pangolin), Su(Hw) and Pita (Marat Sabirov & Artem Bonchuk, 2021). |
| Poc1 | **CECY** | **Proteome of centrioles 1**, centrosome cycle (centrosome duplication and separation) (Blachon et al., 2009). |
| lmgB | **APOP** | **lemming B**, apoptosis pathway (Nagy et al., 2012). |

**Table B. Supplement to Figure 13: Signaling pathways linked to the Myc-CRM P29/P30**

| Gene name | Pathway | Activity/Function |
|---|---|---|
| Larp4B | **MYC; CECY** | **La-related protein Larp4B**, negative regulation of cell growth and translation (Funakoshi et al., 2018). |
| CG9705 | **HH; NOTC** | **Cold shock domain-containing protein CG9705**, dendrite morphogenesis: Hedgehog signaling pathway (Iyer et al., 2013); Notch signaling (Iyer et al., 2013; Mummery-Widmer et al., 2009). |
| Isha (cg4266) | **WNBT** | **Insulator su(Hw) mRNA adaptor binding protein**; Wingless signaling pathway: in complexing of Cp190 with dTCF/Pan (pangolin), Su(Hw) & Pita (Marat Sabirov & Artem Bonchuk, 2021). |
| Slik | **CECY; NOTC; EGFR** | **Sterile20-like kinase, isoform A**, mitotic spindle midzone (Carreno et al., 2008); protein serine/threonine kinase activity, coordinated regulation of epithelial morphology & proliferation (Hughes and Fehon, 2006); regulation of Notch and EGF Receptor signaling pathways (Hughes and Fehon, 2006; Maitra et al., 2006). |
| Arpc3A | **RTK** | **Actin-related protein 2/3 complex, subunit 3A**, structural constituent of cytoskeleton, contributes to actin filament binding activity, involved in Arp2/3 complex-mediated actin nucleation; Receptor Tyrosine Kinases (RTK)-N-WASP/SCAR-Rho-GTPase-Arpc2,3 (Benesh et al., 2002). |
| Vig | **miRNA** | **vasa intronic gene**, involved in RNA interference; ncRNA-mediated post-transcriptional gene silencing; Fragile X-related protein and VIG associate with the RNA interference machinery (Caudy et al., 2002). |
