## Supplemental Tables for "Targeting Regulatory Factors Associated with the *Drosophila Myc cis*-Elements by Reporter Expression, Gel Shift Assay, and Mass Spectrometric Protein Identification": Table C.pdf

**Table C. Supplement to Figure 13: Signaling pathways linked to the Myc-CRM P31/P32**

| Gene name | Pathway | Activity-Function |
|---|---|---|
| Bap55 | **NOTC; EGFR** | **Brahma associated protein 55kD**, Notch signaling (Pillidge & Bray, 2019); response to EGFR signaling in the *Drosophila* wing (Terriente-Félix & de Celis, 2009). |
| caz | **HIPP** | **Chromatin and RNA-binding protein Cabeza** (Mallik et al., 2018; InterPro Project Members, 2004-), ('cabeza' means 'head' in Spanish), a single ortholog of human FUS in *Drosophila*; genetic link exists between Caz & Hippo signaling pathway: Hippo (Hpo) the *Drosophila* ortholog of human mammalian sterile 20-like kinase (MST) 1 rescues Cabezas knockdown-induced eye phenotype- and neuron-specific defects (Azuma et al., 2018; Shimamura et al., 2014; Xia et al., 2012); transcription coregulatory activity (GO Reference Genome Project, 2011-); transcription Factor TFIID. |
| Hmg-2 (HMGB2) | **WNBT** | **High mobility group protein 2,** regulates chondrocyte hypertrophy by mediating Runt-related transcription factor 2 expression & Wnt signaling (Taniguchi et al., 2018); Wnt signaling & HMGB2 regulate articular cartilage surface maintenance (Taniguchi et al., 2009). |
| LKRSDH | **ECDY** | **Lysine ketoglutarate reductase/saccharopine dehydrogenase**, negative regulation of ecdysone receptor signaling pathway (Cakouros et al., 2008). |
| ebi | **EGFR; WNBT; NOTC** | **Ebi**, a nuclear-localized protein, EGFR signaling (Dong et al., 1999); Notch signaling pathway (Tsuda et al., 2002); Wingless signaling pathway (Li J and Wang CY, 2008; Marygold et al., 2011; DasGupta et al., 2005); contributes to multiple processes including wing margin development (Nguyen et al., 2016) & photoreceptor development (Tsuda et al., 2006, Tsuda et al., 2002); negative regulation of JNK cascade; regulation of transcription (Lim et al., 2012). |
| Bin1 (SAP18) | **HH** | **Histone deacetylase complex subunit SAP18**, regulation of Hh signaling by transcription factor Gli in mammals, repression of Gli-mediated transcription by Su(fu), recruitment of the SAP18-mSin3 to promoters containing the Gli-binding element (Yan Cheng and Bisho, 2002). |
| HmgD | **EGFR; ECDY; DPPBMP; WNBT** | **High mobility group protein D, isoform C & High mobility group protein C, isoform D**, EGFR signaling (Anan Ragab & Travers, 2006); Ecdysone/ecdysteroid signaling (Chen et al., 2008); DPP signaling: HMGD binding to the Dpp-responsive enhancer of (*tinman*) as well as to the Tinman protein during *Drosophila* cardiogenesis (Zaffran, 2002); Wnt/TCF signaling (Archbold et al., 2014). |

**Table C. Supplement to Figure 13: Signaling pathways linked to the Myc-CRM P31/P32**

| Gene name | Pathway | Activity-Function |
|---|---|---|
| gw | **miRNA** | **Gawky**, RNAi pathway (miRNA-mediated gene silencing pathway) (Chekulaeva et al., 2010; Chekulaeva et al., 2009; Eulalio et al., 2009; Eulalio et al., 2009; Zekri et al., 2009; Eulalio et al., 2008); miRNA induced silencing complex (Behm-Ansmant et al., 2006; Rehwinkel et al., 2005). |
| larp | **RAS; MAPK** | **La related protein, isoform F**, male meiotic nuclear division, Ras/MAPK signaling pathway (Blagden et al., 2009). |
| Sti | **HIPP; DPPBMP; NOTC; TOLL; WNBT; EGFR** | **Sticky**, negative regulation of Hippo signaling (Tran et al., 2019); TGFb/BMP, Notch, Wingless, and EGFR signaling pathways (Bivik et al., 2015); mitotic cytokinesis (Echard et al., 2004; Naim et al., 2004; Shandala et al., 2004; Rogers et al., 2003). |
| IntS11 | **NOTC**; **EGFR**; **MYC** | **Integrator complex subunit 11**, Notch signaling (Shersher et.al., 2021); Epidermal Growth Factor Receptor pathway (Tilley & Mollet, 2021). |
| (rin) (CG9412) | **RAS; RTK** | **Rasputin**, evolutionarily conserved RNA-binding protein functions as a link between Ras signaling and RNA metabolism (Costa et al., 2013; Pazman et al., 2000); positive regulator of (*orb*) during *Drosophila* oogenesis; Rasputin (*Rin*), the *Drosophila* homologue of Ras-GAP SH3 Binding Protein (G3BP) associates with Orb in a messenger ribonucleoprotein (mRNP) complex; Rho-mediated signaling (Pazman et al., 2000). |
| Ge-1 | **miRNA**; **NFKB; IMD; TOLL** | **Ge-1**, Enhancer of mRNA-decapping protein 4 homolog, RNAi pathway (miRNA-mediated gene silencing) (Eulalio et al., 2009); signaling pathways that activate NF-κB, Toll & Imd pathways (Jin et al., 2008; De Gregorio et al., 2002). |
| sqd | **DPPBMP** | **Squid, isoform E**, Dpp signaling, squid & Hephaestus (but not Hrb27C) are necessary for proper bone morphogenetic protein (BMP) signaling in GSCs; novel roles for RNA binding proteins Squid, Hephaestus,& Hrb27C during *Drosophila* oogenesis (Finger et al., 2022). |
| Pym | **MAPK; RAS** | **Partner of Y14 and Mago**, exon-exon junction complex; cap-dependent translational initiation; The Mago-Tsunagi heterodimer interacts with the EJC key regulator Pym leading to EJC disassembly in the cytoplasm (Ghosh et al., 2014). |

**Table C. Supplement to Figure 13: Signaling pathways linked to the Myc-CRM P31/P32**

| Gene name | Pathway | Activity-Function |
|---|---|---|
| Sep2 | **CECY** | **Septin 2**, regulation of cell cycle (O'Neill and Clark, 2013). |
| Rok, Drok | **WNPCP; CECY; RTK** | **Rho kinase**, *Drosophila* Rho kinase (Drok) (non-specific Ser/Thr protein kinase), activation by Rho1-dependent GTP, phosphorylation & modulation of cytoskeletal protein, particularly myosin II; dynamic regulation in subcellular locals influences cell polarization, movement, & cell shape (Dawes-Hoang et al., 2005) during interphase & mitosis; Fz/Dsh signaling (Fz/Dsh planar cell polarity pathway, cell adhesion & cell planar polarity (PCP) (Winter et al., 2001); mitotic cell cycle; mitotic spindle elongation (Hickson et al., 2006); mitotic cytokinesis (Tsankova et al., 2017; Dean and Spudich, 2006; Hickson et al., 2006; Dean et al., 2005); Rho protein signal transduction (GO Reference Genome Project, 2011-; Mizuno et al., 1999). |
| Dhc64C | **CECY** | **Dynein heavy chain 64C, isoform G**, acentrosomal (centrosome-independent) mitotic spindle assembly pathway; mitotic cell cycle (Maiato et al., 2004). |
| pod1 | **HIPP** | **pod1 coronin**, positive regulators of Hippo signaling pathway (Park et al., 2021). |
| Dap160 | **NOTC** | **Dynamin associated protein 160**; adaptor protein, contributes to endocytosis, regulates the Notch pathway, & mediates the asymmetric accumulation of a number or proteins, including aPKC during neuroblast division; negative regulation of Notch pathway (Tang et al., 2005). |
| Chchd2 | **INR** | **Coiled-coil-helix-coiled-coil-helix domain containing 2**, Insulin/IGF signaling is a key signaling pathway that maintains mitochondrial activity (Meng et al., 2017) |
| spdi | **WNBT** | **split discs**, integrin signaling pathway in cell-migration & adhesion (Saadi et al., 2011). |
| Cno | **HH**; **RAS** | **Canoe**, scaffold protein in adherens junctions, involved in morphogenesis in a variety of tissues; positive regulators of hedgehog signaling pathway (Marada et al., 2016); Ras-association (RA) domains directly bind RanGTP & both the Canoe (RA) domains and RanGTP are required to recruit Mud to the cortex & activate the Pins/Mud/dynein spindle orientation pathway (Wee et al., 2011); regulation of JNK cascade (Boettner et al., 2003). |

**Table C. Supplement to Figure 13: Signaling pathways linked to the Myc-CRM P31/P32**

| Gene name | Pathway | Activity-Function |
|---|---|---|
| Sara | **NOTC** | **Smad anchor for receptor activation**, an endosomal protein (GO Reference Genome Project, 2011-) involved in asymmetric stem cell division, intestinal stem cell homeostasis, and stem cell fate determination; Notch signaling (Montagne and Gonzalez-Gaitan, 2014). |
| Amph | **ARDAMM** | **Amphiphysin**, Amphiphysin-Rho1-Dia/DAAM-Rok pathway: an actomyosin clamp assembled by the Amphiphysin-Rho1-Dia/DAAM-Rok pathway reinforces somatic cell membrane folded around spermatid heads (Kapoor et al., 2021); (DAAM: Diaphanous and Dishevelled Associated Activator of Morphogenesis). |
| CG9231 | **APOP** | Cellular response to hypoxia; positive regulation of apoptosis; positive regulation of release of cytochrome c from mitochondria (Gene Ontology Curators, 2002-). |
| msk | **HH; DPPBMP; MAPK** | **Moleskin**, (Moleskin/Importin-7), an importin, contributes to protein import into the nucleus, is involved in eye development, muscle attachment & wing cell fate specification; Hedgehog, Dpp, MAPK (Vrailas et al., 2006). |
| Cka | **RAS; MAPK; JNK; HIPP** | **Connector of kinase to AP-1**, Hippo (Neal et al., 2022; Neal et al., 2020; Pojer et al., 2021; Gil-Ranedo et al., 2019; Zheng et al., 2017; Liu et al., 2016; Ribeiro et al., 2010), JNK (La Marca et al., 2019; Ashton-Beaucage et al., 2014; Chen et al., 2002), and Ras/MAPK signaling pathways (Ashton-Beaucage et al., 2014); Striatin-Interacting Phosphatase & Kinase Complex (Part of FAR / SIN / STRIPAK complex). |
| Cdk1 | **HH; NOTC; HIPP; JAKSTAT; EGFR; WNBT; JNK; INR; CECY** | **Cyclin-dependent kinase 1**, G1/S transition of mitotic cell cycle (Lehner and O'Farrell, 1990); G2/S transition of mitotic cell cycle (Stern et al., 1993; Lehner and O'Farrell, 1990); follicle cell of egg chamber development, Notch, Hedgehog, EGFR, Wingless, JAK/STAT, Hippo, and JNK pathways; insulin-PI3K pathways (Jia et al., 2015). |
| Gp150 | **NOTC** | **Glycoprotein 150, isoform E,** Transmembrane glycoprotein regulates Notch signaling, involved in compound eye development (Fetchko et al., 2002); transmembrane receptor protein tyrosine phosphatase signaling pathway (Tian and Zinn, 1994). |
| Mpcp2 | **NOTC** | **Phosphate carrier protein, mitochondrial**, Notch signaling wing disc D/V pattern formation (Bejarano et al., 2008). |

**Table C. Supplement to Figure 13: Signaling pathways linked to the Myc-CRM P31/P32**

| Gene name | Pathway | Activity-Function |
|---|---|---|
| udt | **JNK; JAKSTAT** | **Undicht**, wound healing, JNK signaling cascade, transduced by JUN/FOS transcriptional complexes (Ramet et al. 2002; Li et al. 2003; Ting et al. 2003, 2005a,b; Galko and Krasnow 2004; Mace et al. 2005; Campos et al., 2010); involvement of the JAK/STAT signaling cascade in this regenerative process (Mesilaty-Gross et al. 1999). |
| Bap111 | **WNBT**; **HH**; **DPPBMP**; **NOTC** | **Brahma associated protein 111kD**, extensive homology of the Brahma (BRM) complex to SWI/ SNF; Osa/Eyelid (*osa*) shows a strong genetic interaction with (*brm*), suggesting close cooperation with the BRM complex (Treisman et al. 1997; Vazquez et al. 1999); Wnt/TCF, Hh, Dpp, and Notch signaling: Eyelid antagonizes Wingless signaling during *Drosophila* development and has homology to the Bright family of DNA-binding proteins (Treisman et al., 1997; Kal et al., 2000). |
| kis | **HH**; **NOTC; RAS** | **Kismet, isoform F**, a conserved function of the chromatin ATPase Kismet is in the regulation of Hedgehog expression (Terriente-Félix et al., 2011); Notch signaling pathway (Go and Artavanis-Tsakonas, 1998; Verheyen et al., 1996); ( *kis*) mutations have been identified in genetic screens as modifiers of the Ras and Notch signal transduction pathways (Go and Artavanis-Tsakonas, 1998; Therrien.et al., 2000; Melicharek et al., 2010). |
| ebi | **JNK; EGFR**; **WNBT; NOTC** | **F-box-like/WD repeat-containing protein ebi-like**, evolutionarily conserved repressor/silencer; JNK signaling: Ebi/ AP-1 complex (activator protein 1) represses pro-/anti-apoptotic genes; suppresses basal transcription levels of apoptotic genes protecting sensory neurons degeneration (Lim et al., 2012); regulates EGFR (Dong et al., 1999), Notch (Nguyen et al., 2016; Marygold et al., 2011; Tsuda et al., 2002), and Wg signaling; contributes to multiple processes including wing growth, eye development, regulation of transcription & innate immune response. |
| CG1943 | **NOTC** | Wing disc dorsal/ventral pattern formation via Notch signaling (Bejarano et al., 2008); orthologous to human JPT1 (Jupiter microtubule associated homolog 1). |
| sle | **CECY** | **Slender lobes**, part of nucleolus, its depletion alters the aggregation of nucleolar components & results in retardation of proliferation of Kenyon cells; nucleolus organization; (*sle*) gene identified by retarded mushroom body development, is required for proper nucleolar organization in *Drosophila* (Orihara-Ono et al., 2005). |

**Table C. Supplement to Figure 13: Signaling pathways linked to the Myc-CRM P31/P32**

| Gene name | Pathway | Activity-Function |
|---|---|---|
| Grip71 | **CECY** | **Grip71**, mitotic cell cycle (Ducat et al., 2008; Verollet et al., 2006). |
| enc | **CECY** | **Encore**, involved in the regulation of germline mitoses (Hawkins et al., 1996). |
| tral | **piRNA** | **Trailer hitch, isoform D**, piRNA pathway (Liu et al., 2011). |
| Pop2 | **miRNA** | **Pop2**, poly(A)-specific ribonuclease; involved in translation inhibition (Ruscica et al., 2019; Braun et al., 2011); miRNA-mediated mRNA degradation (Braun et al., 2011). |
| CaBP1 | **APOP** | **Calcium-binding protein 1**, (protein disulfide-isomerase A6 homolog), apoptotic pathway (Okada et al., 2012); response to endoplasmic reticulum stress (GO Reference Genome Project, 2011-); protein disulfide isomerase activity (InterPro Project Members, 2004-). |
| CG12384 | **APOP** | Apoptosis signaling pathway (GO Reference Genome Project, 2011-). |
| slf | **HEBI** | **Schlaff**, chitin binding protein involved in wing formation, substrate for Transglutaminase (Tg); a putative C-type lectin needed for the adhesion between the horizontal cuticle layers (Zuber&Moussian, 2019); cooperation with the heme-biosynthesis pathway to stabilize the distribution of the cuticle tyrosinated proteins, exemplified by Resilin, the tyrosinated proteins network needed for correct contact between chitin laminae within the procuticle & the epicuticle (Zuber & Moussian, 2019). |
| Usp7 | **HIPP; HH** | **Ubiquitin-specific protease 7**, negative regulation of Hippo signaling pathway (Sun et al., 2019); positive regulation of Smoothened signaling pathway (Zhou et al., 2015). |
| CG34417 | **TOLL; DPPBMP; HH** | Mesoderm development: Toll, Dpp, Hedgehog signaling pathways (Furlong et al., 2001). |
| Vps4 | **CECY; EGFR** | **Vacuolar protein sorting 4**, cell cycle; EGFR pathway: *Drosophila* Vps4 promotes Epidermal Growth Factor Receptor signaling independent of its role in receptor degradation (Legent et al., 2015); JNK signaling: Disruption of Vps4 and JNK function in *Drosophila* causes tumor growth (Rodahl et al., 2009). |
| Sirt4 | **INR** | **NAD-dependent protein deacylase Sirt4**, insulin signaling pathway (Wood et al., 2018). |

**Table C. Supplement to Figure 13: Signaling pathways linked to the Myc-CRM P31/P32**

| Gene name | Pathway | Activity-Function |
|---|---|---|
| gukh | **RAS** | **GUK-holder**, Wiskott-Aldrich syndrome protein family member, Ras-mediated signaling pathways (Mathew et al., 2002; Chen et al., 1998). |
| eIF4A | **DPPBMP; CECY** | **Eukaryotic translation initiation factor 4A**, an essential DEAD box RNA helicase protein & a canonical translation initiation factor; a component of the eIF4F cap-binding complex, essential for cap-dependent translation of mRNA; SMAD binding; negative regulation of BMP signaling pathway (Li and Li, 2006); mitotic cell cycle (Ducat et al., 2008). |
| CDK2AP1 | **CECY; WNBT; RAS; NOTC; JNK** | **CDK2-associated protein 1**, orthologous to several human genes including CDK2AP1 (cyclin dependent kinase 2 associated protein 1), involved in developmental processes (Reddy et al., 2010). |
| Elp4C (G6907) | **CECY** | **Elongator complex protein 4**, establishment of mitotic spindle asymmetry (Planelles-Herrero et al., 2022); phosphorylase kinase regulator activity (Gene Ontology Curators, 2002-). |
| mod(mdg4) | **APOP** | **modifier of mdg4, isoform AD**, apoptotic process (Harvey et al., 1997). |
| Tailor | **miRNA** | **Tailor**, terminal uridylyl-transferase Tailor, RNA-mediated gene silencing: positive regulation of miRNA catabolic process (Lin et al., 2017); pre-miRNA processing (Reimao-Pinto et al., 2015). |
| abs | **NOTC; CECY** | **Abstrakt**, DEAD-box protein, regulates cell polarity in oocytes & embryos; downregulation of Notch signaling in asymmetric cell division in ganglion mother cell (GMC2-4a) in collaboration with Inscuteable (*Insc*) (Irion et al., 2004). |
| (A4V4A5) Ran | **RAS; CECY; WNBT** | **GTP-binding nuclear protein Ran**, a member of the Ras superfamily (Koizumi et al., 2001); localizes around the microtubule spindle in vivo during mitosis in *Drosophila* embryos (Trieselmann & Wilde, 2002); nuclear import of β-Catenin into nucleus (Yokoya et al., 1999). |
| Mo25 | **WNK** | **Mo25, isoform B**, involved in neuroblast asymmetric cell division (Borkowsky et al., 2023); interactions between chloride & Mo25 regulate WNK [with no lysine (K)] kinases signaling in a transporting renal epithelium; WNK-SPAK/OSR1 (Rodan, 2018). |

**Table C. Supplement to Figure 13: Signaling pathways linked to the Myc-CRM P31/P32**

| Gene name | Pathway | Activity-Function |
|---|---|---|
| sun | **GPCR** | **Stunted,** activation of G-protein coupled receptor Methuselah (*mth*) in vitro, leading to increased intracellular Calcium ion levels; positive regulation of G protein-coupled receptor signaling pathway involved in regulating ageing; associated with longevity in *Drosophila* (Ja et al., 2009; Cvejic et al., 2004). |
| Ufd1-like | **ERAD; HH** | **Ubiquitin fusion degradation protein 1**, ubiquitin-dependent ERAD pathway; VCP-NPL4-UFD1 AAA ATPase complex, polyubiquitin modification-dependent protein binding (GO Reference Genome Project, 2011-); negative regulation of Smoothened signaling pathway (Zhang et al., 2013). |
| Gmd | **NOTC** | **GDP-mannose 4,6 dehydratase**, positive regulation of Notch signaling pathway (Perdigoto et al., 2011); cell fate commitment with the involvement of Notch signaling (Sasamura et al., 2007); involved in the early steps of *Drosophila* oogenesis via Notch signaling (Jagut et al., 2013). |
| PpD3 | **JNK**; **MAPK; CECY** | **Protein phosphatase D3**, conserved protein, involved in JNK and MAPK pathways (Miskei et al., 2011; Morrison et al., 2000); required for mitosis (G2/M progression). |
| clone 2.45 | **ERAD; APOP** | Gene: **BCL2-associated athanogene 6**, Protein: **Large proline-rich protein BAG6,** ubiquitin-dependent ERAD pathway (GO Reference Genome Project, 2011-); negative regulation of apoptotic process (Gene Ontology Curators, 2002-); extracellular release via exosomes, is a ligand of the natural killer/NK cells receptor NCR3, stimulates NK cells cytotoxicity, may thereby trigger NK cells cytotoxicity against neighboring tumor cells & immature myeloid dendritic cells (DC) (UniProt Automatic Annotation); part of BAT3 complex. |
| Spt20 | **JNK**; **NOTC** | **SPT20, isoform C,** Spt-Ada-Gcn5-acetyltransferase (SAGA) complex; JNK and Notch signaling pathway (Weake et al., 2009). |
| apolpp | **WNBT; HH** | **Apolipophorin**, signal transduction activity, required for glycophosphatidylinositol-linked morphogens Wingless (Wg) & Hedgehog (HH) function by acting as vehicles for the movement of (Wg) & (HH) (Pana´kova et al., 2005). |

**Table C. Supplement to Figure 13: Signaling pathways linked to the Myc-CRM P31/P32**

| Gene name | Pathway | Activity-Function |
|---|---|---|
| CG8003 | **NOTC; WNBT** | **Notch-regulated ankyrin repeat protein (NRARP)**, intracellular component of Notch signaling (Lamar et al., 3001), involved in crosstalk between Notch & Wnt signaling during development (Imaoka et al., 2014); involved in the differentiation of neural crest cell via regulation of LEF1 protein stability (Ishitani et al., 2015); controls the coordination between endothelial Notch & Wnt signaling in vessel density during angiogenesis (Phng et al., 2009). |
| Nup50 | **DPPBMP; CECY** | **Nuclear pore complex protein Nup50 (Nucleoporin 50kD)**, an important protein for TGF-beta signal transduction by mediating the nuclear translocation of Mothers against dpp protein (Mad); Association of 400 genes interacting with Nup50 Nucleoporin, among which transcriptionally active genes inside the nucleoplasm are predominantly involved in development & cell cycle regulation (Kalverda et al., 2010). |
| Hcf (dHCF, dHcf1, Hcf1) | **CECY; WNBT** | **Host cell factor**, phosphoprotein, involved in control of cell cycle (Guelman et al., 2006) & several processes including chromatin remodeling, histone acetylation, & positive regulation of transcription; implicated in both activation & repression of transcription, associates with Wingless enhanceosome as a regulator of canonical Wingless signaling pathway during wing imaginal disc vein development (Rodriguez-Jato et al., 2011). |
| piwi | **piRNA; CECY** | **Piwi: P-element induced wimpy testis, isoform B**, piRNA-mediated metabolic process, repression of transposable elements during meiosis by complexes of piRNAs/Piwi containing proteins, methylation & subsequent repression of transposons; piRNA binding (Sienski et al., 2015); chromatin silencing (Brower-Toland et al., 2007; Grimaud et al., 2006; Pal-Bhadra et al., 2004; Pal-Bhadra et al., 2002); female germline stem cell asymmetric division; germline stem cell population maintenance; male germline stem cell asymmetric division (*Cox et al., 1998*); gene silencing by RNA (Le Thomas et al., 2013); heterochromatin organization involved in chromatin silencing (Sienski et al., 2015; Sienski et al., 2012). |
| Nurf-38 | **ECDY** | **Nucleosome remodeling factor - 38kD**, ecdysone receptor-mediated signaling pathway (Badenhorst et al., 2005). |

**Table C. Supplement to Figure 13: Signaling pathways linked to the Myc-CRM P31/P32**

| Gene name | Pathway | Activity-Function |
|---|---|---|
| mre11 | **CECY** | (**meiotic recombination 11**) double-strand break repair protein, G2/M DNA damage checkpoint signaling (Bi et al., 2006); telomere maintenance (Gao et al., 2009; Bi et al., 2005; Bi et al., 2004; Ciapponi et al., 2004). |
| CG9705 | **HH**; **NOTC** | **Cold shock domain-containing protein CG9705**, dendrite morphogenesis: Hedgehog signaling pathway (Iyer et al., 2013); Notch signaling (Iyer et al., 2013; Mummery-Widmer et al., 2009). |
| Isha (cg4266) | **WNBT** | **Insulator su(Hw) mRNA adaptor**; Wingless signaling pathway: complexing of Cp190 with dTCF/Pan (pangolin), Su(Hw) & Pita (Marat Sabirov & Artem Bonchuk, 2021). |
| fzy | **CECY** | **Fizzy**, WD40 domain protein required for the full ubiquitin ligase activity of the anaphase-promoting complex / Cyclosome (APC/C) in mitosis and meiosis; functions to target substrates for destruction and drive metaphase and anaphase transition; anaphase-promoting complex-dependent catabolic process (Swan and Schupbach, 2007); mitotic cell cycle (Ducat et al., 2008). |
| Cp190 | **ECDY**; **WNBT** | **Centrosomal protein 190kD**, involved in the formation of most contact domain boundaries distal to a transcribed promoter; prevention of regulatory cross-talk between specific gene loci patterning the embryo; essential function during early development; response to ecdysone (Pascual-Garcia et al., 2017); Wingless signaling pathway: Cp190 found in complex with dTCF/Pan (pangolin), Su(Hw) and Pita (Marat Sabirov & Artem Bonchuk, 2021). |
| tacc | **CECY** | **Transforming acidic coiled-coil protein**, **mitotic** cell cycle (Gergely et al., 2000). |
| Vig | **miRNA** | **(vasa intronic gene)**, involved in RNA interference; ncRNA-mediated post-transcriptional gene silencing; Fragile X-related protein and VIG associate with the RNA interference machinery (Caudy et al., 2002). |
| Rpt4 | **NOTC; INR; CECY** | **Regulatory particle triple-A ATPase 4**, autophagy via hypoxia signaling (Lőw et al., 2013); Notch-mediated follicle cell differentiation and cell cycle switches, insulin-PI3K pathway (Jia et al., 2015). |
| CG6607 | **VTRA** | **Coiled-coil domain-containing protein 128**, Orthologous to human PPP1R21 (protein phosphatase 1 regulatory subunit 21); Rab protein signal transduction, early endosome, small GTPase binding (Gillingham et al., 2014). |

**Table C. Supplement to Figure 13: Signaling pathways linked to the Myc-CRM P31/P32**

| Gene name | Pathway | Activity-Function |
|---|---|---|
| Mad | **DPPBMP**; **WNBT; EGFR** | **Mothers against decapentaplegic**, BMP signaling pathway core components (Vuilleumier et al., 2022; Guo et al., 2013; Weiss et al., 2010; Kamiya et al., 2008; Yao et al., 2006; Muller et al., 2003; Dai et al., 2000; Das et al., 1998; Inoue et al., 1998); involved in wing development via modulation of EGFR and BMP signaling pathways (Dworkin and Gibson, 2006; Lecuit et al., 1996); negative regulation of salivary gland boundary formation with the involvement of Wnt/Wg signaling pathway (Bradley, P.L., Haberman, A.S., Andrew, D.J. 2001). |
