## Supplemental Tables for "Targeting Regulatory Factors Associated with the *Drosophila Myc cis*-Elements by Reporter Expression, Gel Shift Assay, and Mass Spectrometric Protein Identification": Table D.pdf

**Table D. Supplement to Figure 13: Signaling pathways linked to the Myc-CRM P35/P36**

| Gene name | Pathway | Activity-Function |
|---|---|---|
| kst | **EGFR; JNK; VTRA** | **Karst, isoform F**, EGFR signaling endosome transport via multivesicular body sorting pathway (Tjota et al., 2011); JNK signaling cascade in wound healing & epithelial sheath (Campos et al., 2010). |
| Lig | **JAKSTAT; HIPP** | **Lingerer**, negative regulation of receptor signaling pathway JAK/STAT (Baumgartner et al., 2013); positive regulation of Hippo signaling pathway (Dong et al., 2015). |
| (sw) (CG18000) (E8NH77) | **CECY** | **(short wing)**, non-catalytic intermediate chain subunit cytoplasmic dynein motor complex; contributes to wing, eye, oocyte development & polarity; mitotic cell division (Dzhindzhev et al., 2005); neuronal transport & neurogenesis; *Drosophila* Ser/Thr (MAST) kinase Drop interacts genetically with the components of the dynein/dynactin complex (Hain et al., 2014). |
| Amun | **NOTC** | **Amun, isoform A**, contains a putative DNA glycosylase domain; chaeta, compound eye, & wing disc development (Shalaby et al., 2009). |
| cype | **DPPBMP; HH** | **Cyclope**, cytochrome c oxidase subunit VIc homolog acting as an enhancer of Dpp pathway phenotypes: involved in hair & cell growth, and in ommatidia development; involved in Hh signaling (Chang et al., 2001); significant alteration in the expression of *cype* observed during the progression of prostate cancer (Herrmann et al., 2003). |
| MED17 | **JASAU; RAS; MAPK** | **Mediator of RNA pol II transcription subunit 17**, sex comb development; required for adult cell & segment identity specification; SP1 transcription activation; thyroid hormone receptor complex component; VitD receptor interactor; jasmonate (JA) & Auxin signaling component (Boube et al., 2000); Ras/MAK pathway (Singh and Han 1995). |
| Mpcp2 | **NOTC** | **Phosphate carrier protein, mitochondrial**, Notch signaling involved in wing disc D/V pattern formation (Bejarano et al., 2008). |
| Nurf-38 | **ECDY** | **Nucleosome remodeling factor - 38kD**, ecdysone receptor-mediated signaling pathway (Badenhorst et al., 2005). |
| MED25 | **IMD; NFKB** | **Mediator of RNA poly II transcription subunit 25**, Imd pathway, NF-κB signaling; positive regulation of antibacterial peptide biosynthetic process (Valanne et al., 2010). |

**Table D. Supplement to Figure 13: Signaling pathways linked to the Myc-CRM P35/P36**

| Gene name | Pathway | Activity-Function |
|---|---|---|
| LKRSDH | **ECDY** | **Lysine ketoglutarate reductase/saccharopine dehydrogenase**, negative regulation of ecdysone receptor signaling pathway (Cakouros et al., 2008). |
| PpD3 | **JNK**; **MAPK** | **Protein phosphatase D3**, conserved protein, involved in JNK and MAPK pathways (Miskei et al., 2011; Morrison et al., 2000); required for mitosis (G2/M progression). |
| Idgf2 | **APOP**; **INR**; **IMD**; **JAKSTAT** | **Imaginal disc growth factor 2**, negative regulation of apoptotic process (Broz et al., 2017); insulin receptor pathway (Varela et al., 2002); Imd and JAK/STAT signaling pathways collaborate with the Imaginal Disc Growth Factors 2 and 3 in the *Drosophila* response to nematode infection (Shruti, 2018). |
| Bin1 (SAP18) | **HH** | **Histone deacetylase complex subunit SAP18**, regulation of Hedgehog (Hh) signaling pathway by transcription factor Gli in mammals, repression of Gli-mediated transcription by Su(fu) in cooperation with SAP18 for the recruitment of the SAP18- mSin3 complex to promoters containing the Gli-binding element (Yan Cheng and Bisho, 2002). |
| Cdk5 | **APOP**; **CECY; WNBT** | **Cyclin-dependent kinase 5**, intrinsic apoptotic signaling (Kang et al., 2012); regulation of transcription involved in G1/S transition of mitotic cell cycle (GO Reference Genome Project, 2011-); proapoptotic signaling by CDK5 & MEKK1 (Ryoo HD., 2018); Wingless signaling: human wild-type Tau interacts with Wingless pathway components & produces neurofibrillary pathology in *Drosophila* (Tau-induced neurodegeneration & the Wnt pathway using GSK-3β and 2 additional components, β-Catenin/Armadillo & dTCF) (Jackon & Geschwind ,2002). |
| ebi | **JNK; EGFR**; **WNBT; NOTC** | **F-box-like/WD repeat-containing protein ebi-like**, evolutionarily conserved repressor/silencer; JNK signaling: Ebi/ AP-1 complex (activator protein 1) represses pro-/anti-apoptotic genes; suppresses basal transcription levels of apoptotic genes protecting sensory neurons degeneration (Lim et al., 2012); regulation of EGFR (Dong et al., 1999), Notch (Nguyen et al., 2016; Marygold et al., 2011; Tsuda et al., 2002), and Wg signaling; contributes to multiple processes including wing growth, eye development, regulation of transcription, & innate immune response. |
| Larp4B | **MYC** | **La-related protein Larp4B**, negative regulation of cell growth and translation (Funakoshi et al., 2018). |

**Table D. Supplement to Figure 13: Signaling pathways linked to the Myc-CRM P35/P36**

| Gene name | Pathway | Activity-Function |
|---|---|---|
| Hrb98DE | **PADPRI** | **Heterogeneous nuclear ribonucleoprotein at 98DE**, nuclear RNA-binding protein, controls hnRNA stability, splicing, IRES-dependent translation, & translational repression; represents one of the main targets of the poly(ADP-ribosyl)ation pathway; also regulates tissue polarity patterning & germ-line stem cell fate; negative regulation of RNA splicing (Ji and Tulin, 2009); post-translational modification of hnRNPs such as poly(ADP-ribosyl)ation, is an important mechanism in the regulation of gene expression during development in processes such as eye pattern formation (Ji & Tulin, 2013). |
| pum | **EGFR** | **Pumilio, isoform G**, negative regulation of Epidermal Growth Factor Receptor (EGFR) signaling pathway (Kim et al., 2012). |
| (sm) CG9218 | **ECDY** | **Small ubiquitin-related modifier (Smooth)**, sole family member protein in *Drosophila*, and closest to human heterogeneous nuclear ribonucleoprotein L (hnRNP L) (zur Lage et al. 1997); required for embryonic patterning & mitosis (syncytial blastoderm mitotic cell cycle), and roles in wing patterning, Dpp (Miles et al., 2008) & Ras/MAPK signaling (Nie et al., 2009); localizes to the nucleus during interphase, & to kinetochores & midbodies during mitosis; positive regulation of Smoothened signaling pathway (Zhang et al., 2017); Toll-NF-κB signaling pathway (Kanoh et al., 2015). |
| aub | **miRNA**; **JAKSTAT** | **Aubergine**, piRNA binding (Huang et al., 2021; Webster et al., 2015; Nagao et al., 2010); global gene silencing by mRNA cleavage (Kennerdell et al., 2002); ncRNA-mediated post-transcriptional gene silencing (Tomari et al., 2004); RNA-mediated gene silencing (Bozzetti et al., 2015); JAK/STAT signaling pathway controls host defense in the gut by regulating stem cell proliferation & epithelial cell homeostasis (Cronin et al., 2009). |
| obe | **NOTC**; **WNBT** | **Obelus**, a Ski2-family helicase regulates alternative mRNA splicing of crumb (*crb*); required for cell polarity & adherens junction organization; Notch signaling (*Vichas et al., 2015*); contains EGF repeats with profound effects on cell adhesion (Balzar et al., 2001) and ligand–receptor interactions (Rebay et al., 1991), with involvement in protein localization & proteins glycosylation (*Acar et al., 2008*); involved in Wingless signaling, altering subcellular distribution and dynamics of Armadillo and E-cadherin in third instar larval wing imaginal discs (Somorjai & Martinez-Arias, 2008). |

**Table D. Supplement to Figure 13: Signaling pathways linked to the Myc-CRM P35/P36**

| Gene name | Pathway | Activity-Function |
|---|---|---|
| AGO2 | **miRNA** | **Argonaute 2**, interacts with small interfering RNAs (siRNAs) to form RNA-induced silencing complexes (RISCs), siRNA binding (Goh and Okamura, 2019, Kawamura et al., 2008; Tomari et al., 2007; Rand et al., 2005; Lingel et al., 2003); miRNA-mediated gene silencing (Besnard-Guérin et al., 2015). |
| enc | **CECY** | **Encore** regulation of germline mitoses (Hawkins et al., 1996). |
| Mi-2 | **NOTC; WNBT; ECDY** | **Mi-2**, Notch signaling (Zacharioudaki et al., 2019); wingless and ecdysone pathways (Kon & Nusse; 2005). |
| tral | **piRNA** | **Trailer hitch, isoform D**, piRNA pathway (Liu et al., 2011). |
| Arc1 | **INR** | **Activity-regulated cytoskeleton associated protein 1**, behavioral response to starvation, Insulin receptor (Inr) signaling (Montana and Littleton, 2006; Kremerskothen et al., 2002); adipokinetic hormone (AKH) signaling (Hughson & O'Connor, 2021; Mattaliano et al., 2007). |
| CG9705 | **HH; NOTC** | **Cold shock domain-containing protein CG9705**, dendrite morphogenesis: Hedgehog signaling pathway (Iyer et al., 2013); Notch signaling (Iyer et al., 2013; Mummery-Widmer et al., 2009). |
| msi | **HIF; NOTC** | **Musashi**, negatively regulates the Hypoxia Inducible Factor (HIF) pathway, contributes to cell fate determination, as well as cellular response to normoxic/hypoxic conditions; negative regulation of translation; asymmetric cell fate determination by Notch signaling (Bardin et al., 2004). |
| dgt4 | **CECY** | **dim γ-tubulin 4 (Augmin complex subunit dgt4)**, mitotic cell cycle, plays role in centrosome-independent generation of spindle microtubules (Goshima et al., 2008); mitotic spindle assembly (Goshima et.al., 2007). |
| Sirt4 | **INR** | **NAD-dependent protein deacylase Sirt4**, Insulin receptor (Inr) signaling pathway (Wood et al., 2018). |
| CaBP1 | **APOP** | **Calcium-binding protein 1**, (protein disulfide-isomerase A6 homolog), apoptotic pathway (Okada et al., 2012); response to endoplasmic reticulum stress (GO Reference Genome Project, 2011-); protein disulfide isomerase activity (InterPro Project Members, 2004-). |

**Table D. Supplement to Figure 13: Signaling pathways linked to the Myc-CRM P35/P36**

| Gene name | Pathway | Activity-Function |
|---|---|---|
| smg | **CECY** | **Smaug**, translation repressor activity (Dahanukar et al., 1999); mRNA regulatory element binding translation repressor activity (Dean et al., 2002); cell cycle (Tadros et al., 2007). |
| bel | **ECDY; miRNA** | **Belle**, ecdysone-mediated induction of salivary gland cell autophagy cell death (Ihry and Bashirullah, 2014); ncRNA-mediated post-transcriptional gene silencing (Ulvila et al., 2006); mitotic cell cycle (Pek and Kai, 2011). |
| CG30291 | **NFKB**; **ERK** | NF-κB pathway; ERK1/2 pathways (Utreras & Kulkarni; 2011). |
| rb | **NOTC; VTRA** | **Ruby**, *Drosophila* HOPS & AP-3 complex genes are required for a Deltex (*dx*)-regulated activation of Notch in the endosomal trafficking pathway (Deltex: conserver regulator of Notch signaling), Notch receptor processing (Wilkin et al., 2008). |
| Pop2 | **miRNA** | **Pop2**, poly(A)-specific ribonuclease; involved in translation inhibition (Ruscica et al., 2019; Braun et al., 2011); miRNA-mediated mRNA degradation (Braun et al., 2011). |
| fzy | **CECY** | **Fizzy**, WD40 domain protein required for the full ubiquitin ligase activity of the anaphase-promoting complex / Cyclosome (APC/C) in mitosis and meiosis; functions to target substrates for destruction and drive metaphase and anaphase transition; anaphase-promoting complex-dependent catabolic process (Swan and Schupbach, 2007); mitotic cell cycle (Ducat et al., 2008). |
| Usp7 | **HIPP; HH** | **Ubiquitin-specific protease 7**, negative regulation of Hippo signaling pathway (Sun et al., 2019); positive regulation of Smoothened signaling pathway (Zhou et al., 2015). |
| Usp10 | **NOTC** | **Ubiquitin specific protease 10, ubiquitinyl hydrolase 1**, cysteine-type deubiquitinase activity; positive regulation of Notch signaling (Zhang et al., 2012). |
| SPARC | **DPPBMP** | **SPARC/Osteonectin**, acidic cysteine-rich secreted protein, small $Ca^{2+}$ and growth factor-binding glycoprotein, enriched in basement membranes; expressed in "loser" cells during cell competition to avoid apoptosis mediated by Flower (*fwe*) & Ahuizotl (*azot*); plays role during renal tubule morphogenesis in *Drosophila* (Bunt et al., 2010); cell adhesion (Hynes and Zhao, 2000). |

**Table D. Supplement to Figure 13: Signaling pathways linked to the Myc-CRM P35/P36**

| Gene name | Pathway | Activity-Function |
|---|---|---|
| puf | **WNBT; MYC; TOLL; NFKB; IMD** | **Puffyeye**, Ubiquitin-Specific Protease (USP), an essential deubiquitinating enzyme, acts as a ubiquitin-specific protease, removes ubiquitin polypeptide chains from Myc & CycE, leading to stabilization and increases of their abundance during cell growth & proliferation (Li et al., 2013); TCF dependent signaling in response to WNT. |
| Moe | **CECY; FGFR** | **Moesin**, Moesin/ezrin/radixin protein, involved in mitotic spindle organization (Solinet et al., 2013) & epithelial integrity; FGFR pathway (Slik and the receptor tyrosine kinase Breathless mediate localized activation of Moesin in terminal tracheal cells) (Ukken et al., 2014); maintenance of epithelial integrity via antagonistic interaction with the Rho GTPase pathway (Speck et al., 2003). |
| wrd | **INR; TORC; HIPP** | **(well-rounded)**, one of the two regulatory B' subunits of the protein phosphatase PP2A; influences metabolism and growth via negative regulation of the INR/TORC (Insulin-like Receptor) signaling network (Van Hoof and Goris, 2003). |
| zip (A0A0B4JD95) | **CECY; NOTC; JNK; WNPCP** | **Zipper, isoform H**, microtubule-binding protein involved in cytoskeleton-dependent intracellular transport; *zip* knockout or knockdown has no effect on overall oocyte growth during mid-oogenesis, but the oocyte marker (*orb*) less evenly localized in (*zip*) loss-of-function mutant egg chambers; mitotic cytokinesis (Dean et al., 2005; Rogers et al., 2003); Notch signaling dependent eye-antennal disc morphogenesis: epithelial fold determines boundary formation between developmental fields in the *Drosophila* antenna (Ku and Sun, 2017); JNK signaling: left/right axis specification (Okumura et al., 2010); Fz/Dsh signaling (a newly defined Fz/Dsh cytoskeletal signaling pathway with the involvement of fly myosin VIIA) (Winter et al., 2001). The seven-pass transmembrane receptor frizzled (Vinson et; al., 1989) requires the downstream signaling protein Dishevelled (*Dsh*) (Kling- ensmith et al., 1994; Theisen et al., 1994; Krasnow et al., 1995). Dishevelled & Frizzled also participate in the Wingless (Wg)/Wnt signaling to regulate developmental events including cell proliferation and cell fate specification (re- viewed in Wodarz and Nusse, 1998), however, Wg & planar cell polarity (PCP) distinct pathways downstream of Dishevelled (Axelrod et al., 1998; Boutros et al., 1998). |

**Table D. Supplement to Figure 13: Signaling pathways linked to the Myc-CRM P35/P36**

| Gene name | Pathway | Activity-Function |
|---|---|---|
| lincRNA.1023\CG12702 | **MYC** | (**lincRNA.1023, CG12702**), CIP2A, cancerous inhibitor of PP2A (CIP2A), functions through interactions with many other proteins including MYC, polo like kinase (PLK1), and NIMA protein; inhibitor of tumor suppressor PP2A, binds to PP2A and inhibits phosphatase function resulting in tumorigenic transformation of cells (Junttila et at. 2007; Carlson et al., 2014); CIP2A overexpressed in a number of tumors and expression levels are independent markers for long-term outcomes in many of these tumors (Carlson et at., 2014). |
| spag | **APOP** | **Spaghetti**, negative regulation of motor neuron apoptotic process (Means et al., 2015). |
| mask | **MAPK; JAKSTAT; VEGFR; EGFR; SEVE** | **Multiple ankyrin repeats single KH domain**, mediator of RTK signaling, either downstream of MAPK or signal transducer through a parallel branch of the RTK; EGFR pathway, Sevenless signaling (Smith et al., 2002); positive regulation of receptor signaling via JAK/STAT (Fisher et al., 2018); Vascular Endothelial Growth Factor Receptor signaling pathway (VEGFR) Tsai et al., 2022). |
| Lst8 | **TORC; INR** | **Lst8**, conserved TOR-binding protein; required for "CREB-regulated transcription coactivator2 (Crtc)-dependent regulation of cell growth based on genetic evidence; TORC2 signaling (Kuo et al., 2015; Wang et al., 2012); TORC1 signaling (Wang et al., 2012); positive regulation of Insulin-like receptor signaling (Yang et al., 2006); Target of Rapamycin Complex 2 regulates cell growth via Myc in *Drosophila* (Kuo et al., 2015). |
| CG12384 | **APOP** | Apoptosis signaling pathway (GO Reference Genome Project, 2011-). |
| Sep2 | **CECY** | **Septin 2**, regulation of cell cycle (O'Neill and Clark, 2013). |
| Vps4 | **CECY; EGFR** | **Vacuolar protein sorting 4**, cell cycle; EGFR: *Drosophila* Vps4 promotes Epidermal Growth Factor Receptor signaling (EGFR) independent of its role in receptor degradation (Legent et al., 2015); JNK signaling: Disruption of Vps4 and JNK function in *Drosophila* causes tumor growth (Rodahl et al., 2009). |

**Table D. Supplement to Figure 13: Signaling pathways linked to the Myc-CRM P35/P36**

| Gene name | Pathway | Activity-Function |
|---|---|---|
| slf | **HEBI** | **Schlaff**, chitin binding protein involved in wing formation, substrate for Transglutaminase (Tg); a putative C-type lectin needed for the adhesion between the horizontal cuticle layers (Zuber&Moussian, 2019); cooperation with the heme-biosynthesis pathway to stabilize the distribution of the cuticle tyrosinated proteins, exemplified by Resilin, the tyrosinated proteins network needed for correct contact between chitin laminae within the procuticle & the epicuticle (Zuber & Moussian, 2019). |
| lin-28 | **INR; JAKSTAT** | **Protein lin-28 homolog**, cold shock and RNA-binding protein - regulator of developmental timing - regulator of microRNA maturation; (insulin/IGF signaling (IIS) pathway) (Luhur et al., 2017; Chen et al., 2015); positive regulation of receptor signaling pathway via JAK/STAT (Sreejith et al., 2019). |
| gbb | **DPPBMP** | **(glass bottom boat)**, a BMP ligand in the TGF-β/BMP family of dimeric signaling molecules, binds to a receptor complex to transduce signal through phosphorylation of Mad; stem cell populations maintenance; control of cell fate specification, proliferation, synapse growth, & neuropeptide release (Anderson and Wharton, 2017; Jensen et al., 2009; Shimmi et al., 2005). |
| sxc | **FGFR** | **Super sex combs**, a Polycomb group, encodes a O-GlcNAc transferase, involved in epigenetic gene silencing; positive regulation of Fibroblast Growth Factor Receptor (FGFR) signaling pathway (Mariappa et al., 2011). |
| sun | **GPCR** | **Stunted,** activation of G-protein coupled receptor Methuselah (*mth*) in vitro, leading to increased intracellular Calcium ion levels; positive regulation of G protein-coupled receptor signaling pathway involved in regulating ageing; associated with longevity in *Drosophila* (Ja et al., 2009; Cvejic et al., 2004). |
| rumi | **NOTC** | **EGF-domain serine glucosyltransferase, EGF-domain serine xylosyltransferase**, functions as both O-glucosyltransferase & O-xylosyl-transferase (EGF-domain serine glucosyltransferase) (Takeuchi et al., 2011), an ER enzyme, adds a glucose residue to EGF-like repeats with a specific consensus sequence; modulation of Notch signaling: negatively (Acar et al., 2008) & positively (Servian-Morilla et al., 2016; Ishio et al., 2015; Leonardi et al., 2011; Perdigoto et al., 2011; Acar et al., 2008); regulates the function of other proteins including 'Eyes shut' (*eys*). |

**Table D. Supplement to Figure 13: Signaling pathways linked to the Myc-CRM P35/P36**

| Gene name | Pathway | Activity-Function |
|---|---|---|
| CG4115 | **IMD; NFKB** | Imd pathway (NF-kB signaling), carbohydrate binding (Tanji et al., 2006). |
| RanBPM | **WNBT; JAKSTAT; SAPA** | **Ran-binding protein M, isoform F**, interacts with Armadillo (beta-catenin) (Dansereau and Lasko, 2008); JAK/STAT signaling (Baeg et al., 2005); Sema3A/Plexin-A signaling (Togashi et al., 2006). |
| twf | **TOLL** | **Twinfilin**, highly conserved ubiquitously expressed, actin monomer binding and inhibition of actin filament assembly (GO Reference Genome Project, 2011-); has roles in bristle & neuronal development via Toll pathway (Cai et al., 2022). |
| scrib | **VTRA; EGFR; JAKSTAT; NOTC** | **Scribble**, scaffolding protein, part of conserved machinery regulating apicobasal polarity, interacts with "Discs large" (*dlg1*) & "lethal(2)giant larvae" (*l(2)gl*) to distinguish the basolateral domain of epithelial cells & neuroblasts via reciprocally antagonistic interactions with the aPKC/Par-6 complex that impacts vesicle trafficking; EGFR, JAK and Notch signaling (Li et al., 2009). |
| Gprk1 | **HH** | **G protein-coupled receptor kinase 1**, Hedgehog (Hh) signaling pathway; positive regulation of Smoothened signaling pathway (Cheng et al., 2010). |
| Idgf4 | **INR** | **Imaginal disc growth factor 4**, stimulation of insulin-like peptides for proliferation, polarization & motility of imaginal disk cells; stabilization of the binding of insulin-like peptides to insulin receptor (Kawamura et al., 1999) through a simultaneous interaction with both molecules to form a multiprotein signaling complex. |
| SCAR | **RTK** | **Scar**/WAVE complex pathway (Michael et al., 2013), lamellipodium; cell migration (Georgiou and Baum, 2010); actin filament organization (Sander et al., 2013). |
| cindr | **CECY** | **CIN85 and CD2AP related**, encodes an adaptor protein, links cell surface junctions & adhesion proteins with multiple components of the actin cytoskeleton; regulates cytoskeletal dynamics, eye patterning & endocytosis, cooperates with Scraps (*scra*) to promote intercellular bridge stability during cytokinesis; Cindr/dCortactin promote endocytosis (Quinones et al., 2010); Cindr:ArfGAP:dArf6 regulatory complex conserved across species, both during development & during maintenance of homeostasis (Johnson et al., 2011); Cindr/Anillin control cytokinesis in cleavage furrow (Haglund et al., 2010). |

**Table D. Supplement to Figure 13: Signaling pathways linked to the Myc-CRM P35/P36**

| Gene name | Pathway | Activity-Function |
|---|---|---|
| wmd | **WNBT**; **DPPBMP; EGFR** | **wing morphogenesis defect**; imaginal disc-derived wing morphogenesis with the involvement of Dpp, TGF-β, and Epidermal Growth Factor Receptor pathways (Dworkin and Gibson, 2006). |
| Vps29 | **WNBT** | **Vacuolar protein sorting 29**, Wnt ligand biogenesis & trafficking, retrograde transport endosome to Golgi (Franch-Marro et al., 2008; InterPro Project Members, 2004-). |
| P32 | **MCAM** | **P32**, evolutionarily conserved mitochondrial protein, functions in presynaptic calcium signaling & neurotransmitter release as well as chromatin metabolism, regulation of $Ca^{2+}$-mediated signaling (Lutas et al., 2012). |
| Rcc1 | **CECY; WNBT; APOP** | **Regulator of chromosome condensation 1**, nuclear import & export of β-Catenin (Koyama et at., 2017); regulation of mitotic cell cycle; apoptosis pathway (Trieselmann and Wilde, 2002). |
| mod | **MYC; CECY** | **Modulo**, the *Drosophila* homologue of nucleolin; required for meiosis & spermatid differentiation in male germline (Mikhaylova et al., 2006); MYC pathway (target of MYC selectively required for the growth of proliferative cells) (perinn et al., 2003); involved in chromatin packaging; dominant suppressor of variegation (Bantignies et al., 2002). |
| rush | **VTRA** | **(rush hour)**, Pleckstrin homology domain-containing family F, an endosome-associated protein, directly binds phosphatidylinositol 3-phosphate (PI3P) & the Rab GDP dissociation inhibitor encoded by (*Gdi*); regulates endosomal trafficking by modulating the activity of Rab proteins (Gailite et al., 2012). |
| ImpE1 | **ECDY** | **Ecdysone-inducible gene E1, isoform A**, This protein is similar to low-density lipoprotein LDL receptor; might play a role in the cell rearrangements associated with morphogenesis of the disc epithelium; imaginal disc eversion (Andres et al., 1993); (*ImpE1*) transcription is upregulated by 20-hydroxyecdysone. |
| TotZ | **JAKSTAT** | **Turandot**, belongs to a class of poorly characterized secreted peptides, and is expressed in response to oxidative stress, heat, UV, & bacteria in the fat body by the JAK/STAT pathway (Ekengren and Hultmark, 2001). |

**Table D. Supplement to Figure 13: Signaling pathways linked to the Myc-CRM P35/P36**

| Gene name | Pathway | Activity-Function |
|---|---|---|
| Map60 (CP-60) | **CECY** | **Microtubule-associated protein 60**, cell cycle-dependent Centrosomal localization (Kellogg et al., 1995). |
| Tailor | **miRNA** | **Tailor**, terminal uridylyl-transferase Tailor, RNA-mediated gene silencing; positive regulation of miRNA catabolic process (Lin et al., 2017); pre-miRNA processing (Reimao-Pinto et al., 2015). |
| CG6664 | **CECY** | Establishment of meiotic spindle orientation, spindle pole, and condensed chromosomes (Gene Ontology Curators, 2002-). |
| Ote | **DPPBMP** | **Otefin**, positive regulators of BMP (gbb/BMP) signaling pathway (Jiang et al., 2008). |
| Grip71 | **CECY** | **Grip71**, mitotic cell cycle (Ducat et al., 2008; Verollet et al., 2006). |
| Gp150 | **NOTC** | **Glycoprotein 150, isoform E,** Transmembrane glycoprotein regulates Notch signaling, involved in compound eye development (Fetchko et al., 2002); transmembrane receptor protein tyrosine phosphatase signaling pathway (Tian and Zinn, 1994). |
| e(y)3 (SAYP) | **JAKSTAT** | **Enhancer of yellow 3, isoform D**, coactivator of JAK/STAT pathway (Shidlovskii et al., 2005), required for embryogenesis and oogenesis (Vorobyeva et al., 2009). |
| Mad | **DPPBMP; WNBT; EGFR** | **Mothers against decapentaplegic**, BMP signaling pathway core components (Vuilleumier et al., 2022; Guo et al., 2013; Weiss et al., 2010; Kamiya et al., 2008; Yao et al., 2006; Muller et al., 2003; Dai et al., 2000; Das et al., 1998; Inoue et al., 1998); involved in wing development via modulation of EGFR and BMP signaling pathways (Dworkin and Gibson, 2006; Lecuit et al., 1996); negative regulation of salivary gland boundary formation with the involvement of Wnt/Wg signaling pathway (Bradley, P.L., Haberman, A.S., Andrew, D.J. 2001). |
| Kcmf1 | **RAS; MAPK** | **Potassium channel modulatory factor 1**, negative regulation of the Ras/MAPK signaling pathway in the wing by acting with the E2 enzyme Unc6 and the putative E3 ligases Poe and Ufd4 to mediate the ubiquitination and proteasomal degradation of Rl/MAPK; negative regulation of ERK1 & ERK2 cascade (Ashton-Beaucage et al., 2016). |
