## Supplemental Tables for "Targeting Regulatory Factors Associated with the *Drosophila Myc cis*-Elements by Reporter Expression, Gel Shift Assay, and Mass Spectrometric Protein Identification": Table E.pdf

**Table E. Supplement to Figure 13: Signaling pathways linked to the Myc-CRM P37/P38**

| Gene name | Pathway | Activity-Function |
|---|---|---|
| swm | **HH** | **second mitotic wave missing**, negatively regulates Hedgehog (Hh) protein signal in wing development (Casso & Karnberg, 2008). |
| Mtr4 | **INTERIM** | **Mtr4 helicase**, Interferon immune signaling, RNA helicase involved in the viral defense response (Molleston et al., 2016). |
| rin (CG9412) | **RAS; RTK** | **Rasputin**, evolutionarily conserved RNA-binding protein, functions as a link between Ras signaling and RNA metabolism (Costa et al., 2013; Pazman et al., 2000); positive regulator of *orb* during *Drosophila* oogenesis; Rasputin (*Rin*), the *Drosophila* homologue of Ras-GAP SH3 Binding Protein (G3BP), associates with Orb in a messenger ribonucleoprotein (mRNP) complex; Rho-mediated signaling (Pazman et al., 2000). |
| Ge-1 | **miRNA; NFKB; IMD; TOLL** | **Enhancer of mRNA-decapping protein 4 homolog**, RNAi pathway (miRNA-mediated gene silencing) (Eulalio et al., 2009); signaling cascades that activate NF-κB, Toll & Imd pathways (Jin et al., 2008; De Gregorio et al., 2002). |
| Bre1 | **NOTC; TOLL** | **Bre1**, E3 ubiquitin protein ligase, required for Notch signaling and histone modification (Bray et al., 2005); interaction with E2 ubiquitin ligase Ubc6, ubiquitination of H2B on lysine 120 at most RNA pol II transcripts; Toll pathway: involved in controlling host innate immune response (Cai et al., 2022). |
| awd | **FGFR; WNBT** | **abnormal wing discs**, *Drosophila* homolog of human Nm23, crucial for follicle cells during oogenesis, loss-of-function *awd* mutant cells result in the accumulation & spreading of adherens junction components, such as E-cadherin, beta-catenin/Armadillo, and alpha-spectrin, and disruption of epithelial integrity (Woolworth et al., 2009); regulation of tracheal cell motility by modulating the FGFR levels (Dammai et al., 2003). |
| spdi | **WNBT** | **split discs**, integrin signaling pathway in cell-migration & adhesion (Saadi et al., 2011). |
| sti | **HIPP; DPPBMP; NOTC; TOLL; WNBT; EGFR; CECY** | **Sticky**, negative regulation of Hippo signaling (Tran et al., 2019); regulation of TGF-β/BMP, Notch, Wingless, and EGFR signaling pathways (Bivik et al., 2015); mitotic cytokinesis (Echard et al., 2004; Naim et al., 2004; Shandala et al., 2004; Rogers et al., 2003). |

**Table E. Supplement to Figure 13: Signaling pathways linked to the Myc-CRM P37/P38**

| Gene name | Pathway | Activity-Function |
|---|---|---|
| sqd | **DPPBMP** | **Squid, isoform E**, Dpp signaling, squid & Hephaestus (but not Hrb27C) required for proper bone morphogenetic protein (BMP) signaling in GSCs; novel roles for RNA binding proteins Squid, Hephaestus, and Hrb27C during *Drosophila* oogenesis (Finger et al., 2022). |
| Pak3 | **MAPK; RACP; RAS; ERK** | **p21-activated kinase-1 (Pak1)**, non-specific serine/threonine protein kinase; an effector of Rho family GTPase Rac & Cdc42; regulation of Ras/Erk pathway (Rac GTPase Pak kinase signaling: RACP) (Arias-Romero et al., 2010). |
| Ulp1 | **HH; TOLL; NFKB** | **Ulp1**, negative regulators of Hedgehog signaling pathway (Zhang et al., 2017; Ma et al., 2016); negative regulators of Toll-NF-κB signaling pathway (Anjum et al., 2013). |
| unc-45 | **JAKSTAT** | **(uncoordinated mutant number 45)**, JAK/STAT signaling pathway controls host defense in the gut by regulating stem cell proliferation and thus epithelial cell homeostasis; negative regulation of innate immune response; defense response to Gram-negative bacterium (Cronin et al., 2009); A gradient of JAK/STAT pathway activity patterns the anterior-posterior axis of the follicular epithelium (Xi et al., 2003). |
| tws | **WNBT; HH; TOLL; NFKB; CECY** | **Twins**, a regulatory subunit of protein phosphatase 2A (PP2A), involved in many developmental processes & signaling pathways (Bajpai et al., 2004); Smoothened signaling pathway (Su et al., 2011); Toll signaling, positive regulators of Toll-NF-κB signaling pathway (Kanoh et al., 2021); mitotic cell cycle (Chen et al., 2007). |
| how | **EGFR; MAPK; ERK; DPPBMP; HH; INR; WNBT; NOTC; RAS** | **held out wings**, integrin signaling, apposition of dorsal/ventral imaginal disc-derived wing surfaces (Walsh and Brown, 1998); cell adhesion (Lo and Frasch, 1997); EGFR signaling: glial cell migration; axon ensheatment (Edenfeld et al., 2006); MAPK signaling: germline stem cell population maintenance (Monk et al., 2010), mesoderm spreading via suppression of alternate target gene (*miple1*) (Toledano-Katchalski et al., 2007); ERK1/2 (Nir et al., 2012) muscle development, Hedgehog signaling (Lobbardi et al.,2011); RNA binding, intercellular signal transduction & activation of RNA (STAR) proteins; Insulin receptor; Notch, Ras, and Rho signaling pathways (Lasko, 2003); Dpp signaling pathway (Israeli & Volk, 2007). |

**Table E. Supplement to Figure 13: Signaling pathways linked to the Myc-CRM P37/P38**

| Gene name | Pathway | Activity-Function |
|---|---|---|
| LKRSDH | **ECDY** | **Lysine ketoglutarate reductase/saccharopine dehydrogenase**, negative regulation of ecdysone receptor signaling pathway (Cakouros et al., 2008). |
| sle | **CECY** | **Slender lobes**, a component of nucleolus, its depletion alters the aggregation of nucleolar components & results in retardation of proliferation of Kenyon cells; function: nucleolus organization; identified by retarded mushroom body development, required for proper nucleolar organization in *Drosophila* (Orihara-Ono et al., 2005). |
| Hrb98DE | **PADPRI** | **Heterogeneous nuclear ribonucleoprotein at 98DE**, nuclear RNA-binding protein, controls hnRNA stability, splicing, IRES-dependent translation, & translational repression; represents one of the main targets of the poly(ADP-ribosyl)ation pathway; also regulates tissue polarity patterning and germline stem cell fate; negative regulation of RNA splicing (Ji and Tulin, 2009); post-translational modification of hnRNPs, such as poly(ADP-ribosyl)ation, is an important mechanism in the regulation of gene expression during development, such as eye pattern formation (Ji & Tulin, 2013). |
| Grip71 | **CECY** | **Grip71**, mitotic cell cycle (Ducat et al., 2008; Verollet et al., 2006). |
| enc | **CECY** | **Encore** regulation of germline mitoses (Hawkins et al., 1996). |
| tral | **piRNA** | **Trailer hitch, isoform D**, piRNA pathway (Liu et al., 2011). |
| Pop2 | **miRNA** | **Pop2**, poly(A)-specific ribonuclease; involved in translation inhibition (Ruscica et al., 2019; Braun et al., 2011); miRNA-mediated mRNA degradation (Braun et al., 2011). |
| sm (CG9218) | **ECDY; INR** | **Small ubiquitin-related modifier (Smooth)**, sole family member in *Drosophila*, and closest to human heterogeneous nuclear ribonucleoprotein L (hnRNP L) (zur Lage et al. 1997); required for embryonic patterning & mitosis (syncytial blastoderm mitotic cell cycle), also has roles in wing patterning via Dpp (Miles et al., 2008) and Ras/MAPK signaling pathways (Nie et al., 2009); localization during mitosis: nucleus during interphase, kinetochores, and midbodies; positive regulation of Smoothened signaling pathway (Zhang et al., 2017); involved in Toll-NF-κB signaling pathway (Kanoh et al., 2015). |

**Table E. Supplement to Figure 13: Signaling pathways linked to the Myc-CRM P37/P38**

| Gene name | Pathway | Activity-Function |
|---|---|---|
| CG34417 | **TOLL; DPPBMP; HH** | Mesoderm development via interaction with the Toll, Dpp, and Hedgehog signaling pathways (Furlong et al., 2001). |
| lincRNA.1023CG12702 | **MYC** | **lincRNA.1023, CG12702**, CIP2A, cancerous inhibitor of PP2A (CIP2A), functions via protein binding through interactions with many other proteins including MYC, polo like kinase (PLK1), and NIMA protein; inhibitor of tumor suppressor PP2A: binds to PP2A and inhibits phosphatase function resulting in tumorigenic transformation of cells (Junttila et at. 2007; Carlson et al., 2014); CIP2A overexpressed in a number of tumors and expression levels are independent markers for long-term outcomes in many of these tumors (Carlson et at., 2014). |
| Vps4 | **CECY; EGFR** | **Vacuolar protein sorting 4**, cell cycle; EGFR signaling: *Drosophila* Vps4 promotes Epidermal Growth Factor Receptor signaling independent of its role in receptor degradation (Legent et al., 2015); JNK signaling: Disruption of Vps4 and JNK function in *Drosophila* causes tumor growth (Rodahl et al., 2009). |
| Lst8 | **TORC; INR** | **Protein Lst8**, conserved TOR-binding protein; required for "CREB-regulated transcription coactivator2 (Crtc)-dependent regulation of cell growth based on genetic evidence; TORC2 signaling (Kuo et al., 2015; Wang et al., 2012); TORC1 signaling (Wang et al., 2012); positive regulation of Insulin-like receptor signaling (Yang et al., 2006); Target of Rapamycin Complex 2 regulates cell growth via Myc in *Drosophila* (Kuo et al., 2015). |
| Ns3 | **INR** | **Nucleostemin 3**, GTPase (Hartl et al., 2013); nuclear export of 60S ribosomal subunit by mediating the release of Nmd3 from the 60S ribosomal subunit after export into the cytoplasm (Hartl et al., 2013); regulator of body size: acts in serotonergic neurons to regulate insulin signaling receptor pathway and thus exerts global growth control (Kaplan et al., 2008). |
| RanBPM | **WNBT; JAKSTAT; SAPA** | **Ran-binding protein M, isoform F**, interacts with Armadillo (β-catenin) (Dansereau and Lasko, 2008); JAK/STAT signaling pathway (Baeg et al., 2005); Sema3A/Plexin-A signaling (Togashi et al., 2006). |

**Table E. Supplement to Figure 13: Signaling pathways linked to the Myc-CRM P37/P38**

| Gene name | Pathway | Activity-Function |
|---|---|---|
| Pi3K21B | **INR** | **Pi3K21B**, Insulin receptor signaling pathway (Breitkopf et al., 2016; Britton et al., 2002). |
| pont | **WNBT** | **Pontin**, RuvB-like helicase 1, canonical Wingless/Wnt pathway (Torres & Inestrosa, 2017), interaction with HMG box transcription factors of the lymphoid-enhancing factor-1 (LEF-1/TCF family, positive regulation of canonical Wnt signaling pathway (Bauer et al., 2000); JNK, negative regulation of tumor necrosis factor-mediated (TNFα-Eiger) signaling pathway (Wang et al., 2008). |
| E(bx) | **JAKSTAT** | **Enhancer of bithorax**, negative regulation of receptor signaling pathway via JAK/STAT signaling; negative regulation of innate immune response; spermatid differentiation (Kwon et al., 2009); nuclear receptor binding (Badenhorst et al., 2005); Nucleosome Remodeling Factor. |
| CDK2AP1 | **CECY; WNBT; RAS; NOTC; JNK** | **CDK2-associated protein 1**, orthologous to several human genes including CDK2AP1 (cyclin dependent kinase 2 associated protein 1), involved in developmental processes via Wingless, Ras, Notch, and JNK signaling pathways (Reddy et al., 2010). |
| wds | **NOTC; CECY; APOP** | **Will die slowly, isoform B**, Trithorax-related complex protein (Hollmann et al, 2002), *Drosophila* essential gene codes for a WD-repeat protein with seven repeats; WD40 repeat implicated in a variety of functions ranging from signal transduction and regulation of transcription to cell cycle control and apoptosis; positions the N-terminus of histone H3 for efficient trimethylation at 'Lys-4' (Pascual-Garcia et al., 2014); Mad interacts with the Wds to maintain active transcription by dynamically demethylating intragenic 6mA; maintenance of transcriptional activation for specific sets of genes in neurons together with DNA N6-methyl adenine demethylase Tet (yao et al., 2018); ATAC Complex, TRR complex, COMPASS Complex, TRX Complex. |
| Nurf-38 | **ECDY** | **Nucleosome remodeling factor - 38kD**, ecdysone receptor-mediated signaling pathway (Badenhorst et al., 2005). |

**Table E. Supplement to Figure 13: Signaling pathways linked to the Myc-CRM P37/P38**

| Gene name | Pathway | Activity-Function |
|---|---|---|
| Etl1 | **JAKSTAT** | **Etl1**, belonging to the Etl1 subfamily of the Snf2 family of helicase-related proteins, positive regulation of innate immune response via JAK/STAT signaling, host defense in the gut by regulating stem cell proliferation & epithelial cell homeostasis (Cronin et al., 2009). |
| Su(var)2-10 | **JAKSTAT; HH; IMD** | **Suppressor of variegation 2-10, isoform L**, JAK/STAT pathway regulator, contributes to eye formation & eye determination (Betz et al., 2001); negative regulators of JAK/STAT signaling (Muller et al., 2005; Betz et al., 2001); positive regulators of Smoothened signaling (Zhang et al., 2017; Ma et al., 2016); negative regulators of Imd signaling pathway (Tang et al., 2021; Cronin et al., 2009). |
| tacc | **CECY** | **Transforming acidic coiled-coil protein**, mitotic cell cycle (Gergely et al., 2000). |
| mod(mdg4) | **APOP** | **modifier of mdg4, isoform AD**, regulation of apoptotic process (Harvey et al., 1997). |
| psi | **MYC** | **P-element somatic inhibitor, isoform C**, far upstream binding-element protein 1/2, C-terminal (FUBP1/2) (Davis-Smyth et al., 1996; Ni et al., 2020); dual roles in RNA processing (Labourier et al., 2002; Siebel et al., 1994) and transcriptional regulation; FUBP1required for the activation of MYC transcription by binding to a single-stranded-far upstream sequence element of MYC promoter (Duan et al., 2017; Duncan et al., 1994). |
| Cdk5 | **APOP; CECY; WNBT** | **Cyclin-dependent kinase 5**, intrinsic apoptotic signaling (Kang et al., 2012); regulation of transcription involved in G1/S transition of mitotic cell cycle (GO Reference Genome Project, 2011-); proapoptotic signaling by CDK5 & MEKK1 (Ryoo HD., 2018); Wingless signaling: Human wild-type Tau interacts with Wingless pathway components & produces neurofibrillary pathology in *Drosophila* (Tau-induced neurodegeneration & the Wnt pathway using GSK-3β and 2 additional components, β -Catenin/Armadillo & dTCF (Jackon & Geschwind ,2002). |
| sbb | **NOTC; HH** | **Scribbler**, transcriptional coregulator, acts mainly as a co-repressor, interacts physically & genetically with Grunge (*Gug*) and is used by the repressor Tailless (*tll*) in early embryos, regulates larval behavior, photoreceptor axon target choice, and functions downstream of Hedgehog signaling; wing disc D/V pattern formation via Notch signaling pathway (Bejarano et al., 2008). |

**Table E. Supplement to Figure 13: Signaling pathways linked to the Myc-CRM P37/P38**

| Gene name | Pathway | Activity-Function |
|---|---|---|
| jub | **HIPP** | **Ajuba LIM protein, isoform D**, Hippo signaling pathway (Huang et al., 2016; Fletcher et al., 2015; Rauskolb et al., 2014; Sun and Irvine, 2013; Rauskolb et al., 2011; Das Thakur et al., 2010). |
| baf | **CECY** | **barrier to autointegration factor**, highly conserved in metazoan evolution, plays fundamental roles in nuclear assembly, chromatin organization, gene expression, and gonad development, may potently compress chromatin structure and be involved in membrane recruitment and chromatin decondensation during nuclear assembly; plays role in both the M phase & the interphase of the cell cycle (Nikalayevich and Ohkura, 2015; Furukawa et al., 2003). |
| barr | **CECY; HH; WNBT; NOTC; DPPBMP** | **Barren**, chromatin binding protein, involved in chromatin condensation; Malpighian tubule development & epithelial morphogenesis; tube development (Liu et al., 1999): Hedgehog is required for the completion of bud evagination (Hoch and Pankratz 1996), Wingless is required for cell division & morphogenesis in the tubules (Skaer and Martinez Arias 1992; Harbecke and Lengyel 1995); Notch receptor is required to define the single tip cell at the end of each tubule (Hoch et al. 1994), and elongation of the tubule (Skaer 1989), EGFR (EGF-like) ligand required for the proliferation of the distal cells of the tubule (Baumann and Skaer 1993; Kerber et al. 1998); Dpp expressed in foregut and hindgut, required for morphogenesis of both structures (Pankratz and Hoch 1995; Hoch and Pankratz 1996); cell cycle interphase (Lupo et al., 2001). |
| PpD3 | **JNK**; **MAPK** | **Protein phosphatase D3**, conserved protein, involved in JNK and MAPK pathways (Miskei et al., 2011; Morrison et al., 2000); required for mitosis (G2/M progression). |
| Elp4C (G6907) | **CECY** | **Elongator complex protein 4**, establishment of mitotic spindle asymmetry (Planelles-Herrero et al., 2022); phosphorylase kinase regulator activity (Gene Ontology Curators, 2002-). |
| dgt4 | **CECY** | **dim γ-tubulin 4 (Augmin complex subunit dgt4)**, mitotic cell cycle, plays role in centrosome-independent generation of spindle microtubules (Goshima et al., 2008); mitotic spindle assembly (Goshima et.al., 2007). |
| CNBP | **MYC** | **CNBP/MYC** axis; involved in the control of wing size by regulating Myc levels (Antonucci & Ganettieri, cc-2014). |

**Table E. Supplement to Figure 13: Signaling pathways linked to the Myc-CRM P37/P38**

| Gene name | Pathway | Activity/Function |
|---|---|---|
| eff | **APOP; CECY; HH** | **Effete**, E2 ubiquitin-conjugating enzyme, protein ubiquitination and degradation pathway; roles include female germline stem cell maintenance, eye development, regulation of apoptosis & chromatin organization; mitotic cell cycle (Cenci et al., 1997); negative regulation of Smoothened signaling pathway (Pan et al., 2017). |
| Su(fu) | **HH; DPPBMP** | **Suppressor of fused**, Ser/Thr protein kinase; negative regulator of Hedgehog signaling by forming a complex with the transcription factor Cubitus interruptus (Ci) (Han et al., 2019; Han et al., 2015; Smelkinson et al., 2007; Zhang et al., 2011; Fukumoto et al., 2001; Methot and Basler, 2000); negative regulator of Dpp pathway (Jia et al., 2002). |
| Larp4B | **MYC** | **La-related protein Larp4B**, negative regulation of cell growth and translation (Funakoshi et al., 2018). |
| Fmr1 | **miRNA; piRNA; ERK; WNBT; INR** | **Fragile X messenger ribonucleoprotein 1**, miRNA pathway (Specchia et al., 2017); MAPK/ERK signaling (Leahy et al., 2023); Wg/Wnt pathway; Alk/Jeb(Jelly belly) anterograde pathway: involves presynaptic secretion of Jeb ligand to activate postsynaptic Anaplastic lymphoma kinase (Alk) receptors & drive phosphorylation of substrates such as ERK resulting in modulation of both synaptic structure and function (Friedman et al., 2013); negative regulation of insulin receptor signaling pathway (Luhur et al., 2017; Monyak et al., 2017; Callan et al., 2012). |
| Tailor | **miRNA** | **Tailor**, terminal uridylyl-transferase, RNA-mediated gene silencing: positive regulation of miRNA catabolic process (Lin et al., 2017); pre-miRNA processing (Reimao-Pinto et al., 2015). |
| CaBP1 | **APOP** | **Calcium-binding protein 1**, protein disulfide-isomerase A6 homolog, apoptotic pathway (Okada et al., 2012); response to endoplasmic reticulum stress (GO Reference Genome Project, 2011-); protein disulfide isomerase activity (InterPro Project Members, 2004-). |
| Mi-2 | **NOTC; WNBT; ECDY** | **Mi-2**, Notch signaling (Zacharioudaki et al., 2019); Wingless and ecdysone pathways (Kon & Nusse; 2005). |
| IntS11 | **NOTC**; **EGFR**; **MYC** | **Integrator complex subunit 11**, Notch signaling (Shersher et.al., 2021); Epidermal Growth Factor Pathway (Tilley & Mollet, 2021). |

**Table E. Supplement to Figure 13: Signaling pathways linked to the Myc-CRM P37/P38**

| Gene name | Pathway | Activity-Function |
|---|---|---|
| pzg | **NOTC; ECDY; JAKSTAT; DPPBMP; WNBT** | **Putzig**, involved in chromatin activation of replication related genes & signaling pathways including Notch, ecdysone, & JAK/STAT; regulation of growth, cell death, and various developmental processes (Kugler et al., 2011) |
| Manf | **EGFR; APOP; MAPK** | **Mesencephalic astrocyte-derived neurotrophic factor**, an invertebrate neurotrophic factor supporting dopaminergic neurons, cell-cell signaling between glial cells & neurons during development, EGFR signaling; apoptosis pathway; MAPK signaling (Palgi et al., 2009). |
| Cdk1 | **HH; NOTC; HIPP; JAKSTAT; EGFR; WNBT; JNK; INR; CECY** | **Cyclin-dependent kinase 1**, G1/S transition of mitotic cell cycle (Lehner and O'Farrell, 1990); G2/S transition of mitotic cell cycle (Stern et al., 1993; Lehner and O'Farrell, 1990); follicle cell of egg chamber development, Notch, Hedgehog, EGFR, Wingless, JAK/STAT, Hippo, JNK, and insulin-PI3K pathways (Jia et al., 2015). |
| Ufd1-like | **ERAD; HH** | **Ubiquitin fusion degradation protein 1**, ubiquitin-dependent ERAD pathway; VCP-NPL4-UFD1 AAA ATPase complex, polyubiquitin modification-dependent protein binding (GO Reference Genome Project, 2011-); negative regulation of Smoothened signaling pathway (Zhang et al., 2013). |
| Gmd | **NOTC** | **GDP-mannose 4,6 dehydratase**, positive regulation of Notch signaling pathway (Perdigoto et al., 2011); cell fate commitment with the involvement of Notch signaling (Sasamura et al., 2007); involved in the early steps of *Drosophila* oogenesis via Notch signaling (Jagut et al., 2013). |
| p120ctn | **HGFS; WNBT; RTK** | **Adherens junction protein p120 (p120 catenin)**, an essential Rho-GTPase regulator; an important positive modulator of cell-cell adhesion (Myster et al., 2003); p120 catenin, a cadherin-associated protein, regulates Rho GTPases; Hepatocyte Growth Factor (HGF) HGFS signaling; Receptor Tyrosine Kinase (RTK) signaling; Wnt signaling (Yanagisawa et al., 2008). |
| lin-28 | **INR; JAKSTAT** | **Protein lin-28 homolog**, cold shock and RNA-binding protein - regulator of developmental timing - regulator of microRNA maturation; INR (insulin/IGF signaling (IIS) pathway) (Luhur et al., 2017; Chen et al., 2015); positive regulation of receptor signaling pathway via JAK/STAT (Sreejith et al., 2019). |

**Table E. Supplement to Figure 13: Signaling pathways linked to the Myc-CRM P37/P38**

| Gene name | Pathway | Activity-Function |
|---|---|---|
| Rcc1 | **CECY; WNBT; APOP** | **Regulator of chromosome condensation 1**, nuclear import and export of beta-Catenin (Koyama et at., 2017); regulation of mitotic cell cycle; apoptosis pathway (Trieselmann and Wilde, 2002). |
| Lis-1 | **CECY; DPPBMP** | **Lissencephaly-1 homolog**, required during several dynein- and microtubule-dependent processes such as nuclear migration during cell division, mitotic spindle formation and the removal of mitotic checkpoint proteins from kinetochores at the metaphase to anaphase transition (Siller et al., 2006; Liu & Steward, 1999); positive regulation of BMP signaling pathway (Chen et al., 2010). |
| Sep2 | **CECY** | **Septin 2**, regulation of cell cycle (O'Neill and Clark, 2013). |
| trol | **FGFR; EGFR; PVR; HH; WNBT; SAPA; RAS; TORC** | **Terribly reduced optic lobes**, encodes the extracellular matrix (ECM) molecule Perlecan (Pcan), a phylogenetically conserved secreted heparan sulfate proteoglycan (HSPG) (Schneider et al., 2006); regulator of cell-signaling by multiple growth factors including the Hedgehog (Smoothened signaling) (Park et al., 2003), wingless/WNT, FGF (negative regulation of fibroblast growth factor receptor signaling pathway) (Dragojlovic-Munther and Martinez-Agosto, 2013 ;Park et al., 2003), TGF-beta, EGF, & VEGF families required for the maintenance of epithelial-cell polarity & controlled proliferation of neuronal stem cells in the larval brain, Pcan binds & sequesters FGF & Hedgehog in ECM (Cho et al., 2012; Voigt et al., 2002); modulation of Sema-1a–Plex A signaling (SAPA) (Cho et al., 2012); ECM Heartless signaling in *Drosophila* blood progenitors regulates their differentiation via a Ras/ETS/FOG pathway & target of rapamycin (TOR function (Dragojlovic-Munther and Martinez-Agosto, 2013). |
| mre11 | **CECY** | **meiotic recombination 11** double-strand break repair protein, G2/M DNA damage checkpoint signaling (Bi et al., 2006); telomere maintenance (Gao et al., 2009; Bi et al., 2005; Bi et al., 2004; Ciapponi et al., 2004). |
| shep | **MAPK; ERK; VTRA** | **Protein Alan shepard**, MAPK/ERK pathway phosphorylated ERK (PERK) (Clement et al., 2013); vesicular trafficking: neuropeptide sorting through regulated secretory pathway (dense core vesicles) (Chen et al., 2014). |

**Table E. Supplement to Figure 13: Signaling pathways linked to the Myc-CRM P37/P38**

| Gene name | Pathway | Activity-Function |
|---|---|---|
| cindr | **CECY** | **CIN85 and CD2AP related**: adaptor protein, links cell surface junctions & adhesion proteins with multiple components of the actin cytoskeleton; regulates cytoskeletal dynamics, eye patterning, & endocytosis, cooperates with Scraps (*scra*) to promote intercellular bridge stability during cytokinesis; Cindr/dCortactin promote endocytosis (Quinones et al., 2010); Cindr:ArfGAP:dArf6 regulatory complex conserved across species, both during development and for the maintenance of homeostasis (Johnson et al., 2011); Cindr/Anillin control cytokinesis in cleavage furrow (Haglund et al., 2010). |
| Slik | **CECY; NOTC; EGFR** | **Sterile20-like kinase, isoform A**, mitotic spindle midzone (Carreno et al., 2008); protein serine/threonine kinase activity, coordinated regulation of epithelial morphology & proliferation (Hughes and Fehon, 2006); regulation of Notch and EGF Receptor (EGFR) signaling pathways (Hughes and Fehon, 2006; Maitra et al., 2006). |
| l(2)dtl | **CECY** | **lethal(2)denticleless**, encodes the Cdt2 protein, substrate-specific adapter of a DCX (DDB1-CUL4-X-box) E3 ubiquitin-protein ligase complex; required for cell cycle control and mediation of the polyubiquitination and subsequent degradation of substrates such as E2F during S phase; proteasome-mediated ubiquitin-dependent protein catabolic process (GO Reference Genome Project, 2011-); Cdt2 activity depends on DNA synthesis during S phase or is required after DNA damage involved in genome function and stability (Shibutani et al., 2008). |
| Ote | **DPPBMP** | **Otefin**, positive regulators of BMP (gbb/BMP) signaling pathway (Jiang et al., 2008). |
| lmgB | **APOP** | **lemming B**, apoptotic pathway (Nagy et al., 2012). |
| Cul4 | **HH** | **Cullin 4, isoform A**, molecular scaffold for the CRL4 E3 ubiquitin ligase complex, catalyzes the ubiquitylation & subsequent destruction of proteins functioning in cell growth, proliferation, transcription, replication & repair of the genome; SCF-dependent proteasomal ubiquitin-dependent protein catabolic process; cellular response to DNA damage stimulus (GO Reference Genome Project, 2011-); negative regulation of Smoothened signaling pathway (Li et al., 2018); protein ubiquitination (GO Reference Genome Project, 2011-; Li et al., 2018; Reynolds et al., 2008). |

**Table E. Supplement to Figure 13: Signaling pathways linked to the Myc-CRM P37/P38**

| Gene name | Pathway | Activity-Function |
|---|---|---|
| CG10565 | **WNBT** | **DnaJ homolog subfamily C member 2**, chaperone J-domain superfamily; DnaJ domain; Homeobox-like domain superfamily; J-protein Zuotin/DnaJC2; Myb domain; ribosome-associated complex head domain superfamily; SANT/MYB domain; [Zuotin-related factor 1 (Zrf1) is necessary for chromatin displacement of the Polycomb-repressive complex 1 (PRC1); is required for neural progenitor cells (NPC) specification from ESCs; promotes the expression of NPC markers, including the key regulator Pax6; essential to the establishment & maintenance of Wnt ligand expression levels, necessary for NPC self-renewal (Aloia et al., 2013). |
| Ced-12 | **JNK; PVR; PDGFR; VEGFR** | **Ced-12** (engulfment and cell motility), Pvr signaling: PDGF/VEGF (Platelet-Derived Growth Factor/Vascular Endothelial Growth Factor)-receptor related (Pvr), RTK activated by the binding of PDGF- & VEGF-related factors (Pvf1, Pvf2 or Pvf3), activates canonical Ras/Raf/MAPK (ERK) cascade, PI3K kinase pathway, TORC1 (FBrf0222697), RTK: Rho family small GTPases (Sopko & Perrimon, 2013; Ishimaru et al., 2004); JNK cascade (Sopko & Perrimon, 2013; Tran et al., 2013; Ishimaru et al., 2004). |
| CG9705 | **HH**; **NOTC** | **Cold shock domain-containing protein CG9705**, dendrite morphogenesis via Hedgehog signaling pathway (Iyer et al., 2013); Notch signaling pathway (Iyer et al., 2013; Mummery-Widmer et al., 2009). |
| tyn (CG17131) | **WNBT** | **trynity (SP71, EP1421)**,regulation of embryonic cell shape; Wnt signaling pathway (Fernandes et at., 2010). |
| mbt | **WNBT; MAPK** | **mushroom body tiny**, Ser/Thr-protein kinase PAK (P21-activated kinase), association with Wingless pathway by modulating DE-cadherin mediated cell adhesion via phosphorylation of Armadillo (Menzel et al., 2008); regulation of MAP kinase cascade; protein phosphorylation (GO Reference Genome Project, 2011-; Schneeberger and Raabe, 2003); mushroom body development (Melzig et al., 1998). |
| simj | **WNBT** | **Simjang, isoform D**, *Drosophila* homolog of mammalian p66, a component of the chromatin remodeling complex recruited by methyl-CpG-DNA binding proteins (Kim et al., 2004); Simjang/p66 plays role in *Drosophila* eye development as modifier (repressor) of Wnt signaling with the involvement of Mi-2/NURD complex (Kon et al., 2005); positive regulation of synaptic assembly at neuromuscular junction (Willemsen et al., 2013; Kon et al., 2005). |

**Table E. Supplement to Figure 13: Signaling pathways linked to the Myc-CRM P37/P38**

| Gene name | Pathway | Activity-Function |
|---|---|---|
| asun | **CECY** | **Protein asunder**, mitotic cell cycle (Lee et al., 2005). |
| pum | **EGFR** | **Pumilio, isoform G**, negative regulation of Epidermal Growth Factor Receptor (EGFR) signaling pathway (Kim et al., 2012). |
| Usp10 | **NOTC** | **Ubiquitin specific protease 10, ubiquitinyl hydrolase 1**, cysteine-type deubiquitinase activity; positive regulation of Notch signaling (Zhang et al., 2012). |
| Dmel\Vps29 | **WNBT** | **Vacuolar protein sorting 29**, Wnt ligand biogenesis and trafficking, retrograde transport endosome to Golgi (Franch-Marro et al., 2008; InterPro Project Members, 2004-). |
| Gp150 | **NOTC** | **Glycoprotein 150, isoform E,** transmembrane glycoprotein regulates Notch signaling, involved in compound eye development (Fetchko et al., 2002); transmembrane receptor protein tyrosine phosphatase signaling pathway (Tian and Zinn, 1994). |
| SCAR | **RTK** | **Scar**/WAVE complex pathway (Michael et al., 2013) lamellipodium; cell migration (Georgiou and Baum, 2010); actin filament organization (Sander et al., 2013). |
| Apoltp | **SREBP; WNBT; HH** | **Apolipoprotein lipid transfer particle**, Sterol regulatory-element binding protein; lipid transporter activity (Palm et al., 2012); regulation of Wingless and Hedgehog signaling pathways (Eugster et al., 2007). |
| Poc1 | **CECY** | **Proteome of centrioles 1**, centrosome cycle (centrosome duplication and separation) (Blachon et al., 2009). |
| Mad | **DPPBMP; WNBT; EGFR** | **Mothers against decapentaplegic**, BMP signaling pathway core components (Vuilleumier et al., 2022; Guo et al., 2013; Weiss et al., 2010; Kamiya et al., 2008; Yao et al., 2006; Muller et al., 2003; Dai et al., 2000; Das et al., 1998; Inoue et al., 1998); involved in wing development via modulation of EGFR and BMP signaling pathways (Dworkin and Gibson, 2006; Lecuit et al., 1996); negative regulation of salivary gland boundary formation with the involvement of Wnt/Wg signaling pathway (Bradley, P.L., Haberman, A.S., Andrew, D.J. 2001). |
| Trf2 | **ECDY** | **TATA box-binding protein-like 1**, ecdysone/ecdysteroid signaling pathway (Bashirullah et al., 2007; Shima et al., 2007). |

**Table E. Supplement to Figure 13: Signaling pathways linked to the Myc-CRM P37/P38**

| Gene name | Pathway | Activity-Function |
|---|---|---|
| (A4V4A5) Ran | **RAS; CECY; WNBT** | **GTP-binding nuclear protein Ran**, a member of the Ras superfamily (Koizumi et al., 2001); localizes around the microtubule spindle in vivo during mitosis in *Drosophila* embryos (Trieselmann & Wilde, 2002); regulation of nuclear import of β-Catenin into nucleus (Yokoya et al., 1999). |
