## Supplemental Tables for "Targeting Regulatory Factors Associated with the *Drosophila Myc cis*-Elements by Reporter Expression, Gel Shift Assay, and Mass Spectrometric Protein Identification": Table F.pdf

**Table F. Supplement to Figure 13: Signaling pathways linked to the W-CRM positive control**

| Gene name | Pathway | Activity/Function |
|---|---|---|
| asun | **CECY** | **Protein asunder**, mitotic cell cycle (Lee et al., 2005). |
| glu | **CECY** | **Gluon**, a subunit of the multiprotein complex Condensin, mitotic cell cycle: prometaphase chromosome condensation (Bivik et al., 2015) and sister chromatid segregation, contributes to nervous system development & glucose metabolism; ubiquitous early, expressed in dividing cells throughout embryogenesis—pole cells, neuroblasts in the CNS & the PNS. |
| sle | **CECY** | **Slender lobes**, a component of nucleolus, its depletion alters the aggregation of nucleolar components & results in retardation of proliferation of Kenyon cells; function: nucleolus organization; identified by retarded mushroom body development, is required for proper nucleolar organization in *Drosophila* (Orihara-Ono et al., 2005). |
| Grip71 | **CECY** | **Grip71**, involved in mitotic cell cycle (Ducat et al., 2008; Verollet et al., 2006). |
| tral | **piRNA** | **Trailer hitch, isoform D**, involved in piRNA pathway (*Liu et al., 2011*). |
| Enc | **CECY** | **Encore**, involved in the regulation of germline mitoses (Hawkins et al., 1996). |
| CaBP1 | **APOP** | **Calcium-binding protein 1**, (protein disulfide-isomerase A6 homolog), apoptotic pathway (Okada et al., 2012); response to endoplasmic reticulum stress (GO Reference Genome Project, 2011-); protein disulfide isomerase activity (InterPro Project Members, 2004-). |
| wrd | **INR; TORS; HIPP** | **well-rounded**; one of the two regulatory B' subunits of the protein phosphatase PP2A; influences metabolism and growth via negative regulation of the INR/TORC (Insulin-like Receptor) signaling network (Van Hoof and Goris, 2003). |
| CG30291 | **NFKB**; **ERK** | Involved in the regulation of NF-κB and ERK1/2 signaling pathways (Utreras & Kulkarni; 2011). |
| Cdk1 | **HH; NOTC; HIPP; JAKSTAT; EGFR; WNBT; JNK; INR; CECY** | **Cyclin-dependent kinase 1**, G1/S transition of mitotic cell cycle (Lehner and O'Farrell, 1990); G2/S transition of mitotic cell cycle (Stern et al., 1993; Lehner and O'Farrell, 1990); follicle cell of egg chamber development, Notch, Hedgehog, EGFR, Wingless, JAK/STAT, Hippo, & JNK pathways; insulin-PI3K pathway (Jia et al., 2015). |

**Table F. Supplement to Figure 13: Signaling pathways linked to the W-CRM positive control**

| Gene name | Pathway | Activity/Function |
|---|---|---|
| MED17 | **JASAU; RAS; MAPK** | **Mediator of RNA Pol II transcription subunit 17**, sex comb development; required for adult cell & segment identity specification; SP1 transcription activation; thyroid hormone receptor complex component, Vit D receptor interactor; jasmonate (JA) & Auxin signaling component (Boube et al., 2000); Ras/MAK pathway (Singh and Han 1995). |
| MED24 | **APOP; ECDY** | **Mediator of RNA Pol II transcription subunit 24**, positive regulation of apoptosis pathway (Wang et al., 2008); ecdysone-mediated induction of salivary gland cell autophagic cell death (Ihry and Bashirullah, 2014). |
| pont | **WNBT** | **RuvB-like helicase 1 (Pontin),** act through canonical Wnt/Wg pathway (Torres & Inestrosa, 2017), interaction with HMG box transcription factors of the lymphoid-enhancing factor-1 (LEF-1/T-cell factor (TCF) family; positive regulation of canonical Wnt signaling pathway (Bauer et al., 2000); negative regulation of tumor necrosis factor-mediated (TNFα) through modulation of JNK signaling pathway (Wang et al., 2008). |
| BAP155<br>moira (mor) | EGFR; **NFKB; CECY** | **Brahma associated protein 155 kDa**, imaginal disc-derived wing margin morphogenesis, EGFR (Terriente-Felix & de Celis, 2009); negative regulation of G1/S transition of mitotic cell cycle (Brumby et al., 2002); regulation of innate immune response via NF κB signaling (Bonnay et al., 2014). |
| Snr1 | **EGFR; NOTC; DPPBMP** | **Snf5-related 1**, Swi3 component of the Brahma complex, ATP-dependent chromatin remodeling; a counterpart of yeast SWI/SNF; interaction with the Notch signaling for the maintenance of neuroblast identity (Koe & Wang, 2014; Weng et al., 2010; Bowman et al., 2008; Wang et al., 2006); imaginal disc morphogenesis, EGFR signaling (Terriente-Félix and de Celis, 2009); roles in Notch, EGFR, and Dpp signaling pathways important for wing development (Curtis et al., 2011). |
| Bap111 | **WNBT**; **HH**; **DPPBMP**; **NOTC** | **Brahma associated protein 111kD**, extensive homology of the Brahma (BRM) complex to SWI/ SNF; Osa/Eyelid (*osa/eyelid*) shows a strong genetic interaction with (*brm*), suggesting close cooperation with the BRM complex (Treisman et al. 1997; Vazquez et al. 1999); Wnt/TCF, Hh, Dpp and Notch signaling pathways: Osa/Eyelid antagonizes Wingless signaling during *Drosophila* development and has homology to the novel Bright family of DNA-binding proteins (Treisman et al., 1997; Kal et al., 2000). |

**Table F. Supplement to Figure 13: Signaling pathways linked to the W-CRM positive control**

| Gene name | Pathway | Activity/Function |
|---|---|---|
| Apoltp | **SREBP; WNBT; HH** | **Apolipoprotein lipid transfer particle**, Sterol regulatory-element binding protein; lipid transporter activity (Palm et al., 2012); Wingless and Hedgehog signaling pathways (Eugster et al., 2007). |
| atms/Paf1 | **NOTC** | **Antimeros,** RNA Polymerase II-associated factor 1 **(PAF1)**, alternative mRNA splicing pathway (Lence et al., 2016; Burnette et al., 1999); formation of RNA pol II elongation complex; RNA pol II pre-transcription events; E3 ubiquitin ligases ubiquitinate target proteins; Notch signaling (Lence et al., 2016; Hongay & Orr-Weaver, 2011). |
| lds | **CECY** | **Lodestar** Transcription termination factor 2, involved in transcription termination (Xie, Z., Price, D.H. 1996), microtubule anchoring at centrosome: involved in the regulation of meiosis and mitosis during early embryonic cleavage, oogenesis, and spermatogenesis (Szalontai et al., 2009); DNA repair (GO Reference Genome Project, 2011-); SNF2-like chromatin remodeler. |
| Mapmodulin | **MAPK**; **APOP**; **HH**; **VTRA** | **Mapmodulin, isoform D**, (also named **ANP32A**), involved in nucleocytoplasmic transport (Gene Ontology Curators, 2002-) with implications in proliferation, differentiation, caspase-dependent / independent apoptosis, suppression of transformation, inhibition of protein phosphatase 2A (PP2A) (Costanzo et al., 2006); regulation of mRNA trafficking & stability; inhibition of acetyltransferases as part of the inhibitor of histone acetyltransferases (INHAT) complex; regulation of spermatogenesis & oogenesis via MAPK pathway (Beckmann et al., 2010); regulation of sonic Hedgehog signaling (Gold et al., 2003); vesicular secretory pathway (Lye et al., 2014); Belongs to Acidic Nuclear Phosphoprotein (ANP32) family. |
| Gug | **WNPCP; NOTC; DPPBMP; JNK; EGFR** | **Grunge, isoform C**, (related to human Atrophin-like proteins & is the only Atrophin-like protein in *Drosophila*), activation of Frizzled, Notch, Dpp, & JNK signaling pathways (Zheng et al., 1995; Artavanis-Tsakonas et al., 1999; Zeitlinger & Bohmann, 1999); eye development; negative regulation of Epidermal Growth Factor Receptor (EGFR) signaling pathway (Charroux et al., 2006); negative regulation of Hedgehog signaling pathway (Zhang et al., 2013). |
| HDAC1 | **HH** | **Histone deacetylase 1**, Hedgehog signaling pathway (Zhang et al., 2013). |

**Table F. Supplement to Figure 13: Signaling pathways linked to the W-CRM positive control**

| Gene name | Pathway | Activity/Function |
|---|---|---|
| Su(var)2-10 | **JAKSTAT; HH; IMD** | **Suppressor of variegation 2-10, isoform L**, regulation of JAK/STAT pathway during eye determination and eye formation (Betz et al., 2001); negative regulators of JAK/STAT signaling (Muller et al., 2005; Betz et al., 2001); positive regulators of Smoothened signaling (Zhang et al., 2017; Ma et al., 2016); negative regulators of Imd signaling pathway (Tang et al., 2021; Cronin et al., 2009). |
| HmgD | **EGFR; ECDY; DPPBMP; WNBT** | **High mobility group protein D, isoform C & High mobility group protein C, isoform D**, involved in the regulation of EGFR (Anan Ragab & Travers, 2006), Wnt/TCF (Archbold et al., 2014), and ecdysteroid signaling pathways (Chen et al., 2008); HMGD binding to the Dpp-responsive enhancer of (*tinman*) gene as well as to the Tinman protein during *Drosophila* cardiogenesis (Zaffran, 2002). |
| woc | **ECDY** | **Without children, isoform B**, ecdysone biosynthetic process (Wismar et al., 2000); regulation of transcription by RNA pol II (Font-Burgada et al., 2008). |
| Chi | **NOTC; WNBT** | **Chip, isoform B**, LIM domain-binding protein 2; LIM domain-binding protein/SEUSS; LIM interaction domain; Imaginal disc-derived wing morphogenesis (Rincon-Limas et al., 2000) / leg development; axon guidance (van Meyel et al., 2000). |
| msk | **HH; DPPBMP; MAPK; NOTC; EGFR** | **Moleskin/Importin-7**, (Vrailas et al., 2006); Notch, EGFR signaling (Pepple et al., 2007); regulation of MAPK, Dpp, and Hedgehog signaling pathways (Marenda et al., 2006). |
| 14-3-3zeta | **RAS; MAPK; HIPP; CECY** | **14-3-3 protein zeta**, Ras/Raf/MAPK signaling-dependent photoreceptor development (Kockel et al., 1997); regulation of Yorkie nuclear localization by Hippo (Hpo) signaling pathway (Zhang & Jiang, 2010). |
| net | **EGFR; DPPBMP; HH; WNBT** | **Net, isoform B**; E-box binding (GO Reference Genome Project, 2011-); imaginal disc-derived wing vein specification: EGFR signaling (Brentrup et al., 2000); imaginal disc-derived wing vein morphogenesis (De Celis, 2003); Notch signaling: the expression of E(spl)mβ and intervein-promoting genes *net*, *plexus*, and *bs* occur in the same domains as with the *iro* and *kni* during provein initiation; regulation of Dpp, Hedgehog, and Wingless signaling pathways (De Celis, 2003). |

**Table F. Supplement to Figure 13: Signaling pathways linked to the W-CRM positive control**

| Gene name | Pathway | Activity-Function |
|---|---|---|
| Dhx15 | **MAPK** | **MAPK**, p38 MAPK cascade (Mosallanejad et al., 2014). |
| Cbp80 | **miRNA** | **Cap binding protein 80**, pre-mRNA splicing and RNA-mediated gene silencing (RNAi); miRNA-mediated RNA interference via its interaction with Ars2, required for primary microRNAs (miRNAs) processing (Sabin et al., 2009). |
| tsu | **MAPK; RAS** | **Tsunagi**, rolled/MAPK pathway (Roignant & Treisman, 2010); RAS/MAPK signaling depends on the regulation of MAPK levels by the Exon Junction Complex (EJC) (Ashton-Beaucage & Therrien, 2010); The Mago/Tsunagi heterodimer interacts with the EJC key regulator Pym leading to EJC disassembly in the cytoplasm (Ghosh et al., 2014). |
| (fand) | **FGFR; WNBT; HH; NOTC; DPPBMP** | **'fandango'**, **Pre-mRNA-splicing factor syf1 homolog**, tracheal development, specifically splicing *bnl* transcripts resulting in the activation of the BNL-FGF pathway; regulation of Wnt/Wg, Hedgehog, Notch/Delta (N/Dl) signaling pathways (Hartenstein et al., 1992); TGF-β family (Dpp in *Drosophila*, BMP-2/4 in vertebrates (Fishman and Chien, 1997; Harvey, 1996; Bodmer, 1995). |
| CNBP | **MYC** | **CNBP/MYC** axis; involved in the control of wing size by regulating Myc levels (Antonucci & Ganettieri, cc-2014). |
| Swm | **HH** | **Second mitotic wave missing**, negatively regulates of Hedgehog (Hh) protein signal during wing development (Casso & Karnberg, 2008). |
| l(3)72Ab (Brr2) | **DPPBMP; CECY; ECDY** | **Brr2 U5 snRNP complex subunit, U5 small nuclear ribonucleoprotein 200 kDa helicase**, brr2 and Prp8 control expression of FMRFamide (FMRFa) neuropeptide specifically in six neurons of the VNC (Tv4 neurons); brr2 control is executed by two independent mechanisms, both required for the activation of the BMP retrograde signaling pathway in Tv4 neurons: (1) proper axonal pathfinding to target tissue to receive BMP ligand; (2) proper RNA splicing of 2 genes in BMP pathway: thickveins (*tkv*) gene, encoding a BMP receptor subunit, & Medea gene (a co-Smad) (Monedero Cobeta et al., 2018); mitotic cell cycle (Ducat et al., 2008); steroid hormone ecdysone signaling (Claudius et al., 2014). |

**Table F. Supplement to Figure 13: Signaling pathways linked to the W-CRM positive control**

| Gene name | Pathway | Activity-Function |
|---|---|---|
| Poc1 | **CECY** | **Proteome of centrioles 1**, centrosome cycle (centrosome duplication and separation) (Blachon et al., 2009). |
| IntS11 | **NOTC; EGFR; MYC** | **Integrator complex subunit 11**, Notch signaling (Shersher et.al., 2021); Epidermal Growth Factor Receptor pathway (Tilley & Mollet, 2021). |
| Sirt4 | **INR** | **NAD-dependent protein deacylase Sirt4**, plays role in the regulation of insulin signaling pathway (Wood et al., 2018). |
| aub | **miRNA; JAKSTAT** | **Aubergine**, piRNA binding (Huang et al., 2021; Webster et al., 2015; Nagao et al., 2010); global gene silencing by mRNA cleavage (Kennerdell et al., 2002) ncRNA-mediated post-transcriptional gene silencing (Tomari et al., 2004); RNA-mediated gene silencing (Bozzetti et al., 2015); JAK/STAT signaling pathway controls host defense in the gut by regulating stem cell proliferation & epithelial cell homeostasis (Cronin et al., 2009). |
| Pka-C1 | **HH; HIPP; APOP; ERK; MAPK** | **cAMP-dependent protein kinase catalytic subunit 1**, negative regulation of Smoothened signaling pathway (Ranieri et al., 2014; Jia et al., 2002); positive regulation of Hippo signaling pathway (Yu et al., 2013); regulation of apoptotic pathway (Christiansen et al., 2013); plays role in neuronal plasticity via ERK/MAP kinase signaling (impey et al., 1999). |
| Gprk1 | **HH** | **G protein-coupled receptor kinase 1**, Hedgehog (Hh) signaling pathway: positive regulation of Smoothened signaling pathway (Cheng et al., 2010). |
| CkIIalpha | **INR; TORC; DPPBMP; HH; NFKB; WNBT** | **Casein kinase IIalpha, isoform C**, (inositol kinase PI4KIII, inositol phosphatases CG9784 [IPP-like], Mipp2) metabolic signaling (via insulin receptor), & modulation of the presynaptic cytoskeleton (PP2A, PP4, cdc2, CKI, CK2, Mnb/Dyrk1a, Mast205) as important signaling nodes contributing to the control of synapse stability (Bulat et al., 2014); regulation of yolk catabolism by Atg1-Tor pathway (Kuhn et al., 2015); BMP signaling: regulation of synaptic growth via transcription of downstream targets (Ball et al., 2010; Baines, 2004; Haghighi et al., 2003; Marques et al., 2003); CkIIalpha regulation of Hedgehog signaling by both Smoothened & cubitus interruptus, phosphorylation of the components of Wnt, Akt, and NF-kB pathways by CK2 (Jia et al., 2010). |

**Table F. Supplement to Figure 13: Signaling pathways linked to the W-CRM positive control**

| Gene name | Pathway | Activity-Function |
|---|---|---|
| trol | **FGFR; EGFR; PVR; HH; WNBT; SAPA; RAS; TORC** | **Terribly reduced optic lobes**, encodes the extracellular matrix (ECM) molecule Perlecan (Pcan), a phylogenetically conserved secreted heparan sulfate proteoglycan (HSPG) (Schneider et al., 2006); regulator of cell-signaling by multiple growth factors including the Hedgehog (Smoothened signaling) (Park et al., 2003), Wingless/WNT, FGF (negative regulation of fibroblast growth factor receptor signaling pathway) (Dragojlovic-Munther and Martinez-Agosto, 2013 ;Park et al., 2003), TGF-beta, EGF, & VEGF families required for the maintenance of epithelial-cell polarity & controlled proliferation of neuronal stem cells in the larval brain, Pcan binds & sequesters FGF & Hedgehog in ECM (Cho et al., 2012; Voigt et al., 2002); modulation of Sema-1a–PlexA signaling (SAPA) (Cho et al., 2012); ECM Heartless signaling in *Drosophila* blood progenitors regulates their differentiation via a Ras/ETS/FOG pathway & target of rapamycin (TOR function (Dragojlovic-Munther and Martinez-Agosto, 2013). |
| Idgf3 | **INR** | **Imaginal disc growth factor 3, isoform D**, member of chitinase-like protein family; involved in insulin signaling in imaginal disc cell lines (Kawamura et al., 1999). |
| dlg1 | **EGFR; JNK; ECDY** | **discs large 1**, regulation of epidermal growth factor receptor signaling pathway; JNK pathway: establishment or maintenance of polarity of larval imaginal disc epithelium (Bunker et al., 2015); ecdysone signaling: negative regulation of peptidoglycan recognition protein signaling pathway (Xiong et al., 2016). |
| Sra-1 | **RTK; ERK; VEGFR; APOP; INR; NFKB; MAPK;WNBT** | **Cytoplasmic FMR1 interacting protein (specifically Rac1-associated protein 1)**, Rac1/RhoGTPase activity (linking WAVE complex- Abelson interactor Abi, HSPC300, Kette/Nap1- to Rac1 signaling) (Gohl et al., 2010); role in Rac1/RhoGTPase/FMR1 interaction (Schenck, A., Bardoni, B. et al., 2003); involved in multiple signaling pathways including VEGFA-VEGFR2, ERK, JNK, apoptotic, Insulin-like receptor, MAPK signaling, and NF-kB signaling pathway (Bogdan et al., 2004). |
| mri | **EGFR; RAS; APOP** | **Mrityu, isoform E**, Ras/EGFR and apoptotic pathways during retinal development (Rusconi & Challa, 2007). |
| CG11523 | **WNBT** | Kinase regulator activity and protein kinase A (PKA) binding activity; orthologous to human GSK3B interacting protein (GSKIP); regulation of canonical Wnt signaling pathway (InterPro Project Members, 2004-). |

**Table F. Supplement to Figure 13: Signaling pathways linked to the W-CRM positive control**

| Gene name | Pathway | Activity/Function |
|---|---|---|
| Septin5 (Sep5) | **CECY; APOP; DPPBMP; EGFR; NOTC** | **Septin5**, a member of the septin family of GTP-binding proteins; involved in cytokinesis; cell polarity & membrane rigidity; GTPase activity (GO Reference Genome Project, 2011-); imaginal disc-derived wing morphogenesis (Sun et al., 2021); involved in vein patterning & differentiation via modulation of TGF-β, Epidermal Growth Factor (EGF), & Notch ligands (Sun et al., 2021; De Celis, 2003). positive regulation of apoptosis pathway (Bae et al., 2007); regulation of cell cycle (O'Neill and Clark, 2013). |
| SMC3 | **CECY; NOTC; WNPCP** | **Structural maintenance of chromosomes 3**, a subunit of the cohesin complex, involved in planar cell polarity by regulating the membrane enrichment of the transmembrane cadherin encoded by "starry night" (*stan*); mitotic sister chromatid cohesion; cell division (GO Reference Genome Project, 2011-); collaboration with the Notch and Wnt/Frizzled-PCP signaling during establishment of imaginal disc-derived wing hair orientation and imaginal disc-derived wing morphogenesis (Mouri et al., 2012). |
| rept | **WNBT; MYC; APOP; CECY** | **Reptin**, RuvB-like helicase, transcriptional coactivator in Wingless signaling caused by altered Arm signaling; Pontin/Reptin interfere with the nuclear Arm signaling antagonistically; essential cofactor for the normal function of Myc; required for cellular proliferation and growth; ecdysone-mediated salivary gland cell autophagy cell death (Ihry and Bashirullah, 2014); negative regulation of canonical Wnt signaling pathway (Bauer et al., 2000); negative regulation of cell population proliferation (Bellosta et al., 2005). |
| galla-1 | **CECY; NOTC; HIPPs** | **Galla-1, isoform B**, Crumb (*crb*)-Galla-Xeroderma pigmentosum (*Xpd*), component of the Crb-Galla-Xpd (CGX) complex; essential for proper mitotic chromosome segregation in early embryos (Yeom et al., 2015); CGX complex also required for cell proliferation in developing wing discs (Yeom et al., 2015); galla-1 identified as (*crb*) modifier, loss-of-function or overexpression of galla-1 or (*crb*) show mitotic defects in early embryos; Crumb involved in many developmental pathways including regulation of Hippo signaling (Chen et al., 2010; Ling et al., 2010); regulation of Salvador/Warts/ Hippo signaling (Robinson et al., 2010); Notch signaling pathway (Herranz et al., 2006). |

**Table F. Supplement to Figure 13: Signaling pathways linked to the W-CRM positive control**

| Gene name | Pathway | Activity/Function |
|---|---|---|
| wds | **NOTC; CECY; APOP** | **Will die slowly, isoform B**, Trithorax-related complex protein (Hollmann et al, 2002), *Drosophila* essential gene codes for a WD-repeat protein with seven repeats; WD40 repeat implicated in a variety of functions ranging from signal transduction & transcriptional regulation to cell cycle control & apoptosis; positions the N-terminus of histone H3 for efficient trimethylation at 'Lys-4' (Pascual-Garcia et al., 2014); Mad interacts with the Wds to maintain active transcription by dynamically demethylating intragenic 6mA; maintenance of transcriptional activation for specific sets of genes in neurons together with DNA N6-methyl adenine demethylase Tet (yao et al., 2018); ATAC Complex, TRR complex, COMPASS Complex, TRX Complex. |
| rin (CG9412) | **RAS; RTK** | **Rasputin**, evolutionarily conserved RNA-binding protein functions as a link between Ras signaling and RNA metabolism (Costa et al., 2013; Pazman et al., 2000); positive regulator of (*orb*) during *Drosophila* oogenesis; Rasputin (*Rin*), the *Drosophila* homologue of Ras-GAP SH3 Binding Protein (G3BP), associates with Orb in a messenger ribonucleoprotein (mRNP) complex; Rho-mediated signaling (Pazman et al., 2000). |
| Arpc3A | **RTK** | **Actin-related protein 2/3 complex, subunit 3A**, structural constituent of cytoskeleton, contributes to actin filament binding activity, involved in Arp2/3 complex-mediated actin nucleation; Receptor Tyrosine Kinases (RTK)-N-WASP/SCAR-Rho-GTPase-Arpc2,3 (Benesh et al., 2002). |
| Mcr | **RTK; VTRA; TORC** | **Macroglobulin complement-related**, a transmembrane conserved thioester-containing protein (TEP); defense response to fungus by phagocytosis (Stroschein-Stevenson et al., 2006) with the involvement of RTK (Chimini & Chavrier, 2000); target of rapamycin (TOR) kinase signaling pathway; endomembrane vesicles (shannon et al., 2006); regulation of tube length, open tracheal system (Bäz et al., 2014). |
| Sep1 | **APOP; DPPBMP; EGFR; NOTC** | **Septin-1**, positive regulation of apoptotic pathway (Bae et al., 2007); imaginal disc-derived wing morphogenesis (Sun et al., 2021); involved in imaginal disc-derived wing morphogenesis via modulation of TGF-β, epidermal growth factor (EGF), & Notch ligands (Sun et al., 2021; De Celis, 2003). |

**Table F. Supplement to Figure 13: Signaling pathways linked to the W-CRM positive control**

| Gene name | Pathway | Activity/Function |
|---|---|---|
| Spt20 | **JNK**; **NOTC** | **SPT20, isoform C,** Spt-Ada-Gcn5-acetyltransferase (SAGA) complex; plays role in the regulation of JNK and Notch signaling pathways (Weake et al., 2009). |
| Iswi | **ECDY** | **Imitation SWI**, chromatin-remodeling complex ATPase chain Iswi, ecdysteroid signaling and metamorphosis (Badenhorst et al., 2005; Tsukiyama & Wu, 1995). |
| Mpcp2 | **NOTC** | **Mitochondrial phosphate carrier protein,** , Modulation of Notch signaling pathway during wing disc D/V patterning (Bejarano et al., 2008). |
| CG34159 | **MAPK** | MAPK-regulated corepressor-interacting protein; orthologous to human MCRIP2 (MAPK regulated corepressor interacting protein 2) (Gelbart and Emmert, 2013). |
