## Supplemental Tables for "Targeting Regulatory Factors Associated with the *Drosophila Myc cis*-Elements by Reporter Expression, Gel Shift Assay, and Mass Spectrometric Protein Identification": Table S 1.1-S1.6.pdf

**Table S1.1**

| No | Construct Name | Promoter Type | Promoter Size (kb) Coordinates on X Chr. | Plasmid Size (kb) | Contained Myc-CRMs | *lacZ* activity in Brain, Discs, Emb, Ova |
|---|---|---|---|---|---|---|
| 01 | pJ8 | *MYC* In2 | 8.161; ChrX:3,376,232..3,384,387 | 19.819 | P29/P30; P35/P36 | all tested tissues |
| 02 | pJ8.4 | pJ8 enhancer cluster, upstream of DPE | 0.840; ChrX:3,379,408..3,380,247 | 12.497 | P29/P30 | None |
| 03 | pJ8.5 | 3' end of In2 containing DPE Element | 2.102; ChrX:3,382,334..3,384,387 | 13,760 | P35/P36 | None |
| 04 | pJ2.1 | *MYC* noncoding 5' end | 7.158; ChrX:3,367,866..3,375,005 | 18.816 | P1/P2; P31/P32; P37/P38 | all tested tissues |
| 05 | pJ7 | -100bp + 5'UTR | 1.928; ChrX:3,373,088..3,375,005 | 13.5 | P31/P32; P37/P38 | Emb, Ova |
| 06 | pJ7.1 | 5'UTR, (-100bp contained in pJ7 deleted) | 1.806; ChrX:3,373,200..3,375,005 | 13.4 | P37/P38 | Emb, Ova |
| 07 | pJ7.2 | 5'-UTR (In1+non-coding E2) | 1.298; ChrX:3,373,710..3,375,005 | 13 | P37/P38 | Emb, Ova |
| 08 | pJ7.3 | 5'UTR (TATA region + Exon1) | 0.647; ChrX:3,373,088..3,373,728 | 12.3 | P31/P32 | None |
| 09 | pJ7.4 | pJ2.1 Enhancer region fused to pJ7 promoter | 3.498; composite promoter ChrX:3,369,648..3,371,174 ChrX:3,373,044..3,375,005 | 15.158 | P1/P2; P31/P32; P37/P38 | discs weaker than in pJ2.1 |
| 10 | pJ7.5 | J8 Enhancer region fused to J7 promoter | 2.896; composite promoter ChrX: 3,379,324..3,380,254 ChrX:3,373,044..3,375,005 | 14.518 | P29/P30; P31/P32; P37/P38 | all tested tissues |

**Table S1.1. The pJ2.1, pJ7, and pJ8 constructs and the truncations derived from these plasmids**. Each plasmid contains at least one of the investigated Myc-CRMs that were used in the study. The tested Myc-CRMs included P1/P2, P29/P30, P31/P32, P35/P36, and P37/P38. **Note:** pJ7.4 and pJ7.5 contain composite promoters. **Abbreviations: bps**, base pairs; **DPE**, Downstream Promoter Element; **E2**, Exon 2; **In1**, Intron 1; **UTR**, Untranslated Region; **Emb**, Embryo; **Ova**, Ovary.

**Table S1.2**

| No | | Name | Sequence (5'→3') | Size (bps) | Usage |
|---|---|---|---|---|---|
| 01 | | J7-F | AGCTCCTATTGGTACCACTTAAAGCG<br>AATT C | 31-mer | PCR - pJ7 |
| 02 | | J7-R | CTGGCGAAAGGGGGATGTGCT | 21-mer | |
| 03 | | J1.4-F | AACGCGGCCTTTTTACGGTTC | 21-mer | PCR - pJ7.1-<br>template |
| 04 | | J1.4-R | CACCCACCTTTCGGTACCACTTTCACT<br>TTC AC | 31-mer | |
| 05 | | J7.2-F | TAAATAGAGACGGATACCGCGGCTAT<br>G TTCAG | 32-mer | PCR - pJ7.2 |
| 06 | | J7.2-R | GCTGGCGACCTGCGTTTCAC | 20-mer | |
| 07 | | J7.3-F | CCCGCAGTGAGAGCGCAGAGAG | 22-mer | PCR - pJ7.3 |
| 08 | | J7.3-R | CTCGCTGAACATAGTTTCCGTATCCGT<br>C | 28-mer | |
| 09 | | J7.4-F | TCTAGTGTGCCGTTATAAAACCGCGG<br>CGCTGC | 32-mer | PCR - pJ7.4 |
| 10 | | J7.4-R | GCGGGTACCTGGAAAGTGTGCATGGC<br>GAGAT | 31-mer | |
| 11 | | In2E-F-Not | GCACTCTTAACAAGCGGCCGCATGTT<br>CTTTTT | 32-mer | PCR - pJ7.5 |
| 12 | | In2E-R-Asp | AACAGGTACCACCGCTGTGTTATAAA<br>CAATTGCG | 34-mer | |
| 13 | | SV40-F | GATTAGGGCCGCAAGAAAACTATCC | 25-mer | PCR- SV40<br>poly (A) |
| 14 | | SV40-R | TAGGCACCCCAGGCTTTACACTTTAT | 26-mer | |
| 15 | | SV40 | GTTCAGGGGGAGGTGTGGG | 19 | Sequences<br>SV40 P(A) |
| 16 | | pCAF | GACGGCGATATTTCTGTGGAC | 21-mer | Forward and<br>Reverse<br>sequencing of<br>pCaSpeR4 |
| 17 | | pCAR | CCTTAGCATGTCCGTGGGGTTTGA | 24-mer | |
| 18 | | Acc651_F | GCCGAGAAAATCAAATGGAC | 20-mer | Forward and<br>Reverse<br>sequencing of<br>pJ7.5 |
| 19 | | PZR | CGGGCCTCTTCGCTATTACG | 20-mer | |

**Table S1.2**. **Presented is the list of the analytical primer pairs that were used for the molecular cloning and sequencing of the plasmids**. The oligonucleotides were designed with the Bioinformatics tool Laser Gene module Primer Select and synthesized at Microsynth AG, Switzerland. The primer sets were used for sequencing and amplification of *Myc* non-coding regions as well as for sequencing of cloning vectors and injection plasmids. Orientation of the indicated primer sequences is 5'→3'. The length of primers is in base pairs (bps), PCR forward primers contain the letter 'F' and 'R' stands for the reverse primer.

**Table S1.3**

| No | Target Name | Sequence (5'→3') | Size (bps) | Identity | Modification |
|---|---|---|---|---|---|
| 01 | IR5-Scm1 | TGTGTGTGTGTGCGCACGCGCGTGTGTG<br>TGCGTGCATGCGTGCAT | 45 | Non-Dmel scrambled sequence (origin: Mus musculus *Mdm2* gene) | 5' Dyo 781 |
| 02 | Az5-Scm2-IR3 | ATGCACGCATGCACGCACACACACGCGC<br>GTGCGCACACACACACA | 45 | | 3' Dyo781, 5' Az |
| 03 | Scm2-IR3 | ATGCACGCATGCACGCACACACACGCGC<br>GTGCGCACACACACACA | 45 | The same as above | 3' Dyo781 |
| 04 | IR5-Scm1-Az3 | TGTGTGTGTGTGCGCACGCGCGTGTGTG<br>TGCGTGCATGCGTGCAT | 45 | | 5' Dyo781, 3' Az |
| 05 | IR5-Pos1 | TACGAT**AAGATCAAAGG**TGCAGT**GCCGC**<br>**CA**GTCGTA | 36 | HMG & Helper sites configurations in W-CRM | 5' Dyo781 |
| 06 | Az5-Pos2-IR3 | TACGAC**TGGCGGC**ACTGCA**CCTTTGATCT**<br>**T**ATCGTA | 36 | | 3' Dyo781, 5' Az |
| 07 | Pos2-IR3 | TACGAC**TGGCGGC**ACTGCA**CCTTTGATCT**<br>**T**ATCGTA | 36 | The same as above | 3' Dyo781 |
| 08 | IR5-Pos1-Az3 | TACGAT**AAGATCAAAGG**TGCAGT**GCCGC**<br>**CA**GTCGTA | 36 | | 5' Dyo781, 3' Az |
| 09 | IR5-P1 | GAAGCTTT**TTGTTG**T**TTTTGCAGCTGC**C**G**<br>**TTT**TCGCCCTT | 40 | J2.1 enhancer cluster | 5' Dyo 781 |
| 10 | Az5-P2-IR3 | AAGGGCGA**AAAC**G**GCAGCTGCAAAA**A**C**<br>**AACAA**AAAGCTTC | 40 | | 3' Dyo781, 5' Az |
| 11 | P2-IR3 | AAGGGCGA**AAAC**G**GCAGCTGCAAAA**A**C**<br>**AACAA**AAAGCTTC | 40 | The same as above | 3' Dyo781 |
| 12 | IR5-P1-Az3 | GAAGCTTT**TTGTTG**T**TTTTGCAGCTGC**C**G**<br>**TTT**TCGCCCTT | 40 | | 5' Dyo781, 3' Az |
| 13 | P3 | GAAGCTTT**TTGTTG**T**TTTTGCAGCTGC**C**G**<br>**TTT**TCGCCCTT | 40 | The same as above | 5'-biot |
| 14 | P4 | AAGGGCGA**AAAC**G**GCAGCTGCAAAA**A**C**<br>**AACAA**AAAGCTTC | 40 | | 3'-biot |
| 15 | IR5-P29 | ATG**CGCGTGGGAAAA**TCTT**ACA**GT**GCAG**<br>**CGGAAGCA**A**TGTTT**TTC | 45 | Dead Ringer in J8 enhancer region | 5' Dyo 781 |
| 16 | IR3-P30-Az5 | GAA**AAACA**T**TGCTTCCGCTGC**AC**TGT**AA<br>GA**TTTTCCCACGCG**CAT | 45 | | 3' Dyo781, 5' Az |
| 17 | P30-IR3 | GAA**AAACA**T**TGCTTCCGCTGC**AC**TGT**AA<br>GA**TTTTCCCACGCG**CAT | 45 | Enhancer cluster in J8 enhancer region | 3' Dyo 781 |
| 18 | IR5-P29-Az3 | ATG**CGCGTGGGAAAA**TCTT**ACA**GT**GCAG**<br>**CGGAAGCA**A**TGTTT**TTC | 45 | | 5' Dyo 781, 3' Az |
| 19 | IR5-P31 | GA**GCGCGGC**AGTCTGGTACGATAG**AAAT**<br>**TTTATTTAA**GCCACAG | 44 | GC1-TATA1-box region in J2.1, J7, and J7.3 - J7.5 Promoters | 5' Dyo 781 |
| 20 | Az5-P32-IR3 | CTGTGGC**TTAAATAAAATTT**CTATCGTAC<br>CAGACT**GCCGCGC**TC | 44 | | 3' Dyo781, 5' Az |
| 21 | P32-IR3 | CTGTGGC**TTAAATAAAATTT**CTATCGTAC<br>CAGACT**GCCGCGC**TC | 44 | The same as above | 3' Dyo781 |
| 22 | IR5-P31-Az3 | GA**GCGCGGC**AGTCTGGTACGATAG**AAAT**<br>**TTTATTTAA**GCCACAG | 44 | | 5' Dyo781, 3' Az |
| 23 | IR5-P35 | A**TCATTC**ATTCATTGTCTATCGAAAGCGC<br>GGTGGTGGGCTT**GGTCG**C | 47 | DPE-containing region in J8 | 5' Dyo 781 |
| 24 | Az5-P36-IR3 | G**CGACC**AAGCCCACCACCGCGCTTTCGA<br>TAGACAATGAAT**GAATGA**T | 47 | | 3' Dyo781, 5' Az |
| 25 | P36-IR3 | G**CGACC**AAGCCCACCACCGCGCTTTCGA<br>TAGACAATGAAT**GAATGA**T | 47 | The same as above | 3' Dyo 781 |
| 26 | IR5-P35-Az3 | A**TCATTC**ATTCATTGTCTATCGAAAGCGC<br>GGTGGTGGGCTT**GGTCG**C | 47 | | 5' Dyo 781, 3' Az |

**Table S1.3 continued**

| No | Target Name | Sequence (5'→3') | Size (bp) | Identity | Modification |
|---|---|---|---|---|---|
| 27 | IR5-P37 | A**CAA**CG**ATTT**CCGCCT**TA**TC**TATA**T**TTTT**TCA **GACAGGC**ATA**TAACTCAGGAA**C | 53 | 3' end of In1 & start of Ex2 noncoding region | 5' Dyo781 |
| 28 | Az5-P38-IR3 | G**TTCCTGAGTTA**TATG**CCTGTC**TGA**AAAA**AT **ATA**GA**TAA**GGCGG**AAAT**CG**TTG**T | 53 | | 3' Dyo781, 5' Az |
| 29 | P38-IR3 | G**TTCCTGAGTTA**TATG**CCTGTC**TGA**AAAA**AT **ATA**GA**TAA**GGCGG**AAAT**CG**TTG**T | 53 | The same as above | 3' Dyo781 |
| 30 | IR5-P37-Az3 | A**CAA**CG**ATTT**CCGCCT**TA**TC**TATA**T**TTTT**TCA **GACAGGC**ATA**TAACTCAGGAA**C | 53 | | 5' Dyo781, 3' Az |

**Table S1.3. Listed are the primer pairs prior to annealing to double-stranded oligos and use for the Electrophoretic Mobility Shift Assay (EMSA) and DNA-protein interaction assays followed by protein identification by Mass Spectrometry**. The oligonucleotide sequences for the primer pairs P1/P2 to P37/P38 were taken from the *MYC* non-coding Conserved *cis*-Regulatory Regions (Myc-CRMs). The bold nucleotides in each primer sequence are conserved in all or all but one of the other 5 orthologous *Drosophila* species, *D. sechellia*, *D. Yakuba*, *D. erecta*, *D. willistoni*, or *D. virilis*. Each block of bold bases, highlighted by gray represents a cluster of conserved sequences. Wingless *cis*-Regulatory Module (WCRM) (ARCHBOLD *et al.* 2014) was used as positive control. A non-*Drosophila* scrambled sequence was assembled from the noncoding region of the *Mdm2* gene in *Mus musculus* strain C57BL/6J (gi|372099100|ref|NC_000076.6|:117686685-117712961). The nucleotide search (Blastn) by both the NCBI and the FlyBase did not show any matches for the 45 base pair scrambled sequence with the fly genome. The oligonucleotides were synthesized at Microsynth AG (Balgach, Switzerland). The orientation of the indicated target (primer) sequences is 5'→3'. The lengths of the primers are indicated in base pairs (bps). **Abbreviations**: **biot**, biotinylated oligo; **DPE**, downstream promoter element; **Dyo 781**, Dyomics 781 (Infra-Red Dye); **In**, Intron; **Ex**, Exon; **IR**, Infra-Red; **Pos**, positive; **Scm**, Scrambled; **Az**, Azide moiety end-modification.

**Table S1.4**

| No | Bloomington Stock # | Name | Genotype with FlyBase Link |
|---|---|---|---|
| 01 | 6598 | y w | y$^{1}$ w$^{1118}$ |
| 02 | 8412 | dpp-lacZ | y$^{1}$ w$^{1118}$; P{dpp-lacZ.Exel.2}3 |
| 03 | 2475 | double balancer | w$^{*}$; T(2;3)ap$^{Xa}$, ap$^{Xa}$/In(2L)Cy, In(2R)Cy, Duox$^{Cy}$; TM3, Sb$^{1}$ |
| 04 | 11108 (Kyoto 108991) | blue balancer | Cyo, P{lArB}A66.2F2/b$^{1}$ Adh$^{*}$ cn$^{*}$ l(2)$^{**}$; ry$^{506}$ |
| 05 | 64349 (originally Stk#1) | Canton-S | Canton-S |

**Table S1.4. The list of the fly stocks used in the study**. Embryos taken from the genotype *y[1] w[*1118*]* (Bloomington Stock # 6598) were used for the random P-element mediated transgenesis. Embryos collected from the fly strain Canton-S (Bloomington Stock # 64349, originally Stk#1), 0-72 hours after egg laying (AEL), were used for cytoplasmic and nuclear protein extraction and purification. The fly stock "blue balancer" that expresses *lacZ* in embryos and ovaries, was used as a source for positive control in the embryos and ovaries staining. The dpp-lacZ line that expresses *lacZ* in the pattern of the dpp gene was used as positive control for the staining in third instar larval brain and imaginal discs. The yellow-white flies *y[1] w[*1118*]* were used as negative control.

**Table S1.5**

| No | Annealing Scheme for the Oligo Pairs | Names after Annealing | Coordinates on X Chr. |
|---|---|---|---|
| 01 | ds IR5-Scm1/Az5-Scm2-IR3 | IR-Scm1/Scm2 | NA |
| 02 | ds Scm2-IR3/IR5-Scm1-Az3 | IR-Scm2/Scm1 | |
| 03 | ds IR5-Pos1/Az5-Pos2-IR3 | IR-Pos1/Pos2 | NA |
| 04 | ds Pos2-IR3/IR5-Pos1-Az3 | IR-Pos2/Pos1 | |
| 05 | ds IR5-P1/Az5-P2-IR3 | IR-P1/P2 | ChrX:3,369,776..3,369,815 |
| 06 | ds P2-IR3/IR5-P1-Az3 | IR-P2/P1 | |
| 07 | ds IR5-P29/Az5-P30-IR3 | IR-P29/P30 | ChrX: 3,380,185..3,380,229 |
| 08 | ds P30-IR3/IR5-P29-Az3 | IR-P30/P29 | |
| 09 | ds IR5-P31/Az5-P32-IR3 | IR-P31/P32 | ChrX:3,373,105..3,373,148 |
| 10 | ds P32-IR3/IR5-P31-Az3 | IR-P32/P31 | |
| 11 | ds IR5-P35/Az5-P36-IR3 | IR-P35/P36 | ChrX:3,382,530..3,382,576 |
| 12 | ds P36-IR3/IR5-P35-Az3 | IR-P36/P35 | |
| 13 | ds IR5-P37/Az5-P38-IR3 | IR-P37/P38 | ChrX:3,374,908..3,374,960 |
| 14 | ds P38-IR3/ IR5-P37-Az3 | IR-P38/P37 | |

**Table S1.5. Depicted are the annealing schemes of the single-stranded oligos (Myc-CRMs, positive and negative controls) to double-stranded *Myc* targets and controls**. Each oligonucleotide was annealed with the complementary counterpart to obtain double-stranded targets. The double-stranded oligonucleotides were attached to the Bead-PEG heterodimers via "click" Chemistry reaction, and obtained Bead-PEG-DNA constructs were used in protein binding assays. The coordinates of the genomic loci of the Myc-CRMs are indicated on the X-Chromosome of *D. melanogaster*. **Abbreviations**: **ds**, double-stranded; **Scm**, Scrambled; **IR**, Infra-Red; **NA**, Not Applicable; **P**, **Primer**; **Pos**, Positive; **Az**, Azide. **Abbreviations**: **biot**, biotinylated oligo; **DPE**, downstream promoter element; **Dyo 781**, Dyomics 781 (Infra-Red Dye); **In**, Intron; **Ex**, Exon; **IR**, Infra-Red; **Pos**, positive; **Scm**, Scrambled; **Az**, Azide moiety end-modification.

**Table S1.6**

| No | Original Sample Name | Sample Description | Protein Origin | Sample Name for Mass Spec (MS) Analysis |
|---|---|---|---|---|
| 1 | Beads Only | Dynabeads Magnetic MyOne Carboxylic | None | 20210527_CSSS_ZUH_1 .raw |
| 2 | SCF Only | Soluble Cytoplasmic Fraction | SCF | 20210527_CSSS_ZUH_2 .raw |
| 3 | SNF Only | Soluble Nuclear Fraction | SNF | 20210527_CSSS_ZUH_3 .raw |
| 4 | IR-Scm1/Scm2:SCF | ds IR5-Scm1/IR3-Az5-Scm2, Scm2 strand attached to Beads | SCF | 20210527_CSSS_ZUH_4 .raw |
| 5 | IR-Scm2/Scm1:SCF | ds IR3-Scm2/IR5-Az3-Scm1, Scm1 strand attached to Beads | SCF | 20210527_CSSS_ZUH_5 .raw |
| 6 | IR-Scm1/Scm2:SNF | ds IR5-Scm1/IR3-Az5-Scm2, Scm2 strand attached to Beads | SNF | 20210527_CSSS_ZUH_6 .raw |
| 7 | IR-Scm2/Scm1:SNF | ds IR3-Scm2/IR5-Az3-Scm1, Scm1 strand attached to Beads | SNF | 20210527_CSSS_ZUH_7 .raw |
| 8 | IR-Pos1/Pos2:SNF | ds IR5-Pos1/IR3-Az5-Pos2, Pos2 strand attached to Beads | SNF | 20210527_CSSS_ZUH_8 .raw |
| 9 | IR-Pos2/Pos1:SNF | ds IR3-Pos2/IR5-Az3-Pos1, Pos1 strand attached to Beads | SNF | 20210527_CSSS_ZUH_9 .raw |
| 10 | IR-P1/P2:SCF | ds IR5-P1/IR3-Az5-P2, P2 strand attached to Beads | SCF | 20210527_CSSS_ZUH_1 0.raw |
| 11 | IR-P2/P1:SCF | ds IR3-P2/IR5-Az3-P1, P1 strand attached to Beads | SCF | 20210527_CSSS_ZUH_1 1.raw |
| 12 | IR-P1/P2:SNF | ds IR5-P1/IR3-Az5-P2, P2 strand attached to Beads | SNF | 20210527_CSSS_ZUH_1 2.raw |
| 13 | IR-P2/P1:SNF | ds IR3-P2/IR5-Az3-P1, P1 strand attached to Beads | SNF | 20210527_CSSS_ZUH_1 3.raw |
| 14 | IR-P29/P30:SCF | ds IR5-P29/IR3-Az5-P30, P30 strand attached to Beads | SCF | 20210527_CSSS_ZUH_1 4.raw |
| 15 | IR-P30/P29:SCF | ds IR3-P30/IR5-Az3-P29, P29 strand attached to Beads | SCF | 20210527_CSSS_ZUH_1 5.raw |
| 16 | IR-P29/P30:SNF | ds IR5-P29/IR3-Az5-P30, P30 strand attached to Beads | SNF | 20210527_CSSS_ZUH_1 6.raw |
| 17 | IR-P30/P29:SNF | ds IR3-P30/IR5-Az3-P29, P29 strand attached to Beads | SNF | 20210527_CSSS_ZUH_1 7.raw |

**Table S1.6 continued**

| No | Original Sample Name | Description | Protein Origin | Sample Name for Mass Spec (MS) Analysis |
|---|---|---|---|---|
| 18 | IR-P31/P32:SCF | ds IR5-P31/IR3-Az5-P32, P32 strand attached to Bead | SCF | 20210527_CSSS_ZUH_18.raw |
| 19 | IR-P32/P31:SCF | ds IR3-P32/IR5-Az3-P31, P31 strand attached to Bead | SCF | 20210527_CSSS_ZUH_19.raw |
| 20 | IR-P31/P32:SNF | ds IR5-P31/IR3-Az5-P32, P32 strand attached to Bead | SNF | 20210527_CSSS_ZUH_20.raw |
| 21 | IR-P32/P31:SNF | ds IR3-P32/IR5-Az3-P31, P31 strand attached to Bead | SNF | 20210527_CSSS_ZUH_21.raw |
| 22 | IR-P35/P36:SCF | IR5-P35/IR3-Az5-P36, P36 strand attached to Bead | SCF | 20210527_CSSS_ZUH_22.raw |
| 23 | IR-P36/P35:SCF | ds IR3-P36/IR5-Az3-P35, P35 strand attached to Beads | SCF | 20210527_CSSS_ZUH_23.raw |
| 24 | IR-P35/P36:SNF | IR5-P35/IR3-Az5-P36, P36 strand attached to Bead | SNF | 20210527_CSSS_ZUH_24.raw |
| 25 | IR-P36/P35:SNF | ds IR3-P36/IR5-Az3-P35, P35 strand attached to Beads | SNF | 20210527_CSSS_ZUH_25.raw |
| 26 | IR-P37/P38:SCF | IR5-P37/IR3-Az5-P38, P38 strand attached to Beads | SCF | 20210527_CSSS_ZUH_26.raw |
| 27 | IR-P38/P37:SCF | ds IR3-P38/IR5-Az3-P37, P37 strand attached to Beads | SCF | 20210527_CSSS_ZUH_27.raw |
| 28 | IR-P37/P38:SNF | IR5-P37/IR3-Az5-P38, P38 strand attached to Beads | SNF | 20210527_CSSS_ZUH_28.raw |
| 29 | IR-P38/P37:SNF | ds IR3-P38/IR5-Az3-P37, P37 strand attached to Beads | SNF | 20210527_CSSS_ZUH_29.raw |

**Table S1.6. List of the samples containing DNA-protein complexes for protein identification by Mass Spec analysis**. For each sample two replicates were prepared. Each double-stranded oligo is modified with the infrared dye Dyomics 781 at the 5' end of the top strands, and at the 3' end of the complementary bottom strand. In each replicate only one strand was azide modified for the "click" chemistry reaction. The "click" chemistry reaction took place between the azide moiety of the oligonucleotide and the amine end of the PEG23-amine that was attached to Dynabeads® MyOne™ Carboxylic Acid (Invitrogen by Thermo Fisher cat. #65001) (PEG23-Bead). This strategy helped to reduce the steric effect and crowdedness for the binding of the protein complexes to each DNA (target) that was linked to a Bead-PEG construct. **Abbreviations**: **IR**, Infra-Red; **Dyo 781**, Dyomics 781 (Infra-Red Dye); **P**, Primer; **SCF**, Soluble Cytoplasmic Fraction; **SNF**, Soluble Nuclear Fraction; **Scm**, Scrambled (negative control); **Pos**, Positive (positive control); **Az**, Azide moiety end-modification (in one of the two ds-oligo replicates attached to the "top" strand and in the other one to the "bottom" strand).
