## Supplementary material for "Targeting Regulatory Factors Associated with the *Drosophila Myc cis*-Elements by Reporter Expression, Gel Shift Assay, and Mass Spectrometric Protein Identification": Highlights

- Novel highly selective Solid Surface Magnetic Enrichment Protocol to purify proteins from crude extracts
- *Drosophila Myc* proximal super enhancer expresses eRNAs during early development
- *Drosophila Myc* downstream super enhancer requires the DPE promoter for transcription activation
- Wnt/Wingless signaling pathway is the main regulator of MYC
